# Mechanism-based inhibitors of SIRT2: structure–activity relationship, X-ray structures, target engagement, regulation of α-tubulin acetylation and inhibition of breast cancer cell migration

**DOI:** 10.1101/2020.03.20.000380

**Authors:** Alexander L. Nielsen, Nima Rajabi, Norio Kudo, Kathrine Lundø, Carlos Moreno-Yruela, Michael Bæk, Martin Fontenas, Alessia Lucidi, Andreas S. Madsen, Minoru Yoshida, Christian A. Olsen

## Abstract

Sirtuin 2 (SIRT2) is a protein deacylase enzyme that removes acetyl groups and longer chain acyl groups from post-translationally modified lysine residues. It affects diverse biological functions in the cell and has been considered a drug target in relation to both neurodegenerative diseases and cancer. Therefore, access to well-characterized and robust tool compounds is essential for the continued investigation of the complex functions of this enzyme. Here, we report a collection of probes that are potent, selective, stable in serum, water-soluble, amenable to cell culture experiments, and inhibit both SIRT2 deacetylation and demyristoylation. Compared to the current landscape of SIRT2 inhibitors, this is a unique ensemble of features built into a single compound. We expect the developed chemotypes to find broad application in the interrogation of SIRT2 functions in both healthy and diseased cells, and to provide a foundation for the development of future therapeutics.

## Introduction

The sirtuins are NAD^+^-dependent lysine deacylase enzymes that are highly conserved across species, with seven isoforms (SIRT1–7) present in humans. These enzymes share a common NAD^+^-binding pocket and catalytic core, but have different cellular expression profiles, subcellular localization, and substrate specificities.^1,2^ Sirtuins were originally reported to be ε-*N*-acetyllysine (Kac) hydrolases, but in recent years, it has become evident that a variety of ε-*N*-acyllysine posttranslational modifications (PTMs) can be removed by sirtuins,^3–8^ as well as by zinc-dependent histone deacetylases (HDACs).^9–14^ These findings formed the basis of a paradigm shift in the understanding of lysine modifications and their influence on cell signaling and implication in disease.^15^ Sirtuin 2 is predominantly localized to the cytosol, where it is believed to act mainly as a deacetylase of microtubular proteins such as α-tubulin,^16,17^ serving as a regulator in cell division and proliferation.^18,19^ Moreover, SIRT2 exhibits broad substrate scope, with a preference for long chain acyl groups (C_6_–C_16_) *in vitro*^5,20,21^ and was also recently shown to target lysine benzoylation (Kbz).^22^ Generally, SIRT2 is recognized as a tumor suppressor,^23^ although knockdown and inhibition of SIRT2 also have a broad anticancer effect in human breast cancer cell lines by promoting c-Myc degradation.^24^ Additionally, SIRT2 has been linked to neurodegeneration,^25,26^ and it has been shown to promote lipolysis and prevent differentiation in mature adipocytes,^27^ thus constituting a potential target for treatment of metabolic diseases and obesity.^28,29^ Interestingly, both activation and inhibition of SIRT2 appear to have therapeutic potential, depending on the biology under scrutiny. Accordingly, the complex role of SIRT2 calls for further investigation and development of tool compounds to enable these endeavors.

Numerous SIRT2 inhibitors have been reported recently (Fig. 1),^24, 30–48^ some of which are commercially available (*e.g.,* **SirReal2**, **AGK-2**, **tenovin-6**). However, all of these compounds are endowed with limitations such as lack of isozyme selectivity, lack of potency (revealed by limited ability to inhibit demyristoylase activity), and/or poor solubility. We therefore embarked on a mechanism-based and substrate-mimicking approach to develop novel inhibitors of SIRT2 with improved attributes and efficacy.

**Fig. 1.**
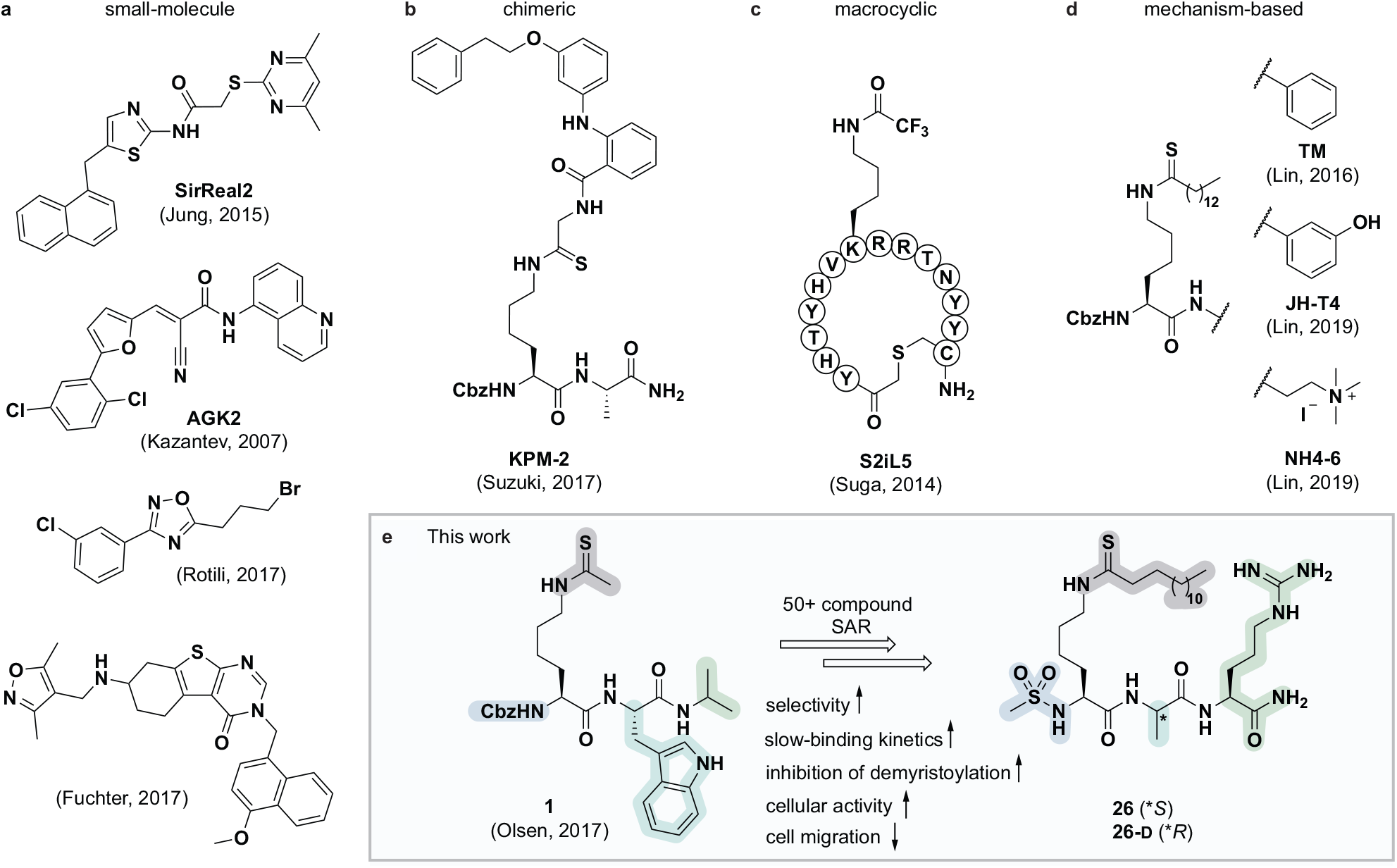
Representative SIRT2 inhibitors. (**a**) Heterocyclic small molecule inhibitors.^30,35,46,47^ (**b**) Chimeric lysine mimic **KPM-2**.^37^ (**c**) Peptide macrocycle **S2iL5**.^34^ (**d**) Mechanism-based inhibitors.^24,32,40^ (**e**) Summary of this study.

Efforts of numerous groups over the last two decades have led to almost 30 X-ray crystal structures, including several co-crystal structures with ligands and inhibitors.^34–40,49–56^ This has provided insight into the binding mechanism and substrate scope of SIRT2 at the molecular level. In the present work, we have built on this knowledge to develop the most potent and selective SIRT2 inhibitors reported to date. These inhibitors were shown to potently inhibit demyristoylation *in vitro*, increase perinuclear acetylation levels of α-tubulin in MCF-7 cells, and exhibit low nanomolar target engagement, as shown by cellular thermal shift assays. The inhibitors also exhibited inhibition of cell migration of MCF-7 breast cancer cells in culture. Finally, valuable insight into the stability of mechanism-based thioamide- and thiourea-containing sirtuin inhibitors in human serum was obtained, which will serve as useful guidance for future studies.

## Methods

For general experimental information, chemical synthesis, compound characterization data, additional methods, and copies of analytical HPLC spectra and NMR spectra of novel compounds, please consult the Supplementary Information.

### Materials and methods for the biochemical assays

SIRT1 (aa 193–741 with *N*-terminal GST-tag, ≥60% purity; cat. #50012), SIRT2 (aa 50–356 with *C*-terminal His-tag, ≥90% purity; cat. #50013), SIRT3 (aa 102–399 with *N-*terminal GST-tag; ≥64% purity; cat. #50014), and SIRT6 (full length with *N*-terminal GST-tag, ≥75% purity; cat. #50017) from BPS Biosciences (San Diego, CA). SIRT5 (aa 37–310 with *N*-terminal His-tag, ≥95% purity; cat. #BML-SE555-0050) from Enzo Life Sciences (Farmingdale, NY). Purities were based on SDS-PAGE and Coomassie blue stain according to the supplier and all enzyme concentrations given were based on the stock concentrations determined by the supplier. Sirtuin substrates were obtained from previous studies: Ac-Gln-Pro-Lys-Lys(Ac)-AMC (QPKKac),^20^ Ac-Glu-Thr-Asp-Lys(Myr)-AMC (ETDKmyr),^57^ Ac-Lys-Gln-Thr-Ala-Arg-Lys(Ac)-Ser-Thr-Gly-Gly-Trp-Trp-NH_2_ (H3K9ac),^20^ and Ac-Leu-Gly-Lys(Suc)-AMC (LGKsuc).^9^ Assay buffer was prepared as described in <u>Biomol International product sheets</u> (BML-KI-143; Tris HCl (50 mM), NaCl (137 mM), KCl (2.7 mM), MgCl_2_ (1 mM), pH 8.0) with addition of BSA (1.0 mg/mL, Sigma; cat. #A2153) unless stated otherwise. Trypsin (10,000 units/mg, TPCK treated from bovine pancreas; cat. #T1426) was purchased from Sigma (Steinheim, Germany). All other chemicals and solvents were of analytical grade and were used as obtained from commercial suppliers without further purification. All reactions were performed in black low binding 96-well microtiter plates (Corning half-area wells; cat. #3964), with duplicate series in each assay and each assay performed at least twice. All reactions were performed in assay buffer, with appropriate concentrations of substrates and inhibitors obtained by dilution from 2–50 mM stock solutions in either water or DMSO. The DMSO concentration in the final assay solution did not exceed 2% (v/v) unless stated otherwise. Control wells without enzyme and inhibitor (negative control) or without inhibitor (positive control) were included in each plate. Plates were analyzed using either a Perkin Elmer Enspire or a BMG Labtech FLUOstar Omega plate reader with excitation at 360 nm and detecting emission at 460 nm. Fluorescence measurements (RFU) were converted to [AMC] concentrations based on an [AMC]-fluorescence standard curve obtained in house, and all data analysis was performed using GraphPad Prism (version 8.1.2).

### End-point SIRT inhibition assays

End-point inhibition assays were performed as previously described.^58^ In brief, the relevant substrate, NAD^+^ (Sigma; cat. #N5755), and inhibitor were added to each well and the experiment was initiated by addition of a freshly prepared solution of relevant sirtuin, for a final volume of 25 μL per well. The following final concentrations were used: SIRT enzyme (100 nM; 600 nM for SIRT6), substrate (50 μM), NAD^+^ (500 μM), and inhibitor (1, 10 or 100 μM *or* fold dilution series for dose–response assays). The plate was incubated at 37 °C for 60 min, then a solution of trypsin and nicotinamide (NAM; 25 μL, 5 mg/mL and 4 mM, respectively; final concentration 2.5 mg/mL and 2 mM, respectively) was added, and the assay development was allowed to proceed for 90 min at RT before fluorescence measurement and calculation of residual activity. For concentration-response assays, IC_50_ values were obtained by fitting the resulting data to the concentration–response equation using GraphPad Prism (version 8.1.2).

### Continuous enzyme inhibition assays

Rate experiments for determination of kinetic parameters were evaluated under varying inhibitor concentrations.^59^ Buffer was prepared as previously described^21^ (HEPES/Na (50 mM), KCl (100 mM), Tween-20 (0.01%), TCEP (0.2 mM), BSA (0.05 mg/mL), pH 7.4). SIRT2 was incubated with substrate, inhibitor, and trypsin in assay buffer, for a final volume of 50 μL per well using the following final concentrations: SIRT2 (40 nM), ETDKmyr (10 μM), inhibitor (100–0.20 μM; 2-fold or 1.5-fold dilution series), NAD^+^ (200 μM), and trypsin (20 ng/μL). *In situ* fluorophore release was monitored immediately by fluorescence readings recorded every 30 seconds for 60 min at 25 °C. Assay progression curves of product concentration [P] vs. time (*t*) were fitted to Equation 1 in order to calculate the apparent first-order rate constant for reaching equilibrium (*k*_obs_) at each inhibitor concentration (*v*_ss_: steady-state velocity, *v*_in_: initial velocity), and kinetic parameters were extracted from either Equation 2 (mechanism A of slow binding, linear dependence of *k*_obs_ on the concentration of inhibitor) or Equation 3 (mechanism B of slow binding, hyperbolic dependence of *k*_obs_ on the concentration of inhibitor) as previously described.^59,60^ (*K*_M_ (ETDKmyr) = 1.8 μM).^21^

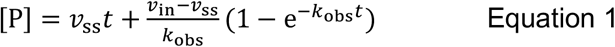

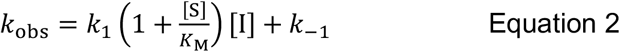

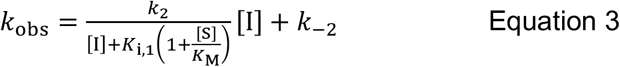

### End-point pre-incubation assays

SIRT2 and inhibitor were pre-incubated with or without NAD^+^ for 30 min at 37 °C in a total volume of 40 μL, prior to addition of substrate (and NAD^+^ if excluded in pre-incubation), for a final volume of 45 μL. For pre-incubation excluding NAD^+^, the following concentrations were used: SIRT1/2 (113 nM during pre-incubation, giving 100 nM after substrate and NAD^+^ addition), inhibitor (ranging from 113/100 μM down to 1.13/1.00 μM), substrate (0/50 μM) and NAD^+^ (0/500 μM); For pre-incubation including NAD^+^, the following concentrations were used: SIRT2 (113 nM during pre-incubation, giving 100 nM after substrate and NAD^+^ addition), inhibitor (ranging from 113/100 μM down to 1.13/1.00 μM), substrate (0/50 μM) and NAD^+^ (563/500 μM). The plate was incubated at 37 °C for 30 min, then a solution of trypsin and NAM (45 μL, 5.0 mg/mL and 4 mM, respectively; final concentration 2.5 mg/mL and 2 mM, respectively) was added, and the assay development was allowed to proceed for 90 min at RT before fluorescence measurement and calculation of residual activity.

### HPLC-based SIRT inhibition assays

All reactions were performed in black low-binding 96-well microtiter plates (Corning half-area wells; cat. #3964), with each assay performed twice, using the same assay buffer as in end-point fluorescence-based assays (Tris, pH 8.0), but with the omission of BSA. Inhibition assays (concentration–response) were performed in a final volume of 40 μL. The following final concentrations were used: SIRT1 or SIRT2 (20 nM), inhibitor (3-fold dilution series), H3K9ac (50 μM) and NAD^+^ (500 μM). After incubating sirtuin, substrate, and inhibitor (when applicable) for 30 or 45 min at 37 °C, the reaction mixture was quenched by the addition of ice-cold MeOH/HCOOH (94:6 (v/v), 20 μL). The samples were analyzed by HPLC on an Agilent 1260 Infinity II system equipped with a diode array detector. A gradient with eluent I (0.1% TFA in water/MeCN 95:5 (v/v)) and eluent II (0.1% TFA in MeCN (v/v)) rising linearly 0–40% during *t* 1.00–5.00 min was applied at a flow rate of 1.2 mL/min. The obtained chromatograms at 280 nm were used to determine reaction progression, by determining area under the curve of the relevant peaks.

### Chemical stability assays

#### Serum stability

400 μL human male serum (Sigma; cat. #H4522) was incubated at 37 °C for 15 min. The serum was spiked with a DMSO-stock solution of the respective inhibitor to reach a final concentration of 150 μM. The mixture was shaken at 750 rpm in an incubator at 37 °C. Samples (45 μL) were taken out at time points (0 min, 15 min, 30 min, 1 h, 2 h, 4 h, 6 h, and 24 h), quenched with aqueous urea (6 M, 50 μL), and incubated for 10 min at 4 °C. Ice-cold acetonitrile (100 μL) was added to each sample, which were incubated for another 10 min at 4 °C. The samples were centrifuged for 90 min at 20,000 g and filtered through 0.50 μm filters before analysis by HPLC and subsequent integration of the peak areas of recovered compound over time. Half-lives (*t*_½_) were determined using GraphPad Prism (version 8.1.2) and fitted to a one-exponential decay equation, assuming first-order kinetics. Each assay was performed at least twice.

#### Cell-media stability

For compounds **25-D** and **26-D**, a similar procedure to the above described was performed using MEM cell-medium (Sigma; cat. #2279) containing FBS (10% (v/v)).

#### Buffer stability

A DMSO-stock solution of compound **20**, **25-D**, or **26-D** was diluted to a final concentration of 100 μM in assay buffer (Tris, pH 8.0). The mixture was shaken at 750 rpm in an incubator at 37 °C, and samples were taken out at several time points (0, 2, 5, and 10 days) and analyzed on by HPLC.

### Immunocytochemistry

MCF-7 cells (150–1000 cells/well) were plated in Labtek Permanox Plastic Chamber slide system (Nunc, Thermo; cat. #177445) and incubated overnight. After 24 h, the cells were treated with inhibitor [**TSA** (5 μM), **26** (5 μM), **TM** (25 μM) or **26-D** (25 μM)] or DMSO (control) for 6 h, after which the cells were fixed in 4% formaldehyde for 15 min at RT. Cells were rinsed three times with PBS (pH 7.4) before blocking for 1 h with blocking buffer [5% goat-serum in PBS-T (PBS + 0.1% Triton-x100)] at RT and were then incubated with acetylated α-tubulin antibody (1:300, Santa Cruz Biotechnology, Dallas, TX; cat. #sc-23950 AC) in 5% goat-serum in PBS-T overnight at 4 °C. The cells were washed in PBS three times and the fluorophore conjugated antibody diluted in blocking buffer (goat-anti mouse Alexa 488, Thermo, Waltham, MA; cat #A11011) was added 1:800 and incubated 1 h in the dark at RT. After three washes with PBS, the slides were mounted using ProLong^®^ gold antifade mountant with DAPI (Thermo; cat. #P36941) and cells were visualized using an inverted fluorescent microscope at 40×. Images were generated using Visiopharm technology (Visiopharm, Hørsholm, DK), processed using ImageJ (version 1.8) and Adobe Photoshop Lightroom (version 5.3).

### Cellular thermal shift assays^61,62^

HEK293T cells were plated in 10 cm plates at 80–90% confluency and incubated overnight. Media was replaced with culture media containing SIRT2 inhibitor (0.01 μM for **26**, 0.10 μM for **26-D** and 10 μM for **TM**) or respective volume of DMSO. The cells were treated for 1 h and the media removed by aspiration. The cells were collected in PBS by scraping, and pelleted by centrifugation (300 g, 5 min). Pellets were resuspended in PBS and spun down again. The washed cell pellets were suspended in PBS supplemented with cOmplete EDTA-free protease inhibitor cocktail (COEDTAF-RO, Sigma, 800 μL/cell treatment). The cell suspensions were aliquoted into PCR tubes (60 μL) and at temperatures ranging from 37.0 °C to 75.0 °C for 3 min and then 3 min at 25 °C in an Eppendorf Mastercycler Nexus thermal cycler. The cellular suspensions were then lysed by three consecutive freeze/thaw cycles, snap-freezing in a dry-ice/acetone bath followed by thawing at 25 °C in the thermal cycler and subsequent vortexing. The suspensions were subjected to centrifugation (20,000 g, 20 min) at 4 °C, and the supernatants were collected as whole-cell lysate. The isolated lysates were resolved by SDS-PAGE in NuPAGE gels (4–12% Bis-Tris, Thermo; cat. #P0322BOX) with MES running buffer (Thermo; cat. #NP000202), and protein bands were transferred onto PVDF membranes (Thermo; cat. #IB24001) using an iBlot 2 gel transfer device. Membranes were blocked in 5% skim milk in TBS-T (TBS + 0.1% Triton-x100) for 1 h at RT. Subsequently, the membranes were washed with TBS-T (3×5 min) followed by incubation with primary antibody in 5% bovine serum albumin in TBS-T (1:1000) overnight at 4 °C. After another three cycles of washing with TBS-T, the membranes were incubated with HRP conjugated secondary antibody in 2% skim milk in TBS-T (1:10,000) for 1 h at RT. After washing with TBS-T (3×5 min) and TBS (1×5 min), membranes were visualized using enhanced chemiluminescent reagents (Pierce ECL Western Blotting Substrate, Thermo; cat. #32106) on a syngene PXi4 image analysis system. Antibodies: mouse anti-SIRT1 (Santa Cruz Biotechnology; cat. #sc-74504), rabbit anti-SIRT2 (Cell Signaling Technology (CST); cat. #12650), rabbit anti-SIRT3 (CST; cat. #5490), Goat anti-rabbit IgG (H+L) HRP-conjugated (Thermo; cat. #31466), Goat anti-mouse poly-HRP (Thermo; cat. #32230).

### Scratch assays

MCF-7 cells were plated in ibidi 2-well culture inserts (ibidi Gmbh, Gräfelfing, Germany; cat. #80209) in 24-well plates at 300,000 cells/well and incubated overnight. After overnight incubation, the inserts were removed and the cells were washed once in PBS and fresh medium was added. Cells were left to settle for 15 min before adding culture medium containing SIRT2 inhibitor [**TM** (5 μM), **26** (0.05 μM), **26-D** (0.5 μM)] or a DMSO control. Brightfield images were taken at time points 0, 2, 4, 6, 8, and 24 h, and the gap areas for each image were calculated using ImageJ (version 1.8) and used to determine the %-closure of the gaps at each time point.

### Crystallization and X-ray co-crystal structures

Crystallization was obtained using the catalytic domain of human SIRT2 lacking loop regions comprising residues 292–303. Final construct: residues (52–291) + (304–356). The SIRT2 catalytic domain (16 mg/mL) and compound **13** or **23** (final concentration 1 mM) were mixed, and crystallization screening was performed using commercial kits (Hampton Research, Aliso Viejo, CA). Crystals for X-ray crystallography were obtained using 0.1 M HEPES, pH 7.0, 30% (v/v) with Jeffamine ED-2001, pH 7.0 for compound **13** or with 0.15 M potassium bromide, 30% (w/v) polyethylene glycol monomethyl ether 2,000 for compound **23** at 20 °C. Crystals were frozen with liquid nitrogen using PEG400 10% (w/v) as cryoprotectant. X-ray diffraction data were collected at 100 K in a nitrogen gas stream at the synchrotron beamlines, PF-BL 5A, 17A at Photon Factory and X06DA at the Swiss Light Source. Data were processed and scaled with the XDS program package.^63^ The crystal structures were determined by the molecular replacement method with MOLREP,^64,65^ using the structure of SIRT2 in complex with H3K9-myr peptide (PDB 4Y6L).^56^ Refinement and model building were performed with REFMAC5^65,66^ and Coot.^67^ Coordinates of inhibitors were calculated in AceDRG.^68^ The geometric quality of the model was assessed with MolProbity.^69^ Data collection and refinement statistics are listed in Table S1. Figures containing structural elements were created using PyMOL Molecular Graphics System (version 1.8.0.6., Schrödinger, LLC).

## Results

### Structure–activity relationship study and X-ray crystallography

During our previous structure–activity relationship (SAR) study that targeted SIRT5,^58^ we found that thioacetamide **1** (Scheme 1a) did not inhibit SIRT5 but rather exhibited inhibition of SIRT2 (14% at 10 μM) and could therefore serve as a starting point for the development of potent substrate-mimicking inhibitors of SIRT2. While thioacetamide-based compounds had already been reported to inhibit SIRT2,^70,71^ we turned our attention to longer-chain acyl groups, inspired by the ε-*N*-thiomyristoyllysine-based inhibitor, **TM**, developed by Lin and co-workers.^24^ Similar to our findings for SIRT5, the initial series of compounds (**2**–**5**, Scheme 1b) revealed that thioamide- and thiourea-based ε-*N*-acyllysine mimics (compounds **2** and **3**, respectively) led to highly potent inhibitors. Interestingly, introduction of analogous hydrazide or inverted amide moieties (compounds **4** and **5**, respectively), which have previously been reported to serve as sirtuin inhibitors,^72,73^ exhibited only limited inhibition in our assays at 100 μM concentration of the compounds. Because the thiourea modifications can be introduced in a late stage of the compound preparation, we chose the thiourea moiety for the subsequent compound series.

We then gained inspiration from the X-ray co-crystal structure of SIRT2 bound to the 14-mer macrocyclic inhibitor **S2iL5**,^34^ which is an analog of macrocyclic inhibitors discovered by Suga and co-workers using mRNA-display technology.^42^ This structure indicated electrostatic interactions between the guanidinium group of the arginine in the *i*+2 position (*i*: position of the modified lysine residue) and two nearby glutamic acid residues of SIRT2 (E116 and E120), prompting us to extend the scaffold by one additional *C*-terminal amino acid residue. This also enabled a solid-phase peptide synthesis (SPPS) approach, using the Rink amide linker^74^ to provide ready access to compounds **6**–**12** (Scheme 1c, for synthesis see Scheme S4). Gratifyingly, potency was improved up to 5-fold by the *i*+2 extension. However, it proved challenging to determine meaningful IC_50_ values for compounds of this potency using our standard deacetylation assay, because stoichiometric inhibition was reached. We therefore determined compound potency in a similar demyristoylation assay using our previously developed myristoylated substrate (Ac-ETDKmyr-AMC),^21,57^ which has a substantially lower *K*_M_ value than acetylated substrates for SIRT2.^21^ The values for deacetylation (shown with an asterisk in Scheme 1), thus remain virtually constant throughout the attempts to improve upon compound affinity, while the ability to inhibit demyristoylation provides a more sensitive read-out of inhibitory potency.

The inhibition of the SIRT2 demyristoylase activity by compounds **6**–**12** revealed equipotency for positively and negatively charged amino acids and lower potency for the neutral residues (**7** and **12**). Thus, we selected compound **9**, containing an arginine residue in the *i*+2 position, for further modification, due to a presumed positive effect on solubility and cell permeability. Reintroduction of a short-chain thioacetyl group (compounds **S1**–**S4**, Scheme S1) led to significantly decreased potency against SIRT2, while displaying respectable inhibition against SIRT1, further emphasizing the importance of a longer chain acyl group to target SIRT2 over SIRT1 and SIRT3.^24,75^

With compound **9** as the starting point, we then investigated the importance of the *i*+1 residue. In some X-ray co-crystal structures of SIRT2, residues surrounding this position are unresolved due to high flexibility and unstructured binding interactions.^6,35^ In particular, amino acids 289–304, which are located at the putative *i*+1 binding region, comprise a SIRT2-specific insertion claimed to be stabilized by—and indeed resolved in the co-crystal structure with—the macrocycle **S2iL5**.^34^ We therefore investigated the importance of the *i*+1 residue by varying its charge, steric bulk, and stereochemistry in analogs **13**–**23** (Scheme 1d). In agreement with the expected flexibility of the binding region, we found that all modifications except proline led to potent compounds against the deacetylation activity of SIRT2. However, the inhibition data against demyristoylation activity revealed that the D-Ala (**21**) or 2-aminobutyric acid (Aib) (**23**) containing analogues were less potent, showing that stereochemistry at this position is more important for binding affinity than side chain functionality.

At this stage, we obtained diffraction-quality crystals of SIRT2 with compounds **13** and **23** bound and were able to solve X-ray co-crystal structures of both complexes at 1.7 Å resolution (Fig. 2). Superimposing the two structures revealed highly similar conformations (Fig. 2a,b) as well as high similarity to a previously solved co-crystal structure of SIRT2 with a thiomyristoylated peptide substrate analog bound (Fig. 2c). More pronounced differences were observed when comparing our structures to a structure of the apo form of SIRT2 and the structure with **S2iL5** bound (Fig. 2d). Not surprisingly, the upper zinc-binding domain and substrate binding pocket adopted a tighter conformation when bound to the macrocyclic peptide compared to our inhibitors, which require accommodation of the long fatty acyl side chain modification in the substrate binding pocket (Fig. 2d), whereas the apo form adopted a more open structure. Generally, the most prominent inhibitor–enzyme interactions were through backbone–backbone hydrogen bonding (Fig. 2e), in agreement with those observed from an X-ray co-crystal structure with a thiomyristoylated peptide bound to SIRT2 (Fig. 2f).^6^ On the other hand, a recent structure of SIRT2 with a glucose-containing analog of **TM** showed a different binding mode (Fig. 2f),^40^ which could help explain the lower potency of **TM** analogs compared to inhibitors with a higher peptide content, due to loss of backbone–backbone interactions.

**Scheme 1.**
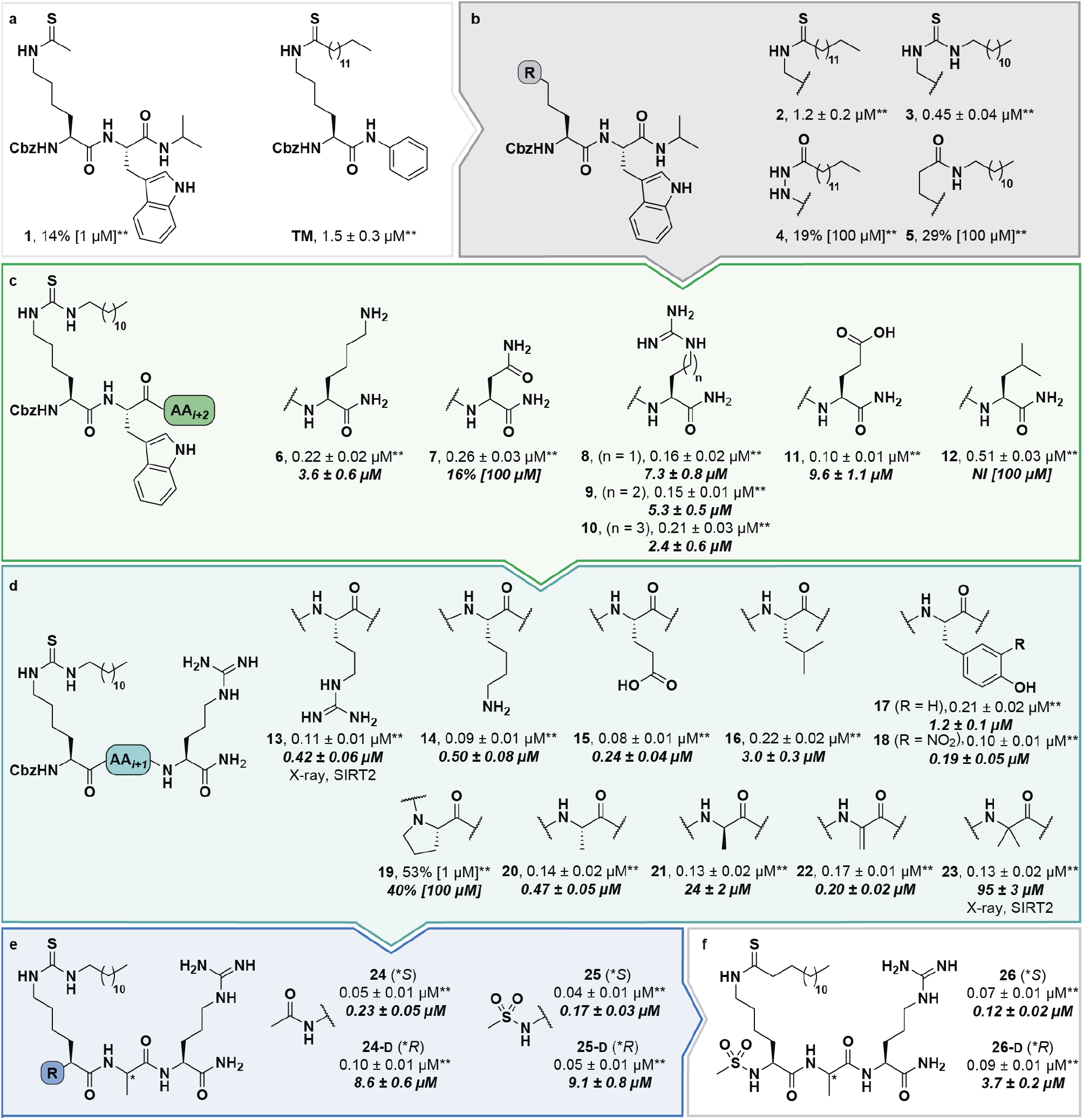
Structure–activity relationship of SIRT2 inhibitors. Potencies for inhibition of the deacylase activity of recombinant SIRT2 (100 nM) against QPKKac (shown with asterisks**) or ETDKmyr (shown in bold/italics) as substrate (50 μM) are given as mean IC_50_ values ± standard deviation (SD) or %-inhibition. (**a**) Lead compounds. (**b**) Amide isosteres. (**c**) Optimization of position *i*+2. (**d**) Optimization of position *i*+1. (**e**) Optimization of *N*-terminal group. (**f**) Final inhibitor series. Data are based on two individual experiments performed in duplicate. See the Supplementary Information (Scheme S1, Fig. S1–S3, and Table S2) for dose–response curves and selectivity profiling of additional inhibitors (**S17**–**S23**) against SIRT1–3. *A single asterisk denotes a stereogenic center.

**Fig. 2.**
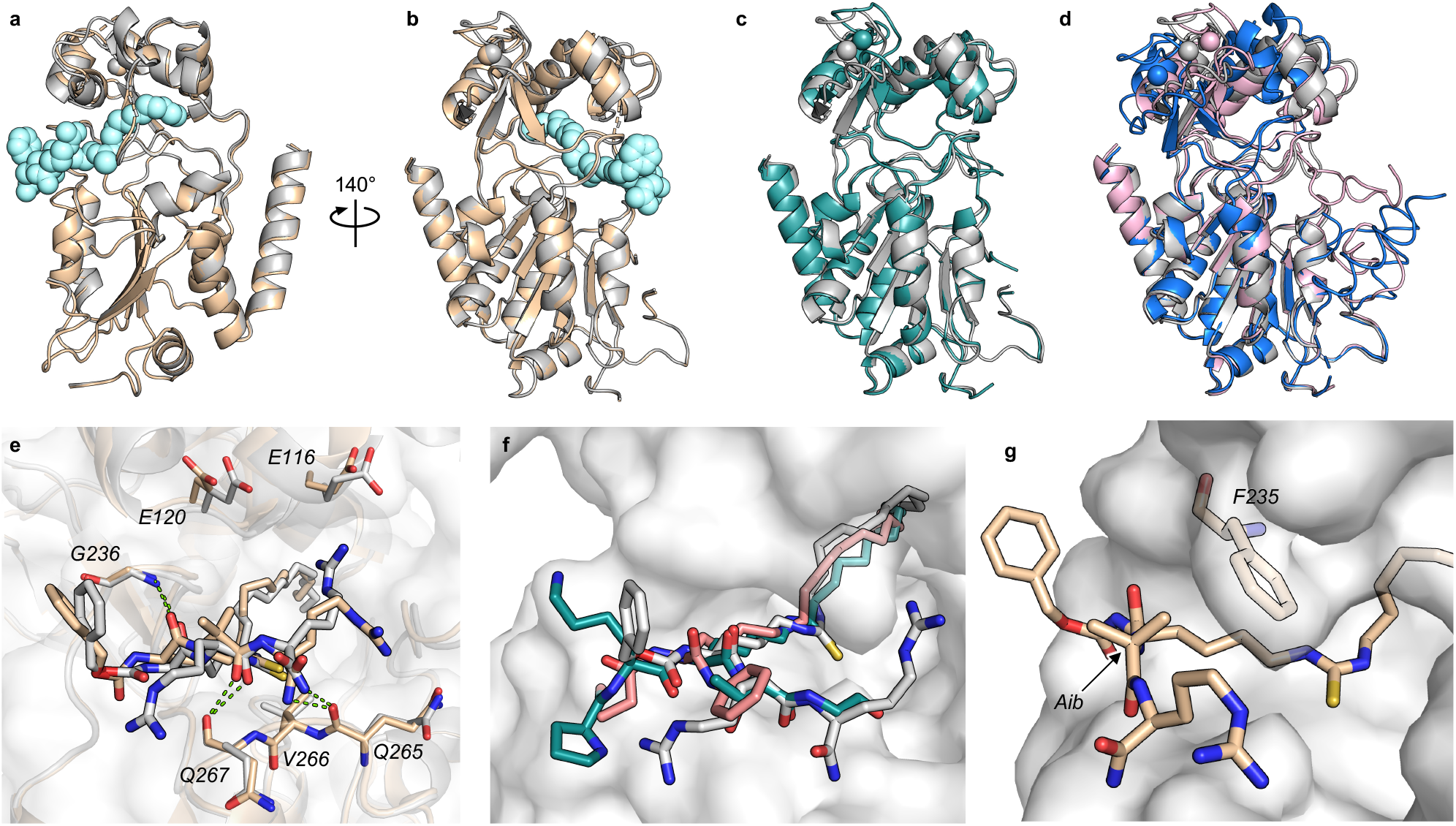
X-ray co-crystal structures of SIRT2 with compounds 13 and 23 bound. (**a**,**b**) Overall structure of SIRT2 co-crystallized with compounds **13** (gray) or **23** (tan). The ligand (**13**) is shown as light cyan spheres. (**c**) Overlay of the SIRT2:**13** structure (gray) with previously solved co-crystal structure of SIRT2 with the thiomyristoylated peptide substrate analog **BHJH-TM1** (PDB 4R8M; teal). (**d**) Overlay of the SIRT2:**13** structure (gray) with previously solved structure of SIRT2 apo form (PDB 3ZGO; blue) and SIRT2 in complex with **S2iL5** (PDB 4L3O; salmon). (**e**) Comparison of the exterior binding pocket of SIRT2 in complex with compound **13** (white) or **23** (tan). Dashed green lines highlight ligand–enzyme hydrogen bonding interactions. (**f**) Surface-view of SIRT2:**13** (white) superimposed with **BHJH-TM1** (teal) and “**glucose-TM**” (PDB 6NR0, pink). (**g**) Surface view of the exterior binding pocket of SIRT2:**23**.

Finally, our structures indicated that both L-configured and α-bis-substituted amino acids could be accommodated at the *i*+1 position (Fig. 2e and g), which was not reflected by the biochemical assay data. The structures also indicated high flexibility regarding the choice of side chains at both *i*+1 and *i*+2 positions, as no direct interactions with the enzyme were visible, which is in agreement with the obtained compound potencies. Most prominently, there was a complete lack of electrostatic interactions between the guanidium group at the *i*+2 position and E120 of SIRT2 (Fig. 2e–g), in contrast to what was observed for **S2iL5**. Again, this observation was in full agreement with the generally subtle differences observed in potency when mutating the *i*+2 position in our SAR. Finally, the structures indicated a significant degree of freedom regarding the choice of *N*-terminal substituents, as also found previously in our optimization of inhibitors of SIRT5.^58^ To address the importance of a select number of substituents at this position, we chose **21** (D-Ala) as the parent compound for the *N*-terminal SAR series, as the more potent compound **20** (L-Ala) already approached stoichiometric inhibition of demyristoylation. Thus, compounds **S5**–**S10**, **24-D**, **25-D** (Scheme 1 and S1) were evaluated for their ability to inhibit both deacetylation and demyristoylation, which revealed an improvement in potency for amides and sulfonamides compared to the carbamate (Cbz group), and flexibility in the degree of steric congestion as expected. To limit the hydrophobicity of our compounds, we therefore continued by synthesizing the L-Ala analogues of **24-D** and **25-D** to give **24** and **25**, respectively.

We then briefly revisited the importance of the ε-*N*-acyllysine substitution. Truncating the acyl chain decreased the potency dramatically and in a more severe manner than observed for small molecule ligand **TM**and its shorter chain analogs^24^ (**S11**–**S13**; Scheme S1). Additionally, introduction of oxygen atoms in the acyl chain to increase aqueous solubility greatly decreased the inhibitory activity (**S14**–**S16**, Scheme S1), again emphasizing the importance of a long hydrophobic PTM to maintain binding affinity. Finally, we reintroduced a thiomyristoyl group to furnish compounds **26** and **26-D** that could serve as an important tool compound for comparing differences in biological effects between thioamides and thioureas.^76^ Both compounds were equipotent to their thiourea homologues.

### Compound selectivity and inhibition kinetics

Despite the number of SIRT2 inhibitors previously reported, only few studies have addressed the inhibition of SIRT2 demyristoylase activity.^21,77,78^ Because *K*_m_ values for ε-*N*-myristoyllysine-based substrates are in the order of 100-fold lower than for corresponding acetylated substrates,^6,21,79^ only high-affinity inhibitors should be able to efficiently outcompete myristoylated substrates. Through our SAR study, we were able to inhibit SIRT2-mediated demyristoylation and we even achieved inhibitor potencies to an extent where the inhibitor–enzyme ratio in the demyristoylation assays approached stoichiometry. We therefore performed continuous, trypsin-coupled progression curve assays^21^ for selected compounds, which can report on the kinetics of inhibition. In addition, such assays may provide estimated *K*_i_ values for inhibitors that do not exhibit the “standard” fast-on/fast-off steady-state kinetics, which prohibits approximation of *K*_i_ values from end-point experiments, using the Cheng-Prusoff equation.

Compounds **TM**, **S2iL5**, **25**, **26**, and **26-D** were chosen for this kinetic evaluation. **TM** did not show inhibition of SIRT2 in this assay, even upon addition of DMSO to help solubilize the inhibitor (10% final concentration; Fig. 3a). Not surprisingly, due to the large peptide ring-size and presence of two Arg residues in the **S2iL5** macrocycle, we found this compound incompatible with the trypsin present in the continuous assays (data not shown). Compounds **25** and **26**, on the other hand, caused a substantial bending of the rate curves (Fig. 3b,c), suggesting slow, tight-binding inhibition kinetics.^80,81^ This was also the case for compound **26-D**; albeit, requiring substantially higher concentrations of the inhibitor (Fig. 3d), as would also be expected based on the end-point inhibition data (Scheme 1). For these three compounds, the apparent first-order rate constants to reach steady state (*k*_obs_) were calculated for each of the bending curves. Fitting the data from the two potent inhibitors (**25** and **26**) revealed hyperbolic relationships between *k*_obs_ and inhibitor concentration, which indicates that the inhibition follows mechanism B of slow-binding,^59^ where the initially formed enzyme–inhibitor complex (EI) undergoes a slow conformational change to give a more stable complex EI* (Fig. 3e). The 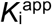 and residence time values could not be estimated for compounds **25** and **26**, as the *k*_−2_ rate constants approached zero in both experiments, revealing slow, tight-binding kinetics. On the other hand, the equilibrium constants for the first fast binding step (*K*_i,1_) were in the same range as the measured IC_50_ values from end-point experiments, correlating with the relatively slow *k*_2_ transition constants (Table S3). The less potent inhibitor **26-D** followed mechanism A of slow-binding, where a single slow binding step is detected (Fig. 3e) and here the estimation of the *k*_−1_ rate constant also approached zero, hampering the estimation of a 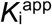 value. The obtained on-rate (*k*_1_), however, was relatively low, which correlates with its limited inhibitory potency.

**Fig. 3.**
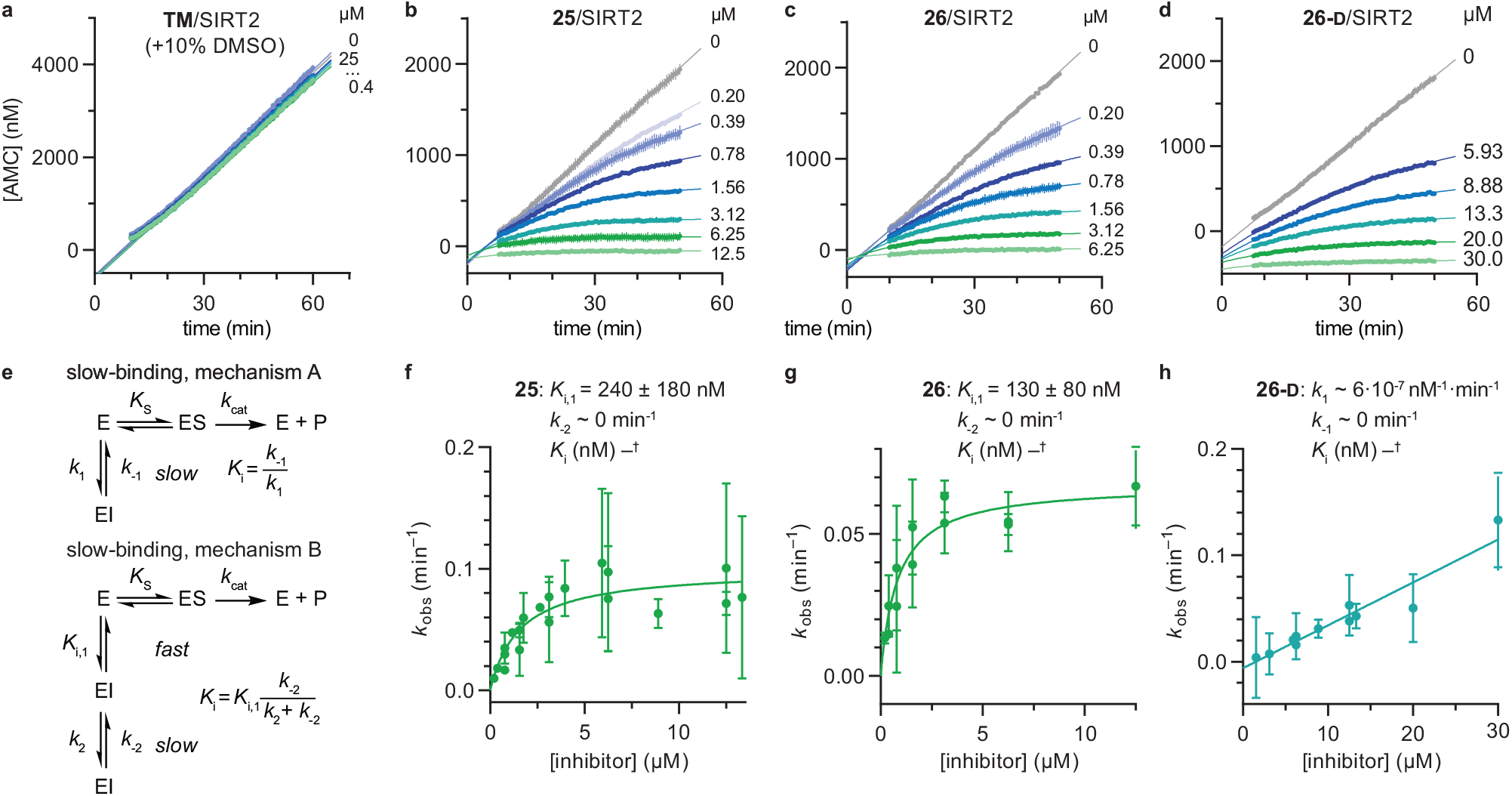
Kinetics of SIRT2 inhibition by compounds 25, 26, and 26-D. Sample rate experiment curves for compounds. (**a**) **TM**, (**b**) **25**, (**c**) **26**, and (**d**) **26-D**, with concentration of inhibitor for each experiment indicated on the right of the curve. (**e**) Common mechanisms of slow-binding inhibitor kinetics, with associated equilibrium and rate constants. Dependence of the first-order rate constant to reach steady state (*k*_obs_) on inhibitor concentration for compounds (**f**) **25** and (**g**) **26**, following mechanism B, and (**h**) **26-D**, which follows mechanism A. Continuous assays were performed with SIRT2 (20 nM), ETDKmyr (20 μM) as substrate, trypsin (20 ng/μL), and at designated inhibitor concentrations. See Table S3 for additional details and complete data fitting.

The potencies of our compounds were then benchmarked against a series of previously reported inhibitors applied as control compounds (Table 1). Compounds **25** and **26** showed superior potency against SIRT2 compared to **SirReal2**, **TM**, **tenovin-6** and **AGK-2**. The macrocycle **S2iL5** reached stoichiometric inhibition in the deacetylation assay but proved to be >10-fold less potent than compounds **25** and **26** when tested in the demyristoylation assay. In addition, no activity against demyristoylation could be recorded for any of the other control compounds at the highest concentration tested (Table 1). Because the most potent inhibitors reached stoichiometry in the SIRT2 deacetylation assay, determination of their selectivity indexes across SIRT1–3 was not possible. However, compounds **25** and **26** were substantially more selective towards SIRT2 over SIRT1 and SIRT3 than **S2iL5**, based on their higher potency against SIRT2 (>10-fold), combined with their higher IC_50_ values against SIRT3 and similar IC_50_ values against SIRT1 (Table 1). In addition, **25** and **26** did not inhibit SIRT5 or SIRT6 significantly (Table S4).

**Table 1.**
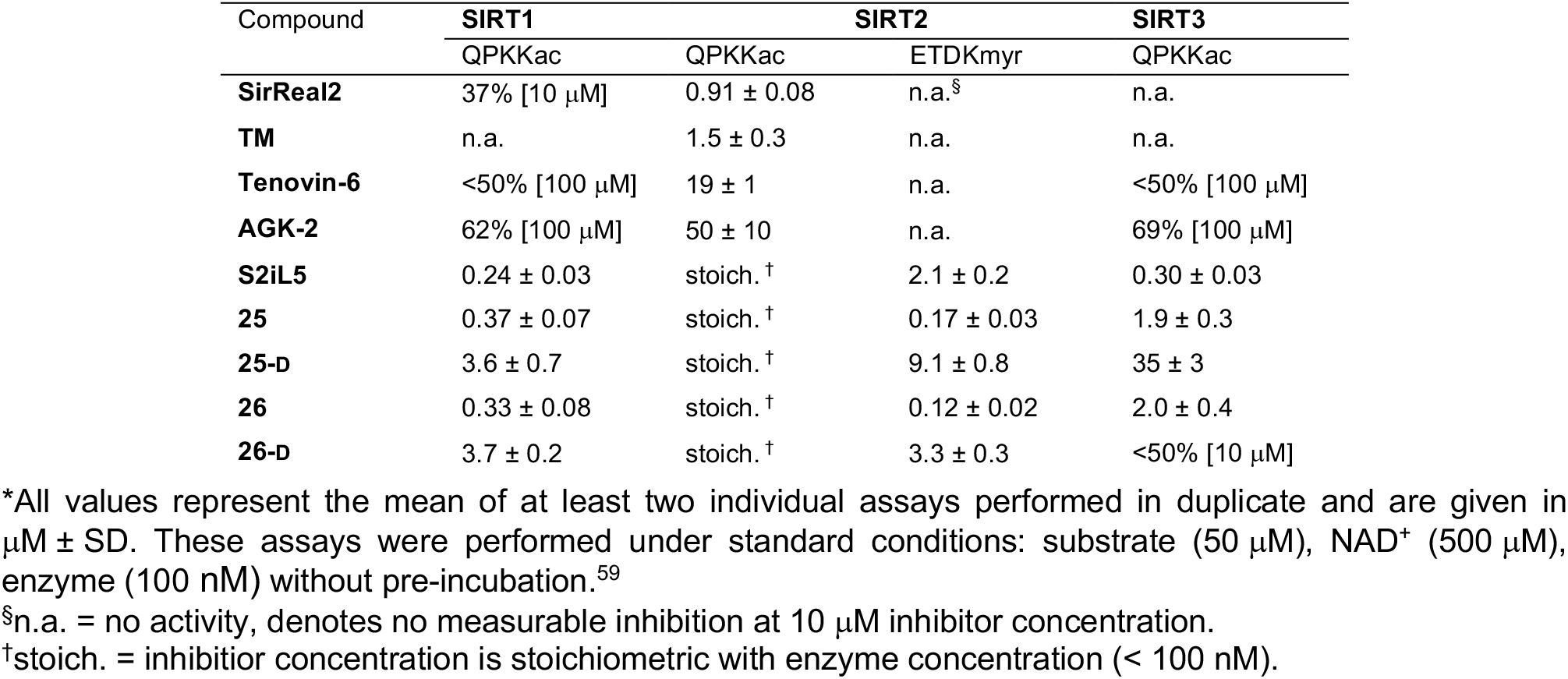
IC_50_ values (μM) or %-inhibition data for final compounds and control compounds*.

Next, we investigated whether the observed inhibition of enzymatic deacylation could be recapitulated with a non-fluorogenic substrate.^82^ To address this, we chose a label-free HPLC-based assay, monitoring inhibition of histone 3 lysine 9 (H3K9) deacetylation, applying a dodecameric H3K9ac-containing peptide (Fig. 4a). We found the IC_50_ values of compounds **26**, **26-D** and **S2iL5** to be in the nanomolar range, with **26** and **26-D** being approximately 3 times more potent than **S2iL5** and approaching stoichiometric inhibition in this assay as well (Fig. 4a and Table S5). Furthermore, **TM** showed more than three orders of magnitude lower potency than **26**(IC_50_ ~40 μM and 0.016 ± 0.004 μM, respectively) when tested in this HPLC-based assay. This is not in accordance with a recent study by Lin,^78^ but corresponds well with another report where **TM**is used as a reference compound.^77^ This may again be a result of the challenging handling of this compound, due to its limited aqueous solubility, which is in accordance with calculated logP and polar surface area (PSA) values, compared to the compounds developed here (Table S6). However, such discrepancies also highlight that IC_50_ values are highly dependent on the experimental conditions applied, which calls for determination in several different assay formats.^83^

**Fig. 4.**
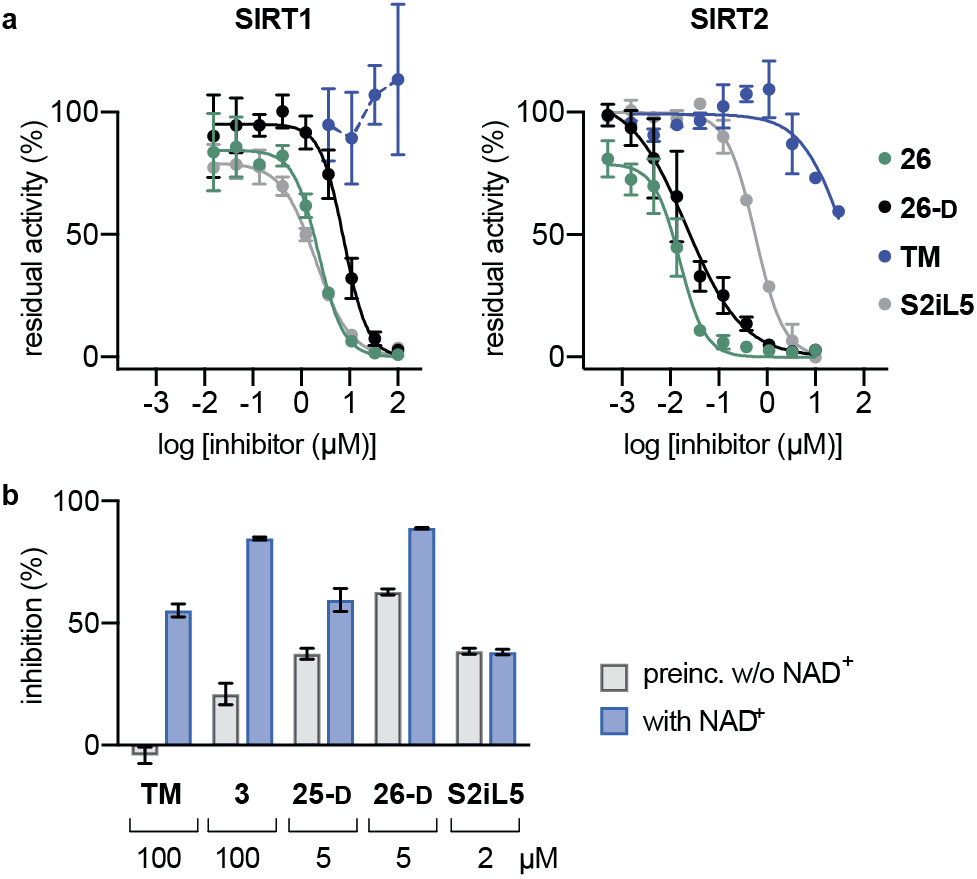
Pre-incubation and HPLC-based sirtuin assays. (**a**) Inhibition of SIRT1 (100 nM) and SIRT2 (100 nM)-mediated deacetylation of the H3K9ac residue in a dodecameric peptide (50 μM) determined by HPLC. Error bars represent mean ± SEM of at least 2 independent experiments. See Table S4 for IC_50_ values against sirtuin mediated H3K9ac deacetylation. (**b**) Pre-incubation (30 min), with or without NAD^+^ (500 μM) for selected compounds against SIRT2 (100 nM) demyristoylation using ETDKmyr (50 μM) as substrate. Mean ± SEM based on at least 2 independent experiments performed in duplicate.

Finally, we also performed assays with pre-incubation of selected inhibitors and SIRT2 with or without addition of NAD^+^, to address whether binding of NAD^+^ to the enzyme is involved in the inhibition mechanism. For compounds **TM**, **3**, **25-D** and **26-D**, substantially increased %-inhibition values were recorded upon pre-incubation including NAD^+^, which suggests a mechanism-based slow-binding mechanism involving the formation of stalled intermediate with adenosine diphosphate ribose (ADPR).^52^ The only tested compound that did not exhibit this behavior was **S2iL5** (Fig. 4b).^58,84^

### Compound stability

Over the last decade, the considerable efforts in development of efficacious sirtuin modulators has furnished a number of potent inhibitors of commercial interest, of which some have entered clinical trials to document safety and efficacy in humans.^85–90^ However, there is a lack of studies addressing the chemical and metabolic stability of sirtuin inhibitors, except for one study of pronase susceptibility of linear vs. cyclic sirtuin inhibitors.^91^ We recently addressed how stability differed greatly between short chain thioamide and thiourea compounds,^76^ which prompted us to investigate stability of selected inhibitors developed in this study. We first addressed the stability of **25-D** (thiourea) and **26-D** (thioamide) in assay buffer or growth cell medium supplemented with fetal bovine serum (FBS). Both compounds showed no sign of degradation in assay buffer for up to 10 days at 37 °C (Fig. S5) and were largely intact in growth medium for 24 h at 37 °C (Fig. S6a). Next, we investigated the stability in human serum, which, in agreement with our previous observations,^76^ revealed that compound half-lives (*t*_½_) of thioamides (**TM**, **2**, **26** and **26-D**) were significantly higher than their thiourea counterparts (**3**, **25** and **25-D**), with the latter series being almost fully degraded in less than two hours (Fig. 5a). Additionally, we measured the stability of three known SIRT2 inhibitors for comparison. The compounds **tenovin-6** and **S2iL5** were rapidly degraded, whereas the small molecule inhibitor **SirReal2**exhibited superior stability with no degradation even after 24 h (Fig. S6c).

**Fig. 5.**
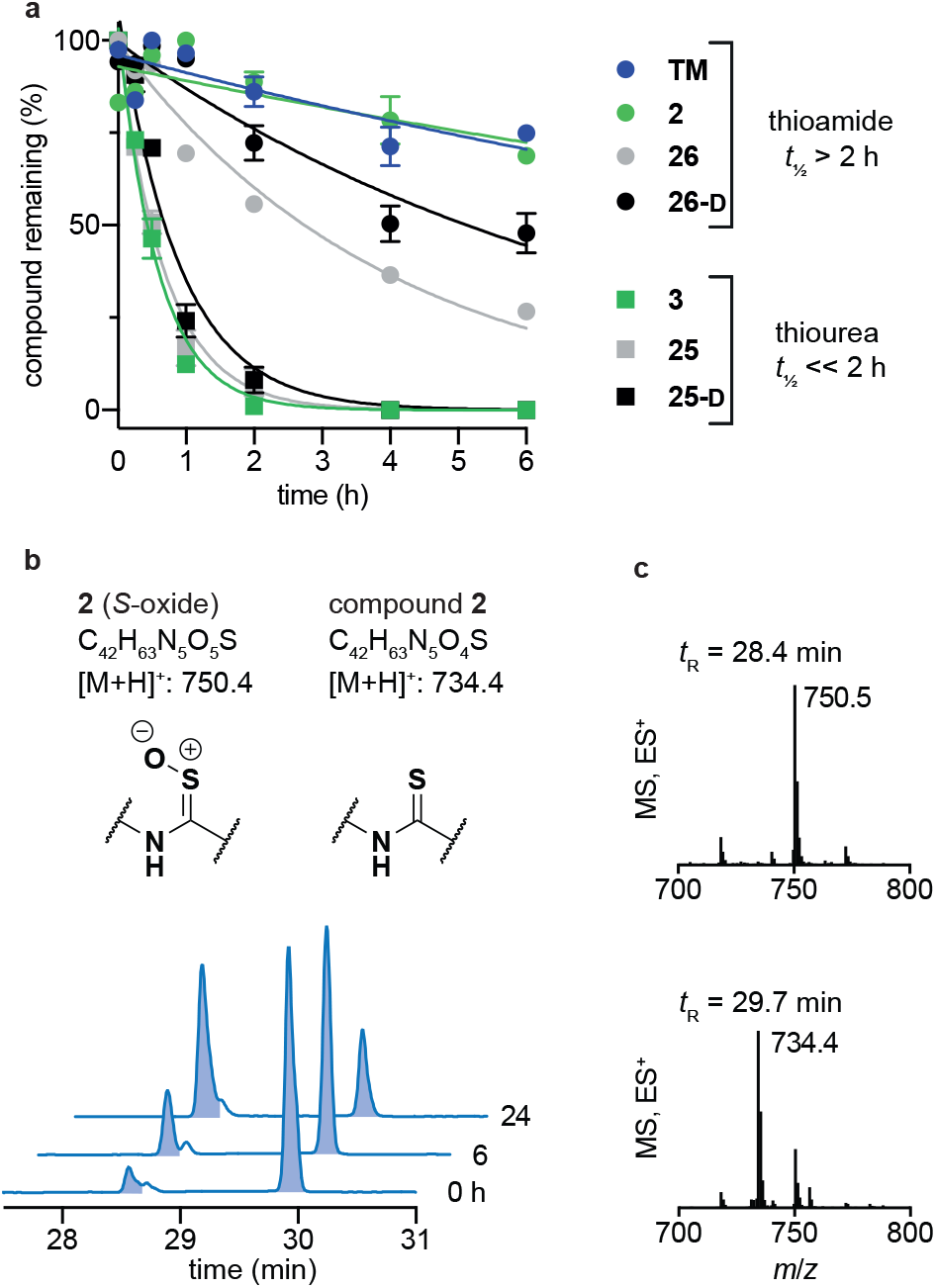
Stability assays in human serum. (**a**) Stability assays of selected compounds in human male serum. (**b**) UV (A_280_) and (**c**) TIC (ES) chromatograms and mass spectra at relevant time points of compound **2** monitoring degradation in serum. Data are shown as mean values relative to the peak intensity at *t* = 0 h ± SEM, n = 2–3. For *t*_½_ values, see Table S7.

The degradation of compounds **2**(thioamide) and **3**(thiourea) could be scrutinized in more detail by UPLC-MS analysis, because they contain tryptophan as a suitable chromophore. We found that the thioamide (**2**) was converted to a species with [M+16]^+^ (*m*/*z*) to a significant extent (Fig. 5b). Because thioamide *S*-oxides are known oxidative metabolites, we hypothesized that this is the major degradation product (Fig. 5b).^92,93^ The rapid degradation of the corresponding thiourea (**3**), on the other hand, led to large number of species that could not be structurally determined (Fig. S7). Release of hydrogen sulfide from both thioamide and thiourea motifs constitutes another possible mechanism of degradation.^94^ These results suggest that enzymes other than peptidases (*i.e.* redox active enzymes) may play a significant role in compound turnover, which raises the question whether these compounds would stay intact *in vivo*. However, the low degree of degradation observed for the compounds in both assay buffer and cell growth medium with FBS render both thiourea and thioamide compounds amenable as probes for *in vitro* enzymatic assays and cell culture experiments.

### Target engagement of SIRT2 inhibitors in cells

With our series of potent inhibitors in hand, we then assessed the cytotoxicity of a selection of >15 compounds, including control inhibitors **TM** and **SirReal2**, in cell culture experiments. We chose a panel of four cell lines, including immortalized human epithelial kidney cells (HEK293T), solid cervical cancer cells (HeLa), breast cancer cells (MCF-7), and T lymphocytes (Jurkat). Despite strong potency against SIRT2 *in vitro*, the compounds displayed lower degree of cytotoxicity than **TM** against all tested cell lines (Fig. S4 and Table S8), begging the question whether the physicochemical properties of **TM** (Table S6) would render it more efficacious in cell-based assays. We therefore investigated the target engagement and selectivity across SIRT1–3, by performing cellular thermal shift assays in HEK293T cells, using Western blots for the subsequent analysis (Fig. 6a and Fig. S9).61,62 Upon treatment with compound **26**, **26-D** or **TM** for 1 hour, a drastic increase in SIRT2 melting temperature was observed for all three compounds compared to the DMSO control. Particularly, compound **26** was found to give rise to >15 °C thermal stabilization of SIRT2 at concentrations down to 10 nM and no stabilization of SIRT1 and SIRT3 was observed under the same conditions (Fig. S9), verifying the selectivity of our compounds *in cellulo*. For **TM**, treatment with 10 μM compound (1000-fold higher concentration) produced a slightly less pronounced thermal shift, rendering our optimized compound (**26**) substantially more efficient in this assay. All tested compounds **26**, **26-D** and **TM** exhibited excellent selectivity for SIRT2 over SIRT1 and SIRT3 (Fig. S9). Thus, these results suggest cellular thermal shift assays as an excellent complementary method to evaluate subtype selectivity across sirtuins along with *in vitro* profiling against recombinant enzymes.

**Fig. 6.**
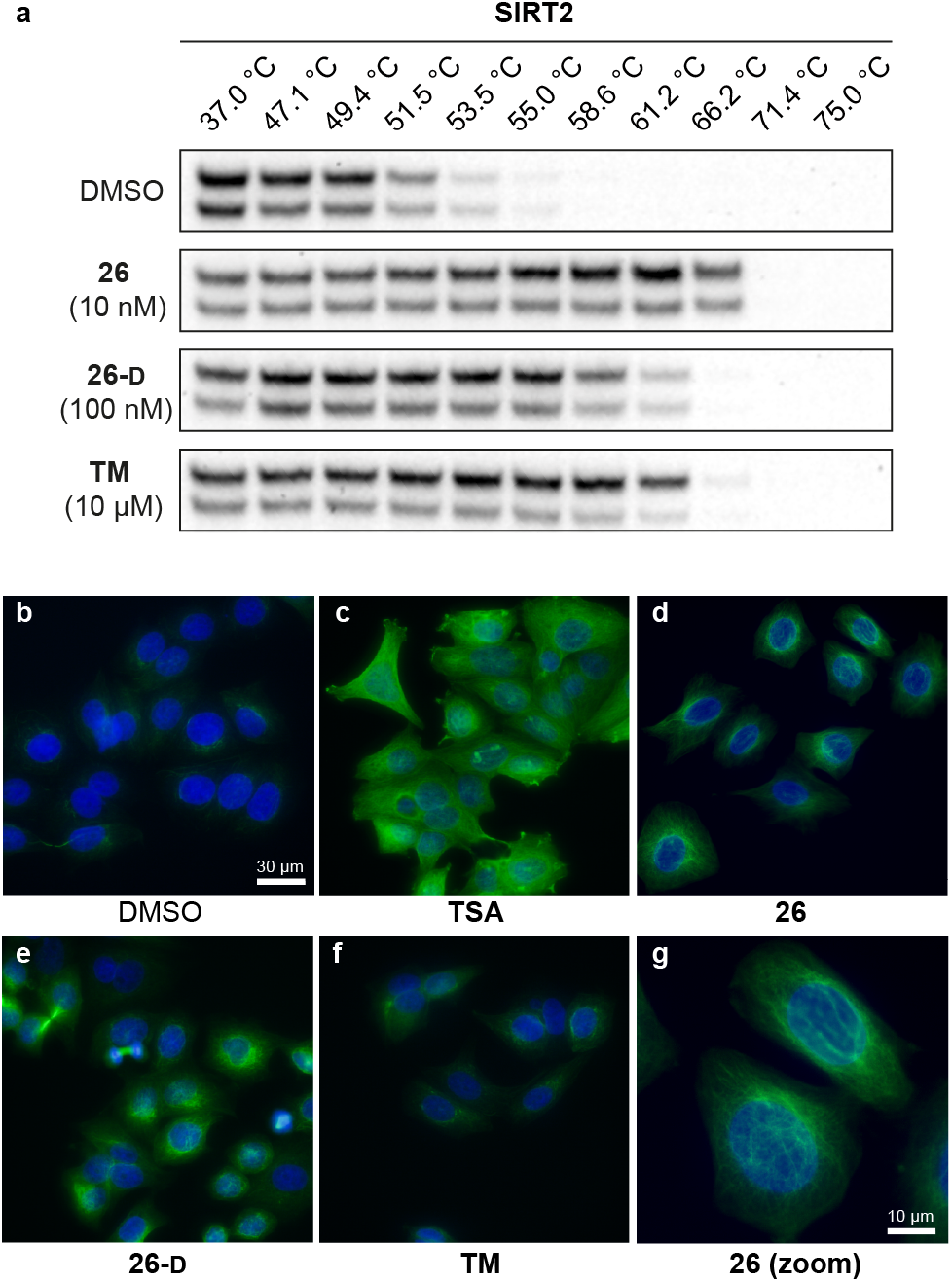
Cellular target engagement and SIRT2i effect on perinuclear α-tubulin acetylation. (**a**) Cellular thermal shift of SIRT2 in HEK293T cells subjected to 1 h treatment with inhibitor at designated concentrations or DMSO (vehicle). Please see Fig. S8 for complete dataset against SIRT1–3 (n = 3). (**b**–**g**) Immunofluorescence images (40×) of MCF-7 cells subjected to 6 h treatment with inhibitor [**TSA** (5 μM), **26** (5 μM), **TM** (25 μM), **26-D** (25 μM)] or DMSO (vehicle). (**g**) Zoomed image for compound **26**. DAPI (blue, nuclear counterstain) and Ac-α-tubulin (green). For additional images (single-filtered and zoomed) for all tested compounds, please see Fig. S9. The data are representative images from two individual experiments.

### Modulation of α-tubulin acetylation levels

To investigate whether the compounds affected the protein acetylation level in cells, we then incubated HeLa and MCF-7 cells under a series of conditions and performed Western blots on cell lysates to monitor changes in acetylation levels of α-tubulin, which is a reported target of SIRT2.^16^ Unfortunately, we failed to observe any reproducible changes in acetylation levels upon treatment with inhibitor or **TM** as the positive control,^24^ including experiments where SIRT2 was overexpressed or HDAC6 was inhibited by co-treatment with trichostatin A (**TSA**)^47,95^ (data not shown). Part of the explanation for these observations may be the differences in acetylation in asynchronous and mitotic cells.^17^ Instead, we applied immunofluorescence as a readout to detect changes in acetylated α-tubulin levels in MCF-7 cells. Upon treatment with inhibitors (5 or 25 μM for 6 hours), we were pleased to observe an increase in cellular fluorescence levels, particularly around the perinuclear microtubules,^35^ which is reported to be the main target of SIRT2 inhibition in cells.^96^ Treatment with the potent HDAC6 inhibitor **TSA**, on the other hand, resulted in hyperacetylation of α-tubulin throughout the entire cell (Fig. 6c). Thus, the SIRT2-mediated effect on α-tubulin acetylation is mainly perinuclear, compared to the global effects mediated by HDAC6 deacetylation,^96^ which may help explain why Western blotting for total α-tubulin acetylation was not an ideal measure of the effect of SIRT2 inhibition in our hands. This is in agreement with previous findings.^35^ However, inhibition of SIRT2 in primary T cells was recently shown to cause large increase in acetylated α-tubulin levels as determined by Western blotting, suggesting that cellular effects are highly cell type dependent.^97^

### Inhibition of breast cancer cell migration

Due to the effects of our compounds on the acylation of the cytoskeleton and the validated target engagement, we next investigated the role of SIRT2 inhibition on breast cancer cell motility. In previous reports, SIRT2 has been implicated in the migration and invasion of gastric, colon, lung, and liver cancers.^98–101^ Therefore we investigated the effect of our lead compounds on cell motility in MCF-7 cells. The effects of **26**, **26-D**, and **TM** inhibitors were investigated in a scratch assay (wound-healing assay) and all inhibitors decreased the degree of migration compared to the vehicle-treated control (Fig. 7a,b). Interestingly, knockdown of SIRT2 was recently shown to have the opposite effect on migration of A549 cells overexpressing RalB in culture,^101^ indicating that pharmacological modulation of the SIRT2 activity may provide a different outcome from enzyme knockdown/overexpression or that the effects are cell type specific. Additional experiments, including comparison with SIRT2 mediated knockdown, are required to investigate this in detail.

**Fig. 7.**
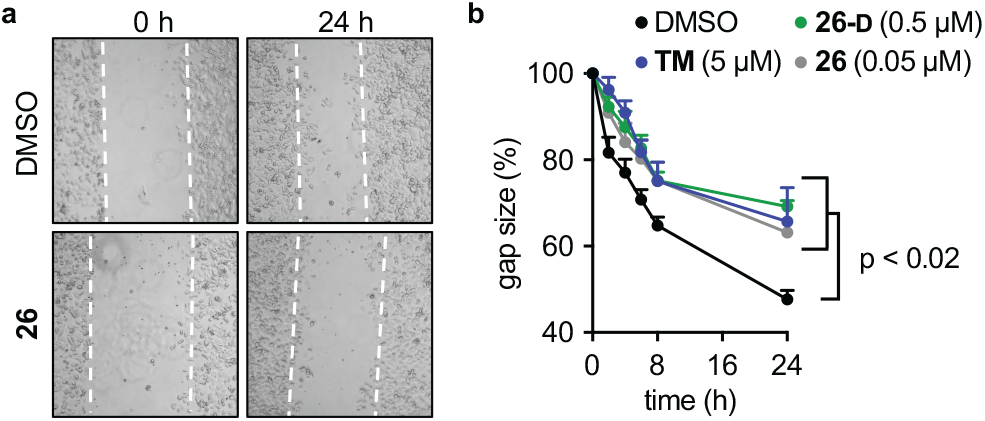
SIRT2 inhibition elicits decreased breast cancer cell motility. (**a**) Representative images from a scratch assay in MCF-7 cells at time points 0 and 24 h (2.5× magnification). (**b**) Quantification of the gap size at different time points treated with inhibitors (**TM**, **26**, and **26-D**) at denoted concentrations relative to control (1% DMSO) treated cells ± SEM. Data are shown as mean values relative to control (1% DMSO) treated cells ± SEM, n = 4. Statistical differences from control treated cells were determined by ordinary (multiple comparison) one-way ANOVA. Statistical p-values compared to DMSO control: **TM** = 0.0042; **26** < 0.0001; **26-D** = 0.0134.

## Discussion

Here, we report an elaborate SAR study, including X-ray co-crystal structures with intermediary compounds bound to SIRT2 to provide structural rationalization of the binding interactions. This furnished the most potent inhibitors of the SIRT2 histone deacetylase enzyme reported to date, which at the same time exhibit improved selectivity for the target compared to structurally similar isoforms, SIRT1 and SIRT3. The compounds were able to inhibit the demyristoylase activity of SIRT2 with unprecedented potencies, which is challenging due to the low *K*_m_ values of Kmyr-containing substrates. This further enabled us to glean insight into the binding kinetics of SIRT2 inhibition by applying continuous assay formats, which has not been achieved with SIRT2 deacetylation assays.^102^ Standard end-point dose–response assays do not take into account the kinetic behavior of inhibitors,^80,81^ which may give rise to IC_50_ values that vary substantially with specific assay conditions, especially if the inhibitor does not exhibit standard fast-on/fast-off kinetics.^103^ It is well documented that mechanism-based thioamide- and thiourea-containing inhibitors may form stalled intermediates with ADPR in the enzyme active sites, which can affect binding kinetics.^58,104,105^ Indeed, evaluation of compounds **25**, **26**, and **26-D** in the continuous assays revealed slow, tight-binding kinetics with two different binding mechanisms. To our satisfaction, the estimated rate constants for the formation of inhibitor–enzyme complexes correlated well with the data obtained in our two other assay formats. We thus provide data from three different assay formats that are all are in agreement.

Furthermore, we provide measures of stability in buffer, cell culture medium, and human serum, which taken together will inform the development of future generations of mechanism-based inhibitors of sirtuins. Thiourea-containing compounds are more readily synthesized than thioamide counterparts, but they proved significantly less stable in human serum, highlighting probe **26** as the preferred choice for biological applications. Due to the similar potencies of inhibitors containing these two functional groups, however, we find the thiourea-based compounds useful during SAR studies, focusing on the optimization of compound potency and selectivity *in vitro*. At a later stage, the resulting lead compounds can then be prepared as their thioamide counterparts for biological studies as demonstrated in this work.

Compound **26**, and also **26-D**, exhibited cellular activity by causing increased levels of perinuclear acetylated α-tubulin in MCF-7 cells in culture, as demonstrated by immunofluorescence experiments. Western blot analyses failed to demonstrate a reproducible effect on overall α-tubulin acetylation in a range of experiments and conditions in our hands, including SIRT2 overexpression and/or treatment with additional HDAC inhibitors. Further evidence for target engagement with SIRT2 in cultured cells was achieved by performing cellular thermal shift assays in HEK293T cells, which showed substantial protein stabilization at just 10 nM inhibitor concentration for compound **26**. Finally, we show that migration of cultured MCF-7 cells is reduced by our SIRT2 inhibitors, which could have implications for inhibiting cancer metastasis.

Thus, the probes reported herein provide an attractive alternative to previously developed SIRT2 inhibitors by exhibiting potent inhibition of SIRT2 and a high degree of selectivity over SIRT1, 3, 5, and 6. Furthermore, the compounds exhibit target engagement and activity in cells, stability in serum, and aqueous solubility that should allow for intravenous dosing of animals without elaborate drug formulation. This combination of features is unique among SIRT2 inhibitors and we expect that the developed probes will therefore be valuable in the continued investigation of the function of SIRT2 as well as the development of novel therapeutics that target this enzyme.

## Abbreviations

ADPR: adenosine diphosphate ribose
Aib: 2-aminoisobutyric acid
AMC: 7-amino-4-methylcoumarin
BSA: bovine serum albumin
Cbz: benzyloxycarbonyl
DAPI: 4’,6’-diamidino-2-phenylindole
DMSO: dimethylsulfoxid
ES: electrospray
FBS: fetal bovine serum
HDAC: histone deacetylase
HEK293T: human embryonic kidney 293 cells
HeLa: Henrietta Lacks (human cervical cancer cells)
HEPES: (4-(2-hydroxyethyl)-1-piperazineethanesulfonic acid)
GFP: green fluorescent protein
GI_50_: half growth inhibitory concentration
HPLC: high-performance liquid chromatography
IC_50_: half maximal inhibitory concentration
LC: liquid chromatography
MCF-7: Michigan cancer foundation
MEM: minimum essential medium eagle
NAD: nicotinamide adenine dinucleotide
NAM: nicotinamide
NMR: nuclear magnetic resonance
PBS: phosphate-buffered saline
PDB: Protein Data Bank
PSA: polar surface area
PTM: post-translational modification
RT: room temperature
SAR: structure-activity relationship
SD: standard deviation
SEM: standard error of the mean
SIRT: sirtuin
SPPS: solid-phase peptide synthesis
TIC: total ion chromatogram
TSA: trichostatin A
HPLC: high performance liquid chromatography.

## Acknowledgements

We thank Dr. Iacopo Galleano, and Dr. Martin Roatsch for donation of building blocks, as well as Julie E. Bolding, Kathrin S. Troelsen and Katrine Krydsfeldt for assistance with cell culture experiments. We thank Professor Thue W. Schwartz for access to equipment for performing immunofluorescence and wound-healing experiments. The X-ray crystallography was performed under the approval of the Photon Factory Program Advisory Committee (Proposal No. 2017G662 and 2019G669 and PSI Proposal No. 20181219 and 20181299). This work was supported by JSPS KAKENHI (JP19H05640; M.Y.), the Lundbeck Foundation (Running cost grant R289-2018-2074; CAO), the Carlsberg Foundation (2013-01-0333, CF15-011, and CF18-0442; C.A.O.), the Novo Nordisk Foundation (NF17OC0029464; C.A.O.), and the European Research Council (ERC-CoG-725172–*SIRFUNCT*; C.A.O.). We thank the COST-Action CM1406 (EPICHEMBIO) for support.

## Author contributions

A.L.N, N.R. and C.A.O conceptualized the study; A.L.N., N.R., M.F. and A.L. performed the chemical synthesis. A.L.N., N.R. and C. M.-Y. performed the biochemical characterization; A.L.N., K.L. and M.B. performed the cell-based assays; A.L.N. performed the formal analysis of raw data; N.K. obtained the crystals and solved the structures; A.L.N. and C.A.O wrote the original draft of the manuscript, and all authors reviewed and edited the final manuscript; C.A.O. acquired funding. A.S.M., M.Y., and C.A.O. supervised the study.

## Additional information

### Accession codes

X-ray diffraction data, coordinates, and structure factors for the X-ray crystal structures are deposited with the PDB (http://www.wwpdb.org) (under the accession numbers 7BOS (SIRT2:**13**) and 7BOT (SIRT2:**23**).

---

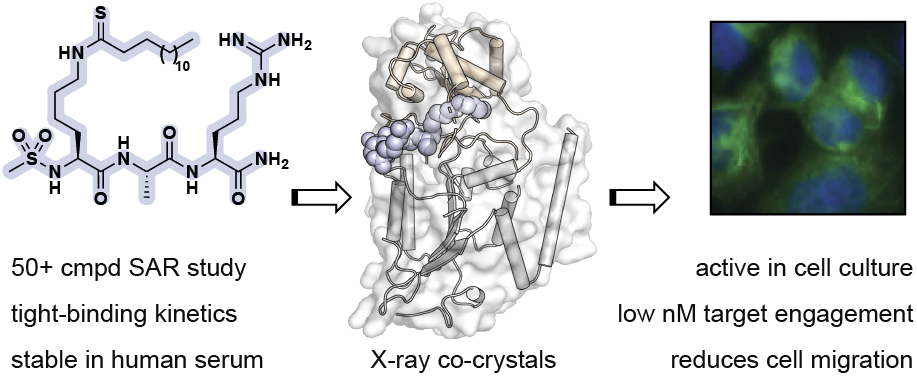
Graphical abstract.

## Notes and references

1 P. Bheda, H. Jing, C. Wolberger and H. Lin, Annu. Rev. Biochem., 2016, 85, 405–429.

2 N. Rajabi, I. Galleano, A. S. Madsen and C. A. Olsen, in Prog. Mol. Biol .Transl. Sci., Academic Press, 2018, vol. 154, pp. 25–69.

3 J. Du, Y. Zhou, X. Su, J. J. Yu, S. Khan, H. Jiang, J. Kim, J. Woo, J. H. Kim, B. H. Choi, B. He, W. Chen, S. Zhang, R. A. Cerione, J. Auwerx, Q. Hao and H. Lin, Science, 2011, 334, 806–809.

4 H. Jiang, S. Khan, Y. Wang, G. Charron, B. He, C. Sebastian, J. Du, R. Kim, E. Ge, R. Mostoslavsky, H. C. Hang, Q. Hao and H. Lin, Nature, 2013, 496, 110–113.

5 J. L. Feldman, J. Baeza and J. M. Denu, J. Biol. Chem., 2013, 288, 31350–31356.

6 Y. Bin Teng, H. Jing, P. Aramsangtienchai, B. He, S. Khan, J. Hu, H. Lin and Q. Hao, Sci. Rep., 2014, 5, 8529.

7 M. Tan, C. Peng, K. A. Anderson, P. Chhoy, Z. Xie, L. Dai, J. Park, Y. Chen, H. Huang, Y. Zhang, J. Ro, G. R. Wagner, M. F. Green, A. S. Madsen, J. Schmiesing, B. S. Peterson, G. Xu, O. R. Ilkayeva, M. J. Muehlbauer, T. Braulke, C. Mühlhausen, D. S. Backos, C. A. Olsen, P. J. McGuire, S. D. Pletcher, D. B. Lombard, M. D. Hirschey and Y. Zhao, Cell Metab., 2014, 19, 605–617.

8 R. A. A. Mathias, T. M. M. Greco, A. Oberstein, H. G. G. Budayeva, R. Chakrabarti, E. A. A. Rowland, Y. Kang, T. Shenk and I. M. M. Cristea, Cell, 2014, 159, 1615–1625.

9 A. S. Madsen and C. A. Olsen, Angew. Chem. Int. Ed., 2012, 51, 9083–9087.

10 P. Aramsangtienchai, N. A. Spiegelman, B. He, S. P. Miller, L. Dai, Y. Zhao and H. Lin, ACS Chem. Biol., 2016, 11, 2685–2692.

11 W. Wei, X. Liu, J. Chen, S. Gao, L. Lu, H. Zhang, G. Ding, Z. Wang, Z. Chen, T. Shi, J. Li, J. Yu and J. Wong, Cell Res., 2017, 27, 898–915.

12 Z. Kutil, Z. Novakova, M. Meleshin, J. Mikesova, M. Schutkowski and C. Barinka, ACS Chem. Biol., 2018, 13, 685–693.

13 C. Moreno-Yruela, I. Galleano, A. S. Madsen and C. A. Olsen, Cell Chem. Biol., 2018, 25, 849–856.

14 J. Cao, L. Sun, P. Aramsangtienchai, N. A. Spiegelman, X. Zhang, W. Huang, E. Seto and H. Lin, Proc. Natl. Acad. Sci. U. S. A., 2019, 116, 5487–5492.

15 C. A. Olsen, ChemMedChem, 2014, 9, 434–437.

16 B. J. North, B. L. Marshall, M. T. Borra, J. M. Denu and E. Verdin, Mol. Cell, 2003, 11, 437–444.

17 T. Nagai, M. Ikeda, S. Chiba, S. I. Kanno and K. Mizuno, J. Cell Sci., 2013, 126, 4369–4380.

18 S. C. Dryden, F. A. Nahhas, J. E. Nowak, A.-S. Goustin and M. A. Tainsky, Mol. Cell. Biol., 2003, 23, 3173–3185.

19 W. Li, B. Zhang, J. Tang, Q. Cao, Y. Wu, C. Wu, J. Guo, E.-A. Ling and F. Liang, J. Neurosci., 2007, 27, 2606–2616.

20 A. S. Madsen, C. Andersen, M. Daoud, K. A. Anderson, J. S. Laursen, S. Chakladar, F. K. Huynh, A. R. Colaço, D. S. Backos, P. Fristrup, M. D. Hirschey and C. A. Olsen, J. Biol. Chem., 2016, 291, 7128–7141.

21 I. Galleano, M. Schiedel, M. Jung, A. S. Madsen and C. A. Olsen, J. Med. Chem., 2016, 59, 1021–1031.

22 H. Huang, D. Zhang, Y. Wang, M. Perez-Neut, Z. Han, Y. G. Zheng, Q. Hao and Y. Zhao, Nat. Commun., 2018, 9, 3374.

23 A. Chalkiadaki and L. Guarente, Nat. Rev. Cancer, 2015, 15, 608–624.

24 H. Jing, J. Hu, B. He, Y. L. Negrón Abril, J. Stupinski, K. Weiser, M. Carbonaro, Y. L. Chiang, T. Southard, P. Giannakakou, R. S. Weiss and H. Lin, Cancer Cell, 2016, 29, 297–310.

25 G. Donmez and T. F. Outeiro, EMBO Mol. Med., 2013, 5, 344–352.

26 G. Donmez, Curr. Drug Targets, 2013, 14, 644–647.

27 B. C. Smith and J. M. Denu, Biochemistry, 2006, 45, 272–282.

28 R. M. de Oliveira, J. Sarkander, A. G. Kazantsev and T. F. Outeiro, Front. Pharmacol., 2012, 3, 82.

29 P. Gomes, T. Fleming Outeiro and C. Cavadas, Trends Pharmacol. Sci., 2015, 36, 756–768.

30 T. F. Outeiro, E. Kontopoulos, S. M. Altmann, I. Kufareva, K. E. Strathearn, A. M. Amore, C. B. Volk, M. M. Maxwell, J.-C. Rochet, P. J. McLean, A. B. Young, R. Abagyan, M. B. Feany, B. T. Hyman and A. G. Kazantsev, Science, 2007, 317, 516–519.

31 G. Eren, A. Bruno, S. Guntekin-Ergun, R. Cetin-Atalay, F. Ozgencil, Y. Ozkan, M. Gozelle, S. G. Kaya and G. Costantino, J. Mol. Graph. Model., 2019, 89, 60–73.

32 N. A. Spiegelman, J. Y. Hong, J. Hu, H. Jing, M. Wang, I. R. Price, J. Cao, M. Yang, X. Zhang and H. Lin, ChemMedChem, 2019, 14, 744–748.

33 T. F. S. Ali, H. I. Ciftci, M. O. Radwan, R. Koga, T. Ohsugi, Y. Okiyama, T. Honma, A. Nakata, A. Ito, M. Yoshida, M. Fujita and M. Otsuka, Bioorg. Med. Chem., 2019, 27, 1767–1775.

34 K. Yamagata, Y. Goto, H. Nishimasu, J. Morimoto, R. Ishitani, N. Dohmae, N. Takeda, R. Nagai, I. Komuro, H. Suga and O. Nureki, Structure, 2014, 22, 345–352.

35 T. Rumpf, M. Schiedel, B. Karaman, C. Roessler, B. J. North, A. Lehotzky, J. Ovadi, K. I. Ladwein, K. Schmidtkunz, M. Gajer, M. Pannek, C. Steegborn, D. A. Sinclair, S. Gerhardt, J. Ovadí, M. Schutkowski, W. Sippl, O. Einsle and M. Jung, Nat. Commun., 2015, 6, 6263.

36 M. Schiedel, T. Rumpf, B. Karaman, A. Lehotzky, J. Oláh, S. Gerhardt, J. Ovádi, W. Sippl, O. Einsle and M. Jung, J. Med. Chem., 2016, 59, 1599–1612.

37 P. Mellini, Y. Itoh, H. Tsumoto, Y. Li, M. Suzuki, N. Tokuda, T. Kakizawa, Y. Miura, J. Takeuchi, M. Lahtela-Kakkonen and T. Suzuki, Chem. Sci., 2017, 8, 6400–6408.

38 N. Kudo, A. Ito, M. Arata, A. Nakata and M. Yoshida, Philos. Trans. R. Soc., 2018, B373, 20170070.

39 L.-L. Yang, H.-L. Wang, L. Zhong, C. Yuan, S.-Y. S. Liu, Z.-J. Yu, S.-Y. S. Liu, Y.-H. Yan, C. Wu, Y. Wang, Z. Wang, Y. Yu, Q. Chen and G.-B. Li, Eur. J. Med. Chem., 2018, 155, 806–823.

40 J. Y. Hong, I. R. Price, J. J. Bai and H. Lin, ACS Chem. Biol., 2019, 14, 1802–1810.

41 S. Lain, J. J. Hollick, J. Campbell, O. D. Staples, M. Higgins, M. Aoubala, A. McCarthy, V. Appleyard, K. E. Murray, L. Baker, A. Thompson, J. Mathers, S. J. Holland, M. J. R. Stark, G. Pass, J. Woods, D. P. Lane and N. J. Westwood, Cancer Cell, 2008, 13, 454–463.

42 J. Morimoto, Y. Hayashi and H. Suga, Angew. Chem. Int. Ed., 2012, 51, 3423–3427.

43 T. Suzuki, M. N. A. Khan, H. Sawada, E. Imai, Y. Itoh, K. Yamatsuta, N. Tokuda, J. Takeuchi, T. Seko, H. Nakagawa and N. Miyata, J. Med. Chem., 2012, 55, 5760–5773.

44 S. Schuster, C. Roessler, M. Meleshin, P. Zimmermann, Z. Simic, C. Kambach, C. Schiene-Fischer, C. Steegborn, M. O. Hottiger and M. Schutkowski, Sci. Rep., 2016, 6, 22643.

45 Y. Huang, J. Liu, L. Yan and W. Zheng, Bioorg. Med. Chem. Lett., 2016, 26, 1612–1617.

46 S. Moniot, M. Forgione, A. Lucidi, G. S. Hailu, A. Nebbioso, V. Carafa, F. Baratta, L. Altucci, N. Giacché, D. Passeri, R. Pellicciari, A. Mai, C. Steegborn and D. Rotili, J. Med. Chem., 2017, 60, 2344–2360.

47 S. Sundriyal, S. Moniot, Z. Mahmud, S. Yao, P. Di Fruscia, C. R. Reynolds, D. T. Dexter, M. J. E. Sternberg, E. W.-F. Lam, C. Steegborn and M. J. Fuchter, J. Med. Chem., 2017, 60, 1928–1945.

48 P. Mellini, Y. Itoh, E. E. Elboray, H. Tsumoto, Y. Li, M. Suzuki, Y. Takahashi, T. Tojo, T. Kurohara, Y. Miyake, Y. Miura, Y. Kitao, M. Kotoku, T. Iida and T. Suzuki, J. Med. Chem., 2019, 62, 5844–5862.

49 T. Rumpf, S. Gerhardt, O. Einsle and M. Jung, Acta Cryst., 2015, F71, 1498–1510.

50 J. Jin, B. He, X. Zhang, H. Lin and Y. Wang, J. Am. Chem. Soc., 2016, 138, 12304–12307.

51 P. Knyphausen, S. De Boor, N. Kuhlmann, L. Scislowski, A. Extra, L. Baldus, M. Schacherl, U. Baumann, I. Neundorf and M. Lammers, J. Biol. Chem., 2016, 291, 14677–14694.

52 Y. Wang, Y. M. E. Fung, W. Zhang, B. He, M. W. H. Chung, J. Jin, J. Hu, H. Lin and Q. Hao, Cell Chem. Biol., 2017, 24, 339–345.

53 L. L. Yang, W. Xu, J. Yan, H. L. Su, C. Yuan, C. Li, X. Zhang, Z. J. Yu, Y. H. Yan, Y. Yu, Q. Chen, Z. Wang, L. Li, S. Qian and G. B. Li, Med. Chem. Comm., 2019, 10, 164–168.

54 M. S. Finnin, J. R. Donigian and N. P. Pavletich, Nat. Struct. Biol., 2001, 8, 621–625.

55 S. Moniot, M. Schutkowski and C. Steegborn, J. Struct. Biol., 2013, 182, 136–143.

56 J. L. Feldman, K. E. Dittenhafer-Reed, N. Kudo, J. N. Thelen, A. Ito, M. Yoshida and J. M. Denu, Biochemistry, 2015, 54, 3037–3050.

57 I. Galleano, J. Nielsen, A. S. Madsen and C. A. Olsen, Synlett, 2017, 28, 2169–2173.

58 N. Rajabi, M. Auth, K. R. Troelsen, M. Pannek, D. P. Bhatt, M. Fontenas, M. D. Hirschey, C. Steegborn, A. S. Madsen and C. A. Olsen, Angew. Chem. Int. Ed., 2017, 56, 14836–14841.

59 C. Moreno-Yruela, A. S. Madsen and C. A. Olsen, Protoc. Exch., , DOI:10.21203/RS.2.13042/V1.

60 B. Kitir, A. R. Maolanon, R. G. Ohm, A. R. Colaço, P. Fristrup, A. S. Madsen and C. A. Olsen, Biochemistry, 2017, 56, 5134–5146.

61 D. M. Molina, R. Jafari, M. Ignatushchenko, T. Seki, E. A. Larsson, C. Dan, L. Sreekumar, Y. Cao, P. Nordlund, D. Martinez Molina, R. Jafari, M. Ignatushchenko, T. Seki, E. A. Larsson, C. Dan, L. Sreekumar, Y. Cao and P. Nordlund, Science, 2013, 341, 84–87.

62 R. Jafari, H. Almqvist, H. Axelsson, M. Ignatushchenko, T. Lundbäck, P. Nordlund and D. M. Molina, Nat. Protoc., 2014, 9, 2100–2122.

63 W. Kabsch, Acta Cryst., 2010, D66, 125–132.

64 A. Vagin and A. Teplyakov, J. Appl. Crystallogr., 1997, 30, 1022–1025.

65 M. D. Winn, C. C. Ballard, K. D. Cowtan, E. J. Dodson, P. Emsley, P. R. Evans, R. M. Keegan, E. B. Krissinel, A. G. W. Leslie, A. McCoy, S. J. McNicholas, G. N. Murshudov, N. S. Pannu, E. A. Potterton, H. R. Powell, R. J. Read, A. Vagin and K. S. Wilson, Acta Cryst., 2011, D67, 235–242.

66 G. N. Murshudov, P. Skubák, A. A. Lebedev, N. S. Pannu, R. A. Steiner, R. A. Nicholls, M. D. Winn, F. Long and A. A. Vagin, Acta Cryst., 2011, D67, 355–367.

67 P. Emsley, B. Lohkamp, W. G. Scott and K. Cowtan, Acta Cryst., 2010, D66, 486–501.

68 F. Long, R. A. Nicholls, P. Emsley, S. Gražulis, A. Merkys, A. Vaitkus and G. N. Murshudov, Acta Cryst., 2017, D73, 112–122.

69 V. B. Chen, W. B. Arendall, J. J. Headd, D. A. Keedy, R. M. Immormino, G. J. Kapral, L. W. Murray, J. S. Richardson and D. C. Richardson, Acta Cryst., 2010, D66, 12–21.

70 B. C. Smith and J. M. Denu, Biochemistry, 2007, 46, 14478–14486.

71 B. C. Smith and J. M. Denu, J. Biol. Chem., 2007, 282, 37256–37265.

72 B. C. R. Dancy, S. A. Ming, R. Papazyan, C. A. Jelinek, A. Majumdar, Y. Sun, B. M. Dancy, W. J. Drury, R. J. Cotter, S. D. Taverna and P. A. Cole, J. Am. Chem. Soc., 2012, 134, 5138–5148.

73 B. M. Hirsch, Z. Du, X. Li, J. A. Sylvester, C. Wesdemiotis, Z. Wang and W. Zheng, Med. Chem. Comm., 2011, 2, 291–299.

74 H. Rink, Tetrahedron Lett., 1987, 28, 3787–3790.

75 A. S. Farooqi, J. Y. Hong, J. Cao, X. Lu, I. R. Price, Q. Zhao, T. Kosciuk, M. Yang, J. J. Bai and H. Lin, J. Med. Chem., 2019, 62, 4131–4141.

76 N. Rajabi, A. L. Nielsen and C. A. Olsen, ACS Med. Chem. Lett., 2020, 11, 1886–1892.

77 M. Kawaguchi, N. Ieda and H. Nakagawa, J. Med. Chem., 2019, 62, 5434–5452.

78 N. A. Spiegelman, I. R. Price, H. Jing, M. Wang, M. Yang, J. Cao, J. Y. Hong, X. Zhang, P. Aramsangtienchai, S. Sadhukhan and H. Lin, ChemMedChem, 2018, 13, 1890–1894.

79 Y. L. Chiang and H. Lin, Org. Biomol. Chem., 2016, 14, 2186–2190.

80 R. L. Stein, in Kinetics of Enzyme Action, 2011, pp. 287–302.

81 C. R.A., in Evaluation of Enzyme Inhibitors in Drug Discovery, 2013, pp. 203–244.

82 B. P. Hubbard, A. P. Gomes, H. Dai, J. Li, A. W. Case, T. Considine, T. V. Riera, J. E. Lee, S. Y. E, D. W. Lamming, B. L. Pentelute, E. R. Schuman, L. A. Stevens, A. J. Y. Ling, S. M. Armour, S. Michan, H. Zhao, Y. Jiang, S. M. Sweitzer, C. A. Blum, J. S. Disch, P. Y. Ng, K. T. Howitz, A. P. Rolo, Y. Hamuro, J. Moss, R. B. Perni, J. L. Ellis, G. P. Vlasuk and D. A. Sinclair, Science, 2013, 339, 1216–1219.

83 K. T. Tran, J. S. Pallesen, S. M. Solbak, D. Narayanan, A. Baig, J. Zang, A. Aguayo-Orozco, R. M. C. Carmona, A. D. Garcia and A. Bach, J. Med. Chem., 2019, 62, 8028–8052.

84 Y. Zhou, H. Zhang, B. He, J. Du, H. Lin, R. A. Cerione and Q. Hao, J. Biol. Chem., 2012, 287, 28307–28314.

85 R. Deziel, J. Rahil, A. Wahhab, M. Allan, N. Nguyen, Sirtuin inhibitors, Patent (WO2009026701A1).

86 W. Zheng, Y. Jiang. Selenourea warhead and building method thereof, Patent (CN106632595A).

87 H. Lin, R. Cerione, Methods for treatment of cancer by targeting SIRT5, Patent (US20190298747).

88 H. Suga, J. Morimoto, Peptide library production method, peptide library, and screening method, Patent (US10195578B2).

89 H. Lin, Thiourea compounds and their us e as inhibitors of Sirt2 or Sirt5, Patent (US10556878B2).

90 H. Dai, D. A. Sinclair, J. L. Ellis and C. Steegborn, Pharmacol Ther., 2018, 188, 140–154.

91 S. Li, B. Wu and W. Zheng, Bioorg. Med. Chem. Lett., 2019, 29, 461–465.

92 A. G. Dodge, J. E. Richman, G. Johnson and L. P. Wackett, Appl. Environ. Microbiol., 2006, 72, 7468–7476.

93 L. L. Poulsen, R. M. Hyslop and D. M. Ziegler, Arch. Biochem. Biophys., 1979, 198, 78–88.

94 E. Zaorska, T. Hutsch, M. Gawryś-Kopczyńska, R. Ostaszewski, M. Ufnal and D. Koszelewski, Bioorg. Chem., 2019, 88, 102941.

95 M. Yoshida, M. Kijima, M. Akita and T. Beppu, J. Biol. Chem., 1990, 265, 17174–17179.

96 R. H. Skoge and M. Ziegler, J. Cell Sci., 2016, 129, 2972–2982.

97 I. Hamaidi, L. Zhang, N. Kim, M. H. Wang, C. Iclozan, B. Fang, M. Liu, J. M. Koomen, A. E. Berglund, S. J. Yoder, J. Yao, R. W. Engelman, B. C. Creelan, J. R. Conejo-Garcia, S. J. Antonia, J. J. Mulé and S. Kim, Cell Metab., 2020, 32, 420–436.e12.

98 F. Hu, X. Sun, G. Li, Q. Wu, Y. Chen, X. Yang, X. Luo, J. Hu and G. Wang, Cell Death Dis., 2019, 10, 9.

99 Y. Li, M. Zhang, R. G. Dorfman, Y. Pan, D. Tang, L. Xu, Z. Zhao, Q. Zhou, L. Zhou, Y. Wang, Y. Yin, S. Shen, B. Kong, H. Friess, S. Zhao, L. Wang and X. Zou, Neoplasia, 2018, 20, 745–756.

100 J. Chen, A. W. H. Chan, K. F. To, W. Chen, Z. Zhang, J. Ren, C. Song, Y. S. Cheung, P. B. S. Lai, S. H. Cheng, M. H. L. Ng, A. Huang and B. C. B. Ko, Hepatology, 2013, 57, 2287–2298.

101 N. A. Spiegelman, X. Zhang, H. Jing, J. Cao, I. B. Kotliar, P. Aramsangtienchai, M. Wang, Z. Tong, K. M. Rosch and H. Lin, ACS Chem. Biol., 2019, 14, 2014–2023.

102 J. M. Tharp, J. T. Hampton, C. A. Reed, A. Ehnbom, P.-H. C. Chen, J. S. Morse, Y. Kurra, L. M. Pérez, S. Xu and W. R. Liu, Nat. Commun., 2020, 11, 1392.

103 R. A. Copeland, Nat. Rev. Drug Discov., 2016, 15, 87–95.

104 Y. Jiang, J. Liu, D. Chen, L. Yan and W. Zheng, Trends Pharmacol. Sci., 2017, 38, 459–472.

105 A. S. Madsen and C. A. Olsen, J. Med. Chem., 2012, 55, 5582–5590.

