## Supplemental Information for "Mechanism-based inhibitors of SIRT2: structure–activity relationship, X-ray structures, target engagement, regulation of α-tubulin acetylation and inhibition of breast cancer cell migration"

### Electronic supplementary information

#### Scheme S1. Additional inhibitors for the structure-activity relationship study

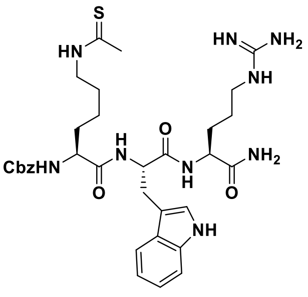

**S1**, 20% [1 μM]**

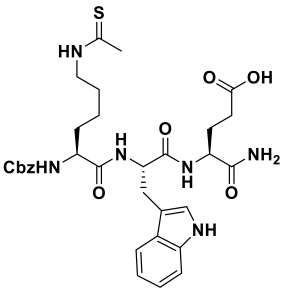

**S2**, 39% [10 μM]**

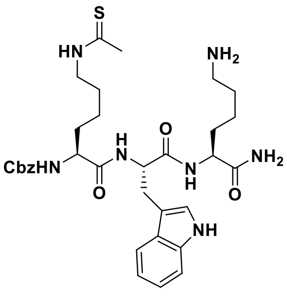

**S3,** 36% [1 μM]**

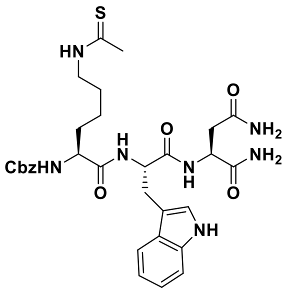

**S4,** 26% [1 μM]**

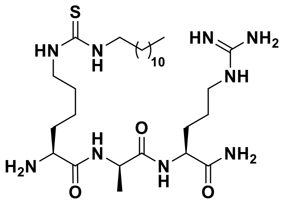

**S5**, 84% [1 μM]**
***37% [100 μM]***

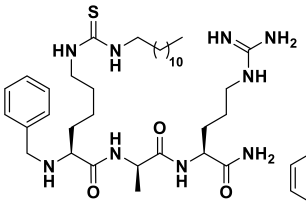

**S6**, 86% [1 μM]**
***31% [100 μM]***

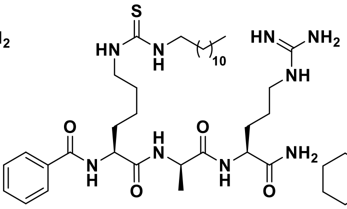

**S7,** 0.17 ± 0.01 μM**
***7.5 0.9 μM***

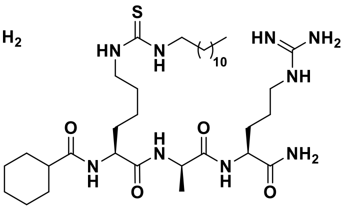

**S8,** 0.16 ± 0.01 μM**
***4.9 1.0 μM***

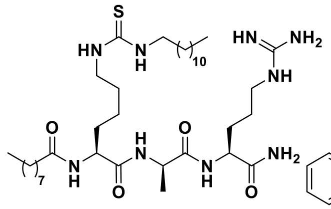

**S9**, 0.51 ± 0.03 μM**
***47% [100 μM]***

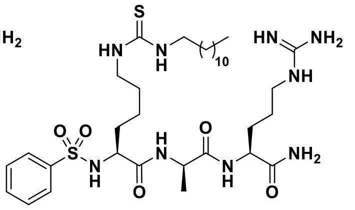

**S10**, 0.19 ± 0.02 μM**
***6.6 0.6 μM***

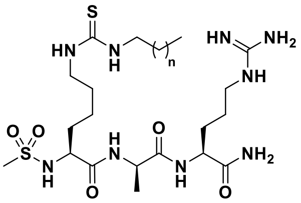

**S11** (n = 6), 0.40 ± 0.05 μM**
**S12** (n = 2), 13% [10 μM]**

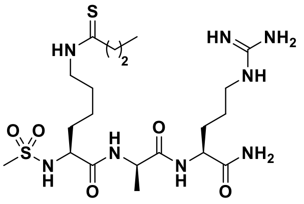

**S13**, 31% [10 μM]**

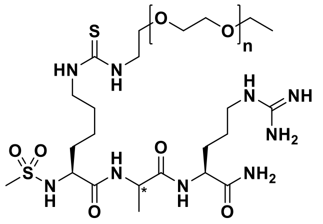

**S14** (**R*, n = 1), 16% [100 μM]**
**S15** (**S*, n = 2), 51 ± 12 μM**
**S16** (**R*, n = 2), 13% [100 μM]**

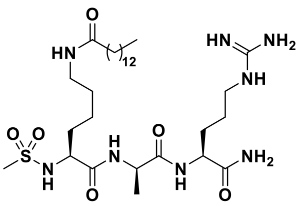

**S17**, 0.74 ± 0.07 μM**

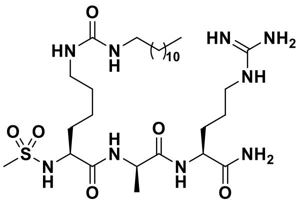

**S18**, 0.75 ± 0.05 μM**

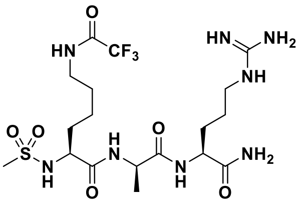

**S19**, No inhib. [100 μM]**

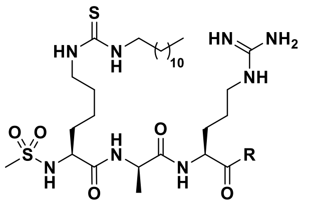

**S20** (R = OH), 0.11 ± 0.01 μM**
**S21** (R = $N(CH_2CH_2)_2O$), 0.09 ± 0.01 μM**
***13 2 μM***

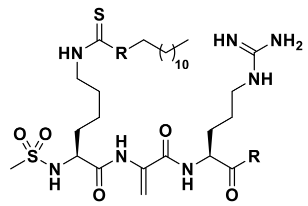

**S22** (R = $NH_2$), 0.03 ± 0.01 μM**
***0.18 0.03 μM***
**S23** (R = $(CH_2)$), 0.03 ± 0.01 μM**
***0.14 0.03 μM***

Potencies against recombinant SIRT2 (100 nM) are given as mean $IC_{50}$ values ± SD or %-inhibition against QPKKac** and ETDKmyr (***bold***) substrates tested at 50 μM.

#### Scheme S2. Synthesis of inhibitors in solution, part I

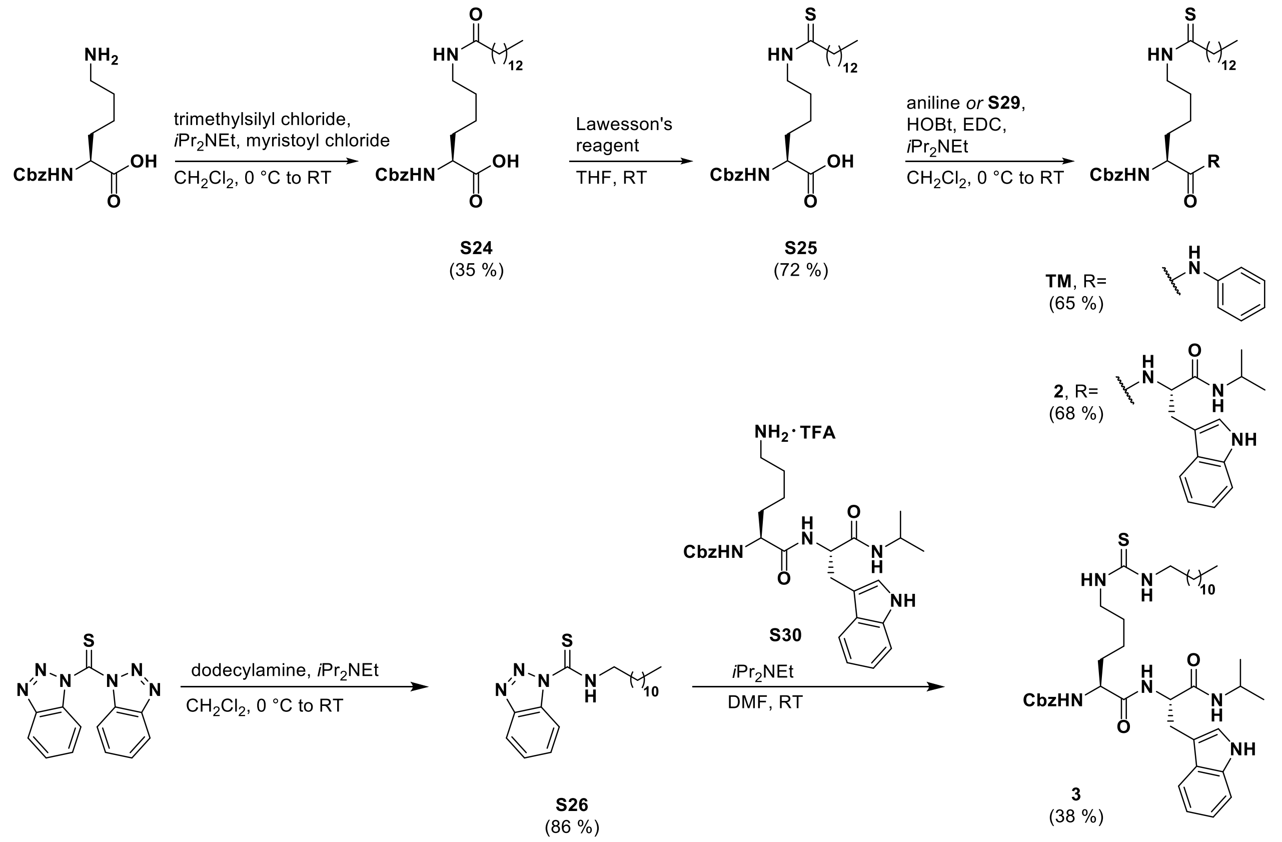

#### Scheme S3. Synthesis of inhibitors in solution, part II

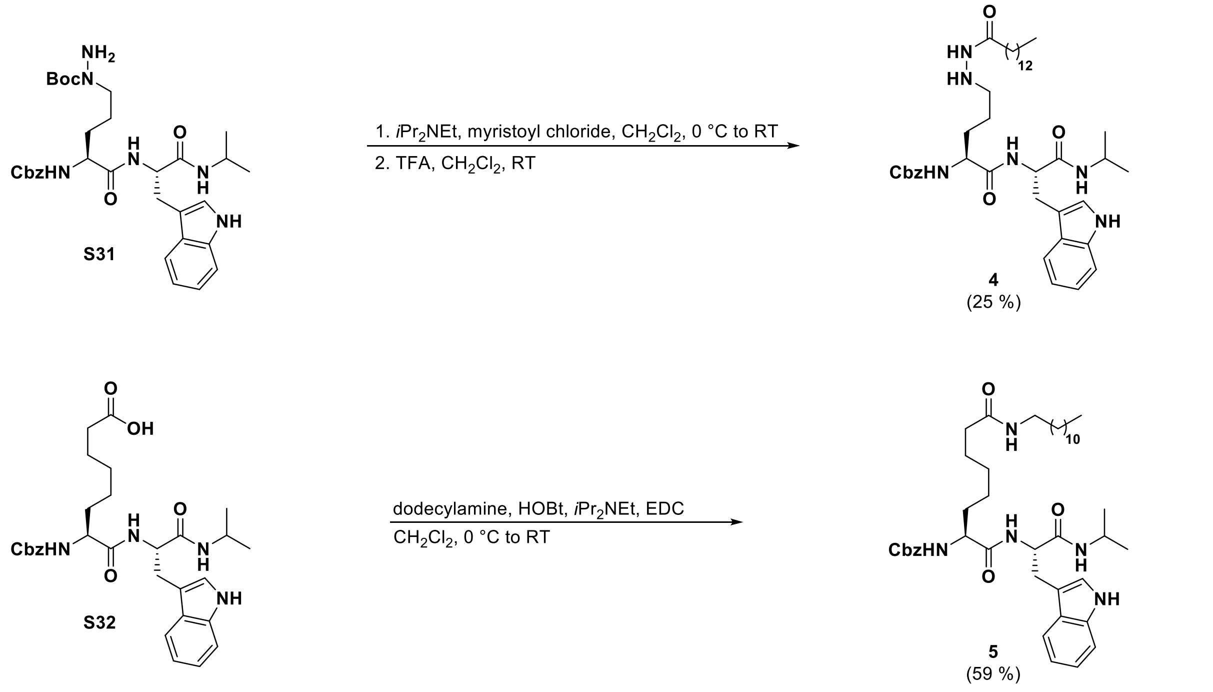

#### Scheme S4. Synthesis of inhibitors on solid-phase, part I

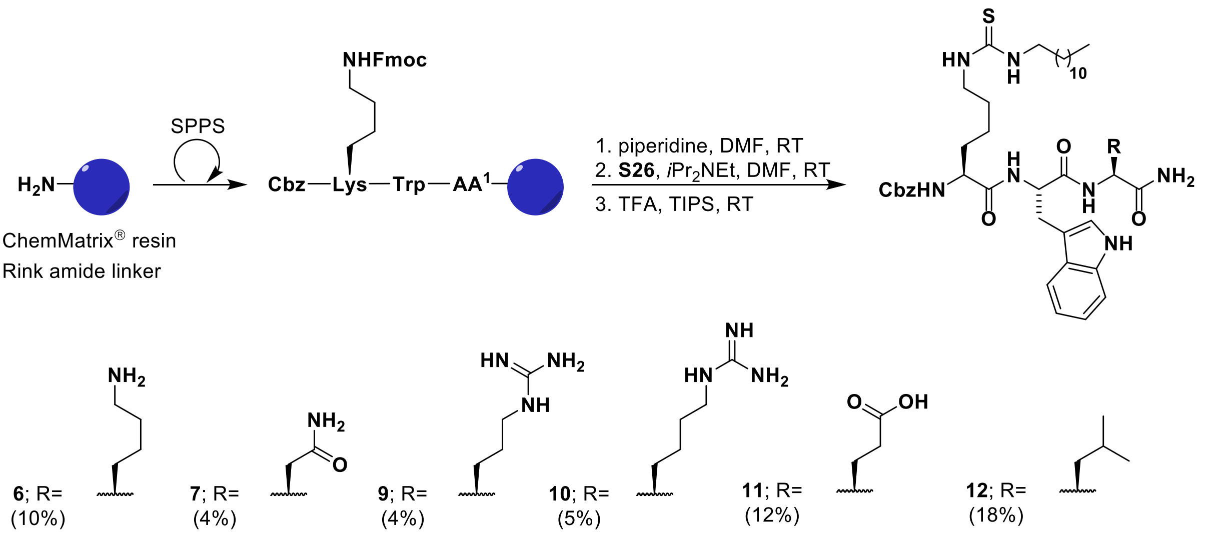

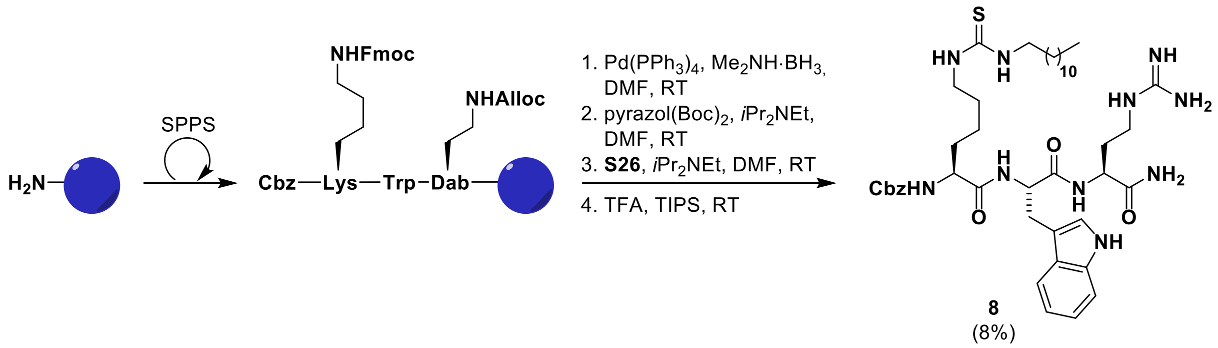

**Note:** Compound **8** was synthesized using an on-resin guanidinylation approach, as attempts loading Fmoc-norArg(Boc)$_2$-OH onto the resin gave rise to poor yields. We suggest this might be caused by an intramolecular cyclization upon activation of the amino acid *C*-terminus.

### Scheme S5. Synthesis of inhibitors on solid-phase, part II

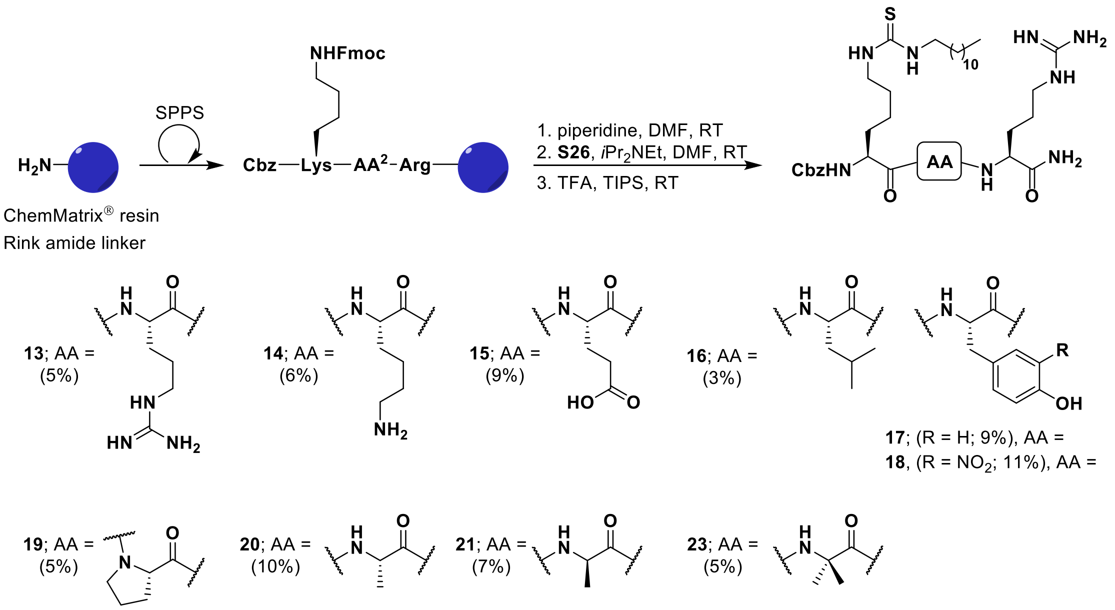

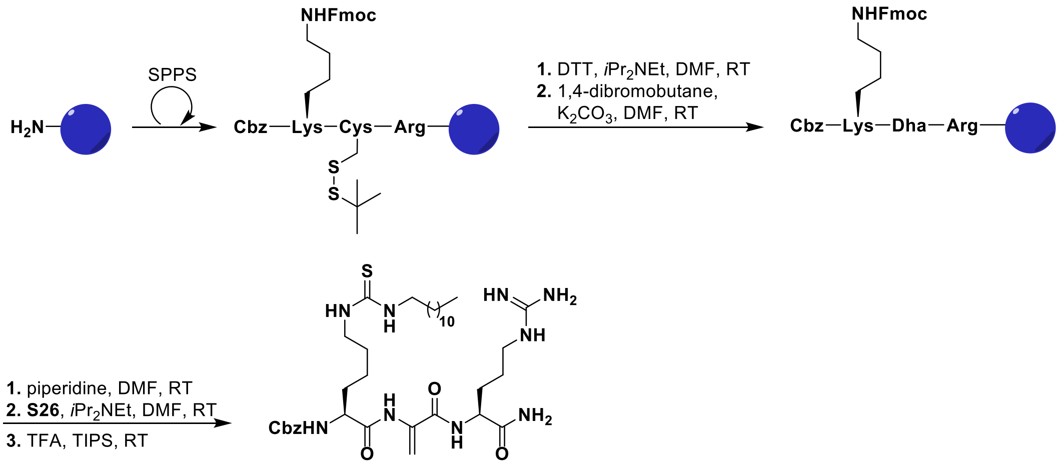

**22**
(5%)

### Scheme S6. Synthesis of inhibitors on solid-phase, part III

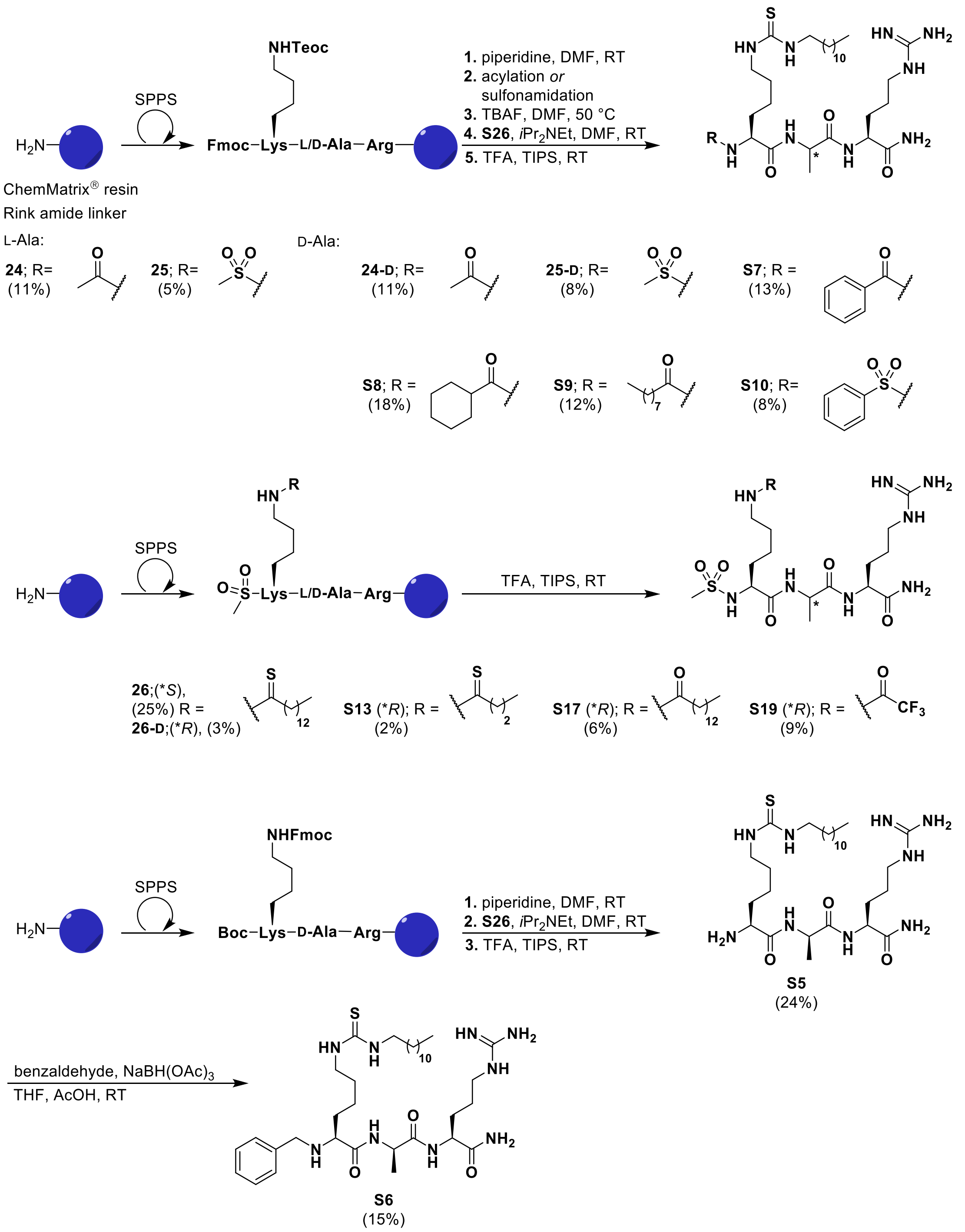

### Scheme S7. Synthesis of inhibitors on solid-phase, part IV

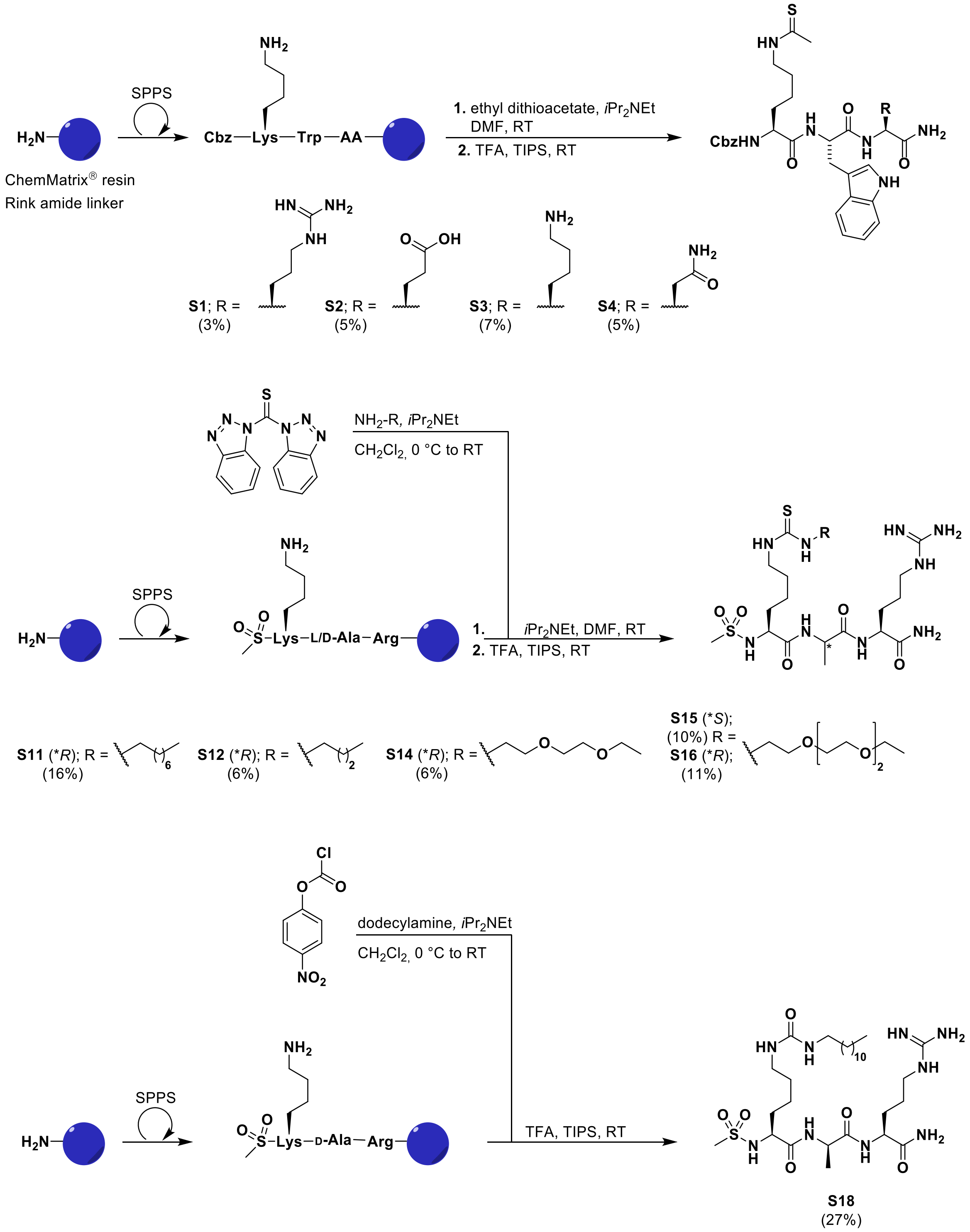

### Scheme S8. Synthesis of inhibitors on solid-phase, part V

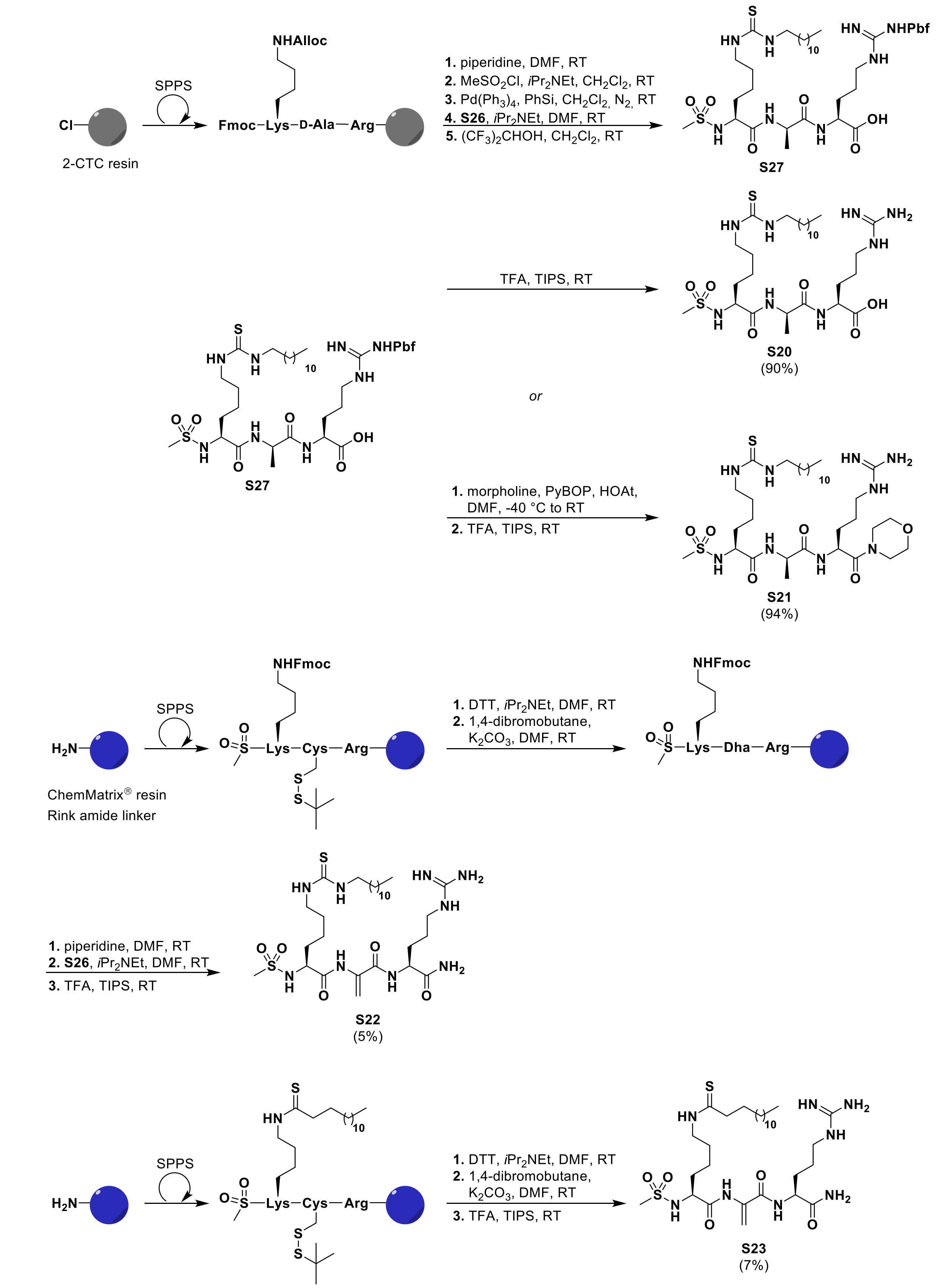

### Scheme S9. Synthesis of Fmoc-hArg(Boc)2-OH

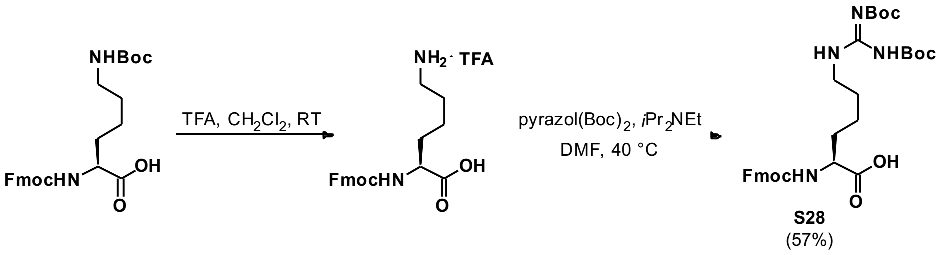

### Scheme S10. Synthesis of tenovin-6 and S2iL5

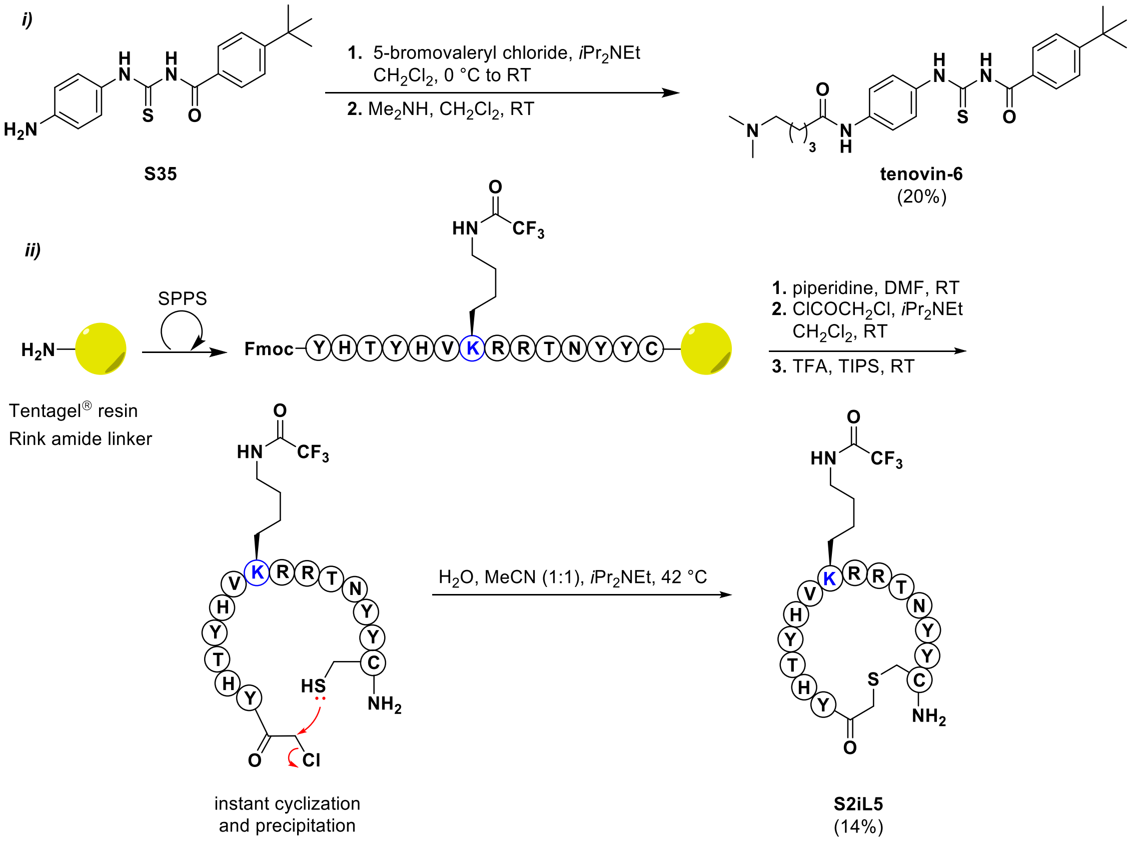

#### Scheme S11. Non-commercial building blocks used in study

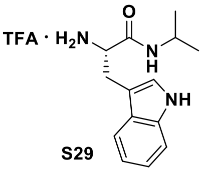

S29

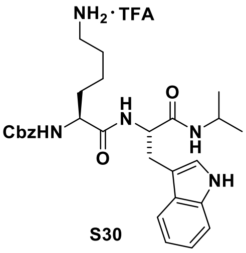

S30

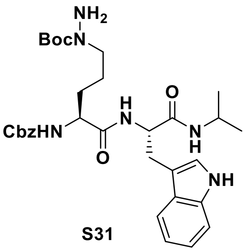

S31

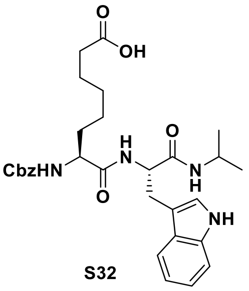

S32

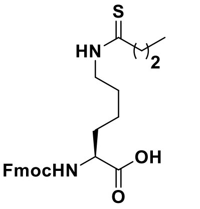

S33

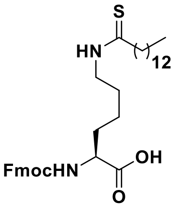

S34

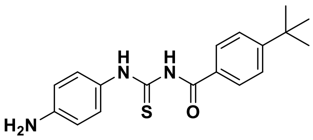

S35

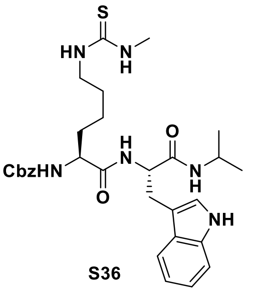

S36

Compounds **S29–S32**,[1] **S33**,[2] **S34**[3] and **S35**[4] were synthesized as previously reported.

### Figure S1. Dose-response curves – SIRT2 deacetylation

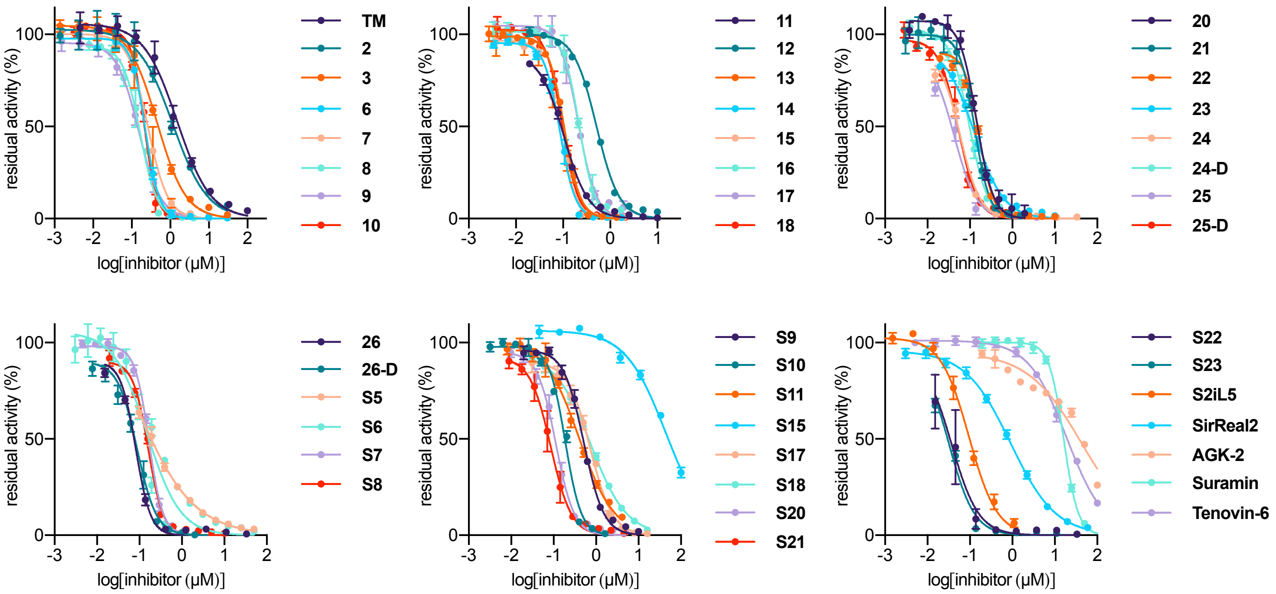

Concentration–response curves against inhibition of SIRT2 deacetylation for representative compounds using QPKKac as substrate. $IC_{50}$ values are reported in Table 1 and Table S1.

### Figure S2. Dose-response curves – SIRT2 demyristoylation

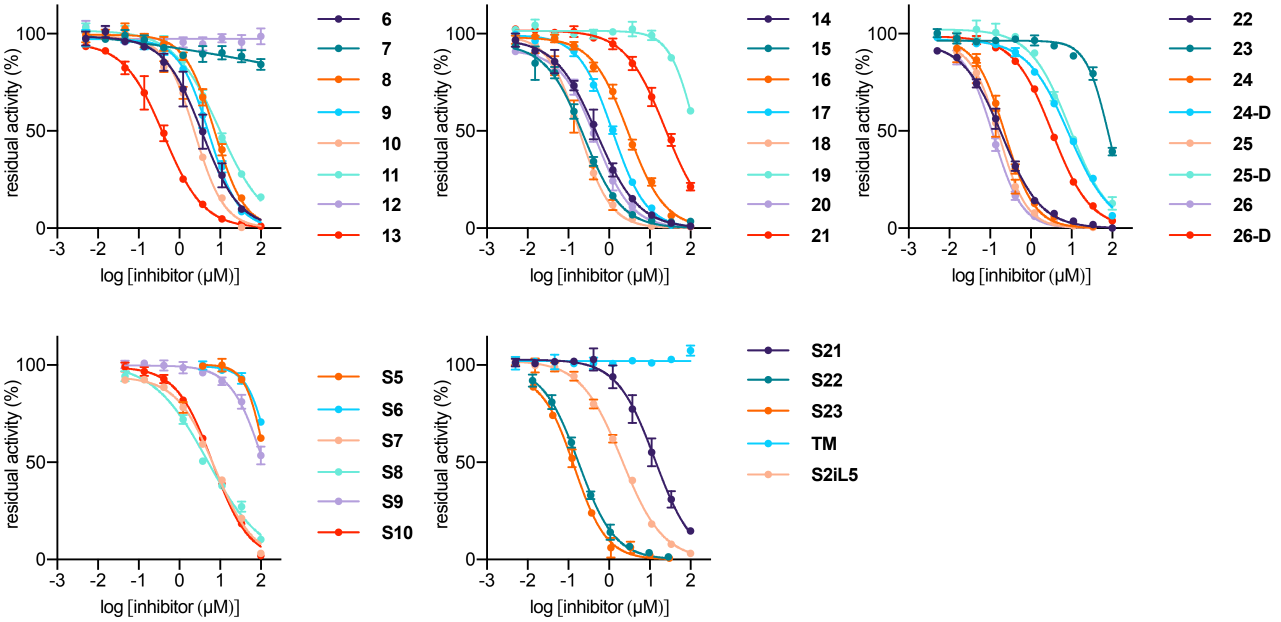

Concentration–response curves against inhibition of SIRT2 demyristoylation for representative compounds using ETDKmyr as substrate. $IC_{50}$ values are reported in Table 1 and Table S1.

### Figure S3. Dose-response curves – SIRT1 and SIRT3 deacetylation

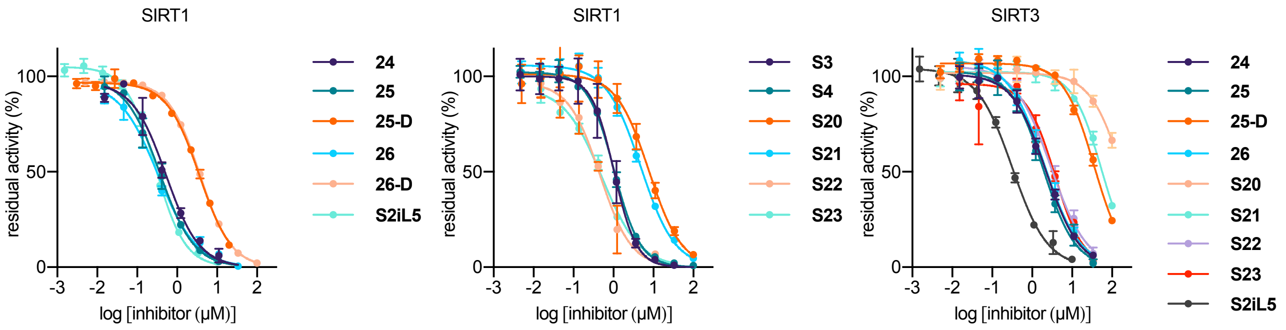

Concentration–response curves against inhibition of SIRT1 and SIRT3 deacetylation for representative compounds using QPKKac as substrate. $IC_{50}$ values are reported in Table S1.

### Figure S4. Dose-response curves from cell viability assays

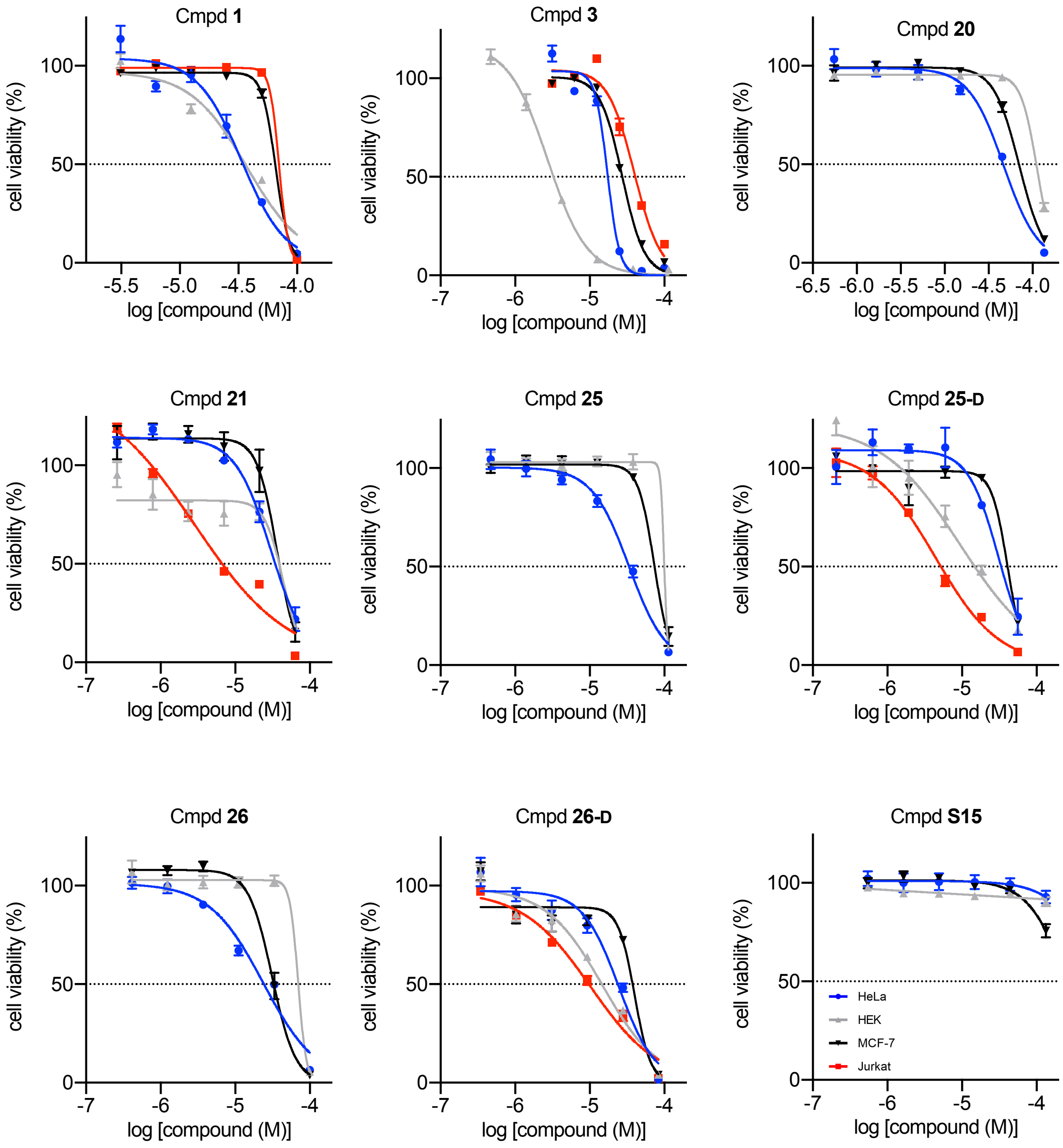

*(continued on next page)*

*(Figure S4, continued)*

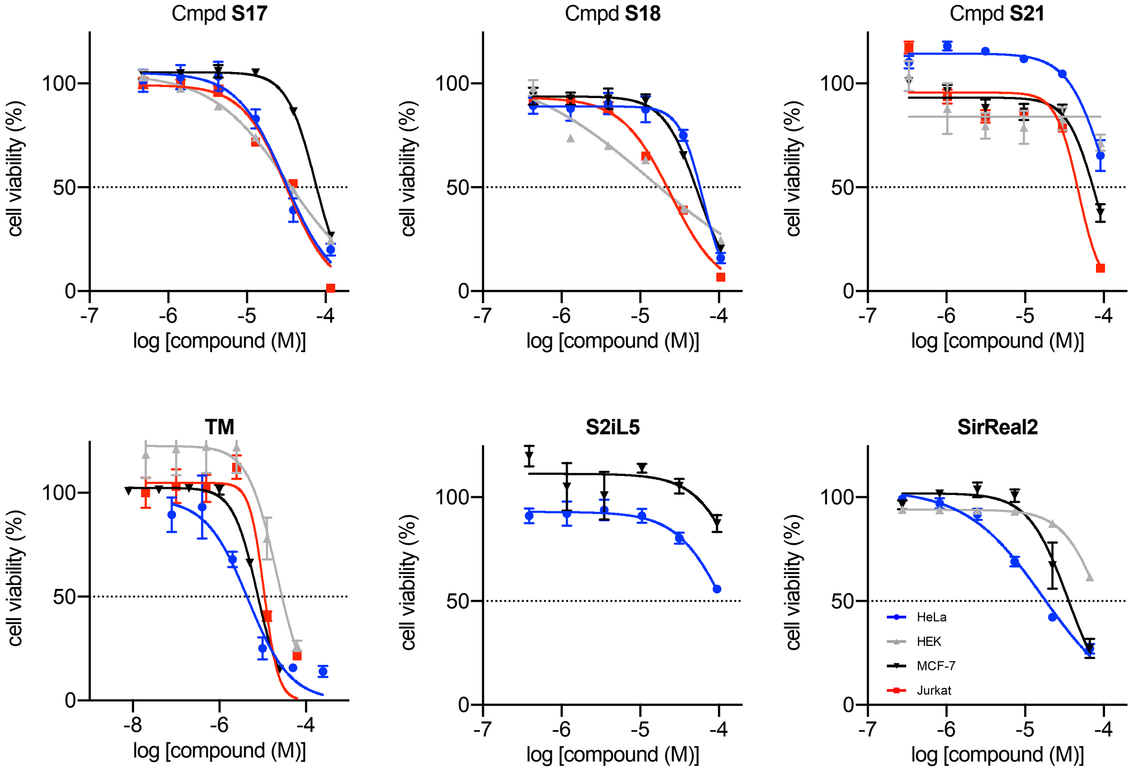

### Figure S5. Chemical stability of selected compounds in assay buffer

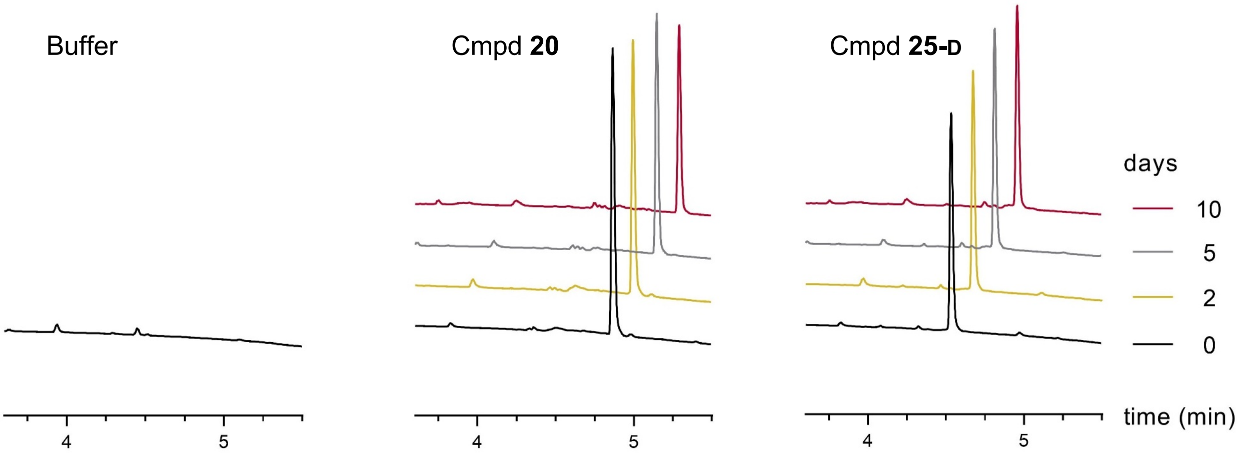

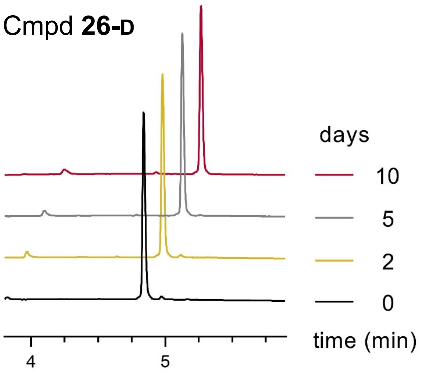

UPLC traces ($A_{214}$) of compounds **20**, **25-D** or **26-D** ($A_{280}$) in SIRT assay buffer at 37 °C, after incubation for up to 10 days showing no inhibitor degradation under assay conditions.

#### Figure S6. Stability of selected compounds in cell media and human serum

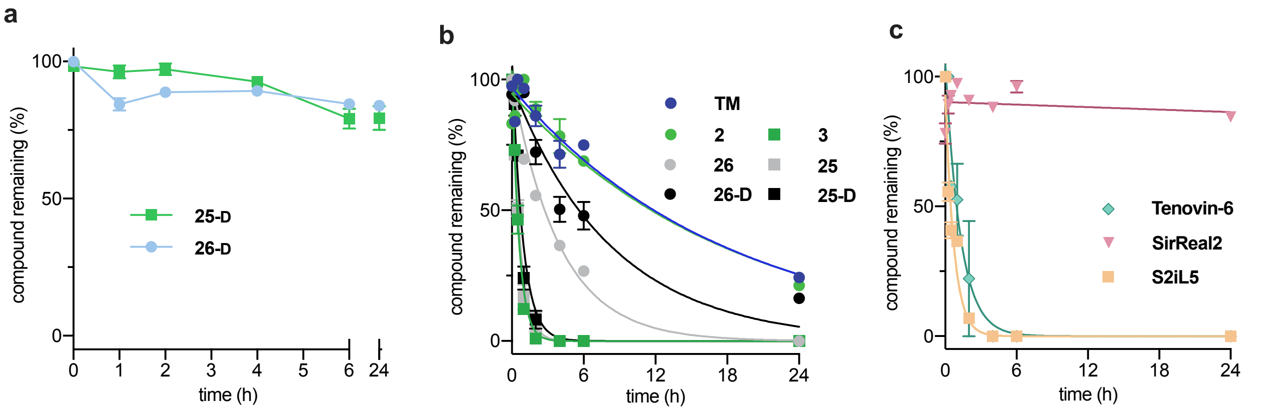

(**a**) Stability of compound **25-D** (thiourea) and **26-D** (thioamide) in cell media (MEM, 10% (v/v) FBS) at 37 °C over 24 h. (**b-c**) Serum stability assays for all tested compounds for up to 24 h. Data are shown as mean values relative to the peak intensity at $t$ = 0 h ± SEM (n = 2–3).

#### Figure S7. Degradation of compound 3 in human serum

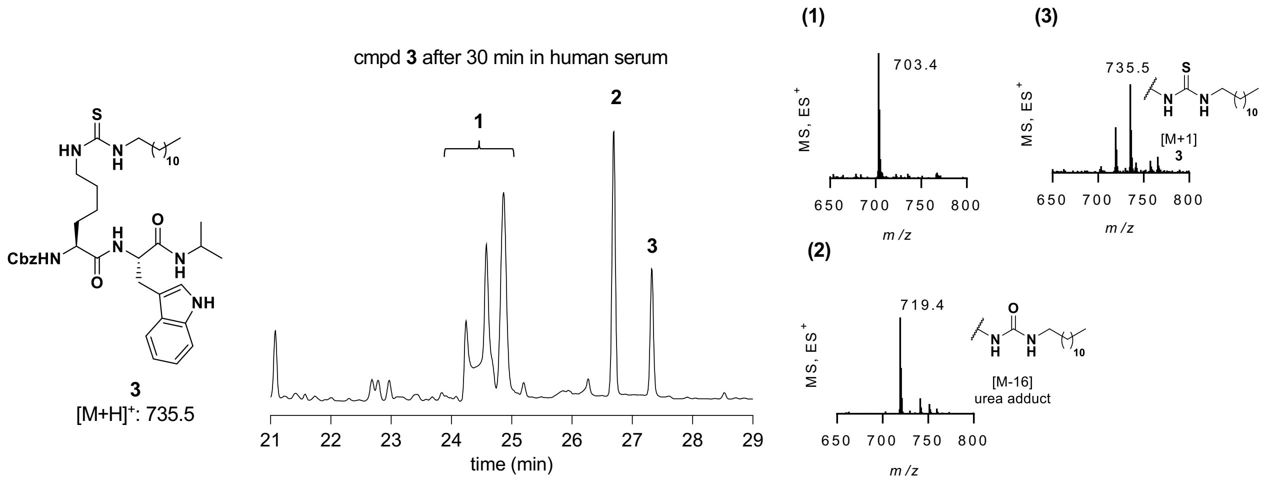

UV ($A_{254}$) and TIC ($ES^+$) chromatogram and mass spectra of compound **3** in human serum after 30 min. Rapid degradation was observed with the formation of several metabolites, in particular, an [M-16] mass adduct (**2**) corresponding to oxourea conversion could be identified.

### Figure S8. Cellular thermal shift assays against SIRT1–3

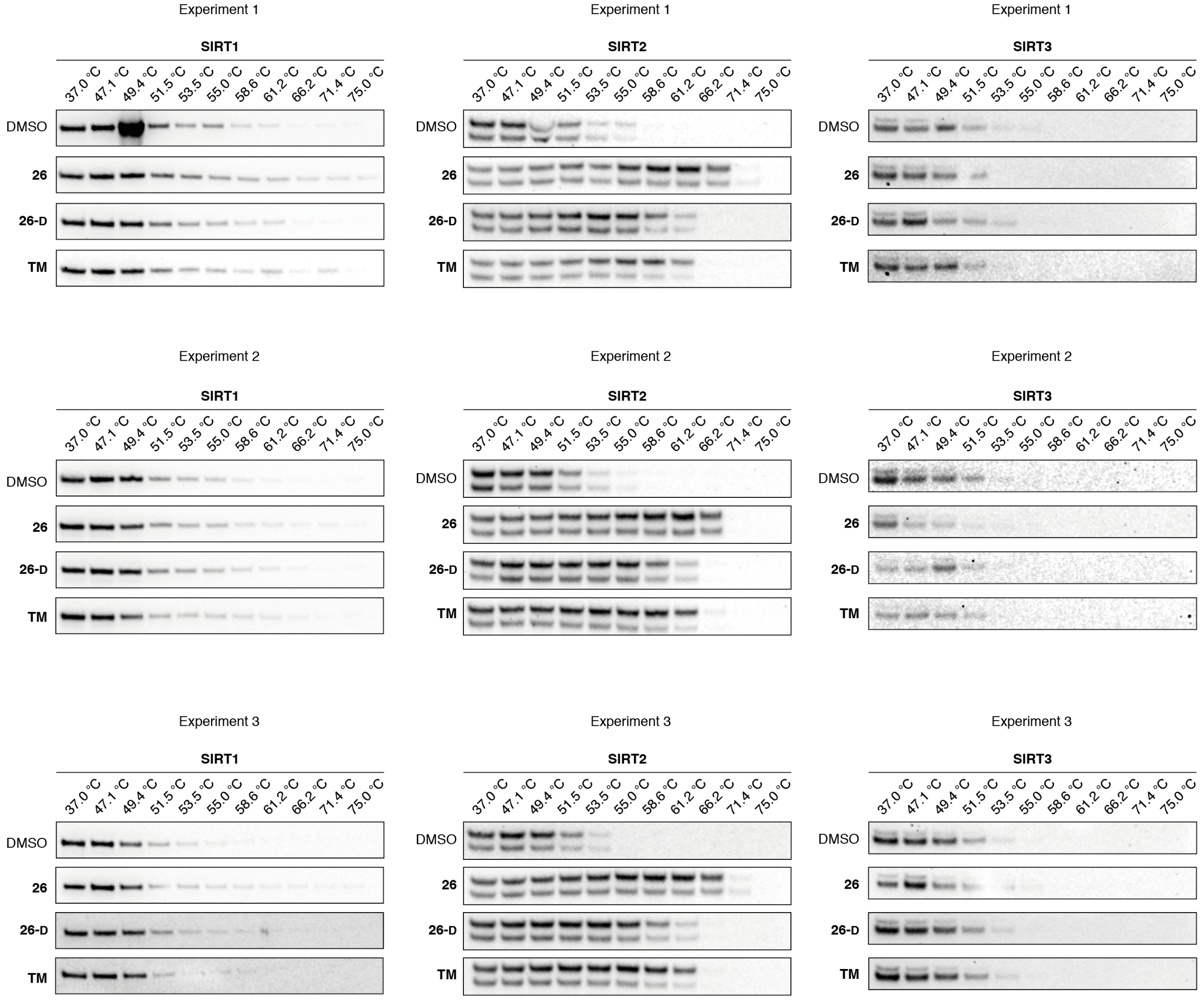

Western blot analysis of whole cell lysates from HEK293T cells after 1 h treatment with **26** (0.01 µM), **26-D** (0.10 µM), **TM** (10 µM) or respective volumes of DMSO followed by heat treatment. For full blots and protein marker see the full Western blot section (n = 3).

#### Figure S9. Supplementary immunofluorescent images

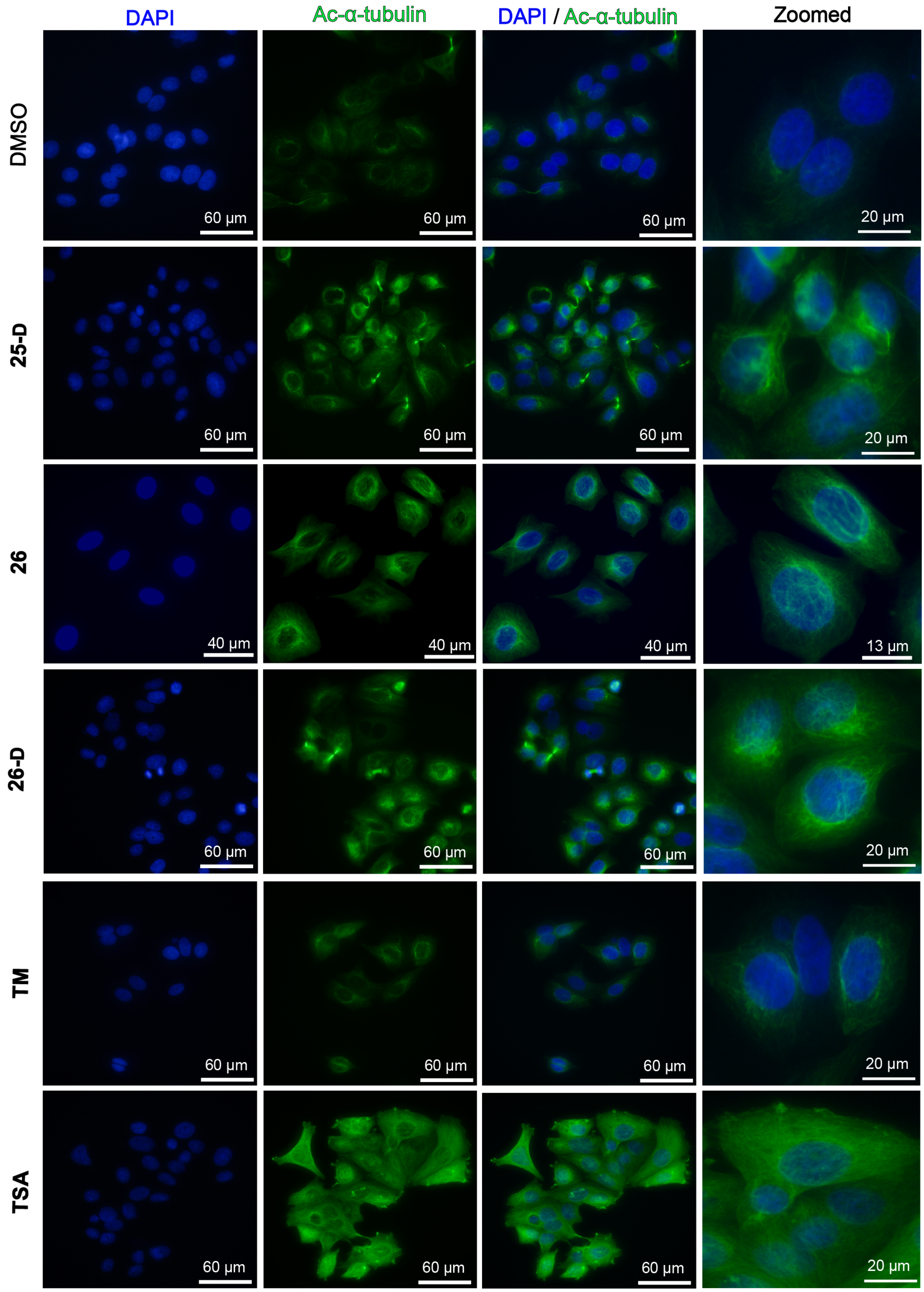

Immunofluorescent images (40×) of MCF-7 cells treated with inhibitor (25 µM, except for **26** and **TSA**: 5 µM) or DMSO (vehicle) for 6 h. The data are representative images from two individual experiments performed in duplicate. DAPI (blue, nuclear counterstain) and Ac-α-tubulin (green).

#### Table S1. Data collection and refinement statistics for X-ray co-crystal structures

| Data collection | | SIRT2:**13** | SIRT2:**23** |
|---|---|---|---|
| X-ray source | | SLS X06DA | SLS X06DA |
| Space group | | $P2_12_12_1$ | $P2_12_12_1$ |
| Unit cell | *a* (Å) | 35.9 | 35.8 |
| | *b* (Å) | 65.3 | 65.5 |
| | *c* (Å) | 114.0 | 114.9 |
| Wavelength (Å) | | 1.000 | 1.000 |
| Resolution (Å) | | 37.9–1.61 (1.64–1.61) | 43.2–1.54 (1.59–1.54) |
| Unique reflections | | 36305 | 40773 |
| Completeness (%) [a] | | 100 (100) | 99.8 (99.1) |
| $R_{merge}$ (%) [a] | | 4.7 (84.8) | 5.1 (95.3) |
| $I/\sigma(I)$[a] | | 9.3 (0.9) | 19.8 (2.2) |
| Wilson *B* (Å$^2$) | | 20.8 | 37.6 |

[a] Values in parentheses are for the highest resolution shell

| Refinement statistics | SIRT2:**13** | SIRT2:**23** |
|---|---|---|
| Resolution range (Å) | 34.2–1.70 | 20.0–1.70 |
| No. of reflections | 28752 | 29045 |
| No. of non-hydrogen atoms | 2428 | 2365 |
| $R_{work}$ (%) | 21.6 | 27.2 |
| $R_{free}$ (%) | 25.6 | 31.7 |
| R.m.s. deviations | | |
| Bond length (Å) | 0.005 | 0.003 |
| Bond angle (degree) | 1.20 | 1.14 |
| *B*-factors (Å$^2$) | 31.1 | 43.9 |
| Protein | 33.3 | 43.9 |
| Inhibitor | 48.4 | 66.1 |
| Waters | 29.9 | 40.1 |
| Ramachandran plot (%) | | |
| Favored region | 98.2 | 97.8 |
| Allowed region | 1.8 | 2.2 |
| Outlier region | 0.0 | 0.0 |

#### Table S2. Inhibitory potencies against SIRT1–3 deacylation

| Compound | SIRT1 | SIRT2 | SIRT3 |
|---|---|---|---|
| **1** | 1.4 ± 0.4 µM | 14% [1 µM] | 15% [10 µM] |
| **2** | NI [10 µM] | 1.2 ± 0.2 µM | NI [10 µM] |
| **3** | 47% [10 µM] | 0.45 ± 0.04 µM (22% [100 µM]) | NI [10 µM] |
| **4** | 10% [100 µM] | 19% [100 µM] | ND |
| **5** | NI [100 µM] | 29% [100 µM] | ND |
| **6** | 71% [10 µM] | 0.22 ± 0.02 µM (3.6 ± 0.6 µM) | 21% [10 µM] |
| **7** | 29% [10 µM] | 0.26 ± 0.03 µM (16% [100 µM]) | NI [10 µM] |
| **8** | 30% [1 µM] | 0.16 ± 0.02 µM (7.3 ± 0.8 µM) | 48% [10 µM] |
| **9** | 52% [1 µM] | 0.15 ± 0.01 µM (5.3 ± 0.5 µM) | 62% [10 µM] |
| **10** | 76% [1 µM] | 0.21 ± 0.03 µM (2.4 ± 0.6 µM) | 20% [10 µM] |
| **11** | 28% [1 µM] | 0.10 ± 0.01 µM (9.6 ± 1.1 µM) | 21% [10 µM] |
| **12** | NI [10 µM] | 0.51 ± 0.03 µM (NI [100 µM]) | 18% [10 µM] |
| **13** | 81% [1 µM] | 0.11 ± 0.01 µM (0.42 ± 0.06 µM) | 48% [1 µM] |
| **14** | 25% [1 µM] | 0.09 ± 0.01 µM (0.50 ± 0.08 µM) | 54% [1 µM] |
| **15** | 56% [1 µM] | 0.08 ± 0.01 µM (0.24 ± 0.04 µM) | 62% [10 µM] |
| **16** | 57% [1 µM] | 0.22 ± 0.02 µM (3.9 ± 0.3 µM) | 63% [10 µM] |
| **17** | 53% [1 µM] | 0.21 ± 0.02 µM (1.2 ± 0.1 µM) | 18% [1 µM] |
| **18** | 56% [1 µM] | 0.09 ± 0.01 µM (0.19 ± 0.05 µM) | 28% [1 µM] |
| **19** | 17% [10 µM] | 53% [1 µM] (40% [100 µM]) | NI [10 µM] |
| **20** | 61% [1 µM] | 0.14 ± 0.02 µM (0.47 ± 0.05 µM) | 20% [1 µM] |
| **21** | 78% [10 µM] | 0.13 ± 0.02 µM (24 ± 2 µM) | 11% [10 µM] |
| **22** | 77% [1 µM] | 0.17 ± 0.01 µM (0.20 ± 0.02 µM) | 56% [1 µM] |

| Compound | SIRT1 | SIRT2 | SIRT3 |
|---|---|---|---|
| **23** | 55% [10 μM] | 0.13 ± 0.02 μM (95 ± 3 μM) | NI [10 μM] |
| **24** | 0.53 ± 0.11 μM | 0.05 ± 0.01 μM (0.23 ± 0.05 μM) | 2.2 ± 0.4 μM |
| **24-D** | 34% [1 μM] | 0.10 ± 0.01 μM (8.6 ± 0.6 μM) | 21% [10 μM] |
| **25** | 0.37 ± 0.07 μM | 0.04 ± 0.01 μM (0.17 ± 0.03 μM) | 1.9 ± 0.3 μM |
| **25-D** | 3.6 ± 0.7 μM | 0.05 ± 0.01 μM (9.1 ± 0.8 μM) | 35 ± 3 μM |
| **26** | 0.33 ± 0.05 μM | 0.07 ± 0.01 μM (0.12 ± 0.02 μM) | 2.0 ± 0.4 μM |
| **26-D** | 3.7 ± 0.2 μM | 0.09 ± 0.01 μM (3.7 ± 0.2 μM) | 31% [10 μM] |
| **S1** | 64% [1 μM] | 20% [1 μM] | 43% [10 μM] |
| **S2** | 83% [1 μM] | 39% [10 μM] | 9% [10 μM] |
| **S3** | 1.0 ± 0.2 μM | 36% [1 μM] | 59% [10 μM] |
| **S4** | 1.0 ± 0.1 μM | 26% [1 μM] | 14% [10 μM] |
| **S5** | 15% [10 μM] | 84% [1 μM] (37% [100 μM]) | NI [10 μM] |
| **S6** | 9% [10 μM] | 86% [1 μM] (31% [100 μM]) | NI [10 μM] |
| **S7** | 49% [1 μM] | 0.17 ± 0.01 μM (7.5 ± 0.9 μM) | 21% [10 μM] |
| **S8** | 44% [1 μM] | 0.16 ± 0.01 μM (4.9 ± 1.0 μM) | 41% [10 μM] |
| **S9** | 55% [10 μM] | 0.51 ± 0.03 μM (47% [100 μM]) | NI [10 μM] |
| **S10** | 21% [1 μM] | 0.19 ± 0.02 μM (6.6 ± 0.6 μM) | 17% [10 μM] |
| **S11** | 71% [10 μM] | 0.40 ± 0.05 μM | NI [10 μM] |
| **S12** | NI [10 μM] | 13% [10 μM] | NI [10 μM] |
| **S13** | 14% [10 μM] | 31% [10 μM] | NI [10 μM] |
| **S14** | 12% [100 μM] | 16% [100 μM] | NI [10 μM] |
| **S15** | ND | 51 ± 12 μM | 16% [100 μM] |
| **S16** | 16% [100 μM] | 13% [100 μM] | NI [10 μM] |
| **S17** | 23% [10 μM] | 0.74± 0.07 μM | 10% [10 μM] |

| Compound | SIRT1 | SIRT2 | SIRT3 |
|---|---|---|---|
| **S18** | 38% [10 µM] | 0.75 ± 0.05 µM | NI [10 µM] |
| **S19** | 14% [100 µM] | NI [100 µM] | 10% [100 µM] |
| **S20** | 7.7 ± 2.9 µM | 0.11 ± 0.01 µM | 34% [100 µM] |
| **S21** | 4.8 ± 0.8 µM | 0.09 ± 0.01 µM<br>(13 ± 2 µM) | 55 ± 5 µM |
| **S22** | 0.46 ± 0.12 µM | 0.03 ± 0.01 µM<br>(0.18 ± 0.03) µM | 2.9 ± 0.5 µM |
| **S23** | 0.51 ± 0.11 µM | 0.03 ± 0.01 µM<br>(0.14 ± 0.03) µM | 3.7 ± 0.9 µM |
| **TM** | NI [100 µM] | 1.5 ± 0.3 µM<br>(NI [100 µM]) | NI [100 µM] |
| **S2iL5** | 0.24 ± 0.03 µM | 0.09 ± 0.01 µM<br>(2.1 ± 0.1 µM) | 0.30 ± 0.03 µM |
| **SirReal2** | 37% [10 µM] | 0.91 ± 0.08 µM<br>(NI [100 µM]) | NI [10 µM] |
| **AGK-2** | 62% [100 µM] | 50 ± 10 µM<br>(NI [10 µM]) | 69% [100 µM] |
| **Suramin** | 74% [10 µM] | 17 ± 1 µM<br>(NI [10 µM]) | 6% [100 µM] |
| **tenovin-6** | 36% [100 µM] | 19 ± 1 µM<br>(NI [10 µM]) | 38% [100 µM] |

Potencies against recombinant SIRT2 (100 nM) are given as mean $IC_{50}$ values ± SD or %-inhibition against QPKKac and ETDKmyr (parenthesis) substrates (50 µM). NI = no inhibition; ND = not determined.
The $K_M$ of QPKKac has been determined by to be >333 µM for SIRT1–3 by Lin and co-workers.[5]

### Table S3. Continous assay kinetic data

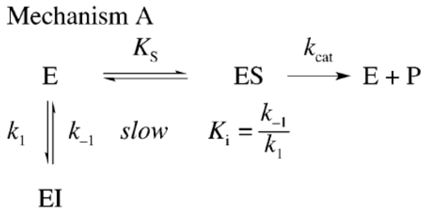

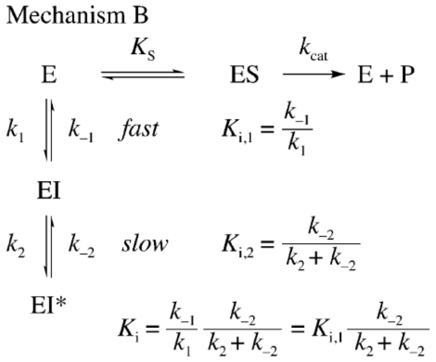

| Mechanism A | 26-D |
|---|---|
| **SIRT2** | |
| $k_1$ (nM·min$^{-1}$) | ~6 × 10$^{-7}$ |
| $k_{-1}$ (min$^{-1}$) | ~0 |
| $K_i$ (nM) | –$^{\dagger}$ |
| Dis. $t_{½}$ (min) | > 10$^5$ |

| Mechanism B | 25 | 26 |
|---|---|---|
| **SIRT2** | | |
| $k_2$ (min$^{-1}$) | 0.10 ± 0.01 | 0.07 ± 0.01 |
| $k_{-2}$ (min$^{-1}$) | ~2 × 10$^{-13}$ | ~6 × 10$^{-9}$ |
| $K_{i,1}$ (nM) | 240 ± 180 | 130 ± 80 |
| $K_i$ (nM) | < 10$^{-3}$ | < 10$^{-3}$ |
| Dis. $t_{½}$ (min) | > 10$^5$ | > 10$^5$ |

$^{\dagger}K_i$ values depend on $k_{-2}$ values and, since $k_{-2}$ values approached 0, $K_i$ could not be determined.

### Table S4. Inhibitory potencies for selected compounds against different sirtuin subtypes

| Compound | SIRT5* | SIRT6$^{\dagger}$ |
|---|---|---|
| **3** | NI [100 μM] | NI [10 μM] |
| **25** | NI [100 μM] | 30% [10 μM] |
| **25-D** | NI [100 μM] | 14% [10 μM] |
| **26** | NI [100 μM] | 41% [10 μM] |
| **26-D** | NI [100 μM] | 29% [10 μM] |
| **TM** | NI [100 μM] | NI [10 μM] |
| **S2iL5** | 25% [100 μM] | NI [10 μM] |
| **SirReal2** | NI [100 μM] | NI [10 μM] |
| **AGK-2** | 15% [100 μM] | NI [10 μM] |
| **Suramin** | 85% [100 μM] | NI [10 μM] |

Potencies against recombinant SIRT5 (100 nM) or SIRT6 (600 nM) given as %-inhibition using *LGKglu as substrate or †ETDKmyr as substrates (50 µM).

#### Table S5. Inhibitory potencies against SIRT1/2 deacetylation of H3K9ac.

| Compound | **26** | **26-D** | **TM** | **S2iL5** |
|---|---|---|---|---|
| SIRT1 | 2.3 ± 0.5 µM | 7.7 ± 1.3 µM | NI [100 µM] | 2.0 ± 0.6 µM |
| SIRT2 | 16 ± 4 nM* | 22 ± 11 nM* | 40 ± 9 µM | 0.57 ± 0.09 |

Potencies against recombinant SIRT1 and SIRT2 (20 nM) using H3K9ac (50 µM) as substrate.
*stoichiometric inhibition.

#### Table S6. cLogP and PSA values.

| **Compound** | **cLogP** | **PSA (Å$^2$)** |
|---|---|---|
| **SirReal2** | 5.14 | 72.0 |
| **TM** | 8.62 | 96.4 |
| **Tenovin-6** | 5.39 | 98.5 |
| **AGK-2** | 5.08 | 77.5 |
| **S2iL5** | – | – |
| **25** | 1.88 | 263 |
| **25-D** | 1.90 | 264 |
| **26** | 2.03 | 252 |
| **26-D** | 2.52 | 253 |

Calculated LogP (cLogP) and polar surface area (PSA) values were calculated using QikProp.[6]

The cLogP values were approximately 3-fold lower for compounds **25**, **25-D**, **26**, and **26-D** than **SirReal2**, **tenovin-6**, and **AGK-2**, while that of **TM** was substantially higher at 8.6. For the PSA, our compounds were generally higher than all control compounds. On the other hand, the higher PSA values for our compounds compared to the small molecule control compounds also gives rise to better aqueous solubility enabling biological studies without the need for drug formulation.

### Table S7. Compound half-lives in human male serum

| Compound | Serum half-life ($t_{½}$) |
|---|---|
| **TM** | 12 ± 3 h |
| **S2iL5** | 0.5 ± 0.1 h |
| **SirReal2** | >>24 h |
| **tenovin-6** | 0.8 ± 0.5 h |
| **2** | 13 ± 2 h |
| **3** | 0.4 ± 0.1 h |
| **25** | 0.5 ± 0.1 h |
| **25-D** | 0.6 ± 0.1 h |
| **26** | 2.8 ± 0.3 h |
| **26-D** | 5.8 ± 1.2 h |

#### Table S8. $EC_{50}$ values of SIRT2 inhibitors in cell viability assays

| Compound | HeLa | MCF-7 | HEK293T | Jurkat |
|---|---|---|---|---|
| **1** | 27.8–40.7 | 62.8–69.0 | 30.8–45.6 | 65.0–76.0 |
| **3** | 15.6–19.2 | 25.6–28.9 | 2.36–3.17 | 34.3–45.4 |
| **20** | 41.4–53.1 | 64.7–76.6 | 102–121 | ND |
| **21** | 25.8–34.9 | 28.7–45.6 | 32.3–60.4 | 1.17–6.89 |
| **25** | 28.9–39.1 | 63.7–84.3 | ~ 100 (wide) | ND |
| **25-D** | 23.6–40.8 | 31.9–53.3 | 4.88–18.3 | 3.24–6.12 |
| **26** | 19.4–29.3 | 28.4–34.3 | 23.4–100+ | ND |
| **26-D** | 18.8–32.1 | 29.0–55.0 | 10.3–21.3 | 7.20–14.7 |
| **S15** | > 100 | > 100 | > 100 | ND |
| **S18** | 23.1–41.4 | 69.2–78.9 | 28.6–38.0 | 25.4–40.8 |
| **S19** | 52.1–75.8 | 47.6–62.8 | 3.68–46.1 | 21.2–28.1 |
| **S21** | 89.8–126 | 64.7–93.7 | >100 | 35.5–64.4 |
| **TM** | 1.87–11.5 | 6.88–8.41 | 12.4–35.9 | 7.27–16.0 |
| **S2iL5** | 90.8–173 | 32.0–162 | ND | ND |
| **SirReal2** | 13.8–20.8 | 27.6–44.9 | 84.5–109 | ND |

$EC_{50}$ values reported from 95% confidence intervals (μM) from cell viability (MTT) assays. Data are based on three individual experiments performed in duplicate. ND = not determined.

#### General experimental methods

All reagents and solvents were of analytical grade and used without further purification as obtained from commercial suppliers. Anhydrous solvents were obtained from a PureSolv-system. Reactions were conducted under an atmosphere of nitrogen whenever anhydrous solvents were used. Reactions were monitored by thin-layer chromatography (TLC) using silica gel coated plates (analytical $SiO_2$-60, F-254) or by LC-MS. TLC plates were visualized under UV light and by dipping in either (a) a solution of potassium permanganate (10 g/L), potassium carbonate (67 g/L) and sodium hydroxide (0.83 g/L) in water, (b) a solution of ninhydrin (3 g/L) in 3% acetic acid in water (v/v), or (c) a solution of molybdate-phosphoric acid (12.5 g/L) and cerium(IV)sulfate (5 g/L) in 3% conc. sulfuric acid in water (v/v) followed by heating with a heat gun. Vacuum liquid chromatography (VLC) was performed with silica gel 60 (particle size 15−40 µm). After column chromatography, appropriate fractions were pooled and dried at high vacuum (<2 mbar) for at least 12 h to give obtained products in high purity (>95%) unless stated otherwise. Evaporation of solvents was carried out under reduced pressure at a temperature below 40 °C. LC−MS analyses were performed on a Phenomenex Kinetex column (1.7 µm, 50×2.10 mm) using a Waters Acquity ultra high-performance liquid chromatography (UPLC) system. Gradient A with eluent I (0.1% HCOOH in $H_2O$) and eluent II (0.1% HCOOH in MeCN) rising linearly from 0% to 95% of II during $t$ = 0.00−5.20 min was applied at a flow rate of 0.6 mL/min. Preparative reversed-phase HPLC purification was performed on a C18 Phenomenex Luna column (5 µm, 100 Å, 250×20 mm) or a C8 Phenomenex Luna column (5 µm, 100 Å, 250×21.2 mm) using an Agilent 1260 LC system equipped with a diode array UV detector and an evaporative light scattering detector (ELSD). Gradient B with eluent III ($H_2O$/MeCN/TFA, 95:5:0.1, v:v) and eluent IV (0.1% TFA in MeCN) rising linearly from 0−30% to 95% of IV during $t$ = 5−45 min at a flow rate of 20 mL/min was applied. Analytical HPLC was performed on a C18 phenomenex Luna column (3 µm, 100 Å, 150×4.60 mm) or a C8 phenomenex Luna column (5 µM, 100 Å, 250×4.60 mm) using an Agilent 1100 series system equipped with a diode array UV detector. Gradient C using eluent III and eluent IV, rising linearly from 0% to 95% of IV during $t$ = 5−20 or $t$ = 5−35 min was applied at a flow rate of 1.2 mL/min. High-resolution mass spectrometry (HRMS) measurements were recorded either on a maXis G3 quadrupole time-of-flight (TOF) mass spectrometer (Bruker Daltonics, Bremen, Germany) equipped with an electrospray ionization (ESI) source or on an Agilent 1290 UHPLC equipped with a diode array detector and coupled to Agilent 6550 QTOF mass spectrometer operated in positive electrospray or on a Bruker Solarix WR by either matrix assisted laser desorption/ionization, or ESI. Nuclear magnetic resonance (NMR) spectra were recorded either on a Bruker Avance III HD equipped with a cryogenically cooled probe ($^{1}$H NMR and $^{13}$C NMR recorded at 600 and 151 MHz, respectively) or a Bruker Avance III ($^{1}$H NMR, $^{13}$C NMR and $^{19}$F NMR recorded at 400, 101, and 377 MHz, respectively). All spectra were recorded at 298 K. Chemical shifts are reported in ppm relative to deuterated solvent as internal standard ($\delta_H$ DMSO-$d_6$

2.50 ppm; $\delta_C$ DMSO-$d_6$ 39.52 ppm; $\delta_H$ $CDCl_3$ 7.26 ppm; $\delta_C$ $CDCl_3$ 77.16 ppm; $\delta_H$ MeOD-$d_4$ 3.31 ppm; $\delta_C$ MeOD 49.0 ppm). Assignments of NMR spectra are based on 2D correlation spectroscopy (COSY, HSQC, TOCSY and HMBC spectra). Compound stock concentrations were determined by quantitative NMR (qNMR) using maleic acid as internal standard.

**General solid phase peptide synthesis procedure (SPPS)**

Peptides were synthesized on a ChemMatrix® or TentaGel®-resin using a Rink amide (RAM) linker by standard solid-phase peptide synthesis. Resin loading was determined spectrophotometrically, quantifying the amount of released fluorene upon cleavage of the Fmoc group from a small sample.[7] Each elongation step was performed by applying the relevant amino acid (3 equiv.), HATU (2.9 equiv.) and *i*$Pr_2NEt$ (6 equiv.) *or* the relevant amino acid (1.5 equiv.), PyOxim (1.5 equiv.) and *i*$Pr_2NEt$ (6 equiv.) in DMF. Fmoc-deprotection was performed by treatment with DMF/piperidine (4:1, v/v, 4 mL; 2 min then 15 min), followed by washing with DMF (3×4 mL). The reaction progress was monitored by Kaiser's tests[8] or by test cleaveage and subsequent UPLC-MS analysis.

**General procedure for global deprotection and cleavage from the resin**

Peptides were cleaved from the resin by TFA/$H_2O$/TIPS (95:2.5:2.5 (v/v), 4 mL; 1 h) and solvent removed under a stream of nitrogen. The resulting crude was triturated with ice-cold diethyl ether and purified by preparative reversed-phase HPLC. Yields were determined based on resin loading.

**General procedure for on-resin Teoc deprotection**

A solution of TBAF trihydrate (10 equiv.) in DMF (2.0 mL/0.1 mmol resin) was added to the fritted syringe containing the resin bound peptide and the reaction mixture was agitated for 1 h at 50 ˚C. The procedure was repeated and the resin was then washed with DMF (3×4 mL) and $CH_2Cl_2$ (3×4 mL).

**General on-resin thioacetylation procedure**

Ethyl dithioacetate (2 equiv.) was dissolved in DMF (2.0 mL/0.1 mmol resin) and *i*$Pr_2NEt$ (2 equiv.) and added to a fritted syringe containing the resin bound peptide and the reaction mixture was agitated for 3-4 h. After washing with DMF (3×4 mL) and $CH_2Cl_2$ (2×4 mL) the reaction progress was evaluated with a Kaiser's test. If the Kaiser's test indicated presence of unreacted amine, the procedure was repeated.

**General on-resin thiourea formation procedure**

Compound **S26** (2 equiv.) and *i*$Pr_2NEt$ (2 equiv.) were dissolved in DMF (2.0 mL/0.1 mmol resin) and added to the fritted syringe containing the resin-bound peptide and the reaction mixture was agitated for 3-4 h. After washing with DMF (3×4 mL) and $CH_2Cl_2$ (3×4 mL) the reaction progress was evaluated with a Kaiser's test. If the Kaiser's test indicated presence of unreacted amine, the procedure was repeated.

For compounds containing other thiourea functionalities (compounds **S11–S12, S14–S16**), a solution of the desired amine (2 equiv.) and *i*$Pr_2NEt$ (3 equiv.) in $CH_2Cl_2$ (3.0 mL) was added dropwise over 5 min to a solution of bis(1-benzotriazolyl)methanethione (2 equiv.) in $CH_2Cl_2$ (1.5 mL) at 0 °C. The reaction mixture was concentrated under reduced pressure and the resulting crude residue and *i*$Pr_2NEt$ (2 equiv.) were dissolved in DMF (2.0 mL/0.1 mmol resin) and then added to the fritted syringe containing the resin bound peptide.

#### Compound syntheses

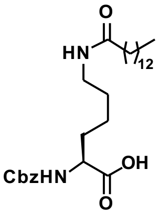

**$N^2$-((benzyloxy)carbonyl)-$N^6$-tetradecanoyl-L-lysine (S24).** Trimethylsilyl chloride (0.27 mL, 2.14 mmol) was added to a solution of Cbz-Lys-OH (300 mg, 1.07 mmol) and *i*$Pr_2NEt$ (0.75 mL, 4.28 mmol) in anh. $CH_2Cl_2$ (20 mL). The reaction mixture was stirred at ambient temperature for 30 min and myristoyl chloride (0.35 mL, 1.28 mmol) was added. After 90 min, the reaction mixture was poured into aq. citric acid (25% (w/w), 50 mL) and extracted with $CH_2Cl_2$ (4×25 mL). The combined organic layers were dried over $Na_2SO_4$, filtered and concentrated under reduced pressure. The crude residue was purified by column chromatography (0→4.5% MeOH and 0.5% AcOH in $CH_2Cl_2$) affording the desired amide **S24** (181 mg, 35%) as a colorless solid. TLC (3% MeOH and 0.5% AcOH in $CH_2Cl_2$): $R_f$ = 0.3. $^1$H NMR (600 MHz, MeOD-$d_4$) δ 7.48–7.18 (m, 5H, $H_{Ar,Cbz}$), 5.09 (s, 2H, $CH_{2,Cbz}$), 4.14 (dd, $J$ = 9.2, 4.7 Hz, 1H, $H_{\alpha,Lys}$), 3.16 (t, $J$ = 6.9 Hz, 2H, $H_{\varepsilon,Lys}$), 2.15 (t, $J$ = 7.5 Hz, 2H, (C=O)$CH_2$), 1.90–1.79 (m, 1H, $H_{\beta,Lys,A}$), 1.75–1.64 (m, 1H, $H_{\beta,Lys,B}$), 1.64–1.36 (m, 6H, $H_{\delta,Lys}$, $H_{\gamma,Lys}$, (C=O)$CH_2C\underline{H}_2$), 1.36–1.22 (m, 20H, $(C\underline{H}_2)_{10}CH_3$), 0.90 (t, $J$ = 7.0 Hz, 3H, $CH_3$). $^{13}$C NMR (151 MHz, MeOD-$d_4$) δ 176.3 ($NH_{\varepsilon,Lys}$CO), 175.9 (COOH), 158.7 ($CO_{Cbz}$), 138.2 ($C1_{Ar,Cbz}$), 129.4 ($C3_{Ar,Cbz}$, $C5_{Ar,Cbz}$), 129.0 ($C4_{Ar,Cbz}$), 128.7 ($C2_{Ar,Cbz}$, $C6_{Ar,Cbz}$), 67.6 ($CH_{2,Cbz}$), 55.3 ($C_{\alpha,Lys}$), 40.0 ($C_{\varepsilon,Lys}$), 37.2 ((C=O)$\underline{C}H_2$), 33.1 ($\underline{C}H_2CH_2CH_3$), 32.4 ($C_{\beta,Lys}$), 30.8–30.3 (8C, $(\underline{C}H_2)_{10}CH_3$), 29.9 ($C_{\delta,Lys}$), 27.1 ((C=O)$CH_2\underline{C}H_2$), 24.3 ($C_{\gamma,Lys}$), 23.7 ($\underline{C}H_2CH_3$), 14.5 ($CH_3$). ESI-MS *m/z* calcd for $C_{28}H_{47}N_2O_5^+$ $[M+H]^+$, 491.3; found 491.4. CAS RN: 213017-44-8.

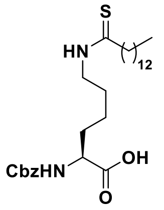

**$N^2$-((benzyloxy)carbonyl)-$N^6$-tetradecanethioyl-L-lysine (S25).** Lawesson's reagent (136 mg, 0.33 mmol) was added to a solution of amide **S24** (160 mg, 0.33 mmol) in anh. THF (10 mL). The reaction mixture was stirred at ambient temperature for 2 h and then poured into aq. HCl (2 M, 50 mL) and extracted with $CH_2Cl_2$ (3×40 mL). The combined organic layers were dried over $Na_2SO_4$, filtered and concentrated under reduced pressure. The crude residue was purified by column chromatography (0→1.5% MeOH and 0.5% AcOH in $CH_2Cl_2$) affording the desired thioamide **S25** (119 mg, 72%) as a colorless solid. TLC (1.5% MeOH and 0.5% AcOH in $CH_2Cl_2$): $R_f$ = 0.4. $^1$H NMR (600 MHz, MeOD-$d_4$) δ 7.43–7.20 (m, 5H, $H_{Ar,Cbz}$), 5.14–5.06 (m, 2H, $CH_{2,Cbz}$), 4.16 (dd, $J$ = 9.2, 4.7 Hz, 1H, $H_{\alpha,Lys}$), 3.58 (t, $J$ = 7.1 Hz, 2H, $H_{\varepsilon,Lys}$), 2.62–2.54 (m, 2H, (C=S)$CH_2$), 1.93–1.81 (m, 1H, $H_{\beta,Lys,A}$), 1.76–1.60 (m, 5H, $H_{\beta,Lys,B}$, $H_{\delta,Lys}$, (C=S)$CH_2C\underline{H}_2$), 1.52–1.38 (m, 2H, $H_{\gamma,Lys}$), 1.30 (d, $J$ = 13.9 Hz, 20H, $(C\underline{H}_2)_{10}CH_3$), 0.90 (t, $J$ = 7.0 Hz, 3H, $CH_3$). $^{13}$C NMR (151 MHz, MeOD-$d_4$) δ 206.5 (C=S), 175.9 (COOH), 158.7 ($CO_{Cbz}$), 138.2 ($C1_{Ar,Cbz}$), 129.5 ($C3_{Ar,Cbz}$, $C5_{Ar,Cbz}$), 129.0 ($C4_{Ar,Cbz}$), 128.7 ($C2_{Ar,Cbz}$, $C6_{Ar,Cbz}$), 67.6 ($CH_{2,Cbz}$), 55.2 ($C_{\alpha,Lys}$), 47.1 ((C=S)$\underline{C}H_2$), 46.6 ($C_{\varepsilon,Lys}$), 33.1 ($\underline{C}H_2CH_2CH_3$), 32.5 ($C_{\beta,Lys}$), 30.8–29.9 (9C, $(\underline{C}H_2)_{11}CH_3$), 28.2 ($C_{\delta,Lys}$), 24.4 ($C_{\gamma,Lys}$), 23.7 ($\underline{C}H_2CH_3$), 14.5 ($CH_3$). ESI-MS *m/z* calcd for $C_{28}H_{45}N_2O_5^+$ $[M+H]^+$, 505.3; found 505.3. CAS RN: 1429749-38-1.

**Benzyl (*S*)-(1-oxo-1-(phenylamino)-6-tetradecanethioamidohexan-2-yl)carbamate (TM).**

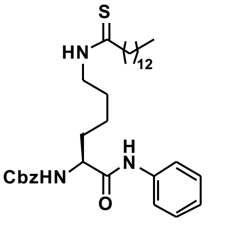

Carboxylic acid **S25** (50 mg, 0.10 mmol), aniline (13 µL, 0.15 mmol), HOBt (20 mg, 0.15 mmol), and *i*$Pr_2NEt$ (34 µL, 0.20 mmol) were dissolved in anh. $CH_2Cl_2$ (2.0 mL) and cooled to 0 °C. EDC (28 mg, 0.15 mmol) was added and the reaction mixture was stirred at 0 °C for 5 min and then overnight at ambient temperature. The reaction mixture was cooled to 0 °C and additional EDC (28 mg, 0.15 mmol) was added and the reaction mixture was stirred at 0 °C for 5 min and then for another 72 h at ambient temperature. The reaction mixture was concentrated under reduced pressure and resuspended in EtOAc (65 mL) and washed with aq. $KHSO_4$ (5%, 3×50 mL), saturated aq. $NaHCO_3$ (3×50 mL) and brine (2×50 mL). The organic phase was dried over $Na_2SO_4$, filtered and concentrated under reduced pressure. The crude residue was purified by column chromatography (0→1.25% MeOH in $CH_2Cl_2$) affording the desired thioamide **TM** (37 mg, 65%) as a colorless solid. $^1$H NMR (600 MHz, DMSO-*d*$_6$) δ 9.99 (s, 1H, $CO_{Lys}NH$), 9.86 (t, *J* = 5.5 Hz, 1H, $NH_{\varepsilon,Lys}$), 7.60 (d, *J* = 8.0 Hz, 2H, $H2_{Ph}$, $H6_{Ph}$), 7.53 (d, *J* = 7.9 Hz, 1H, $NH_{\alpha,Lys}$), 7.42–7.10 (m, 7H, $H_{Ar,Cbz}$, $H3_{Ph}$, $H5_{Ph}$), 7.04 (t, *J* = 7.4 Hz, 1H, $H4_{Ph}$), 5.03 (s, 2H, $CH_{2,Cbz}$), 4.14 (td, *J* = 8.6, 5.2 Hz, 1H, $H_{\alpha,Lys}$), 3.54–3.39 (m, 2H, $H_{\varepsilon,Lys}$), 2.50–2.46 (m, 2H, (C=S)$CH_2$), 1.76–1.48 (m, 6H, $H_{\beta,Lys}$, $H_{\delta,Lys}$, (C=S)$CH_2$<u>$CH_2$</u>), 1.48–1.29 (m, 2H, $H_{\gamma,Lys}$), 1.22 (d, *J* = 2.7 Hz, 20H, (C<u>H</u>$_2$)$_{10}$$CH_3$), 0.85 (t, *J* = 7.0 Hz, 3H, $CH_3$). $^{13}$C NMR (151 MHz, DMSO-*d*$_6$) δ 203.6 (C=S), 171.0 ($CO_{Lys}$), 156.1 ($CO_{Cbz}$), 138.9 ($C1_{Ph}$), 137.0 ($C1_{Ar,Cbz}$), 128.6 ($C3_{Ph}$, $C5_{Ph}$), 128.3 ($C3_{Ar,Cbz}$, $C5_{Ar,Cbz}$), 127.74 ($C4_{Ar,Cbz}$), 127.65 ($C2_{Ar,Cbz}$, $C6_{Ar,Cbz}$), 123.2 ($C4_{Ph}$), 119.2 ($C2_{Ph}$, $C6_{Ph}$), 65.4 ($CH_{2,Cbz}$), 55.3 ($C_{\alpha,Lys}$), 45.0 ($C_{\varepsilon,Lys}$), 44.9 ((C=S)<u>C</u>$H_2$), 31.6 ($C_{\beta,Lys}$), 31.3 (<u>C</u>$H_2CH_2CH_3$), 29.0–28.2 (9C, (<u>C</u>$H_2$)$_{10}$$CH_3$), 26.9 ($C_{\delta,Lys}$), 23.1 ($C_{\gamma,Lys}$), 22.1 (<u>C</u>$H_2CH_3$), 13.9 ($CH_3$). Analytical HPLC gradient 0–95% eluent II in eluent I (C8; 35 min total runtime), $t_R$ 32.9 min (>98%, $UV_{230}$). HRMS calcd for $C_{34}H_{52}N_3O_3^+$ [M+H]$^+$, 582.3724; found 582.3732. CAS RN: 1429749-41-6. The data is in agreement with literature.[9]

**Benzyl ((*S*)-1-(((*S*)-3-(1*H*-indol-3-yl)-1-(isopropylamino)-1-oxopropan-2-yl)amino)-1-oxo-6-tetradecanethioamidohexan-2-yl)carbamate (2).**

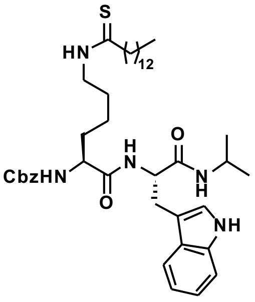

Compounds **S28** (22 mg, 0.04 mmol), **S29** (15 mg, 0.04 mmol), *i*$Pr_2NEt$ (11 µL, 0.06 mmol) and HOBt (9 mg, 0.07 mmol) were dissolved in anh. $CH_2Cl_2$ (3.0 mL) and cooled to 0 °C. EDC (14 mg, 0.07 mmol) was added and the reaction mixture was stirred at 0 °C for 5 min and then overnight at ambient temperature. The reaction mixture was concentrated under reduced pressure and resuspended in EtOAc (30 mL) and washed with aq. $KHSO_4$ (5%, 3×30 mL), saturated aq. $NaHCO_3$ (3×30 mL), and brine (2×30 mL). The organic phase was dried over $Na_2SO_4$, filtered and concentrated under reduced pressure. The crude residue was purified by column chromatography (0→2% MeOH in $CH_2Cl_2$) affording the desired amide **2** (22 mg, 68%) as a colorless solid. TLC (5% MeOH in $CH_2Cl_2$): $R_f$ = 0.5. $^1$H NMR (600 MHz, DMSO-*d*$_6$) δ 10.79 (d, *J* = 2.4 Hz, 1H, $NH_{Indole}$), 9.83 (t, *J* = 5.3 Hz, 1H, $NH_{\varepsilon,Lys}$), 7.86 (d, *J*

= 8.1 Hz, 1H, $NH_{\alpha,Trp}$), 7.66 (d, $J$ = 7.7 Hz, 1H, $CO_{Trp}NH$), 7.56 (d, $J$ = 7.9 Hz, 1H, $H4_{Indole}$), 7.45–7.23 (m, 7H, $NH_{\alpha,Lys}$, $H_{Ar,Cbz}$, $H7_{Indole}$), 7.11 (d, $J$ = 2.4 Hz, 1H, $H2_{Indole}$), 7.08–7.02 (m, 1H, $H6_{Indole}$), 6.96 (t, $J$ = 7.4 Hz, 1H, $H5_{Indole}$), 5.08–4.98 (m, 2H, $CH_{2,Cbz}$), 4.53–4.41 (m, 1H, $H_{\alpha,Trp}$), 3.96 (td, $J$ = 8.5, 5.2 Hz, 1H, $H_{\alpha,Lys}$), 3.84–3.72 (m, 1H, $CH_{iPr}$), 3.47–3.37 (m, 2H, $H_{\varepsilon,Lys}$), 3.06 ($m_{ABX}$, $J$ = 14.5, 6.1 Hz, 1H, $H_{\beta,Trp,A}$), 2.97 ($m_{ABX}$, $J$ = 14.6, 7.6 Hz, 1H, $H_{\beta,Trp,B}$), 2.51–2.48 (m, 2H, (C=S)$CH_2$, overlap with solvent peak), 1.69–1.42 (m, 6H, (C=S)$CH_2C\underline{H}_2$, $H_{\beta,Lys}$, $H_{\delta,Lys}$), 1.38–1.10 (m, 22H, $H_{\gamma,Lys}$, $(C\underline{H}_2)_{10}CH_3$), 1.00 (d, $J$ = 6.6 Hz, 3H, $CH_{3,iPr,A}$), 0.91 (d, $J$ = 6.6 Hz, 3H, $CH_{3,iPr,B}$), 0.85 (t, $J$ = 7.0 Hz, 3H, $CH_2C\underline{H}_3$). $^{13}$C NMR (151 MHz, DMSO-$d_6$) δ 204.0 (C=S), 171.9 ($CO_{Lys}$), 170.5 ($CO_{Trp}$), 156.4 ($CO_{Cbz}$), 137.4 ($C1_{Ar,Cbz}$), 136.4 ($C7a_{Indole}$), 128.8 ($C3_{Ar,Cbz}$, $C5_{Ar,Cbz}$), 128.2 ($C4_{Ar,Cbz}$), 128.1 ($C2_{Ar,Cbz}$, $C6_{Ar,Cbz}$), 127.9 ($C3a_{Indole}$), 124.0 ($C2_{Indole}$), 121.2 ($C6_{Indole}$), 118.9 ($C4_{Indole}$), 118.6 ($C5_{Indole}$), 111.6 ($C7_{Indole}$), 110.3 ($C3_{Indole}$), 65.9 ($CH_{2,Cbz}$), 55.3 ($C_{\alpha,Lys}$), 53.8 ($C_{\alpha,Trp}$), 45.5 ((C=S)$\underline{C}H_2$), 45.4 ($C_{\varepsilon,Lys}$), 40.9 ($CH_{iPr}$), 32.0 ($C_{\beta,Lys}$), 31.8 ($\underline{C}H_2CH_2CH_3$), 29.5–28.7 (9C, $(\underline{C}H_2)_{11}CH_3$), 28.4 ($C_{\beta,Trp}$), 27.3 ($C_{\delta,Lys}$), 23.4 ($C_{\gamma,Lys}$), 22.7 ($CH_{3,iPr,A}$), 22.6 ($CH_{3,iPr,B}$), 22.6 ($\underline{C}H_2CH_3$), 14.4 ($CH_2\underline{C}H_3$). Analytical HPLC gradient 0–95% eluent II in eluent I (C8; 35 min total runtime), $t_R$ 33.8 min (>98%, $UV_{230}$). HRMS calcd for $C_{42}H_{64}N_5O_4S^+$ $[M+H]^+$, 734.4674; found 734.4666.

***N*-dodecyl-1*H*-benzo[*d*][1,2,3]triazole-1-carbothioamide (S26).** A solution of dodecylamine (189 mg, 1.02 mmol) and $i$Pr$_2$NEt (0.2 mL, 1.15 mmol) in anh. $CH_2Cl_2$ (5.0 mL) was added dropwise over 10 min to a solution of bis(1-benzotriazolyl)methanethione (311 mg, 1.11 mmol) in anh. $CH_2Cl_2$ (8 mL) at 0 ˚C. The reaction mixture was stirred overnight at ambient temperature and concentrated under reduced pressure. The crude residue was purified by column chromatography (0→6% EtOAc in heptane), affording the desired compound **S26** (304 mg, 86%) as a colorless solid. TLC (25% EtOAc in heptane): $R_f$ = 0.6. $^1$H NMR (600 MHz, $CDCl_3$) δ 9.07 (br s, 1H, NH), 8.96–8.91 (m, 1H, $H7_{Bt}$), 8.12–8.07 (m, 1H, $H4_{Bt}$), 7.64 (ddd, $J$ = 8.3, 7.0, 1.1 Hz, 1H, $H6_{Bt}$), 7.47 (ddd, $J$ = 8.2, 7.0, 1.0 Hz, 1H, $H5_{Bt}$), 3.84 (td, $J$ = 7.3, 5.5 Hz, 2H, $NHC\underline{H}_2$), 1.81 (p, $J$ = 7.4 Hz, 2H, $NHCH_2C\underline{H}_2$), 1.49–1.42 (m, 2H, $NH(CH_2)_2C\underline{H}_2$), 1.42–1.20 (m, 16H, $(C\underline{H}_2)_8CH_3$), 0.88 (t, $J$ = 7.0 Hz, 3H, $CH_3$). $^{13}$C NMR (151 MHz, $CDCl_3$) δ 174.5 (C=S), 147.2 ($C7a_{Bt}$), 132.6 ($C3a_{Bt}$), 130.4 ($C6_{Bt}$), 125.8 ($C5_{Bt}$), 120.4 ($C4_{Bt}$), 116.2 ($C7_{Bt}$), 45.3 ($NH\underline{C}H_2$), 32.0 ($(\underline{C}H_2)_8CH_3$), 29.8–29.4 (6C, $(\underline{C}H_2)_8CH_3$), 28.3 ($(\underline{C}H_2)_8CH_3$), 27.2 ($NHCH_2\underline{C}H_2$), 22.8 ($NH(CH_2)_2\underline{C}H_2$), 14.3 ($CH_3$). HRMS calcd for $C_{19}H_{30}N_4NaS^+$ $[M+Na]^+$, 369.2089; found 369.2081. **Note**: The crude is sufficiently pure to be used without further purification. Bt = benzotriazole.

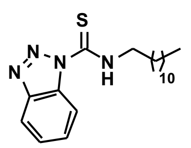

**Benzyl ((5*S*,8*S*)-5-((1*H*-indol-3-yl)methyl)-2-methyl-4,7-dioxo-14-thioxo-3,6,13,15-tetraazaheptacosan-8-yl)carbamate (3).**

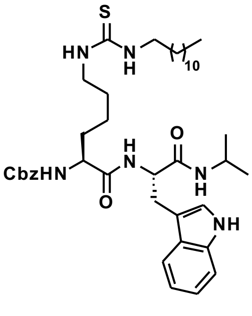

Compounds **S26** (42 mg, 1.21 mmol), **S30** (50 mg, 0.08 mmol), and *i*$Pr_2$NEt (21 µL, 0.12 mmol) were dissolved in anh. DMF (3.0 mL). The reaction mixture was stirred at ambient temperature for 2 h and concentrated under reduced pressure. The crude residue was purified by column chromatography (0→100% EtOAc in heptane), affording the desired thiourea **3** (34 mg, 38%) as a colorless solid. TLC (75% EtOAc in heptane): $R_f$ = 0.6. $^1$H NMR (600 MHz, DMSO-$d_6$) δ 10.79 (br s, 1H, $NH_{Indole}$), 7.86 (d, *J* = 8.1 Hz, 1H, $NH_{\alpha,Trp}$), 7.66 (d, *J* = 7.7 Hz, 1H, $CO_{Trp}$NH), 7.55 (d, *J* = 7.9 Hz, 1H, $H4_{Indole}$), 7.45–7.18 (m, 9H, $NH_{\alpha,Lys}$, $NH_{\varepsilon,Lys}$, $H_{Ar,Cbz}$, $H7_{Indole,}$ N<u>H</u>$CH_2$), 7.10 (d, *J* = 2.4 Hz, 1H, $H2_{Indole}$), 7.04 (t, *J* = 7.5 Hz, 1H, $H6_{Indole}$), 6.95 (t, *J* = 7.4 Hz, 1H, $H5_{Indole}$), 5.08–4.97 (m, 2H, $CH_{2,Cbz}$), 4.51–4.41 (m, 1H, $H_{\alpha,Trp}$), 3.95 (td, *J* = 8.4, 5.1 Hz, 1H, $H_{\alpha,Lys}$), 3.77 (h, *J* = 6.7 Hz, 1H, $CH_{iPr}$), 3.27 (br s, 2H, $H_{\varepsilon,Lys}$, overlap with residual water), 3.06 ($m_{ABX}$, *J* = 14.6, 6.1 Hz, 1H, $H_{\beta,Trp,A}$), 2.97 ($m_{ABX}$, *J* = 14.6, 7.7 Hz, 1H, $H_{\beta,Trp,B}$), 1.61–1.35 (m, 6H, $H_{\beta,Lys}$, $H_{\delta,Lys}$, NHC<u>H</u>$_2$), 1.32–1.13 (m, 22H, $H_{\gamma,Lys}$, (C<u>H</u>$_2$)$_{10}$$CH_3$), 1.00 (d, *J* = 6.6 Hz, 3H, $CH_{3,iPr,A}$), 0.90 (d, *J* = 6.5 Hz, 3H, $CH_{3,iPr,B}$), 0.85 (t, *J* = 6.8 Hz, 3H, $CH_2$C<u>H</u>$_3$). $^{13}$C NMR (151 MHz, DMSO-$d_6$) δ 182.0 (C=S), 171.5 ($CO_{Lys}$), 170.0 ($CO_{Trp}$), 156.0 ($CO_{Cbz}$), 137.0 ($C1_{Ar,Cbz}$), 135.9 ($C7a_{Indole}$), 128.3 ($C3_{Ar,Cbz}$, $C5_{Ar,Cbz}$), 127.6 ($C2_{Ar,Cbz}$, $C4_{Ar,Cbz}$, $C6_{Ar,Cbz}$), 127.4 ($C3a_{Indole}$), 123.5 ($C2_{Indole}$), 120.7 ($C6_{Indole}$), 118.4 ($C4_{Indole}$), 118.1 ($C5_{Indole}$), 111.1 ($C7_{Indole}$), 109.8 ($C3_{Indole}$), 65.4 ($CH_{2,Cbz}$), 54.9 ($C_{\alpha,Lys}$), 53.3 ($C_{\alpha,Trp}$), 43.4 ($C_{\varepsilon,Lys}$), 40.4 ($CH_{iPr}$), 31.6 ($C_{\beta,Lys}$), 31.3 (<u>C</u>$H_2CH_2CH_3$), 29.0–28.5 (9C, (<u>C</u>$H_2$)$_{11}$$CH_3$), $C_{\delta,Lys}$), 27.9 ($C_{\beta,Trp}$), 26.4 ((<u>C</u>$H_2$)$_{11}$$CH_3$), 22.9 ($C_{\gamma,Lys}$), 22.2 (2×$CH_{3,iPr}$), 22.1 (<u>C</u>$H_2CH_3$), 13.9 ($CH_2$<u>C</u>$H_3$). Analytical HPLC gradient 0–95% eluent II in eluent I (C18; 35 min total runtime), $t_R$ 31.0 min (>98%, $UV_{230}$). The peak for $C_{\varepsilon,Lys}$ was broad and of low intensity and the peak for C=S was not visible in $^{13}$C NMR due to fast quadrupolar relaxation via the nearby $^{14}$N-nuclei. HRMS calcd for $C_{41}H_{63}N_6O_4S^+$ [M+H]$^+$, 735.4626; found 735.4617.

**Benzyl ((*S*)-1-(((*S*)-3-(1*H*-indol-3-yl)-1-(isopropylamino)-1-oxopropan-2-yl)amino)-1-oxo-5-(2-tetradecanoylhydrazinyl)pentan-2-yl)carbamate (4).**

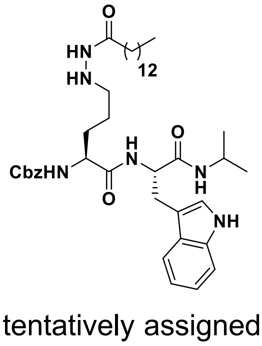

Myristoyl chloride (45 µL, 0.16 mmol) and *i*$Pr_2$NEt (29 µL, 0.16 mmol) were added to a solution of **S31** (50 mg, 0.08 mmol) in anh. $CH_2Cl_2$ (2.0 mL) at 0 °C. The reaction mixture was stirred at ambient temperature overnight and was then concentrated under reduced pressure. The crude residue was purified by column chromatography (0→2% MeOH and 1.0% AcOH in $CH_2Cl_2$) to afford an off-white solid (35 mg), tentatively assigned as *tert*-butyl 1-((*S*)-5-(((*S*)-3-(1*H*-indol-3-yl)-1-(isopropylamino)-1-oxopropan-2-yl)amino)-4-(((benzyloxy)carbonyl)amino)-5-oxopentyl)-2-tetradecanoyl-1,4-diazane-1-carboxylate (ESI-MS *m/z* calcd for $C_{46}H_{71}N_6O_7^+$ [M+H]$^+$, 819.5; found 819.6), which was used without further purification. TFA (1.5 mL) was added to a solution of the colorless solid (35 mg) in anh. $CH_2Cl_2$ (3.0 mL) and stirred for 1 h at ambient temperature. The reaction mixture was then concentrated under reduced pressure and excess TFA was removed by coevaporations: $CH_2Cl_2$/toluene (1:1, 2×25 mL) and $CH_2Cl_2$/MeCN (1:1, 2×25 mL). The crude residue was purified by column chromatography (0→3.5% MeOH and 1.0% AcOH in $CH_2Cl_2$) affording the desired hydrazide **4** (15 mg, 25%) as a colorless fluffy material after lyophilization. TLC (3.5% MeOH and 1% AcOH in $CH_2Cl_2$): $R_f$ = 0.4. $^1$H NMR (600 MHz, DMSO-$d_6$) δ 10.82 (br s, 1H, $NH_{Indole}$), 9.23 (s, 1H, NHN$\underline{H}$CO), 7.95 (d, $J$ = 8.0 Hz, 1H, $NH_{\alpha,Trp}$), 7.70 (d, $J$ = 7.6 Hz, 1H, $CO_{Trp}NH$), 7.55 (d, $J$ = 7.9 Hz, 1H, $H4_{Indole}$), 7.45 (d, $J$ = 7.8 Hz, 1H, $NH_{\alpha,Lys}$), 7.39–7.20 (m, 6H, $H_{Ar,Cbz}$, $H7_{Indole}$), 7.11 (d, $J$ = 2.3 Hz, 1H, $H2_{Indole}$), 7.03 (t, $J$ = 7.3 Hz, 1H, $H6_{Indole}$), 6.95 (t, $J$ = 7.4 Hz, 1H, $H5_{Indole}$), 5.02 (q, $J$ = 12.6 Hz, 2H, $CH_{2,Cbz}$), 4.74 (br s, 1H, N$\underline{H}$NHCO), 4.56–4.38 (m, 1H, $H_{\alpha,Trp}$), 4.09–3.90 (m, 1H, $H_{\alpha,Lys}$), 3.90–3.67 (m, 1H, $CH_{iPr}$), 3.06 ($m_{ABX}$, $J$ = 14.5, 5.9 Hz, 1H, $H_{\beta,Trp,A}$), 2.97 ($m_{ABX}$, $J$ = 14.5, 7.7 Hz, 1H, $H_{\beta,Trp,B}$), 2.66–2.53 (m, 2H, $H_{\delta,Lys}$), 1.99 (t, $J$ = 7.4 Hz, 2H, NHNHCOC$\underline{H}_2$), 1.64–1.40 (m, 4H, $H_{\beta,Lys}$, $H_{\gamma,Lys}$), 1.40–1.08 (m, 24H, $(C\underline{H}_2)_{12}CH_3$), 1.00 (d, $J$ = 6.6 Hz, 3H, $CH_{3,iPr,A}$), 0.91 (d, $J$ = 6.5 Hz, 3H, $CH_{3,iPr,B}$), 0.85 (t, $J$ = 7.0 Hz, 3H, $(CH_2)C\underline{H}_3$). $^{13}$C NMR (151 MHz, DMSO-$d_6$) δ 171.5 ($CO_{Lys}$), 171.0 (NHNHCO), 170.1 ($CO_{Trp}$), 156.0 ($CO_{Cbz}$), 137.0 ($C1_{Ar,Cbz}$), 136.0 ($C7a_{Indole}$), 128.3 ($C3_{Ar,Cbz}$, $C5_{Ar,Cbz}$), 127.7 ($C4_{Ar,Cbz}$), 127.6 ($C2_{Ar,Cbz}$, $C6_{Ar,Cbz}$), 127.4 ($C3a_{Indole}$), 123.5 ($C2_{Indole}$), 120.7 ($C6_{Indole}$), 118.4 ($C4_{Indole}$), 118.1 ($C5_{Indole}$), 111.1 ($C7_{Indole}$), 109.9 ($C3_{Indole}$), 65.4 ($CH_{2,Cbz}$), 54.9 ($C_{\alpha,Lys}$), 53.4 ($C_{\alpha,Trp}$), 50.7 ($C_{\delta,Lys}$), 40.4 ($CH_{iPr}$), 33.5 (NHNHCO$\underline{C}H_2$), 31.3 ($\underline{C}H_2CH_2CH_3$), 29.4 ($C_{\beta,Lys}$), 29.0–28.6 (9C, $(\underline{C}H_2)_{11}CH_3$), 25.2 ($C_{\beta,Trp}$), 23.8 ($C_{\gamma,Lys}$), 22.2 ($\underline{C}H_2CH_3$), 22.1 ($CH_{3,iPr}$), 13.9 ($CH_2\underline{C}H_3$). Analytical HPLC gradient 0–95% eluent II in eluent I (C8; 35 min total runtime), $t_R$ 29.4 min (N/A, $UV_{230}$*). HRMS calcd for $C_{41}H_{63}N_6O_5^+$ $[M+H]^+$, 719.4854; found 719.4871.

*Degrades during reversed-phase HPLC.

**Benzyl ((*S*)-1-(((*S*)-3-(1*H*-indol-3-yl)-1-(isopropylamino)-1-oxopropan-2-yl)amino)-8-(dodecylamino)-1,8-dioxooctan-2-yl)carbamate (5).** Compound **S32** (51 mg, 0.09 mmol), HOBt (19 mg, 0.14 mmol), dodecylamine (25 mg, 0.14 mmol) and $i$Pr$_2$NEt (32 µL, 0.18 mmol) were dissolved in anh. $CH_2Cl_2$ (2.0 mL) and cooled to 0 °C. EDC (26 mg, 0.14 mmol) was added and the reaction mixture was stirred at 0°C for 15 min and then overnight at ambient temperature. The reaction mixture was diluted with EtOAc (50 mL) and washed with aq. $KHSO_4$ (5%, 3×50 mL), saturated aq. $NaHCO_3$ (3×50 mL) and brine (2×50 mL). The organic phase was dried over $Na_2SO_4$, filtered, and concentrated under reduced pressure. The crude residue was purified by column chromatography (0→3% MeOH and 1% AcOH in $CH_2Cl_2$) affording the desired amide **5** (39 mg, 59%) as a colorless solid. TLC (3% MeOH and 1% AcOH in $CH_2Cl_2$): $R_f$ = 0.3. $^1$H NMR (600 MHz, DMSO-$d_6$) δ 10.82 (s, 1H, $NH_{Indole}$), 7.88 (d, $J$ = 8.1 Hz, 1H, $NH_{\alpha,Trp}$), 7.73 (t, $J$ = 5.4 Hz, 1H, CON$\underline{H}$CH$_2$), 7.69 (d, $J$ = 7.7 Hz, 1H, $CO_{Trp}NH$), 7.55 (d, $J$ = 7.9 Hz, 1H, $H4_{Indole}$), 7.44 (d, $J$ = 7.8 Hz, 1H, $NH_{\alpha,Asu}$), 7.39–7.22 (m, 6H, $H_{Ar,Cbz}$, $H7_{Indole}$), 7.10 (d, $J$ = 2.4 Hz, 1H, $H2_{Indole}$), 7.03 (t, $J$ = 7.5 Hz, 1H, $H6_{Indole}$), 6.95 (t, $J$ = 7.4 Hz, 1H $H5_{Indole}$), 5.10–4.95 (m, 2H, $CH_{2,Cbz}$), 4.52–4.38 (m,

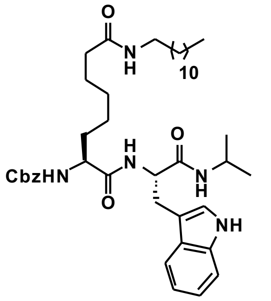

1H, $H_{\alpha,Trp}$), 3.99–3.86 (m, 1H, $H_{\alpha,Asu}$), 3.83–3.68 (m, 1H, $CH_{iPr}$), 3.14–2.89 (m, 4H, $H_{\beta,Trp}$, $CONHC\underline{H}_2$), 2.00 (t, *J* = 7.4 Hz, 2H, $H_{\zeta,Asu}$), 1.55–1.47 (m, 1H, $H_{\beta,Asu,A}$), 1.47–1.30 (m, 5H, $H_{\beta,Asu,B}$, $H_{\gamma,Asu}$, $H_{\varepsilon,Asu}$), 1.29–1.10 (m, 22H, $(C\underline{H}_2)_{10}CH_3$, $H_{\delta,Asu}$), 0.99 (d, *J* = 6.6 Hz, 3H, $CH_{3,iPr,A}$), 0.91 (d, *J* = 6.5 Hz, 3H, $CH_{3,iPr,B}$), 0.85 (t, *J* = 6.9 Hz, 3H, $CH_2C\underline{H}_3$). $^{13}C$ NMR (151 MHz, DMSO-$d_6$) δ 171.9 ($CO_{\alpha,Asu}$), 171.5 ($CO_{\eta,Asu}$), 170.1 ($CO_{Trp}$), 156.0 ($CO_{Cbz}$), 137.0 ($C1_{Ar,Cbz}$), 136.0 ($C7a_{Indole}$), 128.4 ($C3_{Ar,Cbz}$, $C5_{Ar,Cbz}$), 127.8 ($C4_{Ar,Cbz}$), 127.7 ($C2_{Ar,Cbz}$, $C6_{Ar,Cbz}$), 127.4 ($C3a_{Indole}$), 123.5 ($C2_{Indole}$), 120.8 ($C6_{Indole}$), 118.5 ($C4_{Indole}$), 118.1 ($C5_{Indole}$), 111.2 ($C7_{Indole}$), 109.9 ($C3_{Indole}$), 65.4 ($CH_{2,Cbz}$), 55.0 ($C_{\alpha,Asu}$), 53.3 ($C_{\alpha,Trp}$), 40.4 ($C_{\alpha,Lys}$), 38.4 ($CONH\underline{C}H_2$), 35.4 ($C_{\zeta,Asu}$), 31.8 ($C_{\beta,Asu}$), 31.3 ($\underline{C}H_2CH_2CH_3$), 29.2 ($C_{\varepsilon,Asu}$), 29.0 ($(\underline{C}H_2)_{10}CH_3$), 28.8 ($(\underline{C}H_2)_{10}CH_3$), 28.4 ($(\underline{C}H_2)_{10}CH_3$), 27.9 ($C_{\beta,Trp}$), 26.4 ($(\underline{C}H_2)_{10}CH_3$), 25.2 ($C_{\gamma,Asu}$), 25.1 ($C_{\delta,Asu}$), 22.3 ($CH_{3,iPr,A}$), 22.1 ($CH_{3,iPr,B}$), 14.0 ($CH_2\underline{C}H_3$). Analytical HPLC gradient 0–95% eluent II in eluent I (C8; 35 min total runtime), $t_R$ 31.3 min (>98%, $UV_{230}$). HRMS calcd for $C_{42}H_{64}N_5O_5^+$ $[M+H]^+$, 718.4902; found 718.4911. Asu = aminosuberic acid.

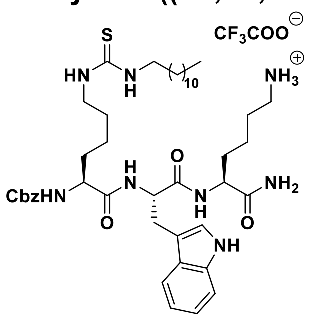

**Benzyl ((5*S*,8*S*,11*S*)-8-((1*H*-indol-3-yl)methyl)-1-amino-5-carbamoyl-7,10-dioxo-17-thioxo-6,9,16,18-tetraazatriacontan-11-yl)carbamate (6).** Starting from Cbz-Lys-Trp-Lys(Boc)-resin (101 mg, estimated loading: 0.42 mmol/g) synthesized from Fmoc-Lys(Boc)-OH, Fmoc-Trp-OH and Cbz-Lys(Fmoc)-OH by SPPS, the title compound was synthesized by on-resin thiourea formation as described in the general procedures. Global deprotection and cleavage from the resin, followed by preparative reversed-phase HPLC purification afforded the desired thiourea **6** (4 mg, 10% based on resin loading) as a colorless fluffy material after lyophilization. $^1H$ NMR (600 MHz, DMSO-$d_6$) δ 10.82 (d, *J* = 2.4 Hz, 1H, $NH_{Indole}$), 7.97 (d, *J* = 7.6 Hz, 1H, $NH_{\alpha,Trp}$), 7.92 (d, *J* = 8.1 Hz, 1H, $NH_{\alpha,Lys}$), 7.70 (s, 3H, $NH_3^+$), 7.57 (d, *J* = 7.9 Hz, 1H, $H4_{Indole}$), 7.45–7.24 (m, 9H, $NH_{\alpha,Lys(Dtu)}$, $NH_{\varepsilon,Lys}$, $H_{Ar,Cbz}$, $H7_{Indole,}$ $N\underline{H}CH_2$), 7.20 (s, 1H, $CONH_{2,A}$), 7.14 (d, *J* = 2.3 Hz, 1H, $H2_{Indole}$), 7.09–7.03 (m, 2H, $H6_{Indole}$, $CONH_{2,B}$), 6.96 (t, *J* = 7.4 Hz, 1H, $H5_{Indole}$), 6.56 (s, 1H, $CO_2H_{TFA}$) 5.09–4.94 (m, 2H, $CH_{2,Cbz}$), 4.54 (dt, *J* = 8.1, 3.9 Hz, 1H, $H_{\alpha,Trp}$), 4.17 (td, *J* = 8.4, 5.1 Hz, 1H, $H_{\alpha,Lys}$), 3.94 (td, *J* = 8.6, 4.8 Hz, 1H, $H_{\alpha,Lys(Dtu)}$), 3.27 (br s, 2H, $H_{\varepsilon,Lys(Dtu)}$, overlap with residual water), 3.15 ($m_{ABX}$, *J* = 14.8, 4.9 Hz, 1H, $H_{\beta,Trp,A}$), 2.99 ($m_{ABX}$, *J* = 14.9, 8.6 Hz, 1H, $H_{\beta,Trp,B}$), 2.80–2.69 (m, 2H, $H_{\varepsilon,Lys}$), 1.72–1.35 (m, 10H, $H_{\beta,Lys(Dtu)}$, $H_{\delta,Lys(Dtu)}$,$H_{\beta,Lys}$, $H_{\delta,Lys}$, $(C\underline{H}_2)_{11}CH_3$), 1.24 (s, 24H, $H_{\gamma,Lys(Dtu)}$, $H_{\gamma,Lys}$, $(C\underline{H}_2)_{11}CH_3$), 0.85 (t, *J* = 6.9 Hz, 3H, $CH_3$). $^{13}C$ NMR (151 MHz, DMSO-$d_6$) δ 173.3 ($CO_{Lys}$), 172.0 ($CO_{Lys(Dtu)}$), 171.2 ($CO_{Trp}$), 158.0 (d, *J* = 33.1 Hz, residual $CO_{TFA}$), 156.0 ($CO_{Cbz}$), 136.9 ($C1_{Ar,Cbz}$), 136.0 ($C7a_{Indole}$), 128.3 ($C3_{Ar,Cbz}$, $C5_{Ar,Cbz}$), 127.8 ($C4_{Ar,Cbz}$), 127.7 ($C2_{Ar,Cbz}$, $C6_{Ar,Cbz}$), 127.3 ($C3a_{Indole}$), 123.6 ($C2_{Indole}$), 120.8 ($C6_{Indole}$), 118.4 ($C4_{Indole}$), 118.2 ($C5_{Indole}$), 117.3 (q, *J* = 283.1 Hz, residual $CF_{3,TFA}$), 111.2 ($C7_{Indole}$), 109.8 ($C3_{Indole}$), 65.5 ($CH_{2,Cbz}$), 54.9 ($C_{\alpha,Lys(Dtu)}$), 53.4 ($C_{\alpha,Trp}$), 52.2 ($C_{\alpha,Lys}$), 43.3 ($C_{\varepsilon,Lys(Dtu)}$), 38.7 ($C_{\varepsilon,Lys}$), 31.6 ($C_{\beta,Lys}$), 31.4 ($C_{\beta,Lys(Dtu)}$), 31.3 ($\underline{C}H_2CH_2CH_3$), 29.0–28.7 (9C, $(\underline{C}H_2)_{11}CH_3$), 27.3 ($C_{\beta,Trp}$), 26.7 ($C_{\delta,Lys(Dtu)}$), 26.4 ($C_{\delta,Lys}$), 23.0 ($C_{\gamma,Lys(Dtu)}$), 22.1 ($C_{\gamma,Lys}$), 22.1 ($\underline{C}H_2CH_3$), 14.0 ($CH_3$). The peak for $C_{\varepsilon,Lys(Dtu)}$ was broad and of low intensity and the peak for

C=S was not visible in $^{13}$C NMR due to fast quadrupolar relaxation via the nearby $^{14}$N-nuclei. Analytical HPLC gradient 0–95% eluent II in eluent I (C18; 35 min total runtime), $t_R$ 25.4 min (>98%, $UV_{230}$). HRMS calcd for $C_{44}H_{69}N_8O_5S^+$ $[M+H]^+$, 821.5106; found 821.5105. Dtu = 1-dodecylthiourea.

**Benzyl ((3S,6S,9S)-6-((1H-indol-3-yl)methyl)-1-amino-3-carbamoyl-1,5,8-trioxo-15-thioxo-4,7,14,16-tetraazaoctacosan-9-yl)carbamate (7).**

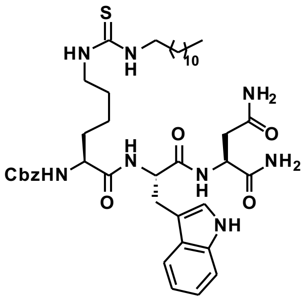

Starting from Cbz-Lys-Trp(Boc)-Asn(Trt)-resin (267 mg, estimated loading: 0.37 mmol/g) synthesized from Fmoc-Asn(Trt)-OH, Fmoc-Trp(Boc)-OH and Cbz-Lys(Fmoc)-OH by SPPS, the title compound was synthesized by on-resin thiourea formation as described in the general procedures. Global deprotection and cleavage from the resin, followed by preparative reversed-phase HPLC purification afforded the desired thiourea **7** (3 mg, 4% based on resin loading) as a colorless fluffy material after lyophilization. $^1$H NMR (600 MHz, DMSO-$d_6$) δ 10.82 (d, $J$ = 2.4 Hz, 1H, $NH_{Indole}$), 8.08 (d, $J$ = 8.0 Hz, 1H, $NH_{\alpha,Trp}$), 8.03 (d, $J$ = 7.4 Hz, 1H, $NH_{\alpha,Asn}$), 7.56 (d, $J$ = 7.9 Hz, 1H, $H4_{Indole}$), 7.40–7.24 (m, 9H, $NH_{\alpha,Lys}$, $NH_{\varepsilon,Lys}$, $H_{Ar,Cbz}$, $H7_{Indole,}$ $N\underline{H}CH_2$), 7.16 (d, $J$ = 2.4 Hz, 1H, $H2_{Indole}$), 7.06–7.01 (m, 2H, $H6_{Indole}$, $CONH_{2,\ Asn,\alpha,B}$), 6.96 (t, $J$ = 7.4 Hz, 1H, $H5_{Indole}$), 6.93 (s, 1H, $CONH_{2,\gamma,A}$), 6.85 (s, 1H, $CONH_{2,\gamma,B}$), 5.04 (d, $J$ = 12.6 Hz, 1H, $CH_{2,Cbz,\ A}$), 4.98 (d, $J$ = 12.5 Hz, 1H, $CH_{2,Cbz,\ B}$), 4.52–4.41 (m, 2H, $H_{\alpha,Trp}$, $H_{\alpha,Asn}$), 3.95 (td, $J$ = 8.7, 4.9 Hz, 1H, $H_{\alpha,Lys}$), 3.31 (br s, 2H, $H_{\varepsilon,Lys}$, overlap with residual water), 3.15 ($m_{ABX}$, $J$ = 14.9, 4.7 Hz, 1H, $H_{\beta,Trp,A}$), 2.97 ($m_{ABX}$, $J$ = 14.9, 8.8 Hz, 1H, $H_{\beta,Trp,B}$), 2.44 (d, $J$ = 6.5 Hz, 2H, $H_{\beta,Asn}$), 1.63–1.34 (m, 6H, $H_{\beta,Lys}$, $H_{\delta,Lys}$, $(C\underline{H}_2)_{11}CH_3$), 1.32–1.11 (m, 22H, $H_{\gamma,Lys}$, $(C\underline{H}_2)_{11}CH_3$), 0.85 (t, $J$ = 7.0 Hz, 3H, $CH_3$). $^{13}$C NMR (151 MHz, DMSO-$d_6$) δ 172.7 ($CO_{\alpha,Asn}$), 172.2 ($CO_{Lys}$), 171.7 ($CO_{Asn,\gamma}$), 171.0 ($CO_{Trp}$), 156.0 ($CO_{Cbz}$), 136.9 ($C1_{Ar,Cbz}$), 136.0 ($C7a_{Indole}$), 128.3 ($C2_{Ar,Cbz}$, $C5_{Ar,Cbz}$), 127.7 ($C2_{Ar,Cbz}$, $C6_{Ar,Cbz}$), 127.3 ($C3a_{Indole}$), 123.6 ($C2_{Indole}$), 120.8 ($C6_{Indole}$), 118.3 ($C4_{Indole}$), 118.2 ($C5_{Indole}$), 111.2 ($C7_{Indole}$), 109.8 ($C3_{Indole}$), 65.6 ($CH_{2,Cbz}$), 54.7 ($C_{\alpha,Lys}$), 53.6 ($C_{\alpha,Trp}$), 49.4 ($C_{\alpha,Asn}$), 43.5 ($C_{\varepsilon,Lys}$), 36.7 ($C_{\beta,Asn}$), 31.5 ($C_{\beta,Lys}$), 31.3 ($\underline{C}H_2CH_2CH_3$), 29.0–28.7 (9C, $(\underline{C}H_2)_{11}CH_3$), 27.3 ($C_{\beta,Trp}$), 26.4 ($C_{\delta,Lys}$), 22.9 ($C_{\gamma,Lys}$), 22.1 ($\underline{C}H_2CH_3$), 13.9 ($CH_3$). The peak for $C_{\varepsilon,Lys}$ was broad and of low intensity and the peak for C=S was not visible in $^{13}$C NMR due to fast quadrupolar relaxation via the nearby $^{14}$N-nuclei. Analytical HPLC gradient 0–95% eluent II in eluent I (C18; 35 min total runtime), $t_R$ 27.3 min (>98%, $UV_{230}$). HRMS calcd for $C_{42}H_{62}N_8O_6SNa^+$ $[M+Na]^+$, 829.4405; found 829.4416.

**Benzyl ((5*S*,8*S*,11*S*)-8-((1*H*-indol-3-yl)methyl)-1-amino-5-carbamoyl-1-imino-7,10-dioxo-17-thioxo-2,6,9,16,18-pentaazatriacontan-11-yl)carbamate (8).**

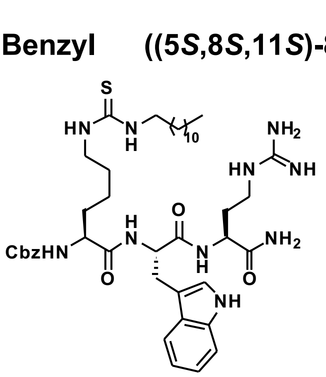

Starting from Cbz-Lys(Fmoc)-Trp-Dab(Alloc)-resin (128 mg, estimated loading: 0.47 mmol/g) synthesized from Fmoc-Dab(Alloc)-OH, Fmoc-Trp-OH and Cbz-Lys(Fmoc)-OH by SPPS, the title compound was synthesized by addition of borane dimethylamine complex (24 mg, 0.40 mmol) and $Pd(PPh_3)_4$ (20 mg, 0.017 mmol) in anh. $CH_2Cl_2$ (2.0 mL) to the resin under agitation for 15 min. The procedure was repeated and the resin was washed with $CH_2Cl_2$ (3×4 mL), DMF (3×4 mL) and $CH_2Cl_2$ (3×4 mL). A solution of pyrazol$(Boc)_2$ (56 mg, 0.18 mmol) and *i*$Pr_2NEt$ (63 µL, 0.32 mmol) in DMF (4 mL) was added to the resin and agitated for 3 h at 37 ˚C. The resin was washed with DMF (3×4 mL) followed by Fmoc-deprotection and subsequent on-resin thiourea formation as described in the general procedures. Global deprotection and cleavage from the resin, followed by preparative reversed-phase HPLC purification afforded the desired thiourea **8** (4 mg, 8% based on resin loading) as a colorless fluffy material after lyophilization. $^{1}$H NMR (600 MHz, DMSO-$d_6$) δ 10.83 (d, $J$ = 2.3, 1H, $NH_{Indole}$), 8.19–8.02 (m, 2H, $NH_{\alpha,Agb}$, $NH_{\alpha,Trp}$), 7.58 (d, $J$ = 7.9 Hz, 1H, $H4_{Indole}$), 7.50 (t, $J$ = 5.8 Hz, 1H, $NH_{\gamma,Agb}$), 7.41 (d, $J$ = 7.8 Hz, 1H, $NH_{\alpha,Lys}$), 7.38–7.27 (m, 8H, $NH_{\varepsilon,Lys}$, $H_{Ar,Cbz}$, $H7_{Indole,}$ $N\underline{H}CH_2$), $CONH_{2,A}$), 7.18–7.10 (m, 3H, $CONH_2$, $H2_{Indole}$), 7.07–7.04 (m, 2H, $H6_{Indole}$), 6.97 (t, $J$ = 7.4 Hz, 1H, $H5_{Indole}$), 5.08–4.92 (m, 2H, $CH_{2,Cbz}$), 4.56–4.46 (m, 1H, $H_{\alpha,Trp}$), 4.27–4.19 (m, 1H, $H_{\alpha,Agb}$), 3.99–3.91 (m, 1H, $H_{\alpha,Lys}$), 3.30 (br s, 2H, $H_{\varepsilon,Lys}$, overlap with residual water), 3.21–3.10 (m, 2H, $H_{\beta,Trp,B}$, $H_{\gamma,Agb,A}$), 3.09–2.97 (m, 2H, $H_{\beta,Trp,B}$, $H_{\gamma,Agb,B}$), 2.89 (s, trace DMF), 2.73 (s, trace DMF), 1.98–1.88 (m, 1H, $H_{\beta,Agb,A}$), 1.73–1.63 (m, 1H, $H_{\beta,Agb,B}$), 1.59–1.55 (m, 1H, $H_{\beta,Lys,A}$), 1.49–1.31 (m, 5H, $H_{\beta,Lys,B}$, $H_{\delta,Lys}$, $(C\underline{H_2})_{11}CH_3$), 1.32–1.09 (m, 24H, $H_{\gamma,Lys}$, $(C\underline{H_2})_{11}CH_3$), 0.85 (t, $J$ = 6.9 Hz, 3H, $CH_3$). $^{13}$C NMR (151 MHz, DMSO-$d_6$) δ 172.8 ($CO_{Agb}$), 172.4 ($CO_{Lys}$), 171.6 ($CO_{Trp}$), 156.7 ($NHC(=NH)NH_2$), 158.3 (q, $J$ = 32.6 Hz, residual $CO_{TFA}$), 156.1 ($CO_{Cbz}$), 136.9 ($C1_{Ar,Cbz}$), 136.0 ($C7a_{Indole}$), 128.3 ($C3_{Ar,Cbz}$, $C5_{Ar,Cbz}$), 127.8 ($C4_{Ar,Cbz}$), 127.7 ($C2_{Ar,Cbz}$, $C6_{Ar,Cbz}$), 127.3 ($C3a_{Indole}$), 123.6 ($C2_{Indole}$), 120.9 ($C6_{Indole}$), 118.4 ($C4_{Indole}$), 118.2 ($C5_{Indole}$), 111.3 ($C7_{Indole}$), 109.8 ($C3_{Indole}$), 65.5 ($CH_{2,Cbz}$), 54.8 ($C_{\alpha,Lys}$), 53.8 ($C_{\alpha,Trp}$), 50.2 ($C_{\alpha,Agb}$), 43.4 ($C_{\varepsilon,Lys}$), 37.8 ($C_{\gamma,Agb}$), 31.5 ($C_{\beta,Lys,}$ $C_{\beta,Agb}$), 31.3 ($\underline{C}H_2CH_2CH_3$), 29.0–28.5 (10C, $C_{\delta,Agb}$ $(\underline{C}H_2)_{11}CH_3$), 27.0 ($C_{\beta,Trp}$), 26.4 ($C_{\delta,Lys}$), 23.0 ($C_{\gamma,Lys}$), 22.1 ($\underline{C}H_2CH_3$), 13.9 ($CH_3$). The peak for $C_{\varepsilon,Lys}$ was broad and of low intensity and the peak for C=S was not visible in $^{13}$C NMR due to fast quadrupolar relaxation via the nearby $^{14}$N-nuclei. Analytical HPLC gradient 0–95% eluent II in eluent I (C8; 35 min total runtime), $t_R$ 26.3 min (>95%, $UV_{254}$). HRMS calcd for $C_{43}H_{67}N_{10}O_5S^+$ $[M+H]^+$, 835.5011; found 835.5005. Agb = 2-amino-guanidinobutyric acid.

**Benzyl ((6*S*,9*S*,12*S*)-9-((1*H*-indol-3-yl)methyl)-1-amino-6-carbamoyl-1-imino-8,11-dioxo-18-thioxo-2,7,10,17,19-pentaazahentriacontan-12-yl)carbamate (9).**

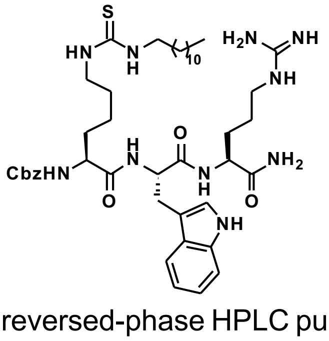

Starting from Cbz-Lys-Trp-Arg(Pbf)-resin (101 mg, estimated loading: 0.42 mmol/g) synthesized from Fmoc-Arg(Pbf)-OH, Fmoc-Trp-OH and Cbz-Lys(Fmoc)-OH by SPPS, the title compound was synthesized by on-resin thiourea formation as described in the general procedures. Global deprotection and cleavage from the resin, followed by preparative reversed-phase HPLC purification afforded the desired thiourea **9** (1 mg, 4% based on resin loading), as a colorless fluffy material after lyophilization. $^{1}$H NMR (600 MHz, DMSO-$d_6$) δ 10.81 (br s, 1H, $NH_{Indole}$), 8.01–7.91 (m, 2H, $NH_{\alpha,Arg}$, $NH_{\alpha,Trp}$), 7.56 (d, $J$ = 7.9 Hz, 1H, $H4_{Indole}$), 7.54–7.48 (m, 1H, $NH_{\delta,Arg}$), 7.42–7.27 (m, 9H, $NH_{\alpha,Lys}$, $NH_{\varepsilon,Lys}$, $H_{Ar,Cbz}$, $H7_{Indole,}$ $N\underline{H}CH_2$), 7.25 (s, 1H, $CONH_{2,A}$), 7.14 (d,

$J$ = 2.3 Hz, 1H, $H2_{Indole}$), 7.10 (s, 1H, $CONH_{2,B}$), 7.07–7.03 (m, 1H, $H6_{Indole}$), 6.96 (t, $J$ = 7.4 Hz, 1H, $H5_{Indole}$), 6.54 (s, 2H, residual $CO_2H_{TFA}$), 5.08–4.95 (m, 2H, $CH_{2,Cbz}$), 4.54 (td, $J$ = 8.0, 4.7 Hz, 1H, $H_{\alpha,Trp}$), 4.21 (td, $J$ = 8.0, 5.6 Hz, 1H, $H_{\alpha,Arg}$), 3.94 (td, $J$ = 8.7, 4.8 Hz, 1H, $H_{\alpha,Lys}$), 3.27 (br s, 2H, $H_{\varepsilon,Lys}$, overlap with residual water), 3.15 ($m_{ABX}$, $J$ = 14.8, 4.7 Hz, 1H, $H_{\beta,Trp,A}$), 3.11–3.04 (m, 2H, $H_{\delta,Arg}$), 2.99 ($m_{ABX}$, $J$ = 14.9, 8.7 Hz, 1H, $H_{\beta,Trp,B}$), 1.75–1.63 (m, 1H, $H_{\beta,Arg,A}$), 1.61–1.32 (m, 9H, $H_{\beta,Arg,B}$, $H_{\gamma,Arg}$, $H_{\beta,Lys}$, $H_{\delta,Lys}$, $(C\underline{H}_2)_{11}CH_3$), 1.24 (s, 22H, $H_{\gamma,Lys}$, $(C\underline{H}_2)_{11}CH_3$), 0.85 (t, $J$ = 7.0 Hz, 3H, $CH_3$). $^{13}C$ NMR (151 MHz, DMSO-$d_6$) δ 173.1 ($CO_{Arg}$), 172.0 ($CO_{Lys}$), 171.2 ($CO_{Trp}$), 157.9 (q, $J$ = 30.9 Hz, residual $CO_{TFA}$), 156.7 ($NHC(=NH)NH_2$), 156.0 ($CO_{Cbz}$), 136.9 ($C1_{Ar,Cbz}$), 136.0 ($C7a_{Indole}$), 128.3 ($C3_{Ar,Cbz}$, $C5_{Ar,Cbz}$), 127.8 ($C4_{Ar,Cbz}$), 127.7 ($C2_{Ar,Cbz}$, $C6_{Ar,Cbz}$), 127.3 ($C3a_{Indole}$), 123.6 ($C2_{Indole}$), 120.8 ($C6_{Indole}$), 118.4 ($C4_{Indole}$), 118.2 ($C5_{Indole}$), 117.3 (q, $J$ = 300.1 Hz, residual $CF_{3,TFA}$), 111.2 ($C7_{Indole}$), 109.8 ($C3_{Indole}$), 65.5 ($CH_{2,Cbz}$), 54.8 ($C_{\alpha,Lys}$), 53.4 ($C_{\alpha,Trp}$), 51.9 ($C_{\alpha,Arg}$), 43.4 ($C_{\varepsilon,Lys}$), 40.4 ($C_{\delta,Arg}$), 31.6 ($C_{\beta,Lys}$), 31.3 ($\underline{C}H_2CH_2CH_3$), 29.2 ($C_{\beta,Arg}$), 29.0–28.5 (9C, $(\underline{C}H_2)_{11}CH_3$), 27.4 ($C_{\beta,Trp}$), 26.4 ($C_{\delta,Lys}$), 25.0 ($C_{\gamma,Arg}$), 23.0 ($C_{\gamma,Lys}$), 22.1 ($\underline{C}H_2CH_3$), 13.9 ($CH_3$). The peak for $C_{\varepsilon,Lys}$ was broad and of low intensity and the peak for C=S was not visible in $^{13}C$ NMR due to fast quadrupolar relaxation via the nearby $^{14}N$-nuclei. Analytical HPLC gradient 0–95% eluent II in eluent I (C18; 35 min total runtime), $t_R$ 25.8 min (>95%, $UV_{230}$). HRMS calcd for $C_{44}H_{69}N_{10}O_5S^+$ [M+H]$^+$, 849.5168; found 849.5189.

**Benzyl ((7*S*,10*S*,13*S*)-10-((1*H*-indol-3-yl)methyl)-1-amino-7-carbamoyl-1-imino-9,12-dioxo-19-thioxo-2,8,11,18,20-pentaazadotriacontan-13-yl)carbamate (10).**

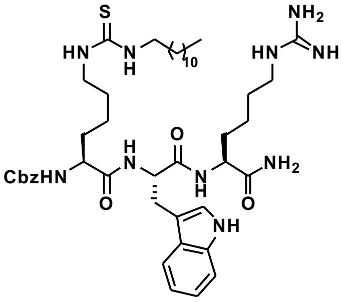

Starting from Cbz-Lys-Trp(Boc)-hArg($Boc_2$)-resin (267 mg, estimated loading: 0.37 mmol/g) synthesized from Fmoc-hArg($Boc_2$)-OH (**S28**), Fmoc-Trp(Boc)-OH and Cbz-Lys(Fmoc)-OH by SPPS, the title compound was synthesized by on-resin thiourea formation as described in the general procedures. Global deprotection and cleavage from the resin, followed by preparative reversed-phase HPLC purification afforded the desired thiourea **10** (4 mg, 5% based on resin loading), as a colorless fluffy material after lyophilization. $^1H$ NMR (600 MHz, DMSO-$d_6$) δ 10.81 (d, $J$ = 2.4, 1H, $NH_{Indole}$), 8.01–7.89 (m, 2H, $NH_{\alpha,hArg}$, $NH_{\alpha,Trp}$), 7.57 (d, $J$ = 7.9 Hz, 1H, $H4_{Indole}$), 7.47–7.43 (m, 1H, $NH_{\varepsilon,hArg}$), 7.40 (d, $J$ = 7.9 Hz, 1H, $NH_{\alpha,Lys}$), 7.38–7.26 (m, 8H, $NH_{\varepsilon,Lys}$, $H_{Ar,Cbz}$, $H7_{Indole,}$ $N\underline{H}CH_2$), 7.24 (br s, 1H, $CONH_{2,A}$), 7.14 (d, $J$ = 2.4 Hz, 1H, $H2_{Indole}$), 7.07–7.02 (m, 2H, $H6_{Indole}$, $CONH_{2,B}$), 6.96 (t, $J$ = 7.3 Hz, 1H, $H5_{Indole}$), 5.04 (d, $J$ = 12.6, 1H, $CH_{2,Cbz,A}$), 4.98 (d, $J$ = 12.6, 1H, $CH_{2,Cbz,B}$), 4.54 (td, $J$ = 8.0, 4.8 Hz, 1H, $H_{\alpha,Trp}$), 4.19 (td, $J$ = 8.2, 5.5 Hz, 1H, $H_{\alpha,hArg}$), 3.96–3.91 (m, 1H, $H_{\alpha,Lys}$), 3.29 (br s, 2H, $H_{\varepsilon,Lys}$, overlap with residual water), 3.15 ($m_{ABX}$, $J$ = 14.8, 4.8 Hz, 1H, $H_{\beta,Trp,A}$), 3.08–3.03 (m, 2H, $H_{\varepsilon,hArg}$), 2.99 ($m_{ABX}$, $J$ = 14.9, 8.6 Hz, 1H, $H_{\beta,Trp,B}$), 1.70–1.62 (m, 1H, $H_{\beta,hArg,A}$), 1.58–1.35 (m, 9H, $H_{\beta,hArg,B}$, $H_{\gamma,hArg}$, $H_{\beta,Lys}$, $H_{\delta,Lys}$, $(CH_2)_{11}CH_3$), 1.32–1.19 (m, 24H, $H_{\gamma,Lys}$, $H_{\delta,hArg}$, $(C\underline{H}_2)_{11}CH_3$), 0.85 (t, $J$ = 7.0 Hz, 3H, $CH_3$). $^{13}C$ NMR (151 MHz, DMSO-$d_6$) δ 173.3 ($CO_{hArg}$), 172.0 ($CO_{Lys}$), 171.1 ($CO_{Trp}$), 156.6 ($NHC(=NH)NH_2$), 156.0 ($CO_{Cbz}$), 136.9 ($C1_{Ar,Cbz}$), 136.0 ($C7a_{Indole}$), 128.3 ($C3_{Ar,Cbz}$, $C5_{Ar,Cbz}$), 127.8 ($C4_{Ar,Cbz}$), 127.7 ($C2_{Ar,Cbz}$, $C6_{Ar,Cbz}$), 127.3 ($C3a_{Indole}$), 123.6

($C2_{Indole}$), 120.8 ($C6_{Indole}$), 118.4 ($C4_{Indole}$), 118.2 ($C5_{Indole}$), 111.2 ($C7_{Indole}$), 109.8 ($C3_{Indole}$), 65.5 ($CH_{2,Cbz}$), 54.9 ($C_{\alpha,Lys}$), 53.3 ($C_{\alpha,Trp}$), 52.2 ($C_{\alpha,hArg}$), 43.4 ($C_{\varepsilon,Lys}$), 40.4 ($C_{\varepsilon,hArg}$), 31.6 ($C_{\beta,hArg}$), 31.6 ($C_{\beta,Lys}$), 31.3 ($\underline{C}H_2CH_2CH_3$), 29.0–28.5 (10C, $C_{\delta,hArg}$, ($\underline{C}H_2)_{11}CH_3$), 27.4 ($C_{\beta,Trp}$), 26.4 ($C_{\delta,Lys}$), 23.0 ($C_{\gamma,Lys}$), 22.3 ($C_{\gamma,hArg}$), 22.1 ($\underline{C}H_2CH_3$), 13.9 ($CH_3$). The peak for $C_{\varepsilon,Lys}$ was broad and of low intensity and the peak for C=S was not visible in $^{13}C$ NMR due to fast quadrupolar relaxation via the nearby $^{14}N$-nuclei. Analytical HPLC gradient 0–95% eluent II in eluent I (C18; 20 min total runtime), $t_R$ 14.0 min (>98%, $UV_{230}$). HRMS calcd for $C_{45}H_{71}N_{10}O_5S^+$ $[M+H]^+$, 863.5324; found 863.5313. hArg = homoarginine.

**(4*S*,7*S*,10*S*)-7-((1*H*-indol-3-yl)methyl)-10-(((benzyloxy)carbonyl)amino)-4-carbamoyl-6,9-dioxo-16-thioxo-5,8,15,17-tetraazanonacosanoic acid (11).**

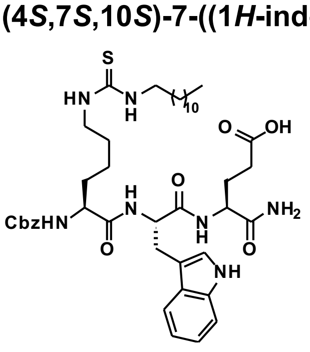

Starting from Cbz-Lys-Trp-Glu(O*t*Bu)-resin (87 mg, estimated loading: 0.51 mmol/g) synthesized from Fmoc-Glu(O*t*Bu)-OH, Fmoc-Trp(Boc)-OH and Cbz-Lys(Fmoc)-OH by SPPS, the title compound was synthesized by on-resin thiourea formation as described in the general procedures. Global deprotection and cleavage from the resin, followed by preparative reversed-phase HPLC purification afforded the desired thiourea **11** (5 mg, 12% based on resin loading) as a colorless fluffy material after lyophilization. $^1H$ NMR (600 MHz, DMSO-$d_6$) δ 12.10 (br s, COOH), 10.80 (br s, 1H, $NH_{Indole}$), 8.00 (d, J = 7.7 Hz, 1H, $NH_{\alpha,Trp}$), 7.90 (d, *J* = 8.0 Hz, 1H, $NH_{Glu}$), 7.56 (d, *J* = 8.0 Hz, 1H, $H4_{Indole}$), 7.41 (d, *J* = 7.7 Hz, 1H, $NH_{\alpha,Lys}$), 7.38–7.20 (m, 8H, $NH_{\varepsilon,Lys}$, $H_{Ar,Cbz}$, $H7_{Indole,}$ $N\underline{H}CH_2$), 7.19–7.14 (m, 2H, $H2_{Indole}$, $CONH_{2,A}$), 7.11–7.02 (m, 2H, $CONH_{2,B}$, $H6_{Indole}$), 6.96 (t, *J* = 7.3 Hz, 1H, $H5_{Indole}$), 5.07–4.94 (m, 2H, $CH_{2,Cbz}$), 4.53 (td, *J* = 8.1, 4.8 Hz, 1H, $H_{\alpha,Trp}$), 4.20 (td, *J* = 8.2, 5.0 Hz, 1H, $H_{\alpha,Glu}$), 3.93 (td, *J* = 8.5, 4.8 Hz, 1H, $H_{\alpha,Lys}$), 3.29 (br s, 2H, $H_{\varepsilon,Lys}$, overlap with residual water), 3.16 ($m_{ABX}$, *J* = 14.9, 4.9 Hz, 1H, $H_{\beta,Trp,A}$), 2.99 ($m_{ABX}$, *J* = 14.9, 8.6 Hz, 1H, $H_{\beta,Trp,B}$), 2.26–2.14 (m, 2H, $H_{\gamma,Glu}$), 1.97–1.85 (m, 1H, $H_{\beta,Glu,A}$), 1.97–1.85 (m, 1H, $H_{\beta,Glu,A}$), 1.70–1.60 (m, 1H, $H_{\beta,Glu,B}$), 1.60–1.04 (m, 28H, $H_{\beta,Lys}$, $H_{\gamma,Lys}$, $H_{\delta,Lys}$, $(C\underline{H}_2)_{11}CH_3$), 0.85 (t, *J* = 6.9 Hz, 3H, $CH_3$). $^{13}C$ NMR (151 MHz, DMSO-$d_6$) δ 174.0 ($COOH_{Glu}$), 172.9 ($CO_{Glu}$), 172.0 ($CO_{Lys}$), 171.2 ($CO_{Trp}$), 156.1 ($CO_{Cbz}$), 136.9 ($C1_{Ar,Cbz}$), 136.0 ($C7a_{Indole}$), 128.3 ($C3_{Ar,Cbz}$, $C5_{Ar,Cbz}$), 127.74 ($C4_{Ar,Cbz}$), 127.69 ($C2_{Ar,Cbz}$, $C6_{Ar,Cbz}$), 127.3 ($C3a_{Indole}$), 123.5 ($C2_{Indole}$), 120.8 ($C6_{Indole}$), 118.4 ($C4_{Indole}$), 118.2 ($C5_{Indole}$), 111.2 ($C7_{Indole}$), 109.9 ($C3_{Indole}$), 65.5 ($CH_{2,Cbz}$), 54.9 ($C_{\alpha,Lys}$), 53.4 ($C_{\alpha,Trp}$), 51.8 ($C_{\alpha,Glu}$), 43.4 ($C_{\varepsilon,Lys}$), 31.5 ($C_{\beta,Lys}$), 31.3 ($\underline{C}H_2CH_2CH_3$), 30.1 ($C_{\gamma,Glu}$), 29.0–28.7 (9C, $(\underline{C}H_2)_{11}CH_3$), 27.4 ($C_{\beta,Trp}$), 27.2 ($C_{\beta,Glu}$), 26.4 ($C_{\delta,Lys}$), 23.0 ($C_{\gamma,Lys}$), 22.1 ($\underline{C}H_2CH_3$), 13.9 ($CH_3$). The peak for $C_{\varepsilon,Lys}$ was broad and of low intensity and the peak for C=S was not visible in $^{13}C$ NMR due to fast quadrupolar relaxation via the nearby $^{14}N$-nuclei. Analytical HPLC gradient 0–95% eluent II in eluent I (C8; 35 min total runtime), $t_R$ 26.7 min (>98%, $UV_{210}$). HRMS calcd for $C_{43}H_{64}N_7O_7S^+$ $[M+H]^+$, 822,4582; found 822.4569.

**Benzyl ((4*S*,7*S*,10*S*)-7-((1*H*-indol-3-yl)methyl)-4-carbamoyl-2-methyl-6,9-dioxo-16-thioxo-5,8,15,17-tetraazanonacosan-10-yl)carbamate (12).**

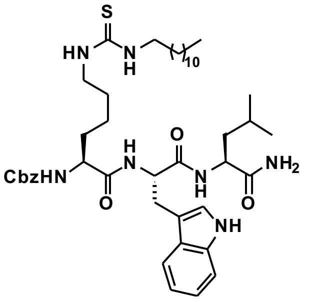

Starting from Cbz-Lys-Trp-Leu-resin (76 mg, estimated loading: 0.51 mmol/g) synthesized from Fmoc-Leu-OH, Fmoc-Trp(Boc)-OH and Cbz-Lys(Fmoc)-OH by SPPS, the title compound was synthesized by on-resin thiourea formation as described in the general procedures. Global deprotection and cleavage from the resin, followed by preparative reversed-phase HPLC purification afforded the desired thiourea **12** (6 mg, 18% based on resin loading) as a colorless fluffy material after lyophilization. $^{1}$H NMR (600 MHz, DMSO-$d_6$) δ 10.84 (d, *J* = 2.4, 1H, $NH_{Indole}$), 8.01 (d, J = 7.9 Hz, 1H, $NH_{\alpha,Trp}$), 7.85 (d, *J* = 8.0 Hz, 1H, $NH_{Leu}$), 7.56 (d, *J* = 7.9 Hz, 1H, $H4_{Indole}$), 7.45–7.19 (m, 9H, $NH_{\alpha,Lys}$, $NH_{\varepsilon,Lys}$, $H_{Ar,Cbz}$, $H7_{Indole,}$ $N\underline{H}CH_2$), 7.18–7.08 (m, 2H, $H2_{Indole}$, $CONH_{2,A}$), 7.08–7.00 (m, 1H, $H6_{Indole}$), 6.99–6.91 (m, 2H, $CONH_{2,B}$, $H5_{Indole}$), 5.08–4.94 (m, 2H, $CH_{2,Cbz}$), 4.55 (td, *J* = 8.1, 5.3 Hz, 1H, $H_{\alpha,Trp}$), 4.22 (q, *J* = 7.8, 1H, $H_{\alpha,Leu}$), 3.95 (td, *J* = 8.4, 4.9 Hz, 1H, $H_{\alpha,Lys}$), 3.29 (br s, 2H, $H_{\varepsilon,Lys}$, overlap with residual water), 3.14 ($m_{ABX}$, *J* = 14.8, 5.3 Hz, 1H, $H_{\beta,Trp,A}$), 2.97 ($m_{ABX}$, *J* = 14.8, 8.3 Hz, 1H, $H_{\beta,Trp,B}$), 1.59–1.50 (m, 2H, $H_{\beta,Lys}$, $H_{\gamma,Leu}$), 1.49–1.10 (m, 29H, $H_{\beta,Lys,B}$, $H_{\gamma,Lys}$, $H_{\delta,Lys}$, $H_{\beta,Leu}$, $(C\underline{H}_2)_{11}CH_3$), 0.88–0.83 (m, 6H, $H_{\delta,Leu,1}$, $CH_2\underline{CH_3}$), 0.81 (d, *J* = 6.5 Hz, 3H, $H_{\delta,Leu,2}$). $^{13}$C NMR (151 MHz, DMSO-$d_6$) δ 173.9 ($CO_{Leu}$),zz 172.0 ($CO_{Lys}$), 171.0 ($CO_{Trp}$), 156.0 ($CO_{Cbz}$), 136.9 ($C1_{Ar,Cbz}$), 136.0 ($C7a_{Indole}$), 128.3 ($C3_{Ar,Cbz}$, $C5_{Ar,Cbz}$), 127.75 ($C4_{Ar,Cbz}$), 127.66 ($C2_{Ar,Cbz}$, $C6_{Ar,Cbz}$), 127.3 ($C3a_{Indole}$), 123.5 ($C2_{Indole}$), 120.8 ($C6_{Indole}$), 118.4 ($C4_{Indole}$), 118.2 ($C5_{Indole}$), 111.2 ($C7_{Indole}$), 109.9 ($C3_{Indole}$), 65.4 ($CH_{2,Cbz}$), 54.8 ($C_{\alpha,Lys}$), 53.3 ($C_{\alpha,Trp}$), 50.9 ($C_{\alpha,Glu}$), 43.4 ($C_{\varepsilon,Lys}$), 41.0 ($C_{\beta,Leu}$), 31.6 ($C_{\beta,Lys}$), 31.3 ($\underline{C}H_2CH_2CH_3$), 29.0–28.7 (9C, $(\underline{C}H_2)_{11}CH_3$), 27.2 ($C_{\beta,Trp}$), 26.4 ($C_{\delta,Lys}$), 24.1 ($C_{\gamma,Leu}$), 23.0 ($C_{\gamma,Lys}$), 22.9 ($C_{\delta,Leu,1}$), 22.1 ($\underline{C}H_2CH_3$), 21.6 ($C_{\delta,Leu,2}$), 13.9 ($CH_2\underline{C}H_3$). The peak for $C_{\varepsilon,Lys}$ was broad and of low intensity and the peak for C=S was not visible in $^{13}$C NMR due to fast quadrupolar relaxation via the nearby $^{14}$N-nuclei. Analytical HPLC gradient 0–95% eluent II in eluent I (C18; 35 min total runtime), $t_R$ 27.9 min (>98%, $UV_{230}$). HRMS calcd for $C_{44}H_{68}N_7O_5S^+$ $[M+H]^+$, 806.4997; found 806.4987.

**Benzyl ((6*S*,9*S*,12*S*)-1-amino-6-carbamoyl-9-(3-guanidinopropyl)-1-imino-8,11-dioxo-18-thioxo-2,7,10,17,19-pentaazahentriacontan-12-yl)carbamate (13).**

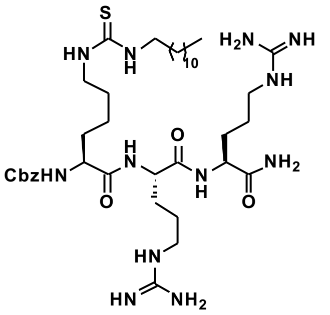

Starting from Cbz-Lys-Arg(Pbf)-Arg(Pbf)-resin (87 mg, estimated loading: 0.46 mmol/g) synthesized from Fmoc-Arg(Pbf)-OH and Cbz-Lys(Fmoc)-OH by SPPS, the title compound was synthesized by on-resin thiourea formation as described in the general procedures. Global deprotection and cleavage from the resin, followed by preparative reversed-phase HPLC purification afforded the desired thiourea **13** (2 mg, 6% based on resin loading) as a colorless fluffy material after lyophilization. $^{1}$H NMR (600 MHz, DMSO-$d_6$) δ 8.01 (d, *J* = 7.7 Hz, 1H, $NH_{Arg,1}$), 7.92 (d, *J* = 7.9 Hz, 1H, $NH_{Arg\ 2}$), 7.56–7.48 (m, 2H, $NH_{\delta,Arg\ 1}$, $NH_{\delta,Arg\ 2}$), 7.43 (d, *J* = 7.9 Hz, 1H,

$NH_{\alpha,Lys}$), 7.42–7.26 (m, 8H, $NH_{\varepsilon,Lys}$, $N\underline{H}CH_2$, $H_{Ar,Cbz}$, $CONH_{2,A}$), 7.09 (s, 1H, $CONH_{2,B}$), 5.09–4.95 (m, 2H, $CH_{2,Cbz}$), 4.28 (td, $J$ = 7.9, 5.5 Hz, 1H, $H_{\alpha,Arg\ 1}$), 4.20 (td, $J$ = 7.8, 5.7 Hz, 1H, $H_{\alpha,Arg\ 2}$), 3.99 (td, $J$ = 8.8, 4.5 Hz, 1H, $H_{\alpha,Lys}$), 3.30 (br s, 2H, $H_{\varepsilon,Lys}$, overlap with residual water), 3.14–3.03 (m, 4H, $H_{\delta,Arg\ 1}$, $H_{\delta,Arg\ 2}$), 1.74–1.57 (m, 3H, $H_{\beta,Arg\ 1,A}$, $H_{\beta,Arg\ 2,A}$, $H_{\beta,Lys,A}$), 1.57–1.38 (m, 11H, $H_{\beta,Arg\ 1,B}$, $H_{\beta,Arg\ 2,B}$, $H_{\beta,Lys,B}$, $H_{\gamma,Arg\ 1}$, $H_{\gamma,Arg\ 2}$, $(C\underline{H}_2)_{11}CH_3$), 1.37–1.19 (m, 22H, $H_{\gamma,Lys}$, $H_{\delta,Lys}$, $(C\underline{H}_2)_{11}CH_3$), 0.85 (t, $J$ = 6.9 Hz, 3H, $CH_3$). $^{13}C$ NMR (151 MHz, DMSO-$d_6$) δ 173.1 ($CO_{Arg\ 2}$), 172.1 ($CO_{Arg\ 1}$), 171.1 ($CO_{Lys}$), 156.6 ($NHC(=NH)NH_{2,Arg\ 1}$, $NHC(=NH)NH_{2,Arg\ 2}$), 156.0 ($CO_{Cbz}$), 136.9 ($C1_{Ar,Cbz}$), 128.4 ($C3_{Ar,Cbz}$, $C5_{Ar,Cbz}$), 127.8 ($C4_{Ar,Cbz}$), 127.7 ($C2_{Ar,Cbz}$, $C6_{Ar,Cbz}$), 65.4 ($CH_{2,Cbz}$), 54.7 ($C_{\alpha,Lys}$), 52.1 ($C_{\alpha,Arg\ 1}$), 51.9 ($C_{\alpha,Arg\ 2}$), 43.4 ($C_{\varepsilon,Lys}$), 40.5 ($C_{\delta,Arg\ 1}$), 40.4 ($C_{\delta,Arg\ 2}$), 31.6 ($C_{\beta,Lys}$), 31.3 ($\underline{C}H_2CH_2CH_3$), 29.2–28.7 (11C, $C_{\beta,Arg,1}$, $C_{\beta,Arg,2}$, $(\underline{C}H_2)_{11}CH_3$), 26.4 ($C_{\delta,Lys}$), 25.0 ($C_{\gamma,Arg\ 2}$), 24.9 ($C_{\gamma,Arg\ 1}$), 23.1 ($C_{\gamma,Lys}$), 22.1 ($\underline{C}H_2CH_3$), 13.9 ($CH_3$). The peak for $C_{\varepsilon,Lys}$ was broad and of low intensity and the peak for C=S was not visible in $^{13}C$ NMR due to fast quadrupolar relaxation via the nearby $^{14}N$-nuclei. Analytical HPLC gradient 0–95% eluent II in eluent I (C8; 35 min total runtime), $t_R$ 24.7 min (>98%, $UV_{230}$). HRMS calcd for $C_{39}H_{71}N_{12}O_5S^+$ $[M+H]^+$, 819.5386; found 819.5374.

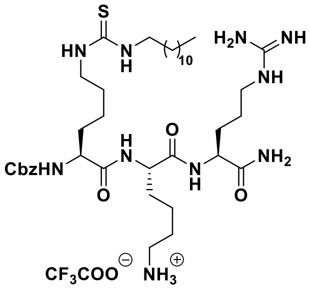

**Benzyl ((6*S*,9*S*,12*S*)-1-amino-9-(4-aminobutyl)-6-carbamoyl-1-imino-8,11-dioxo-18-thioxo-2,7,10,17,19-pentaazahentriacontan-12-yl)carbamate (14).** Starting from Cbz-Lys-Lys(Boc)-Arg(Pbf)-resin (87 mg, estimated loading: 0.46 mmol/g) synthesized from Fmoc-Arg(Pbf)-OH, Fmoc-Lys(Boc)-OH and Cbz-Lys(Fmoc)-OH by SPPS, the title compound was synthesized by on-resin thiourea formation as described in the general procedures. Global deprotection and cleavage from the resin, followed by preparative reversed-phase HPLC purification afforded the desired thiourea **14** (2 mg, 6% based on resin loading) as a colorless fluffy material after lyophilization. $^1H$ NMR (600 MHz, DMSO-$d_6$) δ 7.98 (d, $J$ = 7.8 Hz, 1H, $NH_{\alpha,Lys}$), 7.89 (d, $J$ = 7.8 Hz, 1H, $NH_{\alpha,Arg}$), 7.62–7.35 (m, 3H, $NH_3{}^{+}{}_{,\varepsilon,Lys}$), 7.55 (t, $J$ = 5.8 Hz, 1H, $NH_{\delta,Arg}$), 7.42 (d, $J$ = 7.9 Hz, 1H, $NH_{\alpha,Lys(Dtu)}$), 7.40–7.29 (m, 8H, $NH_{\varepsilon,Lys(Dtu)}$, $N\underline{H}CH_2$, $H_{Ar,Cbz}$, $CONH_{2,A}$), 7.09 (s, 1H, $CONH_{2,B}$), 5.06–4.98 (m, 2H, $CH_{2,Cbz}$), 4.24 (td, $J$ = 8.4, 5.0 Hz, 1H, $H_{Lys}$), 4.19 (td, $J$ = 7.9, 5.8 Hz, $1H_{\alpha,Arg}$), 4.03–3.93 (m, 1H, $H_{\alpha,Lys(Dtu)}$), 3.30 (br s, 2H, $H_{\varepsilon,Lys(Dtu)}$, overlap with residual water), 3.09 (q, $J$ = 6.7 Hz, 2H, $H_{\delta,Arg}$), 2.80–2.69 (m, 2H, $H_{\delta,Lys}$), 1.71–1.58 (m, 3H, $H_{\beta,Arg,A}$, $H_{\beta,Lys,A}$, $H_{\beta,Lys(Dtu),A}$), 1.57–1.39 (m, 11H, $H_{\beta,Arg,B}$, $H_{\gamma,Arg}$, $H_{\beta,Lys,B}$, $H_{\beta,Lys(Dtu),B}$, $H_{\delta,Lys(Dtu)}$, $(C\underline{H}_2)_{11}CH_3$), 1.37–1.19 (m, 24H, $H_{\gamma,Lys}$, $H_{\delta,Lys}$, $H_{\gamma,Lys(Dtu)}$, $(C\underline{H}_2)_{11}CH_3$), 0.85 (t, $J$ = 7.0 Hz, 3H, $CH_3$). $^{13}C$ NMR (151 MHz, DMSO-$d_6$) δ 173.2 ($CO_{Arg}$), 172.1 ($CO_{Lys}$), 171.3 ($CO_{Lys(Dtu)}$), 156.7 ($NHC(=NH)NH_2$), 156.0 ($CO_{Cbz}$), 136.9 ($C1_{Ar,Cbz}$), 128.3 ($C3_{Ar,Cbz}$, $C5_{Ar,Cbz}$), 127.8 ($C4_{Ar,Cbz}$), 127.6 ($C2_{Ar,Cbz}$, $C6_{Ar,Cbz}$), 65.4 ($CH_{2,Cbz}$), 54.7 ($C_{\alpha,Lys(Dtu)}$), 52.2 ($C_{\alpha,Lys}$), 51.8 ($C_{\alpha,Arg}$), 43.4 ($C_{\varepsilon,Lys(Dtu)}$), 40.4 ($C_{\delta,Arg}$), 31.58 ($C_{\beta,Lys}$), 31.56 ($C_{\beta,Lys(Dtu)}$), 31.3 ($\underline{C}H_2CH_2CH_3$), 29.2–28.7 (10C, $C_{\beta,Arg}$, $(\underline{C}H_2)_{11}CH_3$), 26.6 ($C_{\delta,Lys}$), 26.4 ($C_{\delta,Lys(Dtu)}$), 25.0 ($C_{\gamma,Arg}$), 23.0 ($C_{\gamma,Lys(Dtu)}$), 22.10 ($C_{\gamma,Lys}$), 22.06 ($\underline{C}H_2CH_3$), 13.9 ($CH_3$). The peak for $C_{\varepsilon,Lys(Dtu)}$ was broad and of low intensity and the peak for C=S was not visible in $^{13}C$ NMR due to fast quadrupolar relaxation via the nearby $^{14}$N-nuclei. Analytical HPLC gradient 0–95% eluent II in eluent I (C18; 35 min total runtime), $t_R$ 25.7 min (>95%, $UV_{254}$). HRMS calcd for $C_{39}H_{71}N_{10}O_5S^+$ $[M+H]^+$, 791.5324; found 791.5313. Dtu = 1-dodecylthiourea.

**(*S*)-5-(((*S*)-1-amino-5-guanidino-1-oxopentan-2-yl)amino)-4-((*S*)-2-(((benzyloxy)carbonyl)-amino)-6-(3-dodecylthioureido)hexanamido)-5-oxopentanoic acid (15).**

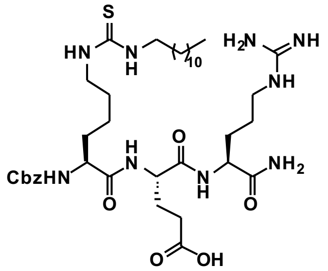

Starting from Cbz-Lys-Glu(O*t*Bu)-Arg(Pbf)-resin (87 mg, estimated loading: 0.46 mmol/g) synthesized from Fmoc-Arg(Pbf)-OH, Fmoc-Glu(O*t*Bu)-OH and Cbz-Lys(Fmoc)-OH by SPPS, the title compound was synthesized by on-resin thiourea formation as described in the general procedures. Global deprotection and cleavage from the resin, followed by preparative reversed-phase HPLC purification afforded the desired thiourea **15** (3 mg, 9% based on resin loading), as a colorless fluffy material after lyophilization. $^1$H NMR (600 MHz, DMSO-$d_6$) δ 12.11 (s, 1H, $COOH_{Glu}$), 8.02 (d, $J$ = 7.6 Hz, 1H. $NH_{Glu}$), 7.89 (d, $J$ = 7.9 Hz, 1H, $NH_{\alpha,Arg}$), 7.53 (t, $J$ = 5.8 Hz, 1H, $NH_{\delta,Arg}$), 7.43 (d, $J$ = 7.9 Hz, 1H, $NH_{\alpha,Lys}$), 7.40–7.26 (m, 8H, $NH_{\varepsilon,Lys}$, $N\underline{H}CH_2$, $H_{Ar,Cbz}$, $CONH_{2,A}$), 7.08 (s, 1H, $CONH_{2,B}$), 5.08–4.97 (m, 2H, $CH_{2,Cbz}$), 4.25 (td, $J$ = 8.2, 5.3 Hz, 1H, $H_{\alpha,Glu}$), 4.18 (td, $J$ = 7.9, 5.6 Hz, 1H, $H_{\alpha,Arg}$), 3.98 (td, $J$ = 8.9, 4.7 Hz, 1H, $H_{\alpha,Lys}$), 3.31 (br s, 2H, $H_{\varepsilon,Lys}$, overlap with residual water), 3.12–3.06 (q, $J$ = 6.7 Hz, 2H, $H_{\delta,Arg}$), 2.32–2.18 (m, 2H, $H_{\gamma,Glu}$), 1.96–1.87 (m, 1H, $H_{\beta,Glu,A}$), 1.80–1.72 (m, 1H, $H_{\beta,Glu,B}$), 1.71–1.58 (m, 2H, $H_{\beta,Arg,A}$, $H_{\beta,Lys,A}$), 1.56–1.37 (m, 8H, $H_{\beta,Arg,B}$, $H_{\beta,Lys,B}$, $H_{\gamma,Arg}$, $_2(C\underline{H}_2)_{11}CH_3$), 1.36–1.10 (m, 22H, $H_{\gamma,Lys}$, $H_{\delta,Lys}$, $(C\underline{H}_2)_{10}CH_3$), 0.85 (t, $J$ = 6.9 Hz, 3H, $CH_3$). $^{13}$C NMR (151 MHz, DMSO-$d_6$) δ 174.0 ($CO_{\delta,Glu}$), 173.1 ($CO_{Arg}$), 172.1 ($CO_{Lys}$), 170.9 ($CO_{\alpha,Glu}$), 156.6 ($NHC(=NH)NH_2$), 156.0 ($CO_{Cbz}$), 136.9 ($C1_{Ar,Cbz}$), 128.3 ($C3_{Ar,Cbz}$, $C5_{Ar,Cbz}$), 127.8 ($C4_{Ar,Cbz}$), 127.7 ($C2_{Ar,Cbz}$, $C6_{Ar,Cbz}$), 65.4 ($CH_{2,Cbz}$), 54.7 ($C_{\alpha,Lys}$), 52.0 ($C_{\alpha,Glu}$), 51.9 ($C_{\alpha,Arg}$), 43.4 ($C_{\varepsilon,Lys}$), 40.4 ($C_{\delta,Arg}$), 31.5 ($C_{\beta,Lys}$), 31.3 ($\underline{C}H_2CH_2CH_3$), 30.1 ($C_{\gamma,Glu}$), 29.1–28.7 (10C, $C_{\beta,Arg}$, $(\underline{C}H_2)_{11}CH_3$), 27.3 ($C_{\beta,Glu}$), 26.4 ($C_{\delta,Lys}$), 25.0 ($C_{\gamma,Arg}$), 23.0 ($C_{\gamma,Lys}$), 22.1 ($\underline{C}H_2CH_3$), 14.0 ($CH_3$). The peak for $C_{\varepsilon,Lys}$ was broad and of low intensity and the peak for C=S was not visible in $^{13}$C NMR due to fast quadrupolar relaxation via the nearby $^{14}$N-nuclei. Analytical HPLC gradient 0–95% eluent II in eluent I (C18; 20 min total runtime), $t_R$ 13.0 min (>96%, $UV_{230}$). HRMS calcd for $C_{38}H_{66}N_9O_7S^+$ $[M+H]^+$, 792.4800; found 792.4791.

**Benzyl ((6*S*,9*S*,12*S*)-1-amino-6-carbamoyl-1-imino-9-isobutyl-8,11-dioxo-18-thioxo-2,7,10,17,19-pentaazahentriacontan-12-yl)carbamate (16).**

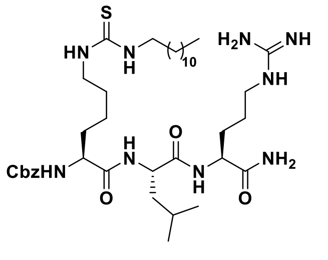

Starting from Cbz-Lys-Leu-Arg(Pbf)-resin (87 mg, estimated loading: 0.46 mmol/g) synthesized from Fmoc-Arg(Pbf)-OH, Fmoc-Leu-OH and Cbz-Lys(Fmoc)-OH by SPPS, the title compound was synthesized by the general on-resin thiourea formation as described in the general procedures. Global deprotection and cleavage from the resin, followed by preparative reversed-phase HPLC purification afforded the desired thiourea **16** (1 mg, 3% based on resin loading) as a colorless fluffy material after lyophilization. $^{1}$H NMR (600 MHz, DMSO-$d_6$) δ 7.96 (d, $J$ = 7.9 Hz, 1H, $NH_{Leu}$), 7.83 (d, $J$ = 8.0 Hz, 1H, $NH_{\alpha,Arg}$), 7.49 (t, $J$ = 6.0 Hz, 1H, $NH_{\delta,Arg}$), 7.41 (d, $J$ = 8.0 Hz, 1H, $NH_{\alpha,Lys}$), 7.39–7.25 (m, 8H, $NH_{\varepsilon,Lys}$, N<u>H</u>CH, $H_{Ar,Cbz}$, $CONH_{2,A}$), 7.08 (s, 1H, $CONH_{2,B}$), 5.07–4.99 (m, 2H, $CH_{2,Cbz}$), 4.31–4.23 (m, 1H, $H_{\alpha,Leu}$), 4.18 (td, $J$ = 8.0, 5.6 Hz, 1H, $H_{\alpha,Arg}$), 3.98 (td, $J$ = 8.6, 4.9 Hz, 1H, $H_{\alpha,Lys}$), 3.30 (br s, 2H, $H_{\varepsilon,Lys}$, overlap with residual water), 3.12–3.04 (m, 2H, $H_{\delta,Arg}$), 1.71–1.58 (m, 3H, $H_{\beta,Arg,A}$, $H_{\gamma,Leu}$, $H_{\beta,Lys,A}$), 1.55–1.37 (m, 10H, $H_{\beta,Arg,B}$, $H_{\beta,Lys,B}$, $H_{\gamma,Arg}$, $H_{\beta,Leu}$, $(C\underline{H}_2)_{11}CH_3$), 1.35–1.19 (m, 22H, $H_{\gamma,Lys}$, $H_{\delta,Lys}$, $(C\underline{H}_2)_{11}CH_3$), 0.91–0.79 (m, 9H, $H_{\delta,Leu}$, $CH_2C\underline{H}_3$). $^{13}$C NMR (151 MHz, DMSO-$d_6$) δ 173.1 ($CO_{Arg}$), 172.0 ($CO_{Leu}$), 171.8 ($CO_{Lys}$), 156.6 (NHC(=NH)$NH_2$), 156.0 ($CO_{Cbz}$), 137.0 ($C1_{Ar,Cbz}$), 128.3 ($C3_{Ar,Cbz}$, $C5_{Ar,Cbz}$), 127.8 ($C4_{Ar,Cbz}$), 127.7 ($C2_{Ar,Cbz}$, $C6_{Ar,Cbz}$), 65.4 ($CH_{2,Cbz}$), 54.7 ($C_{\alpha,Lys}$), 51.8 ($C_{\alpha,Arg}$), 51.2 ($C_{\alpha,Leu}$), 43.4 ($C_{\varepsilon,Lys}$), 40.5 ($C_{\beta,Leu}$), 40.4 ($C_{\delta,Arg}$), 31.6 ($C_{\beta,Lys}$), 31.3 ($\underline{C}H_2CH_2CH_3$), 29.2–28.7 (10C, $C_{\beta,Arg}$, $(\underline{C}H_2)_{11}CH_3$), 26.4 ($C_{\delta,Lys}$), 25.0 ($C_{\gamma,Arg}$), 24.1 ($C_{\gamma,Leu}$), 23.1 ($C_{\delta,Leu,1}$), 23.0 ($C_{\gamma,Lys}$), 22.1 ($\underline{C}H_2CH_3$), 21.5 ($C_{\delta,Leu,2}$), 13.9 ($CH_2\underline{C}H_3$). The peak for $C_{\varepsilon,Lys}$ was broad and of low intensity and the peak for C=S was not visible in $^{13}$C NMR due to fast quadrupolar relaxation via the nearby $^{14}$N-nuclei. Analytical HPLC gradient 0–95% eluent II in eluent I (C18; 20 min total runtime), $t_R$ 13.9 min (>98%, $UV_{230}$). HRMS calcd for $C_{39}H_{70}N_9O_5S^+$ [M+H]$^+$, 776.5215; found 776.5211.

**Benzyl ((6*S*,9*S*,12*S*)-1-amino-6-carbamoyl-9-(4-hydroxybenzyl)-1-imino-8,11-dioxo-18-thioxo-2,7,10,17,19-pentaazahentriacontan-12-yl)carbamate (17).**

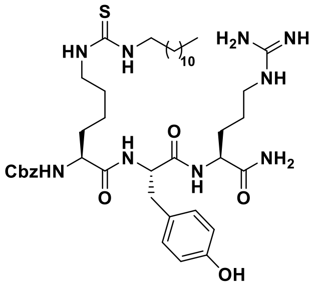

Starting from Cbz-Lys-Tyr(*t*Bu)-Arg(Pbf)-resin (87 mg, estimated loading: 0.46 mmol/g) synthesized from Fmoc-Arg(Pbf)-OH, Fmoc-Tyr(*t*Bu)-OH and Cbz-Lys(Fmoc)-OH by SPPS, the title compound was synthesized by on-resin thiourea formation as described in the general procedures. Global deprotection and cleavage from the resin, followed by preparative reversed-phase HPLC purification afforded the desired thiourea **17** (3 mg, 9% based on resin loading) as a colorless fluffy material after lyophilization. $^{1}$H NMR (600 MHz, DMSO-$d_6$) δ 9.15 (s, 1H, $OH_{Tyr}$), 7.99 (d, $J$ = 8.1Hz, 1H, $NH_{Arg}$), 7.86 (d, $J$ = 7.9 Hz, 1H, $NH_{Tyr}$), 7.51 (t, $J$ = 5.8 Hz, 1H, $NH_{\delta,Arg}$), 7.41–7.27 (m, 8H, $NH_{\alpha,Lys}$, $NH_{\varepsilon,Lys}$, N<u>H</u>CH$_2$, $H_{Ar,Cbz}$), 7.24 (br s, 1H, $CONH_{2,A}$), 7.10 (br s, 1H, $CONH_{2,B}$), 7.00 (d, $J$ = 8.1 Hz, 2H, $H2_{Ar,Tyr}$, $H6_{Ar,Tyr}$), 6.62 (d, $J$ = 8.3 Hz, 2H, $H3_{Ar,Tyr}$, $H5_{Ar,Tyr}$), 5.08–4.96 (m, 2H, $CH_{2,Cbz}$), 4.43 (td, $J$ = 8.3, 4.6 Hz, 1H, $H_{\alpha,Tyr}$), 4.20 (td, $J$ = 8.0, 5.6 Hz, 1H, $H_{\alpha,Arg}$), 3.96–3.88 (m, 1H, $H_{\alpha,Lys}$), 3.30 (br s, 2H, $H_{\varepsilon,Lys}$, overlap with residual water), 3.09 (q, $J$ = 6.7 Hz, 2H, $H_{\delta,Arg}$), 2.91 (dd, $J$ = 14.1, 4.6 Hz, 1H, $H_{\beta,Tyr,A}$), 2.71 (dd, $J$ = 14.0, 9.0 Hz, 1H, $H_{\beta,Tyr,B}$), 1.73–1.64 (m, 1H, $H_{\beta,Arg,A}$), 1.58–1.48 (m, 2H, $H_{\beta,Arg,B}$, $H_{\beta,Lys,A}$), 1.47–1.35 (m, 7H, $H_{\beta,Lys,B}$, $H_{\gamma,Arg}$, $(CH_2)_{11}CH_3$), 1.29–1.10 (m, 22H, $H_{\gamma,Lys}$, $H_{\delta,Lys}$, $(C\underline{H}_2)_{11}CH_3$), 0.85 (t, $J$ = 6.9 Hz, 3H, $CH_3$).$^{13}$C NMR (151 MHz, DMSO-$d_6$) δ 173.1 ($CO_{Arg}$), 171.8 ($CO_{Lys}$), 170.9 ($CO_{Tyr}$), 156.7 (NHC(=NH)$NH_2$), 155.9 ($CO_{Cbz}$), 155.8 ($C4_{Ar,Tyr}$), 136.9 ($C1_{Ar,Cbz}$), 130.1 ($C2_{Ar,Tyr}$, $C6_{Ar,Tyr}$), 128.4 ($C3_{Ar,Cbz}$, $C5_{Ar,Cbz}$), 127.8 ($C4_{Ar,Cbz}$), 127.7 ($C2_{Ar,Cbz}$, $C6_{Ar,Cbz}$), 127.5 ($C1_{Ar,Tyr}$), 114.8 ($C3_{Ar,Tyr}$, $C5_{Ar,Tyr}$), 65.5 ($CH_{2,Cbz}$), 54.8 ($C_{\alpha,Lys}$), 54.1 ($C_{\alpha,Tyr}$), 51.9

($C_{\alpha,Arg}$), 43.4 ($C_{\varepsilon,Lys}$), 40.4 ($C_{\delta,Arg}$), 36.5 ($C_{\beta,Tyr}$), 31.6 ($C_{\beta,Lys}$), 31.3 ($\underline{C}H_2CH_2CH_3$), 29.2–28.7 (10C, $C_{\beta,Arg}$, $(\underline{C}H_2)_{11}CH_3$), 26.4 ($C_{\delta,Lys}$), 25.0 ($C_{\gamma,Arg}$), 23.0 ($C_{\gamma,Lys}$), 22.1 ($\underline{C}H_2CH_3$), 13.9 ($CH_3$). The peak for $C_{\varepsilon,Lys}$ was broad and of low intensity and the peak for C=S was not visible in $^{13}$C NMR due to fast quadrupolar relaxation via the nearby $^{14}$N-nuclei. Analytical HPLC gradient 0–95% eluent II in eluent I (C18; 20 min total runtime), $t_R$ 26.6 min (>98%, $UV_{230}$). HRMS calcd for $C_{42}H_{68}N_9O_5S^+$ $[M+H]^+$, 826.5008; found 826.5003.

**Benzyl ((6*S*,9*S*,12*S*)-1-amino-6-carbamoyl-9-(4-hydroxy-3-nitrobenzyl)-1-imino-8,11-dioxo-18-thioxo-2,7,10,17,19-pentaazahentriacontan-12-yl)carbamate (18).**

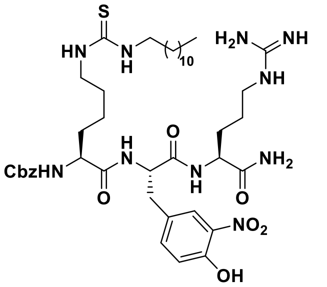

Starting from Cbz-Lys-Tyr(3-$NO_2$)-Arg(Pbf)-resin (87 mg, estimated loading: 0.46 mmol/g) synthesized from Fmoc-Arg(Pbf)-OH, Fmoc-Tyr(3-$NO_2$)-OH and Cbz-Lys(Fmoc)-OH by SPPS, the title compound was synthesized by on-resin thiourea formation as described in the general procedures. Global deprotection and cleavage from the resin, followed by preparative reversed-phase HPLC purification afforded the desired thiourea **18** (4 mg, 11% based on resin loading), as a yellow fluffy material after lyophilization. $^1$H NMR (600 MHz, DMSO-$d_6$) δ 9.15 (s, 1H, $Ar\underline{OH}_{NTyr}$), 8.12 (d, $J$ = 8.0 Hz, 1H, $NH_{\alpha,Arg}$), 7.94 (d, $J$ = 8.2 Hz, 1H, $NH_{NTyr}$), 7.78 (d, $J$ = 2.2 Hz, 1H, $H2_{Ar,NTyr}$), 7.51 (t, $J$ = 5.8 Hz, 1H, $NH_{\delta,Arg}$), 7.42 (dd, $J$ = 8.5, 2.2 Hz, 1H, $H6_{Ar,NTyr}$), 7.39–7.22 (m, 9H, $NH_{\alpha,Lys}$, $NH_{\varepsilon,Lys}$, $N\underline{H}CH_2$, $H_{Ar,Cbz}$, $CONH_{2,A}$), 7.10 (br s, 1H, $CONH_{2,B}$), 7.01 (d, $J$ = 8.5 Hz, 1H, $H5_{Ar,NTyr}$), 5.05–4.94 (m, 2H, $CH_{2,Cbz}$), 4.53 (td, $J$ = 8.8, 4.2 Hz, 1H, $H_{\alpha,NTyr}$), 4.22 (td, $J$ = 7.9, 5.6 Hz, 1H, $H_{\alpha,Arg}$), 3.88 (td, $J$ = 8.8, 4.9 Hz, 1H, $H_{\alpha,Lys}$), 3.28 (br s, 2H, $H_{\varepsilon,Lys}$, overlap with residual water), 3.10 (q, $J$ = 6.6 Hz, 2H, $H_{\delta,Arg}$), 3.02 (dd, $J$ = 14.0, 4.2 Hz, 1H, $H_{\beta,NTyr,A}$), 2.77 (dd, $J$ = 14.0, 9.6 Hz, 1H, $H_{\beta,NTyr,B}$), 1.73–1.65 (m, 1H, $H_{\beta,Arg,A}$), 1.59–1.33 (m, 9H, $H_{\beta,Arg,B}$, $H_{\beta,Lys}$, $H_{\gamma,Arg}$, $(C\underline{H}_2)_{11}CH_3$), 1.31–1.10 (m, 22H, $H_{\gamma,Lys}$, $H_{\delta,Lys}$, $(C\underline{H}_2)_{11}CH_3$), 0.85 (t, $J$ = 7.0 Hz, 3H, $CH_3$). $^{13}$C NMR (151 MHz, DMSO-$d_6$) δ 173.0 ($CO_{Arg}$), 171.9 ($CO_{Lys}$), 170.5 ($CO_{NTyr}$),156.6 ($NHC(=NH)NH_2$), 155.8 ($CO_{Cbz}$), 150.9 ($C4_{Ar,NTyr}$), 136.9 ($C1_{Ar,Cbz}$), 136.5 ($C6_{Ar,NTyr}$), 136.0 ($C3_{Ar,NTyr}$), 128.8 ($C1_{Ar,NTyr}$), 128.4 ($C3_{Ar,Cbz,}$ $C5_{Ar,Cbz}$), 127.8 ($C4_{Ar,Cbz}$), 127.7 ($C2_{Ar,Cbz,}$ $C6_{Ar,Cbz}$), 125.5 ($C2_{Ar,NTyr}$), 118.7 ($C5_{Ar,NTyr}$), 65.4 ($CH_{2,Cbz}$), 54.9 ($C_{\alpha,Lys}$), 53.4 ($C_{\alpha,NTyr}$), 52.0 ($C_{\alpha,Arg}$), 43.4 ($C_{\varepsilon,Lys}$), 40.4 ($C_{\delta,Arg}$), 36.2 ($C_{\beta,NTyr}$), 31.7 ($C_{\beta,Lys}$), 31.3 ($\underline{C}H_2CH_2CH_3$), 29.2–28-7 (10C, $C_{\beta,Arg}$, $(\underline{C}H_2)_{11}CH_3$), 26.4 ($C_{\delta,Lys}$), 25.0 ($C_{\gamma,Arg}$), 23.0 ($C_{\gamma,Lys}$), 22.1 ($\underline{C}H_2CH_3$), 13.9 ($CH_3$). The peak for $C_{\varepsilon,Lys}$ was broad and of low intensity and the peak for C=S was not visible in $^{13}$C NMR due to fast quadrupolar relaxation via the nearby $^{14}$N-nuclei. Analytical HPLC gradient 0–95% eluent II in eluent I (C18; 20 min total runtime), $t_R$ 13.9 min (>97%, $UV_{230}$). HRMS calcd for $C_{42}H_{67}N_{10}O_8S^+$ $[M+H]^+$, 871.4859; found 871.4849.

**Benzyl ((*S*)-1-((*S*)-2-(((*S*)-1-amino-5-guanidino-1-oxopentan-2-yl)carbamoyl)pyrrolidin-1-yl)-6-(3-dodecylthioureido)-1-oxohexan-2-yl)carbamate (19).** 

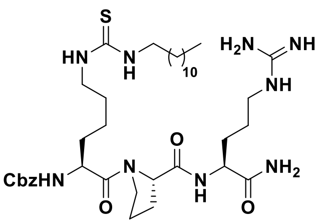

Starting from Cbz-Lys-Pro-Arg(Pbf)-resin (87 mg, estimated loading: 0.46 mmol/g) synthesized from Fmoc-Arg(Pbf)-OH, Fmoc-Pro-OH and Cbz-Lys(Fmoc)-OH by SPPS, the title compound was synthesized by on-resin thiourea formation as described in the general procedures. Global deprotection and cleavage from the resin, followed by preparative reversed-phase HPLC purification afforded the desired thiourea **19** (3 mg, 10% based on resin loading) as a colorless fluffy material after lyophilization. $^1$H NMR (600 MHz, DMSO-$d_6$) δ 7.88 (d, *J* = 8.0 Hz, 1H, $NH_{\alpha,Arg}$), 7.52 (t, *J* = 5.9 Hz, 1H, $NH_{\delta,Arg}$), 7.47 (d, *J* = 7.8 Hz, 1H, $NH_{\alpha,Lys}$), 7.41–7.27 (m, 7H, $NH_{\varepsilon,Lys}$, $N\underline{H}CH_2$, $H_{Ar,Cbz}$), 7.25 (br s, 1H, $CONH_{2,A}$), 7.06 (br s, 1H, $CONH_{2,B}$), 5.06–4.98 (m, 2H, $CH_{2,Cbz}$), 4.34 (dd, *J* = 8.5, 4.4 Hz, 1H, $H_{\alpha,Pro}$), 4.26–4.20 (m, 1H, $H_{\alpha,Lys}$), 4.16 (td, *J* = 8.1, 5.3 Hz, 1H, $H_{\alpha,Arg}$), 3.72–3.63 (m, 1H, $H_{\delta,Pro,A}$), 3.60–3.53 (m, 1H, $H_{\delta,Pro,B}$), 3.31 (br s, 2H, $H_{\varepsilon,Lys}$, overlap with residual water), 3.09 (q, *J* = 6.8 Hz, 2H, $H_{\delta,Arg}$), 2.10–2.03 (m, 1H, $H_{\beta,Pro,A}$), 1.96–1.77 (m, 3H, $H_{\beta,Pro,B}$, $H_{\gamma,Pro}$), 1.75–1.68 (m, 1H, $H_{\beta,Arg,A}$), 1.65–1.57 (m, 1H, $H_{\beta,Lys,A}$), 1.58–1.40 (m, 8H, $H_{\beta,Arg,B}$, $H_{\gamma,Arg}$, $H_{\beta,Lys,B}$, $(C\underline{H}_2)_{11}CH_3$), 1.39–1.31 (m, 2H, $H_{\gamma,Lys}$), 1.30–1.20 (m, 20H, $H_{\delta,Lys}$, $(C\underline{H}_2)_{11}CH_3$), 0.86 (t, *J* = 7.0 Hz, 3H, $CH_3$). $^{13}$C NMR (151 MHz, DMSO-$d_6$) δ 173.3 ($CO_{Arg}$), 171.3 ($CO_{Pro}$), 170.8 ($CO_{Lys}$), 156.7 (NHC(=NH)$NH_2$), 156.0 ($CO_{Cbz}$), 137.0 ($C1_{Ar,Cbz}$), 128.3 ($C3_{Ar,Cbz,}$ $C5_{Ar,Cbz}$), 127.8 ($C4_{Ar,Cbz}$), 127.7 ($C2_{Ar,Cbz,}$ $C6_{Ar,Cbz}$), 65.4 ($CH_{2,Cbz}$), 59.6 ($C_{\alpha,Pro}$), 52.4 ($C_{\alpha,Lys}$), 51.8 ($C_{\alpha,Arg}$), 46.9 ($C_{\delta,Pro}$), 43.4 ($C_{\varepsilon,Lys}$), 40.4 ($C_{\delta,Arg}$), 31.3 ($\underline{C}H_2CH_2CH_3$), 30.5 ($C_{\beta,Lys}$), 29.0–28.7 (10C, $C_{\beta,Arg}$, $(\underline{C}H_2)_{11}CH_3$), 26.4 ($C_{\delta,Lys}$), 25.0 ($C_{\gamma,Arg}$), 24.6 ($C_{\gamma,Pro}$), 22.6 ($C_{\gamma,Lys}$), 22.1 ($\underline{C}H_2CH_3$), 13.9 ($CH_3$). The peak for $C_{\varepsilon,Lys}$ was broad and of low intensity and the peak for C=S was not visible in $^{13}$C NMR due to fast quadrupolar relaxation via the nearby $^{14}$N-nuclei. Two sets of signals (approximately 10:1) were detectable due to rotamers. Only peaks for the major rotamer is given. Analytical HPLC gradient 0–95% eluent II in eluent I (C18; 35 min total runtime), $t_R$ 26.4 min (>98%, $UV_{230}$). HRMS calcd for $C_{38}H_{66}N_9O_5S^+$ $[M+H]^+$, 760.4902; found 760.4900.

**Benzyl ((6*S*,9*S*,12*S*)-1-amino-6-carbamoyl-1-imino-9-methyl-8,11-dioxo-18-thioxo-2,7,10,17,19-pentaazahentriacontan-12-yl)carbamate (20).** 

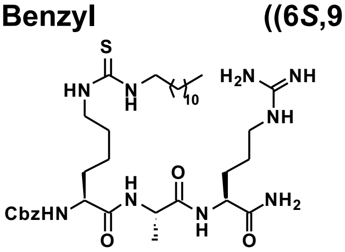

Starting from Cbz-Lys-Ala-Arg(Pbf)-resin (87 mg, estimated loading: 0.46 mmol/g) synthesized from Fmoc-Arg(Pbf)-OH, Fmoc-Ala-OH and Cbz-Lys(Fmoc)-OH by SPPS, the title compound was synthesized by on-resin thiourea formation as described in the general procedures. Global deprotection and cleavage from the resin, followed by preparative reversed-phase HPLC purification afforded the desired thiourea **20** (3 mg, 10% based on resin loading), as a colorless fluffy material after lyophilization. $^1$H NMR (600 MHz, DMSO-$d_6$) δ 8.05 (d, *J* = 7.0 Hz, 1H, $NH_{Ala}$), 7.83 (d, *J* = 8.1 Hz, 1H, $NH_{\alpha,Arg}$), 7.52 (t, *J* = 5.8 Hz, 1H, $NH_{\delta,Arg}$), 7.42 (d, *J* = 8.0 Hz, 1H, $NH_{\alpha,Lys}$), 7.38–7.29 (m, 8H, $NH_{\varepsilon,Lys}$, $N\underline{H}CH_2$, $H_{Ar,Cbz}$, $CONH_{2,A}$), 7.08 (s, 1H, $CONH_{2,B}$), 5.06–4.98 (m, 2H, $CH_{2,Cbz}$), 4.25 (p, *J* = 7.1 Hz, 1H, $H_{\alpha,Ala}$), 4.18 (td, *J* = 8.0, 5.6

Hz, 1H, $H_{\alpha,Arg}$), 3.97 (ddd, *J* = 9.5, 7.9, 4.7 Hz, 1H, $H_{\alpha,Lys}$), 3.21 (br s, 2H, $H_{\varepsilon,Lys}$, overlap with residual water), 3.12–3.05 (m, 2H, $H_{\delta,Arg}$), 1.73–1.59 (m, 2H, $H_{\beta,Arg,A}$, $H_{\beta,Lys,A}$), 1.56–1.39 (m, 8H, $H_{\beta,Arg,B}$, $H_{\beta,Lys,B}$, $H_{\gamma,Arg}$, $(C\underline{H}_2)_{11}CH_3$), 1.36–1.24 (m, 22H, $H_{\gamma,Lys}$, $H_{\delta,Lys}$, $(C\underline{H}_2)_{11}CH_3$), 1.22 (d, *J* = 7.0 Hz, 3H, $H_{\beta,Ala}$), 0.85 (t, *J* = 6.9 Hz, 3H, $CH_2C\underline{H}_3$). $^{13}$C NMR (151 MHz, DMSO-$d_6$) δ 173.1 ($CO_{Arg}$), 172.0 ($CO_{Ala}$), 171.8 ($CO_{Lys}$), 156.6 ($NHC(=NH)NH_2$), 156.0 ($CO_{Cbz}$), 137.0 ($C1_{Ar,Cbz}$), 128.3 ($C3_{Ar,Cbz,}$ $C5_{Ar,Cbz}$), 127.8 ($C4_{Ar,Cbz}$), 127.6 ($C2_{Ar,Cbz,}$ $C6_{Ar,Cbz}$), 65.4 ($CH_{2,Cbz}$), 54.6 ($C_{\alpha,Lys}$), 51.8 ($C_{\alpha,Arg}$), 48.3 ($C_{\alpha,Ala}$), 43.4 ($C_{\varepsilon,Lys}$), 40.4 ($C_{\delta,Arg}$), 31.6 ($C_{\beta,Lys}$), 31.3 ($\underline{C}H_2CH_2CH_3$), 29.2–28.7 (10C, $C_{\beta,Arg}$, $(\underline{C}H_2)_{11}CH_3$), 26.4 ($C_{\delta,Lys}$), 25.0 ($C_{\gamma,Arg}$), 23.0 ($C_{\gamma,Lys}$), 22.1 ($\underline{C}H_2CH_3$), 18.0 ($C_{\beta,Ala}$), 13.9 ($CH_2\underline{C}H_3$). The peak for $C_{\varepsilon,Lys}$ was broad and of low intensity and the peak for C=S was not visible in $^{13}$C NMR due to fast quadrupolar relaxation via the nearby $^{14}$N-nuclei. Analytical HPLC gradient 0–95% eluent II in eluent I (C18; 20 min total runtime), $t_R$ 13.3 min (>96%, $UV_{230}$). HRMS calcd for $C_{36}H_{64}N_9O_5S^+$ $[M+H]^+$, 734.4746; found 734.4743.

**Benzyl ((6*S*,9*R*,12*S*)-1-amino-6-carbamoyl-1-imino-9-methyl-8,11-dioxo-18-thioxo-2,7,10,17,19-pentaazahentriacontan-12-yl)carbamate (21).** 

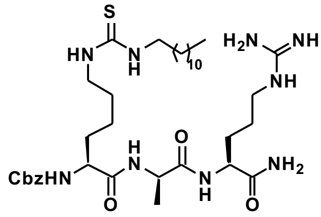

Starting from Cbz-Lys-D-Ala-Arg(Pbf)-resin (87 mg, estimated loading: 0.46 mmol/g) synthesized from Fmoc-Arg(Pbf)-OH, Fmoc-D-Ala-OH and Cbz-Lys(Fmoc)-OH by SPPS, the title compound was synthesized by on-resin thiourea formation as described in the general procedures. Global deprotection and cleavage from the resin, followed by preparative reversed-phase HPLC purification afforded the desired thiourea **21** (2 mg, 7% based on resin loading) as a colorless fluffy material after lyophilization. $^1$H NMR (600 MHz, DMSO-$d_6$) δ 8.15 (d, *J* = 7.2 Hz, 1H, $NH_{D\text{-}Ala}$), 7.93 (d, *J* = 8.4 Hz, 1H, $NH_{\alpha,Arg}$), 7.50 (t, *J* = 5.8 Hz, 1H, $NH_{\delta,Arg}$), 7.44 (d, *J* = 7.5 Hz, 1H, $NH_{\alpha,Lys}$), 7.40–7.26 (m, 8H, $NH_{\varepsilon,Lys}$, $N\underline{H}CH_2$, $H_{Ar,Cbz}$, $CONH_{2,A}$), 7.10 (s, 1H, $CONH_{2,B}$), 5.10–4.95 (m, 2H, $CH_{2,Cbz}$), 4.28 (p, *J* = 7.1 Hz, 1H, $H_{\alpha,D\text{-}Ala}$), 4.19 (td, *J* = 8.6, 5.1 Hz, 1H, $H_{\alpha,Arg}$), 3.97 (q, *J* = 7.6, 1H, $H_{\alpha,Lys}$), 3.30 (br s, 2H, $H_{\varepsilon,Lys}$, overlap with residual water), 3.11–3.02 (m, 2H, $H_{\delta,Arg}$), 1.80–1.70 (m, 1H, $H_{\beta,Arg,A}$), 1.65–1.57 (m, 1H, $H_{\beta,Lys,A}$), 1.56–1.37 (m, 8H, $H_{\beta,Arg,B}$, $H_{\beta,Lys,B}$, $H_{\gamma,Arg}$, $CH_2(C\underline{H}_2)_{11}CH_3$), 1.36–1.08 (m, 25H, $H_{\gamma,Lys}$, $H_{\delta,Lys}$, $(C\underline{H}_2)_{11}CH_3$, $H_{\beta,D\text{-}Ala}$), 0.85 (t, *J* = 6.8 Hz, 3H, $CH_2C\underline{H}_3$). $^{13}$C NMR (151 MHz, DMSO-$d_6$) δ 173.2 ($CO_{Arg}$), 172.0 ($CO_{D\text{-}Ala}$), 171.8 ($CO_{Lys}$), 156.6 ($NHC(=NH)NH_2$), 156.0 ($CO_{Cbz}$), 136.9 ($C1_{Ar,Cbz}$), 128.3 ($C3_{Ar,Cbz,}$ $C5_{Ar,Cbz}$), 127.8 ($C4_{Ar,Cbz}$), 127.7 ($C2_{Ar,Cbz,}$ $C6_{Ar,Cbz}$), 65.4 ($CH_{2,Cbz}$), 54.7 ($C_{\alpha,Lys}$), 51.8 ($C_{\alpha,Arg}$), 48.4 ($C_{\alpha,D\text{-}Ala}$), 43.3 ($C_{\varepsilon,Lys}$), 40.3 ($C_{\delta,Arg}$), 31.5 ($C_{\beta,Lys}$), 31.3 ($\underline{C}H_2CH_2CH_3$), 29.0–28.7 (10C, $C_{\beta,Arg}$ , $(\underline{C}H_2)_{11}CH_3$), 26.4 ($C_{\delta,Lys}$), 25.0 ($C_{\gamma,Arg}$), 22.9 ($C_{\gamma,Lys}$), 22.1 ($\underline{C}H_2CH_3$), 18.2 ($C_{\beta,Ala}$), 13.9 ($CH_2\underline{C}H_3$). The peak for $C_{\varepsilon,Lys}$ was broad and of low intensity and the peak for C=S was not visible in $^{13}$C NMR due to fast quadrupolar relaxation via the nearby $^{14}$N-nuclei Analytical HPLC gradient 0–95% eluent II in eluent I (C18; 35 min total runtime), $t_R$ 13.3 min (>98%, $UV_{254}$). HRMS calcd for $C_{36}H_{64}N_9O_5S^+$ $[M+H]^+$, 734.4746; found 734.4740.

**Benzyl ((6*S*,12*S*)-1-amino-6-carbamoyl-1-imino-9-methylene-8,11-dioxo-18-thioxo-2,7,10,17,19-pentaazahentriacontan-12-yl)carbamate (22).**

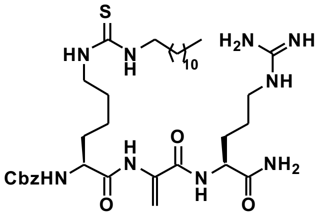

Starting from Cbz-Lys(Fmoc)-Cys(StBu)-Arg(Pbf)-resin (166 mg, estimated loading: 0.33 mmol/g) synthesized from Fmoc-Arg(Pbf)-OH, Fmoc-Cys(StBu)-OH and Cbz-Lys(Fmoc)-OH by SPPS, the title compound was synthesized by addition of dithiotreitol (DTT, 77 mg, 0.50 mmol) and *i*$Pr_2NEt$ (173 µL, 1.00 mmol) in anh. DMF (2.0 mL) to the resin under agitation for 6 h. The resin was washed with DMF (3×4 mL) after which 1,4-dibromobutane (65 mg, 0.30 mmol) in 1.5 mL anh. DMF and $K_2CO_3$ (69 mg, 0.5 mmol) were added. The resin was agitated for 6 h and washed with DMF, $H_2O$, MeOH and $CH_2Cl_2$ (3×4 mL each) followed by subsequent Fmoc-deprotection and general on-resin thiourea formation as described in the general procedures. Global deprotection and cleavage from the resin, followed by preparative reversed-phase HPLC purification afforded the desired thiourea **22** (1 mg, 2% based on resin loading) as a white fluffy material after lyophilization. $^{1}$H NMR (600 MHz, DMSO-$d_6$) δ 9.23 (s, 1H, $NH_{Dha}$), 8.29 (d, *J* = 8.0 Hz, 1H, $NH_{\alpha,Arg}$), 7.73 (d, *J* = 7.6 Hz, 1H, $NH_{\alpha,Lys}$), 7.49 (t, *J* = 5.8 Hz, 1H, $NH_{\delta,Arg}$), 7.39–7.28 (m, 8H, $NH_{\varepsilon,Lys}$, $N\underline{H}CH_2$, $H_{Ar,Cbz}$, $CONH_{2,A}$), 7.10 (d, *J* = 2.0 Hz, 1H, $CONH_{2,B}$), 6.08 (s, 1H, $H_{\beta,Dha,A}$), 5.59 (s, 1H, $H_{\beta,Dha,B}$), 5.09–5.01 (m, 2H, $CH_{2,Cbz}$), 4.29–4.20 (m, 1H, $H_{\alpha,Arg}$), 3.97 (dt, *J* = 11.6, 7.6 Hz, 1H, $H_{\alpha,Lys}$), 3.30 (br s, 2H, $H_{\varepsilon,Lys}$, overlap with residual water), 3.13–3.06 (m, 2H, $H_{\delta,Arg}$), 1.83–1.74 (m, 1H, $H_{\beta,Arg,A}$), 1.72–1.40 (m, 10H, $H_{\beta,Arg,B}$, $H_{\beta,Lys}$, $H_{\gamma,Arg}$, $H_{\gamma,Lys,A}$, $C\underline{H}_2(CH_2)_{10}CH_3$, $CH_2(C\underline{H}_2)_{10}CH_3$), 1.38–1.20 (m, 21H, $H_{\gamma,Lys,B}$, $H_{\delta,Lys}$, $CH_2(C\underline{H}_2)_{10}CH_3$), 0.85 (t, *J* = 6.9 Hz, 3H, $CH_3$). $^{13}$C NMR (151 MHz, DMSO-$d_6$) δ 173.2 ($CO_{Arg}$), 171.6 ($CO_{Lys}$), 163.8 ($CO_{Dha}$), 156.6 (NHC(=NH)$NH_2$), 156.2 ($CO_{Cbz}$), 136.9 ($C1_{Ar,Cbz}$), 135.0 ($C_{\alpha,Dha}$), 128.4 ($C3_{Ar,Cbz,}$ $C5_{Ar,Cbz}$), 127.8 ($C4_{Ar,Cbz}$), 127.6 ($C2_{Ar,Cbz,}$ $C6_{Ar,Cbz}$), 104.1 ($C_{\beta,Dha}$), 65.6 ($CH_{2,Cbz}$), 55.6 ($C_{\alpha,Lys}$), 52.9 ($C_{\alpha,Arg}$), 43.4 ($C_{\varepsilon,Lys}$), 40.4 ($C_{\delta,Arg}$), 31.3 ($\underline{C}H_2CH_2CH_3$), 31.0 ($C_{\beta,Lys}$), 29.0–28.5 (10C, $C_{\beta,Arg}$, $(\underline{C}H_2)_{11}CH_3$), 26.4 ($C_{\delta,Lys}$), 25.4 ($C_{\gamma,Arg}$), 23.1 ($C_{\gamma,Lys}$), 22.1 ($\underline{C}H_2CH_3$), 13.9 ($CH_3$). The peak for $C_{\varepsilon,Lys}$ was broad and of low intensity and the peak for C=S was not visible in $^{13}$C NMR due to fast quadrupolar relaxation via the nearby $^{14}$N-nuclei. Analytical HPLC gradient 0–95% eluent II in eluent I (C8; 35 min total runtime), $t_R$ 26.7 min (>97%, $UV_{230}$). HRMS calcd for $C_{36}H_{62}N_9O_5S^+$ [M+H]$^+$, 732.4589; found 732.4585. Dha = dehydroalanine.

**Benzyl ((6*S*,12*S*)-1-amino-6-carbamoyl-1-imino-9,9-dimethyl-8,11-dioxo-18-thioxo-2,7,10,17,19-pentaazahentriacontan-12-yl)carbamate (23).**

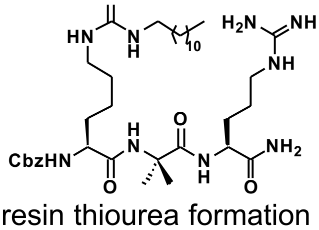

Starting from Cbz-Lys-Aib-Arg(Pbf)resin (258 mg, estimated loading: 0.33 mmol/g) synthesized from Fmoc-Arg(Pbf)-OH, Fmoc-Aib-OH and Cbz-Lys(Fmoc)-OH by SPPS, the title compound was synthesized by on-resin thiourea formation as described in the general procedures. Global deprotection and cleavage from the resin, followed by preparative reversed-phase HPLC purification afforded the desired thiourea **23** (3 mg, 5% based on resin loading) as a white fluffy material after lyophilization. $^{1}$H NMR

(600 MHz, DMSO-$d_6$) δ 8.26 (s, 1H, $NH_{Aib}$), 7.52–7.46 (m, 2H, $NH_{α,Lys}$, $NH_{δ,Arg}$), 7.39–7.28 (m, 8H, $NH_{α,Lys}$, $NH_{ε,Lys}$, $N\underline{H}CH_2$, $H_{Ar,Cbz}$), 7.18 (s, 1H, $CONH_{2,A}$), 7.07 (s, 1H, $CONH_{2,B}$), 5.08–4.97 (m, 2H, $CH_{2,Cbz}$), 4.10 (td, $J$ = 8.8, 4.5 Hz, 1H, $H_{α,Arg}$), 3.93 (dt, $J$ = 8.7, 6.0 Hz, 1H, $H_{α,Lys}$), 3.30 (br s, 2H, $H_{ε,Lys}$, overlap with residual water), 3.07–3.00 (m, 2H, $H_{δ,Arg}$), 1.81–1.73 (m, 1H, $H_{β,Arg,A}$), 1.67–1.58 (m, 1H, $H_{β,Lys,A}$), 1.49–1.39 (m, 8H, $H_{β,Arg,B}$, $H_{β,Lys,B}$, $H_{γ,Arg}$, $(C\underline{H}_2)_{11}CH_3$), 1.37–1.30 (m, 7H, $H_{β,Aib}$, $H_{γ,Lys,A}$), 1.29–1.19 (m, 21H, $H_{γ,Lys,B}$, $H_{δ,Lys}$, $(C\underline{H}_2)_{11}CH_3$), 0.85 (t, $J$ = 7.0 Hz, 3H, $CH_2C\underline{H}_3$). $^{13}C$ NMR (151 MHz, DMSO-$d_6$) δ 173.7 ($CO_{Aib}$), 173.4 ($CO_{Arg}$), 172.3 ($CO_{Lys}$), 156.6 ($NHC(=NH)NH_2$), 156.2 ($CO_{Cbz}$), 136.9 ($C1_{Ar,Cbz}$), 128.3 ($C3_{Ar,Cbz,}$ $C5_{Ar,Cbz}$), 127.8 ($C4_{Ar,Cbz}$), 127.6 ($C2_{Ar,Cbz,}$ $C6_{Ar,Cbz}$), 65.5 ($CH_{2,Cbz}$), 56.0 ($C_{α,Aib}$), 55.1 ($C_{α,Lys}$), 52.0 ($C_{α,Arg}$), 43.4 ($C_{ε,Lys}$), 40.3 ($C_{δ,Arg}$), 31.3 ($\underline{C}H_2CH_2CH_3$), 31.0 ($C_{β,Lys}$), 29.0–28.4 (10C, $C_{β,Arg}$, $(\underline{C}H_2)_{11}CH_3$), 26.4 ($C_{δ,Lys}$), 25.2 ($C_{β,Aib,1}$), 24.9 ($C_{γ,Arg}$), 24.7 ($C_{β,Aib,2}$), 22.9 ($C_{γ,Lys}$), 22.1 ($\underline{C}H_2CH_3$), 13.9 ($CH_2\underline{C}H_3$). The peak for $C_{ε,Lys}$ was broad and of low intensity and the peak for C=S was not visible in $^{13}C$ NMR due to fast quadrupolar relaxation via the nearby $^{14}N$-nuclei. Analytical HPLC gradient 0–95% eluent II in eluent I (C18; 35 min total runtime), $t_R$ 25.6 min (>98%, $UV_{230}$). HRMS calcd for $C_{37}H_{66}N_9O_5S^+$ [M+H]$^+$, 748.4902; found 748.4898.

**(*S*)-2-acetamido-*N*-((*S*)-1-(((*S*)-1-amino-5-guanidino-1-oxopentan-2-yl)amino)-1-oxopropan-2-yl)-6-(3-dodecylthioureido)hexanamide (24).**

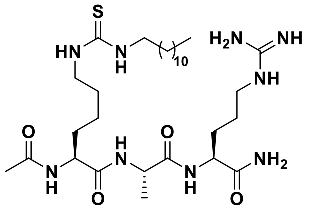

Starting from H-Lys(Teoc)-Ala-Arg(Pbf)-resin (104 mg, estimated loading: 0.49 mmol/g) synthesized from Fmoc-Arg(Pbf)-OH, Fmoc-Ala-OH and Fmoc-Lys(Teoc)-OH by SPPS, the title compound was synthesized by addition of $Ac_2O$:DMF (1:3, v/v, 1.5 mL) to the resin under agitation for 2 h. The resin was washed with DMF (3×4 mL) and $CH_2Cl_2$ (3×4 mL) followed by Teoc deprotection and on-resin thiourea formation as described in the general procedures. Global deprotection and cleavage from the resin, followed by preparative reversed-phase HPLC purification afforded the desired thiourea **24** (4 mg, 11% based on resin loading), as a colorless fluffy material after lyophilization. $^1H$ NMR (600 MHz, DMSO-$d_6$) δ 8.06 (d, $J$ = 7.0 Hz, 1H, $NH_{Ala}$), 8.02 (d, $J$ = 7.6 Hz, 1H, $NH_{α,Lys}$), 7.76 (d, $J$ = 8.0 Hz, 1H, $NH_{α,Arg}$), 7.57 (t, $J$ = 5.8 Hz, 1H, $NH_{δ,Arg}$), 7.38–7.26 (m, 3H, $NH_{ε,Lys}$, $N\underline{H}CH_2$, $CONH_{2,A}$), 7.08 (s, 1H, $CONH_{2,B}$), 4.26–4.12 (m, 3H, $H_{α,Ala}$, $H_{α,Lys}$, $H_{α,Arg}$), 3.32 (br s, 2H, 2H, $H_{ε,Lys}$, overlap with residual water), 3.09 (q, $J$ = 6.7 Hz, 2H, $H_{δ,Arg}$), 1.85 (s, 3H, $COCH_3$), 1.73–1.60 (m, 2H, $H_{β,Arg,A}$. $H_{β,Lys,A}$), 1.57–1.40 (m, 8H, $H_{β,Arg,B}$, $H_{β,Lys,B}$, $H_{γ,Arg}$, $(C\underline{H}_2)_{11}CH_3$), 1.35–1.19 (m, 25H, $H_{γ,Lys}$, $H_{δ,Lys}$, $(C\underline{H}_2)_{11}CH_3$, $H_{β,Ala}$), 0.85 (t, $J$ = 6.9 Hz, 3H, $CH_2C\underline{H}_3$). $^{13}C$ NMR (151 MHz, DMSO-$d_6$) δ 173.2 ($CO_{Arg}$), 172.0 ($CO_{Ala}$), 171.9 ($CO_{Lys}$), 169.6 ($\underline{C}OCH_3$), 156.7 ($NHC(=NH)NH_2$), 52.8 ($C_{α,Lys}$), 51.9 ($C_{α,Arg}$), 48.4 ($C_{α,Ala}$), 43.4 ($C_{ε,Lys}$), 40.4 ($C_{δ,Arg}$), 31.5 ($C_{β,Lys}$) , 31.3 ($\underline{C}H_2CH_2CH_3$), 29.1–28.7 (10C, $C_{β,Arg}$, $(\underline{C}H_2)_{11}CH_3$), 26.4 ($C_{δ,Lys}$), 25.0 ($C_{γ,Arg}$), 22.9 ($C_{γ,Lys}$), 22.5 ($CO\underline{C}H_3$), 22.1 ($\underline{C}H_2CH_3$), 17.8 ($C_{β,Ala}$), 13.9 ($CH_2\underline{C}H_3$). The peak for $C_{ε,Lys}$ was broad and of low intensity and the peak for C=S was not visible in $^{13}C$ NMR due to fast quadrupolar relaxation via the nearby $^{14}N$-nuclei. Analytical HPLC gradient 0–95% eluent II in eluent

I (C8; 35 min total runtime), $t_R$ 23.9 min (>98%, $UV_{230}$). HRMS calcd for $C_{30}H_{59}N_9NaO_4S^+$ $[M+Na]^+$, 664.4302; found 664.4299.

**(*S*)-2-acetamido-*N*-((*R*)-1-(((*S*)-1-amino-5-guanidino-1-oxopentan-2-yl)amino)-1-oxopropan-2-yl)-6-(3-dodecylthioureido)hexanamide (24-D).**

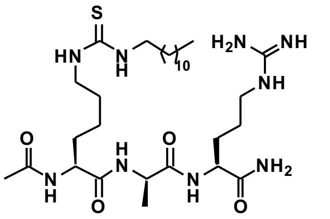

Starting from H-Lys(Teoc)-D-Ala-Arg(Pbf)-resin (168 mg, estimated loading: 0.33 mmol/g) synthesized from Fmoc-Arg(Pbf)-OH, Fmoc-D-Ala-OH and Fmoc-Lys(Teoc)-OH by SPPS, the title compound was synthesized by addition of $Ac_2O$:$CH_2Cl_2$ (1:3, v/v, 1.5 mL) to the resin under agitation for 2 h. The resin was washed with DMF (3×4 mL) and $CH_2Cl_2$ (3×4 mL) followed by Teoc deprotection and on-resin thiourea formation as described in the general procedures. Global deprotection and cleavage from the resin, followed by preparative reversed-phase HPLC purification afforded the desired thiourea **24-D** (4 mg, 11% based on resin loading), as a colorless fluffy material after lyophilization. $^1H$ NMR (600 MHz, DMSO-$d_6$) δ 8.26 (d, $J$ = 7.3 Hz, 1H, $NH_{D\text{-}Ala}$), 8.09 (d, $J$ = 7.0 Hz, 1H, $NH_{\alpha,Lys}$), 7.87 (d, $J$ = 8.4 Hz, 1H, $NH_{\alpha,Arg}$), 7.51 (t, $J$ = 5.7 Hz, 1H, $NH_{\delta,Arg}$), 7.35–7.29 (m, 2H, $NH_{\varepsilon,Lys}$, $N\underline{H}(CH_2)_{11}CH_3$), 7.28 (d, $J$ = 2.1 Hz, 1H, $CONH_{2,A}$), 7.09 (d, $J$ = 2.1 Hz, 1H, $CONH_{2,B}$), 4.24 (p, $J$ = 7.2 Hz, 1H, $H_{\alpha,D\text{-}Ala}$), 4.19–4.09 (m, 2H, $H_{\alpha,Lys}$, $H_{\alpha,Arg}$), 3.31 (br s, 2H, $H_{\varepsilon,Lys}$, overlap with residual water), 3.06 (q, $J$ = 6.8 Hz, 2H, $H_{\delta,Arg}$), 1.84 (s, 3H, $COC\underline{H}_3$), 1.77–1.68 (m, 1H, $H_{\beta,Arg,A}$), 1.64–1.53 (m, 1H, $H_{\beta,Lys,A}$), 1.55–1.36 (m, 8H, $H_{\beta,Arg,B}$, $H_{\beta,Lys,B}$, $H_{\gamma,Arg}$, $C\underline{H}_2(CH_2)_{10}CH_3$, $CH_2(C\underline{H}_2)_{10}CH_3$), 1.29–1.19 (m, 25H, $H_{\gamma,Lys}$, $H_{\delta,Lys}$, $CH_2(C\underline{H}_2)_{10}CH_3$, $H_{\beta,D\text{-}Ala}$), 0.85 (t, $J$ = 7.0Hz, 3H, $CH_2C\underline{H}_3$). $^{13}C$ NMR (151 MHz, DMSO-$d_6$) δ 173.3 ($CO_{Arg}$), 172.0 ($CO_{D\text{-}Ala}$), 171.9 ($CO_{Lys}$), 169.7 ($\underline{C}OCH_3$), 156.6 ($NHC(=NH)NH_2$), 53.2 ($C_{\alpha,Lys}$), 52.0 ($C_{\alpha,Arg}$), 48.3 ($C_{\alpha,D\text{-}Ala}$), 43.3 ($C_{\varepsilon,Lys}$), 40.4 ($C_{\delta,Arg}$), 31.3 ($C_{\beta,Lys}$, $\underline{C}H_2CH_2CH_3$), 29.0–28.7 (10C, $C_{\beta,Arg}$, $(\underline{C}H_2)_{11}CH_3$), 26.4 ($C_{\delta,Lys}$), 25.1 ($C_{\gamma,Arg}$), 22.8 ($C_{\gamma,Lys}$), 22.4 ($CO\underline{C}H_3$), 22.1 ($\underline{C}H_2CH_3$), 17.9 ($C_{\beta,D\text{-}Ala}$), 14.0 ($CH_2\underline{C}H_3$). The peak for $C_{\varepsilon,Lys}$ was broad and of low intensity and the peak for C=S was not visible in $^{13}C$ NMR due to fast quadrupolar relaxation via the nearby $^{14}N$-nuclei. Analytical HPLC gradient 0–95% eluent II in eluent I (C8; 35 min total runtime), $t_R$ 23.1 min (>98%, $UV_{230}$). HRMS calcd for $C_{30}H_{60}N_9O_4S^+$ $[M+H]^+$, 642.4483; found 642.4479.

**(*S*)-*N*-((*S*)-1-(((*S*)-1-amino-5-guanidino-1-oxopentan-2-yl)amino)-1-oxopropan-2-yl)-6-(3-dodecylthioureido)-2-(methylsulfonamido)hexanamide (25).**

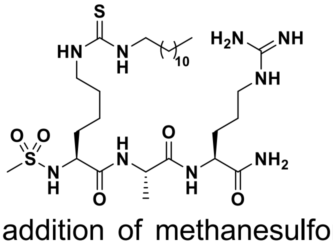

Starting from H-Lys(Teoc)-Ala-Arg(Pbf)-resin (163 mg, estimated loading: 0.49 mmol/g) synthesized from Fmoc-Arg(Pbf)-OH, Fmoc-Ala-OH and Fmoc-Lys(Teoc)-OH by SPPS, the title compound was synthesized by addition of methanesulfonyl chloride (18 μL, 0.24 mmol) and $i$$Pr_2NEt$ (83 μL, 0.48 mmol) in anh. $CH_2Cl_2$ (2.0 mL) to the resin under overnight agitation. The resin was washed with DMF (3×4 mL) and $CH_2Cl_2$ (3×4 mL) followed by Teoc deprotection and on-resin thiourea formation as described in the general procedures. Global deprotection and cleavage from the resin, followed by preparative reversed-phase HPLC purification afforded the desired thiourea **25** (3 mg, 5% based on resin loading), as a colorless fluffy material after lyophilization. $^{1}$H NMR (600 MHz, DMSO-$d_6$) δ 8.17 (d, $J$ = 7.1 Hz, 1H, $NH_{Ala}$), 7.90 (d, $J$ = 8.0 Hz, 1H, $NH_{\alpha,Arg}$), 7.54 (t, $J$ = 5.8 Hz, $NH_{\delta,Arg}$), 7.38–7.28 (m, 4H, $NH_{\alpha,Lys}$, N$\underline{H}$CH$_2$, $NH_{\varepsilon,Lys}$, $CONH_{2,A}$), 7.07 (d, $J$ = 2.2 Hz, 1H, $CONH_{2,B}$), 4.29 (p, $J$ = 7.1 Hz, 1H, $H_{\alpha,Ala}$), 4.18 (td, $J$ = 8.0, 5.6 Hz, 1H, $H_{\alpha,Arg}$), 3.78 (td, $J$ = 8.8, 5.2 Hz, 1H, $H_{\alpha,Lys}$), 3.31 (br s, 2H, $H_{\varepsilon,Lys}$, overlap with residual water), 3.12–3.06 (m, 2H, $H_{\delta,Arg}$), 2.85 (s, 3H, $CH_3SO_2$), 1.72–1.58 (m, 2H, $H_{\beta,Arg,A}$, $H_{\beta,Lys,A}$), 1.55–1.19 (m, 33H, $H_{\beta,Arg,B}$, $H_{\gamma,Arg}$, $H_{\beta,Lys,B}$, $H_{\gamma,Lys}$, $H_{\delta,Lys}$, (C$\underline{H}_2$)$_{11}$CH$_3$), $H_{\beta,Ala}$), 0.85 (t, $J$ = 6.9 Hz, 3H, CH$_2$C$\underline{H}_3$). $^{13}$C NMR (151 MHz, DMSO-$d_6$) δ 173.1 ($CO_{Arg}$), 171.8 ($CO_{Ala}$), 171.3 ($CO_{Lys}$), 156.7 (NHC(=NH)NH$_2$), 56.3 ($C_{\alpha,Lys}$), 51.9 ($C_{\alpha,Arg}$), 48.3 ($C_{\alpha,Ala}$), 43.3 ($C_{\varepsilon,Lys}$), 40.7 ($CH_3SO_2$), 40.4 ($C_{\delta,Arg}$), 32.4 ($C_{\beta,Lys}$), 31.3 ($\underline{C}$H$_2$CH$_2$CH$_3$), 29.2–28.4 (10C, $C_{\beta,Arg}$, ($\underline{C}$H$_2$)$_{11}$CH$_3$), 26.4 ($C_{\delta,Lys}$), 25.0 ($C_{\gamma,Arg}$), 22.7 ($C_{\gamma,Lys}$), 22.1 ($\underline{C}$H$_2$CH$_3$), 18.1 ($C_{\beta,Ala}$), 13.9 (CH$_2\underline{C}$H$_3$).The peak for $C_{\varepsilon,Lys}$ was broad and of low intensity and the peak for C=S was not visible in $^{13}$C NMR due to fast quadrupolar relaxation via the nearby $^{14}$N-nuclei. Analytical HPLC gradient 0–95% eluent II in eluent I (C8; 35 min total runtime), $t_R$ 24.1 min (>97%, UV$_{230}$). HRMS calcd for $C_{29}H_{59}N_9NaO_5S_2^+$ [M+Na]$^+$, 700.3972; found 700.3964.

**(*S*)-*N*-((*R*)-1-(((*S*)-1-amino-5-guanidino-1-oxopentan-2-yl)amino)-1-oxopropan-2-yl)-6-(3-dodecylthioureido)-2-(methylsulfonamido)hexanamide (25-D).**

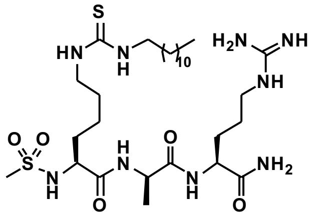

Starting from H-Lys(Teoc)-D-Ala-Arg(Pbf)-resin (168 mg, estimated loading: 0.33 mmol/g) synthesized from Fmoc-Arg(Pbf)-OH, Fmoc-D-Ala-OH and Fmoc-Lys(Teoc)-OH by SPPS, the title compound was synthesized by addition of methanesulfonyl chloride (13 µL, 0.17 mmol) and *i*Pr$_2$NEt (59 µL, 0.34 mmol) in anh. DMF (1.0 mL) to the resin under overnight agitation. The resin was washed with DMF (3×4 mL) and CH$_2$Cl$_2$ (3×4 mL) followed by Teoc deprotection and on-resin thiourea formation as described in the general procedures. Global deprotection and cleavage from the resin, followed by preparative reversed-phase HPLC purification afforded the desired thiourea **25-D** (3 mg, 8% based on resin loading) as a colorless fluffy material after lyophilization. $^{1}$H NMR (600 MHz, DMSO-$d_6$) δ 8.37 (d, $J$ = 6.5 Hz, 1H, $NH_{D\text{-}Ala}$), 8.10 (d, $J$ = 8.3 Hz, 1H, $NH_{\alpha,Arg}$), 7.53 (t, $J$ = 5.8 Hz, $NH_{\delta,Arg}$), 7.36 (d, $J$ = 9.0 Hz, 1H, $NH_{\alpha,Lys}$), 7.32 (br s, 2H, N$\underline{H}$CH$_2$, $NH_{\varepsilon,Lys}$), 7.27 (d, $J$ = 2.2 Hz, 1H, $CONH_{2,A}$), 7.12 (d, $J$ = 2.2 Hz, 1H, $CONH_{2,B}$), 6.84 (s, 1H, residual $CO_2H_{TFA}$), 4.25 (p, $J$ = 7.0 Hz, 1H, $H_{\alpha,D\text{-}Ala}$), 4.15 (td, $J$ = 8.4, 5.0 Hz, 1H, $H_{\alpha,Arg}$), 3.83 (td, $J$ = 8.8, 5.5 Hz, 1H, $H_{\alpha,Lys}$), 3.31 (br s, 2H, $H_{\varepsilon,Lys}$, overlap with residual water), 3.11–3.04 (m, 2H, $H_{\delta,Arg}$), 2.79 (s, 3H, $CH_3SO_2$), 1.79–1.70 (m, 1H, $H_{\beta,Arg,A}$), 1.61–1.54 (m, 1H, $H_{\beta,Lys,A}$), 1.52–1.31 (m, 10H, $H_{\beta,Arg,B}$, $H_{\beta,Lys,B}$, $H_{\gamma,Arg}$, $H_{\gamma,Lys}$, (C$\underline{H}_2$)$_{11}$CH$_3$), 1.30–1.23 (m, 20H, $H_{\delta,Lys}$, (C$\underline{H}_2$)$_{11}$CH$_3$), 1.22 (t, $J$ = 7.0 Hz, 3H, $H_{\beta,D\text{-}Ala}$), 0.85 (t, $J$ = 7.0 Hz, 3H, CH$_2$C$\underline{H}_3$). $^{13}$C NMR (151 MHz, DMSO-$d_6$) δ 173.2 ($CO_{Arg}$), 172.3 ($CO_{D\text{-}Ala}$), 171.6 ($CO_{Lys}$), 158.1 (q, $J$ = 31.0 Hz, residual $CO_{TFA}$), 156.6 (NHC(=NH)NH$_2$), 117.3 (q, $J$ = 300.2 Hz, residual $CF_{3,TFA}$), 56.0 ($C_{\alpha,Lys}$), 51.8 ($C_{\alpha,Arg}$), 48.7 ($C_{\alpha,D\text{-}Ala}$), 43.4 ($C_{\varepsilon,Lys}$), 40.7 ($CH_3SO_2$), 40.3 ($C_{\delta,Arg}$), 32.5 ($C_{\beta,Lys}$), 31.3 ($\underline{C}$H$_2$CH$_2$CH$_3$), 29.0–28.7 (10C, $C_{\beta,Arg}$, ($\underline{C}$H$_2$)$_{11}$CH$_3$), 26.4 ($C_{\delta,Lys}$), 25.1 ($C_{\gamma,Arg}$), 22.7 ($C_{\gamma,Lys}$), 22.1 ($\underline{C}$H$_2$CH$_3$), 17.8 ($C_{\beta,D\text{-}}$

$_{Ala}$), 14.0 ($CH_2\underline{C}H_3$). The peak for $C_{\varepsilon,Lys}$ was broad and of low intensity and the peak for C=S was not visible in $^{13}C$ NMR due to fast quadrupolar relaxation via the nearby $^{14}N$-nuclei. Analytical HPLC gradient 0–95% eluent II in eluent I (C8; 35 min total runtime), $t_R$ 22.9 min (>98%, $UV_{230}$). HRMS calcd for $C_{29}H_{60}N_9O_5S_2^+$ $[M+H]^+$, 678.4153; found 678.4145.

**(*S*)-*S*-((*R*)-1-(((*S*)-1-amino-5-guanidino-1-oxopentan-2-yl)amino)-1-oxopropan-2-yl)-2-(methylsulfonamido)-6-tetradecanethioamidohexanamide (26).**

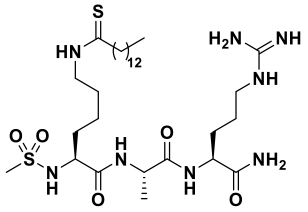

Starting from H-Lys(thiomyristoyl)-Ala-Arg(Pbf)-resin (204 mg, estimated loading: 0.49 mmol/g) synthesized from Fmoc-Arg(Pbf)-OH, Fmoc-Ala-OH and Fmoc-Lys(thiomyristoyl)-OH (**S34**) by SPPS, the title compound was synthesized by addition of methanesulfonyl chloride (25 µL, 0.32 mmol) and *i*$Pr_2NEt$ (111 µL, 0.64 mmol) in 3 mL anh. $CH_2Cl_2$ to the resin under overnight agitation. The resin was washed with DMF (3×4 mL) and $CH_2Cl_2$ (3×4 mL) followed by global deprotection, cleavage from the resin, followed by preparative reversed-phase HPLC purification afforded the desired thioamide **26** (17 mg, 25% based on resin loading), as a colorless fluffy material after lyophilization. $^1H$ NMR (600 MHz, DMSO) δ 9.88 (t, $J$ = 5.3 Hz, 1H, $NH_{\varepsilon,Lys}$), 8.16 (d, $J$ = 7.1 Hz, 1H, $NH_{Ala}$), 7.91 (d, $J$ = 8.1 Hz, 1H, $NH_{\alpha,Arg}$), 7.51 (t, $J$ = 5.8 Hz, 1H, $NH_{\delta,Arg}$), 7.35 (d, $J$ = 8.7 Hz, 1H, $NH_{\alpha,Lys}$), 7.32 (d, $J$ = 2.2 Hz, 1H, $CONH_{2,A}$), 7.07 (d, $J$ = 2.1 Hz, 1H, $CONH_{2,B}$), 4.30 (p, $J$ = 7.1 Hz, 1H, $H_{\alpha,Ala}$), 4.22–4.14 (m, 1H, $H_{\alpha,Arg}$), 3.82–3.76 (m, 1H, $H_{\alpha,Lys}$), 3.50–3.42 (m, 2H, $H_{\varepsilon,Lys}$, overlap with residual water), 3.09 (q, $J$ = 6.7 Hz, 2H, $H_{\delta,Arg}$), 2.85 (s, 3H, $CH_3SO_2$), 2.49 (s, 1H), 1.71–1.18 (m, 35H, $H_{\beta,Lys}$, $H_{\gamma,Lys}$, $H_{\delta,Lys}$, $H_{\beta,Arg}$, $H_{\gamma,Arg}$, $H_{\beta,Ala}$, $(C\underline{H}_2)_{11}CH_3$), 0.85 (t, J = 7.0 Hz, 3H, $CH_2C\underline{H}_3$). $^{13}C$ NMR (151 MHz, DMSO-$d_6$) δ 203.6 (C=S), 173.1 ($CO_{Arg}$), 172.3 ($CO_{Ala}$), 171.2 ($CO_{Lys}$), 156.6 ($NHC(=NH)NH_2$), 56.2 ($C_{\alpha,Lys}$), 51.8 ($C_{\alpha,Arg}$), 48.3 ($C_{\alpha,Ala}$), 45.02 ($CS\underline{C}H_2$), 44.99 ($C_{\varepsilon,Lys}$), 40.7 ($CH_3SO_2$), 40.4 ($C_{\delta,Arg}$), 32.3 ($C_{\beta,Lys}$), 31.3 ($\underline{C}H_2CH_2CH_3$), 29.2–28.7 (10C, $C_{\beta,Arg}$ , $(\underline{C}H_2)_{11}CH_3$), 26.7 ($C_{\delta,Lys}$), 25.0 ($C_{\gamma,Arg}$), 22.8 ($C_{\gamma,Lys}$), 22.1 ($\underline{C}H_2CH_3$), 18.1 ($C_{\beta,Ala}$), 13.9 ($CH_2\underline{C}H_3$). Analytical HPLC gradient 0–95% eluent II in eluent I (C8; 35 min total runtime), $t_R$ 25.5 min (>96%, $UV_{254}$). HRMS calcd for $C_{30}H_{60}N_8NaO_4S_2^+$ $[M+Na]^+$, 699.4020; found 699.4016.

**(*S*)-*N*-((*R*)-1-(((*S*)-1-amino-5-guanidino-1-oxopentan-2-yl)amino)-1-oxopropan-2-yl)-2-(methylsulfonamido)-6-tetradecanethioamidohexanamide (26-D).**

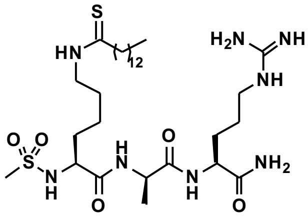

Starting from H-Lys(thiomyristoyl)-D-Ala-Arg(Pbf)-resin (228 mg, estimated loading: 0.47 mmol/g) synthesized from Fmoc-Arg(Pbf)-OH, Fmoc-D-Ala-OH and Fmoc-Lys(thiomyristoyl)-OH (**S34**) by SPPS, the title compound was synthesized by addition of methanesulfonyl chloride (25 µL, 0.32 mmol) and *i*$Pr_2NEt$ (111 µL, 0.64 mmol) in 2.0 mL anh. DMF to the resin under overnight agitation. The resin was washed with DMF (3×4 mL) and $CH_2Cl_2$ (3×4 mL) followed by global deprotection, cleavage from the resin, followed by preparative reversed-phase HPLC purification afforded the desired thioamide **26-D** (2 mg, 3% based on resin loading), as a colorless fluffy material after lyophilization. $^1H$ NMR

(600 MHz, DMSO-$d_6$) δ 9.88 (t, $J$ = 5.4 Hz, 1H, $NH_{\varepsilon,Lys}$), 8.37 (d, $J$ = 6.5 Hz, 1H, $NH_{D\text{-}Ala}$), 8.10 (d, $J$ = 8.3 Hz, 1H, $NH_{\alpha,Arg}$), 7.55 (t, $J$ = 5.8 Hz, 1H, $NH_{\delta,Arg}$), 7.36 (d, $J$ = 8.9 Hz, 1H, $NH_{\alpha,Lys}$), 7.27 (d, $J$ = 2.2 Hz, 1H, $CONH_{2,A}$), 7.12 (d, $J$ = 2.2 Hz, 1H, $CONH_{2,B}$), 4.28–4.21 (m, 1H, $H_{\alpha,D\text{-}Ala}$), 4.15 (td, $J$ = 8.5, 5.0 Hz, 1H, $H_{\alpha,Arg}$), 3.84 (td, $J$ = 8.8, 5.5 Hz, 1H, $H_{\alpha,Lys}$), 3.50–3.40 (m, 2H, $H_{\varepsilon,Lys}$), 3.12–3.04 (m, 2H, $H_{\delta,Arg}$), 2.79 (s, 3H, $CH_3SO_2$), 2.50–2.46 (m, 2H, $CSC\underline{H}_2$, overlap with solvent peak), 1.79–1.71 (m, 1H, $H_{\beta,Arg,A}$), 1.67–1.10 (m, 34H, $H_{\beta,Arg,B}$, $H_{\gamma,Arg}$, $H_{\beta,Lys}$, $H_{\gamma,Lys}$, $H_{\delta,Lys}$, $H_{\beta,D\text{-}Ala}$, $(C\underline{H}_2)_{11}CH_3$ ), 0.85 (t, $J$ = 7.0 Hz, 3H, $CH_2C\underline{H}_3$). $^{13}$C NMR (151 MHz, DMSO-$d_6$) δ 203.6 (C=S), 173.2 ($CO_{Arg}$), 172.3 ($CO_{D\text{-}Ala}$), 171.6 ($CO_{Lys}$), 156.7 ($NHC(=NH)NH_2$), 55.9 ($C_{\alpha,Lys}$), 51.9 ($C_{\alpha,Arg}$), 48.7 ($C_{\alpha,D\text{-}Ala}$), 45.0 ($CS\underline{C}H_2$), 44.9 ($C_{\varepsilon,Lys}$), 40.7 ($CH_3SO_2$), 40.4 ($C_{\delta,Arg}$), 32.4 ($C_{\beta,Lys}$), 31.3 ($\underline{C}H_2CH_2CH_3$), 29.0–28.7 (10C, $C_{\beta,Arg}$, $(\underline{C}H_2)_{11}CH_3$), 26.7 ($C_{\delta,Lys}$), 25.1 ($C_{\gamma,Arg}$), 22.8 ($C_{\gamma,Lys}$), 22.1 ($\underline{C}H_2CH_3$), 17.8 ($C_{\beta,D\text{-}Ala}$), 13.9 ($CH_2\underline{C}H_3$). Analytical HPLC gradient 0–95% eluent II in eluent I (C8; 35 min total runtime), $t_R$ 25.2 min (>98%, $UV_{254}$). HRMS calcd for $C_{30}H_{61}N_8O_4S_2^+$ [M+H]$^+$, 677.4201; found 677.4201. Note: A large amount of oxo-byproduct was observed upon acidic cleaveage from the resin, which could be collected on reversed-phase HPLC to afford compound **S17**.

**Benzyl ((6*S*,9*S*,12*S*)-9-((1*H*-indol-3-yl)methyl)-1-amino-6-carbamoyl-1-imino-8,11-dioxo-18-thioxo-2,7,10,17-tetraazanonadecan-12-yl)carbamate (S1).** Starting from Cbz-Lys-Trp-Arg(Pbf)-resin (100 mg, estimated loading: 0.38 mmol/g) synthesized from Fmoc-Arg(Pbf)-OH, Fmoc-Trp-OH and Cbz-Lys(Fmoc)-OH by SPPS, the title compound was synthesized by on-resin thioacetylation as described in the general procedures. Global deprotection and cleavage from the resin, followed by preparative reversed-phase HPLC purification afforded the desired thioamide **S1** (1 mg, 3% based on resin loading) as a colorless fluffy material after lyophilization. $^1$H NMR (600 MHz, DMSO-$d_6$) δ 10.80 (s, 1H, $NH_{Indole}$), 9.91 (t, $J$ = 5.4 Hz, 1H, $NH_{\varepsilon,Lys}$), 8.04–7.92 (m, 2H, $NH_{\alpha,Trp}$, $NH_{\alpha,Arg}$), 7.57 (d, $J$ = 7.9 Hz, 1H, $H4_{Indole}$), 7.48 (t, $J$ = 5.8 Hz, 1H, $NH_{\delta,Arg}$), 7.43–7.28 (m, 7H, $NH_{\alpha,Trp}$, $H_{Ar,Cbz}$, $H7_{Indole}$), 7.25 (s, 1H, $CONH_{2,A}$), 7.14 (d, $J$ = 2.4 Hz, 1H, $H2_{Indole}$), 7.09 (s, 1H, $CONH_{2,B}$), 7.05 (t, $J$ = 7.5 Hz, 1H, $H6_{Indole}$), 6.96 (t, $J$ = 7.5 Hz, 1H, $H5_{Indole}$), 6.53 (s, residual $COOH_{TFA}$), 5.08–4.94 (m, 2H, $CH_{2,Cbz}$), 4.55 (td, $J$ = 8.2, 4.8 Hz, 1H, $H_{\alpha,Trp}$), 4.21 (td, $J$ = 7.5, 5.6 Hz, 1H, $H_{\alpha,Lys}$), 3.95 (td, $J$ = 8.5, 4.8 Hz, 1H, $H_{\alpha,Arg}$), 3.44–3.36 (m, 2H, $H_{\varepsilon,Lys}$, overlap with residual water), 3.15 ($m_{ABX}$, $J$ = 14.9, 4.8 Hz, 1H, $H_{\beta,Trp,A}$), 3.11–3.04 (m, 2H, $H_{\delta,Arg}$) 2.99 ($m_{ABX}$, $J$ = 14.9, 8.8 Hz, 1H, $H_{\beta,Trp,B}$), 2.37 (s, 3H, $CH_3$), 1.73–1.40 (m, 8H, $H_{\beta,Lys}$, $H_{\delta,Lys}$, $H_{\beta,Arg}$, $H_{\gamma,Arg}$), 1.29–1.20 (m, 2H, $H_{\gamma,Lys}$). $^{13}$C NMR (151 MHz, DMSO-$d_6$) δ 198.7 (C=S), 173.1 ($CO_{Arg}$), 171.9 ($CO_{Lys}$), 171.2 ($CO_{Trp}$), 157.8 (q, $J$ = 31.2 Hz, residual $CO_{TFA}$), 156.6 ($NH(CNH)NH_2$), 156.0 ($CO_{Cbz}$), 136.9 ($C1_{Ar,Cbz}$), 136.0 ($C7a_{Indole}$), 128.3 ($C3_{Ar,Cbz}$, $C5_{Ar,Cbz}$), 127.8 ($C4_{Ar,Cbz}$), 127.7 ($C2_{Ar,Cbz}$, $C6_{Ar,Cbz}$), 127.3 ($C3a_{Indole}$), 123.6 ($C2_{Indole}$), 120.8 ($C6_{Indole}$), 118.4 ($C4_{Indole}$), 118.2 ($C5_{Indole}$), 117.3 (q, $J$ = 306.1 Hz, residual $CF_{3,TFA}$), 111.2 ($C7_{Indole}$), 109.8 ($C3_{Indole}$), 65.5 ($CH_{2,Cbz}$), 54.7 ($C_{\alpha,Lys}$), 53.4 ($C_{\alpha,Trp}$), 51.9 ($C_{\alpha,Arg}$), 45.3 ($C_{\varepsilon,Lys}$), 32.8 ($C_{\delta,Arg}$), 31.5 ($C_{\beta,Lys}$), 29.2 ($C_{\beta,Arg}$),

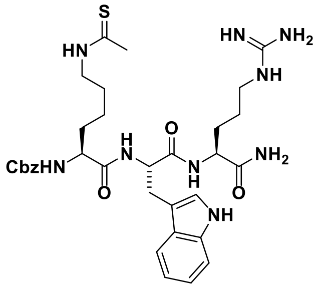

27.4 ($C_{\beta,Trp}$), 26.9 ($C_{\delta,Lys}$), 25.0 ($C_{\gamma,Arg}$), 23.1 ($C_{\gamma,Lys}$). Analytical HPLC gradient 0–95% eluent II in eluent I (C18; 35 min total runtime), $t_R$ 16.2 min (>95%, $UV_{230}$). HRMS calcd for $C_{33}H_{46}N_9O_5S^+$ $[M+H]^+$, 680.3337; found 680.3329. TA = thioacetyl.

**(5*S*,8*S*,11*S*)-8-((1*H*-indol-3-yl)methyl)-11-carbamoyl-5-(4-ethanethioamidobutyl)-3,6,9-trioxo-1-phenyl-2-oxa-4,7,10-triazatetradecan-14-oic acid (S2).**

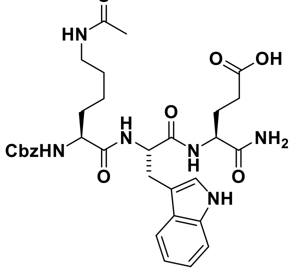

Starting from Cbz-Lys-Trp-Glu(*t*Bu)-resin (100 mg, estimated loading: 0.42 mmol/g) synthesized from Fmoc-Glu(O*t*Bu)-OH, Fmoc-Trp-OH and Cbz-Lys(Fmoc)-OH by SPPS, the title compound was synthesized by on-resin thioacetylation as described in the general procedures. Global deprotection and cleavage from the resin, followed by preparative reversed-phase HPLC purification afforded the desired thioamide **S2** (1 mg, 5% based on resin loading), as a colorless fluffy material after lyophilization. $^1$H NMR (600 MHz, DMSO-$d_6$) δ 12.06 (s, 1H, COOH), 10.80 (d, $J$ = 2.5 Hz, 1H, $NH_{Indole}$), 9.89 (t, $J$ = 5.3 Hz, 1H, $NH_{\varepsilon,Lys}$), 7.99 (d, $J$ = 7.7 Hz, 1H, $NH_{\alpha,Trp}$), 7.89 (d, $J$ = 8.0 Hz, 1H, $NH_{\alpha,Glu}$), 7.56 (d, $J$ = 7.9 Hz, 1H, $H4_{Indole}$), 7.45–7.25 (m, 7H, $NH_{\alpha,Lys}$, $H_{Ar,Cbz}$, $H7_{Indole}$), 7.19–7.12 (m, 2H, $CONH_{2,A}$, $H2_{Indole}$), 7.09–7.01 (m, 2H, $CONH_{2,B}$, $H6_{Indole}$), 6.96 (t, $J$ = 7.4 Hz, 1H, $H5_{Indole}$), 5.08–4.94 (m, 2H, $CH_{2,Cbz}$), 4.53 (td, $J$ = 8.1, 4.9 Hz, 1H, $H_{\alpha,Trp}$), 4.19 (td, $J$ = 8.2, 5.1 Hz, 1H, $H_{\alpha,Glu}$), 3.94 (td, $J$ = 8.6, 5.0 Hz, 1H, $H_{\alpha,Lys}$), 3.48–3.28 (m, 2H, $H_{\varepsilon,Lys}$, overlap with residual water), 3.16 ($m_{ABX}$, $J$ = 14.8, 5.0 Hz, 1H, $H_{\beta,Trp,A}$), 2.99 ($m_{ABX}$, $J$ = 14.8, 8.6 Hz, 1H, $H_{\beta,Trp,B}$), 2.36 (s, 3H, $CH_3$), 2.25–2.13 (m, 2H, $H_{\gamma,Glu}$), 1.96–1.89 (m, 1H, $H_{\beta,Glu,A}$), 1.79–1.70 (m, 1H, $H_{\beta,Glu,A}$), 1.63–1.39 (m, 4H, $H_{\beta,Lys}$, $H_{\delta,Lys}$), 1.33–1.13 (m, 2H, $H_{\gamma,Lys}$). $^{13}$C NMR (151 MHz, DMSO-$d_6$) δ 198.7 (C=S), 173.9 (COOH), 172.9 ($CO_{\alpha,Glu}$), 172.0 ($CO_{Lys}$), 171.2 ($CO_{Trp}$), 156.0 ($CO_{Cbz}$), 136.9 ($C1_{Ar,Cbz}$), 136.0 ($C7a_{Indole}$), 128.3 ($C3_{Ar,Cbz}$, $C5_{Ar,Cbz}$), 127.8 ($C4_{Ar,Cbz}$), 127.7 ($C2_{Ar,Cbz}$, $C6_{Ar,Cbz}$), 127.3 ($C3a_{Indole}$), 123.5 ($C2_{Indole}$), 120.8 ($C6_{Indole}$), 118.4 ($C4_{Indole}$), 118.2 ($C5_{Indole}$), 111.2 ($C7_{Indole}$), 109.9 ($C3_{Indole}$), 65.5 ($CH_{2,Cbz}$), 54.8 ($C_{\alpha,Lys}$), 53.4 ($C_{\alpha,Trp}$), 51.8 ($C_{\alpha,Glu}$), 45.3 ($C_{\varepsilon,Lys}$), 32.8 ($CH_3$), 31.5 ($C_{\beta,Lys}$), 30.1 ($C_{\gamma,Glu}$), 27.4 ($C_{\beta,Glu}$), 27.3 ($C_{\beta,Trp}$), 26.9 ($C_{\delta,Lys}$), 23.1 ($C_{\gamma,Lys}$). Analytical HPLC gradient 0–95% eluent II in eluent I (C18; 35 min total runtime), $t_R$ 17.0 min (>97%, $UV_{230}$). HRMS calcd for $C_{32}H_{40}N_6O_7S^+$ $[M+H]^+$, 675.2571; found 675.2572. TA = thioacetyl.

**Benzyl ((*S*)-1-(((*S*)-1-(((*S*)-1,6-diamino-1-oxohexan-2-yl)amino)-3-(1*H*-indol-3-yl)-1-oxopropan-2-yl)amino)-6-ethanethioamido-1-oxohexan-2-yl)carbamate (S3).**

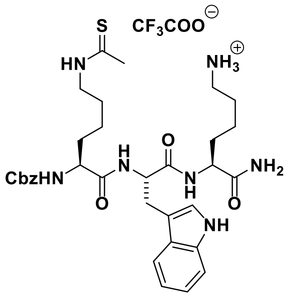

Starting from Cbz-Lys-Trp-Lys(Boc)-resin (101 mg, estimated loading: 0.42 mmol/g) synthesized from Fmoc-Lys(Boc)-OH, Fmoc-Trp-OH and Cbz-Lys(Fmoc)-OH by SPPS, the title compound was synthesized by on-resin thioacetylation as described in the general procedures. Global deprotection and cleavage from the resin, followed by preparative reversed-phase HPLC purification afforded the desired thioamide **S3** (2 mg, 7% based on resin loading) as a colorless fluffy material after lyophilization. $^1$H NMR (600 MHz, DMSO-$d_6$) δ 10.83 (d, $J$ = 2.4 Hz, 1H,

NH$_{Indole}$), 9.96 (t, *J* = 5.3 Hz, 1H, NH$_{ε,Lys(TA)}$), 8.00 (d, *J* = 7.6 Hz, 1H, NH$_{α,Trp}$), 7.93 (d, *J* = 8.1 Hz, 1H, NH$_{α,Lys(TA)}$), 7.77 (br s, 3H, $NH_3^+$), 7.57 (d, *J* = 7.9 Hz, 1H, H4$_{Indole}$), 7.53–7.17 (m, 8H, H$_{Ar,Cbz}$, H7$_{Indole}$, NH$_{α,Lys}$, CONH$_{2,A}$), 7.15 (d, *J* = 2.4 Hz, 1H, H2$_{Indole}$), 7.08–7.01 (m, 2H, H6$_{Indole}$, CONH$_{2,B}$), 6.99–6.93 (m, 1H, H5$_{Indole}$), 6.56 (s, residual COOH$_{TFA}$), 5.13–4.91 (m, 2H, CH$_{2,Cbz}$), 4.54 (td, *J* = 8.1, 5.0 Hz, 1H, H$_{α,Trp}$), 4.17 (td, *J* = 8.4, 5.1 Hz, 1H, H$_{α,Lys(TA)}$), 4.03–3.87 (m, 1H, H$_{α,Lys}$), 3.44–3.37 (m, 2H, H$_{ε,Lys(TA)}$, overlap with residual water), 3.15 (m$_{ABX}$, *J* = 14.8, 4.9 Hz, 1H, H$_{β,Trp,A}$), 2.99 (m$_{ABX}$, *J* = 14.9, 8.6 Hz, 1H, H$_{β,Trp,B}$), 2.80–2.66 (m, 2H, ), 2.37 (s, 3H, $CH_3$), 1.73–1.37 (m, 8H, H$_{β,Lys(TA)}$, H$_{δ,Lys(TA)}$, H$_{β,Lys}$, H$_{δ,Lys}$), 1.36–1.13 (m, 4H, H$_{γ,Lys(TA)}$, H$_{γ,Lys}$). $^{13}$C NMR (151 MHz, DMSO-*d*$_6$) δ 198.8 (C=S), 173.3 (CO$_{Lys(TA)}$), 172.0 (CO$_{Trp}$), 171.2 (CO$_{Lys}$), 157.9 (q, *J* = 32.0 Hz, CO$_{TFA}$), 156.0 (CO$_{Cbz}$), 136.9 (C1$_{Ar,Cbz}$), 136.0 (C7a$_{Indole}$), 128.4 (C3$_{Ar,Cbz}$, C5$_{Ar,Cbz}$), 127.8 (C4$_{Ar,Cbz}$), 127.7 (C2$_{Ar,Cbz}$, C6$_{Ar,Cbz}$), 127.3 (C3a$_{Indole}$), 123.6 (C2$_{Indole}$), 120.8 (C6$_{Indole}$), 118.4 (C4$_{Indole}$), 118.2 (C5$_{Indole}$), 117.3 (q, *J* = 302.2 Hz, CF$_{3,TFA}$), 111.2 (C7$_{Indole}$), 109.8 (C3$_{Indole}$), 65.5 (CH$_{2,Cbz}$), 54.8 (C$_{α,Lys}$), 53.4 (C$_{α,Trp}$), 52.2 (C$_{α,Lys(TA)}$), 45.3 (C$_{ε,Lys(TA)}$), 38.7 (C$_{ε,Lys}$), 32.8 ($CH_3$), 31.5 (C$_{β,Lys}$), 31.4 (C$_{β,Lys(TA)}$), 27.3 (C$_{β,Trp}$), 26.9 (C$_{δ,Lys(TA)}$), 26.6 (C$_{δ,Lys}$), 23.1 (C$_{γ,Lys(TA)}$), 22.1 (C$_{γ,Lys}$). Analytical HPLC gradient 0–95% eluent II in eluent I (C18; 35 min total runtime), $t_R$ 16.0 min (>96%, UV$_{230}$). HRMS calcd for $C_{33}H_{46}N_7O_5S^+$ [M+H]$^+$, 652.3276; found 652.3279. TA = thioacetyl.

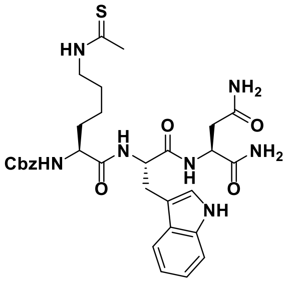

**Benzyl ((*S*)-1-(((*S*)-1-(((*S*)-1,4-diamino-1,4-dioxobutan-2-yl)amino)-3-(1*H*-indol-3-yl)-1-oxopropan-2-yl)amino)-6-ethanethioamido-1-oxohexan-2-yl)carbamate (S4)**. Starting from Cbz-Lys-Trp-Asn(Trt)-resin (80 mg, estimated loading: 0.42 mmol/g) synthesized from Fmoc-Ans(Trt)-OH, Fmoc-Trp-OH and Cbz-Lys(Fmoc)-OH by SPPS, the title compound was synthesized by on-resin thioacetylation as described in the general procedures. Global deprotection and cleavage from the resin, followed by preparative reversed-phase HPLC purification afforded the desired thioamide **S4** (2 mg, 7% based on resin loading) as a colorless fluffy material after lyophilization. $^1$H NMR (600 MHz, DMSO-*d*$_6$) δ 10.79 (d, *J* = 2.5 Hz, 1H, NH$_{Indole}$), 9.90 (t, *J* = 5.2 Hz, 1H, NH$_{ε,Lys}$), 8.08 (d, *J* = 8.0 Hz, 1H, NH$_{α,Asn}$), 8.03 (d, *J* = 7.3 Hz, 1H, NH$_{α,Trp}$), 7.56 (d, *J* = 7.9 Hz, 1H, H4$_{Indole}$), 7.46–7.21 (m, 8H, H$_{Ar,Cbz}$, H7$_{Indole}$, NH$_{α,Lys}$, CONH$_{2,Asn,α,A}$), 7.16 (d, *J* = 2.4 Hz, 1H, H2$_{Indole}$), 7.07–7.01 (m, 2H, H6$_{Indole}$, CONH$_{2,Asn,α,B}$), 6.99–6.89 (m, 2H, H5$_{Indole}$, CONH$_{2,Asn,γ,A}$), 6.84 (s, 1H, CONH$_{2,Asn,γ,A}$), 5.12–4.92 (m, 2H, CH$_{2,Cbz}$), 4.55–4.38 (m, 2H, H$_{α,Trp}$, H$_{α,Asn}$), 3.96 (td, *J* = 8.7, 4.9 Hz, 1H, H$_{α,Lys}$), 3.51–3.27 (m, 2H, H$_{ε,Lys}$), 3.15 (m$_{ABX}$, *J* = 14.9, 4.9 Hz, 1H, H$_{β,Trp,A}$), 2.97 (m$_{ABX}$, *J* = 14.9, 8.7 Hz, 1H, H$_{β,Trp,B}$), 2.48–2.41 (m, 2H, H$_{δ,Asn}$), 2.36 (s, 3H, $CH_3$), 1.67–1.39 (m, 4H, H$_{β,Lys}$, H$_{δ,Lys}$), 1.35–1.14 (m, 2H, H$_{γ,Lys}$). $^{13}$C NMR (151 MHz, DMSO-*d*$_6$) δ 198.7 (C=S), 172.7 (CO$_{Asn,α}$), 172.2 (CO$_{Lys}$), 171.7 (CO$_{Asn,γ}$), 171.1 (CO$_{Trp}$), 156.0 (CO$_{Cbz}$), 136.9 (C1$_{Ar,Cbz}$), 136.0 (C7a$_{Indole}$), 128.3 (C3$_{Ar,Cbz}$, C5$_{Ar,Cbz}$), 127.8 (C4$_{Ar,Cbz}$), 127.7 (C2$_{Ar,Cbz}$, C6$_{Ar,Cbz}$), 127.3 (C3a$_{Indole}$), 123.6 (C2$_{Indole}$), 120.8 (C6$_{Indole}$), 118.3 (C4$_{Indole}$), 118.2 (C5$_{Indole}$), 111.2 (C7$_{Indole}$), 109.8 (C3$_{Indole}$), 65.5 (CH$_{2,Cbz}$), 54.7 (C$_{α,Lys}$), 53.7 (C$_{α,Trp}$), 49.6 (C$_{α,Asn}$), 45.3 (C$_{ε,Lys}$),

36.8 ($C_{\beta,Asn}$), 32.8 ($CH_3$), 31.5 ($C_{\beta,Lys}$), 27.3 ($C_{\beta,Trp}$), 26.9 ($C_{\delta,Lys}$), 23.1 ($C_{\gamma,Lys}$). Analytical HPLC gradient 0–95% eluent II in eluent I (C18; 35 min total runtime), $t_R$ 16.6 min (>95%, $UV_{230}$). HRMS calcd for $C_{31}H_{39}N_7O_6SNa^+$ $[M+Na]^+$, 660.2575; found 660.2569. TA = thioacetyl.

**(*S*)-2-amino-*N*-((*R*)-1-(((*S*)-1-amino-5-guanidino-1-oxopentan-2-yl)amino)-1-oxopropan-2-yl)-**

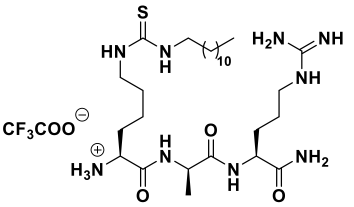

**6-(3-dodecylthioureido)hexanamide (S5).** Starting from Boc-Lys-D-Ala-Arg(Pbf)-resin (666 mg, estimated loading: 0.33 mmol/g) synthesized from Fmoc-Arg(Pbf)-OH, Fmoc-D-Ala-OH and Boc-Lys(Fmoc)-OH by SPPS, the title compound was synthesized by on-resin thiourea formation as described in the general procedures. Global deprotection and cleavage from the resin, followed by preparative reversed-phase HPLC purification afforded the desired thiourea **S5** (31 mg, 24% based on resin loading), as a colorless fluffy material after lyophilization. $^1H$ NMR (600 MHz, DMSO-$d_6$) δ 8.67 (d, $J$ = 7.5 Hz, 1H, $NH_{D\text{-}Ala}$), 8.25 (d, $J$ = 8.4 Hz, 1H, $NH_{\alpha,Arg}$), 8.08 (br s, 3H, $NH_3^+{}_{\alpha,Lys}$), 7.74–7.69 (m, 1H, $NH_{\delta,Arg}$), 7.43–7.34 (m, 3H, $NH_{\varepsilon,Lys}$, $N\underline{H}CH_2$, $CONH_{2,A}$), 7.09 (d, $J$ = 2.2 Hz, 1H, $CONH_{2,B}$), 4.48 (p, $J$ = 7.1 Hz, 1H, $H_{\alpha,D\text{-}Ala}$), 4.23 (td, $J$ = 8.4, 5.4 Hz, 1H, $H_{\alpha,Arg}$), 3.81 (t, $J$ = 6.5, 1H, $H_{\alpha,Lys}$), 3.32 (br s, 2H, $H_{\varepsilon,Lys}$, overlap with residual water), 3.12–3.04 (m, 2H, $H_{\delta,Arg}$), 1.74–1.65 (m, 3H, $H_{\beta,Arg,A}$, $H_{\beta,Lys}$), 1.55–1.39 (m, 7H, $H_{\beta,Arg,B}$, $H_{\gamma,Arg}$, $(C\underline{H}_2)_{11}CH_3$), 1.32–1.17 (m, 25H, $H_{\gamma,Lys}$, $H_{\delta,Lys}$, $(C\underline{H}_2)_{11}CH_3$, $H_{\beta,D\text{-}Ala}$), 0.85 (t, $J$ = 6.9 Hz, 3H, $CH_2C\underline{H}_3$). $^{13}C$ NMR (151 MHz, DMSO-$d_6$) δ 173.2 ($CO_{Arg}$), 171.5 ($CO_{D\text{-}Ala}$), 168.1 ($CO_{Lys}$), 158.3 (q, $J$ = 31.1 Hz, residual $CO_{TFA}$), 156.8 (NHC(=NH)$NH_2$), 118.2 (q, $J$ = 299.2 Hz, residual $CF_{3,TFA}$), 52.0 ($C_{\alpha,Lys}$), 51.7 ($C_{\alpha,Arg}$), 48.3 ($C_{\alpha,D\text{-}Ala}$), 43.2 ($C_{\varepsilon,Lys}$), 40.3 ($C_{\delta,Arg}$), 31.3 ($\underline{C}H_2CH_2CH_3$), 31.0 ($C_{\beta,Lys}$), 29.2–28.7 (10C, $C_{\beta,Arg}$, $(\underline{C}H_2)_{11}CH_3$), 26.4 ($C_{\delta,Lys}$), 25.1 ($C_{\gamma,Arg}$), 22.1 ($\underline{C}H_2CH_3$), 21.6 ($C_{\gamma,Lys}$), 18.9 ($C_{\beta,D\text{-}Ala}$), 13.9 ($CH_2\underline{C}H_3$). The peak for $C_{\varepsilon,Lys}$ was broad and of low intensity and the peak for C=S was not visible in $^{13}C$ NMR due to fast quadrupolar relaxation via the nearby $^{14}N$-nuclei. Analytical HPLC gradient 0–95% eluent II in eluent I (C18; 35 min total runtime), $t_R$ 21.9 min (>98%, $UV_{230}$). HRMS calcd for $C_{28}H_{58}N_9O_3S^+$ $[M+H]^+$, 600.4378; found 600.4373.

**(*S*)-*N*-((*R*)-1-(((*S*)-1-amino-5-guanidino-1-oxopentan-2-yl)amino)-1-oxopropan-2-yl)-2-**

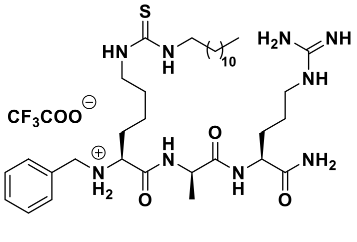

**(benzylamino)-6-(3-dodecylthioureido)hexanamide (S6).** Compound **S5** (6 mg, 0.010 mmol) was dissolved in anh. THF (1.0 mL) followed by addition of benzaldehyde (1 µL, 0.010 mmol), $NaBH(OAc)_3$ (103 mg, 0.049 mmol) and AcOH (2–3 drops) to the reaction mixture, which was stirred overnight at ambient temperature. The reaction mixture was neutralized with NaOH (2 M, aq) and directly subjected to preparative reversed-phase HPLC purification to afford the desired thiourea **S6** (1 mg, 15%) as a colorless fluffy material after lyophilization. Unreacted starting material (**S5**) could be recovered and reused. $^1H$ NMR (600 MHz, DMSO-$d_6$) δ 9.28 (br s, 1H, $NH_{\alpha,Lys,A}$), 9.20 (br s, 1H, $NH_{\alpha,Lys,B}$), 8.91 (br s, 1H, $NH_{D\text{-}Ala}$), 8.29 (d, $J$ = 8.4 Hz, 1H, $NH_{\alpha,Arg}$), 7.62 (t, $J$ = 5.8 Hz, 1H, $NH_{\delta,Arg}$), 7.50–7.30 (m, 8H, $NH_{\varepsilon,Lys}$,

N$\underline{H}$CH$_2$, CONH$_{2,A}$, H$_{Ar,Bn}$), 7.17 (d, *J* = 2.3 Hz, 1H, CONH$_{2,B}$), 4.47–4.41 (m, 1H, H$_{α,D\text{-}Ala}$), 4.25 (td, *J* = 8.3, 5.3 Hz, 1H, H$_{α,Arg}$), 4.05 (br s, 1H, CH$_{2,Bn,A}$), 3.98 (br s, 1H, CH$_{2,Bn,B}$), 3.77 (br s, 1H, H$_{α,Lys}$), 3.32 (br s, 2H, H$_{ε,Lys}$, overlap with residual water), 3.10 (q, *J* = 6.6 Hz, 2H, H$_{δ,Arg}$), 1.83–1.67 (m, 3H, H$_{β,Arg,A}$, H$_{β,Lys}$), 1.55–1.41 (m, 7H, H$_{β,Arg,B}$, H$_{γ,Arg}$, (C$\underline{H}_2$)$_{11}$CH$_3$), 1.28–1.17 (m, 25H, H$_{γ,Lys}$, H$_{δ,Lys}$, CH$_2$($\underline{CH_2}$)$_{10}$CH$_3$, H$_{β,D\text{-}Ala}$), 0.85 (t, *J* = 7.0 Hz, 3H, CH$_3$). $^{13}$C NMR (151 MHz, DMSO-*d*$_6$) δ 173.2 (CO$_{Arg}$), 171.6 (CO$_{D\text{-}Ala}$), 167.1 (CO$_{Lys}$), 158.0 (q, *J* = 30.9 Hz, CO$_{TFA}$), 156.7 (NHC(=NH)NH$_2$), 138.0 (C1$_{Ar,Bn}$), 130.3 (C4$_{Ar,Bn}$), 129.1 (C3$_{Ar,Bn}$, C5$_{Ar,Bn}$), 128.7 (C2$_{Ar,Bn}$, C6$_{Ar,Bn}$), 117.3 (q, *J* = 299.2 Hz, CF$_{3,TFA}$), 58.5 (C$_{α,Lys}$), 51.8 (C$_{α,Arg}$), 48.8 (CH$_{2,Bn}$), 48.7 (C$_{α,D\text{-}Ala}$), 43.3 (C$_{ε,Lys}$), 40.4 (C$_{δ,Arg}$), 31.3 ($\underline{C}$H$_2$CH$_2$CH$_3$), 29.2–28.5 (11C, C$_{β,Lys}$, C$_{β,Arg}$, ($\underline{C}$H$_2$)$_{11}$CH$_3$), 26.4 (C$_{δ,Lys}$), 25.1 (C$_{γ,Arg}$), 22.1 ($\underline{C}$H$_2$CH$_3$), 21.6 (C$_{γ,Lys}$), 18.1 (C$_{β,Ala}$), 13.9 (CH$_3$). The peak for C$_{ε,Lys}$ was broad and of low intensity and the peak for C=S was not visible in $^{13}$C NMR due to fast quadrupolar relaxation via the nearby $^{14}$N-nuclei. Analytical HPLC gradient 0–95% eluent II in eluent I (C8; 35 min total runtime), $t_R$ 23.1 min (>95%, UV$_{230}$). HRMS calcd for $C_{35}H_{64}N_9O_3S^+$ [M+H]$^+$, 690.4847; found 690.4853.

***N*-((6*S*,9*R*,12*S*)-1-amino-6-carbamoyl-1-imino-9-methyl-8,11-dioxo-18-thioxo-2,7,10,17,19-pentaazahentriacontan-12-yl)benzamide (S7).**

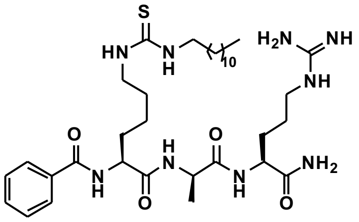

Starting from H-Lys(Teoc)-D-Ala-Arg(Pbf)-resin (168 mg, estimated loading: 0.33 mmol/g) synthesized from Fmoc-Arg(Pbf)-OH, Fmoc-D-Ala-OH and Fmoc-Lys(Teoc)-OH by SPPS, the title compound was synthesized by addition of benzoyl chloride (13 µL, 0.11 mmol) and *i*Pr$_2$NEt (59 µL, 0.34 mmol) in 1.0 mL anh. CH$_2$Cl$_2$ to the resin under agitation for 2 h. The resin was washed with DMF (3×4 mL) and CH$_2$Cl$_2$ (3×4 mL) followed by Teoc deprotection and on-resin thiourea formation as described in the general procedures. Global deprotection and cleavage from the resin, followed by preparative reversed-phase HPLC purification afforded the desired thiourea **S7** (5 mg, 13% based on resin loading), as a colorless fluffy material after lyophilization. $^1$H NMR (600 MHz, DMSO-*d*$_6$) δ 8.52 (d, *J* = 7.0 Hz, 1H, NH$_{α,Lys}$), 8.30 (d, *J* = 7.3 Hz, 1H, NH$_{D\text{-}Ala}$), 7.93 (d, *J* = 8.4 Hz, 1H, NH$_{α,Arg}$), 7.91–7.87 (m, 2H, H2$_{Ar}$, H6$_{Ar}$), 7.57–7.43 (m, 4H, H3$_{Ar}$, H4$_{Ar}$, H5$_{Ar}$, NH$_{δ,Arg}$), 7.35–7.29 (m, 3H, NH$_{ε,Lys}$, N$\underline{H}$CH$_2$, CONH$_{2,A}$), 7.11 (d, *J* = 2.1 Hz, 1H, CONH$_{2,B}$), 4.35 (q, *J* = 7.2 Hz, 1H, H$_{α,Lys}$), 4.28 (p, *J* = 7.1 Hz, 1H, H$_{α,D\text{-}Ala}$), 4.17 (td, *J* = 8.8, 5.1 Hz, 1H, H$_{α,Arg}$), 3.31 (br s, 2H, H$_{ε,Lys}$, overlap with residual water), 3.12–3.04 (m, 2H, H$_{δ,Arg}$), 1.79–1.70 (m, 3H, H$_{β,Lys}$, H$_{β,Arg,A}$), 1.57–1.36 (m, 8H, H$_{β,Arg,B}$, H$_{γ,Arg}$, (C$\underline{H}_2$)$_{11}$CH$_3$, H$_{γ,Lys,A}$), 1.34–1.19 (m, 24H, H$_{γ,Lys,B}$, H$_{δ,Lys}$, CH$_2$(C$\underline{H}_2$)$_{10}$CH$_3$, H$_{β,D\text{-}Ala}$), 0.85 (t, *J* = 7.0 Hz, 3H, CH$_2$C$\underline{H}_3$). $^{13}$C NMR (151 MHz, DMSO-*d*$_6$) δ 173.3 (CO$_{Arg}$), 172.0 (CO$_{D\text{-}Ala}$), 171.9 (CO$_{Lys}$), 166.7 ($\underline{C}$OPh), 156.6 (NHC(=NH)NH$_2$), 133.8 (C1$_{Ar}$), 131.4 (C4$_{Ar}$), 128.2 (C3$_{Ar}$, C5$_{Ar}$), 127.5 (C2$_{Ar}$, C6$_{Ar}$), 54.1 (C$_{α,Lys}$), 51.9 (C$_{α,Arg}$), 48.4 (C$_{α,D\text{-}Ala}$), 43.3 (C$_{ε,Lys}$), 40.3 (C$_{δ,Arg}$), 31.3 ($\underline{C}$H$_2$CH$_2$CH$_3$), 31.0 (C$_{β,Lys}$), 29.0–28.7 (10C, C$_{β,Arg}$, ($\underline{C}$H$_2$)$_{11}$CH$_3$), 26.4 (C$_{δ,Lys}$), 25.0 (C$_{γ,Arg}$), 23.2 (C$_{γ,Lys}$), 22.1 ($\underline{C}$H$_2$CH$_3$), 18.0 (C$_{β,D\text{-}Ala}$), 13.9 (CH$_2\underline{C}$H$_3$). The peak for C$_{ε,Lys}$ was broad and of low intensity and the peak for C=S was not visible in $^{13}$C NMR due to fast quadrupolar relaxation via the nearby $^{14}$N-nuclei. Analytical HPLC gradient 0–95% eluent II in eluent I (C8; 35 min total runtime), $t_R$ 24.5 min (>98%, $UV_{230}$). HRMS calcd for $C_{35}H_{62}N_9O_4S^+$ $[M+H]^+$, 704.4640; found 704.4638.

***N*-((6*S*,9*R*,12*S*)-1-amino-6-carbamoyl-1-imino-9-methyl-8,11-dioxo-18-thioxo-2,7,10,17,19-pentaazahentriacontan-12-yl)cyclohexanecarboxamide (S8).**

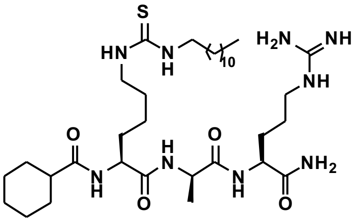

Starting from H-Lys(Teoc)-D-Ala-Arg(Pbf)-resin (168 mg, estimated loading: 0.33 mmol/g) synthesized from Fmoc-Arg(Pbf)-OH, Fmoc-D-Ala-OH and Fmoc-Lys(Teoc)-OH by SPPS, the title compound was synthesized by addition of cyclohexanecarbonyl chloride (23 µL, 0.17 mmol) and *i*$Pr_2NEt$ (59 µL, 0.34 mmol) in anh. $CH_2Cl_2$ (1.0 mL) to the resin under agitation for 2 h. The resin was washed with DMF (3×4 mL) and $CH_2Cl_2$ (3×4 mL) followed by Teoc deprotection and on-resin thiourea formation as described in the general procedures. Global deprotection and cleavage from the resin, followed by preparative reversed-phase HPLC purification afforded the desired thiourea **S8** (7 mg, 18% based on resin loading), as a colorless fluffy material after lyophilization $^1$H NMR (600 MHz, DMSO-$d_6$) δ 8.11 (d, $J$ = 7.4 Hz, 1H, $NH_{D\text{-}Ala}$), 7.94 (d, $J$ = 8.4 Hz, 1H, $NH_{\alpha,Arg}$), 7.88 (d, $J$ = 7.0 Hz, 1H, $NH_{\alpha,Lys}$), 7.58 (t, $J$ = 5.7 Hz, 1H, $NH_{\delta,Arg}$), 7.35–7.28 (m, 3H, $NH_{\varepsilon,Lys}$, $N\underline{H}(CH_2)_{11}CH_3$, $CONH_{2,A}$), 7.10 (d, $J$ = 2.1 Hz, 1H, $CONH_{2,B}$), 4.25 (p, $J$ = 7.1 Hz, 1H, $H_{\alpha,D\text{-}Ala}$), 4.16 (td, $J$ = 8.8, 5.1 Hz, 1H, $H_{\alpha,Arg}$), 4.12–4.05 (m, 1H, $H_{\alpha,Lys}$), 3.31 (br s, 2H, $H_{\varepsilon,Lys}$, overlap with residual water), 3.06 (q, $J$ = 6.8 Hz, 2H, $H_{\delta,Arg}$), 2.20 (tt, $J$ = 11.5, 3.2 Hz, 1H, $CH_{Cy}$), 1.80–1.35 (m, 16H, $H_{\beta,Arg}$, $H_{\gamma,Arg}$, $H_{\gamma,Lys,A}$, $H_{\beta,Lys}$, $CH_{2,Cy}$, $(C\underline{H}_2)_{11}CH_3$), 1.35–1.10 (m, 30H, $H_{\delta,Lys,B}$, $(C\underline{H}_2)_{11}CH_3$, $CH_{2,Cy}$, $H_{\beta,D\text{-}Ala}$), 0.85 (t, $J$ = 6.9 Hz, 3H, $CH_2C\underline{H}_3$). $^{13}$C NMR (151 MHz, DMSO-$d_6$) δ 175.7 ($\underline{C}OC_6H_{11}$), 173.3 ($CO_{Arg}$), 172.0 ($CO_{D\text{-}Ala}$), 171.9 ($CO_{Lys}$), 158.3 (q, $J$ = 31.2 Hz, residual $CO_{TFA}$), 156.7 ($NHC(=NH)NH_2$), 117.1 (q, $J$ = 299.2 Hz, residual $CF_{3,TFA}$), 53.0 ($C_{\alpha,Lys}$), 52.0 ($C_{\alpha,Arg}$), 48.2 ($C_{\alpha,D\text{-}Ala}$), 43.5 ($C1_{Cy}$), 43.4 ($C_{\varepsilon,Lys}$), 40.3 ($C_{\delta,Arg}$), 31.3 ($\underline{C}H_2CH_2CH_3$), 31.2 ($C_{\beta,Lys}$), 29.4 ($C2_{Cy}$, $C6_{Cy}$), 29.0–28.7 (10C, $C_{\beta,Arg}$, $(\underline{C}H_2)_{11}CH_3$), 26.4 ($C_{\delta,Lys}$), 25.4 ($C3_{Cy}$), 25.3 ($C5_{Cy}$), 25.2 ($C_{\gamma,Arg}$), 25.1 ($C4_{Cy}$), 22.8 ($C_{\gamma,Lys}$), 22.1 ($\underline{C}H_2CH_3$), 18.0 ($C_{\beta,D\text{-}Ala}$), 13.9 ($CH_2\underline{C}H_3$). The peak for C=S was not visible in $^{13}$C NMR due to fast quadrupolar relaxation via the nearby $^{14}$N-nuclei. Analytical HPLC gradient 0–95% eluent II in eluent I (C18; 35 min total runtime), $t_R$ 24.6 min (>98%, $UV_{230}$). HRMS calcd for $C_{35}H_{68}N_9O_4S^+$ $[M+H]^+$, 710.5109; found 710.5106. Cy = cyclohexane.

***N*-((6*S*,9*R*,12*S*)-1-amino-6-carbamoyl-1-imino-9-methyl-8,11-dioxo-18-thioxo-2,7,10,17,19-pentaazahentriacontan-12-yl)nonanamide (S9).**

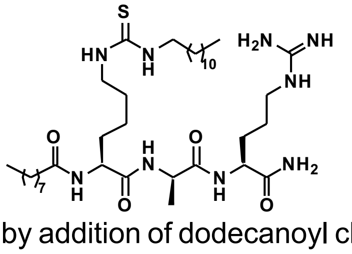

Starting from H-Lys(Teoc)-D-Ala-Arg(Pbf)-resin (168 mg, estimated loading: 0.33 mmol/g) synthesized from Fmoc-Arg(Pbf)-OH, Fmoc-D-Ala-OH and Fmoc-Lys(Teoc)-OH by SPPS, the title compound was synthesized by addition of dodecanoyl chloride (40 µL, 0.17 mmol) and *i*$Pr_2NEt$ (59 µL, 0.34 mmol) in anh. $CH_2Cl_2$ (1.0 mL) to the resin under agitation for 2 h. The resin was washed with DMF (3×4 mL) and $CH_2Cl_2$ (3×4 mL) followed by Teoc deprotection and on-resin thiourea formation as described in the general procedures. Global deprotection and cleavage from the resin, followed by preparative reversed-phase HPLC purification afforded the desired thiourea **S9** (5 mg, 12% based on resin loading), as a colorless fluffy material after lyophilization. $^1$H NMR (600 MHz, DMSO-$d_6$) δ 8.18 (d, $J$ = 7.4 Hz, 1H, $NH_{D\text{-}Ala}$), 7.99 (d, $J$ = 7.0 Hz, 1H, $NH_{\alpha,Lys}$), 7.92 (d, $J$ = 8.4 Hz, 1H, $NH_{\alpha,Arg}$), 7.51 (t, $J$ = 5.7 Hz, 1H, $NH_{\delta,Arg}$), 7.36–7.27 (m, 3H, $NH_{\varepsilon,Lys}$, $N\underline{H}(CH_2)_{11}CH_3$, $CONH_{2,A}$), 7.09 (d, $J$ = 2.1 Hz, 1H, $CONH_{2,B}$), 4.25 (p, $J$ = 7.1 Hz, 1H, $H_{\alpha,D\text{-}Ala}$), 4.20–4.08 (m, 2H, $H_{\alpha,Arg}$, $H_{\alpha,Lys}$), 3.31 (br s, 2H, $H_{\varepsilon,Lys}$, overlap with residual water), 3.09–3.02 (m, 2H, $H_{\delta,Arg}$), 2.18–2.05 (m, 2H, $CH_2CO$), 1.78–1.70 (m, 1H, $H_{\beta,Arg,A}$), 1.64–1.57 (m, 1H, $H_{\beta,Lys,A}$), 1.56–1.36 (m, 10H, $H_{\beta,Arg,B}$, $H_{\gamma,Arg}$, $H_{\beta,Lys,B}$, $H_{\gamma,Lys,A}$, $(C\underline{H}_2)_{11}CH_3$, $C\underline{H}_2CH_3CO$), 1.32–1.17 (m, 35H, $H_{\gamma,Lys}$, $H_{\delta,Lys}$, $(C\underline{H}_2)_{11}CH_3$, $COCH_2(C\underline{H}_2)_6CH_3$, $H_{\beta,D\text{-}Ala}$), 0.85 (t, $J$ = 6.9 Hz, 6H, $CH_2C\underline{H}_{3,A}$, $CH_2C\underline{H}_{3,B}$). $^{13}$C NMR (151 MHz, DMSO-$d_6$) δ 173.3 ($CO_{Arg}$), 172.7 ($\underline{C}OCH_2$), 172.0 ($CO_{D\text{-}Ala}$), 171.9 ($CO_{Lys}$), 158.0 (q, $J$ = 31.0 Hz, residual $CO_{TFA}$), 156.6 (NHC(=NH)$NH_2$), 117.3 (q, $J$ = 300.5 Hz, residual $CF_{3,TFA}$), 53.1 ($C_{\alpha,Lys}$), 51.9 ($C_{\alpha,Arg}$), 48.3 ($C_{\alpha,D\text{-}Ala}$), 43.4 ($C_{\varepsilon,Lys}$), 40.3 ($C_{\delta,Arg}$), 35.0 ($\underline{C}H_2CH_2CO$), 31.28 ($\underline{C}H_2CH_2CH_3$), 31.25 ($C_{\beta,Lys}$), 29.0–28.6 (16C $C_{\beta,Arg}$, $(\underline{C}H_2)_6CH_3$, $(\underline{C}H_2)_{11}CH_3$), 26.4 ($C_{\delta,Lys}$), 25.2 ($C_{\gamma,Arg}$), 25.1 ($\underline{C}H_2CH_{3,A}$), 22.8 ($C_{\gamma,Lys}$), 22.1 ($\underline{C}H_2CH_{3,B}$), 18.0 ($C_{\beta,D\text{-}Ala}$), 13.94 ($CH_2\underline{C}H_{3,A}$), 13.93 ($CH_2\underline{C}H_{3,B}$). The peak for $C_{\varepsilon,Lys}$ was broad and of low intensity and the peak for C=S was not visible in $^{13}$C NMR due to fast quadrupolar relaxation via the nearby $^{14}$N-nuclei. Analytical HPLC gradient 0–95% eluent II in eluent I (C8; 35 min total runtime), $t_R$ 27.3 min (>98%, $UV_{230}$). HRMS calcd for $C_{37}H_{73}N_9O_4S^+$ $[M+H]^+$, 740.5579; found 740.5591.

**(*S*)-*N*-((*R*)-1-(((*S*)-1-amino-5-guanidino-1-oxopentan-2-yl)amino)-1-oxopropan-2-yl)-6-(3-dodecylthioureido)-2-(phenylsulfonamido)hexanamide (S10).**

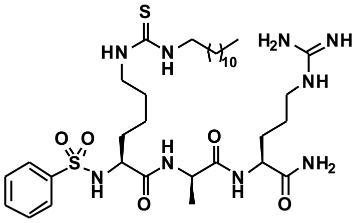

Starting from H-Lys(Teoc)-D-Ala-Arg(Pbf)-resin (168 mg, estimated loading: 0.33 mmol/g) synthesized from Fmoc-Arg(Pbf)-OH, Fmoc-D-Ala-OH and Fmoc-Lys(Teoc)-OH by SPPS, the title compound was synthesized by addition of benzene sulfonyl chloride (22 µL, 0.17 mmol) and $i$Pr$_2$NEt (59 µL, 0.34 mmol) in 1.0 mL anh. DMF to the resin under overnight agitation. The resin was washed with DMF (3×4 mL) and $CH_2Cl_2$ (3×4 mL) followed by Teoc deprotection and on-resin thiourea formation as described in the general procedures. Global deprotection and cleavage from the resin, followed by preparative reversed-phase HPLC purification afforded the desired thiourea **S10** (3 mg, 8% based on resin loading), as a colorless fluffy material after lyophilization. $^1$H NMR (600 MHz, DMSO-$d_6$) δ 8.04 (d, $J$ = 7.3 Hz, 1H, $NH_{D\text{-}Ala}$), 7.97 (d, $J$ = 8.3 Hz, 1H, $NH_{\alpha,Arg}$), 7.95 (d, $J$ = 8.3 Hz, 1H, $NH_{\alpha,Lys}$), 7.78–7.73 (m, 2H, $H2_{Ar}$, $H6_{Ar}$), 7.61–7.55 (m, 1H, $H4_{Ar}$), 7.54–7.47 (m, 3H, $H3_{Ar}$, $H5_{Ar}$, $NH_{\delta,Arg}$), 7.38 (d, $J$ = 2.3 Hz, 1H, $CONH_{2,A}$), 7.29 (br s, 1H, $N\underline{H}CH_2$), 7.22 (br s, 1H, $NH_{\varepsilon,Lys}$), 7.14 (d, $J$ = 2.2 Hz, 1H, $CONH_{2,B}$), 4.23 (td, $J$ = 8.5, 5.2 Hz, 1H, $H_{\alpha,Arg}$), 4.16 (p, $J$ = 7.2 Hz, 1H, $H_{\alpha,D\text{-}Ala}$), 3.70 (td, $J$ = 8.3, 5.6 Hz, 1H, $H_{\alpha,Lys}$), 3.19 (br s, 2H, $H_{\varepsilon,Lys}$), 3.12–3.04 (m, 2H, $H_{\delta,Arg}$), 1.76–1.67 (m, 1H, $H_{\beta,Arg,A}$), 1.54–1.35 (m, 7H, $H_{\beta,Arg,B}$, $H_{\gamma,Arg}$, $H_{\beta,Lys}$, $(CH_2)_{11}CH_3$), 1.34–0.99 (m, 27H, $H_{\gamma,Lys}$, $H_{\delta,Lys}$, $(C\underline{H}_2)_{11}CH_3$, $H_{\beta,D\text{-}Ala}$), 0.85 (t, $J$ = 6.9 Hz, 3H, $CH_2C\underline{H}_3$).

$^{13}C$ NMR (151 MHz, DMSO-$d_6$) δ 173.2 ($CO_{Arg}$), 171.7 ($CO_{D\text{-}Ala}$), 170.5 ($CO_{Lys}$), 156.6 (NHC(=NH)$NH_2$), 140.9 ($C1_{Ar}$), 132.4 ($C4_{Ar}$), 129.0 ($C3_{Ar}$, $C5_{Ar}$), 126.4 ($C2_{Ar}$, $C6_{Ar}$), 56.3 ($C_{\alpha,Lys}$), 51.7 ($C_{\alpha,Arg}$), 48.0 ($C_{\alpha,D\text{-}Ala}$), 43.1 ($C_{\varepsilon,Lys}$), 40.4 ($C_{\delta,Arg}$), 32.2 ($C_{\beta,Lys}$), 31.3 ($\underline{C}H_2CH_2CH_3$), 29.1–28.7 (C10, $C_{\beta,Arg}$, $(\underline{C}H_2)_{11}CH_3$), 26.4 ($C_{\delta,Lys}$), 25.1 ($C_{\gamma,Arg}$), 22.4 ($C_{\gamma,Lys}$), 22.1 ($\underline{C}H_2CH_3$), 18.6 ($C_{\beta,D\text{-}Ala}$), 14.0 ($CH_2\underline{C}H_3$). The peak for $C_{\varepsilon,Lys}$ was broad and of low intensity and the peak for C=S was not visible in $^{13}C$ NMR due to fast quadrupolar relaxation via the nearby $^{14}N$-nuclei. Analytical HPLC gradient 0–95% eluent II in eluent I (C8; 35 min total runtime), $t_R$ 24.9 min (>98%, $UV_{230}$). HRMS calcd for $C_{34}H_{62}N_9O_5S_2^+$ $[M+H]^+$, 740.4310; found 740.4302.

**(*S*)-*N*-((*R*)-1-(((*S*)-1-amino-5-guanidino-1-oxopentan-2-yl)amino)-1-oxopropan-2-yl)-2-(methylsulfonamido)-6-(3-octylthioureido)hexanamide (S11).**

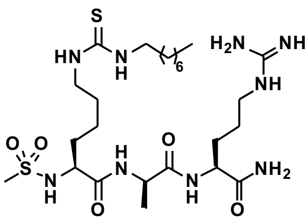

Starting from H-Lys(Teoc)-D-Ala-Arg(Pbf)-resin (168 mg, estimated loading: 0.33 mmol/g) synthesized from Fmoc-Arg(Pbf)-OH, Fmoc-D-Ala-OH and Fmoc-Lys(Teoc)-OH by SPPS, the title compound was synthesized by addition of methanesulfonyl chloride (13 µL, 0.17 mmol) and *i*$Pr_2NEt$ (59 µL, 0.34 mmol) in anh. DMF (2.0 mL) to the resin under overnight agitation. The resin was washed with DMF (3×4 mL) and $CH_2Cl_2$ (3×4 mL) followed by Teoc deprotection and on-resin thiourea formation as described in the general procedures using octylamine instead of dodecylamine. Global deprotection and cleavage from the resin, followed by preparative reversed-phase HPLC purification afforded the desired thiourea **S11** (6 mg, 16% based on resin loading) as a colorless fluffy material after lyophilization. $^1H$ NMR (600 MHz, DMSO-$d_6$) δ 8.37 (d, $J$ = 6.5 Hz, 1H, $NH_{D\text{-}Ala}$), 8.10 (d, $J$ = 8.3 Hz, 1H, $NH_{\alpha,Arg}$), 7.56 (t, $J$ = 5.8 Hz, $NH_{\delta,Arg}$), 7.36 (d, $J$ = 9.0 Hz, 1H, $NH_{\alpha,Lys}$), 7.32 (br s, 2H, $N\underline{H}CH_2$, $NH_{\varepsilon,Lys}$), 7.27 (d, $J$ = 2.2 Hz, 1H, $CONH_{2,A}$), 7.12 (d, $J$ = 2.2 Hz, 1H, $CONH_{2,B}$), 4.24 (p, $J$ = 7.0 Hz, 1H, $H_{\alpha,D\text{-}Ala}$), 4.15 (td, $J$ = 8.4, 5.0 Hz, 1H, $H_{\alpha,Arg}$), 3.83 (td, $J$ = 8.8, 5.5 Hz, 1H, $H_{\alpha,Lys}$), 3.31 (br s, 2H, $H_{\varepsilon,Lys}$, overlap with residual water), 3.12–3.05 (m, 2H, $H_{\delta,Arg}$), 2.79 (s, 3H, $CH_3SO_2$), 1.79–1.70 (m, 1H, $H_{\beta,Arg,A}$), 1.61–1.54 (m, 1H, $H_{\beta,Lys,A}$), 1.54–1.40 (m, 9H, $H_{\beta,Arg,B}$, $H_{\gamma,Arg}$, $H_{\beta,Lys,B}$, $H_{\gamma,Lys,A}$ $(C\underline{H}_2)_7CH_3$,), 1.38–1.32 (m, 1H, $H_{\gamma,Lys,B}$), 1.30–1.25 (m, 15H, $H_{\delta,Lys}$, $(C\underline{H}_2)_7CH_3$, $H_{\beta,D\text{-}Ala}$), 0.85 (t, $J$ = 7.0 Hz, 3H, $CH_2C\underline{H}_3$). $^{13}C$ NMR (151 MHz, DMSO-$d_6$) δ 173.2 ($CO_{Arg}$), 172.3 ($CO_{D\text{-}Ala}$), 171.7 ($CO_{Lys}$), 158.2 (q, $J$ = 32.2 Hz, residual $CO_{TFA}$), 156.7 (NHC(=NH)$NH_2$), 116.9 (q, $J$ = 298.2 Hz, residual $CF_{3,TFA}$), 56.0 ($C_{\alpha,Lys}$), 51.9 ($C_{\alpha,Arg}$), 48.7 ($C_{\alpha,D\text{-}Ala}$), 43.3 ($C_{\varepsilon,Lys}$), 40.7 ($CH_3SO_2$), 40.4 ($C_{\delta,Arg}$), 32.5 ($C_{\beta,Lys}$), 31.2 ($\underline{C}H_2CH_2CH_3$), 28.73 ($(\underline{C}H_2)_7CH_3$), 28.69 ($(\underline{C}H_2)_7CH_3$), 28.4 ($C_{\beta,Arg}$), 26.4 ($C_{\delta,Lys}$), 25.1 ($C_{\gamma,Arg}$), 22.7 ($C_{\gamma,Lys}$), 22.1 ($\underline{C}H_2CH_3$), 17.8 ($C_{\beta,D\text{-}Ala}$), 13.9 ($CH_3$). The peak for $C_{\varepsilon,Lys}$ was broad and of low intensity and the peak for C=S was not visible in $^{13}C$ NMR due to fast quadrupolar relaxation via the nearby $^{14}N$-nuclei. Analytical HPLC gradient 0–95% eluent II in eluent I (C18; 35 min total runtime), $t_R$ 17.9 min (>98%, $UV_{230}$). HRMS calcd for $C_{25}H_{52}N_9O_5S_2^+$ $[M+H]^+$, 622.3527; found 622.3524.

**(*S*)-*N*-((*R*)-1-(((*S*)-1-amino-5-guanidino-1-oxopentan-2-yl)amino)-1-oxopropan-2-yl)-6-(3-butylthioureido)-2-(methylsulfonamido)hexanamide (S12).**

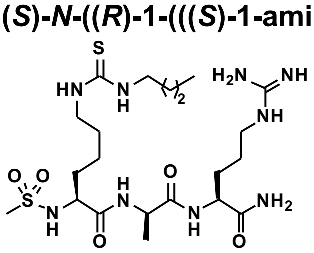

Starting from H-Lys(Teoc)-D-Ala-Arg(Pbf)-resin (168 mg, estimated loading: 0.33 mmol/g) synthesized from Fmoc-Arg(Pbf)-OH, Fmoc-D-Ala-OH and Fmoc-Lys(Teoc)-OH by SPPS, the title compound was synthesized by addition of methanesulfonyl chloride (13 µL, 0.17 mmol) and *i*$Pr_2NEt$ (59 µL, 0.34 mmol) in anh. DMF (2.0 mL) to the resin under overnight agitation. Washing with DMF (3×4 mL) and $CH_2Cl_2$ (3×4 mL) was followed by Teoc deprotection and on-resin thiourea formation as described in the general procedures using butylamine instead of dodecylamine. Global deprotection and cleavage from the resin, followed by preparative reversed-phase HPLC purification afforded the desired thiourea **S12** (2 mg, 6% based on resin loading) as a colorless fluffy material after lyophilization. $^1$H NMR (600 MHz, DMSO-$d_6$) δ 8.37 (d, *J* = 6.5 Hz, 1H, $NH_{D\text{-}Ala}$), 8.10 (d, *J* = 8.2 Hz, 1H, $NH_{\alpha,Arg}$), 7.52 (t, *J* = 5.7 Hz, $NH_{\delta,Arg}$), 7.36 (d, *J* = 8.9 Hz, 1H, $NH_{\alpha,Lys}$), 7.32 (br s, 2H, N$\underline{H}$$CH_2$, $NH_{\varepsilon,Lys}$), 7.27 (d, *J* = 2.2 Hz, 1H, $CONH_{2,A}$), 7.12 (d, *J* = 2.2 Hz, 1H, $CONH_{2,B}$), 4.24 (p, *J* = 7.0 Hz, 1H, $H_{\alpha,D\text{-}Ala}$), 4.15 (td, *J* = 8.5, 5.0 Hz, 1H, $H_{\alpha,Arg}$), 3.83 (td, *J* = 8.8, 5.5 Hz, 1H, $H_{\alpha,Lys}$), 3.31 (br s, 2H, $H_{\varepsilon,Lys}$, overlap with residual water), 3.12–3.04 (m, 2H, $H_{\delta,Arg}$), 2.80 (s, 3H, $CH_3SO_2$), 1.79–1.70 (m, 1H, $H_{\beta,Arg,A}$), 1.62–1.32 (m, 11H, $H_{\beta,Arg,B}$, $H_{\beta,Lys}$, $H_{\gamma,Arg}$, $(C\underline{H}_2)_3CH_3$, $H_{\gamma,Lys}$), 1.31–1.23 (m, 4H, $H_{\delta,Lys}$, $(C\underline{H}_2)_3CH_3$), 1.22 (d, *J* = 7.0 Hz, 3H, $H_{\beta,D\text{-}Ala}$), 0.88 (t, *J* = 7.4 Hz, 3H, $CH_2C\underline{H}_3$). $^{13}$C NMR (151 MHz, DMSO-$d_6$) δ 173.2 ($CO_{Arg}$), 172.2 ($CO_{D\text{-}Ala}$), 171.6 ($CO_{Lys}$), 156.7 (NHC(=NH)$NH_2$), 56.0 ($C_{\alpha,Lys}$), 51.8 ($C_{\alpha,Arg}$), 48.7 ($C_{\alpha,D\text{-}Ala}$), 43.3 ($C_{\varepsilon,Lys}$), 40.7 ($CH_3SO_2$), 40.3 ($C_{\delta,Arg}$), 32.5 ($C_{\beta,Lys}$), 30.9 ($\underline{C}H_2CH_2CH_3$), 28.7 ($C_{\delta,Lys}$), 28.39 (NH$\underline{C}H_2$), 28.38 ($C_{\beta,Arg}$), 25.1 ($C_{\gamma,Arg}$), 22.7 ($C_{\gamma,Lys}$), 19.5 ($\underline{C}H_2CH_3$), 17.8 ($C_{\beta,D\text{-}Ala}$), 13.7 ($CH_2\underline{C}H_3$). The peak for $C_{\varepsilon,Lys}$ was broad and of low intensity and the peak for C=S was not visible in $^{13}$C NMR due to fast quadrupolar relaxation via the nearby $^{14}$N-nuclei. Analytical HPLC gradient 0–95% eluent II in eluent I (C18; 35 min total runtime), $t_R$ 12.4 min (>98%, $UV_{230}$). HRMS calcd for $C_{21}H_{44}N_9O_5S_2^+$ [M+H]$^+$, 566.2901; found 566.2906.

**(*S*)-*N*-((*R*)-1-(((*S*)-1-amino-5-guanidino-1-oxopentan-2-yl)amino)-1-oxopropan-2-yl)-6-butanethioamido-2-(methylsulfonamido)hexanamide (S13).**

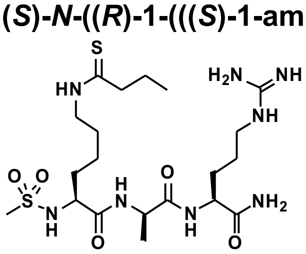

Starting from H-Lys(thiobutyryl)-D-Ala-Arg(Pbf)-resin (228 mg, estimated loading: 0.47 mmol/g) synthesized from Fmoc-Arg(Pbf)-OH, Fmoc-D-Ala-OH and Fmoc-Lys(thiobuturyl)-OH (**S33**) by SPPS, the title compound was synthesized by addition of methanesulfonyl chloride (25 µL, 0.32 mmmol) and *i*$Pr_2NEt$ (111 µL, 0.64 mmol) in anh. $CH_2Cl_2$ (2.0 mL) to the resin under agitation for 30 min. The resin was washed with DMF (3×4 mL) and $CH_2Cl_2$ (3×4 mL) followed by global deprotection and cleavage from the resin, and subsequent preparative reversed-phase HPLC purification to afford the desired thioamide **S13** (1 mg, 2% based on resin loading) as a colorless fluffy material after lyophilization. $^1$H NMR (600 MHz, DMSO-$d_6$) δ 9.89 (t, *J* = 5.4 Hz, 1H, $NH_{\varepsilon,Lys}$), 8.37 (d, *J* = 6.5 Hz, 1H, $NH_{D\text{-}Ala}$), 8.10 (d, *J*

= 8.3 Hz, 1H, $NH_{\alpha,Arg}$), 7.48 (t, $J$ = 5.9 Hz, 1H, $NH_{\delta,Arg}$), 7.36 (d, $J$ = 8.9 Hz, 1H, $NH_{\alpha,Lys}$), 7.27 (d, $J$ = 2.3 Hz, 1H, $CONH_{2,A}$), 7.13 (d, $J$ = 2.2 Hz, 1H, $CONH_{2,B}$), 4.25 (p, $J$ = 7.0 Hz, 1H, $H_{\alpha,D\text{-}Ala}$), 4.15 (td, $J$ = 8.4, 5.0 Hz, 1H, $H_{\alpha,Arg}$), 3.84 (td, $J$ = 8.8, 4.3 Hz, 1H, $H_{\alpha,Lys}$), 3.50–3.40 (m, 2H, $H_{\varepsilon,Lys}$, overlap with residual water), 3.11–3.04 (m, 2H, $H_{\delta,Arg}$), 2.80 (s, 3H, $CH_3SO_2$), 2.50–2.46 (m, 2H, $CSC\underline{H}_2$, overlap with solvent peak), 1.79–1.71 (m, 1H, $H_{\beta,Arg,A}$), 1.70–1.25 (m, 11H, $H_{\beta,Arg,B}$, $H_{\gamma,Arg}$, $H_{\beta,Lys,}$ $H_{\gamma,Lys}$,$H_{\delta,Lys}$, $C\underline{H}_2CH_3$ ), 1.22 (d, $J$ = 7.0 Hz, 3H, $H_{\beta,D\text{-}Ala}$), 0.85 (t, $J$ = 7.4 Hz, 3H, $CH_2C\underline{H}_3$). $^{13}C$ NMR (151 MHz, DMSO-$d_6$) δ 203.4 (C=S), 173.2 ($CO_{Arg}$), 172.3 ($CO_{D\text{-}Ala}$), 171.6 ($CO_{Lys}$), 156.6 ($NHC(=NH)NH_2$), 55.9 ($C_{\alpha,Lys}$), 51.8 ($C_{\alpha,Arg}$), 48.7 ($C_{\alpha,D\text{-}Ala}$), 46.9 ($CS\underline{C}H_2$), 44.9 ($C_{\varepsilon,Lys}$), 40.7 ($CH_3SO_2$), 40.4 ($C_{\delta,Arg}$), 32.4 ($C_{\beta,Lys}$), 28.7 ($C_{\beta,Arg}$), 26.7 ($C_{\delta,Lys}$), 25.1 ($C_{\gamma,Arg}$), 22.8 ($C_{\gamma,Lys}$), 22.3 ($\underline{C}H_2CH_3$), 17.8 ($C_{\beta,D\text{-}Ala}$), 13.1 ($CH_2\underline{C}H_3$). Analytical HPLC gradient 0–95% eluent II in eluent I (C8; 35 min total runtime), $t_R$ 14.4 min (>98%, $UV_{280}$). HRMS calcd for $C_{20}H_{41}N_8O_5S_2^+$ $[M+H]^+$, 537.2636; found 537.2634.

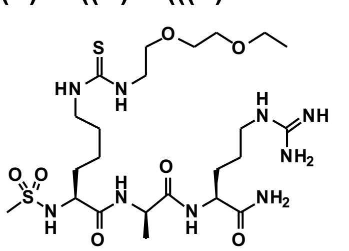

**(*S*)-*N*-((*R*)-1-(((*S*)-1-amino-5-guanidino-1-oxopentan-2-yl)amino)-1-oxopropan-2-yl)-16-(methylsulfonamido)-10-thioxo-3,6-dioxa-9,11-diazaheptadecan-17-amide (S14).** Starting from H-Lys(Teoc)-D-Ala-Arg(Pbf)-resin (168 mg, estimated loading: 0.33 mmol/g) synthesized from Fmoc-Arg(Pbf)-OH, Fmoc-D-Ala-OH and Fmoc-Lys(Teoc)-OH by SPPS, the title compound was synthesized by addition of methanesulfonyl chloride (13 µL, 0.17 mmol) and $i\text{Pr}_2\text{NEt}$ (59 µL, 0.34 mmol) in anh. DMF (2.0 mL) to the resin under overnight agitation. Washing with DMF (3×4 mL) and $CH_2Cl_2$ (3×4 mL) was followed by Teoc deprotection and on-resin thiourea formation as described in the general procedures using 2-(2-ethoxyethoxy)ethanamine instead of dodecylamine. Global deprotection and cleavage from the resin, followed by preparative reversed-phase HPLC purification afforded the desired thiourea **S4** (2 mg, 6% based on resin loading) as a white sticky material after lyophilization. $^1H$ NMR (600 MHz, DMSO-$d_6$) δ 8.37 (d, $J$ = 6.5 Hz, 1H, $NH_{D\text{-}Ala}$), 8.10 (d, $J$ = 8.3 Hz, 1H, $NH_{\alpha,Arg}$), 7.54 (t, $J$ = 5.8 Hz, 1H, $NH_{\delta,Arg}$), 7.50 (br s, 1H, $NH_{\varepsilon,Lys}$), 7.37–7.33 (m, 2H, $NH_{\alpha,Lys}$, $N\underline{H}(PEG)_2$), 7.27 (d, $J$ = 2.2 Hz, 1H, $CONH_{2,A}$), 7.12 (d, $J$ = 2.1 Hz, 1H, $CONH_{2,B}$), 4.24 (p, $J$ = 7.0 Hz, 1H, $H_{\alpha,D\text{-}Ala}$), 4.15 (td, $J$ = 8.4, 5.0 Hz, 1H, $H_{\alpha,Arg}$), 3.88–3.80 (m, 1H, $H_{\alpha,Lys}$), 3.53–3.45 (m, 8H, $NH(\underline{PEG})_2$), 3.43 (q, $J$ = 7.0 Hz, $OC\underline{H}_2CH_3$), 3.33 (br s, 2H, $H_{\varepsilon,Lys}$), 3.11–3.04 (m, 2H, $H_{\delta,Arg}$), 2.79 (s, 3H, $CH_3SO_2$), 1.79–1.70 (m, 1H, $H_{\beta,Arg,A}$), 1.62–1.32 (m, 8H, $H_{\beta,Arg,B}$, $H_{\gamma,Arg}$, $H_{\beta,Lys,}$ , $H_{\gamma,Lys,A}$, $H_{\delta,Lys}$), 1.30–1.19 (m, 4H, $H_{\beta,D\text{-}Ala}$, $H_{\gamma,Lys,B}$), 1.10 (t, $J$ = 7.0 Hz, 3H, $CH_2C\underline{H}_3$). $^{13}C$ NMR (151 MHz, DMSO-$d_6$) δ 173.2 ($CO_{Arg}$), 172.3 ($CO_{D\text{-}Ala}$), 171.6 ($CO_{Lys}$), 158.2 (q, $J$ = 32.0 Hz, residual $CO_{TFA}$), 156.7 ($NHC(=NH)NH_2$), 69.7 ($PEG_2$), 69.1, ($PEG_2$), 69.0 ($PEG_2$), 65.5 ($OC\underline{H}_2CH_3$), 56.0 ($C_{\alpha,Lys}$), 51.8 ($C_{\alpha,Arg}$), 48.7 ($C_{\alpha,D\text{-}Ala}$), 43.4 ($C_{\varepsilon,Lys}$), 40.7 ($CH_3SO_2$), 40.3 ($C_{\delta,Arg}$), 32.5 ($C_{\beta,Lys}$), 28.7 ($C_{\beta,Arg}$), 28.3 ($C_{\delta,Lys}$), 25.1 ($C_{\gamma,Arg}$), 22.8 ($C_{\gamma,Lys}$), 17.8 ($C_{\beta,D\text{-}Ala}$), 15.1 ($CH_2\underline{C}H_3$). The peak for $C_{\varepsilon,Lys}$ was broad and of low intensity and the peak for C=S was not visible in $^{13}C$ NMR due to fast quadrupolar relaxation via the nearby $^{14}N$-nuclei. Analytical HPLC gradient 0–95% eluent II in eluent I (C18; 35 min total runtime), $t_R$ 11.3 min (>98%, $UV_{230}$). HRMS calcd for $C_{23}H_{48}N_9O_7S_2^+$ $[M+H]^+$, 626.3113; found 626.3110. PEG = polyethylene glycol.

**(*S*)-*N*-((*S*)-1-(((*S*)-1-amino-5-guanidino-1-oxopentan-2-yl)amino)-1-oxopropan-2-yl)-19-(methylsulfonamido)-13-thioxo-3,6,9-trioxa-12,14-diazaicosan-20-amide (S15).**

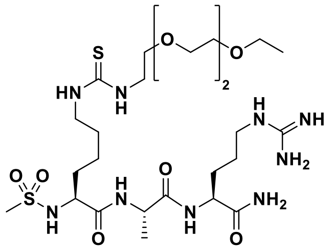

Starting from H-Lys(Teoc)-Ala-Arg(Pbf)-resin (102 mg, estimated loading: 0.49 mmol/g) synthesized from Fmoc-Arg(Pbf)-OH, Fmoc-Ala-OH and Fmoc-Lys(Teoc)-OH by SPPS, the title compound was synthesized by addition of methanesulfonyl chloride (12 µL, 0.15 mmol) and *i*$Pr_2NEt$ (52 µL, 0.30 mmol) in anh. $CH_2Cl_2$ (2.0 mL) to the resin under overnight agitation. Washing with DMF (3×4 mL) and $CH_2Cl_2$ (3×4 mL) was followed by Teoc deprotection and on-resin thiourea formation as described in the general procedures using 2-(2-(2-ethoxyethoxy) ethoxy)ethanamine instead of dodecylamine to form the benzotriazole coupling reagent. Global deprotection and cleavage from the resin, followed by preparative reversed-phase HPLC purification afforded the desired thiourea **S15** (4 mg, 10% based on resin loading) as a white sticky material after lyophilization. $^1$H NMR (600 MHz, DMSO-$d_6$) δ 8.17 (d, *J* = 7.1 Hz, 1H, $NH_{Ala}$), 7.90 (d, *J* = 8.0 Hz, 1H, $NH_{\alpha,Arg}$), 7.58–7.48 (m, 2H, $NH_{\delta,Arg}$, $NH_{\varepsilon,Lys}$), 7.39–7.30 (m, 3H, $NH_{\alpha,Lys}$, N<u>H</u>$(PEG)_3$, $CONH_{2,A}$), 7.07 (d, *J* = 2.1 Hz, 1H, $CONH_{2,B}$), 4.29 (p, *J* = 7.1 Hz, 1H, $H_{\alpha,Ala}$), 4.18 (td, *J* = 8.0, 5.6 Hz, 1H, $H_{\alpha,Arg}$), 3.78 (td, *J* = 8.7, 5.1 Hz, 1H, $H_{\alpha,Lys}$), 3.43–3.45 (m, 12H, NH(<u>PEG</u>)$_3$), 3.42 (q, *J* = 7.0 Hz, OC<u>H</u>$_2$CH$_3$, overlap with residual water), 3.33 (br s, 2H, $H_{\varepsilon,Lys}$, overlap with residual water), 3.12–3.06 (m, 2H, $H_{\delta,Arg}$), 2.85 (s, 3H, $CH_3SO_2$), 1.71–1.58 (m, 2H, $H_{\beta,Arg,A}$, $H_{\beta,Lys,A}$), 1.55–1.26 (m, 8H, $H_{\beta,Arg,B}$, $H_{\gamma,Arg}$, $H_{\beta,Lys,}$ $H_{\gamma,Lys}$, $H_{\delta,Lys,}$), 1.23 (d, *J* = 7.1 Hz, 3H, $H_{\beta,Ala}$), 1.09 (t, *J* = 7.0 Hz, 3H, CH$_2$C<u>H</u>$_3$). $^{13}$C NMR (151 MHz, DMSO-$d_6$) δ 173.1 ($CO_{Arg}$), 171.9 ($CO_{Ala}$), 171.3 ($CO_{Lys}$), 158.3 (q, *J* = 33.7 Hz, residual $CO_{TFA}$), 156.7 ($NHC(=NH)NH_2$), 116.4 (q, *J* = 295.5 Hz, residual $CF_{3,TFA}$), 69.8 ($PEG_3$), 69.7 ($PEG_3$), 69.6 ($PEG_3$), 69.2, ($PEG_3$), 69.0 ($PEG_3$), 65.5 (O<u>C</u>H$_2$CH$_3$), 56.3 ($C_{\alpha,Lys}$), 51.9 ($C_{\alpha,Arg}$), 48.3 ($C_{\alpha,Ala}$), 43.4 ($C_{\varepsilon,Lys}$), 40.7 ($CH_3SO_2$), 40.4 ($C_{\delta,Arg}$), 32.4 ($C_{\beta,Lys}$), 29.2 ($C_{\beta,Arg}$), 28.4 ($C_{\delta,Lys}$), 25.0 ($C_{\gamma,Arg}$), 22.7 ($C_{\gamma,Lys}$), 18.1 ($C_{\beta,Ala}$), 15.1 (CH$_2$<u>C</u>H$_3$). The peak for $C_{\varepsilon,Lys}$ was broad and of low intensity and the peak for C=S was not visible in $^{13}$C NMR due to fast quadrupolar relaxation via the nearby $^{14}$N-nuclei. Analytical HPLC gradient 0–95% eluent II in eluent I (C8; 35 min total runtime), $t_R$ 13.9 min (>98%, $UV_{230}$). HRMS calcd for $C_{25}H_{50}N_9NaO_8S_2^+$ $[M+Na]^+$, 692.3193; found 692.3195. PEG = polyethylene glycol.

**(*S*)-*N*-((*R*)-1-(((*S*)-1-amino-5-guanidino-1-oxopentan-2-yl)amino)-1-oxopropan-2-yl)-19-(methylsulfonamido)-13-thioxo-3,6,9-trioxa-12,14-diazaicosan-20-amide (S16).**

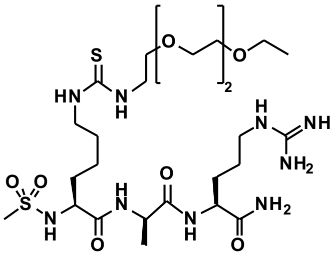

Starting from H-Lys(Teoc)-D-Ala-Arg(Pbf)-resin (168 mg, estimated loading: 0.33 mmol/g) synthesized from Fmoc-Arg(Pbf)-OH, Fmoc-D-Ala-OH and Fmoc-Lys(Teoc)-OH by SPPS, the title compound was synthesized by addition of methanesulfonyl chloride (13 µL,

0.17 mmol) and *i*$Pr_2NEt$ (59 µL, 0.34 mmol) in anh. DMF (2.0 mL) to the resin under overnight agitation. Washing with DMF (3×4 mL) and $CH_2Cl_2$ (3×4 mL) was followed by Teoc deprotection and on-resin thiourea formation as described in the general procedures using 2-(2-(2-ethoxyethoxy) ethoxy)ethanamine instead of dodecylamine to form the benzotriazole coupling reagent. Global deprotection and cleavage from the resin, followed by preparative reversed-phase HPLC purification afforded the desired thiourea **S16** (4 mg, 11% based on resin loading), as a white sticky powder after lyophilization. $^1$H NMR (600 MHz, DMSO-$d_6$) δ 8.37 (d, $J$ = 6.5 Hz, 1H, $NH_{D\text{-}Ala}$), 8.10 (d, $J$ = 8.3 Hz, 1H, $NH_{\alpha,Arg}$), 7.54–7.44 (m, 2H, $NH_{\delta,Arg}$, $NH_{\varepsilon,Lys}$), 7.37–7.33 (m, 2H, $NH_{\alpha,Lys}$, $N\underline{H}(PEG)_3$), 7.27 (d, $J$ = 2.2 Hz, 1H, $CONH_{2,A}$), 7.12 (d, $J$ = 2.2 Hz, 1H, $CONH_{2,B}$), 4.24 (p, $J$ = 7.0 Hz, 1H, $H_{\alpha,D\text{-}Ala}$), 4.15 (td, $J$ = 8.4, 5.0 Hz, 1H, $H_{\alpha,Arg}$), 3.83 (td, $J$ = 8.8, 5.5 Hz, 1H, $H_{\alpha,Lys}$), 3.53–3.46 (m, 12H, $NH(\underline{PEG})_3$), 3.43 (q, $J$ = 7.0 Hz, $OC\underline{H_2}CH_3$, overlap with residual water), 3.33 (br s, 2H, $H_{\varepsilon,Lys}$, overlap with residual water), 3.11–3.04 (m, 2H, $H_{\delta,Arg}$), 2.80 (s, 3H, $CH_3SO_2$), 1.79–1.70 (m, 1H, $H_{\beta,Arg,A}$), 1.62–1.32 (m, 8H, $H_{\beta,Arg,B}$, $H_{\gamma,Arg}$, $H_{\beta,Lys}$, $H_{\gamma,Lys,A}$, $H_{\delta,Lys}$), 1.30–1.19 (m, 4H, $H_{\gamma,Lys,B}$, $H_{\beta,D\text{-}Ala}$), 1.10 (t, $J$ = 7.0 Hz, 3H, $CH_2C\underline{H}_3$). $^{13}$C NMR (151 MHz, DMSO-$d_6$) δ 173.2 ($CO_{Arg}$), 172.3 ($CO_{D\text{-}Ala}$), 171.6 ($CO_{Lys}$), 158.1 (q, $J$ = 31.5 Hz, residual $CO_{TFA}$), 156.6 ($NHC(=NH)NH_2$), 69.8 ($PEG_3$), 69.7 ($PEG_3$), 69.6 ($PEG_3$), 69.2, ($PEG_3$), 69.0 ($PEG_3$), 65.5 ($O\underline{C}H_2CH_3$), 56.0 ($C_{\alpha,Lys}$), 51.8 ($C_{\alpha,Arg}$), 48.7 ($C_{\alpha,D\text{-}Ala}$), 43.4 ($C_{\varepsilon,Lys}$), 40.7 ($CH_3SO_2$), 40.3 ($C_{\delta,Arg}$), 32.5 ($C_{\beta,Lys}$), 28.7 ($C_{\beta,Arg}$), 28.4 ($C_{\delta,Lys}$), 25.1 ($C_{\gamma,Arg}$), 22.8 ($C_{\gamma,Lys}$), 17.8 ($C_{\beta,D\text{-}Ala}$), 15.1 ($CH_2\underline{C}H_3$). The peak for $C_{\varepsilon,Lys}$ was broad and of low intensity and the peak for C=S was not visible in $^{13}$C NMR due to fast quadrupolar relaxation via the nearby $^{14}$N-nuclei. Analytical HPLC gradient 0–95% eluent II in eluent I (C18; 35 min total runtime), $t_R$ 11.7 min (>98%, $UV_{230}$). HRMS calcd for $C_{25}H_{51}N_9O_8S_2^+$ $[M+H]^+$, 670.3375; found 670.3371. PEG = polyethylene glycol.

***N*-((*S*)-6-(((*R*)-1-(((*S*)-1-amino-5-guanidino-1-oxopentan-2-yl)amino)-1-oxopropan-2-yl)amino)-5-(methylsulfonamido)-6-oxohexyl)tetradecanamide (S17).** In the process of synthesizing compound **26-D**, a byproduct was observed upon acid cleavage from the resin, which was isolated by preparative reversed-phase HPLC purification to afford the amide **S17** (4 mg, 6% based on resin loading of **26-D**) as a colorless fluffy material after lyophilization. $^1$H NMR (600 MHz, DMSO-$d_6$) δ 8.37 (d, $J$ = 6.5 Hz, 1H, $NH_{D\text{-}Ala}$), 8.10 (d, $J$ = 8.3 Hz, 1H, $NH_{\alpha,Arg}$), 7.72 (t, $J$ = 5.6 Hz, 1H, $NH_{\varepsilon,Lys}$), 7.63 (t, $J$ = 5.7 Hz, 1H, $NH_{\delta,Arg}$), 7.33 (d, $J$ = 9.0 Hz, 1H, $NH_{\alpha,Lys}$), 7.27 (br s, 1H, $CONH_{2,A}$), 7.12 (br s, 1H, $CONH_{2,B}$), 4.24 (p, $J$ = 7.1 Hz, 1H, $H_{\alpha,D\text{-}Ala}$), 4.14 (td, $J$ = 8.4, 4.9 Hz, 1H, $H_{\alpha,Arg}$), 3.82 (td, $J$ = 8.7, 5.6 Hz, 1H, $H_{\alpha,Lys}$), 3.12–3.04 (m, 2H, $H_{\delta,Arg}$), 3.03–2.95 (m, 2H, $H_{\varepsilon,Lys}$), 2.79 (s, 3H, $CH_3SO_2$), 2.02 (t, $J$ = 7.5 Hz, 2H, $COC\underline{H_2}$), 1.79–1.71 (m, 1H, $H_{\beta,Arg,A}$), 1.67–1.15 (m, 36H, $H_{\beta,Arg,B}$, $H_{\gamma,Arg}$, $H_{\beta,Lys}$, $H_{\gamma,Lys}$, $H_{\delta,Lys}$, $H_{\beta,D\text{-}Ala}$, $(C\underline{H}_2)_{12}CH_3$), 0.85 (t, $J$ = 6.9 Hz, 3H, $CH_2C\underline{H}_3$). $^{13}$C NMR (151 MHz, DMSO-$d_6$) δ 173.3 ($CO_{Arg}$), 172.3 ($CO_{D\text{-}Ala}$), 171.9 ($\underline{C}OCH_2$), 171.7 ($CO_{Lys}$), 156.7 ($NHC(=NH)NH_2$), 55.9 ($C_{\alpha,Lys}$), 51.9 ($C_{\alpha,Arg}$), 48.7 ($C_{\alpha,D\text{-}Ala}$), 40.7 ($CH_3SO_2$), 40.4 ($C_{\delta,Arg}$), 38.2 ($C_{\varepsilon,Lys}$), 35.4 ($CO\underline{C}H_2$), 32.4 ($C_{\beta,Lys}$), 31.3 ($\underline{C}H_2CH_2CH_3$), 29.0–28.7 (10C, $C_{\beta,Arg}$, $(\underline{C}H_2)_{10}CH_3$), 25.3 ($C_{\delta,Lys}$), 25.1 ($C_{\gamma,Arg}$), 22.7 ($C_{\gamma,Lys}$), 22.1

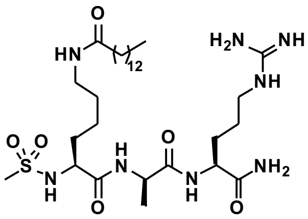

($\underline{C}H_2CH_3$), 17.8 ($C_{\beta,D\text{-}Ala}$), 13.9 ($CH_2\underline{C}H_3$). Analytical HPLC gradient 0–95% eluent II in eluent I (C8; 35 min total runtime), $t_R$ 24.1 min (>95%, $UV_{210}$). HRMS calcd for $C_{30}H_{60}N_8O_6S^+$ $[M+H]^+$, 661.4429; found 661.4428.

**(*S*)-*N*-((*R*)-1-(((*S*)-1-amino-5-guanidino-1-oxopentan-2-yl)amino)-1-oxopropan-2-yl)-6-(3-dodecylureido)-2-(methylsulfonamido)hexanamide (S18).**

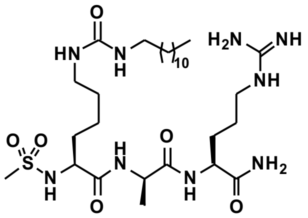

Starting from H-Lys(Teoc)-D-Ala-Arg(Pbf)-resin (322 mg, estimated loading: 0.23 mmol/g) synthesized from Fmoc-Arg(Pbf)-OH, Fmoc-D-Ala-OH and Fmoc-Lys(Teoc)-OH by SPPS, the title compound was synthesized by addition of methanesulfonyl chloride (17 µL, 0.22 mmol) and *i*$Pr_2NEt$ (76 µL, 0.44 mmol) in anh. DMF (2.0 mL) to the resin, which was agitated for 30 min. Washing with DMF (3×4 mL) and $CH_2Cl_2$ (3×4 mL) was followed by Teoc deprotection as described in the general procedures. Alongside, a solution of dodecylamine (46 mg, 0.25 mmol) and *i*$Pr_2NEt$ (130 µL, 0.75 mmol) in anh. $CH_2Cl_2$ (4 mL) were added dropwise to a solution of 4-nitrophenyl chloroformate (42 mg, 0.21 mmol) in 1.0 mL anh. $CH_2Cl_2$ and stirred at 0 ˚C for 10 min. The reaction mixture was added to the resin and agitated for 1 h followed by washing with DMF (3×4 mL) and $CH_2Cl_2$ (3×4 mL). Global deprotection and cleavage from the resin, followed by preparative reversed-phase HPLC purification afforded the desired urea **S18** (13 mg, 27% based on resin loading) as a colorless fluffy material after lyophilization. $^1$H NMR (600 MHz, DMSO-$d_6$) δ 8.36 (d, $J$ = 6.5 Hz, 1H, $NH_{D\text{-}Ala}$), 8.10 (d, $J$ = 8.2 Hz, 1H, $NH_{\alpha,Arg}$), 7.54 (t, $J$ = 5.7 Hz, $NH_{\delta,Arg}$), 7.34 (d, $J$ = 8.9 Hz, 1H, $NH_{\alpha,Lys}$), 7.27 (d, $J$ = 2.3 Hz, 1H, $CONH_{2,A}$), 7.13 (d, $J$ = 2.1 Hz, 1H, $CONH_{2,B}$), 6.86 (br s, 1H, $NH_{\varepsilon,Lys}$), 5.79–5.68 (m, 2H, $N\underline{H}C(=O)N\underline{H}$), 4.24 (p, $J$ = 7.0 Hz, 1H, $H_{\alpha,D\text{-}Ala}$), 4.15 (td, $J$ = 8.5, 5.0 Hz, 1H, $H_{\alpha,Arg}$), 3.82 (td, $J$ = 8.8, 5.5 Hz, 1H, $H_{\alpha,Lys}$), 3.11–3.04 (m, 2H, $H_{\delta,Arg}$), 2.97–2.90 (m, 4H, $H_{\varepsilon,Lys}$, $NHC\underline{H}_2$), 2.79 (s, 3H, $CH_3SO_2$), 1.79–1.70 (m, 1H, $H_{\beta,Arg,A}$), 1.61–1.10 (m, 32H, $H_{\beta,Arg,B}$, $H_{\gamma,Arg}$, $H_{\beta,Lys}$, $H_{\gamma,Lys}$, $H_{\delta,Lys}$ $H_{\beta,D\text{-}Ala}$, $(C\underline{H}_2)_{10}CH_3$), 0.85 (t, $J$ = 6.9 Hz, 3H, $CH_2C\underline{H}_3$). $^{13}$C NMR (151 MHz, DMSO-$d_6$) δ 173.2 ($CO_{Arg}$), 172.3 ($CO_{D\text{-}Ala}$), 171.7 ($CO_{Lys}$), 158.1 (NHC(=O)NH), 156.7 ($NHC(=NH)NH_2$), 56.0 ($C_{\alpha,Lys}$), 51.8 ($C_{\alpha,Arg}$), 48.7 ($C_{\alpha,D\text{-}Ala}$), 40.6 ($CH_3SO_2$), 40.3 ($C_{\delta,Arg}$), 39.1 ($C_{\varepsilon,Lys}$, $NH\underline{C}H_2$, overlap with residual solvent), 43.4 ($C_{\varepsilon,Lys}$), 32.5 ($C_{\beta,Lys}$), 31.3 ($\underline{C}H_2CH_2CH_3$), 30.0 ($(\underline{C}H_2)_{10}CH_3$), 29.6 ($CH_2(\underline{C}H_2)_{10}CH_3$), 29.0–28.7 (9C, $C_{\beta,Arg}$, $(\underline{C}H_2)_{10}CH_3$), 26.4 ($C_{\delta,Lys}$), 25.1 ($C_{\gamma,Arg}$), 22.7 ($C_{\gamma,Lys}$), 22.1 ($\underline{C}H_2CH_3$), 17.8 ($C_{\beta,D\text{-}Ala}$), 13.9 ($CH_2\underline{C}H_3$). Analytical HPLC gradient 0–95% eluent II in eluent I (C8; 35 min total runtime), $t_R$ 23.4 min (>95%, $UV_{210}$). HRMS calcd for $C_{29}H_{60}N_9O_6S^+$ $[M+H]^+$, 662.4382; found 662.4385.

**(*S*)-*N*-((*R*)-1-(((*S*)-1-amino-5-guanidino-1-oxopentan-2-yl)amino)-1-oxopropan-2-yl)-2-(methylsulfonamido)-6-(2,2,2-trifluoroacetamido)hexanamide (S19).**

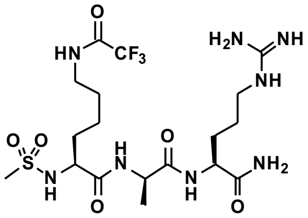

Starting from Lys(tfa)-D-Ala-Arg(Pbf)-resin (228 mg, estimated loading: 0.47 mmol/g) synthesized from Fmoc-Arg(Pbf)-OH, Fmoc-D-Ala-OH and Fmoc-Lys(tfa)-OH by SPPS, the title compound was synthesized by addition of methanesulfonyl chloride (25 µL, 0.32 mmmol) and *i*$Pr_2NEt$ (111 µL, 0.64 mmol) in anh.

$CH_2Cl_2$ (2.0 mL) to the resin under agitation for 30 min. The resin was washed with DMF (3×4 mL) and $CH_2Cl_2$ (3×4 mL) followed by global deprotection and cleavage from the resin, and subsequent preparative reversed-phase HPLC purification to afford the desired fluorinated amide **S19** (5 mg, 9% based on resin loading) as a colorless fluffy material after lyophilization. $^1$H NMR (600 MHz, DMSO-$d_6$) δ 9.41 (t, $J$ = 5.7 Hz, 1H, $NH_{ε,Lys}$), 8.38 (d, $J$ = 6.6 Hz, 1H, $NH_{D\text{-}Ala}$), 8.11 (d, $J$ = 8.3 Hz, 1H, $NH_{α,Arg}$), 7.62 (t, $J$ = 5.7 Hz, 1H, $NH_{δ,Arg}$), 7.36 (d, $J$ = 9.0 Hz, 1H, $NH_{α,Lys}$), 7.27 (d, $J$ = 2.2 Hz, 1H, $CONH_{2,A}$), 7.12 (d, $J$ = 2.2. Hz, 1H, $CONH_{2,B}$), 4.24 (p, $J$ = 7.0 Hz, 1H, $H_{α,D\text{-}Ala}$), 4.14 (td, $J$ = 8.5, 5.0 Hz, 1H, $H_{α,Arg}$), 3.84 (td, $J$ = 8.8, 5.5 Hz, 1H, $H_{α,Lys}$), 3.19–3.12 (m, 2H, $H_{ε,Lys}$), 3.11–3.04 (m, 2H, $H_{δ,Arg}$), 2.79 (s, 3H, $CH_3SO_2$), 2.07 (residual MeCN), 1.78–1.71 (m, 1H, $H_{β,Arg,A}$), 1.64–1.24 (m, 9H, $H_{β,Arg,B}$, $H_{γ,Arg}$, $H_{β,Lys}$, $H_{γ,Lys}$, $H_{δ,Lys}$), 1.21 (d, $J$ = 7.1 Hz, 3H, $H_{β,D\text{-}Ala}$). $^{13}$C NMR (151 MHz, DMSO-$d_6$) δ 173.3 ($CO_{Arg}$), 172.3 ($CO_{D\text{-}Ala}$), 171.6 ($CO_{Lys}$), 158.4 (q, $J$ = 32.3 Hz, residual $CO_{TFA}$), 156.7 (NHC(=NH)$NH_2$), 156.1 (q, $J$ = 35.7 Hz, <u>C</u>O$CF_3$), 116.9 (q, $J$ = 298.0 Hz, residual $CF_{3,TFA}$), 116.0 (q, $J$ = 288.7 Hz, CO<u>C</u>$F_3$), 55.8 ($C_{α,Lys}$), 51.9 ($C_{α,Arg}$), 48.7 ($C_{α,D\text{-}Ala}$), 40.7 ($CH_3SO_2$), 40.4 ($C_{δ,Arg}$), 39.1 ($C_{ε,Lys}$, overlap with solvent peak), 32.3 ($C_{β,Lys}$), 28.7 ($C_{β,Arg}$), 27.8 ($C_{δ,Lys}$), 25.1 ($C_{γ,Arg}$), 22.5 ($C_{γ,Lys}$), 17.8 ($C_{β,D\text{-}Ala}$). $^{19}$F NMR (376 MHz, DMSO-$d_6$) δ -74.3 (s, $CF_{3,TFA}$), -74.7 (s, $COCF_3$). Analytical HPLC gradient 0-50% eluent II in eluent I (C8; 35 min total runtime), $t_R$ 15.0 min (>98%, $UV_{210}$). HRMS calcd for $C_{18}H_{34}F_3N_8O_6S^+$ $[M+H]^+$, 547.2269; found 547.2267.

***N*⁶-(dodecylcarbamothioyl)-*N*²-(methylsulfonyl)-L-lysyl-D-alanyl-L-arginine (S20).** H-Lys(Alloc)-D-Ala-Arg(Pbf)-resin (0.86 mmol) was synthesized on 2-chlorotrityl chloride (2-CTC) resin using Fmoc-Arg(Pbf)-OH, Fmoc-D-Ala-OH and Fmoc-Lys(Alloc)-OH standard chlorotrityl SPPS procedures described previously.[10] A solution of *i*$Pr_2NEt$ (0.90 mL, 5.2 mmol) in anh. $CH_2Cl_2$ (5.0 mL) was added to the resin followed by dropwise addition of methanesulfonyl chloride (0.2 mL, 2.6 mmol in 2.0 mL anh. $CH_2Cl_2$) under magnetic stirring for 10 min. The resin was washed with $CH_2Cl_2$ (3×4.0 mL), DMF (3×4.0 mL) and $CH_2Cl_2$ (3×4.0 mL) followed by Alloc deprotection as previously described.[11] On-resin thiourea formation was performed as described in the general procedures followed by protective cleaveage from the resin with $(CF_3)_2CHOH/CH_2Cl_2$ (2×5.0 mL, 1:4, v/v, 2×30 min). Solvent was removed under a stream of nitrogen followed by preparative reversed-phase HPLC purification to afford the intermediate **S27** (30 mg, 4% based on resin loading) as a colorless fluffy material after lyophilization. ESI-MS *m/z* calcd for $C_{42}H_{75}N_8O_9S_3^+$ $[M+H]^+$, 931.48; found 931.50. To compound **S27** (3 mg, 0.004 mmol) was added TFA:TIPS (2.0 mL, 99:1, v/v) and the reaction mixture was stirred for 1 h at ambient temperature. Solvent was removed under a stream of nitrogen, followed by preparative reversed-phase HPLC purification to afford the desired thiourea **S20** (2 mg, 90%) as a colorless fluffy material after lyophilization. $^1$H NMR (600 MHz, DMSO-$d_6$) δ 12.69 (br s, 1H, $COOH_{Arg}$), 8.29 (d, $J$ = 7.6 Hz, 1H, $NH_{D\text{-}Ala}$), 8.12 (d, $J$ = 8.1 Hz, 1H, $NH_{α,Arg}$), 7.54 (t, $J$ = 5.7 Hz, 1H, $NH_{δ,Arg}$), 7.37 (d, $J$ = 8.8 Hz, 1H, $NH_{α,Lys}$), 7.31 (br s, 2H, N<u>H</u>$CH_2$, $NH_{ε,Lys}$), 4.34 (p,

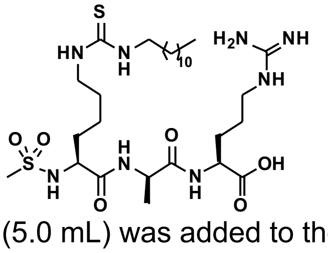

$J$ = 7.1 Hz, 1H, $H_{\alpha,D\text{-}Ala}$), 4.24–4.17 (m, 1H, $H_{\alpha,Arg}$), 3.83 (td, $J$ = 8.7, 5.6 Hz, 1H, $H_{\alpha,Lys}$), 3.31 (br s, 2H, $H_{\varepsilon,Lys}$, overlap with residual water), 3.13–3.05 (m, 2H, $H_{\delta,Arg}$), 2.82 (s, 3H, $CH_3SO_2$), 1.80–1.71 (m, 1H, $H_{\beta,Arg,A}$), 1.62–1.53 (m, 2H, $H_{\beta,Arg,B}$, $H_{\beta,Lys,A}$), 1.52–1.40 (m, 7H, $H_{\beta,Lys,B}$, $H_{\gamma,Arg}$, $(CH_2)_{11}CH_3$), 1.39–1.31 (m, 1H, $H_{\gamma,Lys,A}$), 1.30–1.18 (m, 24H, $H_{\gamma,Lys,B}$, $H_{\delta,Lys}$, $(C\underline{H}_2)_{11}CH_3$, $H_{\beta,D\text{-}Ala}$), 0.85 (t, $J$ = 7.0 Hz, 3H, $CH_2C\underline{H}_3$). $^{13}C$ NMR (151 MHz, DMSO-$d_6$) δ 173.1 ($CO_{Arg}$), 172.1 ($CO_{D\text{-}Ala}$), 171.2 ($CO_{Lys}$), 158.0 (q, $J$ = 31.2 Hz, residual $CO_{TFA}$), 156.7 ($NHC(=NH)NH_2$), 117.1 (q, $J$ = 299.0 Hz, residual $CF_{3,TFA}$), 56.0 ($C_{\alpha,Lys}$), 51.3 ($C_{\alpha,Arg}$), 48.1 ($C_{\alpha,D\text{-}Ala}$), 43.4 ($C_{\varepsilon,Lys}$), 40.5 ($CH_3SO_2$), 40.2 ($C_{\delta,Arg}$), 32.4 ($C_{\beta,Lys}$), 31.3 ($\underline{C}H_2CH_2CH_3$), 29.0–28.7 (9C, $(\underline{C}H_2)_{11}CH_3$), 28.2 ($C_{\beta,Arg}$), 26.4 ($C_{\delta,Lys}$), 25.0 ($C_{\gamma,Arg}$), 22.7 ($C_{\gamma,Lys}$), 22.1 ($\underline{C}H_2CH_3$), 18.5 ($C_{\beta,D\text{-}Ala}$), 13.9 ($CH_2\underline{C}H_3$). The peak for $C_{\varepsilon,Lys}$ was broad and of low intensity and the peak for C=S was not visible in $^{13}C$ NMR due to fast quadrupolar relaxation via the nearby $^{14}N$-nuclei. Analytical HPLC gradient 0–95% eluent II in eluent I (C18; 35 min total runtime), $t_R$ 22.7 min (>99%, $UV_{230}$). HRMS calcd for $C_{29}H_{59}N_9O_5S_2^+$ [M+H]$^+$, 679.3993; found 679.3992.

**(*S*)-6-(3-dodecylthioureido)-*N*-((*R*)-1-(((S)-5-guanidino-1-morpholino-1-oxopentan-2-yl)amino)-1-oxopropan-2-yl)-2-(methylsulfonamido)hexanamide (S21).** Compound **S30** (3 mg, 0.004 mmol) was dissolved in anh. DMF (2.0 mL) and cooled to -40 ˚C. Morpholine (3 µL, 0.04 mmol), HOAt (1 mg, 0.01 mmol) and then PyBOP (3 mg, 0.01 mmol) was added to the reaction mixture and stirred at -40 ˚C for 30 min and then for 15 min at ambient temperature. $H_2O$ (10 mL) was added and excess solvent was removed by lyophilization. TFA:TIPS (2.0 mL, 99:1, v/v) was added to the residue and stirred for 1 h at ambient temperature. Solvent was removed under a stream of nitrogen followed by preparative reversed-phase HPLC purification to afford the desired thiourea **S21** (2 mg, 94% in two steps) as a colorless fluffy material after lyophilization.

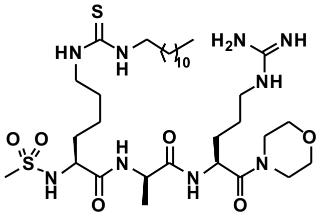

$^1H$ NMR (600 MHz, DMSO-$d_6$) δ 8.26 (d, $J$ = 7.2 Hz, 1H, $NH_{D\text{-}Ala}$), 8.24 (d, $J$ = 8.6 Hz, 1H, $NH_{\alpha,Arg}$), 7.59 (t, $J$ = 5.7 Hz, 1H, $NH_{\delta,Arg}$), 7.34 (d, $J$ = 9.0 Hz, 1H, $NH_{\alpha,Lys}$), 7.32 (br s, 2H, $N\underline{H}CH_2$, $NH_{\varepsilon,Lys}$), 4.68 (td, $J$ = 8.6, 5.2 Hz, 1H, $H_{\alpha,Arg}$), 4.31–4.23 (m, 1H, $H_{\alpha,D\text{-}Ala}$), 3.83 (td, $J$ = 8.8, 5.5 Hz, 1H, $H_{\alpha,Lys}$), 3.58–3.50 (m, 4H, $N(CH_2C\underline{H}_2)_2O$), 3.49–3.39 (m, 4H, $N(C\underline{H}_2CH_2)_2O$), 3.31 (br s, 2H, $H_{\varepsilon,Lys}$, overlap with residual water), 3.13–3.03 (m, 2H, $H_{\delta,Arg}$), 2.81 (s, 3H, $CH_3SO_2$), 1.67–1.53 (m, 2H, $H_{\beta,Arg,A}$, $H_{\beta,Lys,A}$), 1.52–1.31 (m, 10H, $H_{\beta,Arg,B}$, $H_{\beta,Lys,B}$, $H_{\gamma,Arg}$, $(C\underline{H}_2)_{11}CH_3$, $H_{\gamma,Lys}$), 1.30–1.19 (m, 23H, $H_{\delta,Lys}$, $(C\underline{H}_2)_{11}CH_3$, $H_{\beta,D\text{-}Ala}$), 0.85 (t, $J$ = 7.0 Hz, 3H, $CH_2C\underline{H}_3$). $^{13}C$ NMR (151 MHz, DMSO-$d_6$) δ 171.8 ($CO_{Arg}$), 171.3 ($CO_{D\text{-}Ala}$), 169.2 ($CO_{Lys}$),158.1 (q, $J$ = 30.9 Hz, residual $CO_{TFA}$), 156.7 ($NHC(=NH)NH_2$), 117.1 (q, $J$ = 299.0 Hz, residual $CF_{3,TFA}$), 66.2 ($N(CH_2\underline{C}H_2)_2O$), 66.1 ($N(CH_2\underline{C}H_2)_2O$), 56.0 ($C_{\alpha,Lys}$), 48.3 ($C_{\alpha,Arg}$), 47.5 ($C_{\alpha,D\text{-}Ala}$), 45.4 ($N(\underline{C}H_2CH_2)_2O$), 43.3 ($C_{\varepsilon,Lys}$), 42.0 ($N(\underline{C}H_2CH_2)_2O$), 40.5 ($CH_3SO_2$), 40.4 ($C_{\delta,Arg}$), 32.5 ($C_{\beta,Lys}$), 31.3 ($\underline{C}H_2CH_2CH_3$), 29.0–28.7 (10C, $C_{\beta,Arg}$, $(\underline{C}H_2)_{11}CH_3$), 26.4 ($C_{\delta,Lys}$), 24.6 ($C_{\gamma,Arg}$), 22.7 ($C_{\gamma,Lys}$), 22.1 ($\underline{C}H_2CH_3$), 18.3 ($C_{\beta,D\text{-}Ala}$), 13.9 ($CH_2\underline{C}H_3$). The peak for $C_{\varepsilon,Lys}$ was broad and of low intensity and the peak for C=S was not visible in $^{13}C$ NMR due to fast quadrupolar relaxation via the nearby $^{14}N$-nuclei. Analytical HPLC gradient 0–95% eluent II in eluent I (C8; 35 min total runtime), $t_R$ 24.3 min (>95%, $UV_{230}$). HRMS calcd for $C_{33}H_{66}N_9O_6S_2^+$ $[M+H]^+$, 748.4572; found 748.4565.

**(*S*)-*N*-(3-(((*S*)-1-amino-5-guanidino-1-oxopentan-2-yl)amino)-3-oxoprop-1-en-2-yl)-6-(3-dodecylthioureido)-2-(methylsulfonamido)hexanamide (S22).** 

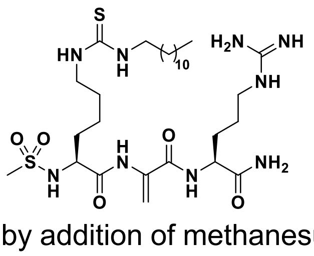

Starting from H-Lys(Teoc)-Cys(S*t*Bu)-Arg(Pbf)-resin (204 mg, estimated loading: 0.49 mmol/g) synthesized from Fmoc-Arg(Pbf)-OH, Fmoc-Cys(S*t*Bu)-OH and Fmoc-Lys(Teoc)-OH by SPPS, the title compound was synthesized by addition of methanesulfonyl chloride (23 µL, 0.30 mmol) and *i*$Pr_2NEt$ (104 µL, 0.60 mmol) in anh. $CH_2Cl_2$ (2.0 mL) to the resin under overnight agitation. The resin was washed with DMF (3×4 mL) and $CH_2Cl_2$ (3×4 mL) followed by dehydroalanine (Dha) formation as described for compound **22**. Subsequent Teoc deprotection and on-resin thiourea formation was performed as described in the general procedures. Global deprotection and cleavage from the resin, followed by preparative reversed-phase HPLC purification afforded the desired thiourea **S22** (5 mg, 5% based on resin loading), as a colorless fluffy material after lyophilization. $^1H$ NMR (600 MHz, DMSO-$d_6$) δ 9.43 (s, , 1H, $NH_{Dha}$), 8.34 (d, $J$ = 7.9 Hz, 1H, $NH_{\alpha,Arg}$), 7.67 (d, $J$ = 7.5 Hz, 1H, $NH_{\alpha,Lys}$), 7.57 (t, $J$ = 5.7 Hz, $NH_{\delta,Arg}$), 7.41–7.28 (m, 3H, N$\underline{H}$$CH_2$, $NH_{\varepsilon,Lys}$, $CONH_{2,A}$), 7.10 (br s, 1H, $CONH_{2,B}$), 6.10 (s, 1H, $H_{\beta,Dha,A}$), 5.61 (s, 1H, $H_{\beta,Dha,B}$), 4.23 (td, $J$ = 8.7, 5.2 Hz, 1H, $H_{\alpha,Arg}$), 3.78 (td, $J$ = 8.2, 4.9 Hz, 1H, $H_{\alpha,Lys}$), 3.31 (br s, 2H, $H_{\varepsilon,Lys}$, overlap with residual water), 3.13–3.06 (m, 2H, $H_{\delta,Arg}$), 2.92 (s, 3H, $CH_3SO_2$), 1.83–1.75 (m, 1H, $H_{\beta,Arg,A}$), 1.72–1.19 (m, 31H, $H_{\beta,Arg,B}$, $H_{\gamma,Arg}$, $H_{\beta,Lys}$, $H_{\gamma,Lys}$, $H_{\delta,Lys}$, $(C\underline{H}_2)_{11}CH_3$)), 0.85 (t, $J$ = 6.9 Hz, 3H, $CH_3$). $^{13}C$ NMR (151 MHz, DMSO-$d_6$) δ 173.2 ($CO_{Arg}$), 171.1 ($CO_{Lys}$), 163.7 ($CO_{Dha}$), 158.7 (q, $J$ = 31.0 Hz, residual $CO_{TFA}$), 156.7 ($NHC(=NH)NH_2$), 134.9 ($C_{\alpha,Dha}$), 104.0 ($C_{\beta,Dha}$), 57.0 ($C_{\alpha,Lys}$), 52.9 ($C_{\alpha,Arg}$), 43.3 ($C_{\varepsilon,Lys}$), 40.42 ($CH_3SO_2$), 40.37 ($C_{\delta,Arg}$), 32.0 ($C_{\beta,Lys}$), 31.3 ($\underline{C}H_2CH_2CH_3$), 29.0–28.4 (10C, $C_{\beta,Arg}$, $(\underline{C}H_2)_{11}CH_3$), 26.4 ($C_{\delta,Lys}$), 25.4 ($C_{\gamma,Arg}$), 22.7 ($C_{\gamma,Lys}$), 22.1 ($\underline{C}H_2CH_3$), 13.9 ($CH_3$). Analytical HPLC gradient 0–95% eluent II in eluent I (C8; 35 min total runtime), $t_R$ 24.5 min (>98%, $UV_{230}$). HRMS calcd for $C_{29}H_{58}N_9O_5S_2^+$ $[M+H]^+$, 676.3997; found 676.3984.

**(*S*)-*N*-(3-(((*S*)-1-amino-5-guanidino-1-oxopentan-2-yl)amino)-3-oxoprop-1-en-2-yl)-2-(methylsulfonamido)-6-tetradecanethioamidohexanamide (S23).**

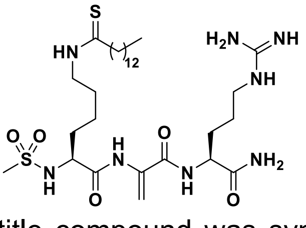

Starting from H-Lys(thiomyristoyl)-Cys(S*t*Bu)-Arg(Pbf)-resin (228 mg, estimated loading: 0.49 mmol/g) synthesized from Fmoc-Arg(Pbf)-OH, Fmoc-Cys(S*t*Bu)-OH and Fmoc-Lys(thiomyristoyl)-OH (**S34**) by SPPS, the title compound was synthesized by addition of methanesulfonyl chloride (23 µL, 0.30 mmol) and *i*$Pr_2NEt$ (104 µL, 0.60 mmol) in 2.0 mL anh. $CH_2Cl_2$ to the resin under overnight agitation. The resin was washed with DMF (3×4 mL) and $CH_2Cl_2$ (3×4 mL) followed by dehydroalanine (Dha) formation as described for compound **22.** Global deprotection, cleavage from the resin, followed by preparative reversed-phase HPLC purification afforded the desired thioamide **S23** (6 mg, 7% based on resin loading) as a colorless fluffy material after lyophilization $^1H$ NMR (600 MHz, DMSO-$d_6$) δ 9.89 (t, $J$ =

5.4 Hz, 1H, $NH_{\varepsilon,Lys}$), 9.44 (s, 1H, $NH_{Dha}$), 8.34 (d, $J$ = 7.9 Hz, 1H, $NH_{\alpha,Arg}$), 7.68 (d, $J$ = 7.5 Hz, 1H, $NH_{\alpha,Lys}$), 7.62 (t, $J$ = 5.8 Hz, 1H, $NH_{\delta,Arg}$), 7.38 (d, $J$ = 2.0 Hz, 1H, $CONH_{2,A}$), 7.10 (d, $J$ = 2.1 Hz, 1H, $CONH_{2,B}$), 6.10 (s, 1H, $H_{\beta,Dha,A}$), 5.62 (s, 1H, $H_{\beta,Dha,A}$), 4.15 (td, $J$ = 8.6, 5.2 Hz, 1H, $H_{\alpha,Arg}$), 3.94 (td, $J$ = 8.2, 4.9 Hz, 1H, $H_{\alpha,Lys}$), 3.50–3.40 (m, 2H, $H_{\varepsilon,Lys}$, overlap with residual water), 3.13–3.07 (m, 2H, $H_{\delta,Arg}$), 2.92 (s, 3H, $CH_3SO_2$), 2.50–2.46 (m, 2H, $CSC\underline{H}_2$, overlap with solvent peak), 1.84–1.75 (m, 1H, $H_{\beta,Arg,A}$), 1.74–1.18 (m, 31H, $H_{\beta,Arg,B}$, $H_{\gamma,Arg}$, $H_{\beta,Lys}$, $H_{\gamma,Lys}$, $H_{\delta,Lys}$, $(C\underline{H}_2)_{11}CH_3$ ), 0.85 (t, $J$ = 6.9 Hz, 3H, $CH_3$). $^{13}$C NMR (151 MHz, DMSO-$d_6$) δ 203.6 (C=S), 173.2 ($CO_{Arg}$), 171.1 ($CO_{Lys}$), 163.7 ($CO_{Dha}$), 156.7 ($NHC(=NH)NH_2$), 134.9 ($C_{\alpha,Dha}$), 104.0 ($C_{\beta,Dha}$), 56.9 ($C_{\alpha,Lys}$), 52.9 ($C_{\alpha,Arg}$), 45.0 ($CS\underline{C}H_2$), 44.9 ($C_{\varepsilon,Lys}$),40.44 ($CH_3SO_2$), 40.37 ($C_{\delta,Arg}$), 31.9 ($C_{\beta,Lys}$), 31.3 ($\underline{C}H_2CH_2CH_3$), 29.0–28.2 (10C, $C_{\beta,Arg}$, $(\underline{C}H_2)_{11}CH_3$), 26.8 ($C_{\delta,Lys}$), 25.4 ($C_{\gamma,Arg}$), 22.8 ($C_{\gamma,Lys}$), 22.1 ($\underline{C}H_2CH_3$), 13.9 ($CH_3$). Analytical HPLC gradient 0–95% eluent II in eluent I (C8; 35 min total runtime), $t_R$ 26.0 min (>98%, $UV_{230}$). HRMS calcd for $C_{30}H_{59}N_8O_5S_2^+$ $[M+H]^+$, 675.4044; found 675.4043.

**Fmoc-hArg(Boc)$_2$-OH (S28).** The compound was synthesized as previously described,[12] apart from

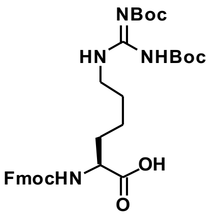

elevating the temperature to 40 °C. Isolated yield 311 mg, 57% in two steps from Fmoc-Lys(Boc)-OH. $^1$H NMR (600 MHz, $CDCl_3$) δ 11.46 (br s, 1H, COOH), 8.52 (s, 1H, $NH_{\varepsilon,hArg}$), 7.75 (d, $J$ = 7.5 Hz, 2H, $H_{Ar,Fmoc}$), 7.63–7.53 (m, 2H, $H_{Ar,Fmoc}$), 7.38 (tt, $J$ = 7.5, 0.9 Hz, 2H, $H_{Ar,Fmoc}$), 7.30 (td, $J$ = 7.5, 1.2 Hz, 2H, $H_{Ar,Fmoc}$), 5.57 (d, $J$ = 7.9 Hz, 1H, $NH_{\alpha,hArg}$), 4.59–4.32 (m, 3H, $CH_{2,Fmoc}$, $H_{\alpha,hArg}$), 4.22 (t, $J$ = 7.0 Hz, 1H, $CH_{Fmoc}$), 3.58–3.21 (m, 2H, $H_{\varepsilon,hArg}$), 2.00–1.89 (m, 1H, $H_{\beta,hArg,A}$), 1.89–1.78 (m, 1H, $H_{\beta,hArg,B}$), 1.69–1.21 (m, 22H, $H_{\gamma,hArg}$, $H_{\delta,hArg}$, $C(CH_3)_{3,A}$, $C(CH_3)_{3,B}$). $^{13}$C NMR (151 MHz, $CDCl_3$) δ 174.6 (COOH), 156.17 ($HN(\underline{C}N)NHBoc_2$ or $CO_{Fmoc}$), 156.15 ($HN(\underline{C}N)NHBoc_2$ or $CO_{Fmoc}$), 153.3 ($CO_{Boc}$), 144.0 ($C_{Ar,Fmoc}$), 143.9 ($C_{Ar,Fmoc}$), 141.4 ($2C_{Ar,Fmoc}$), 127.8 ($C_{Ar,Fmoc}$), 127.2 ($C_{Ar,Fmoc}$), 125.3 ($2C_{Ar,Fmoc}$), 120.1 ($2C_{Ar,Fmoc}$), 83.8 ($\underline{C}(CH_3)_{3,A}$, $\underline{C}(CH_3)_{3,B}$), 67.2 ($CH_{2,Fmoc}$), 53.4 ($C_{\alpha,hArg}$), 47.3 ($CH_{Fmoc}$), 41.0 ($C_{\delta,hArg}$), 32.0 ($C_{\beta,hArg}$), 29.2 ($C_{\delta,hArg}$), 28.3 ($C(\underline{C}H_3)_{3,A}$), 28.2 ($C(\underline{C}H_3)_{3,A}$), 22.8 ($C_\gamma$). ESI-MS $m/z$ calcd for $C_{33}H_{44}N_6O_{13}^+$ $[M+H]^+$, 611.3; found 611.4. CAS RN: 158478-81-0. The data is in agreement with literature with minor deviations in reported chemical shifts.[12]

**4-(*tert-butyl*)-*N*-((4-(5-(dimethylamino)pentanamido)phenyl)carbamothioyl)benzamide**

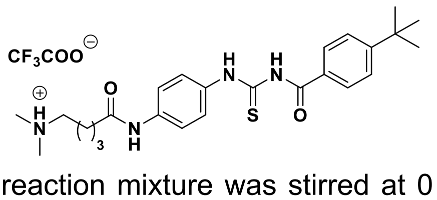

**(tenovin-6).** To a stirring solution of **S35** (100 mg, 0.31 mmol) in 2.0 mL anh. $CH_2Cl_2$ at 0 ˚C was added 5-bromovaleryl chloride (43 µL, 0.32 mmol), followed by $i$$Pr_2NEt$ (106 µL, 0.61 mmol). The reaction mixture was stirred at 0 ˚C for 10 min and then overnight at ambient temperature. Next, dimethylamine (100 µL, 0.56 mmol, 33% in abs. ethanol (~5.6 M) was added and the reaction was stirred for an additional 6 h at ambient temperature. The mixture was diluted with $CH_2Cl_2$ (25 mL) and the resulting organic layer washed with aq. NaOH (1 M, 25 mL) and brine (25 mL), dried over $Na_2SO_4$, filtered and concentrated under reduced pressure to afford the crude product. Preparative reversed-phase HPLC purification afforded **tenovin-6** (28 mg, 20% over 2 steps) as an off-white fluffy material after lyophilisation. $^{1}$H NMR (600 MHz, DMSO-$d_6$) δ 12.60 (s, 1H, CON<u>H</u>Ar), 11.41 (s, 1H, ArN<u>H</u>(C=S)), 10.07 (s, 1H, (C=S)NHCO), 9.47 (s, 1H, N<u>H</u>$^+$(CH3)$_2$), 8.00–7.88 (m, 2H, $H_{Ar}$), 7.73–7.52 (m, 6H, $H_{Ar}$), 3.07 (q, $J$ = 5.4, 3.1 Hz, 2H, C<u>H</u>$_2$N($CH_3$)$_2$), 2.77 (s, 6H, N($CH_3$)$_2$), 2.39 (t, $J$ = 6.8 Hz, 2H, $CH_2$CO), 1.71–1.59 (m, 4H, ($CH_2$)$_2$$CH_2$CO), 1.32 (s, 9H, ($CH_3$)$_3$). $^{13}$C NMR (151 MHz, DMSO-$d_6$) δ 179.0 (C=S), 170.7 ($CH_2$<u>C</u>O), 168.1 (NH<u>C</u>OAr), 157.9 (q, $J$ = 31.0 Hz, residual $CO_{TFA}$), 156.3 (<u>C</u>C($CH_3$)$_3$), 137.3 ($C_{Ar}$), 132.9 ($C_{Ar}$), 129.3 ($C_{Ar}$), 128.6 ($C_{Ar}$), 125.3 ($C_{Ar}$), 124.8 ($C_{Ar}$), 119.0 ($C_{Ar}$), 117.2 (q, $J$ = 298.9 Hz, residual $CF_{3,TFA}$), 56.4 (<u>C</u>H$_2$N($CH_3$)$_2$), 42.2 N($CH_3$)$_2$), 35.5 (<u>C</u>H$_2$CO), 34.9 (<u>C</u>($CH_3$)$_3$), 30.8 (C(<u>C</u>H$_3$)$_3$), 23.4 (N$CH_2$<u>C</u>H$_2$), 21.8 (<u>C</u>H$_2$$CH_2$CO). Analytical HPLC gradient 0–95% eluent II in eluent I (C8; 35 min total runtime), $t_R$ 21.9 min (>98%, $UV_{230}$). ESI-MS $m/z$ calcd for $C_{25}H_{35}N_4O_2S^+$ [M+H]$^+$, 455.2; found 455.3. CAS RN: 1011557-82-6. The data is in agreement with literature.[4]

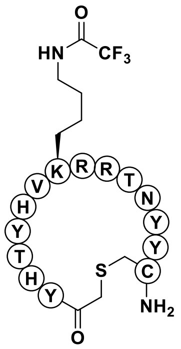

**S2iL5.** The macrocyclic peptide, S2iL5, was synthesized with standard Fmoc-based SPPS using Rink amide linker on TentaGel® resin (167 mg, estimated loading: 0.24 mmol/g), inspired by prior synthesis.[13] The resulting $N$-terminal α-amino group was incubated with a 0.2 M solution of chloroacetyl chloride in DMF (4 mL) and agitated for 40 min at ambient temperature and subsequently washed with DMF (3×5 mL) and $CH_2Cl_2$ (3×5 mL). Global deprotection and cleavage from the resin was performed with a solution of TFA/DODT/TIPS/$H_2O$ (5.0 mL, 92.5:2.5:2.5:2.5; v/v) under agitation at ambient temperature for 3 h. The solution was collected, and the resin was washed with an additional TFA (2.0 mL). Solvent was removed under a stream of nitrogen the peptide was triturated in ice-cold ether. The resulting pellet was dissolved in 20 mL $H_2O$:MeCN (1:1) and $NEt_3$ (2–3 drops) was added and the reaction stirred for 30 min at 42 °C resulting in the formation of an off-white precicipitate. The solution was acidified with a few drops of TFA and concentrated under reduced pressure followed by preparative reversed-phase HPLC purification to afford the macrocycle **S2iL5** (11 mg, 14% based on resin loading) as colorless fluffy material after lyophilization. Analytical HPLC gradient 0–95% eluent II in eluent I (C8; 35 min total runtime), $t_R$ 14.3 min (>98%, $UV_{230}$). HRMS $m/z$ calcd for $C_{90}H_{124}F_3N_{27}O_{23}S^{2+}$ [M+2H]$^{2+}$, 1019.9513; found 1019.9509. CAS RN: 1707311-44-1.

#### Drug treatment for analysis of cytotoxicity in cells

**Cell culture.** All cell culture media contained 10% (v/v) FBS (Thermo Fisher Scientific; cat. #26140079) and 1% penicillin-streptomycin (Sigma-Aldrich; cat. #P4333) unless stated otherwise and cultured at 37 °C with 5% $CO_2$ in a humidified incubator. MCF-7 (Sigma-Aldrich; cat. #86012803) and HeLa (Sigma-Aldrich; cat. #93021013) cells were maintained in Minimum Essential Medium Eagle (MEM, Sigma-Aldrich; cat. #M2279) supplemented with L-glutamine (2.0 mM, Sigma-Aldrich; cat. #G7513) and MEM non-essential amino acid solution (1%, Sigma-Aldrich; cat. #P7145). Jurkat (Sigma-Aldrich; cat. #88042803) cells were maintained in Roswell Park Memorial Institute medium (RPMI-1640, Sigma-Aldrich; cat. #R0883). HEK293T (ATCC; cat. #CRL-1573) cells were maintained in Dulbecco's modified Eagle's medium (DMEM, Thermo Scientific; cat. #11965118). Cell lines were sub-cultured every 2-4 days.

**Cell viability assays.** Cell viability was assessed using MTT[14] cell growth kits (Merck Millipore; cat. #CT02) as previously described.[15] In short, cells were seeded into flat 96-well plates (Corning, cat. #3596) at 5,000 cells/well (HEK293T and HeLa) or 10,000 cells/well (Jurkat and MCF-7). After 24 h, test compounds were added to final concentrations ranging from 100–0.02 µM and incubated for 72 h. Cell viability was measured following the manufacturer's protocol. The relative cell viability in presence of test compounds was measured at 570 nm normalized to the DMSO-treated controls after background subtraction at 630 nm on a plate reader. All viability assays were performed as duplicates of triplicates. GraphPad Prism (vers. 8.1.2) was used to determine $EC_{50}$ values.

#### Supporting references

1 N. Rajabi, M. Auth, K. R. Troelsen, M. Pannek, D. P. Bhatt, M. Fontenas, M. D. Hirschey, C. Steegborn, A. S. Madsen and C. A. Olsen, *Angew. Chem. Int. Ed.*, 2017, **56**, 14836–14841.

2 N. Rajabi, A. L. Nielsen and C. A. Olsen, *ACS Med. Chem. Lett.*, 2020, **11**, 1886–1892.

3 M. Bæk, P. Martín-Gago, J. S. Laursen, J. L. H. Madsen, S. Chakladar and C. A. Olsen, *Chem. – A Eur. J.*, 2020, **26**, 3862–3869.

4 A. R. McCarthy, L. Pirrie, J. J. Hollick, S. Ronseaux, J. Campbell, M. Higgins, O. D. Staples, F. Tran, A. M. Z. Slawin, S. Lain and N. J. Westwood, *Bioorganic Med. Chem.*, 2012, **20**, 1779–1793.

5 Y. L. Chiang and H. Lin, *Org. Biomol. Chem.*, 2016, **14**, 2186–2190.

6

7 M. Gude, J. Ryf and P. D. White, *Int. J. Pept. Res. Ther.*, 2002, **9**, 203–206.

8 E. Kaiser, R. L. Colescott, C. D. Bossinger and P. I. Cook, *Anal. Biochem.*, 1970, **34**, 595–598.

9 H. Jing, J. Hu, B. He, Y. L. Negrón Abril, J. Stupinski, K. Weiser, M. Carbonaro, Y. L. Chiang, T. Southard, P. Giannakakou, R. S. Weiss and H. Lin, *Cancer Cell*, 2016, **29**, 297–310.

10 I. Galleano, J. Nielsen, A. S. Madsen and C. A. Olsen, *Synlett*, 2017, **28**, 2169–2173.

11 H. Zhang, A. L. Nielsen, M. W. Boesgaard, K. Harpsøe, N. L. Daly, X.-F. Xiong, C. R. Underwood, L. M. Haugaard-Kedström, H. Bräuner-Osborne, D. E. Gloriam and K. Strømgaard, *Eur. J. Med. Chem.*, 2018, **156**, 847–860.

12 S. Frølund, A. Bella, A. S. Kristensen, H. L. Ziegler, M. Witt, C. A. Olsen, K. Strømgaard, H. Franzyk and J. W. Jaroszewski, *J. Med. Chem.*, 2010, **53**, 7441–7451.

13 K. Yamagata, Y. Goto, H. Nishimasu, J. Morimoto, R. Ishitani, N. Dohmae, N. Takeda, R. Nagai, I. Komuro, H. Suga and O. Nureki, *Structure*, 2014, **22**, 345–352.

14 T. Mosmann, *J. Immunol. Methods*, 1983, **65**, 55–63.

15 B. Kitir, A. R. Maolanon, R. G. Ohm, A. R. Colaço, P. Fristrup, A. S. Madsen and C. A. Olsen, *Biochemistry*, 2017, **56**, 5134–5146.

### Full Western blot images

DMSO control CETSA experiment 1

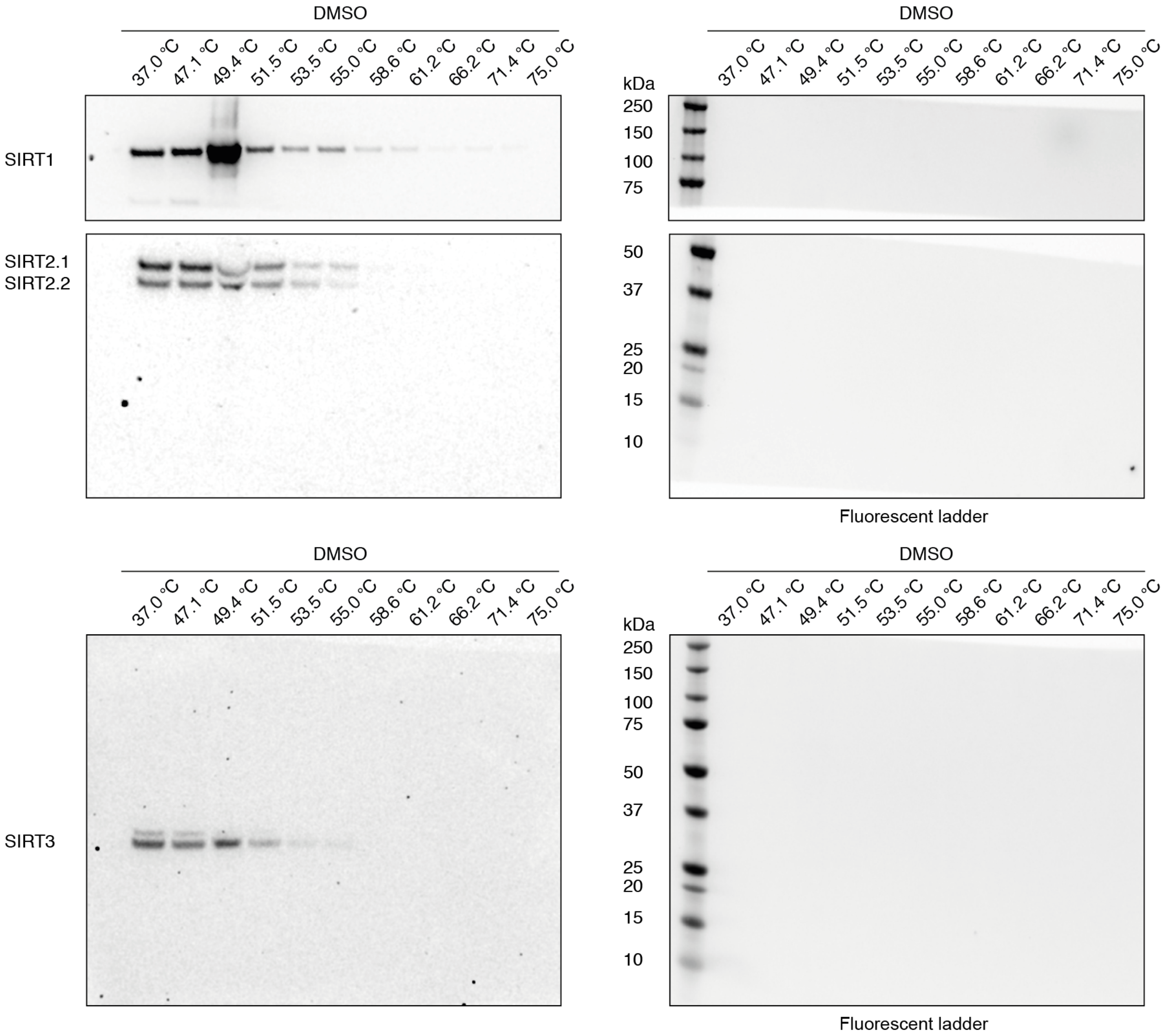

DMSO control CETSA experiment 2

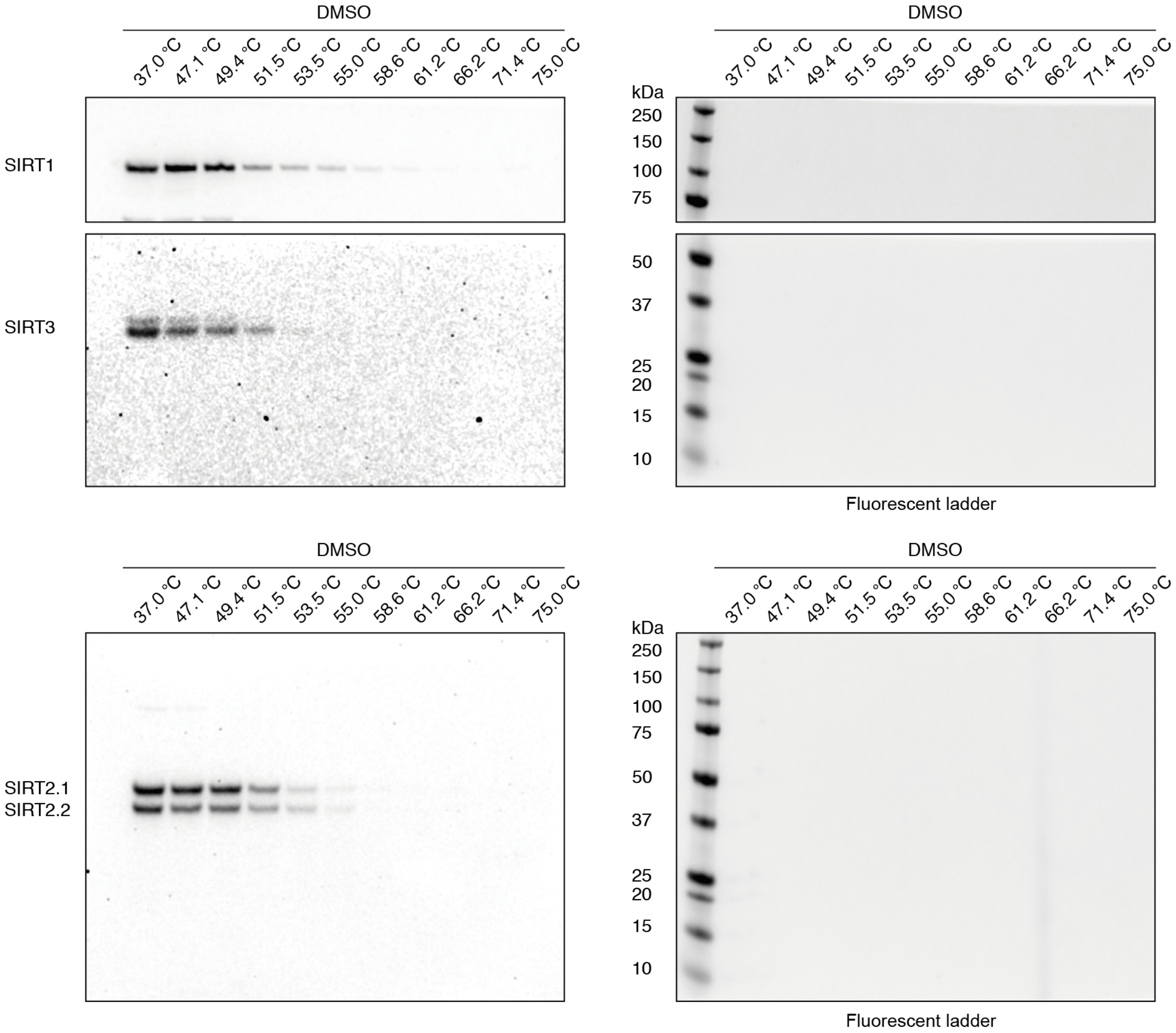

DMSO control CETSA experiment 3

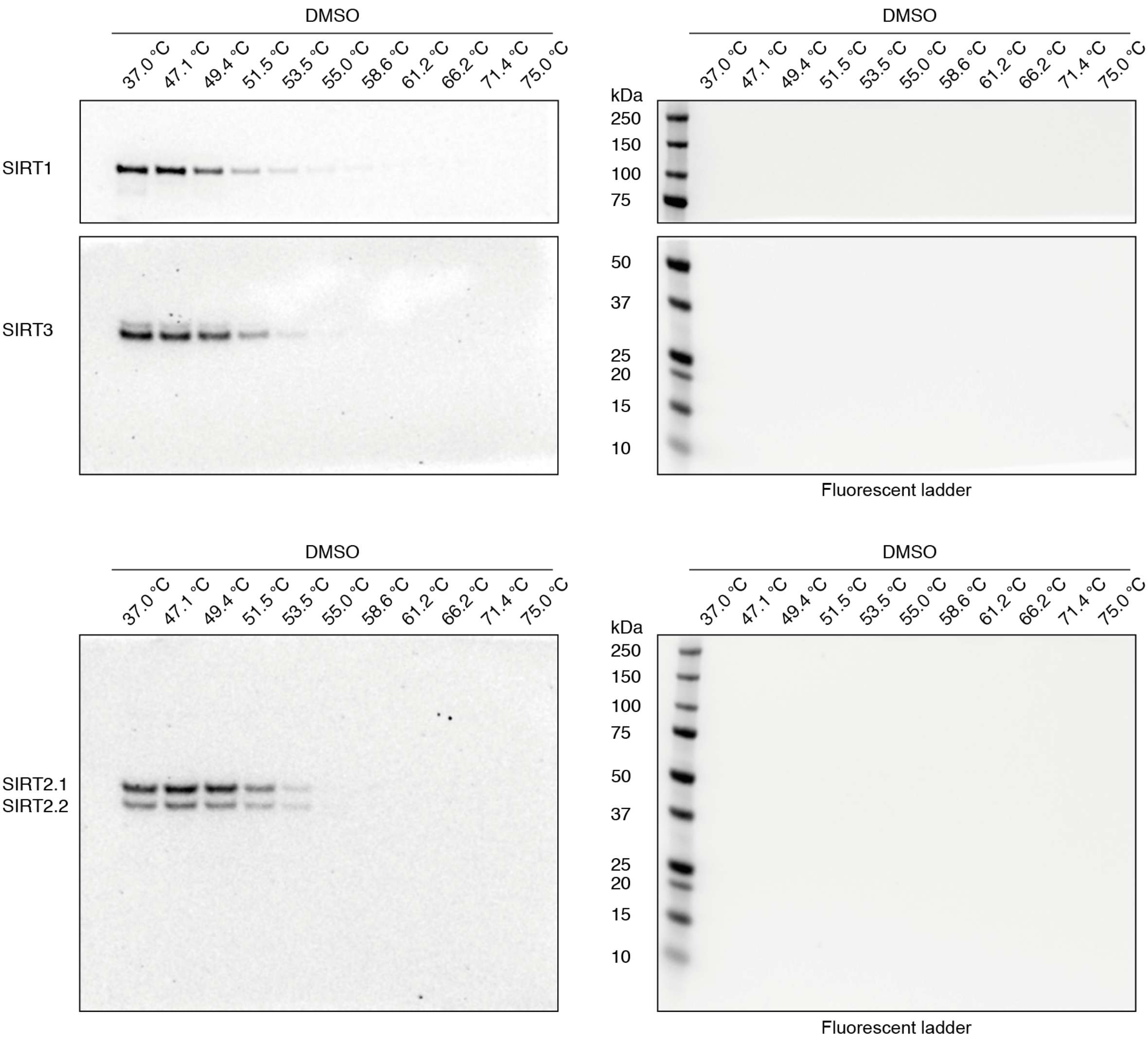

CETSA experiment 1: Compound **26**

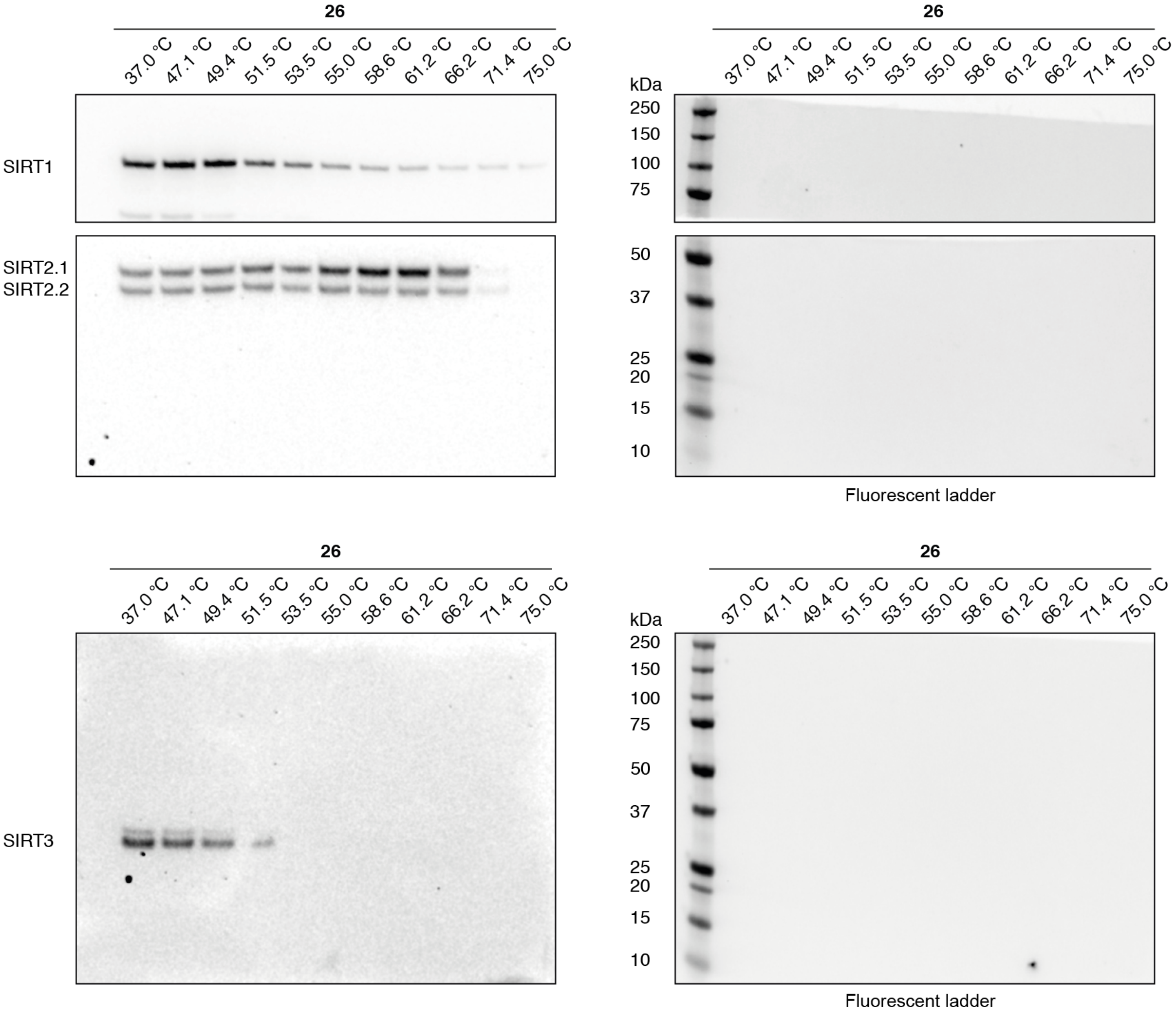

CETSA experiment 2: Compound **26**

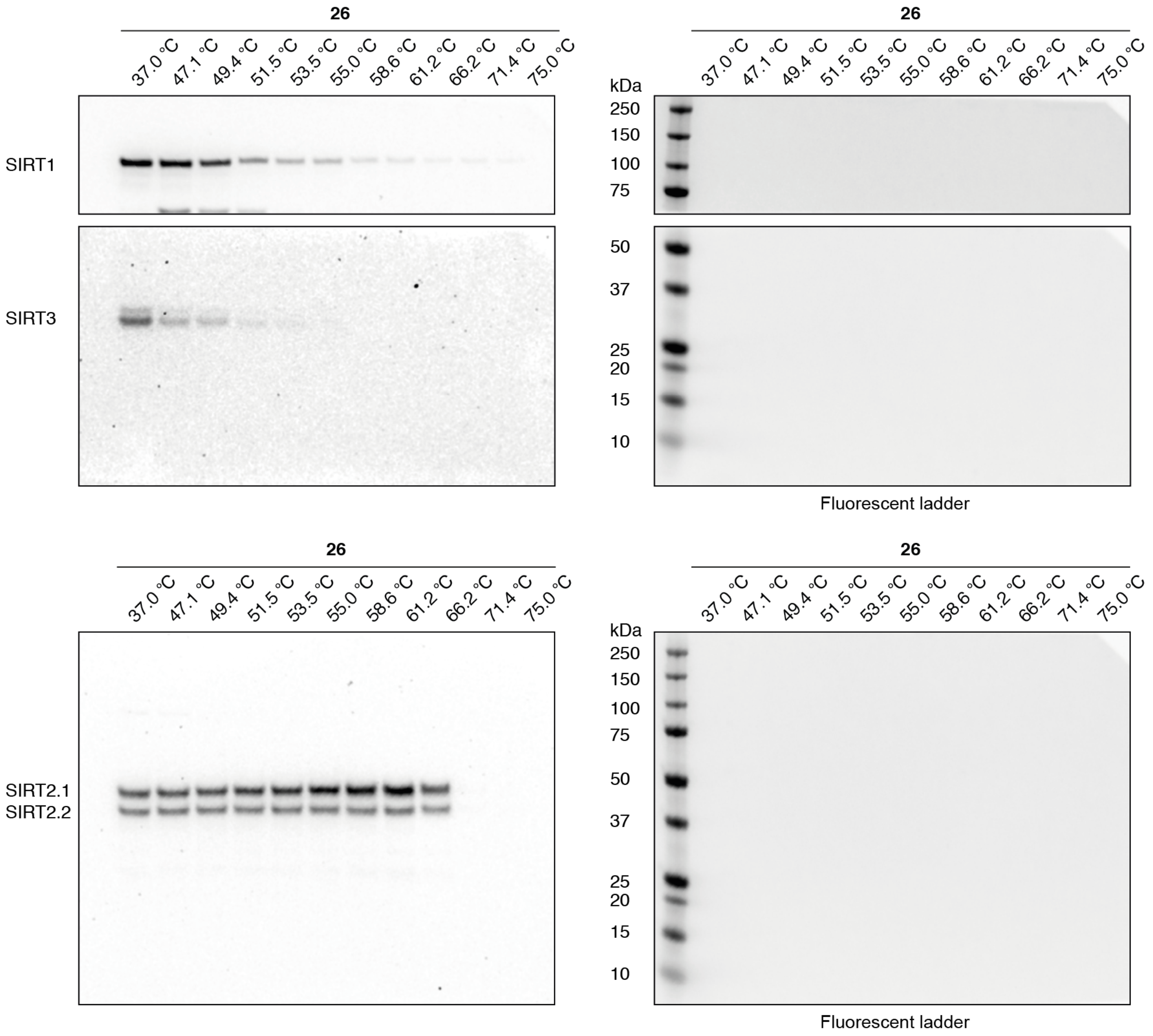

CETSA experiment 3: Compound **26**

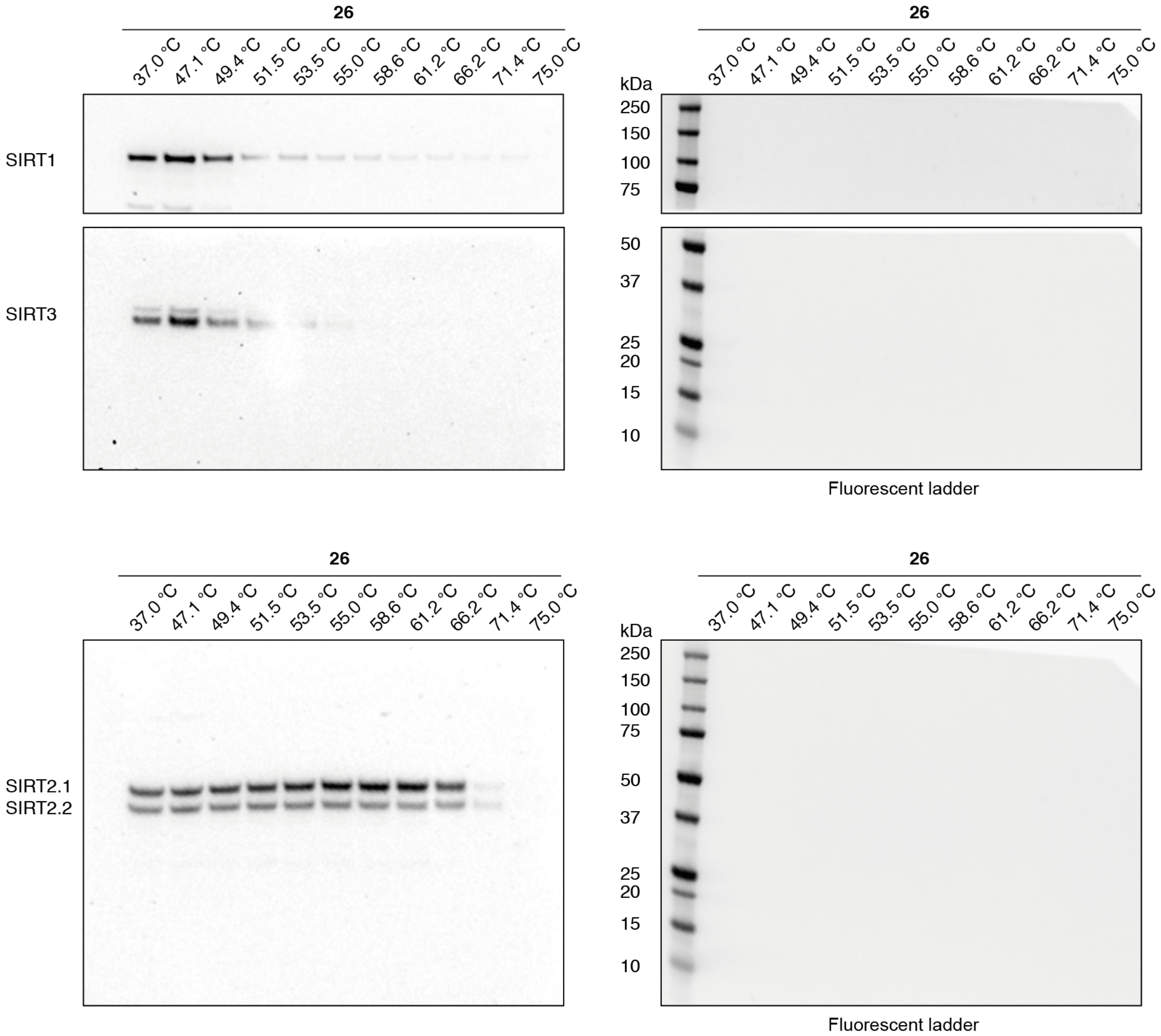

CETSA experiment 1: Compound **26-D**

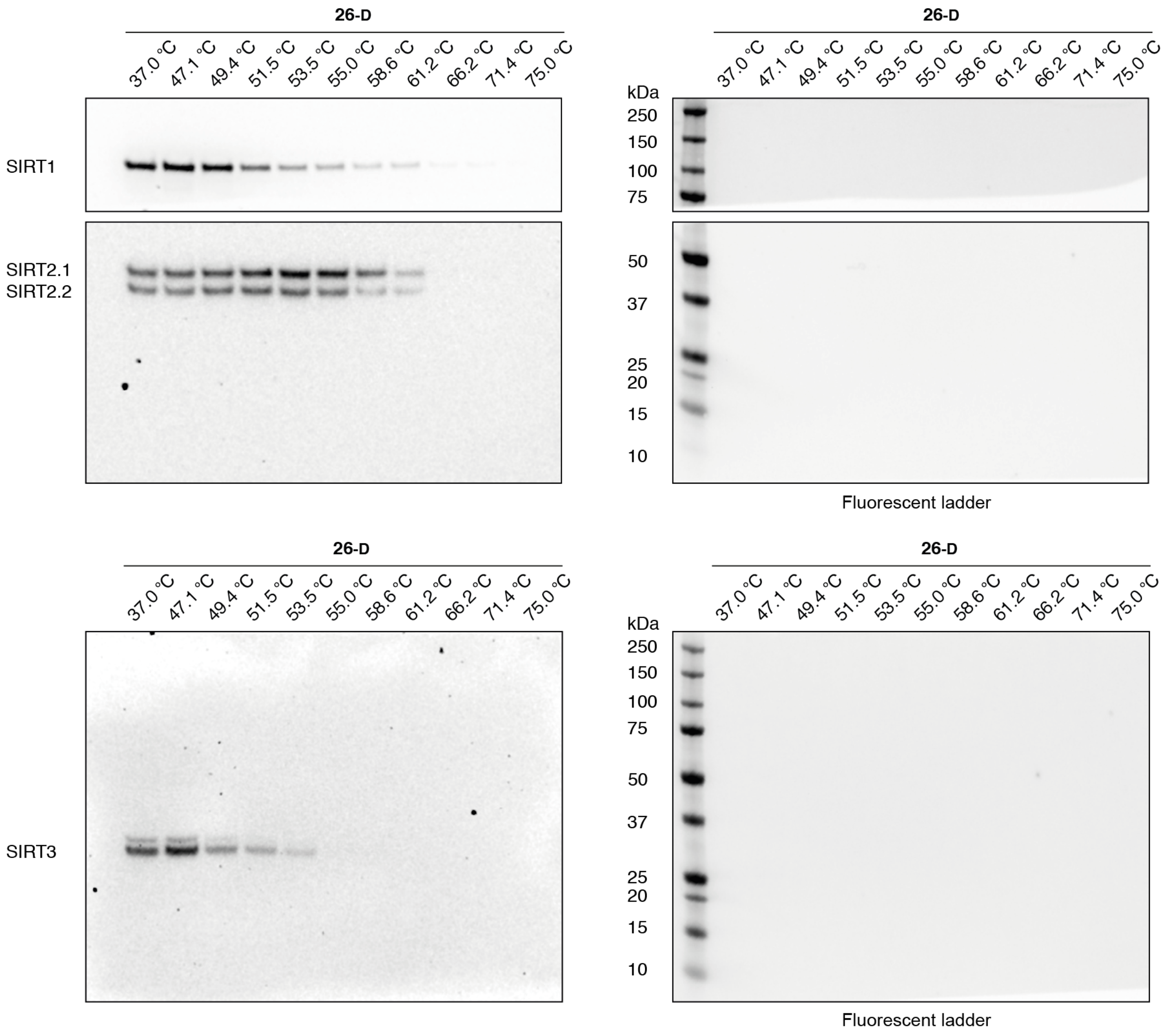

CETSA experiment 2: Compound **26-D**

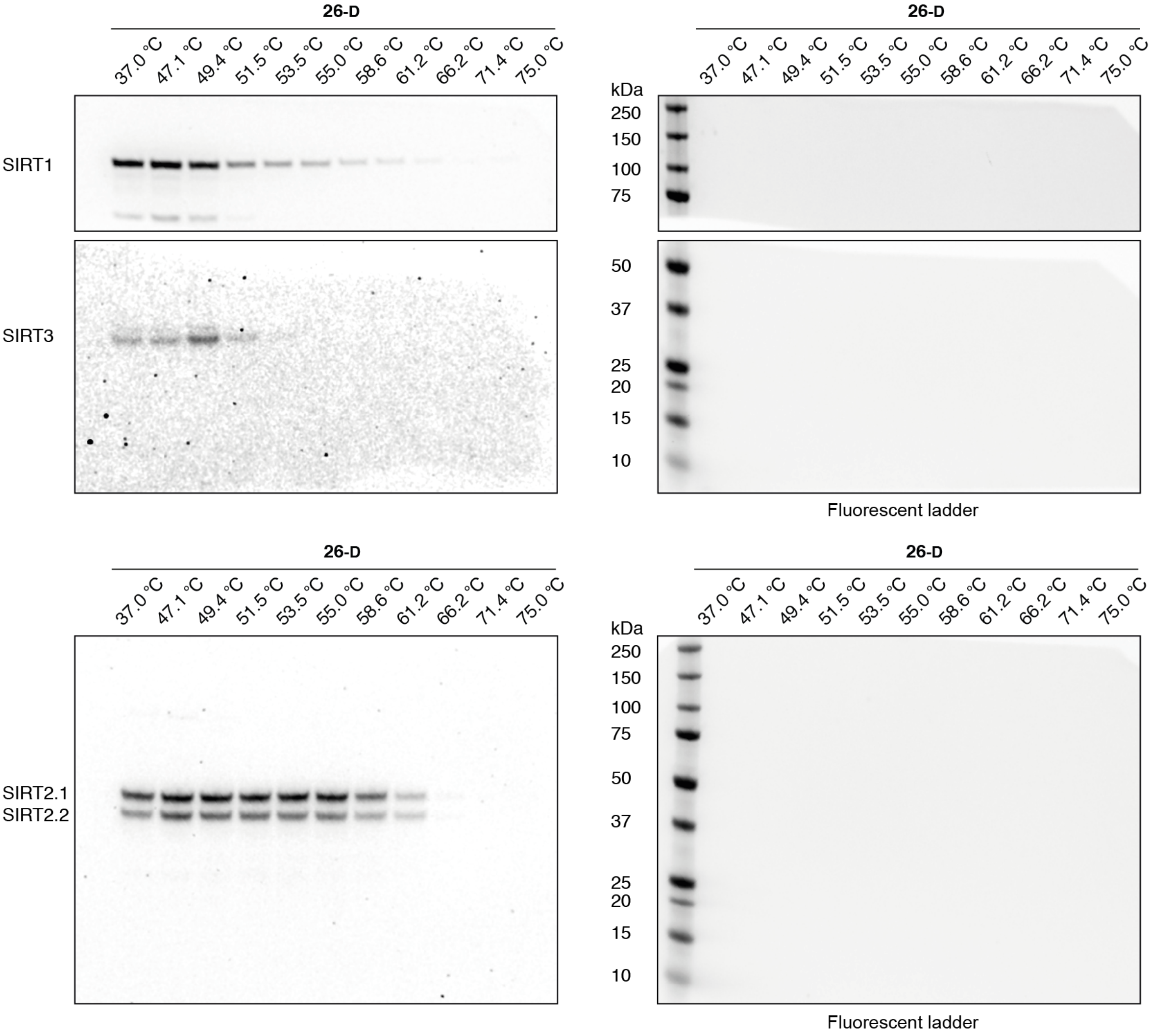

CETSA experiment 3: Compound **26-D**

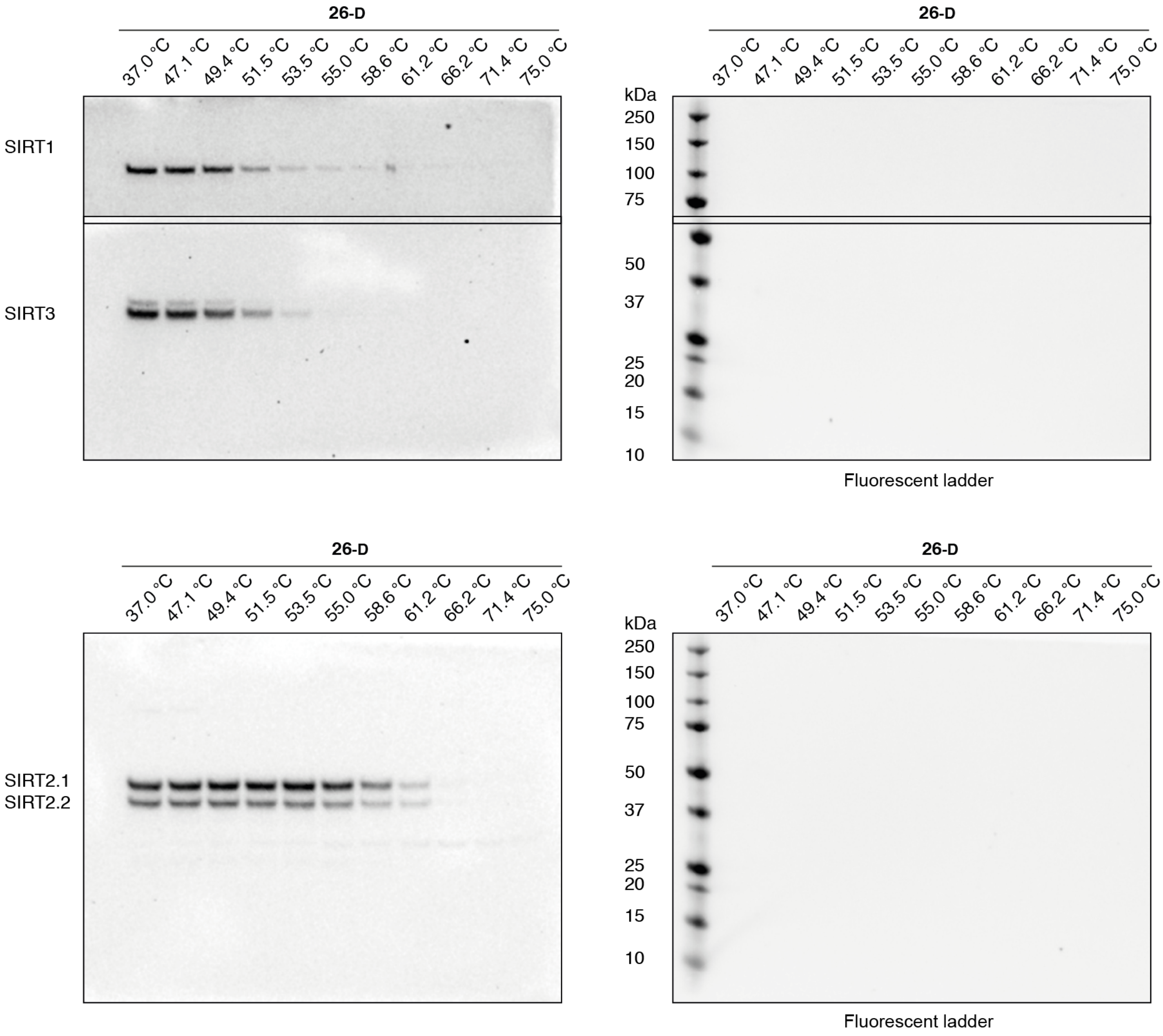

CETSA experiment 1: Compound **TM**

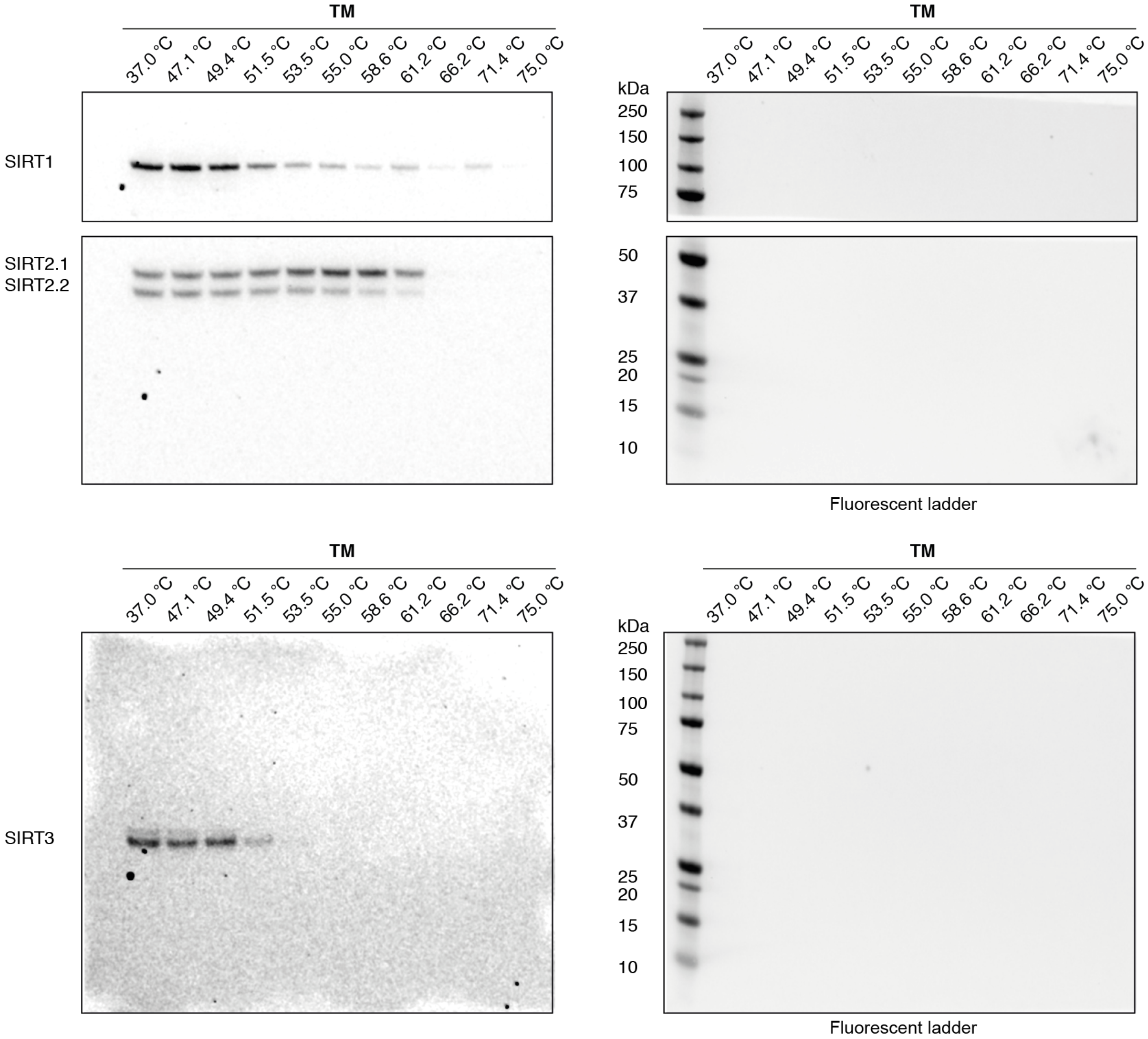

CETSA experiment 2: Compound **TM**

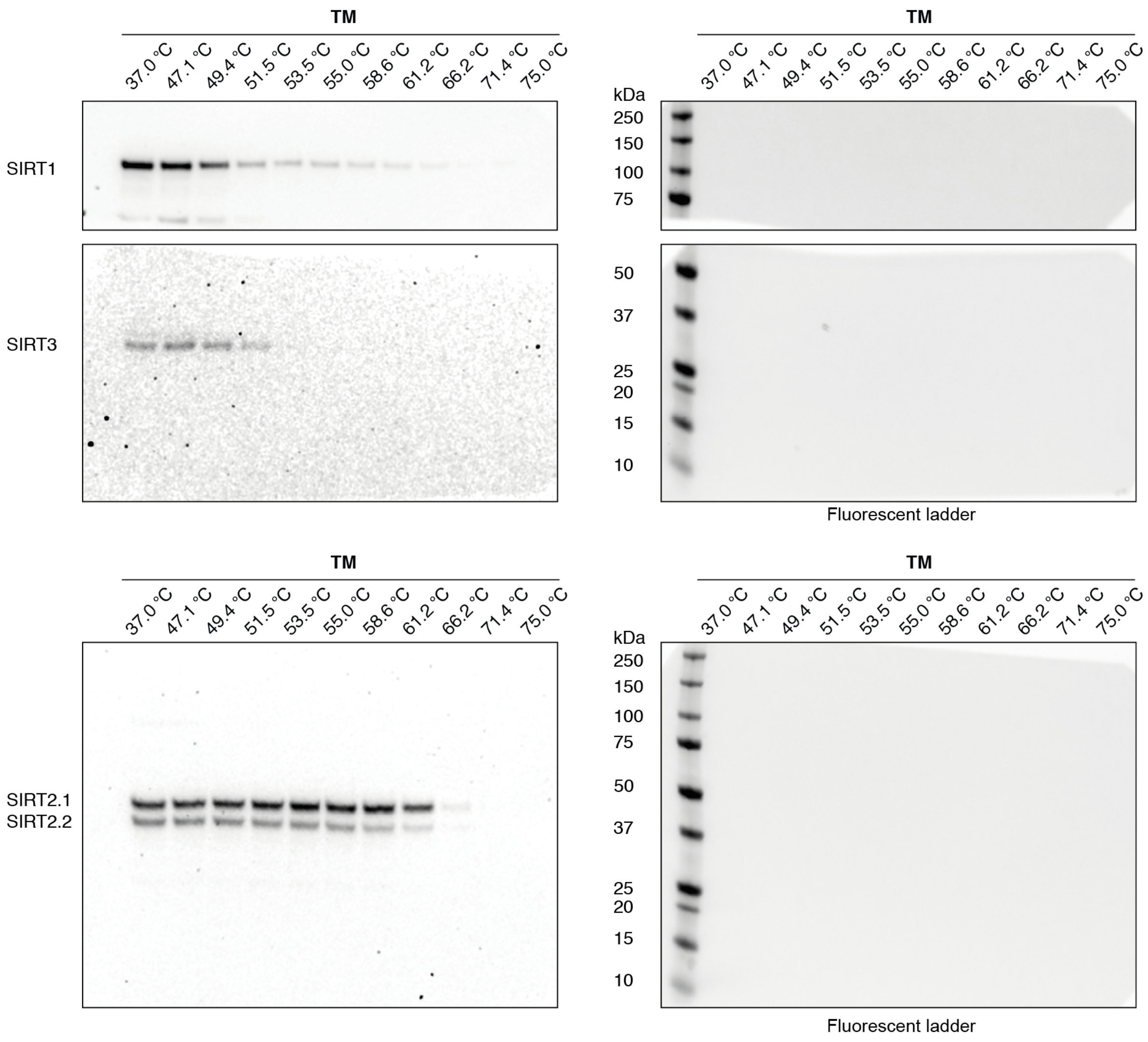

CETSA experiment 3: Compound **TM**

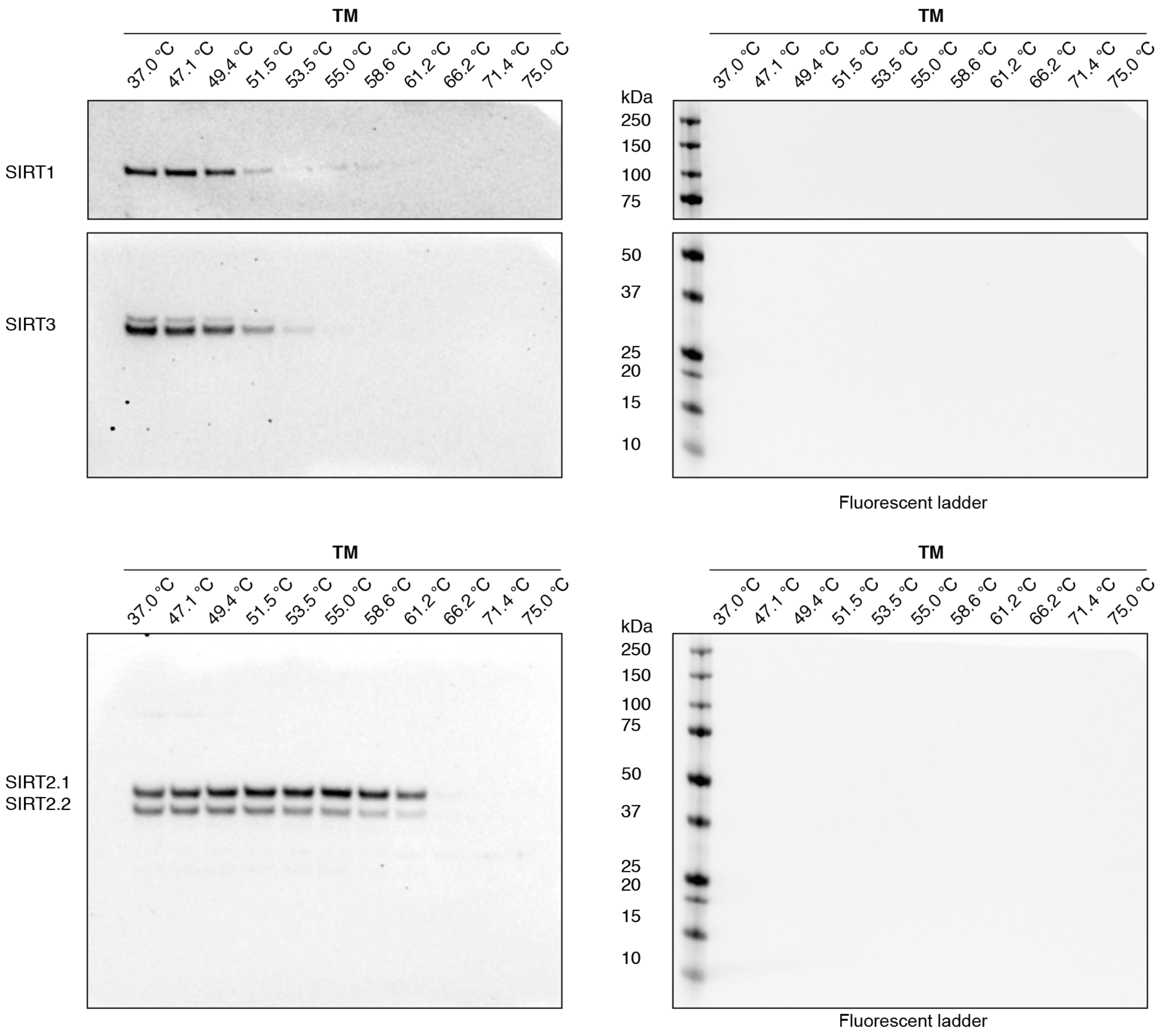

### Analytical HPLC spectra of final compounds

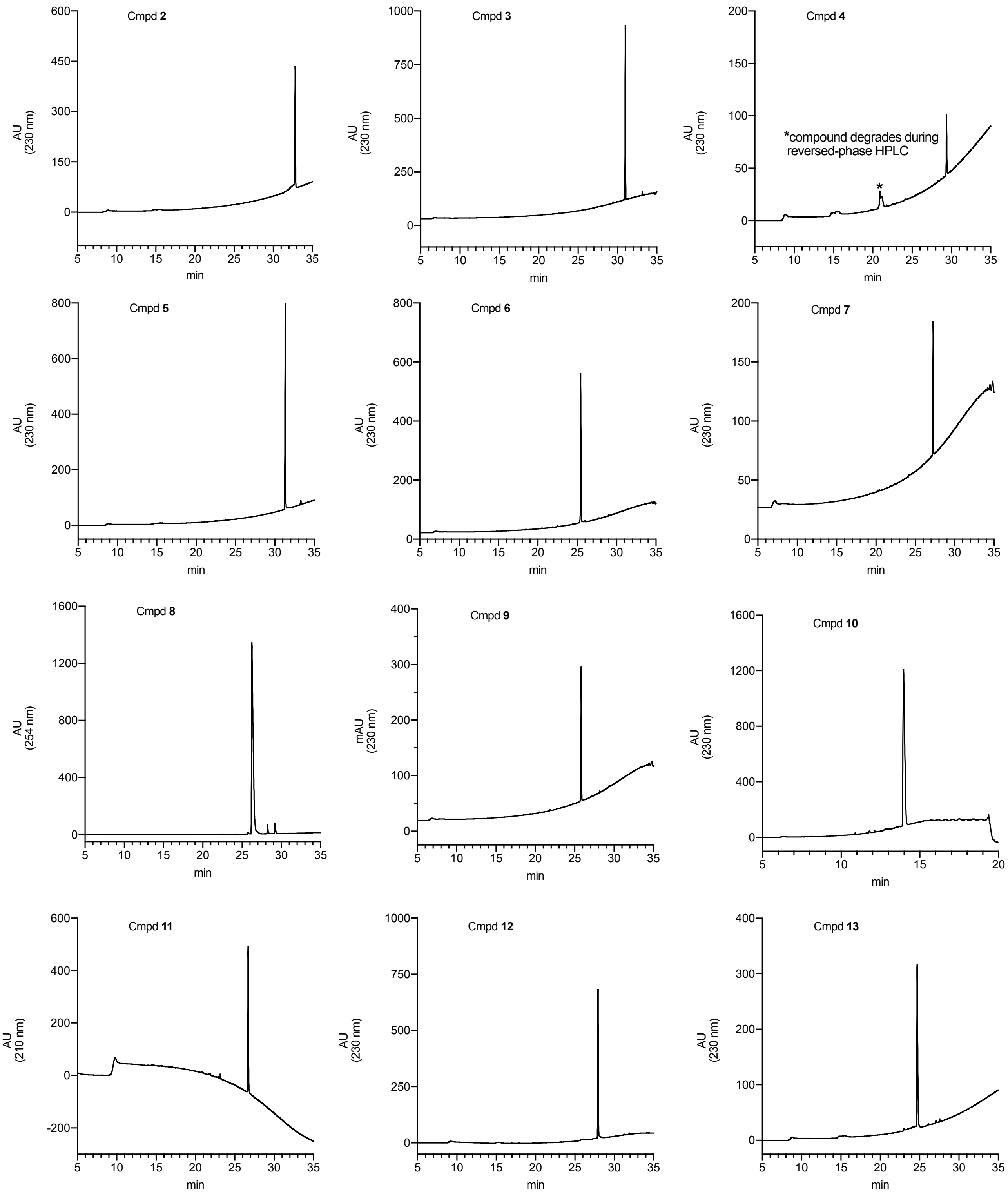

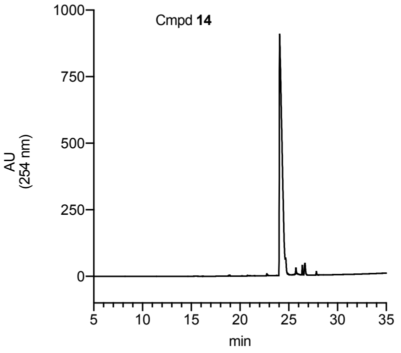
Cmpd **14**
AU (254 nm)
min

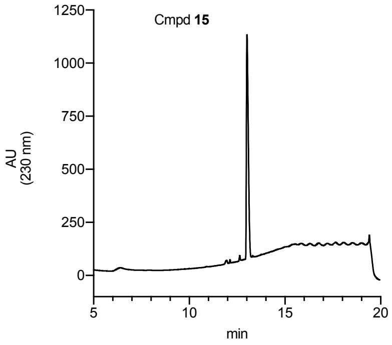
Cmpd **15**
AU (230 nm)
min

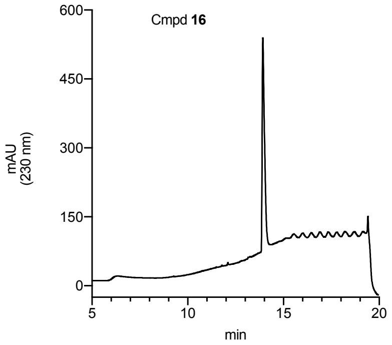
Cmpd **16**
mAU (230 nm)
min

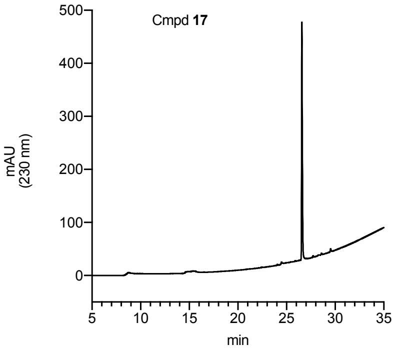
Cmpd **17**
mAU (230 nm)
min

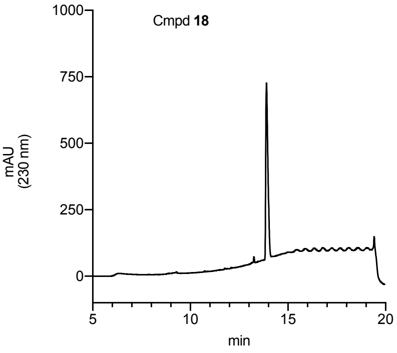
Cmpd **18**
mAU (230 nm)
min

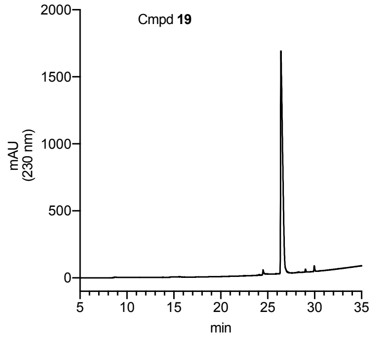
Cmpd **19**
mAU (230 nm)
min

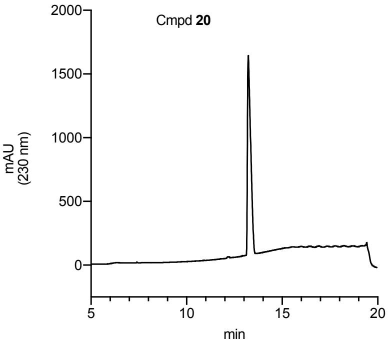
Cmpd **20**
mAU (230 nm)
min

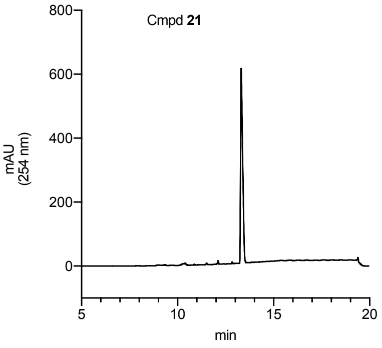
Cmpd **21**
mAU (254 nm)
min

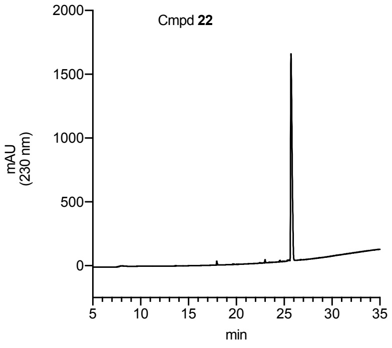
Cmpd **22**
mAU (230 nm)
min

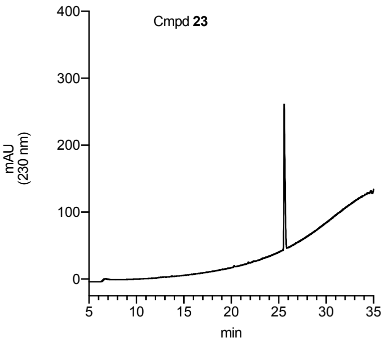
Cmpd **23**
mAU (230 nm)
min

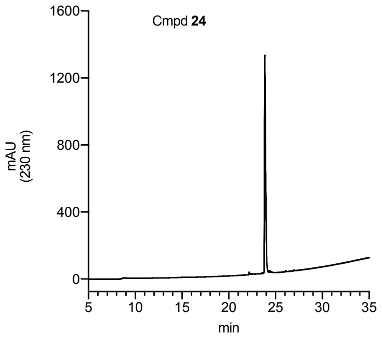
Cmpd **24**
mAU (230 nm)
min

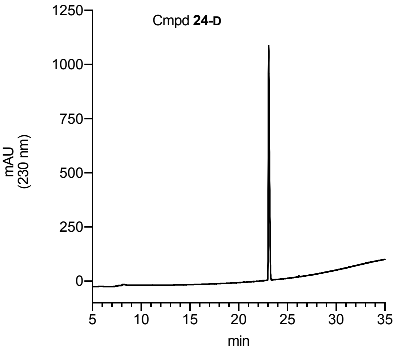
Cmpd **24-D**
mAU (230 nm)
min

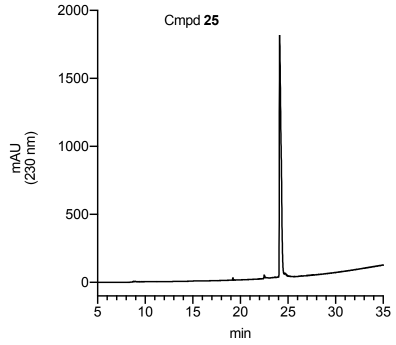
Cmpd **25**
mAU (230 nm)
min

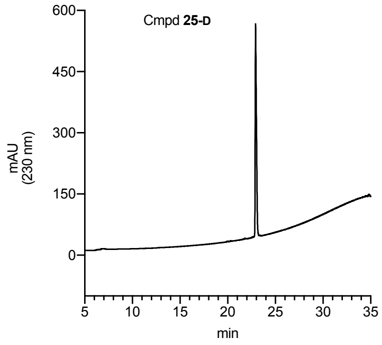
Cmpd **25-D**
mAU (230 nm)
min

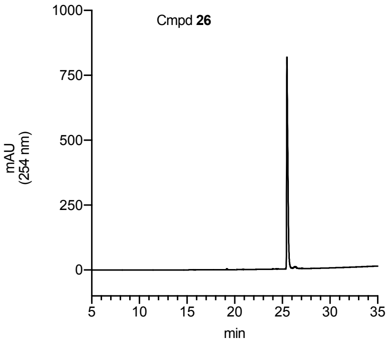
Cmpd **26**
mAU (254 nm)
min

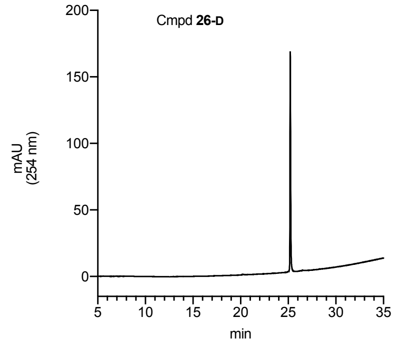
Cmpd **26-D**
mAU (254 nm)
min

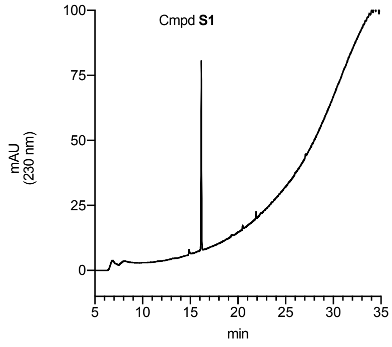
Cmpd **S1**
mAU (230 nm)
min

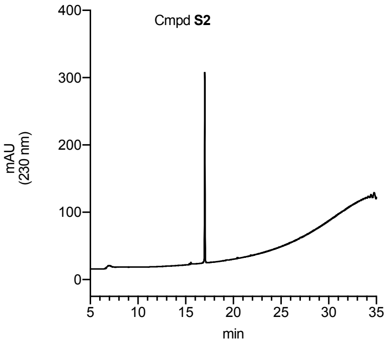
Cmpd **S2**
mAU (230 nm)
min

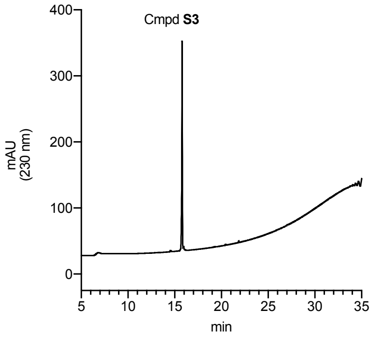
Cmpd **S3**
mAU (230 nm)
min

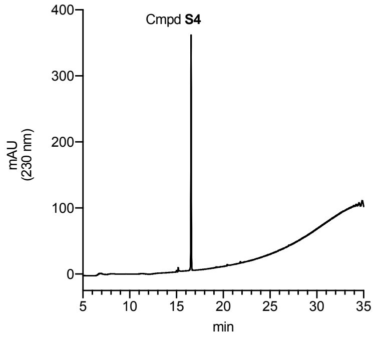
Cmpd **S4**
mAU (230 nm)
min

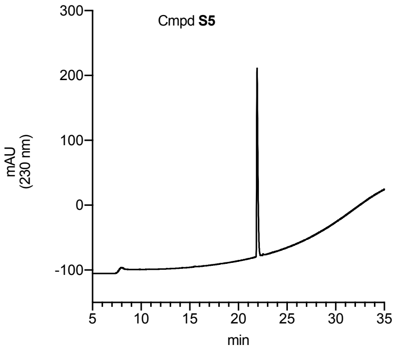
Cmpd **S5**
mAU (230 nm)
min

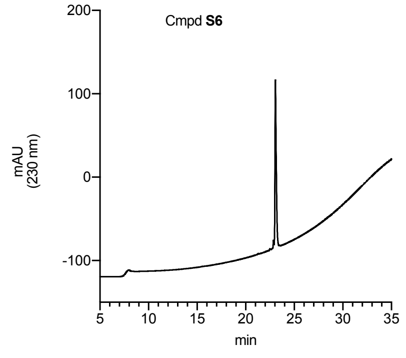
Cmpd **S6**
mAU (230 nm)
min

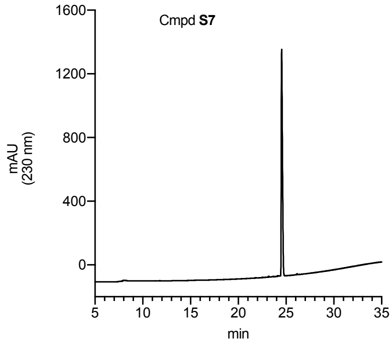
Cmpd **S7**
mAU (230 nm)
min

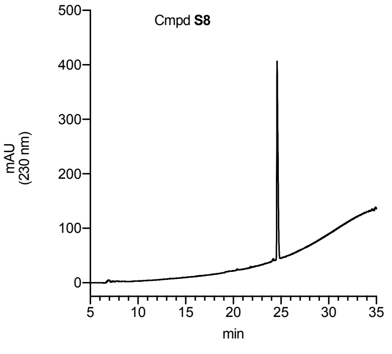
Cmpd **S8**
mAU (230 nm)
min

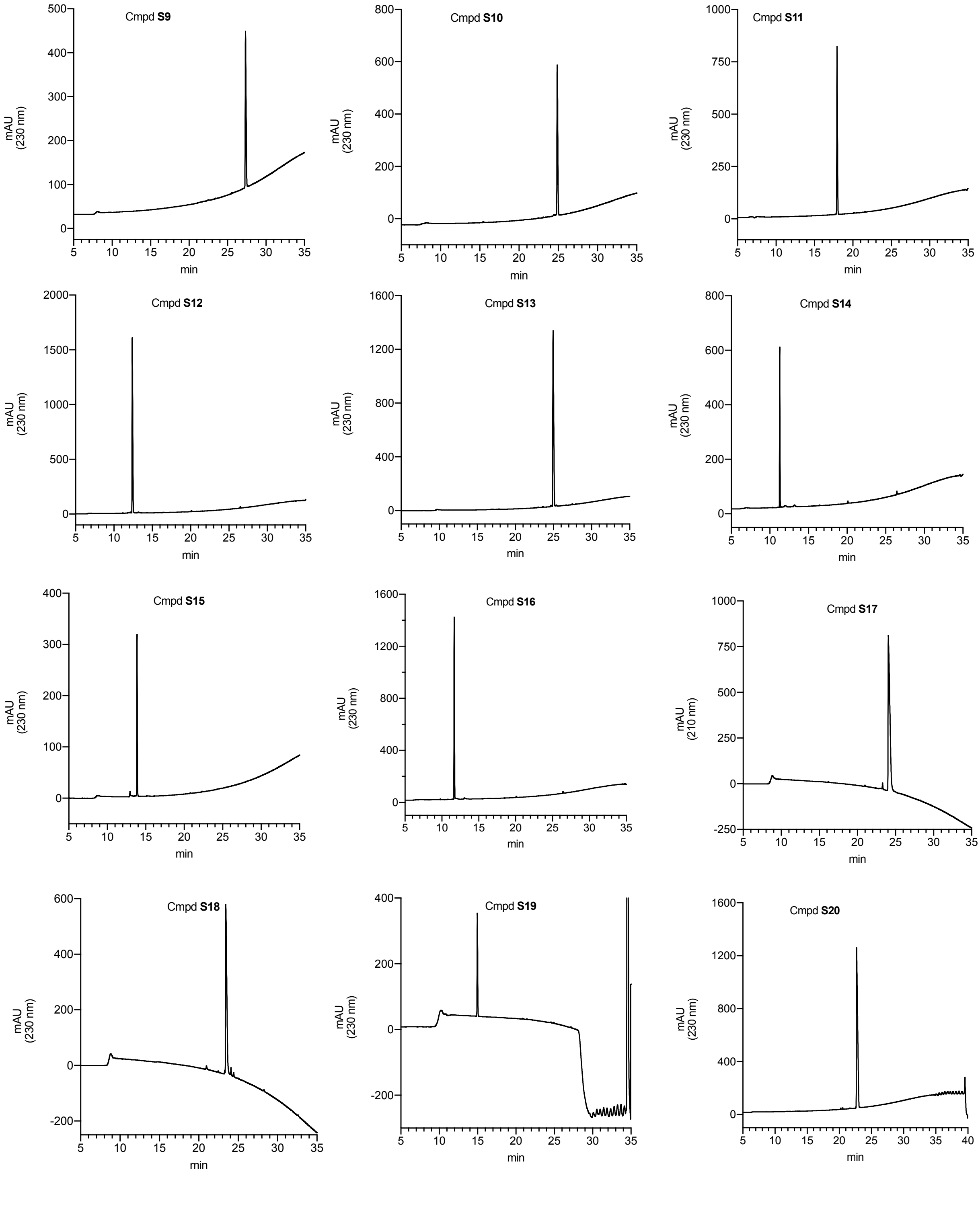

Cmpd S9
Cmpd S10
Cmpd S11
Cmpd S12
Cmpd S13
Cmpd S14
Cmpd S15
Cmpd S16
Cmpd S17
Cmpd S18
Cmpd S19
Cmpd S20
mAU (230 nm)
mAU (210 nm)
min

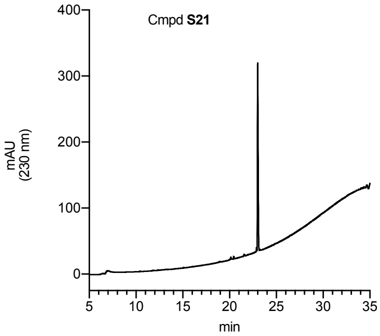
Cmpd S21
mAU (230 nm)
400
300
200
100
0
5 10 15 20 25 30 35
min

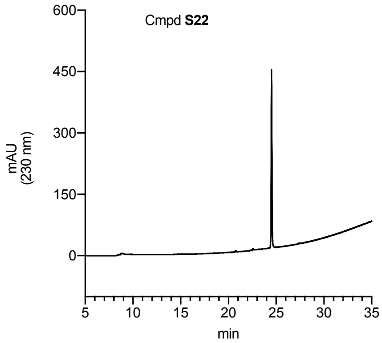
Cmpd S22
mAU (230 nm)
600
450
300
150
0
5 10 15 20 25 30 35
min

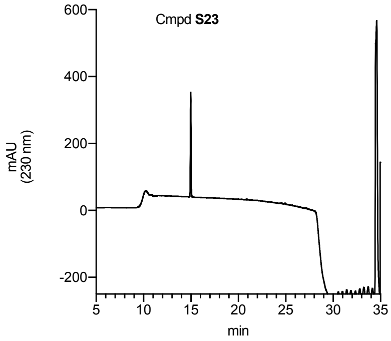
Cmpd S23
mAU (230 nm)
600
400
200
0
-200
5 10 15 20 25 30 35
min

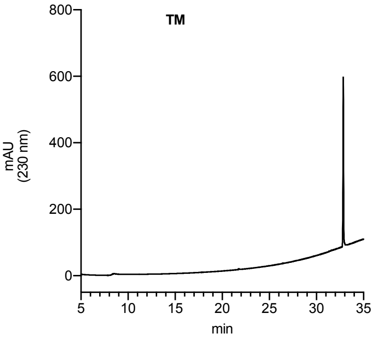
TM
mAU (230 nm)
800
600
400
200
0
5 10 15 20 25 30 35
min

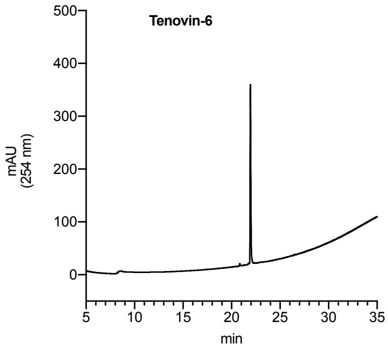
Tenovin-6
mAU (254 nm)
500
400
300
200
100
0
5 10 15 20 25 30 35
min

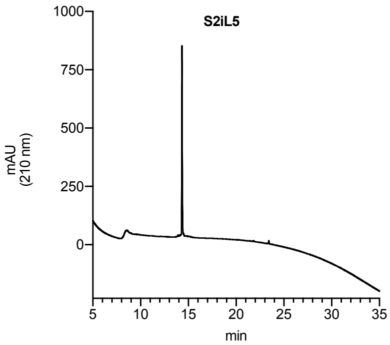
S2iL5
mAU (210 nm)
1000
750
500
250
0
5 10 15 20 25 30 35
min

### NMR spectra

#### $^{1}H$ and $^{13}C$ spectra of compound **2**

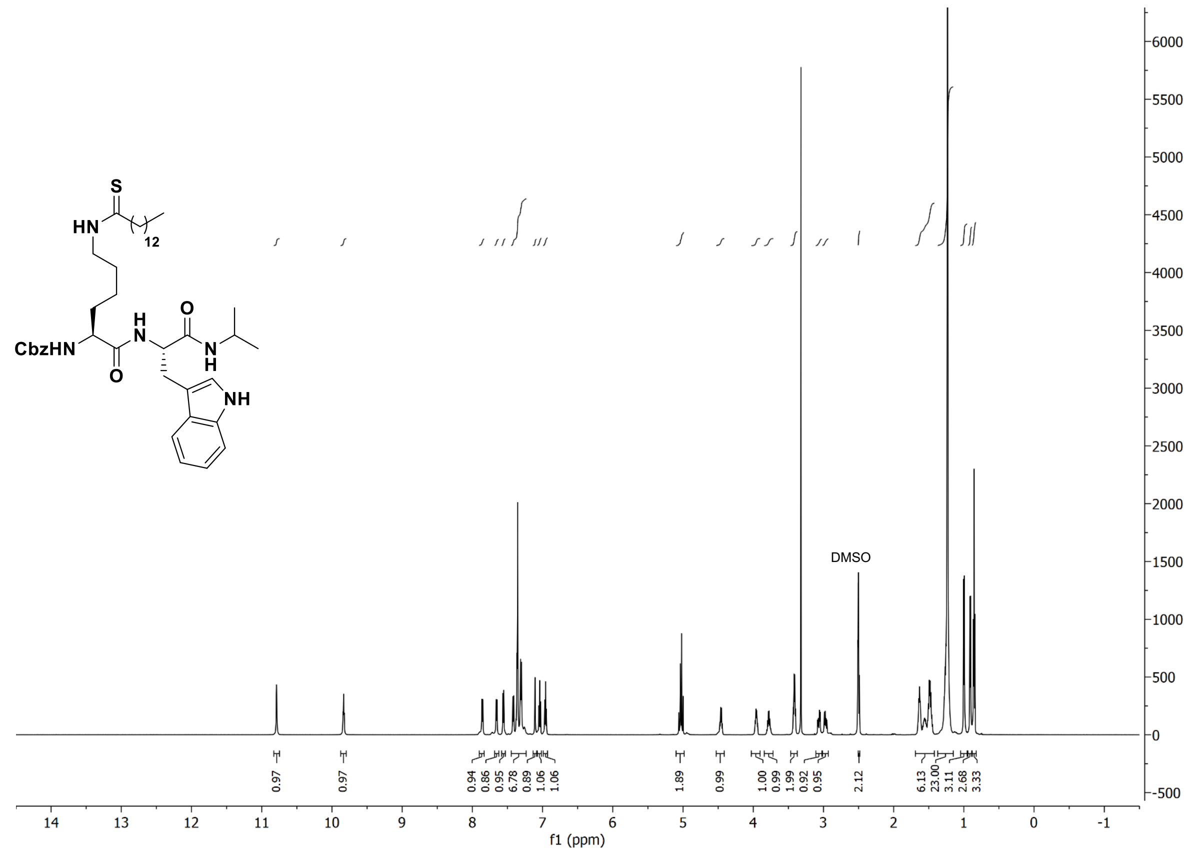

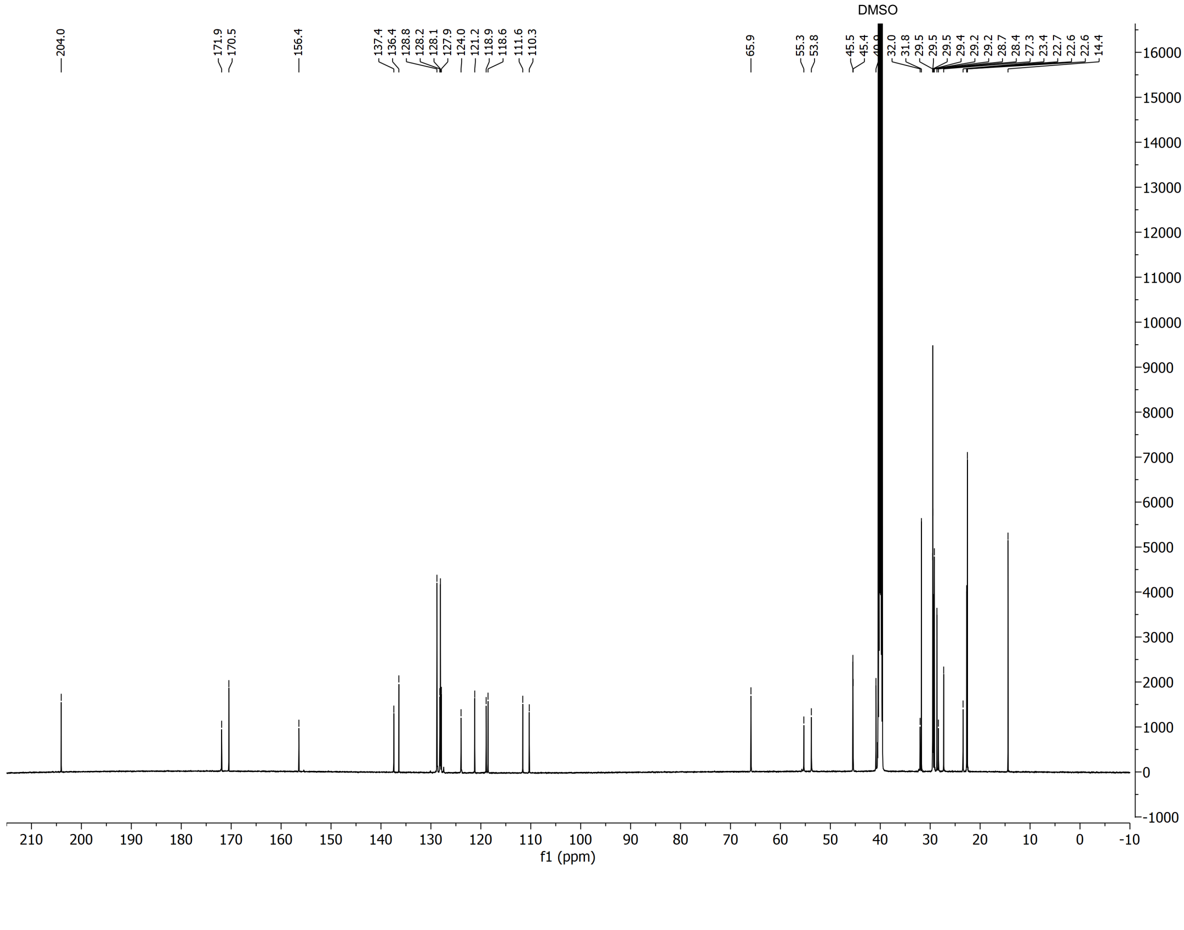

[1]H and [13]C spectra of compound **3**

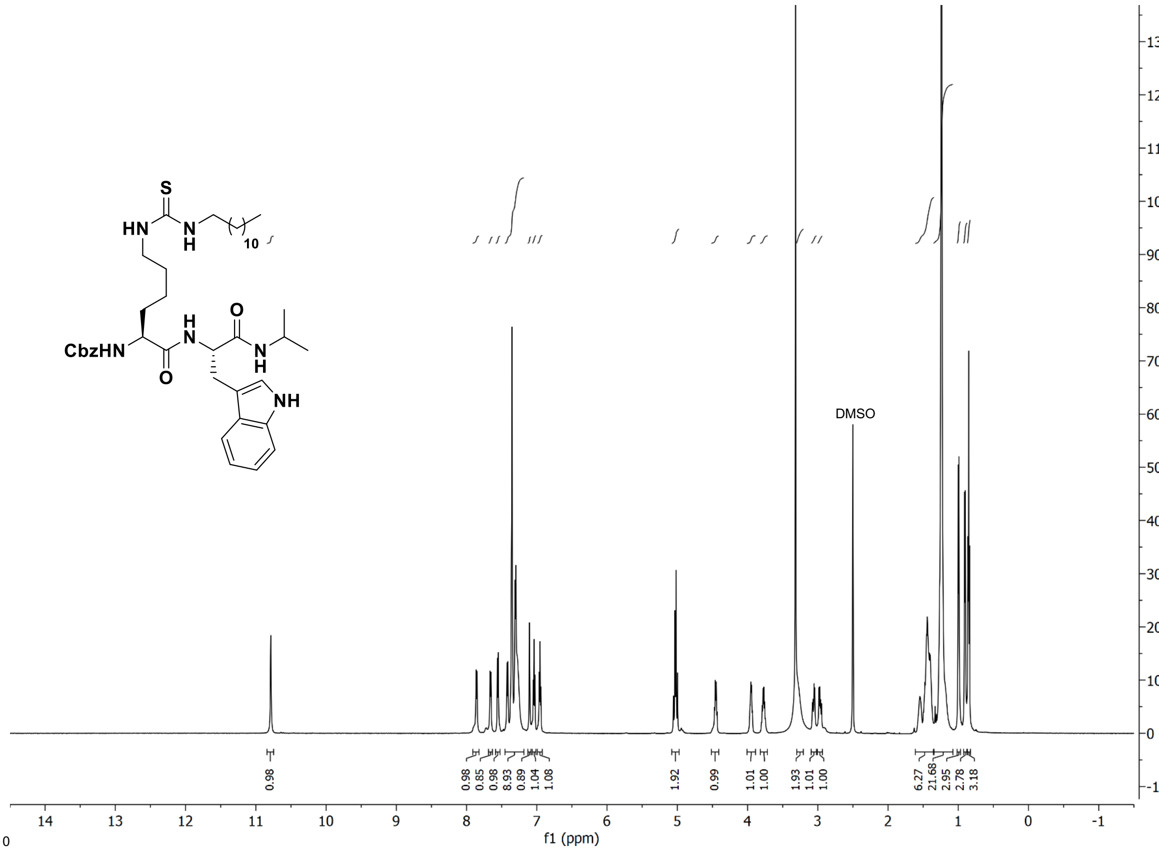

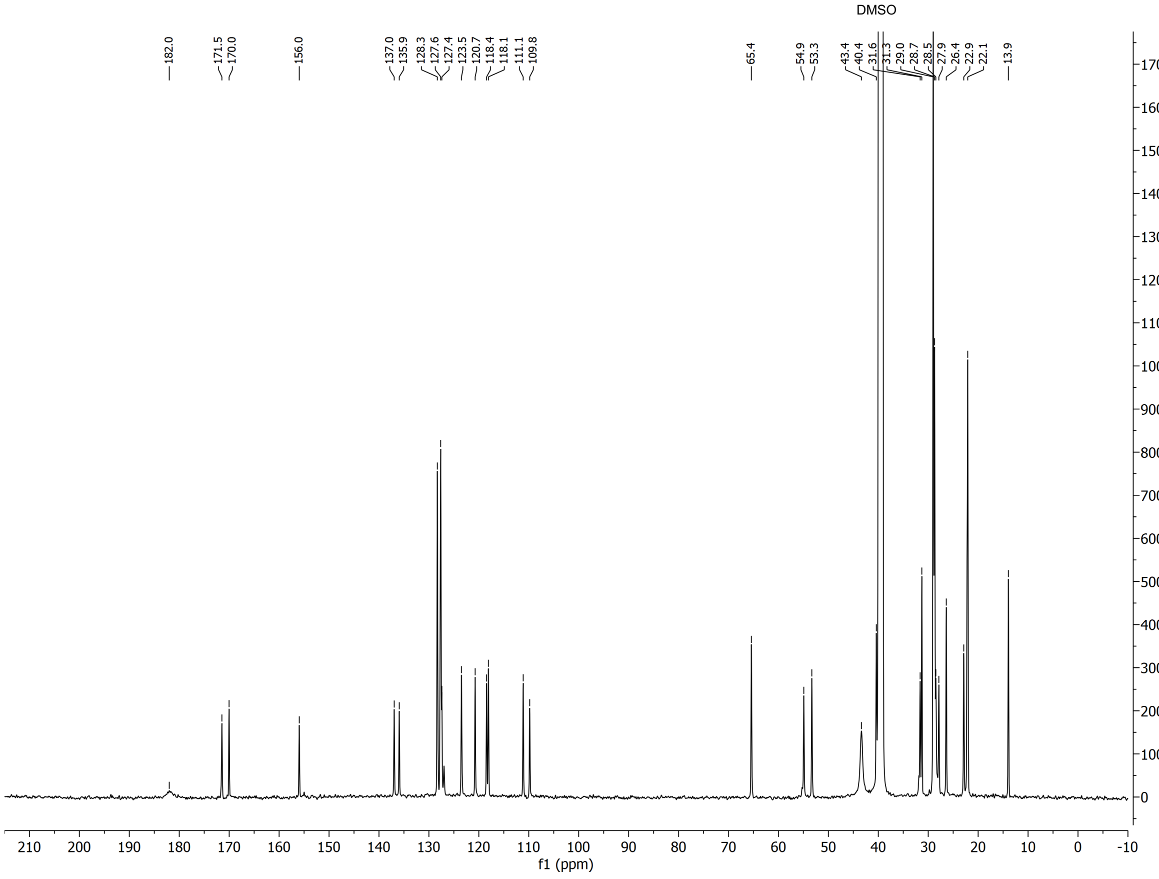

$^{1}H$ and $^{13}C$ spectra of compound **4**

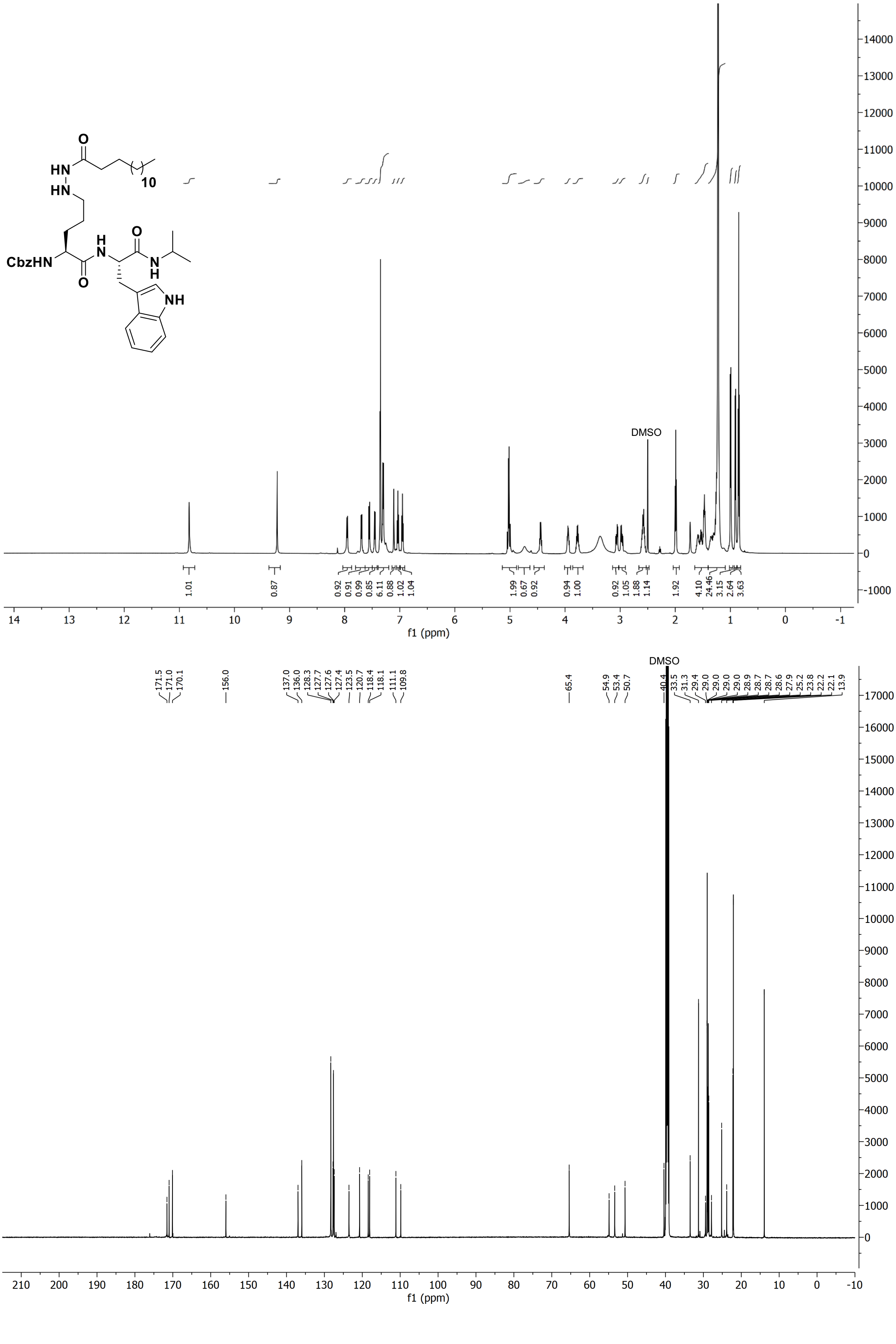

#### $^{1}H$ and $^{13}C$ spectra of compound **5**

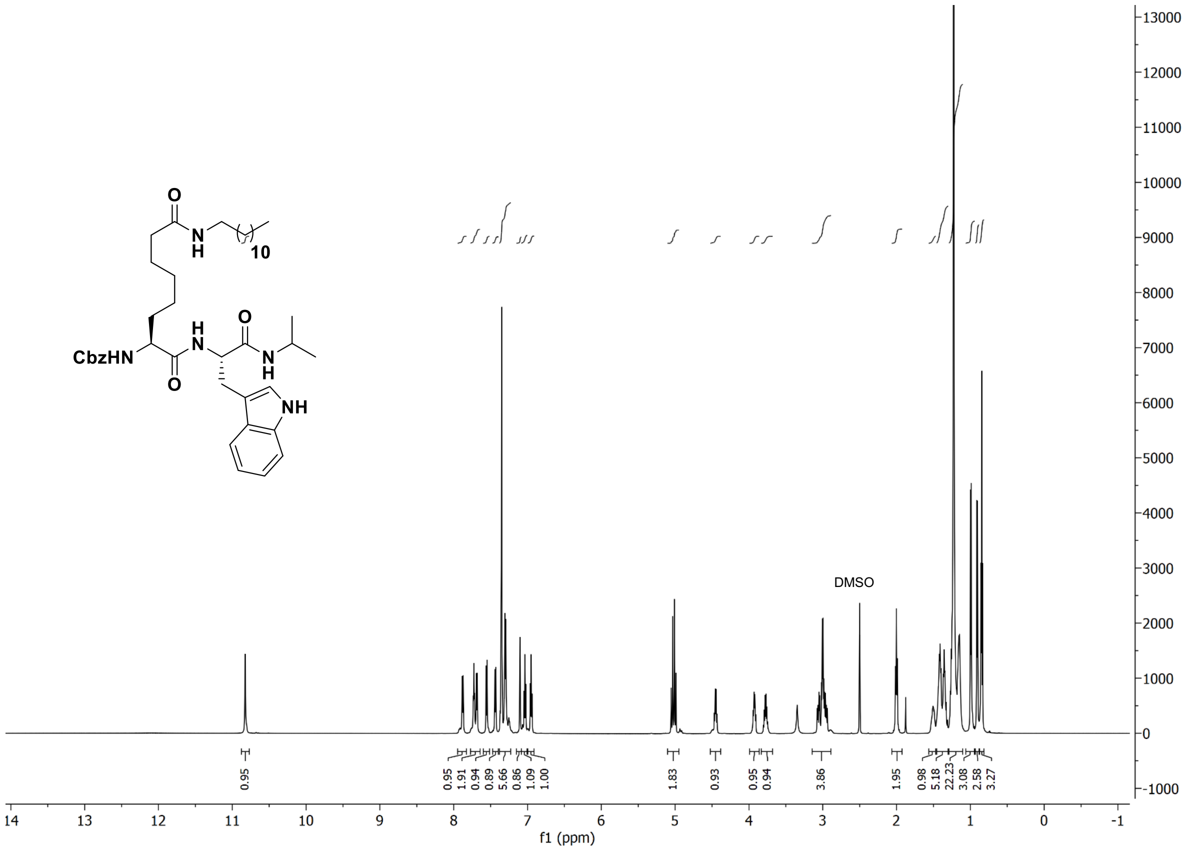

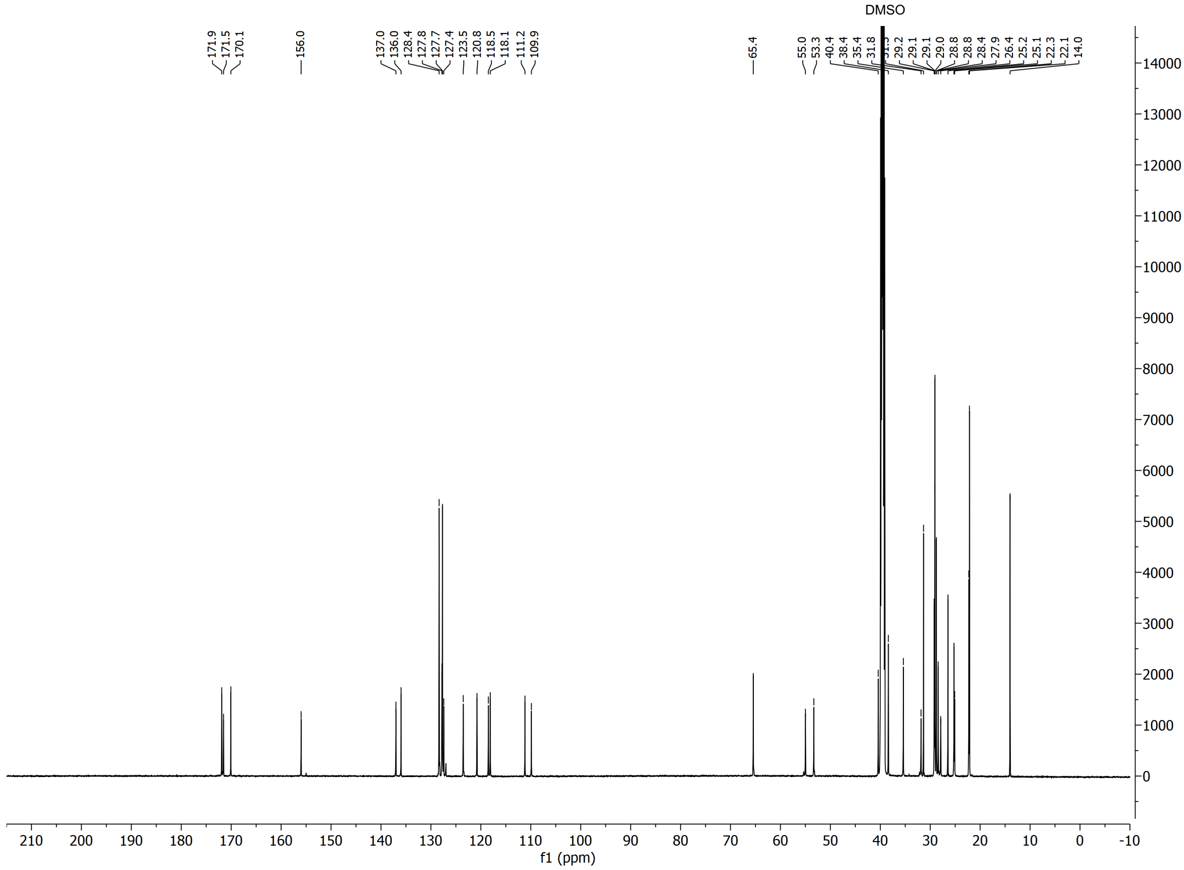

#### $^1$H and $^{13}$C spectra of compound **6**

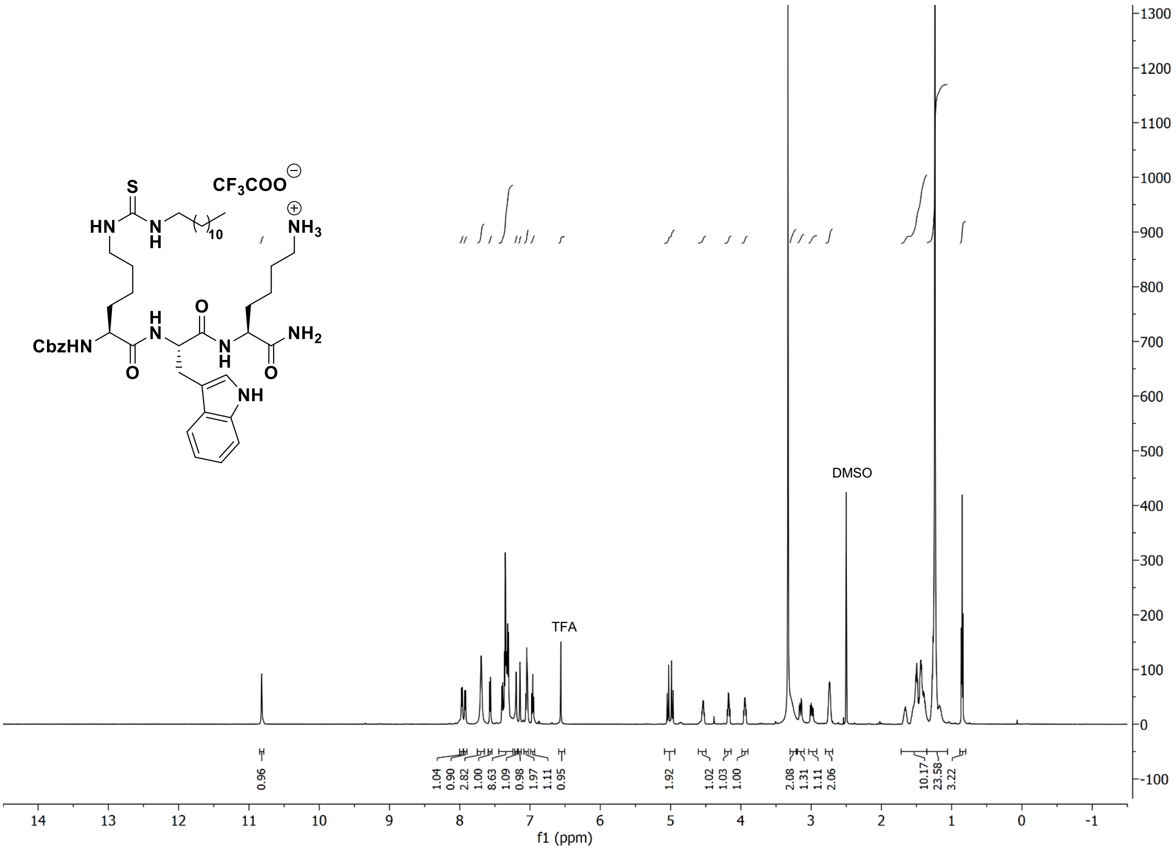

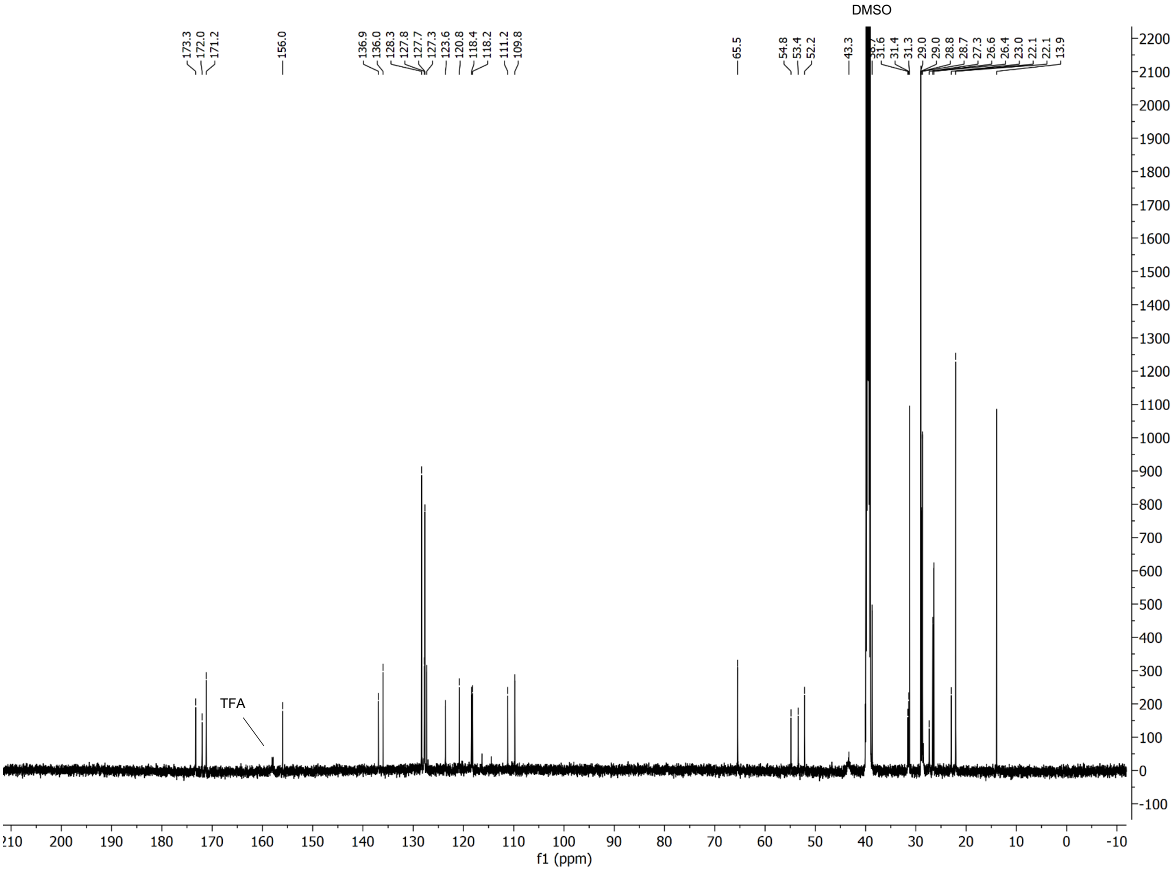

#### $^1$H and $^{13}$C spectra of compound **7**

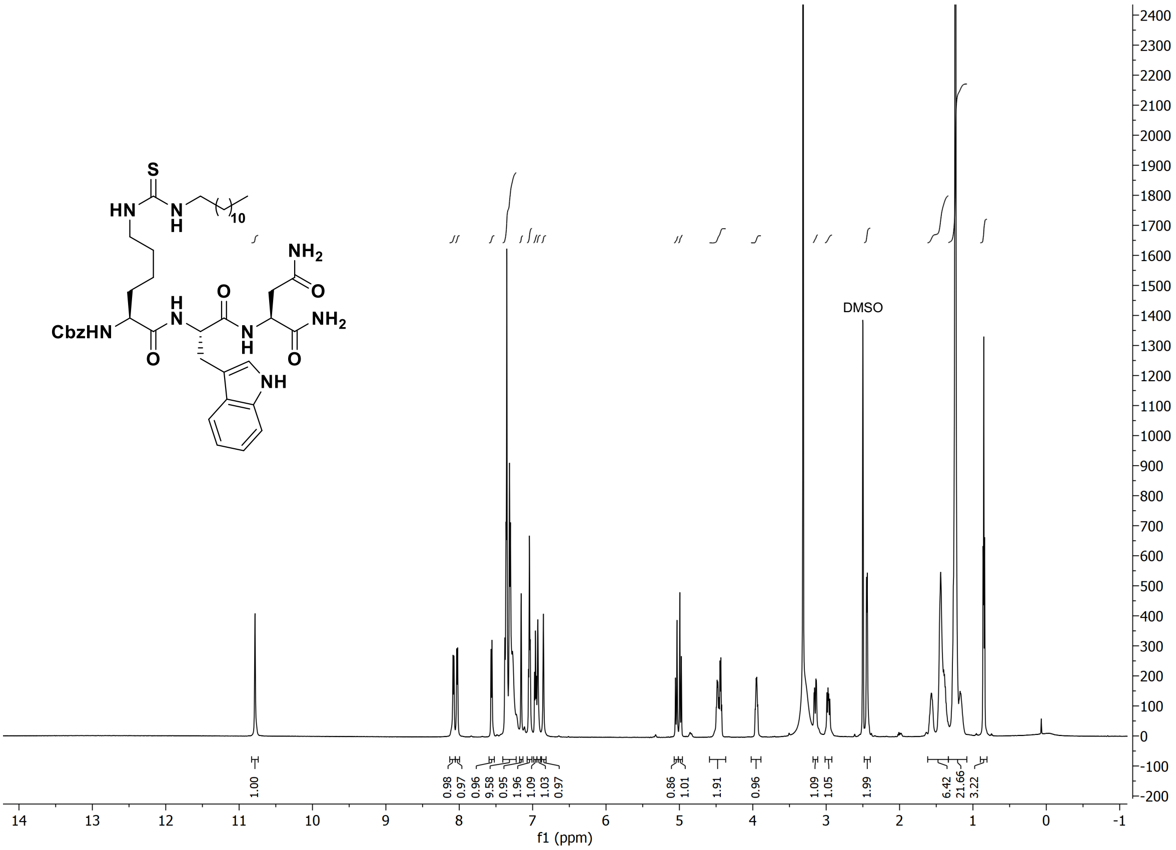

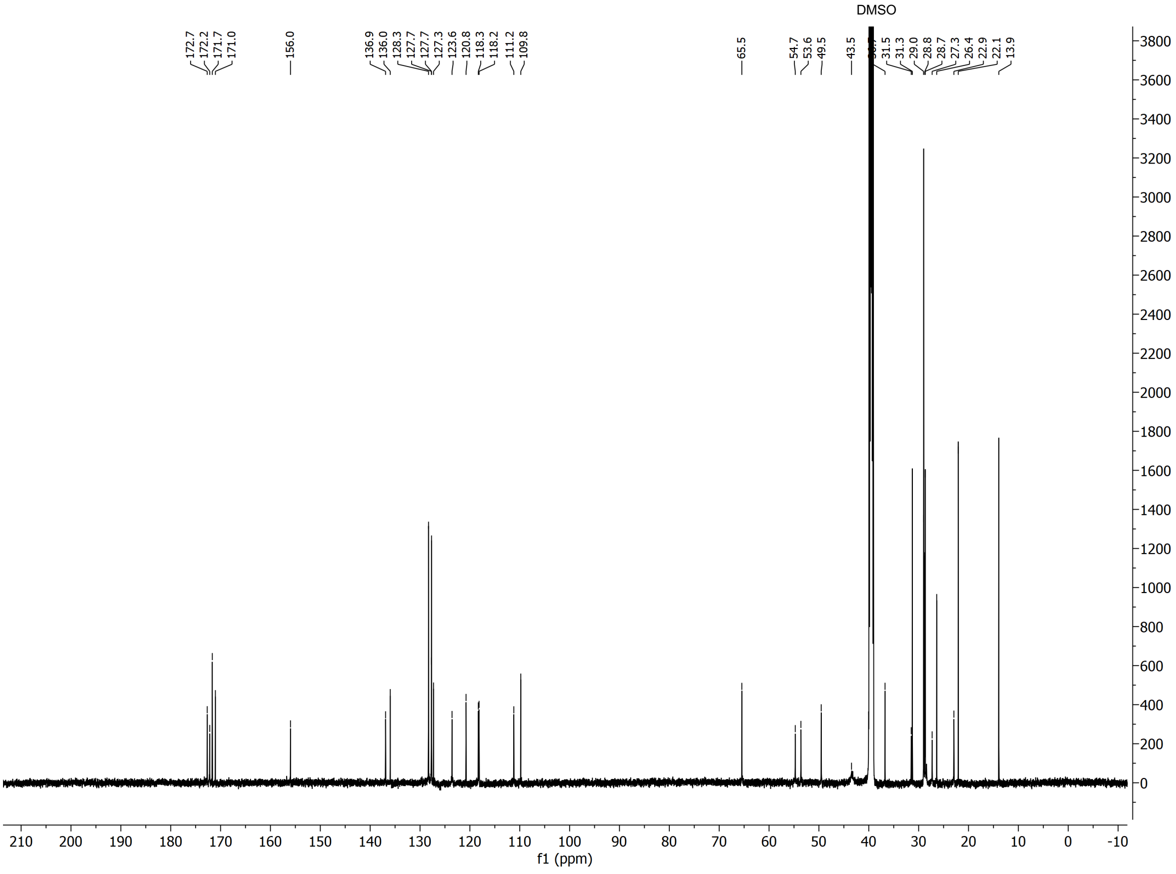

### $^{1}$H and $^{13}$C spectra of compound **8**

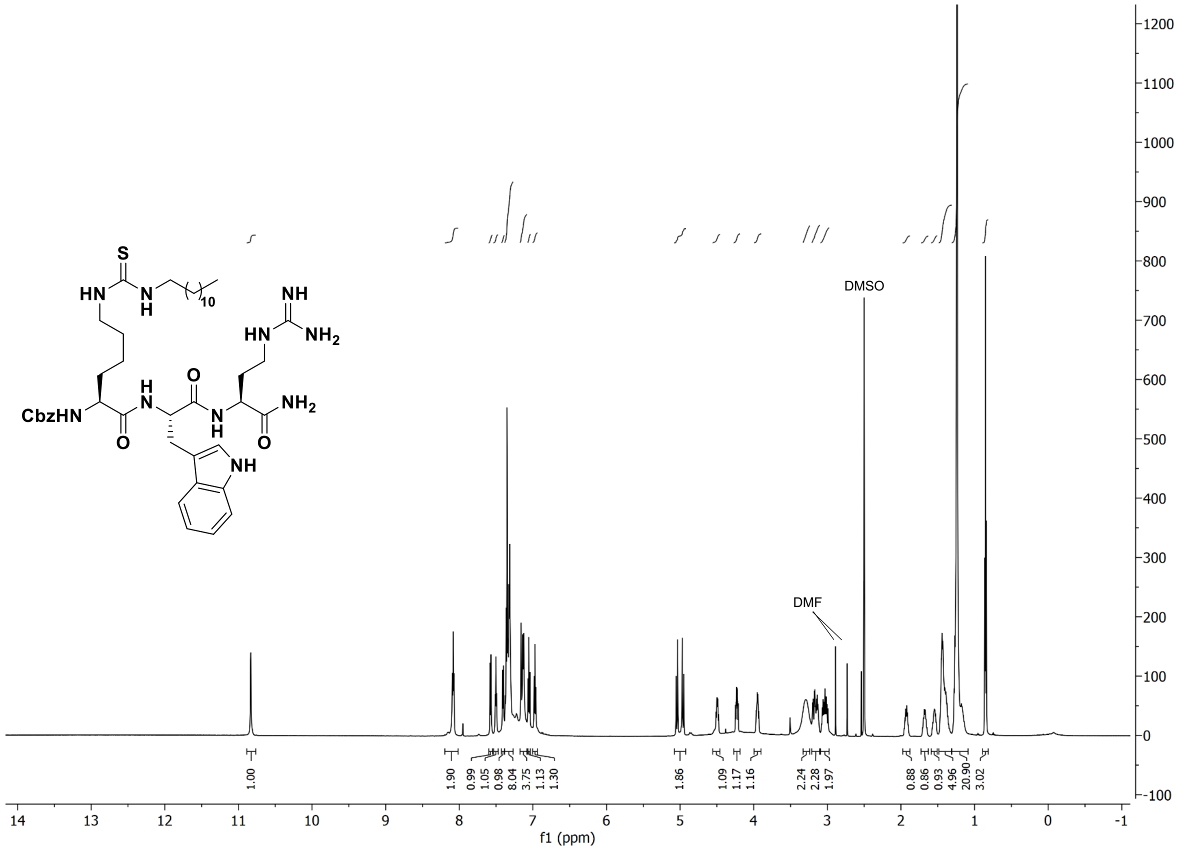

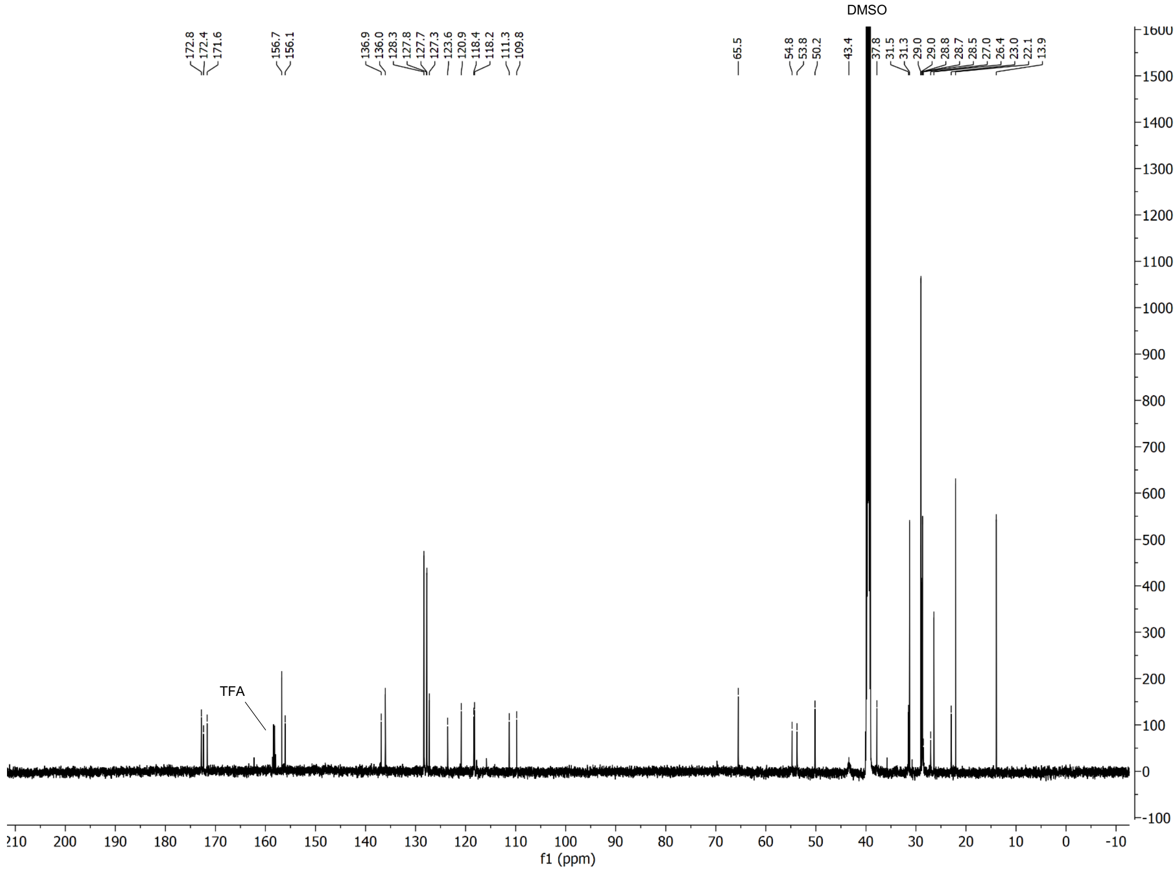

$^{1}$H and $^{13}$C spectra of compound **9**

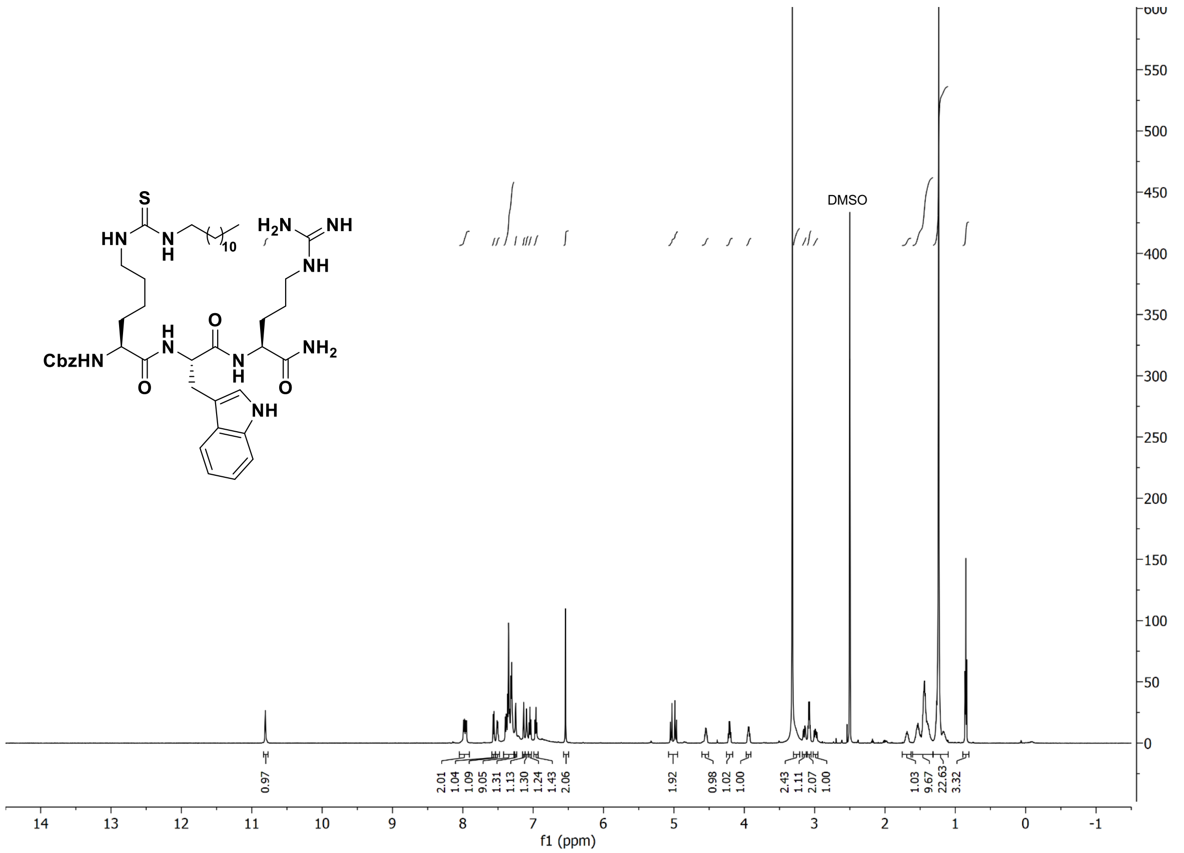

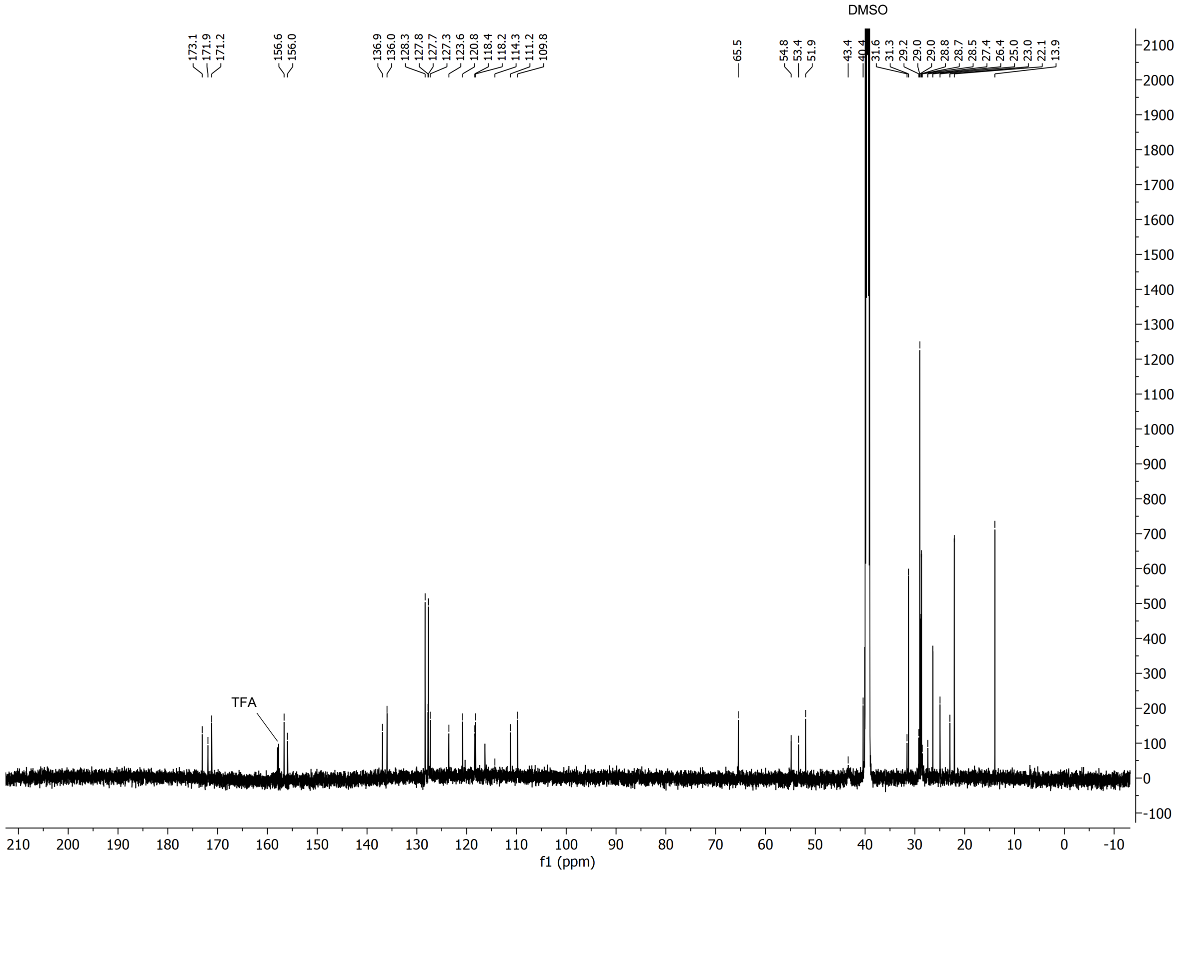

### $^{1}H$ and $^{13}C$ spectra of compound **10**

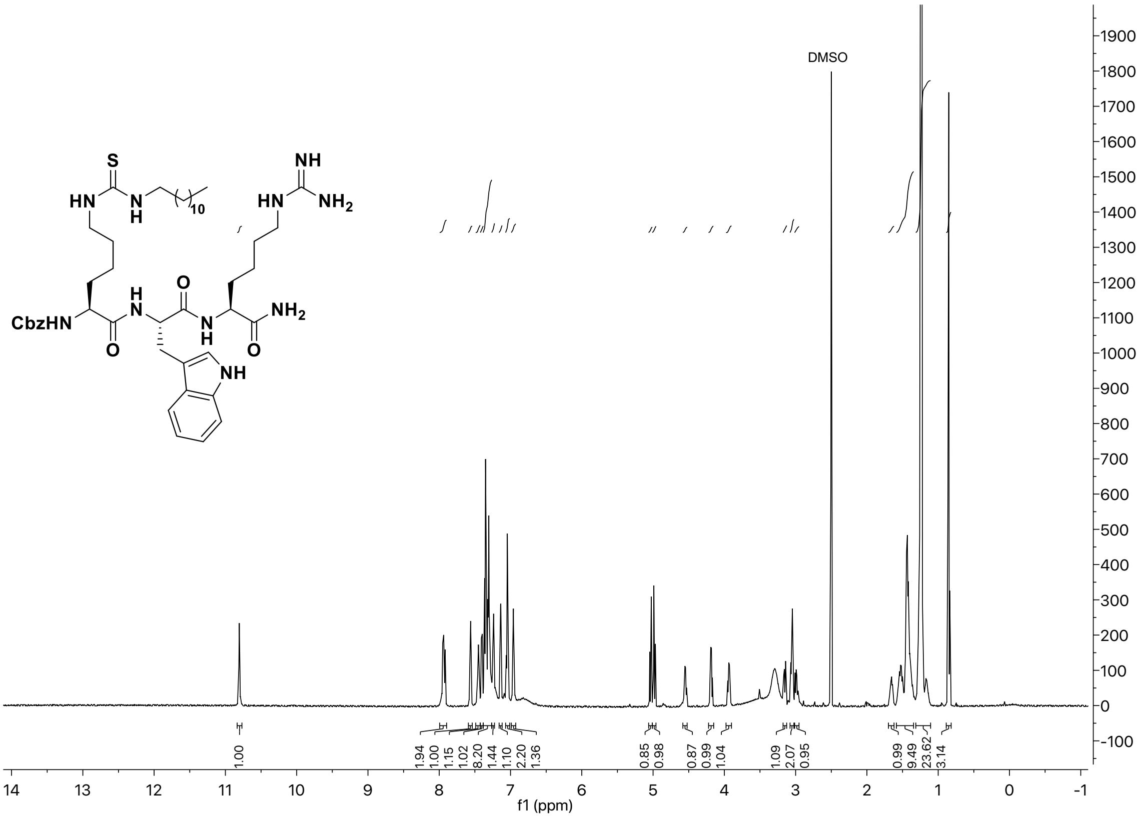

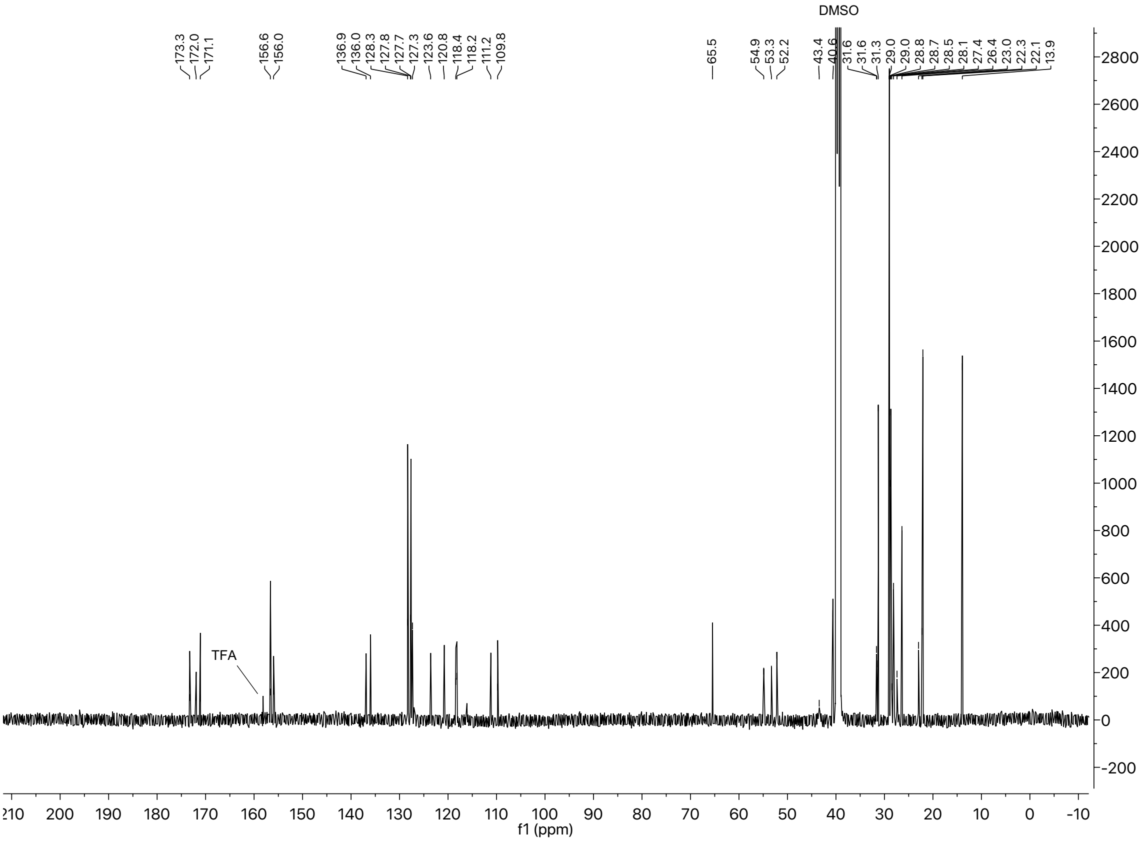

[1]H and [13]C spectra of compound **11**

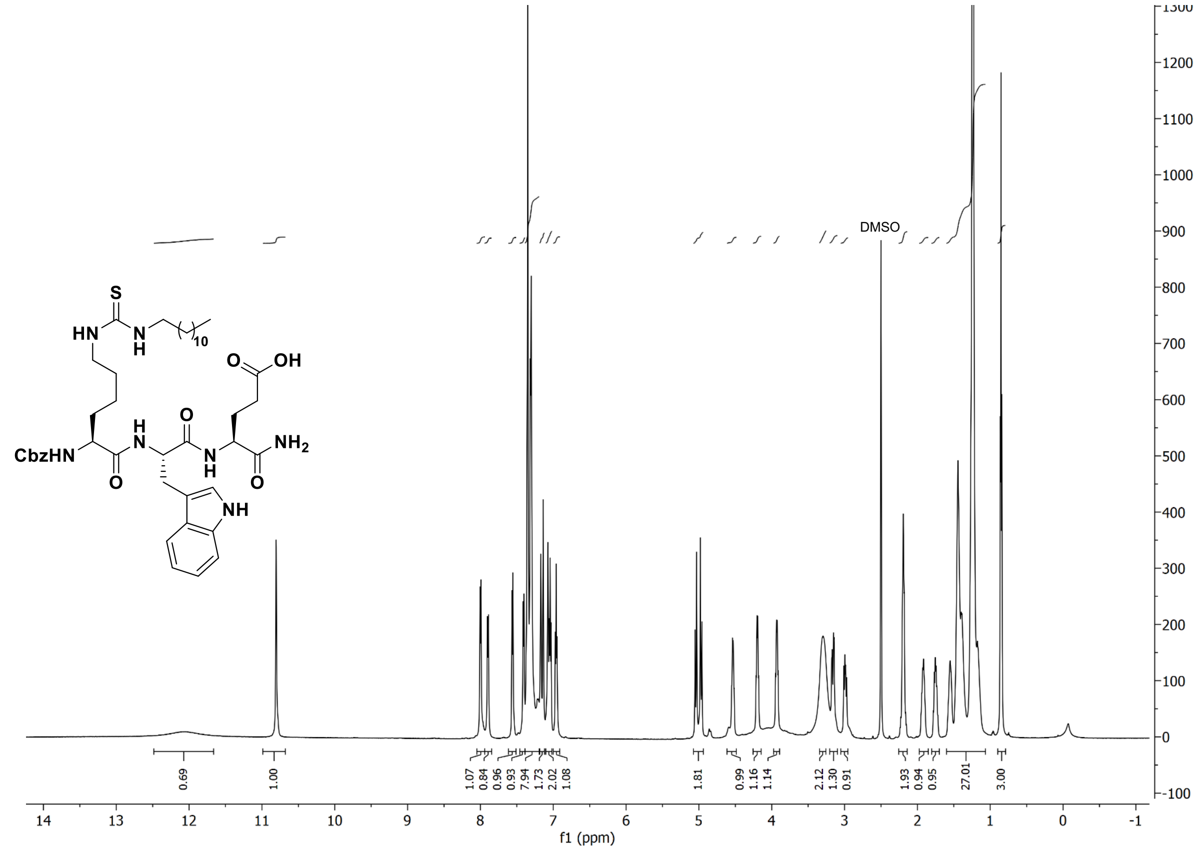

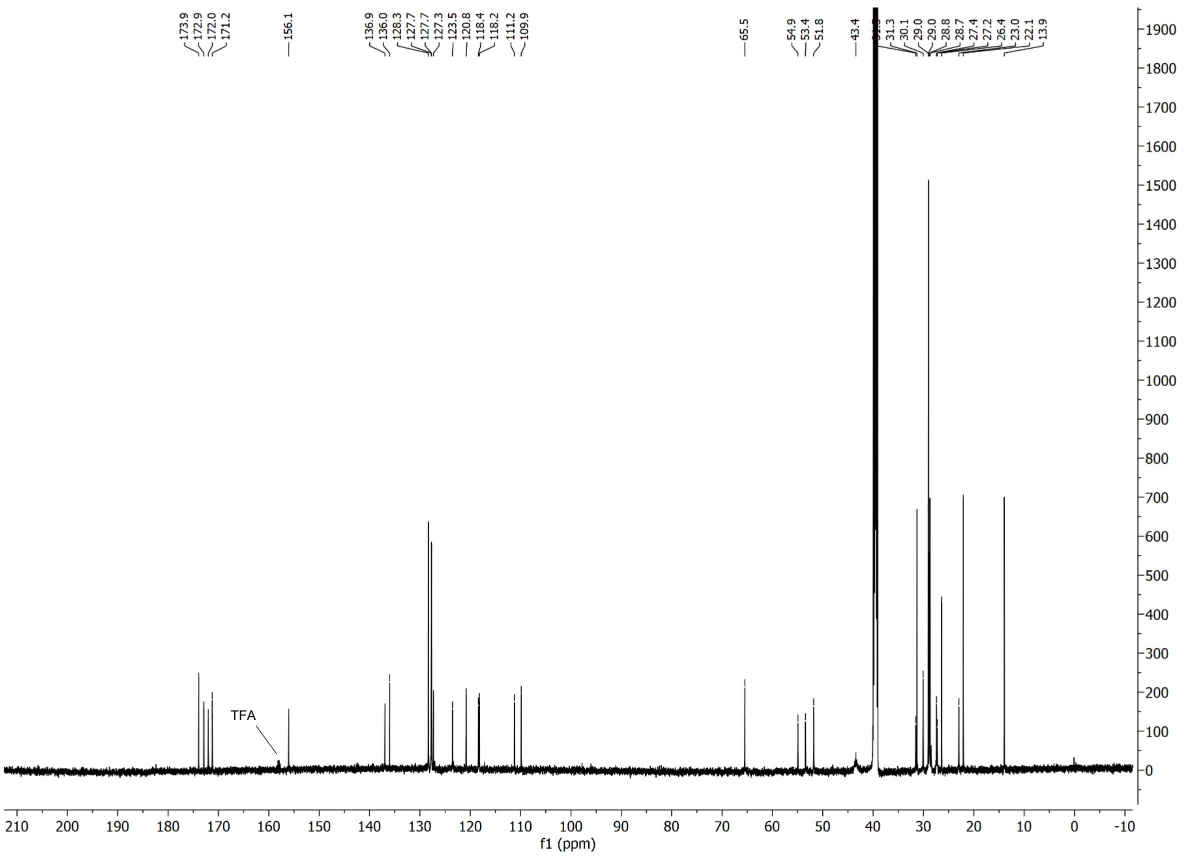

### $^{1}H$ and $^{13}C$ spectra of compound **12**

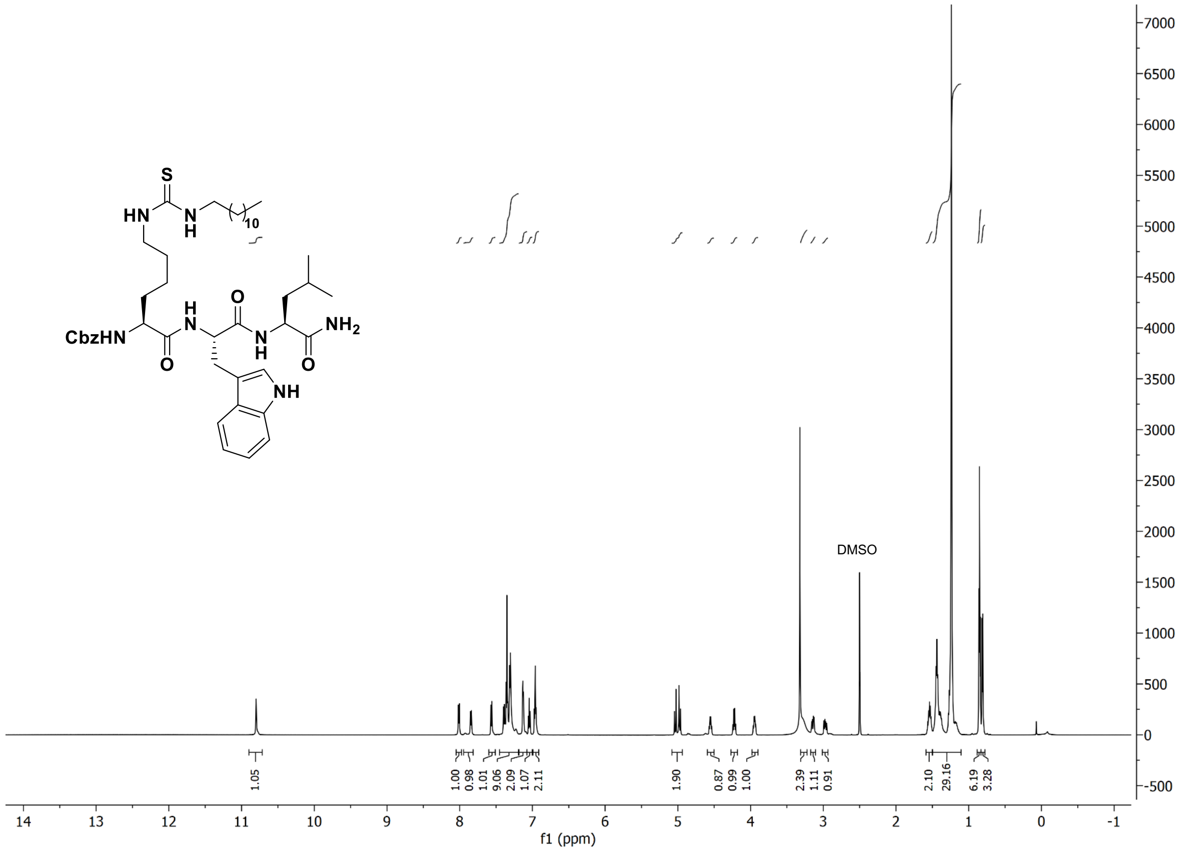

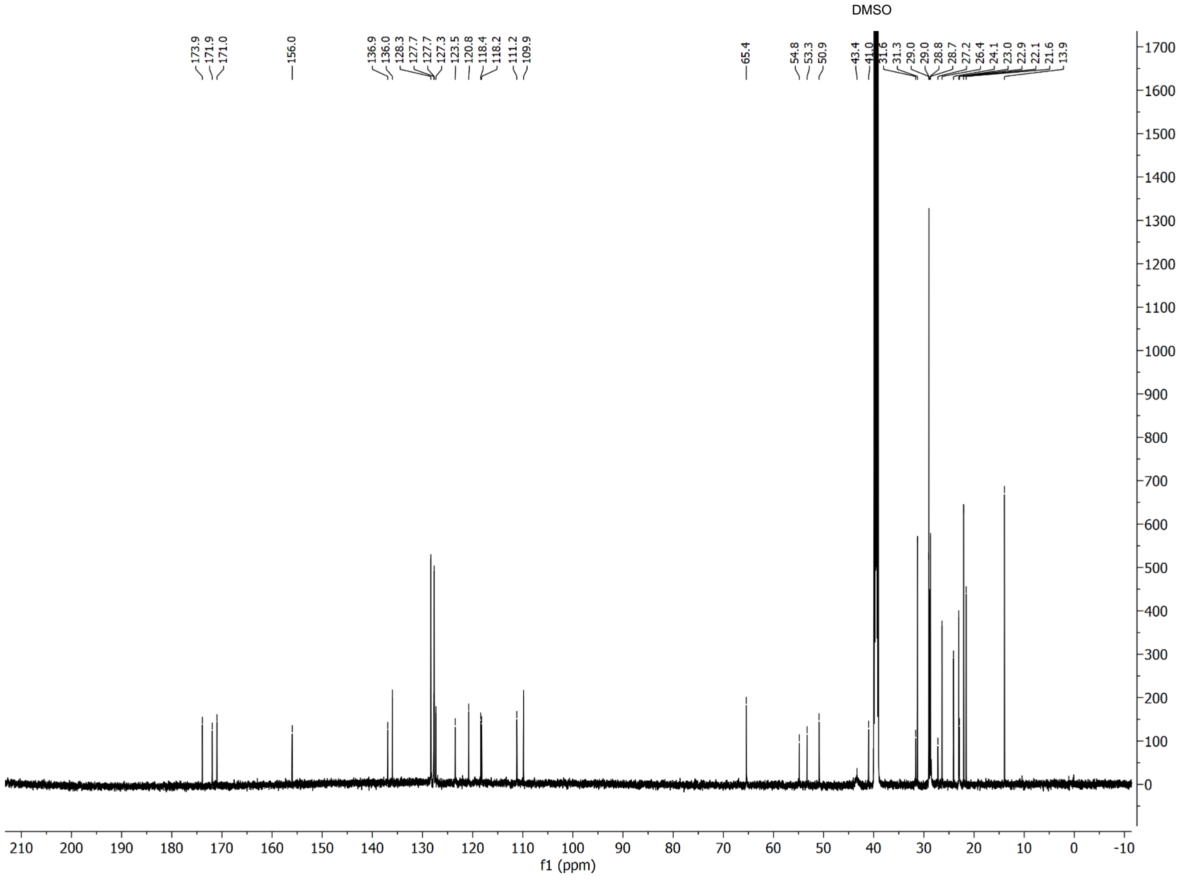

[1]H and [13]C spectra of compound **13**

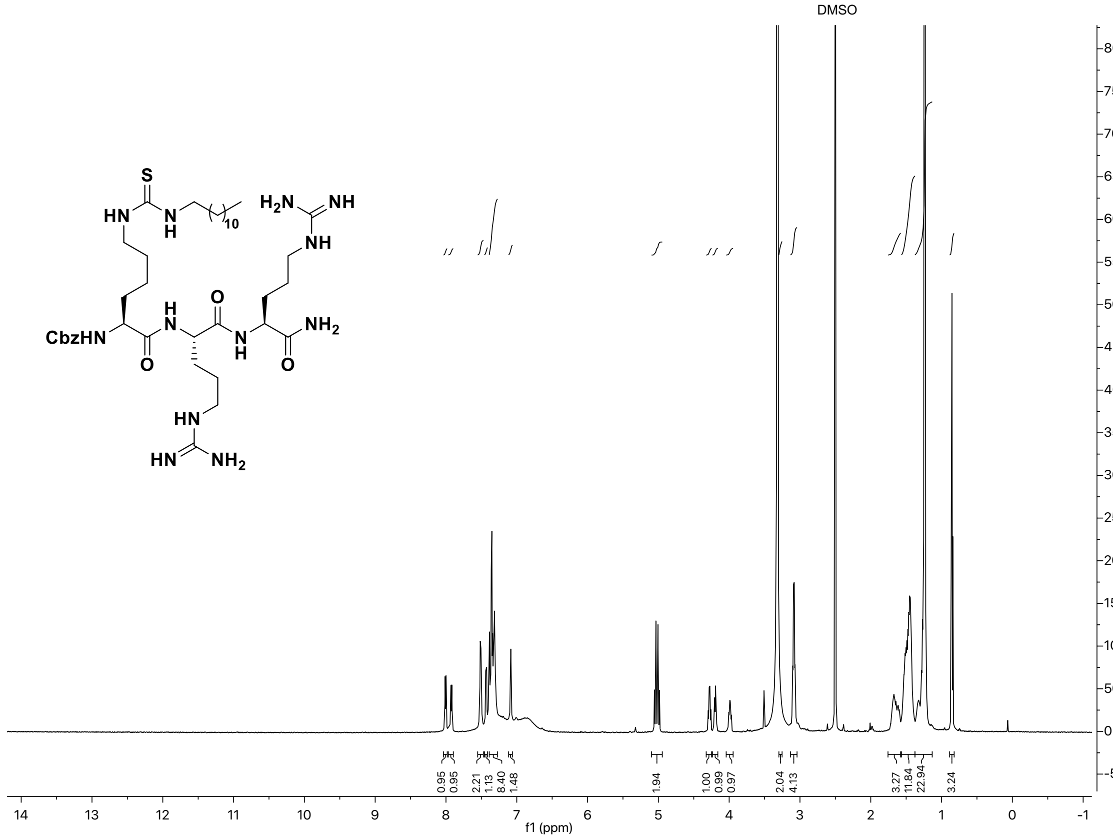

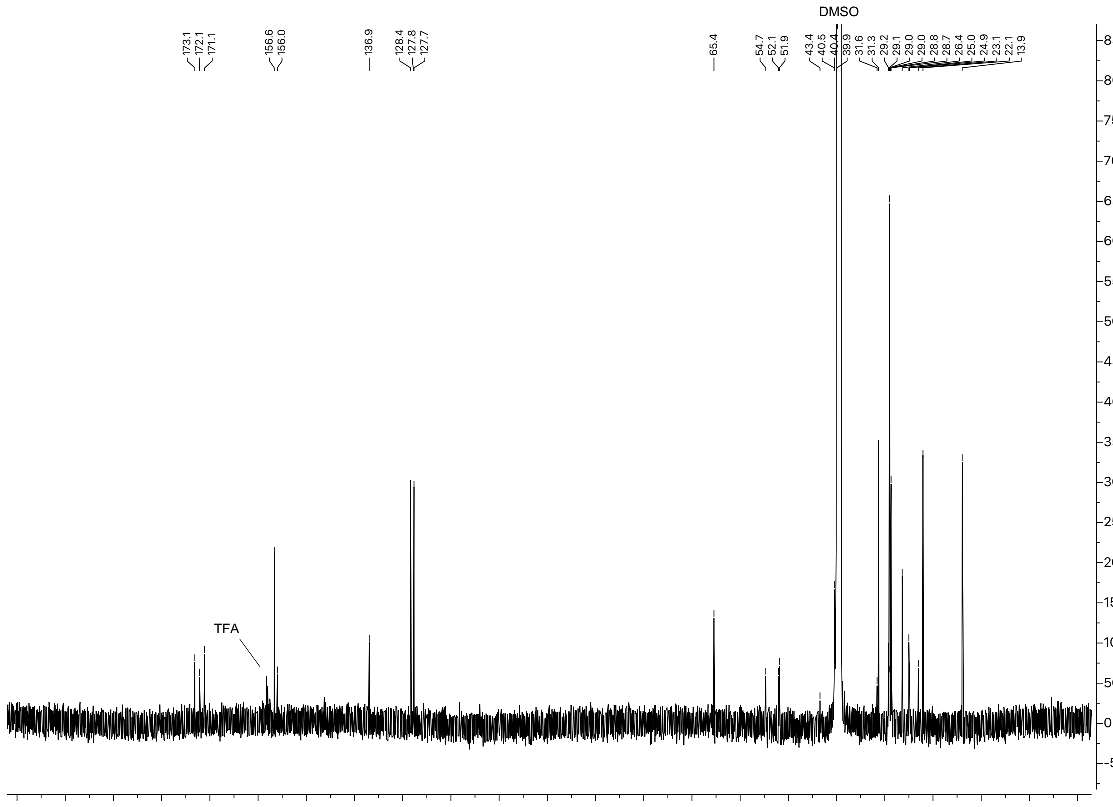

### $^{1}H$ and $^{13}C$ spectra of compound **14**

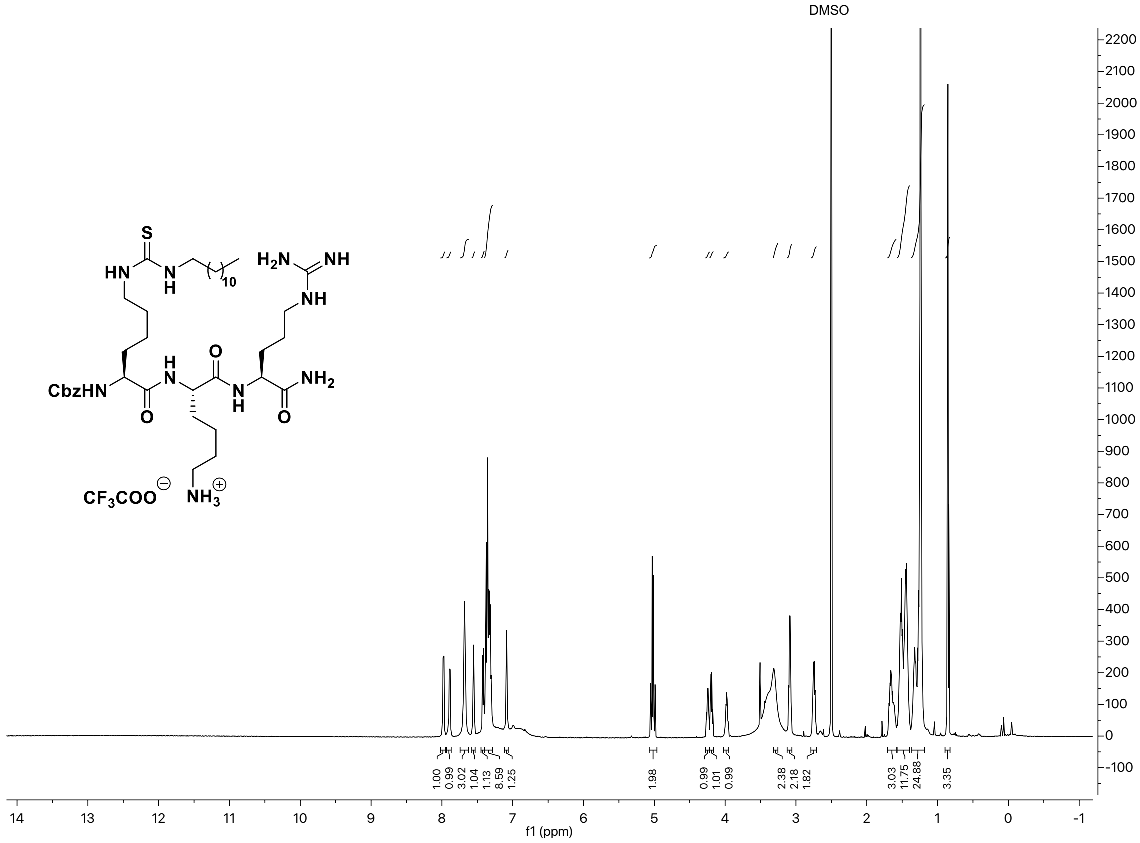

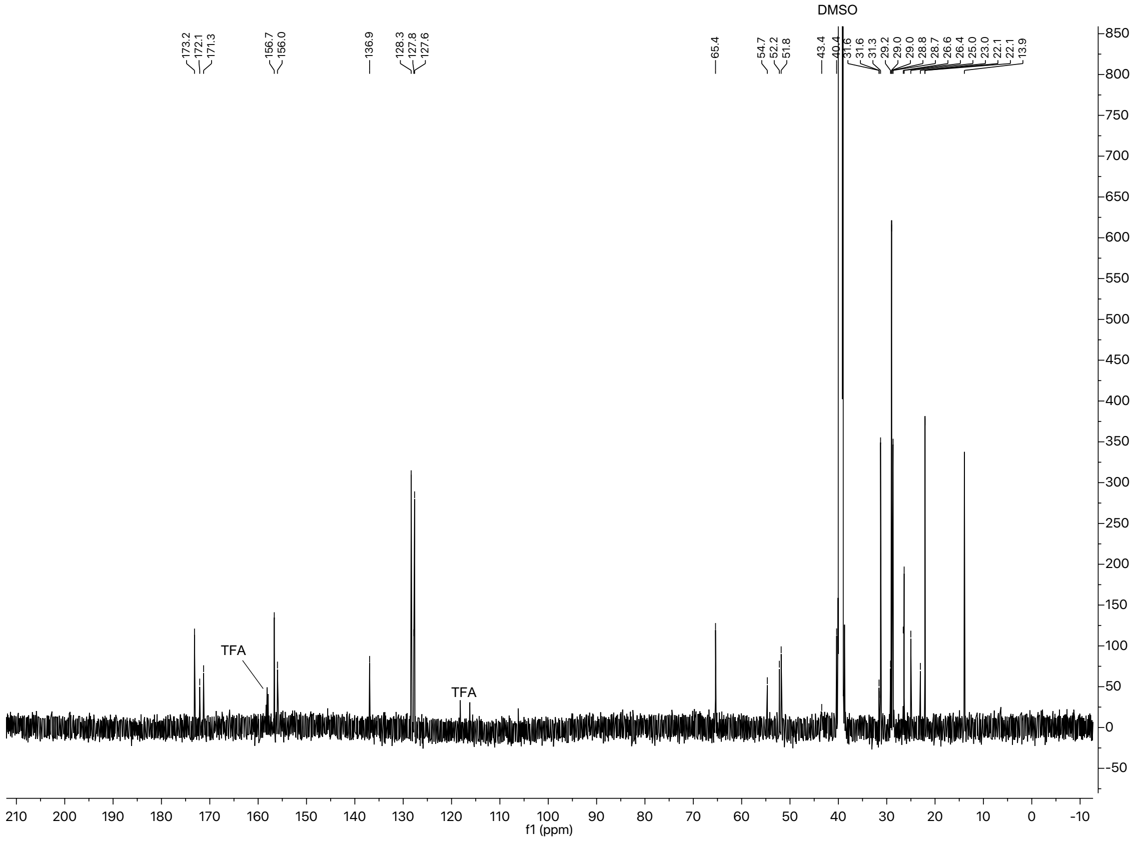

#### $^{1}H$ and $^{13}C$ spectra of compound **15**

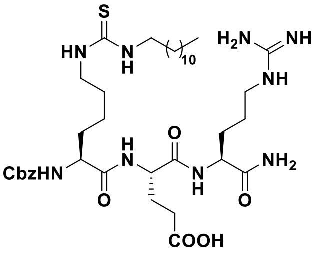

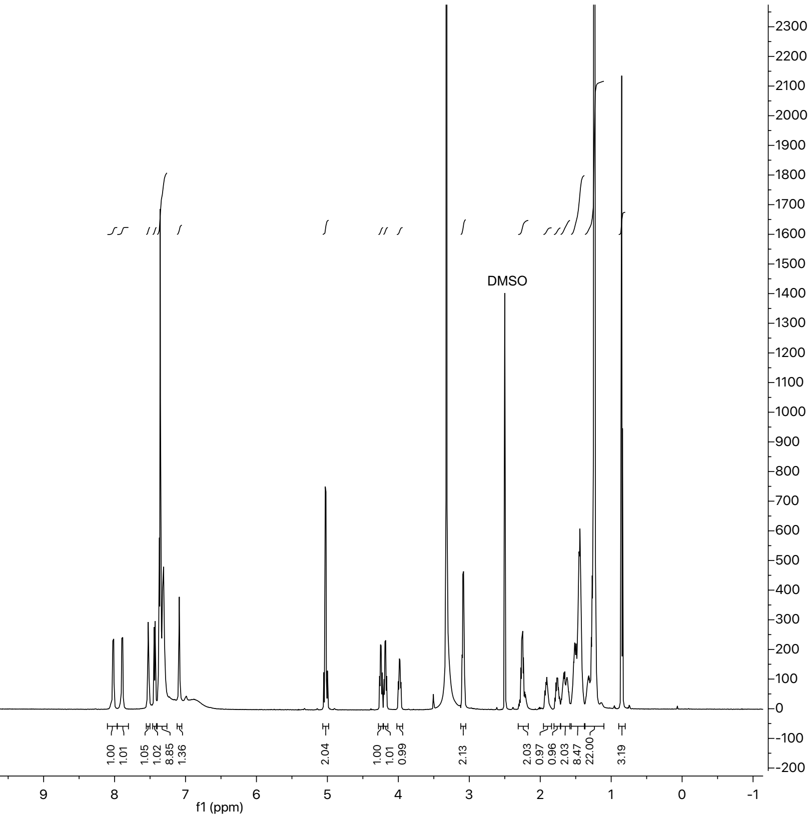

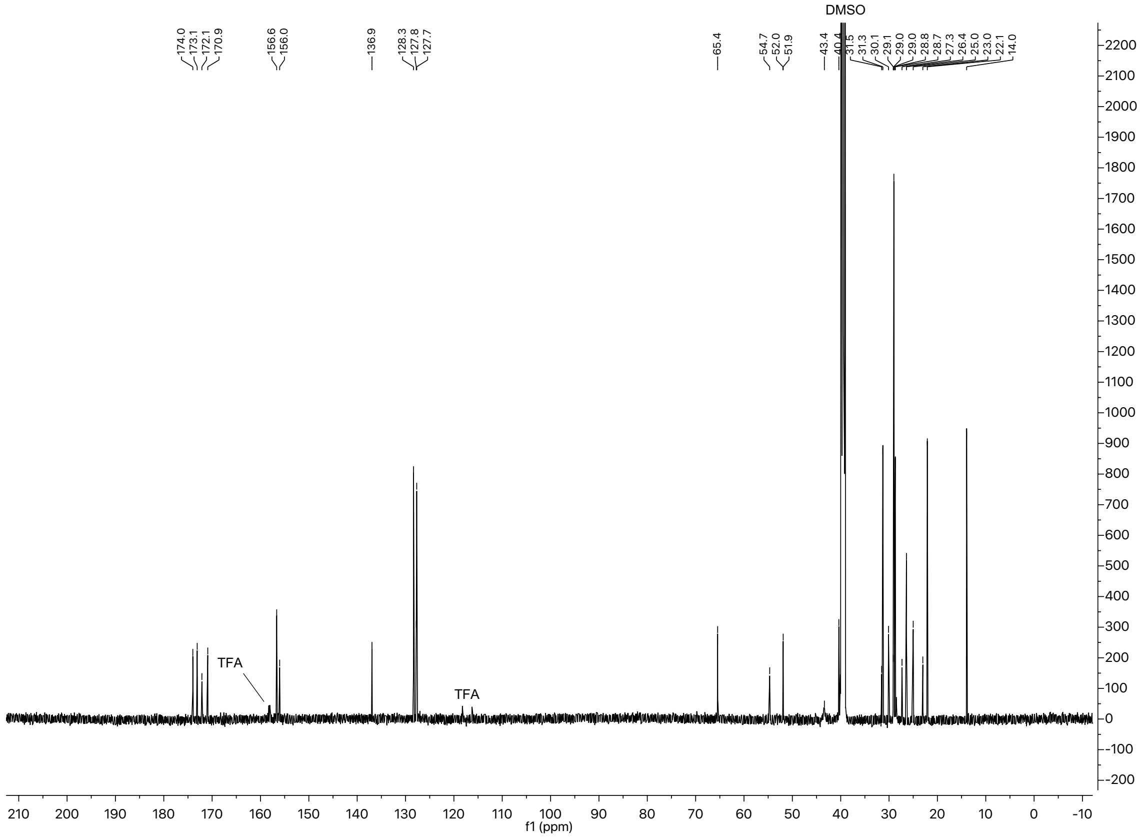

### $^{1}H$ and $^{13}C$ spectra of compound **16**

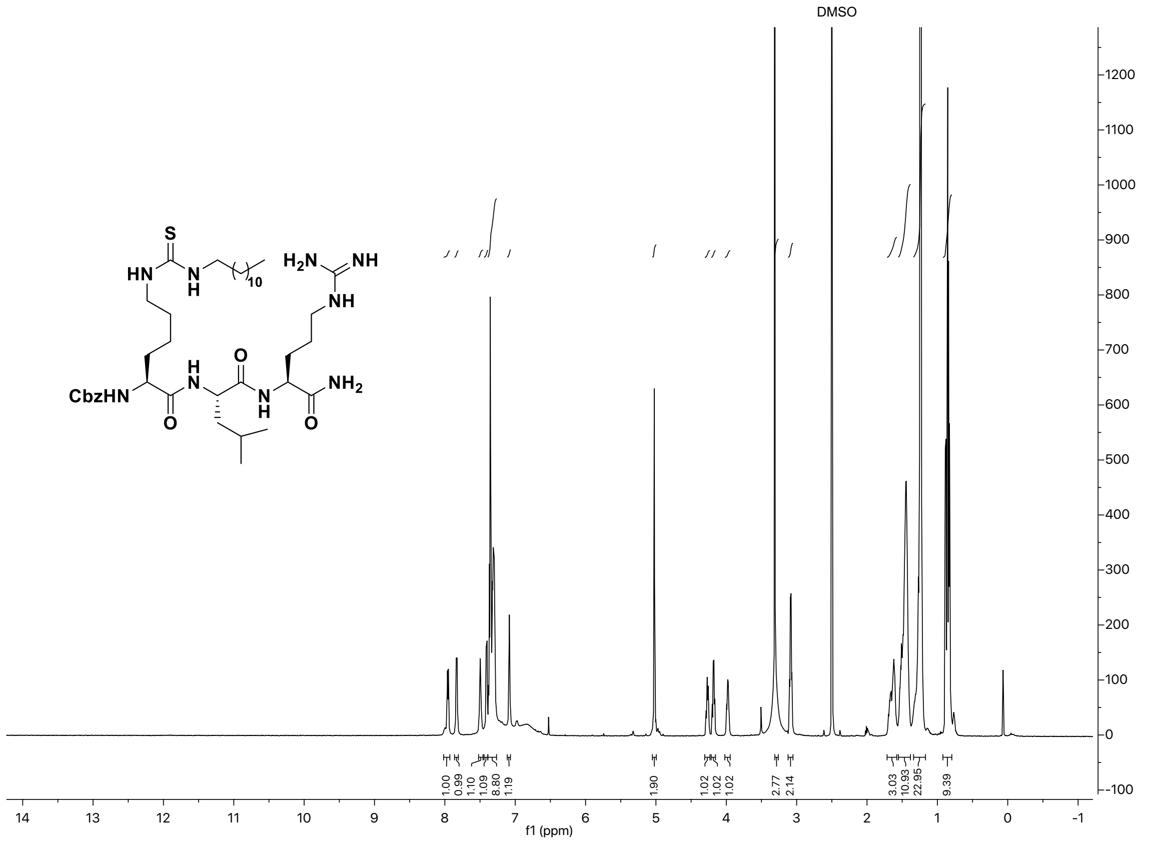

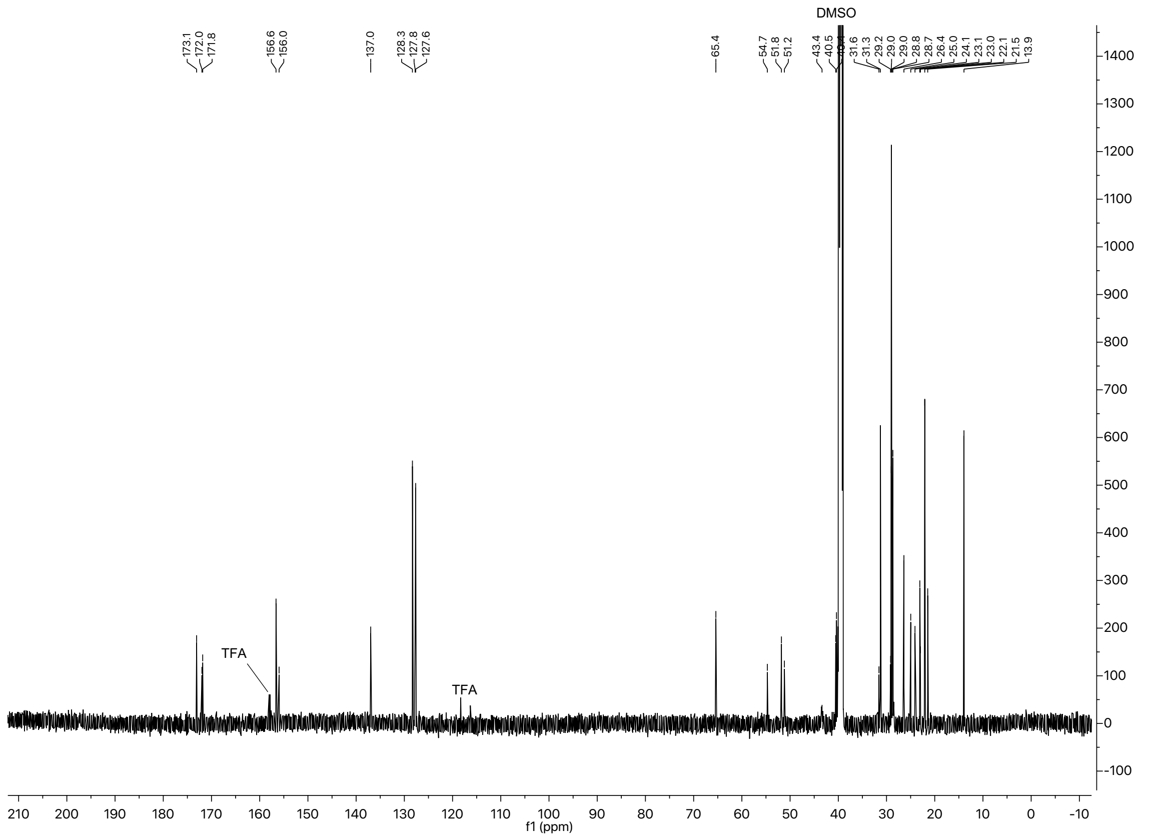

#### [1]H and [13]C spectra of compound **17**

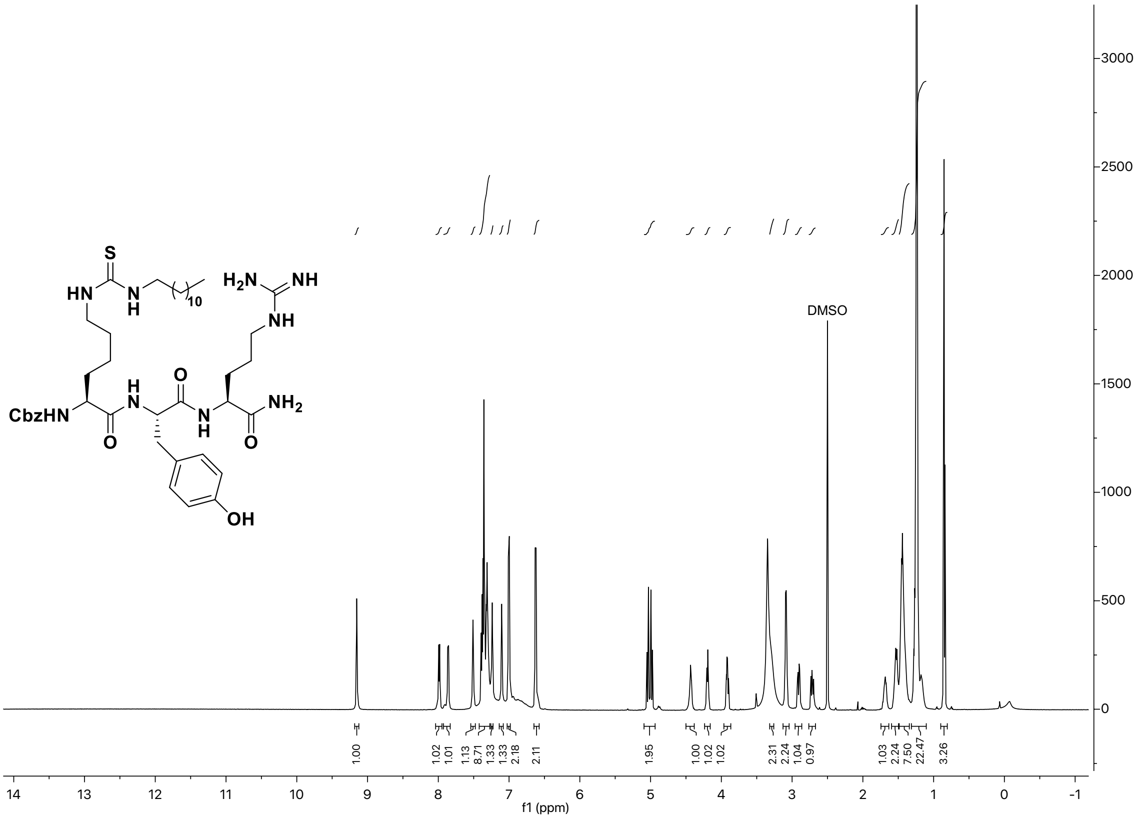

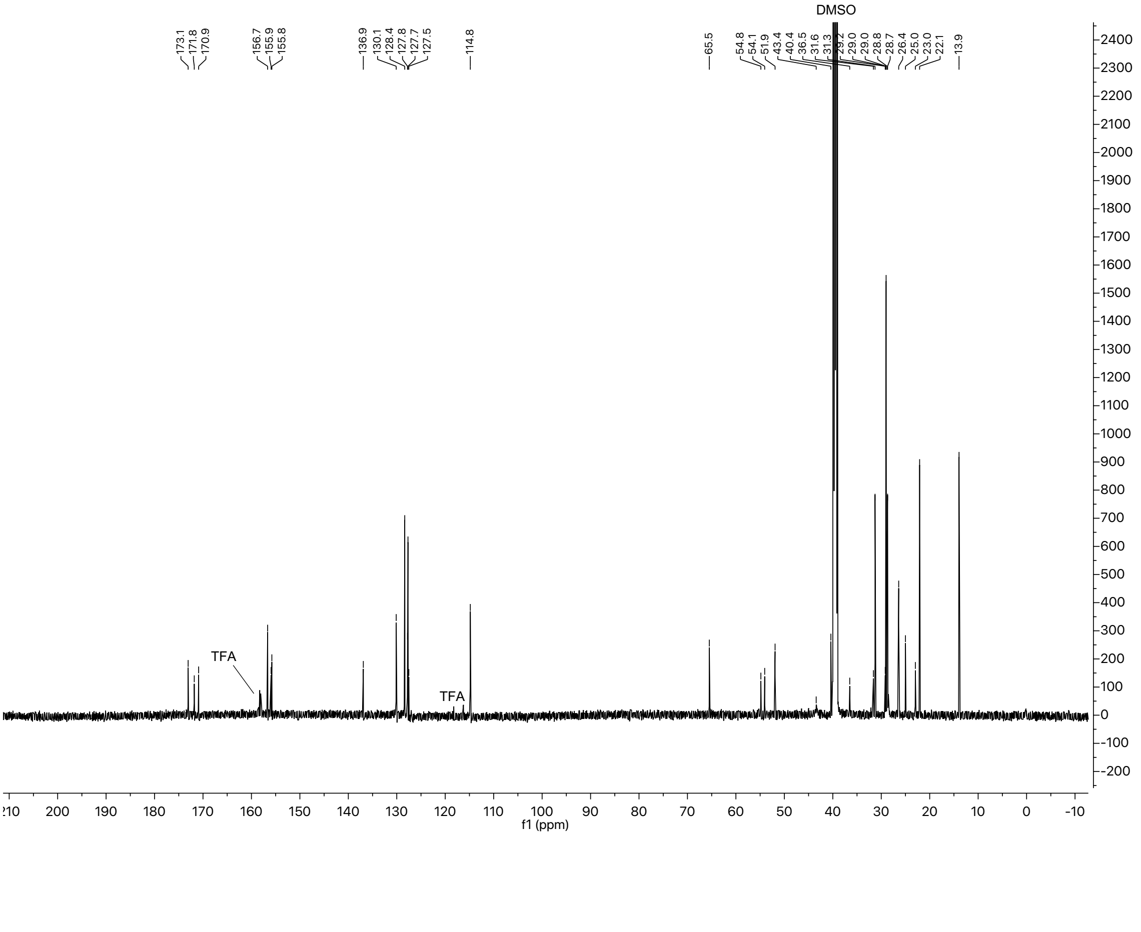

#### $^{1}H$ and $^{13}C$ spectra of compound **18**

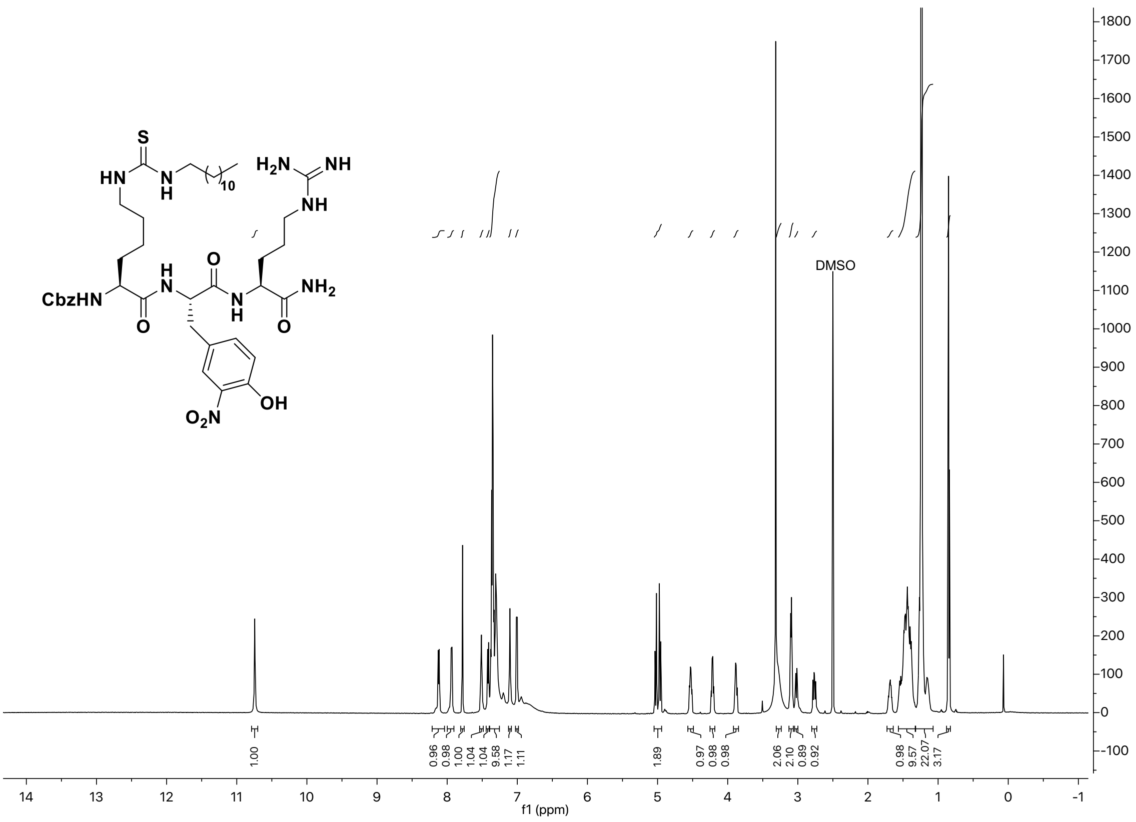

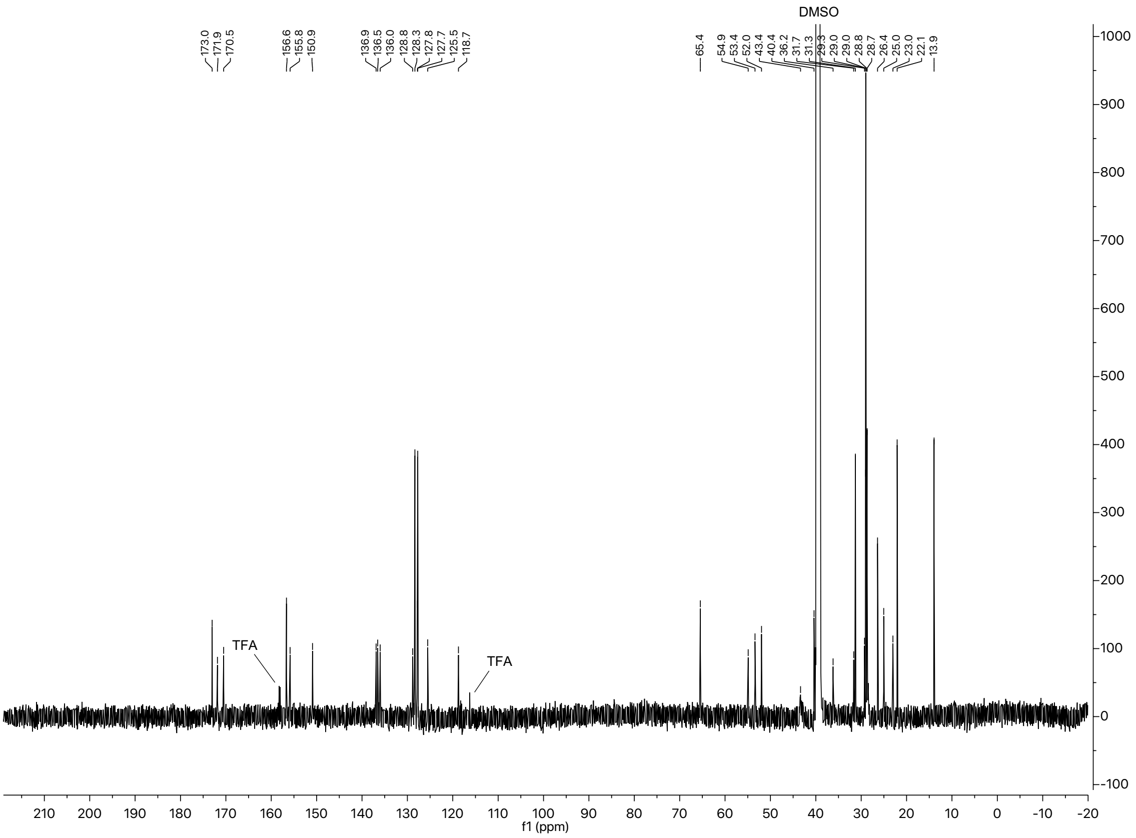

### $^{1}H$ and $^{13}C$ spectra of compound **19**

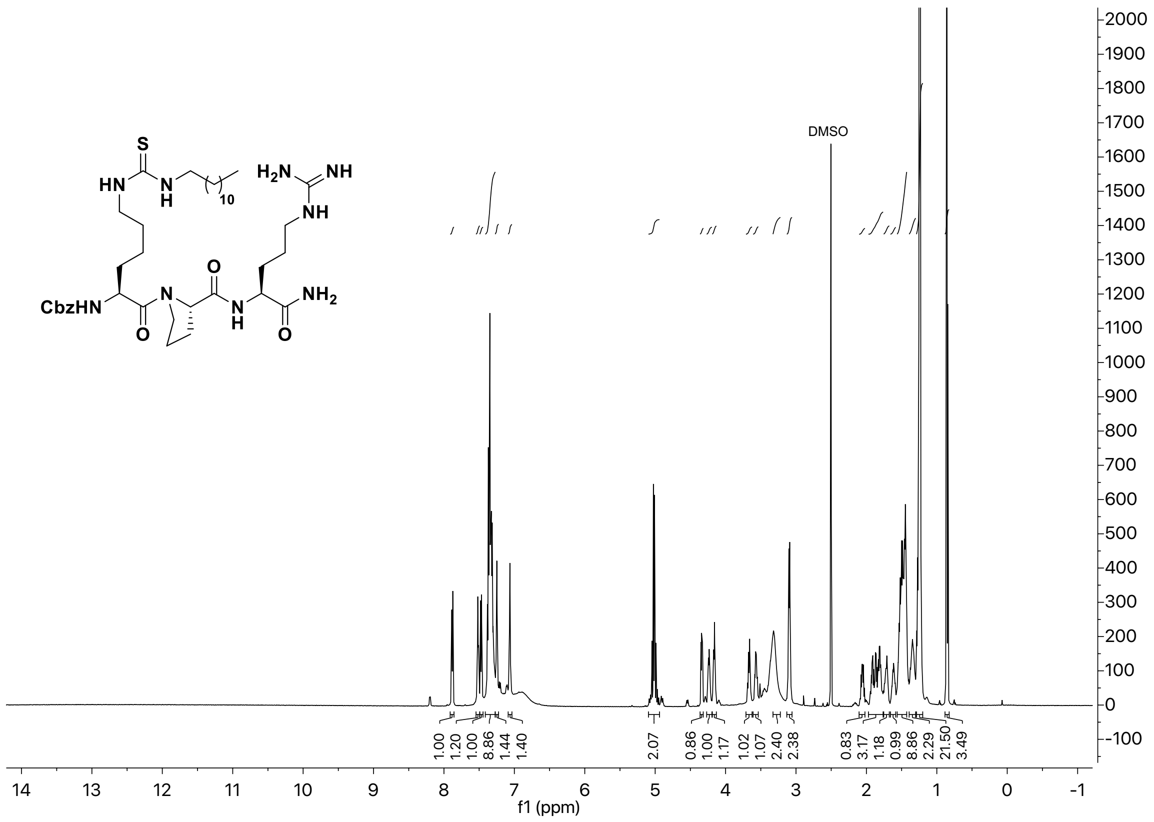

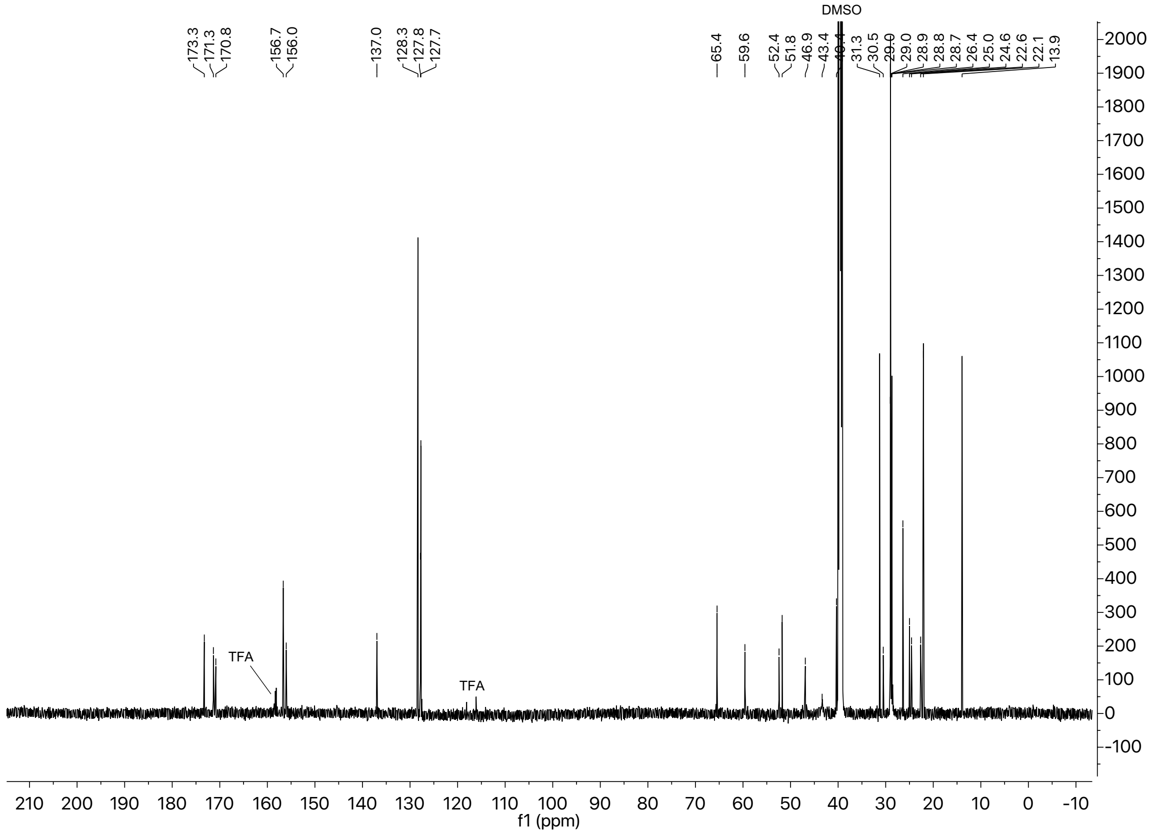

#### [1]H and [13]C spectra of compound **20**

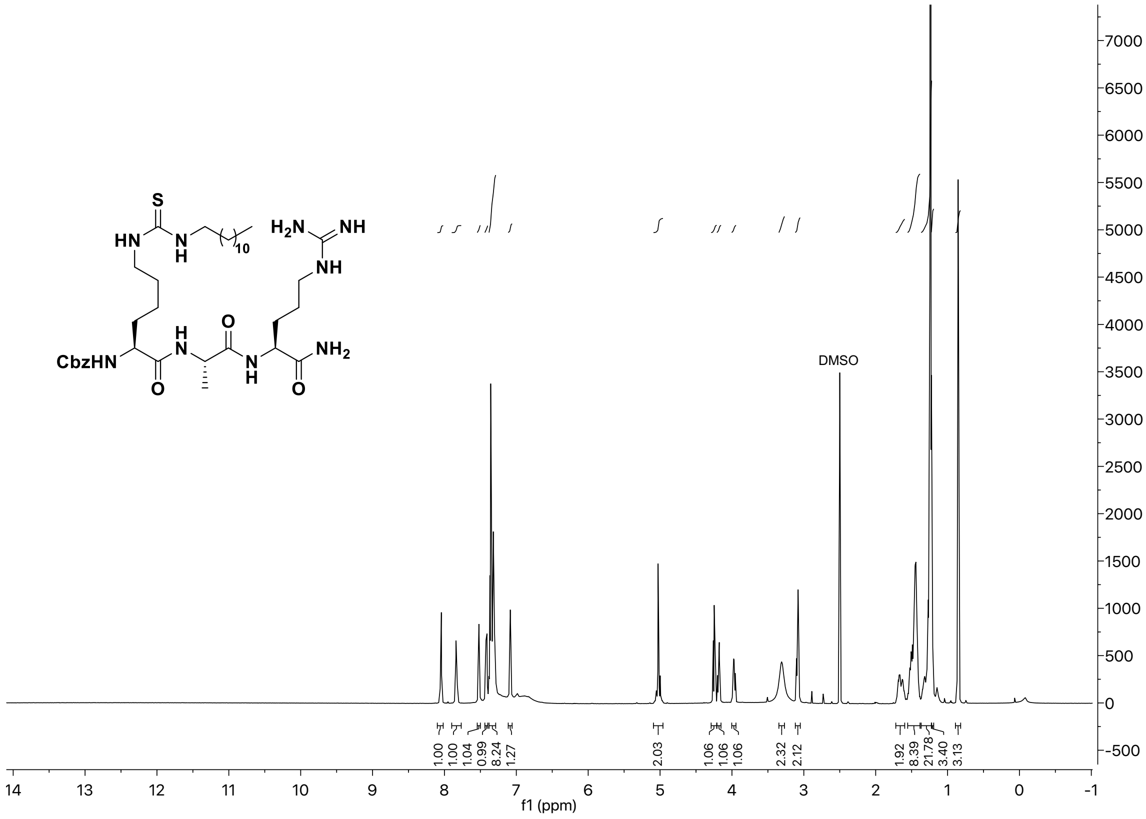

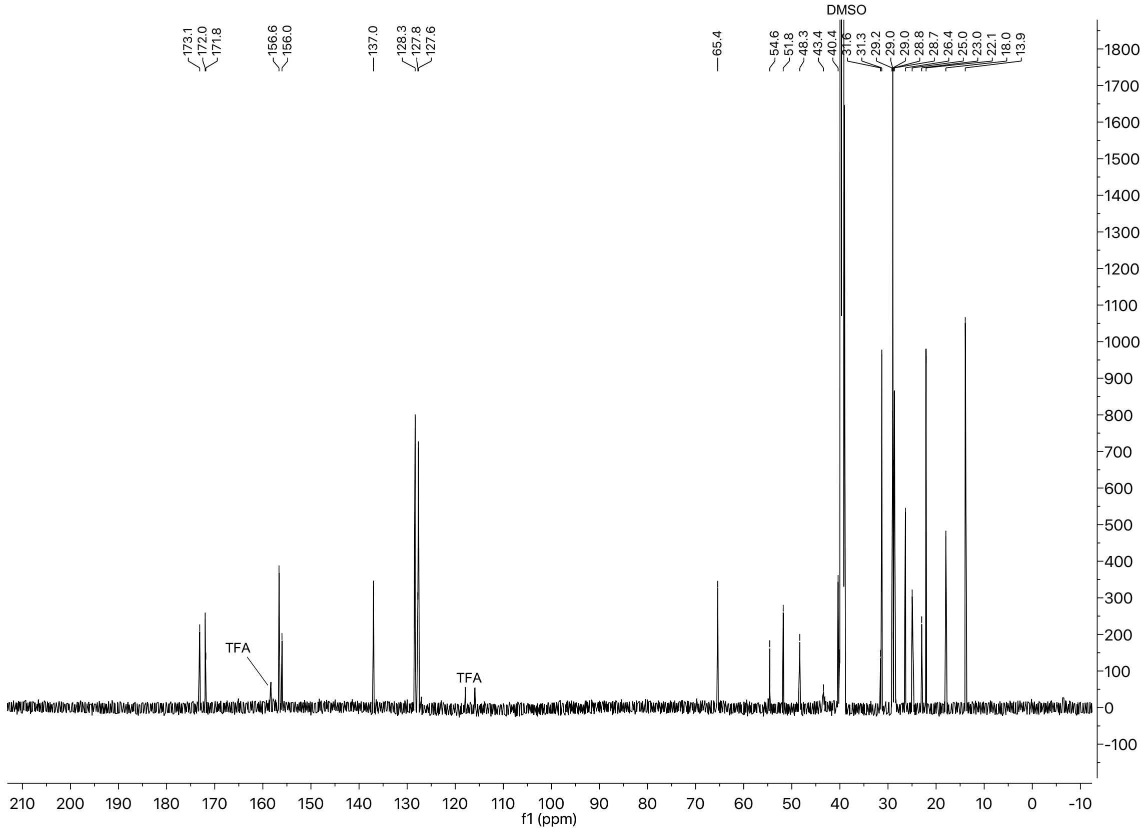

### $^{1}H$ and $^{13}C$ spectra of compound **21**

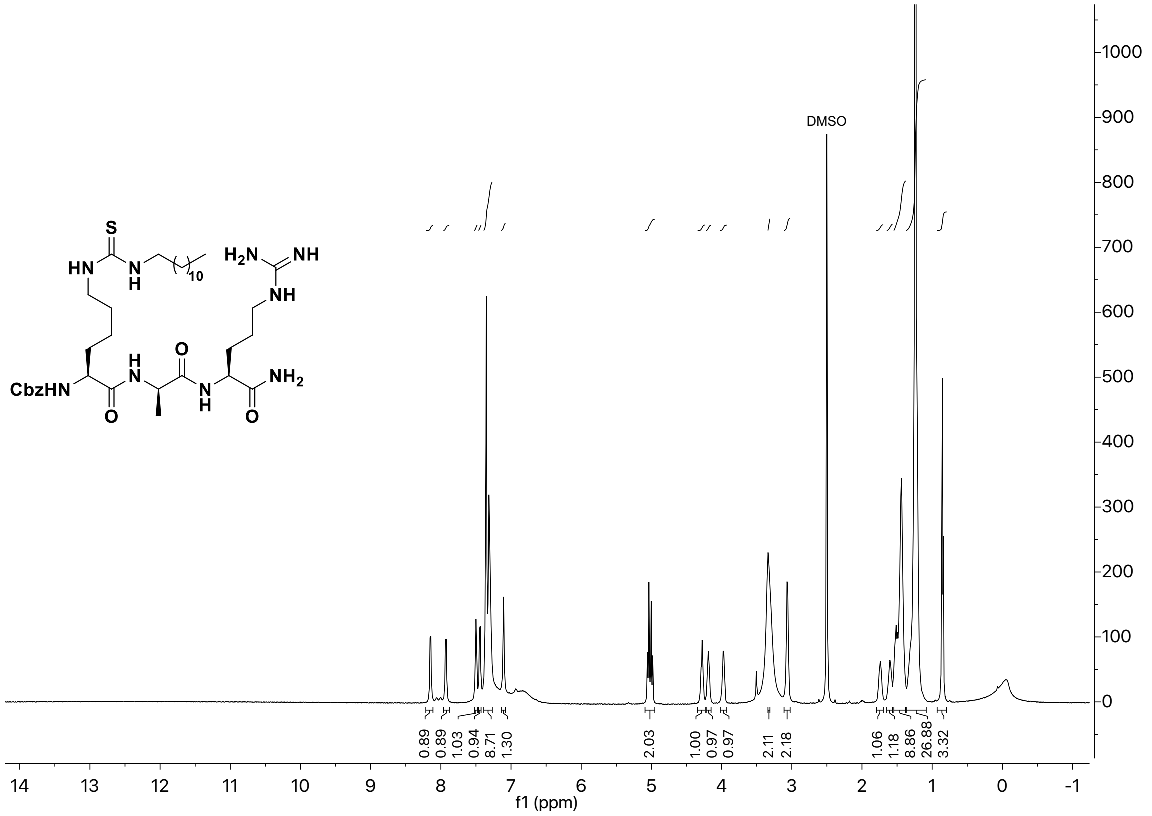

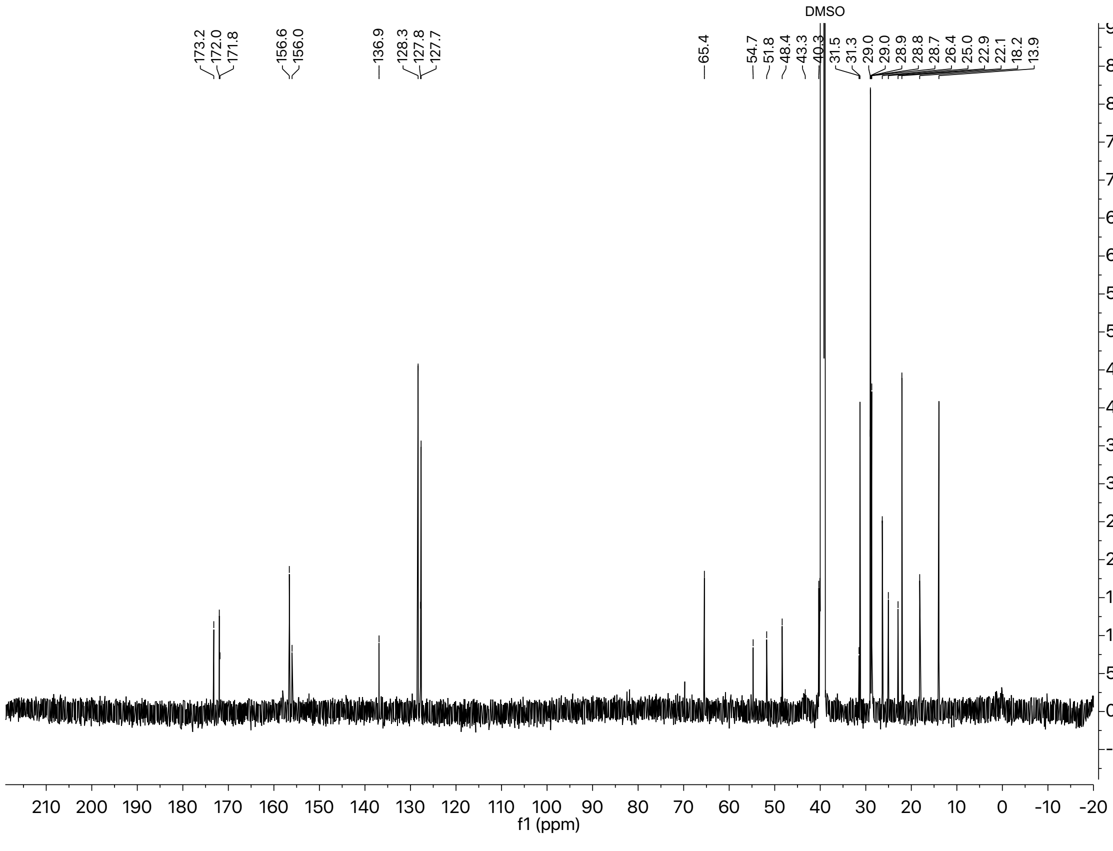

#### $^{1}$H and $^{13}$C spectra of compound **22**

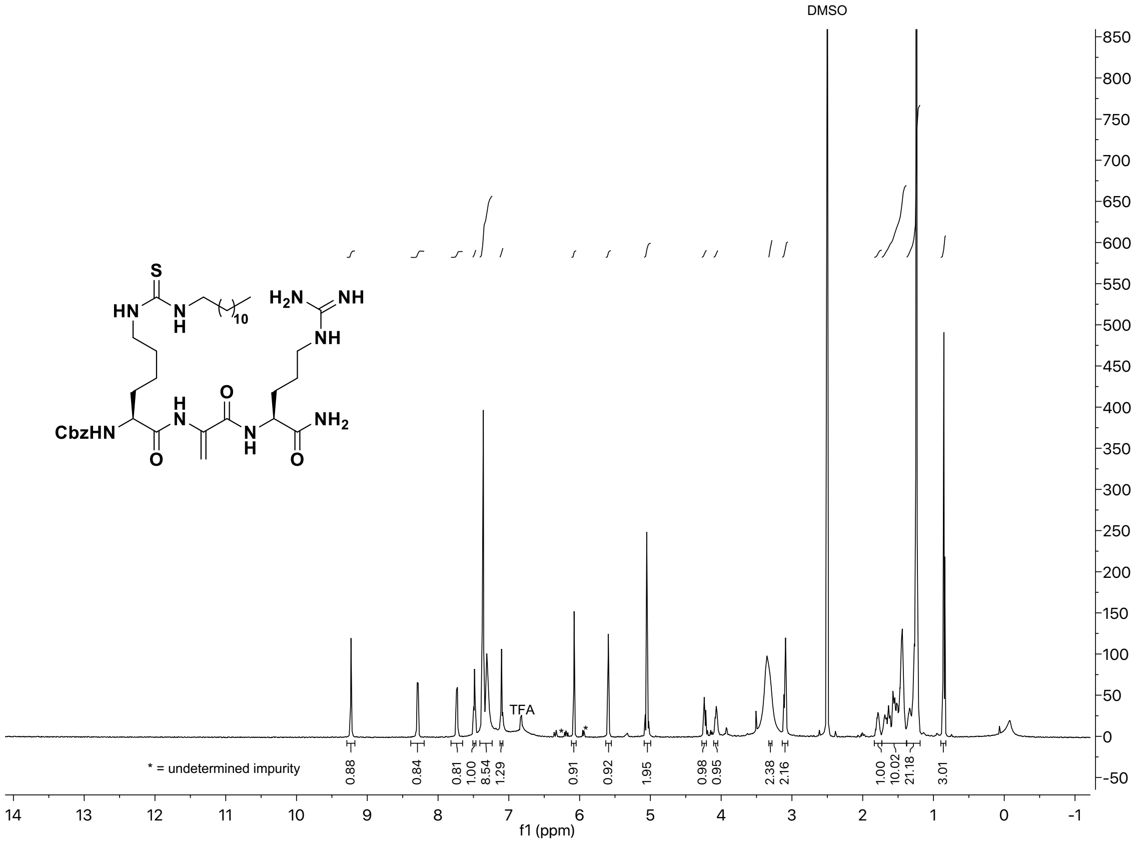

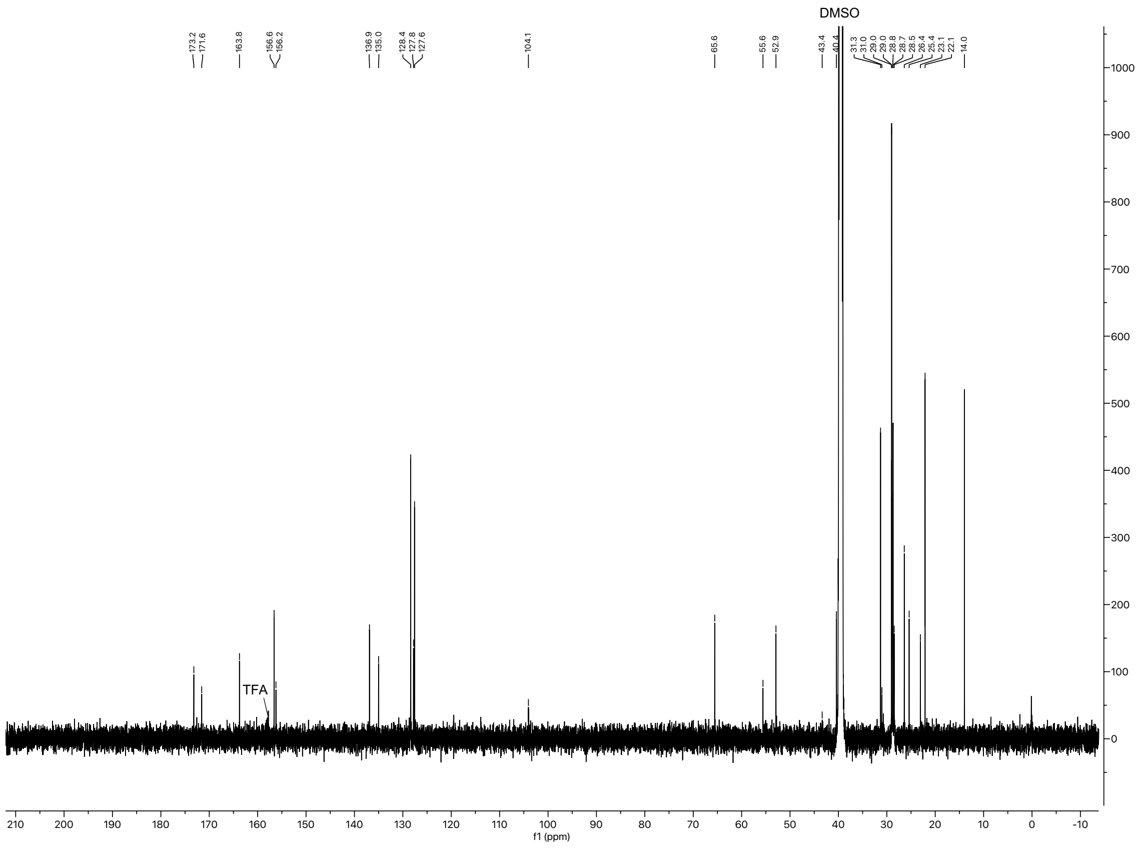

$^{1}H$ and $^{13}C$ spectra of compound **23**.

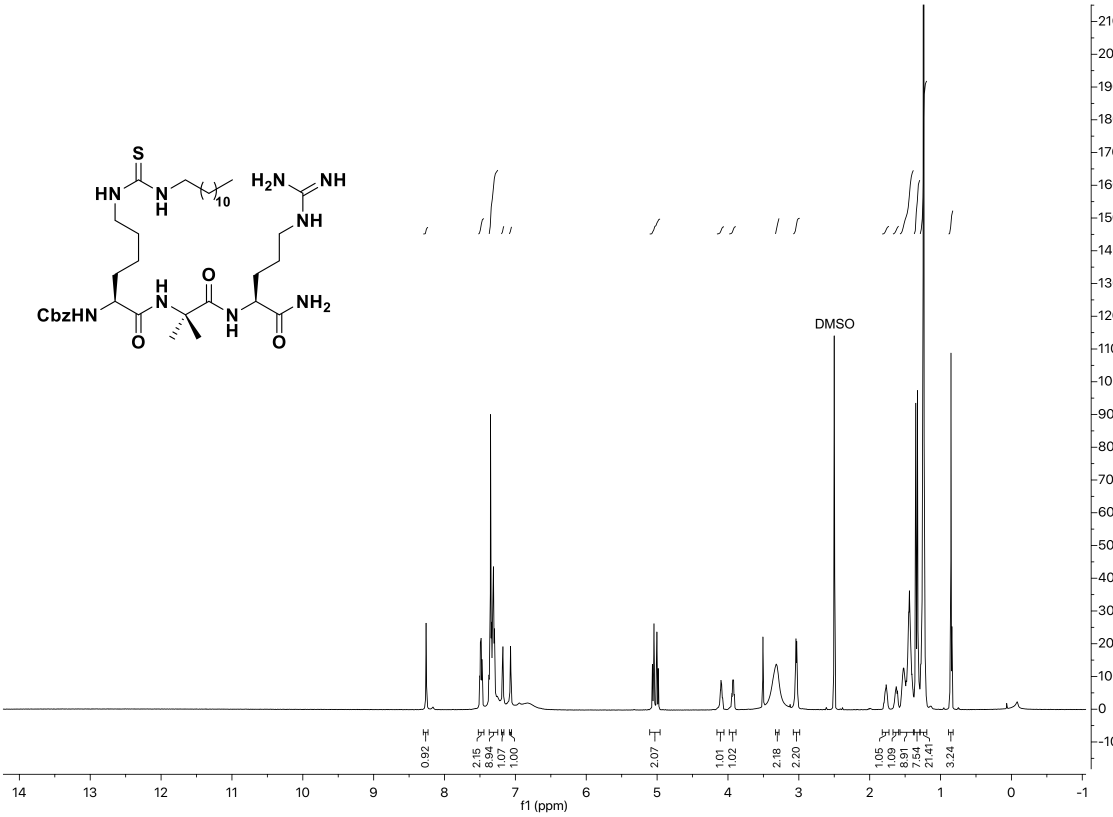

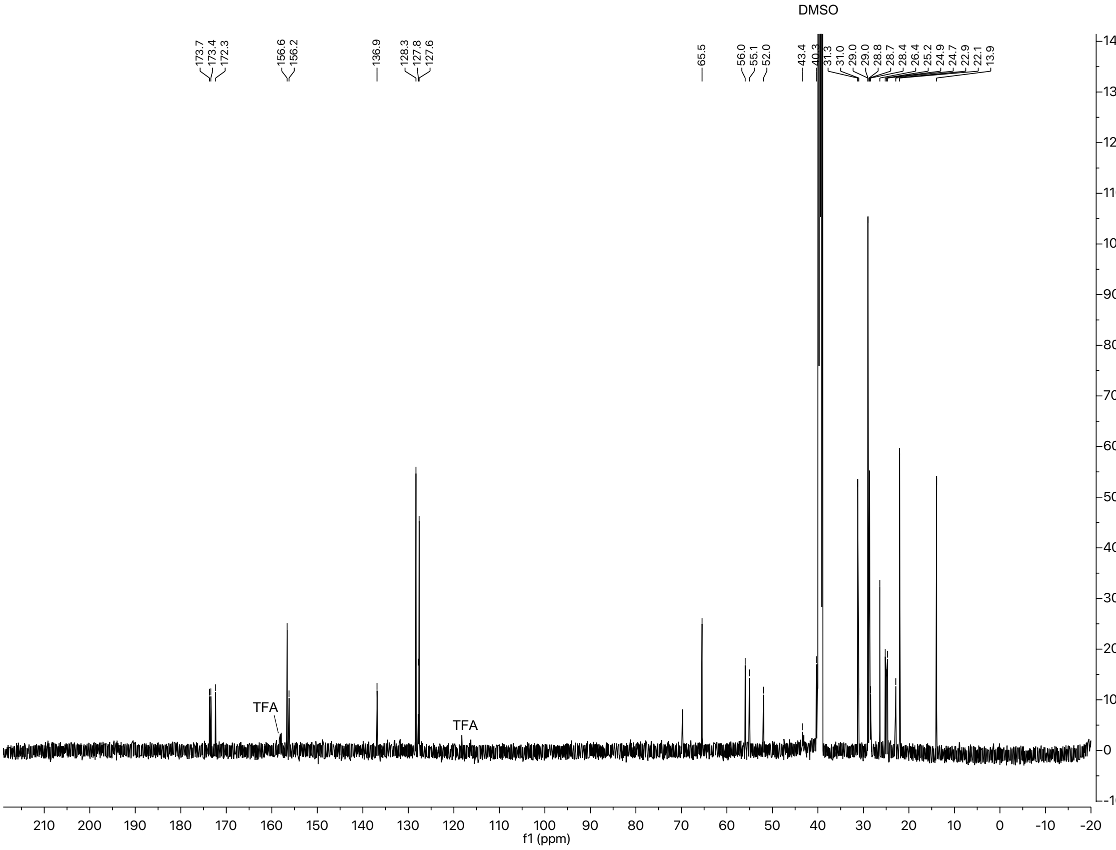

#### $^{1}H$ and $^{13}C$ spectra of compound **24**

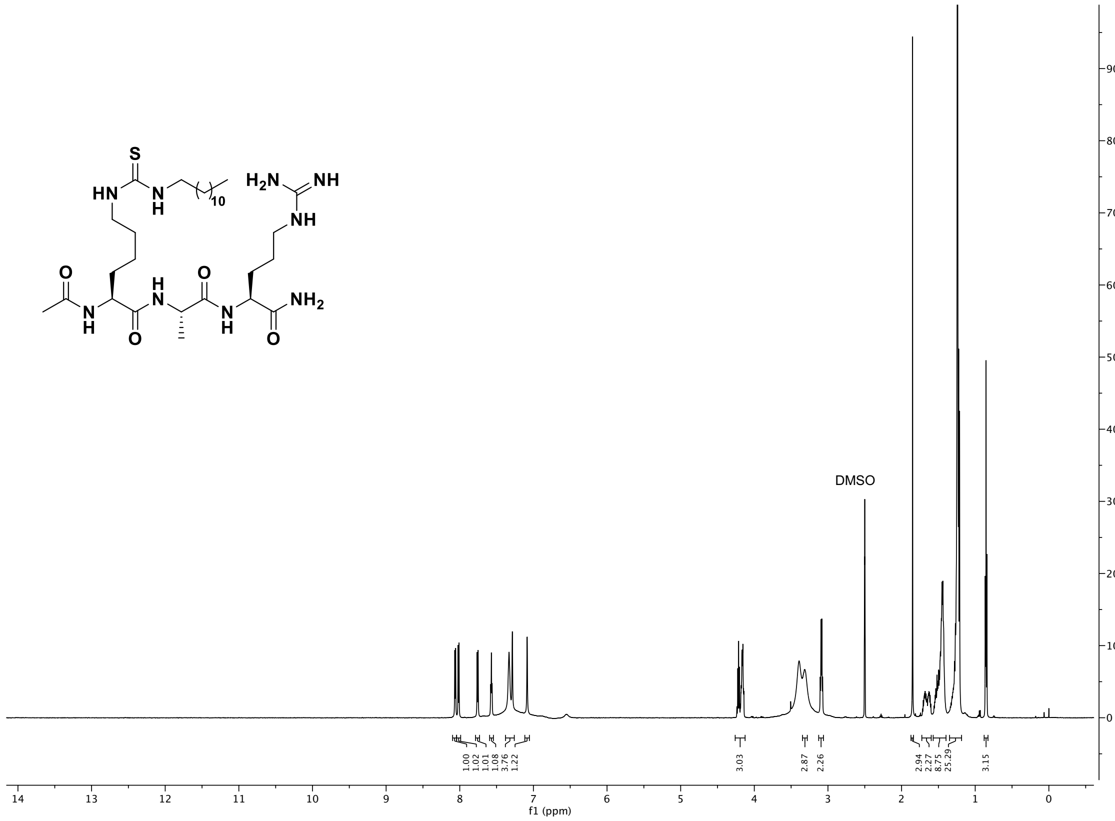

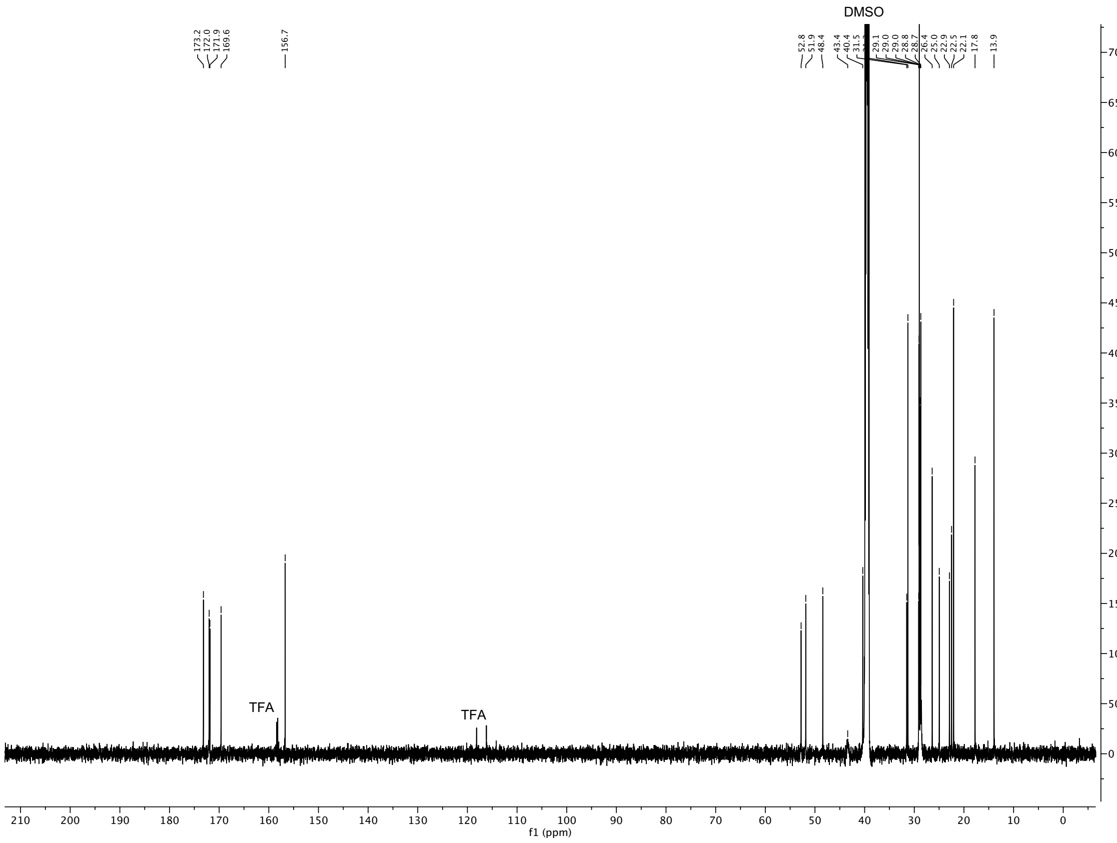

#### $^{1}H$ and $^{13}C$ spectra of compound **24-D**

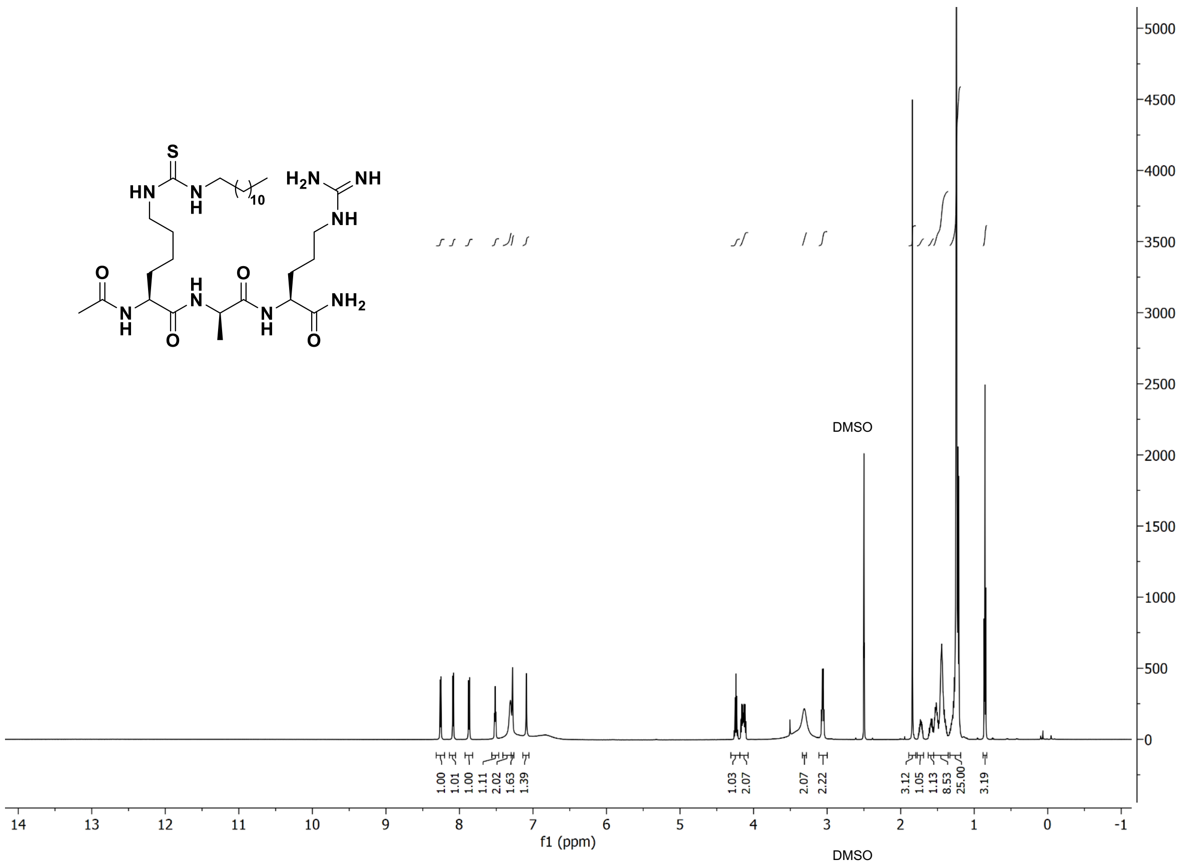

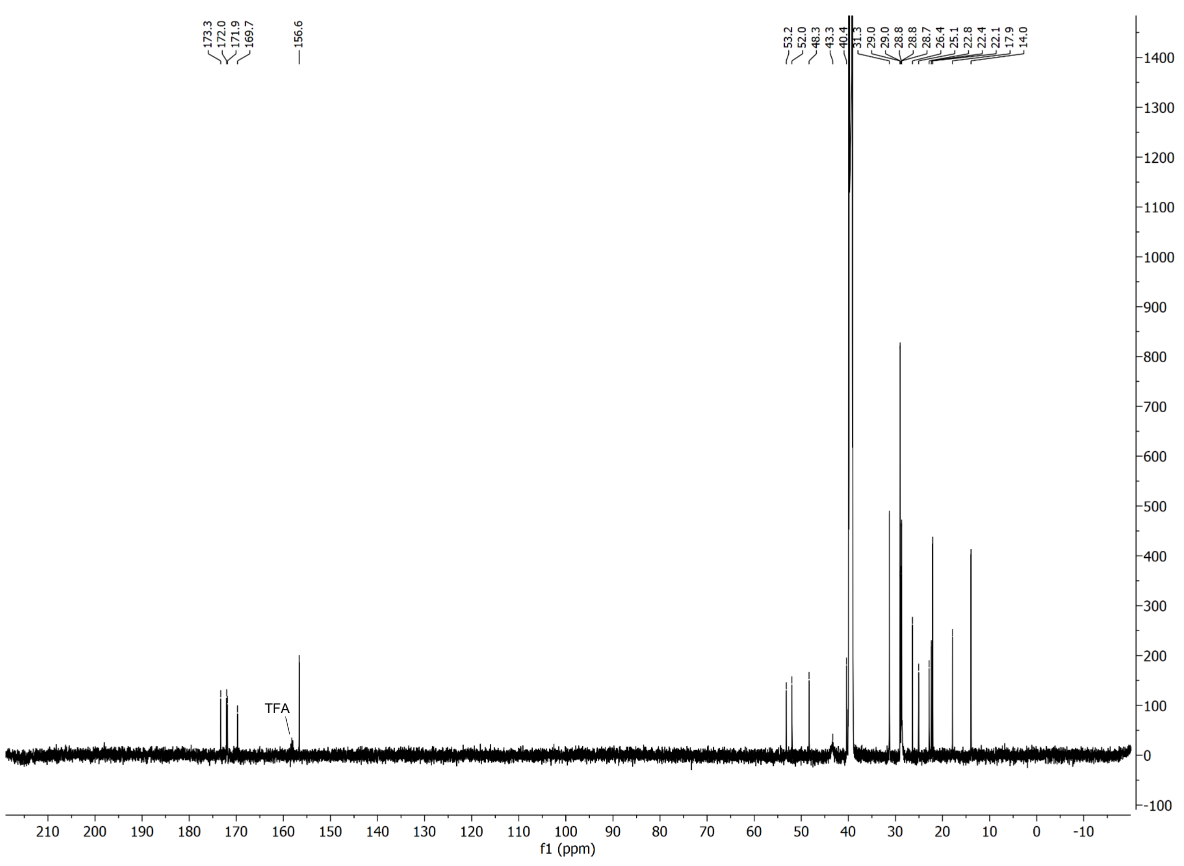

### $^{1}H$ and $^{13}C$ spectra of compound **25**

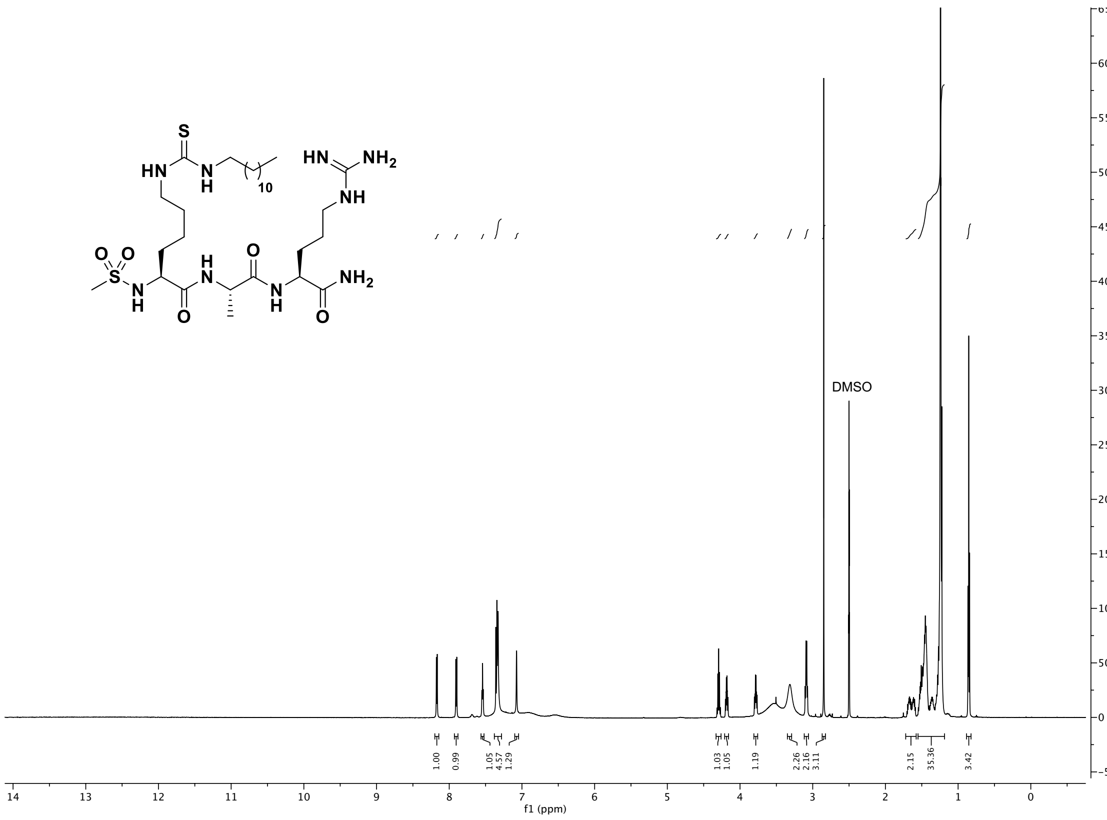

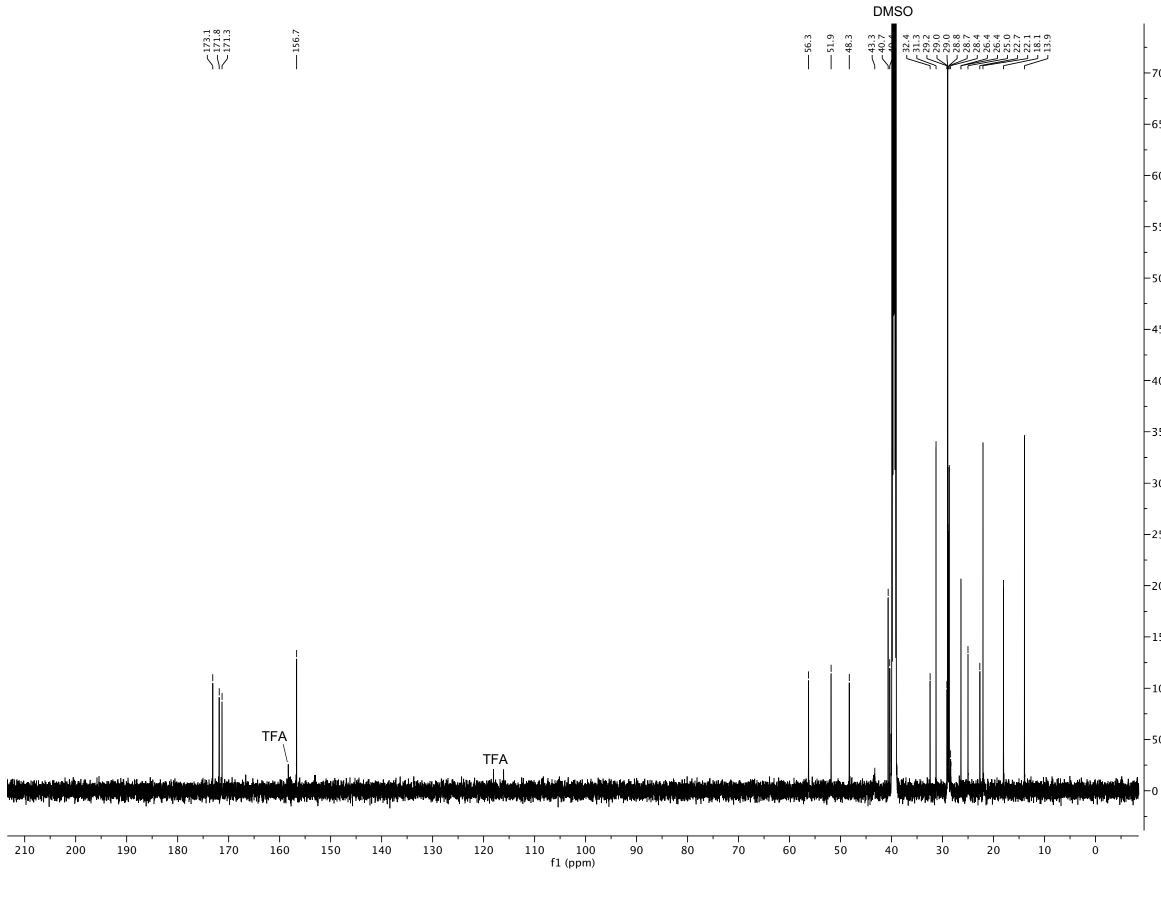

<sup></sup>

[1]H-[1]H COSY and [1]H-[13]C HSQC spectra of compound **25**

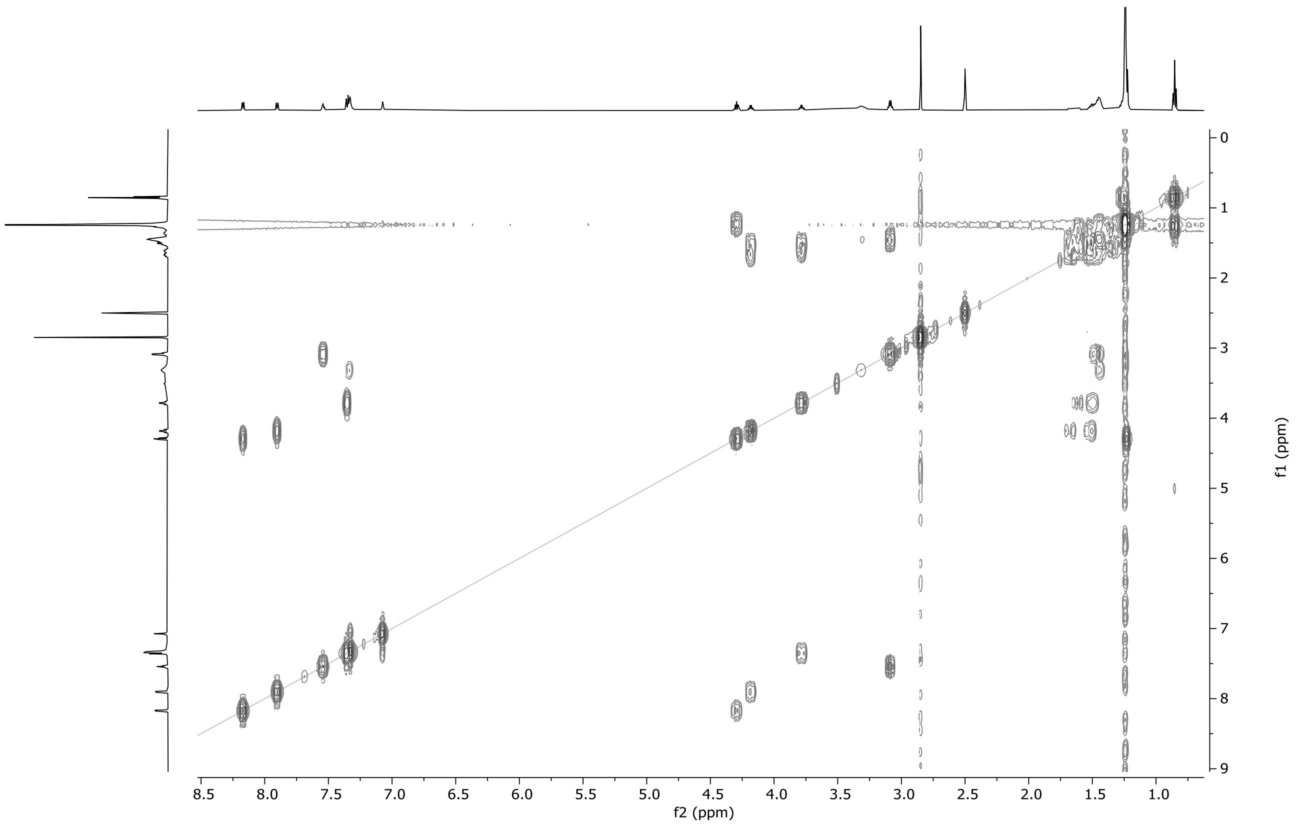

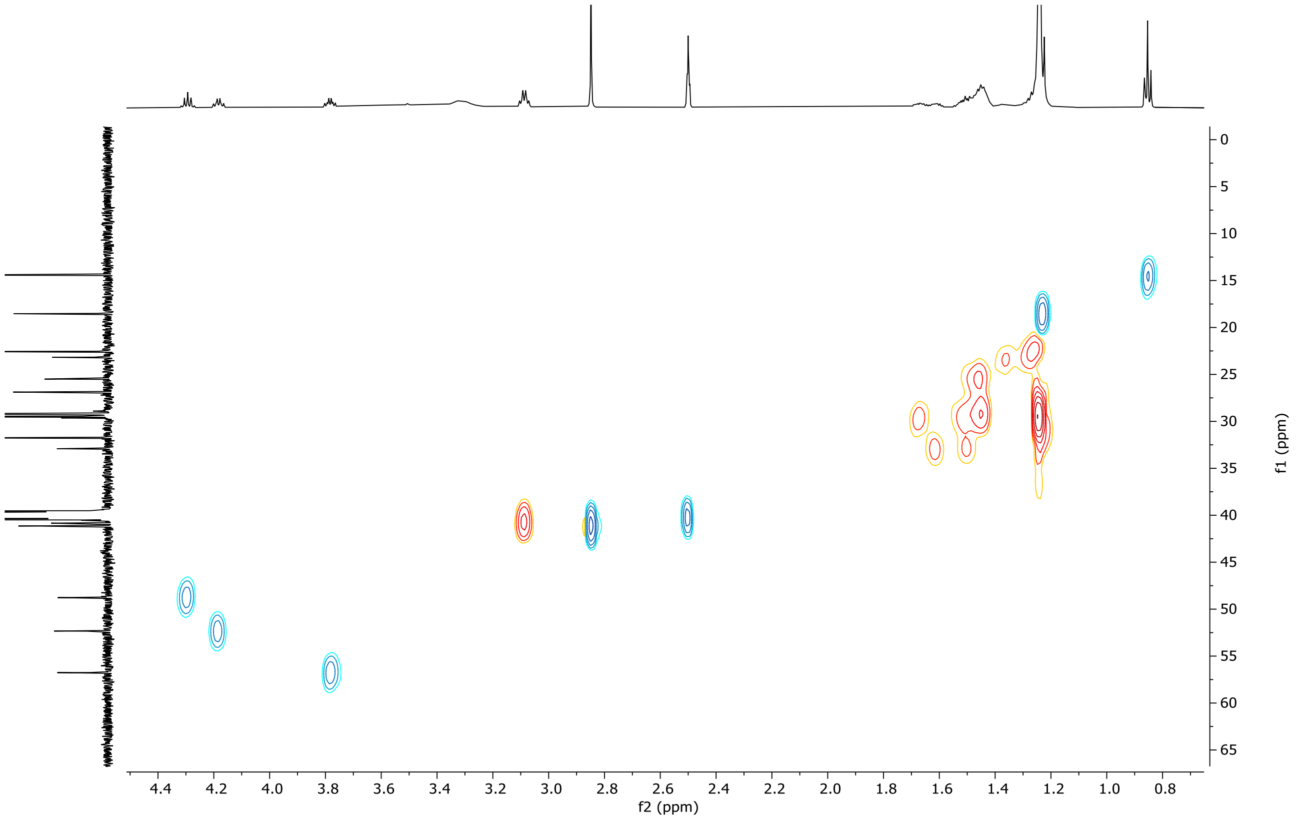

$^{1}$H-$^{13}$C HMBC spectrum of compound **25**

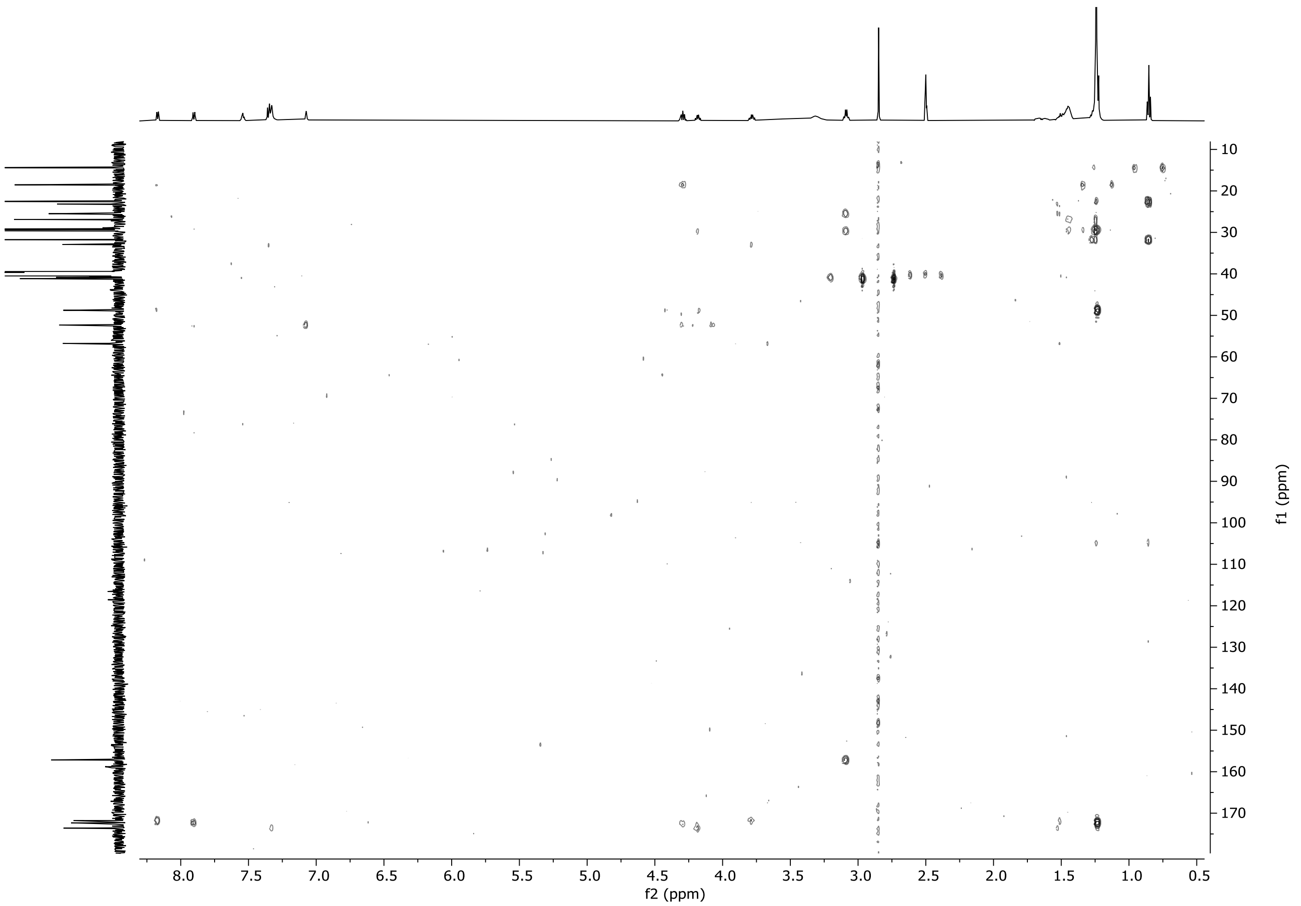

[1]H and [13]C spectra of compound **25-D**

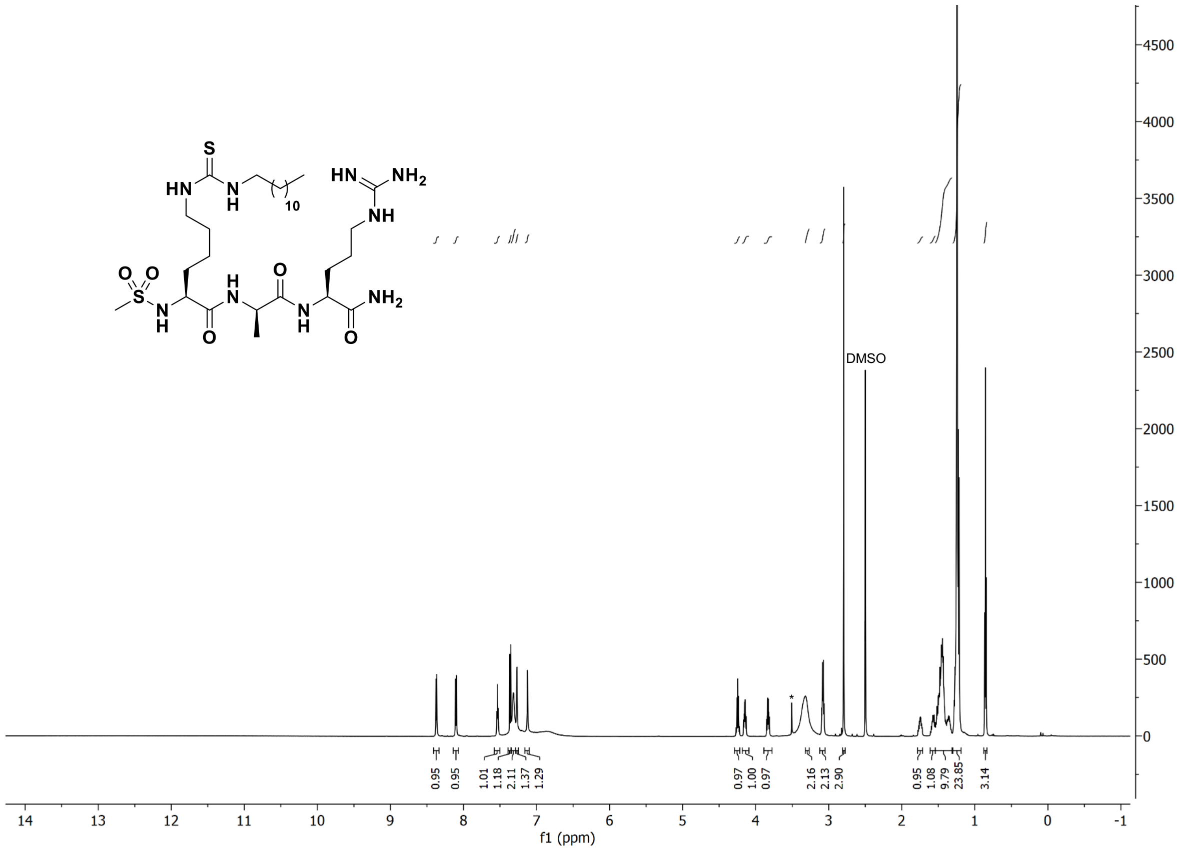

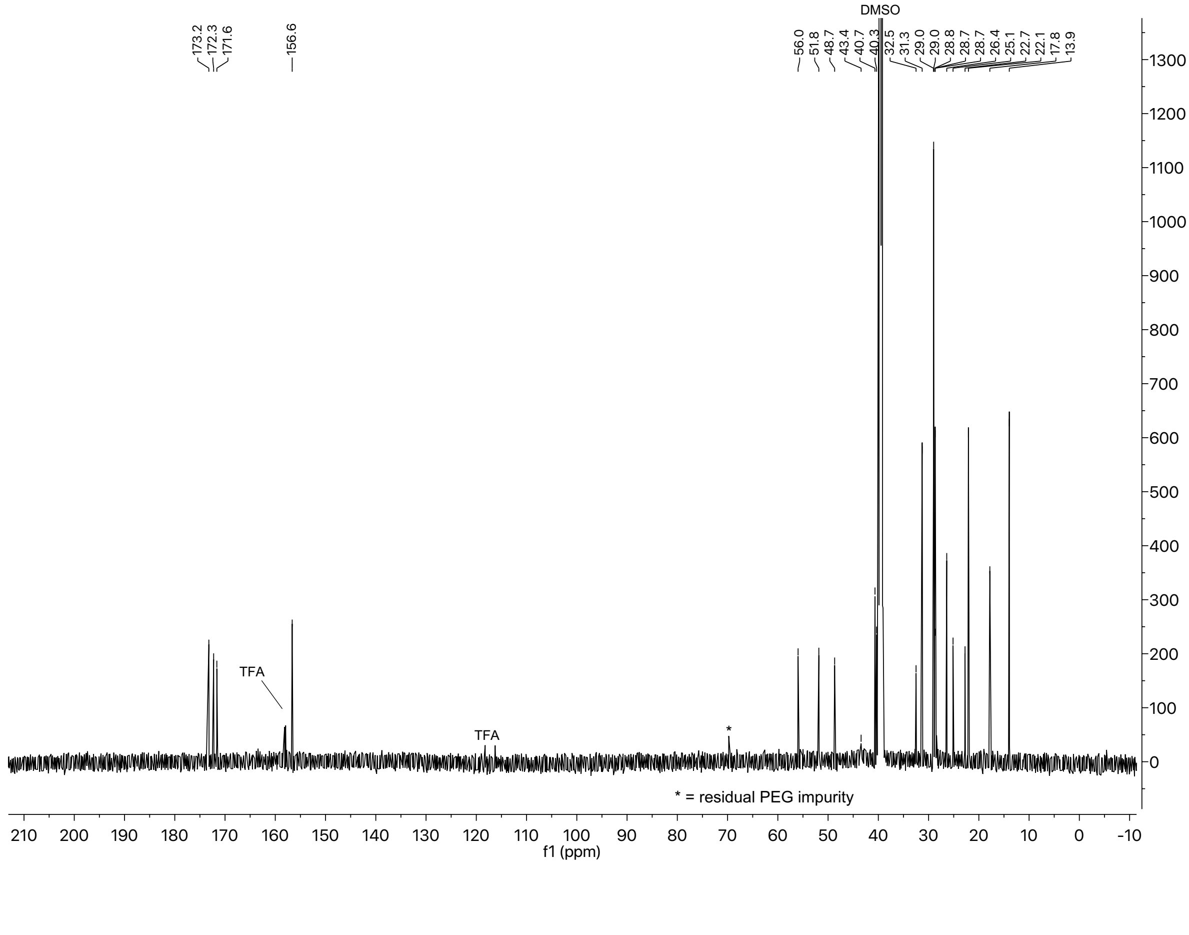

$^{1}H$-$^{1}H$ COSY and $^{1}H$-$^{13}C$ HSQC spectra of compound **25-D**

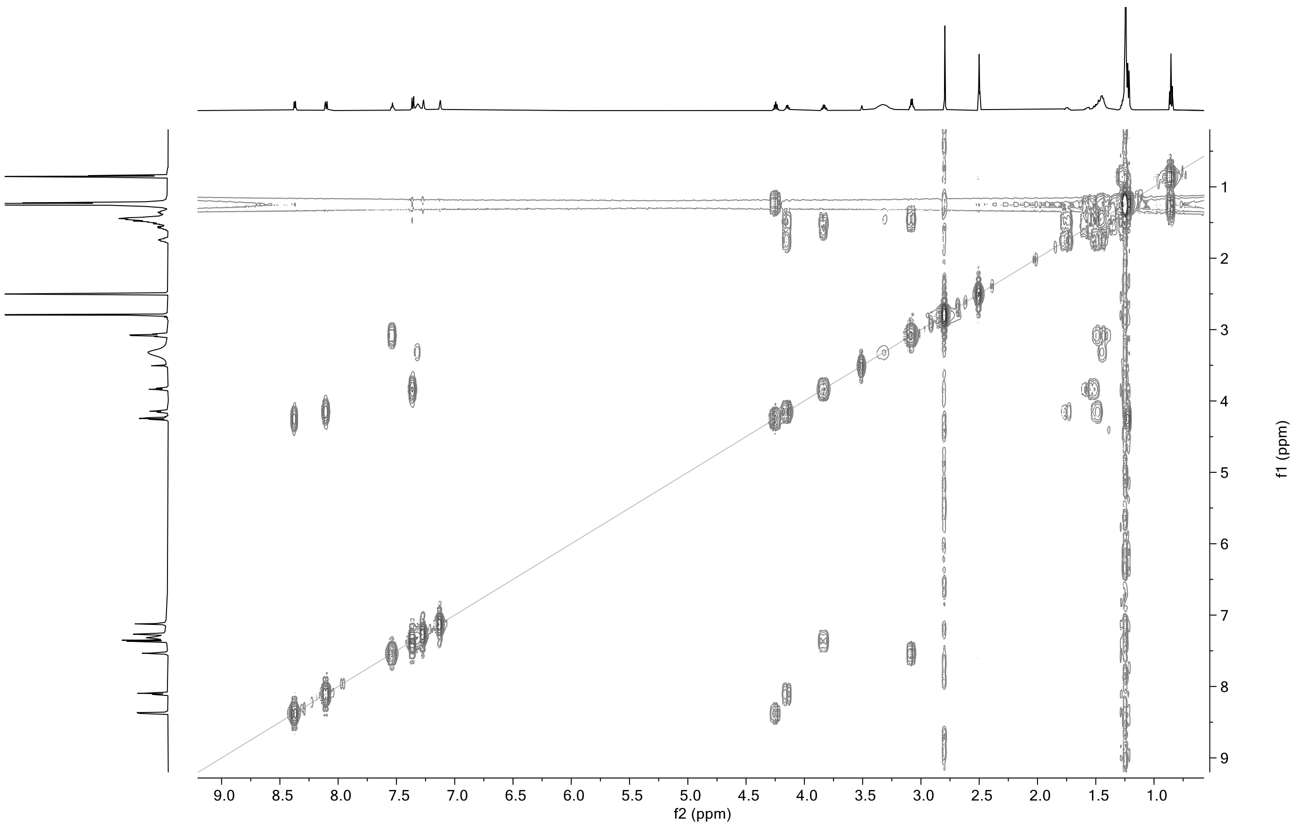

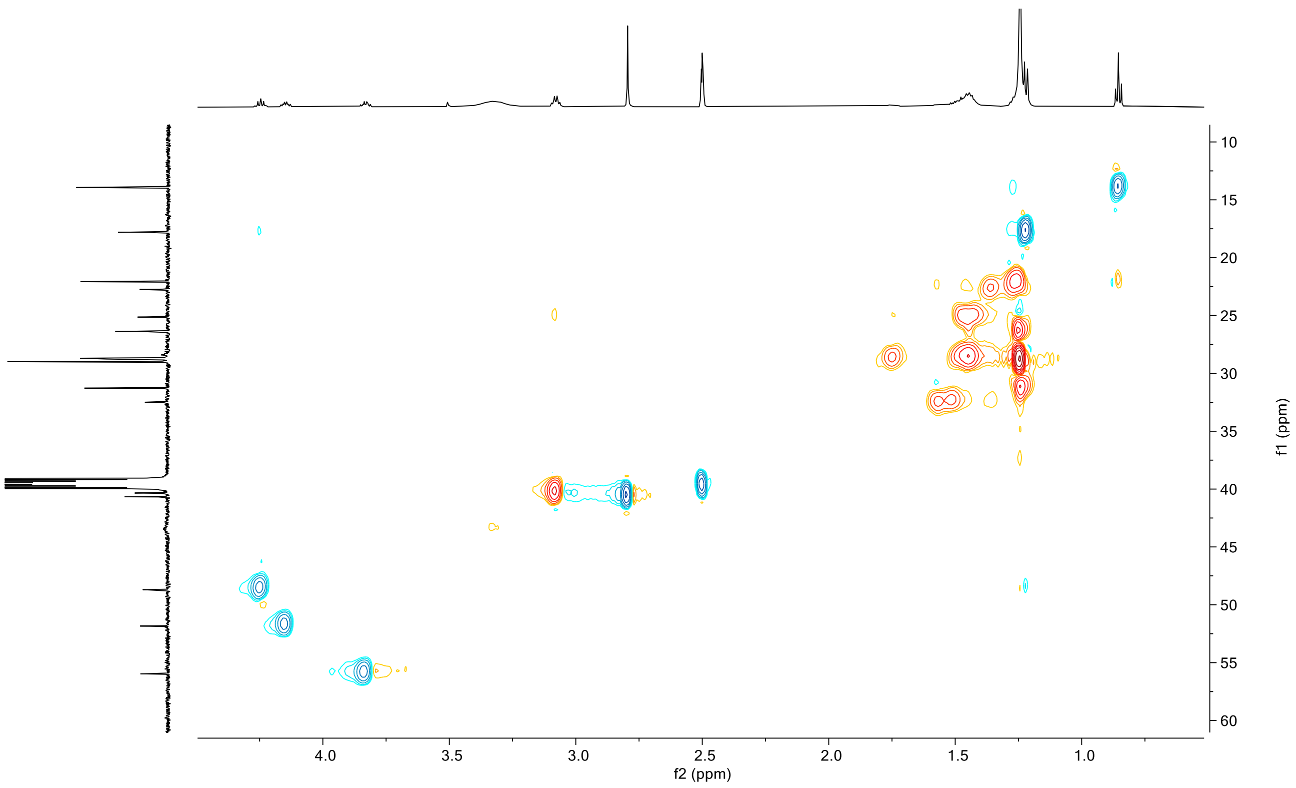

[1]H-[13]C HMBC spectrum of compound **25-D**

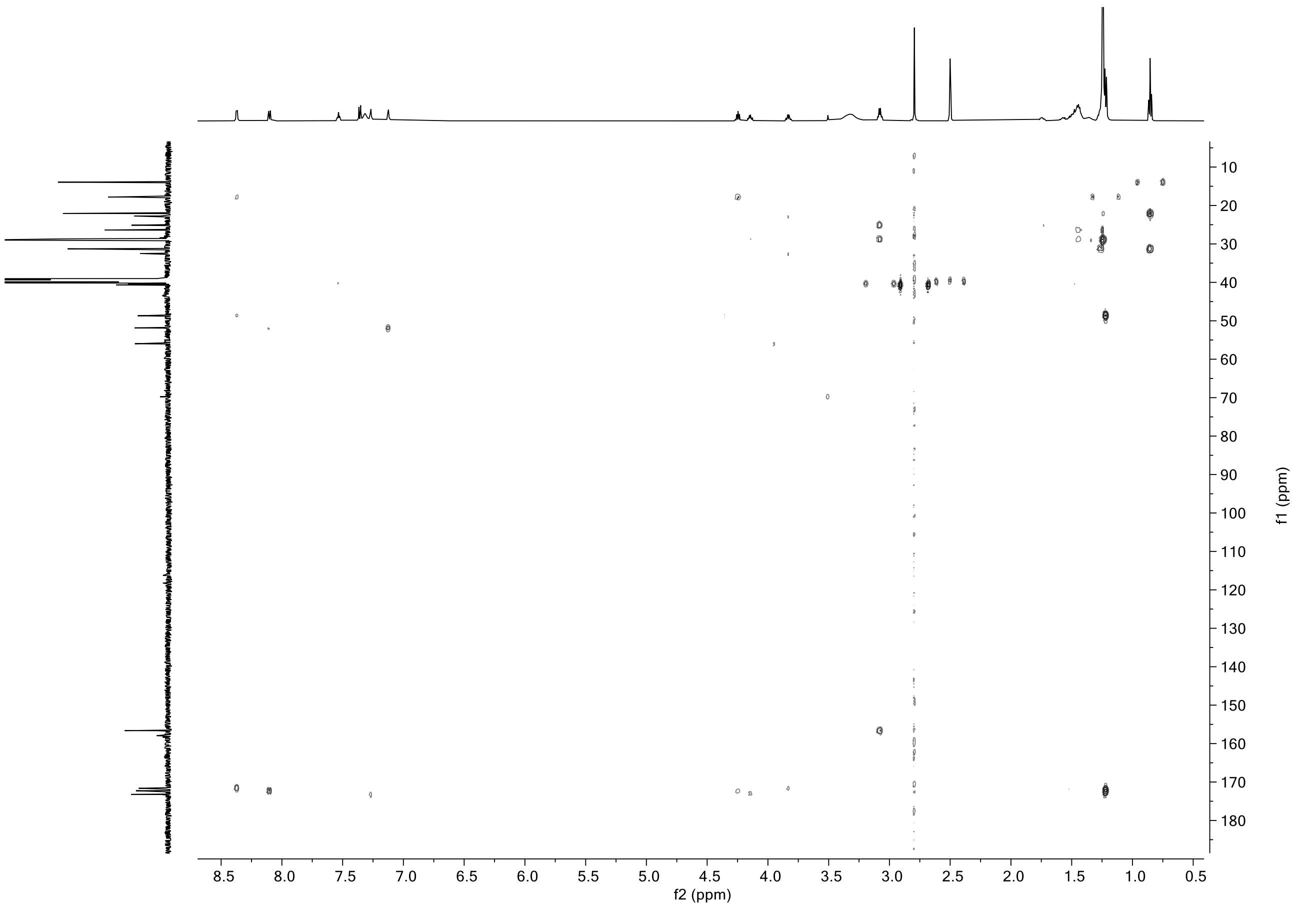

### $^{1}H$ and $^{13}C$ spectra of compound **26**

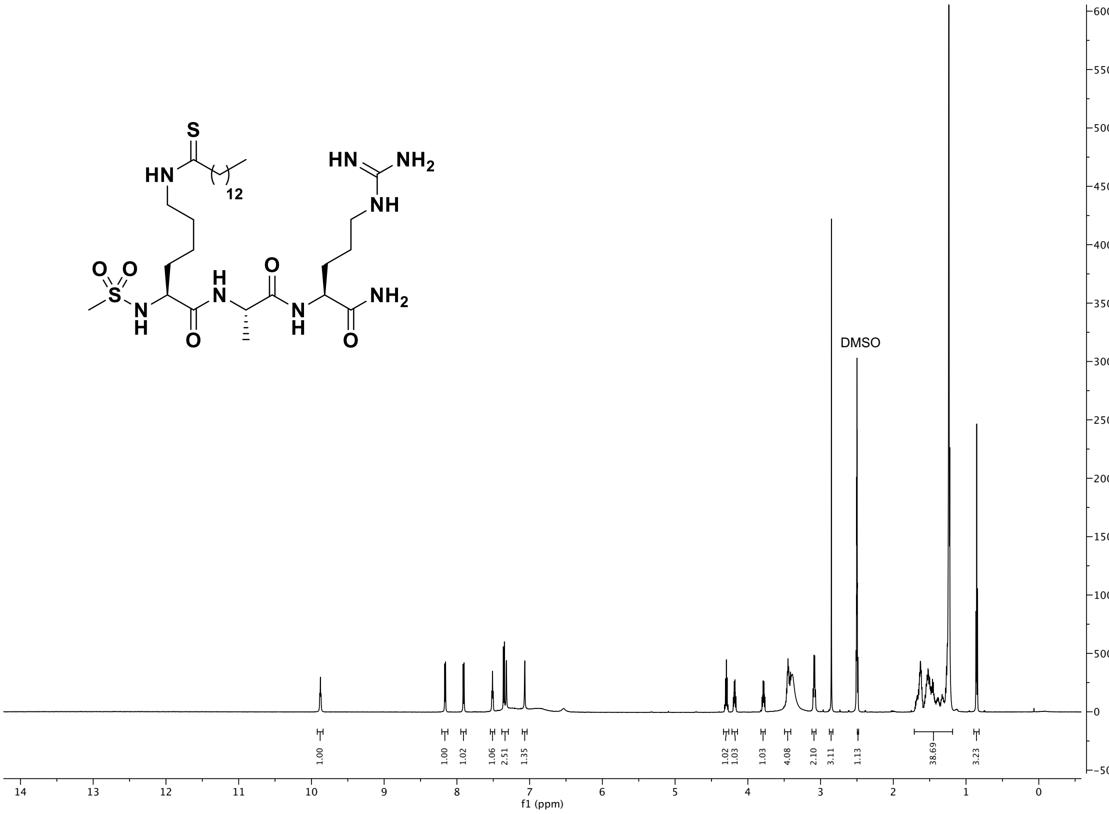

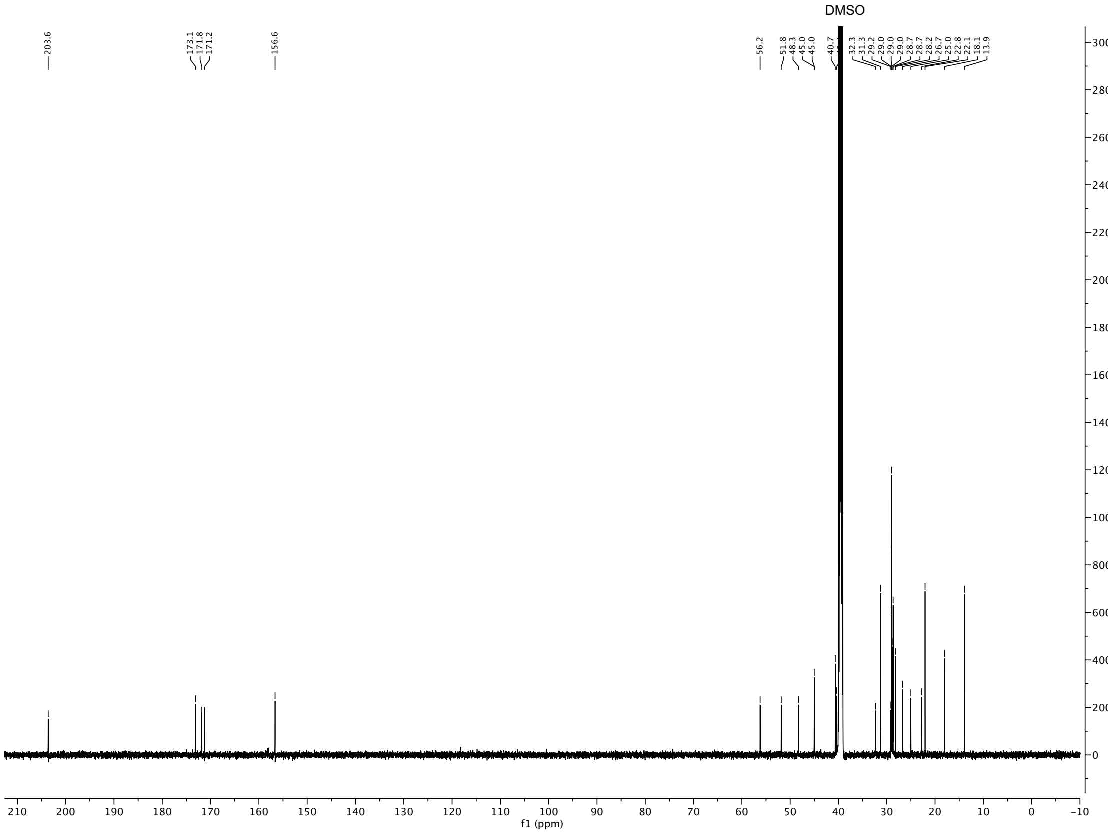

### $^{1}H$-$^{1}H$ COSY and $^{1}H$-$^{13}C$ HSQC spectra of compound **26**

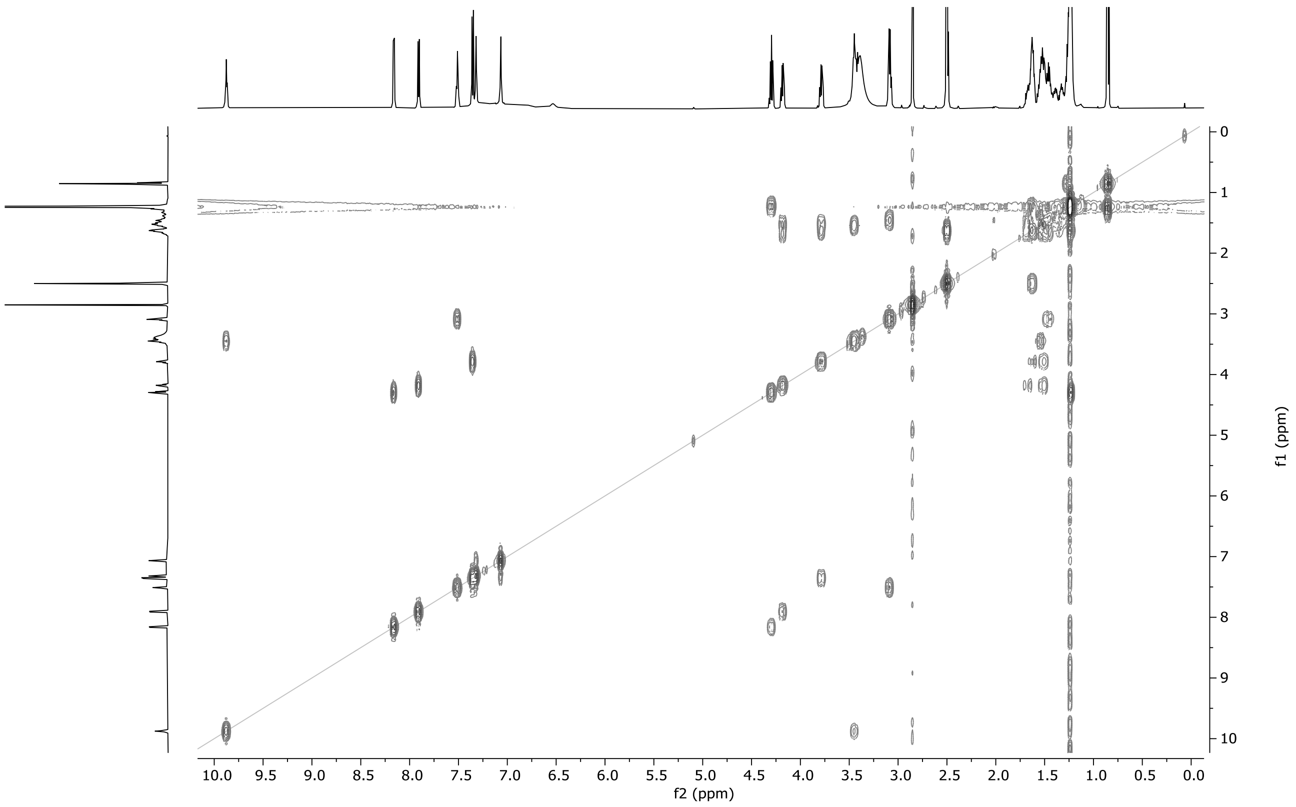

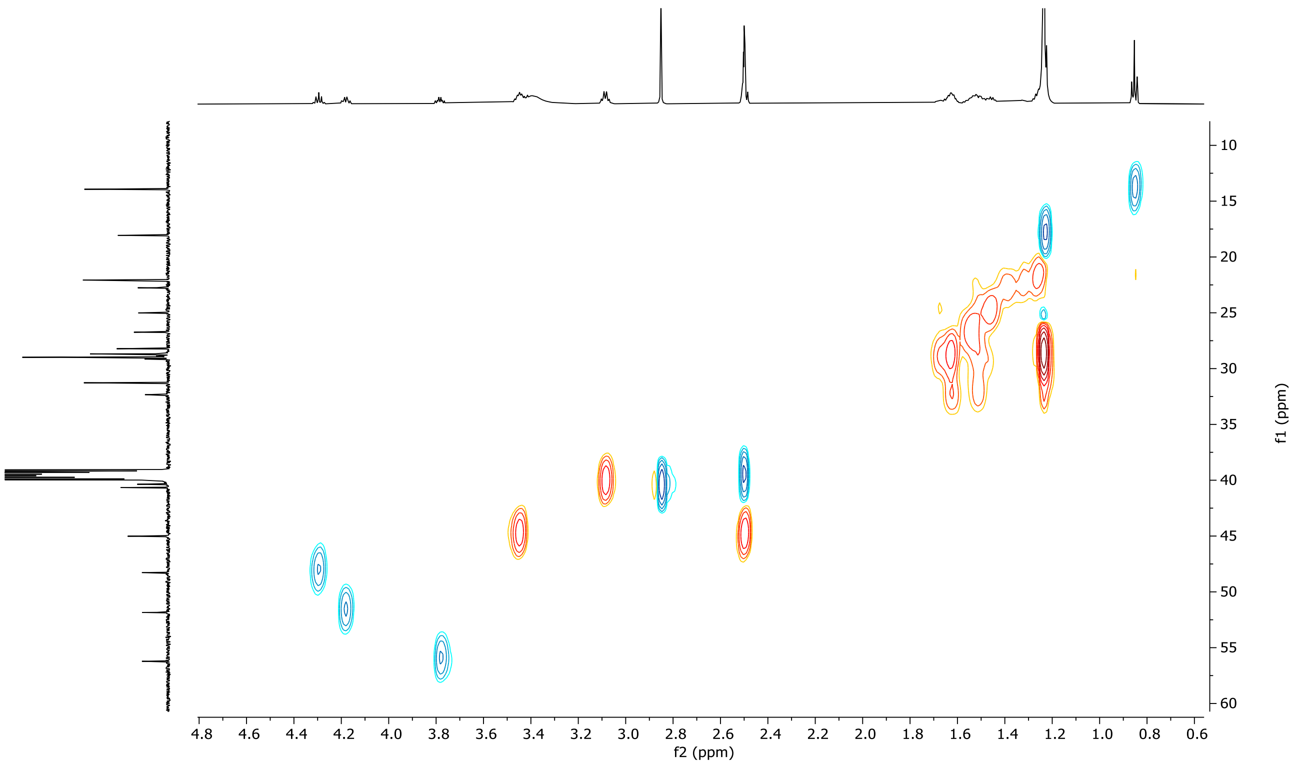

$^1$H-$^{13}$C HMBC spectrum of compound **26**

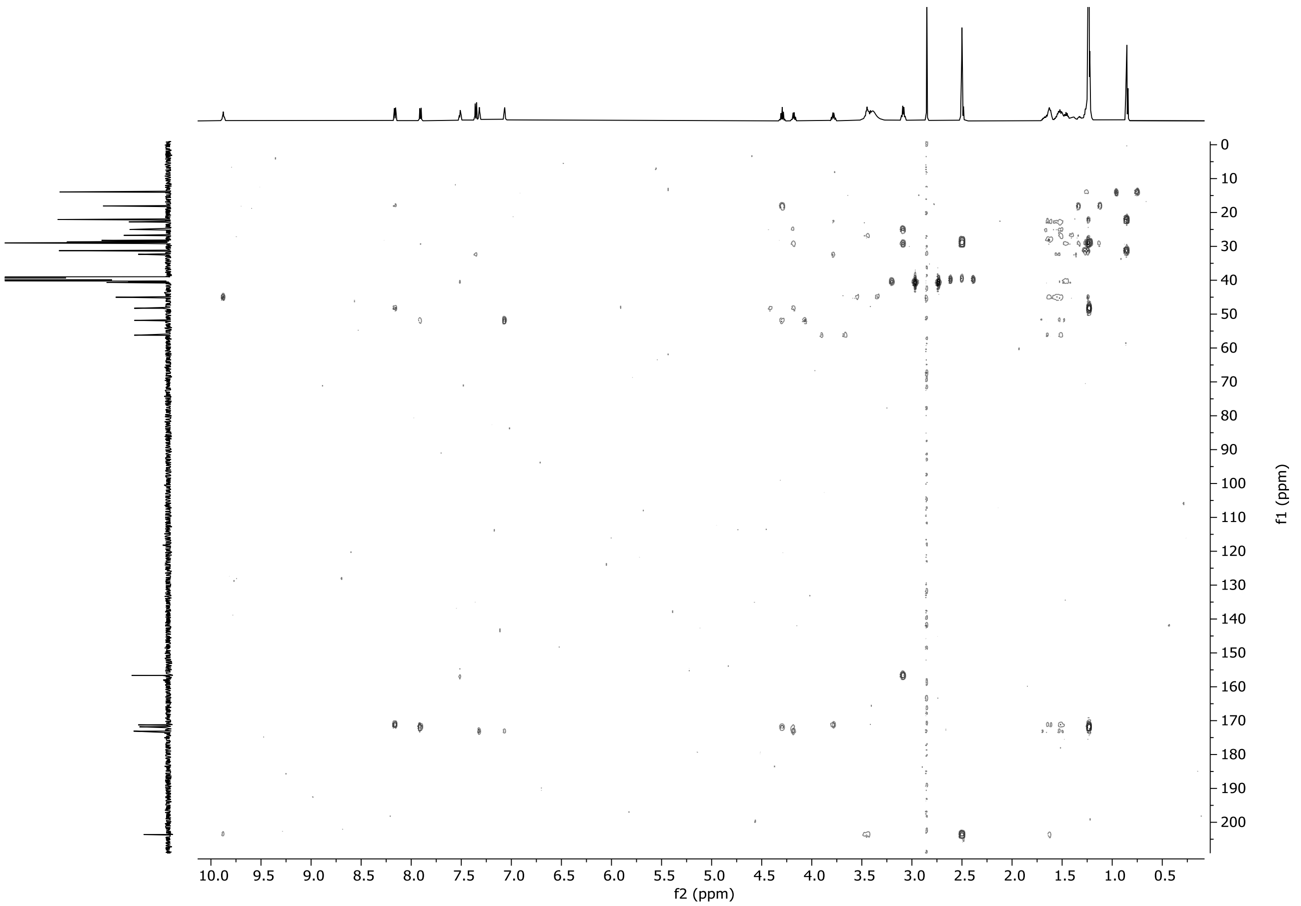

#### [1]H and [13]C spectra of compound **26-D**

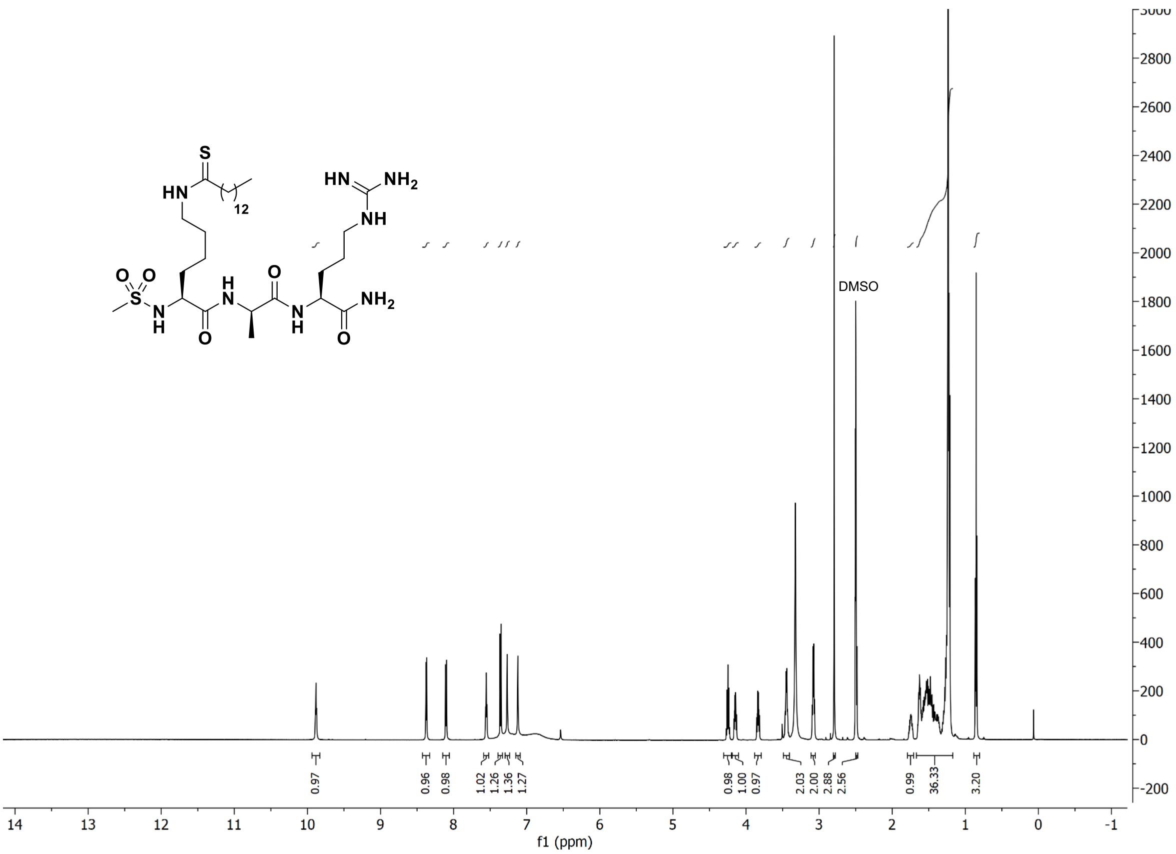

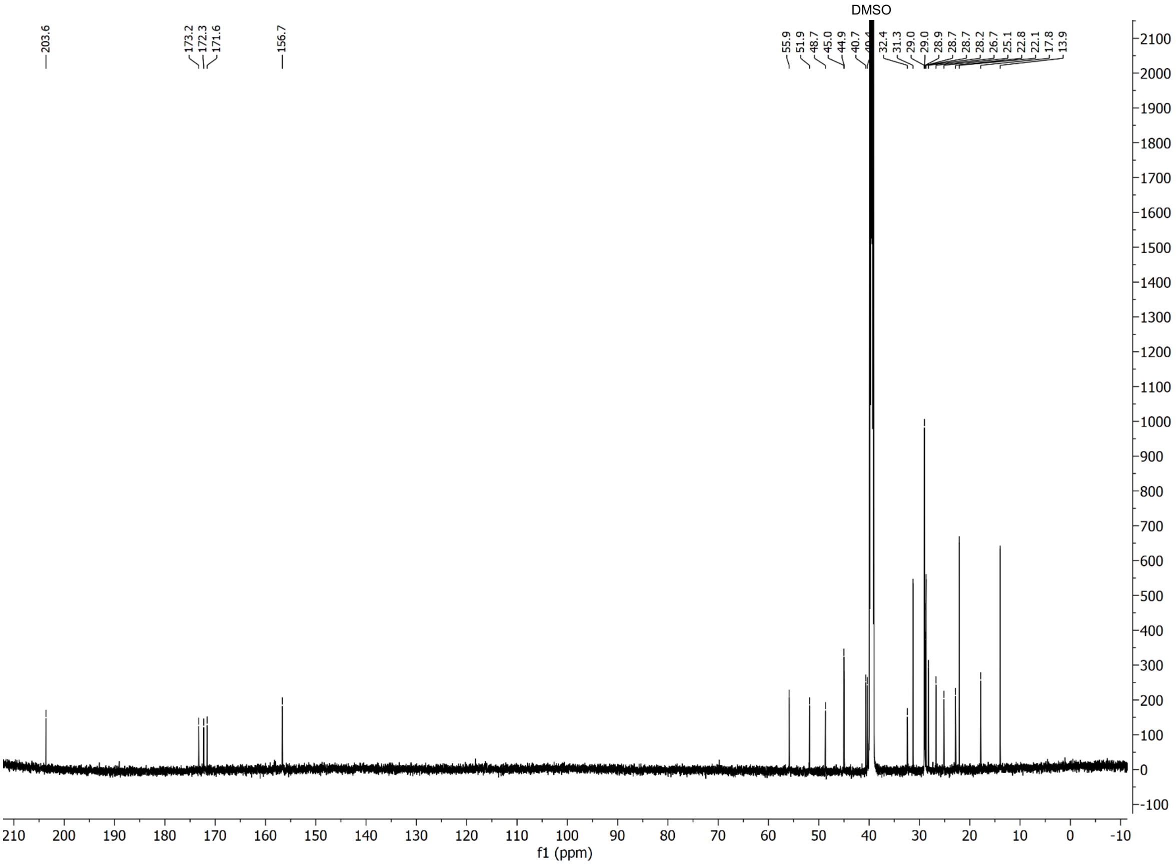

$^{1}H$-$^{1}H$ COSY and $^{1}H$-$^{13}C$ HSQC spectra of compound **26-D**

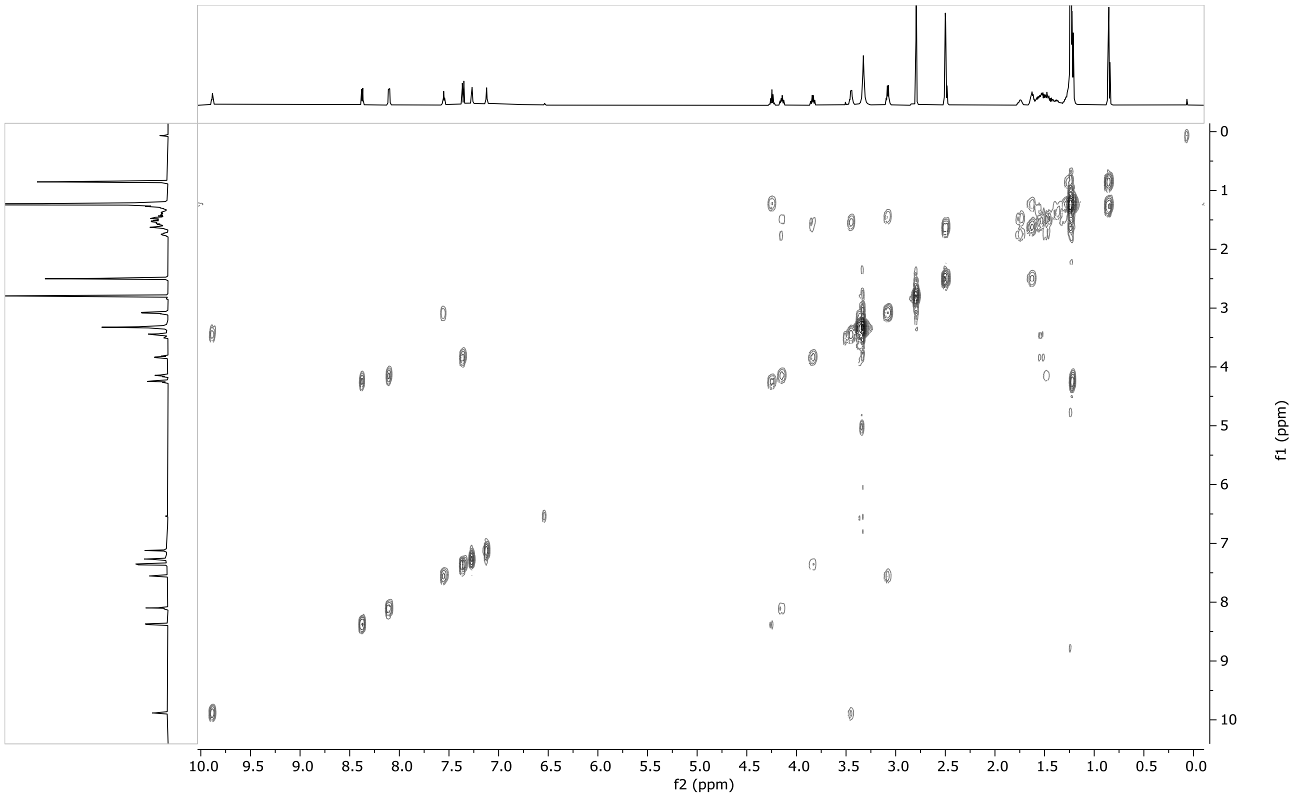

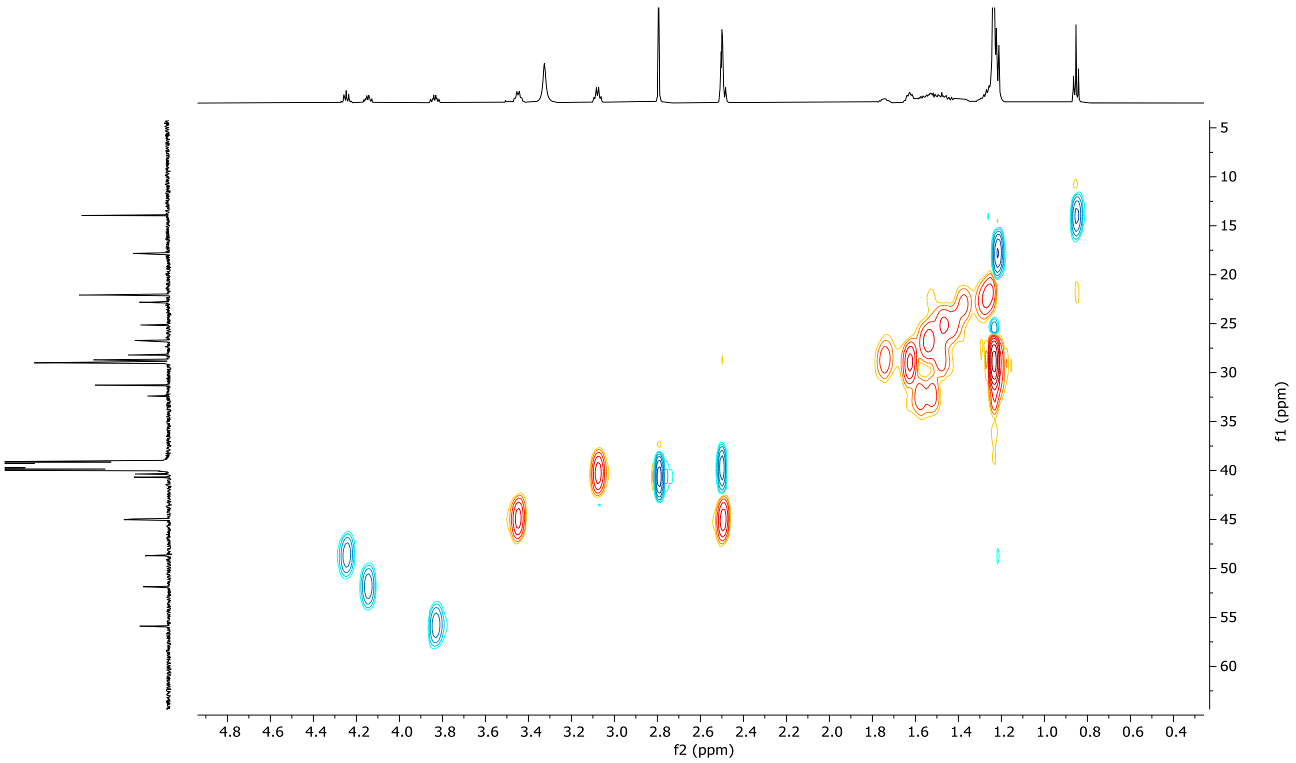

$^{1}$H-$^{13}$C HMBC spectrum of compound **26-D**

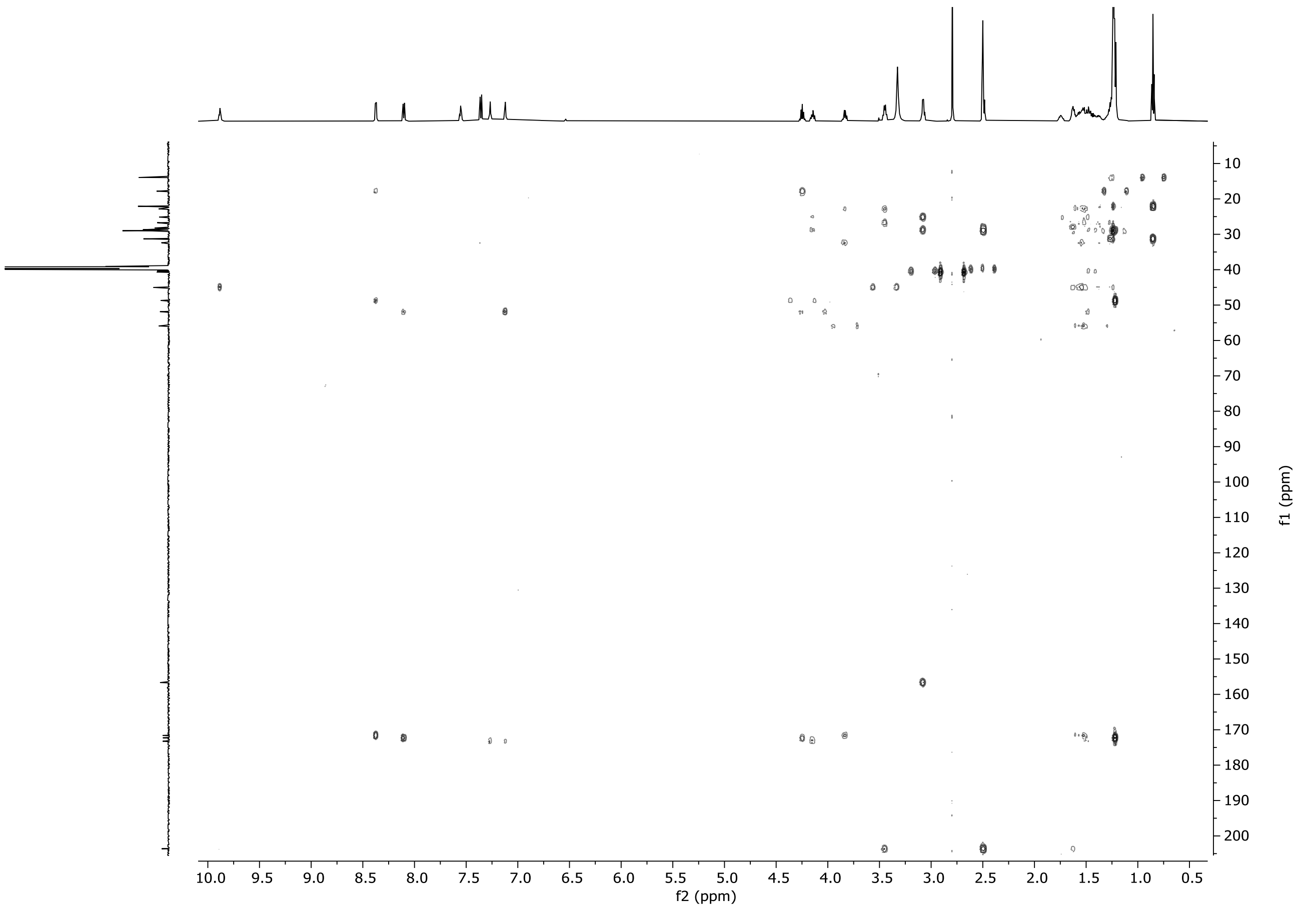

### [1]H and [13]C spectra of compound **S1**

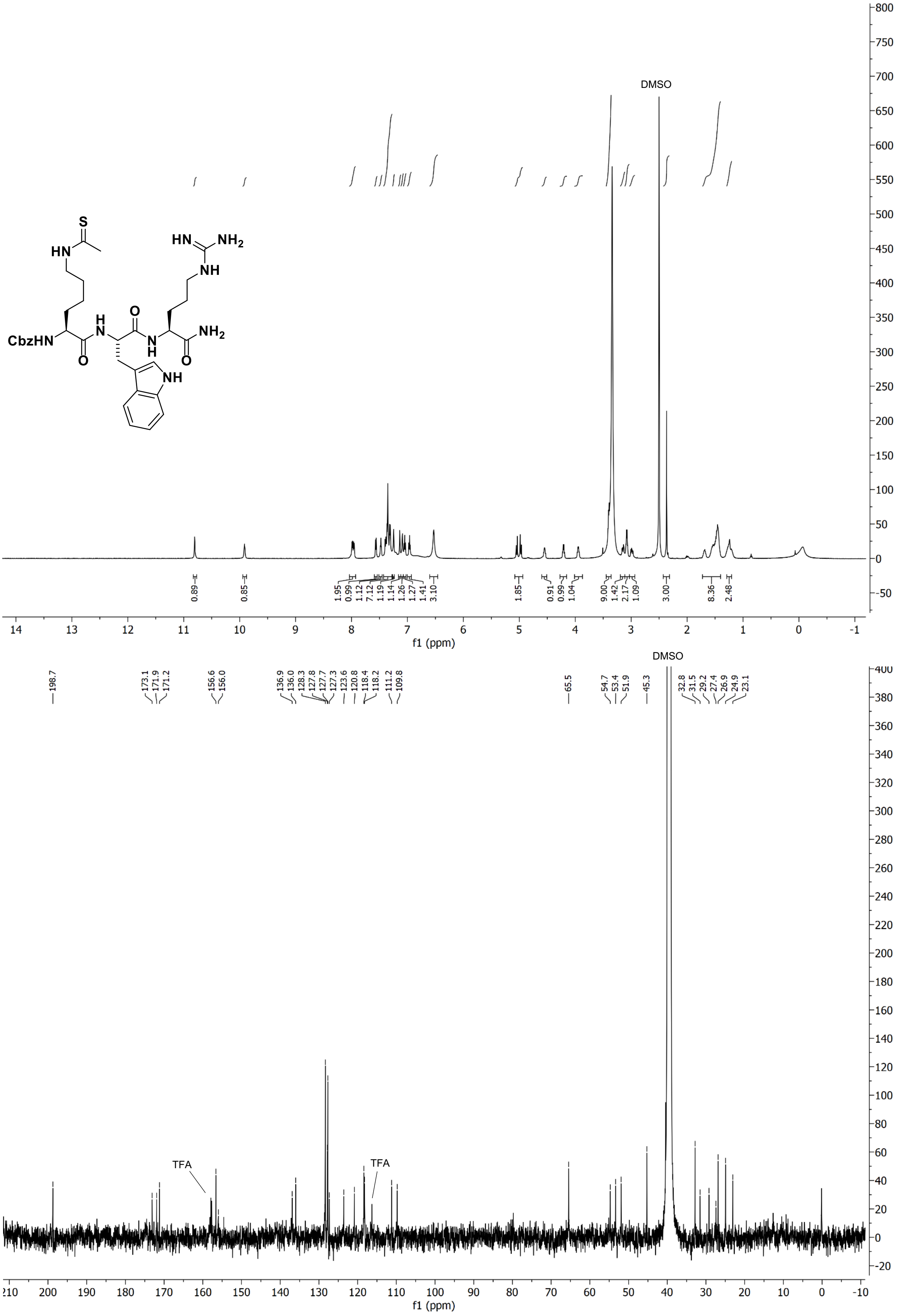

$^{1}H$ and $^{13}C$ spectra of compound **S2**

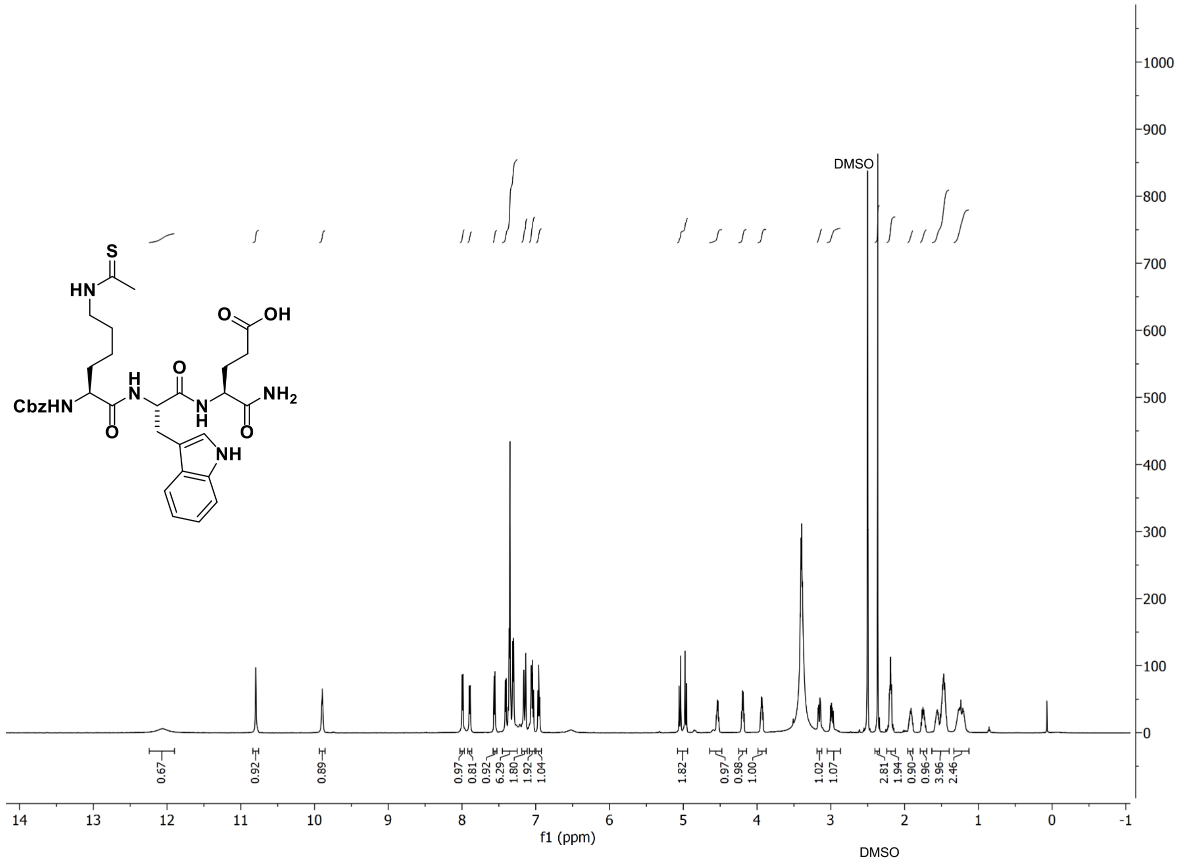

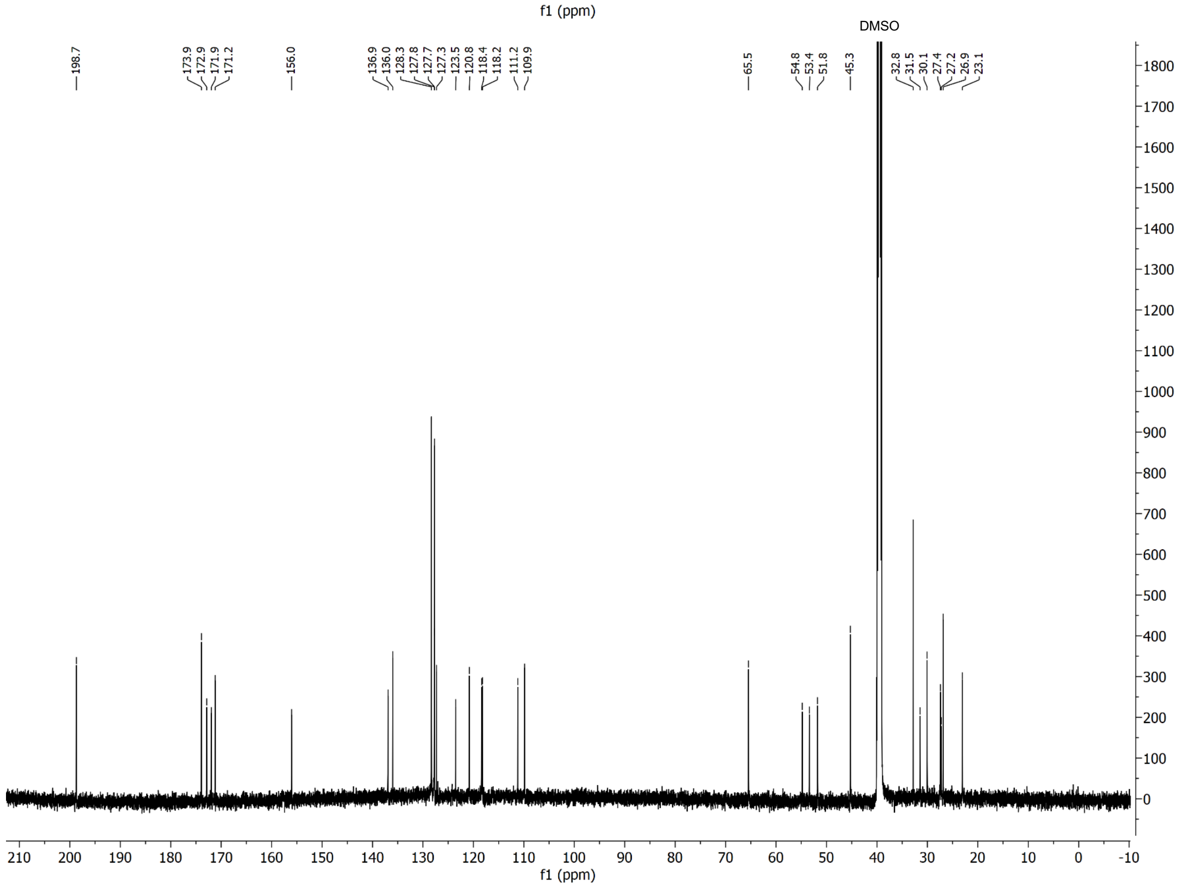

### $^{1}H$ and $^{13}C$ spectra of compound **S3**

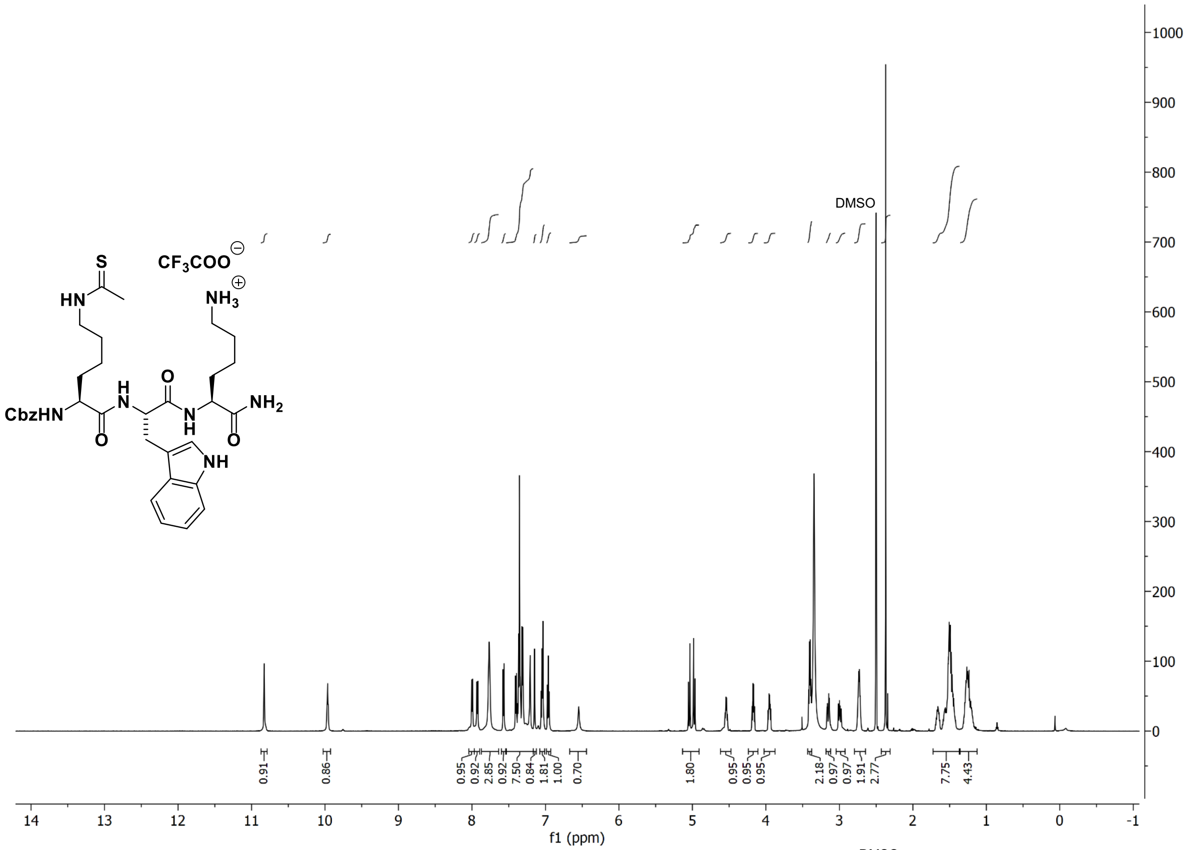

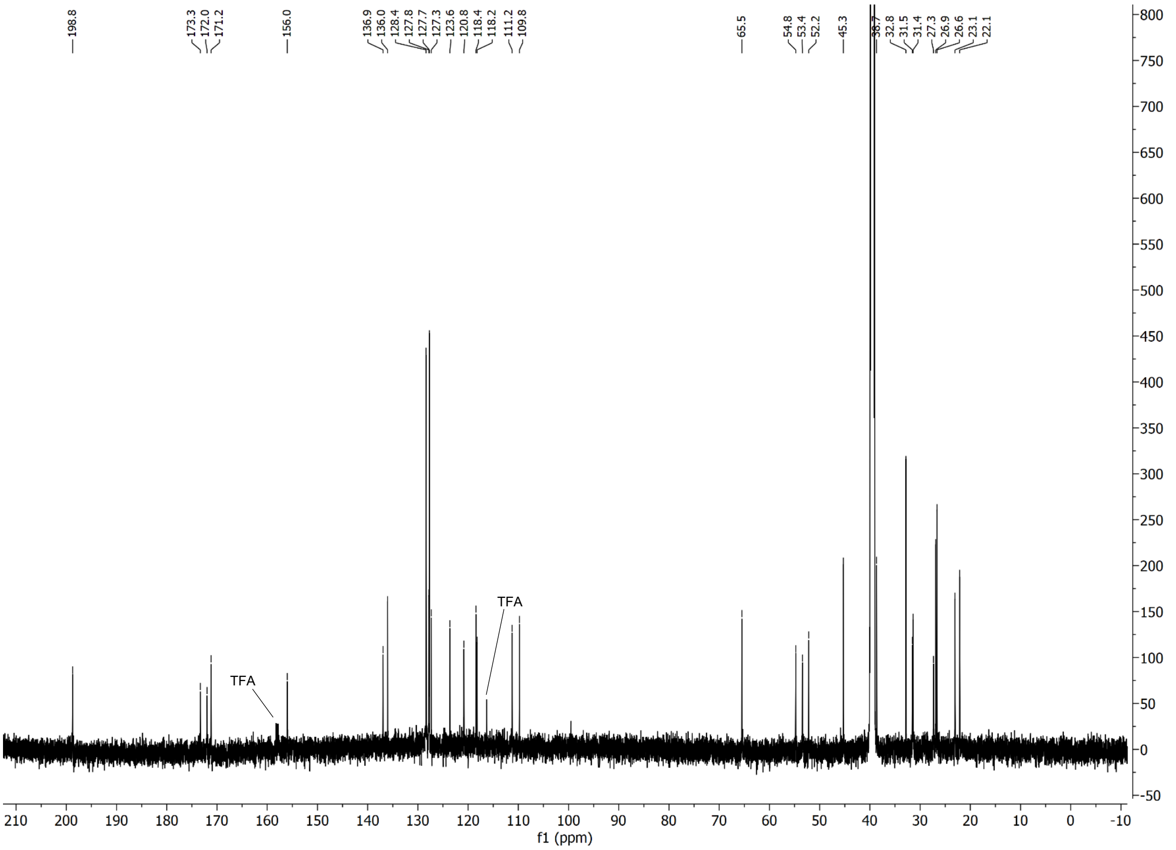

$^{1}H$ and $^{13}C$ spectra of compound **S4**

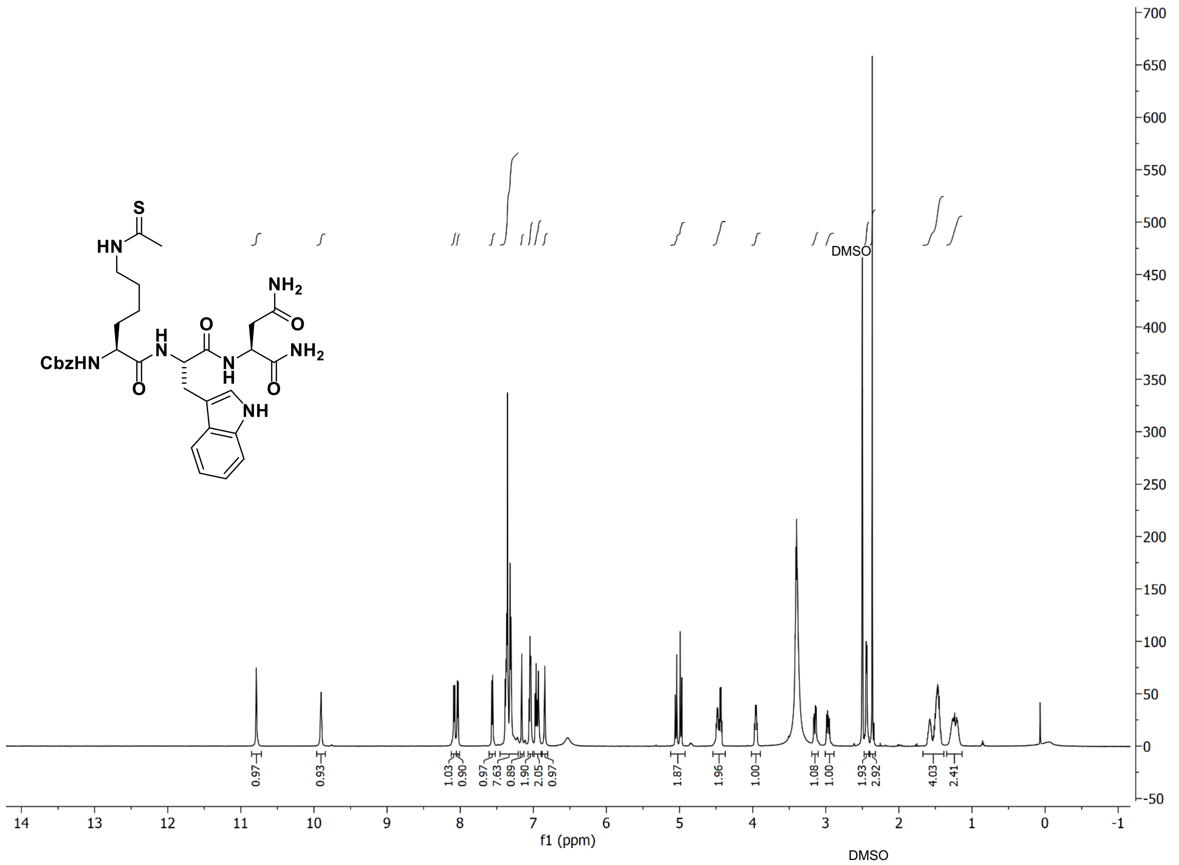

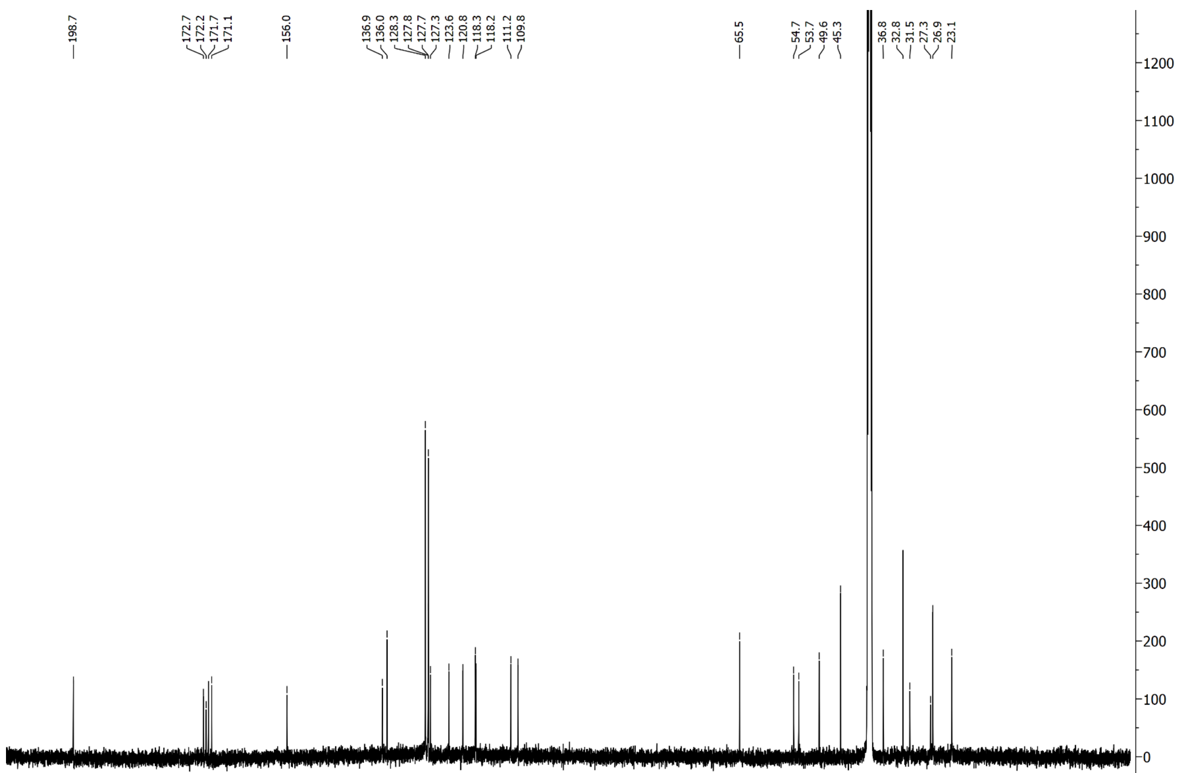

#### $^{1}H$ and $^{13}C$ spectra of compound **S5**

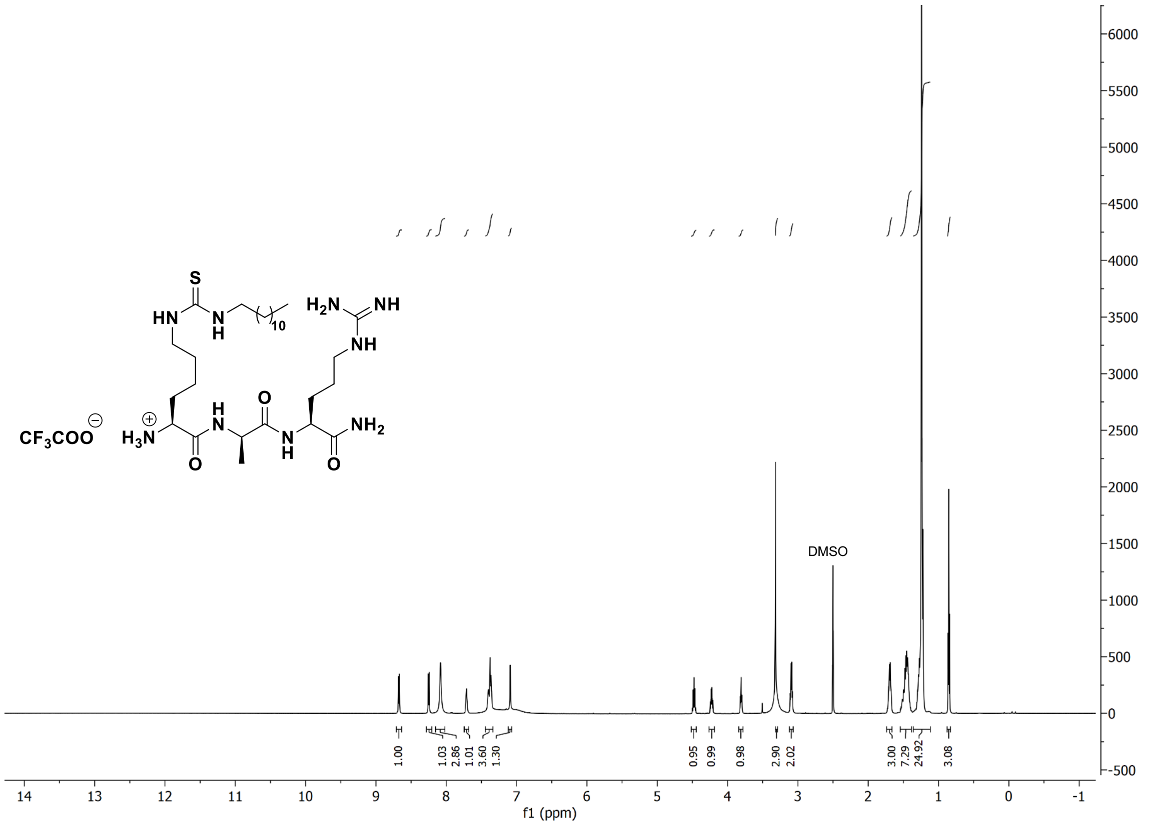

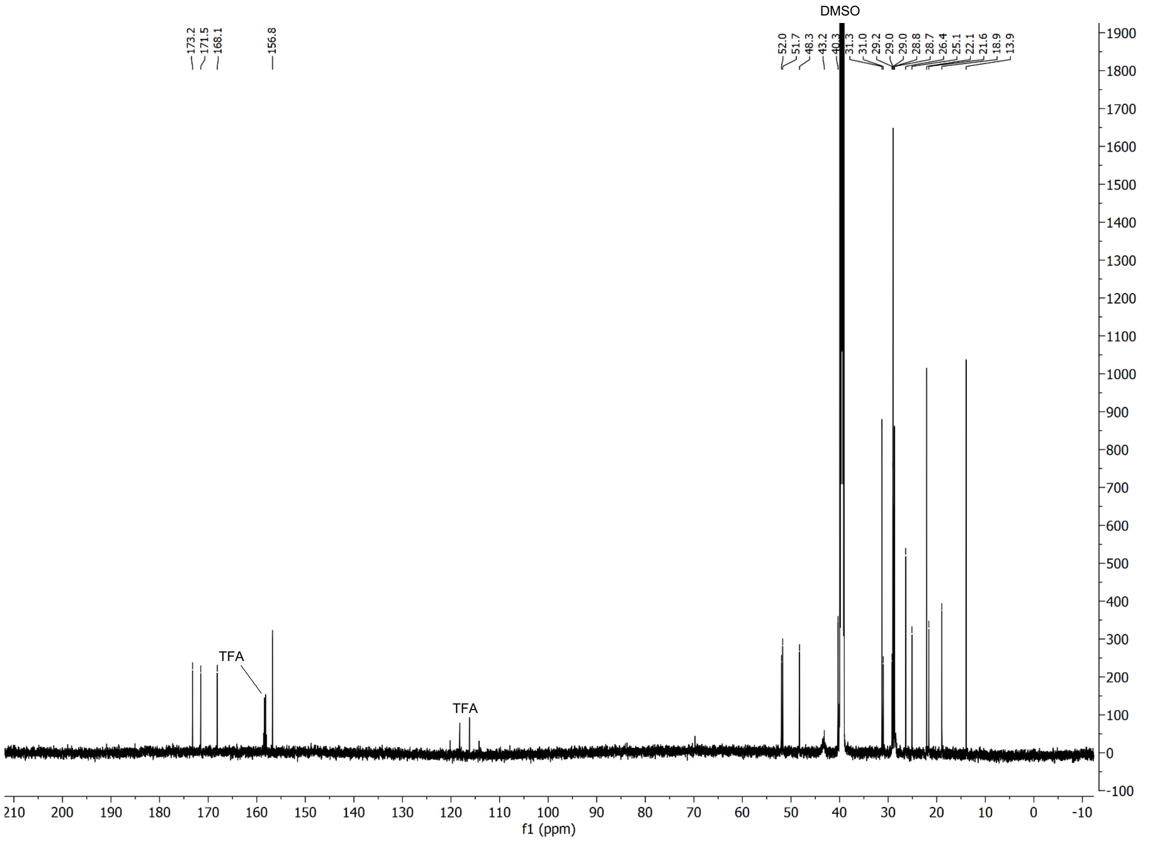

#### $^{1}$H and $^{13}$C spectra of compound **S6**

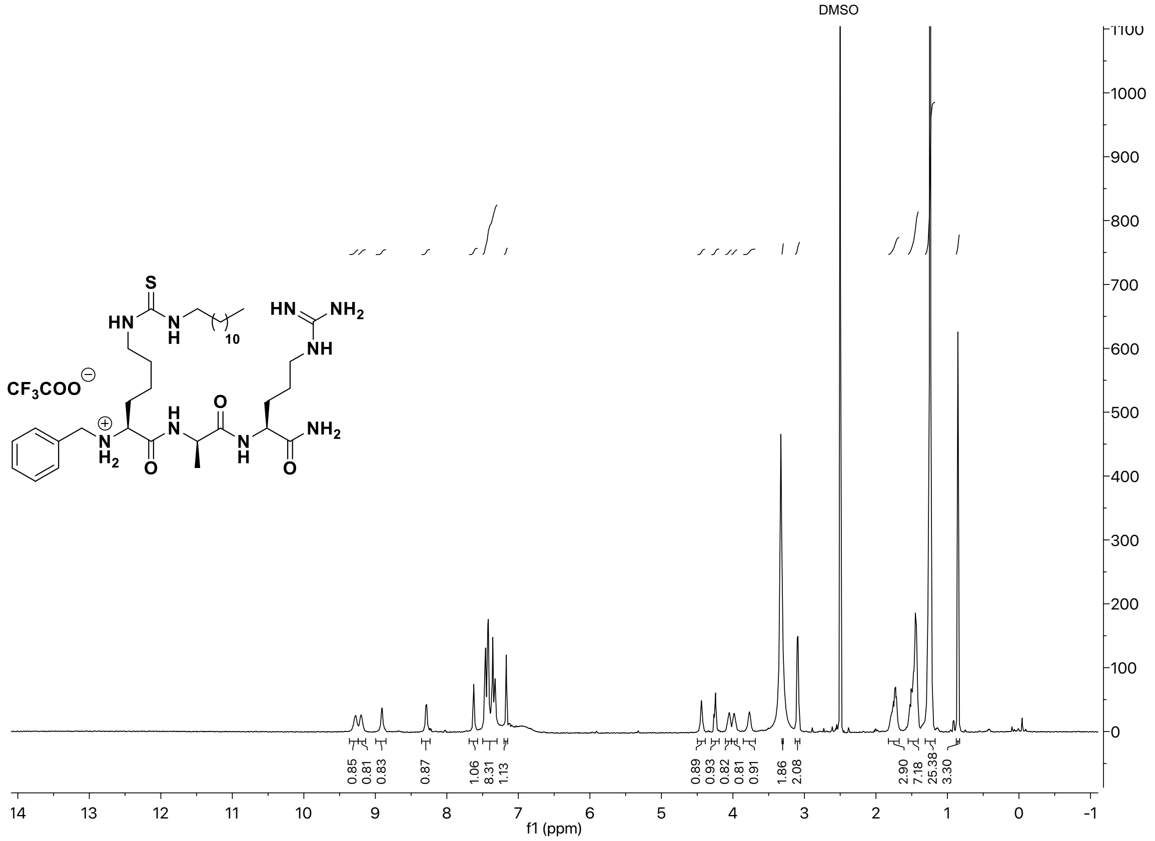

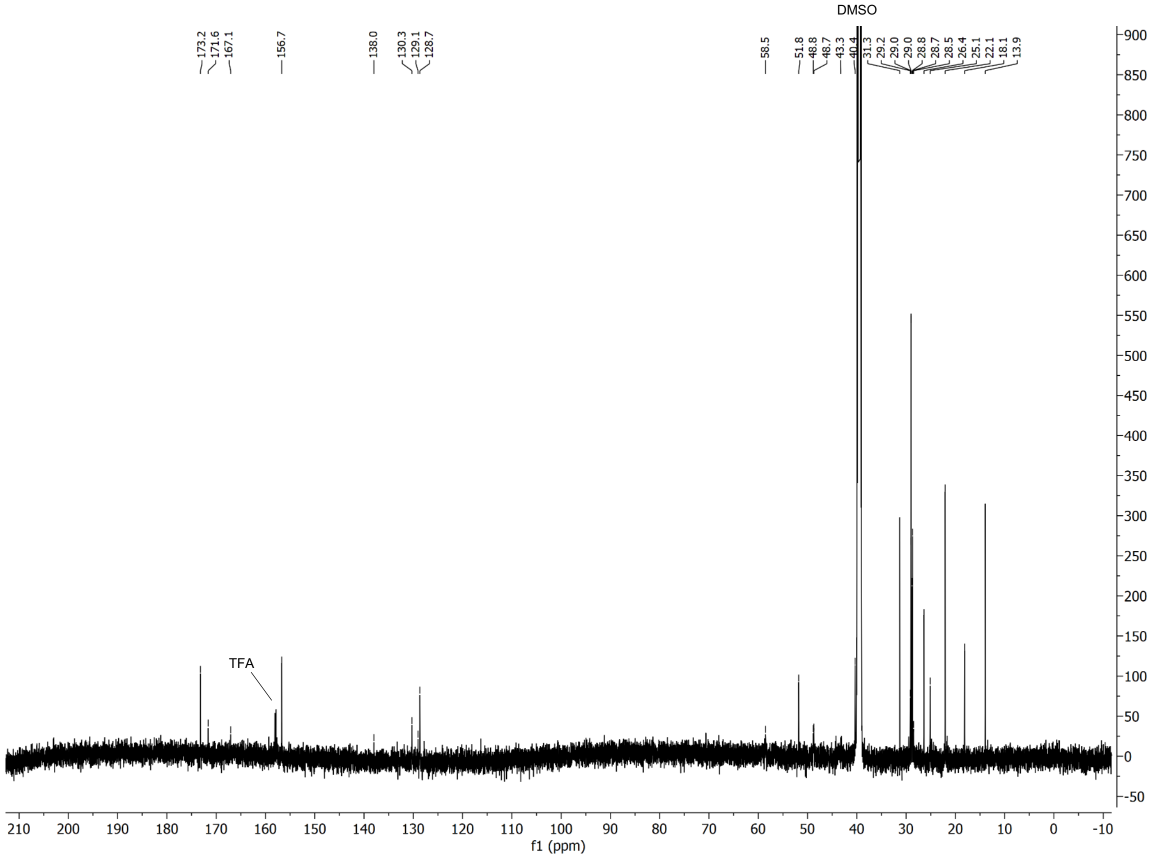

### $^{1}H$ and $^{13}C$ spectra of compound **S7**

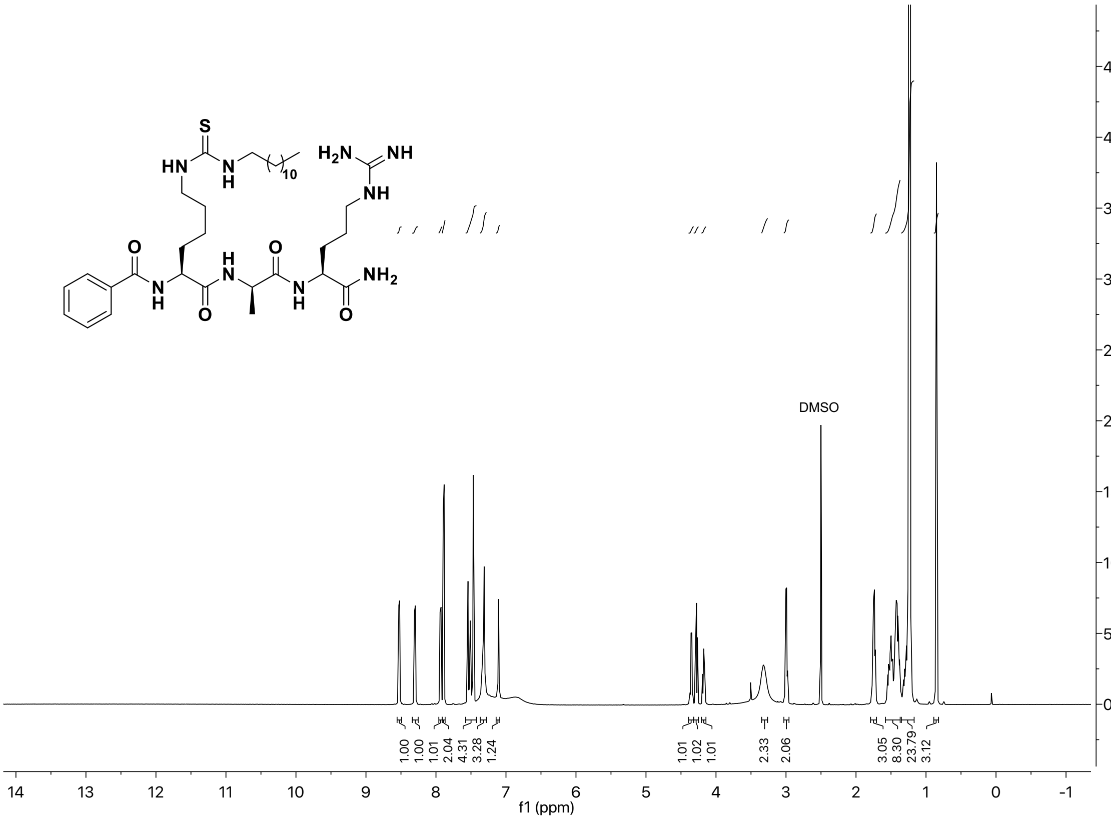

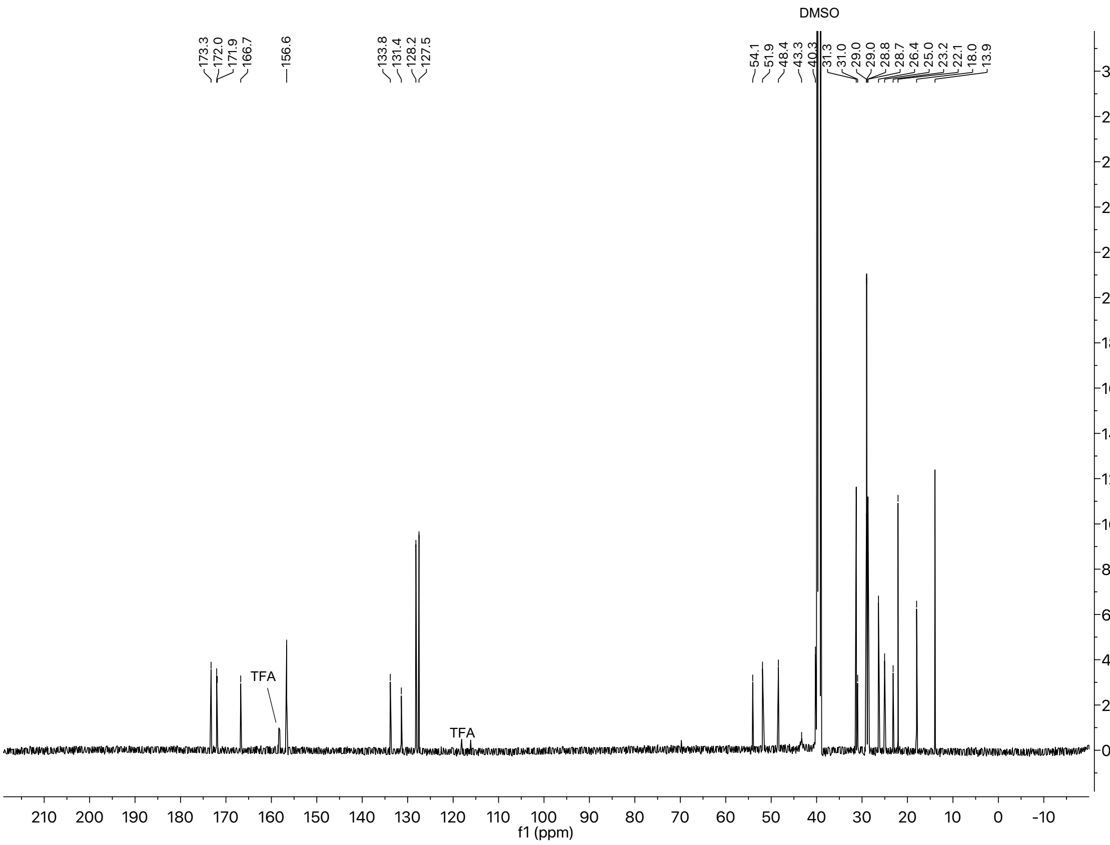

#### [1]H and [13]C spectra of compound **S8**

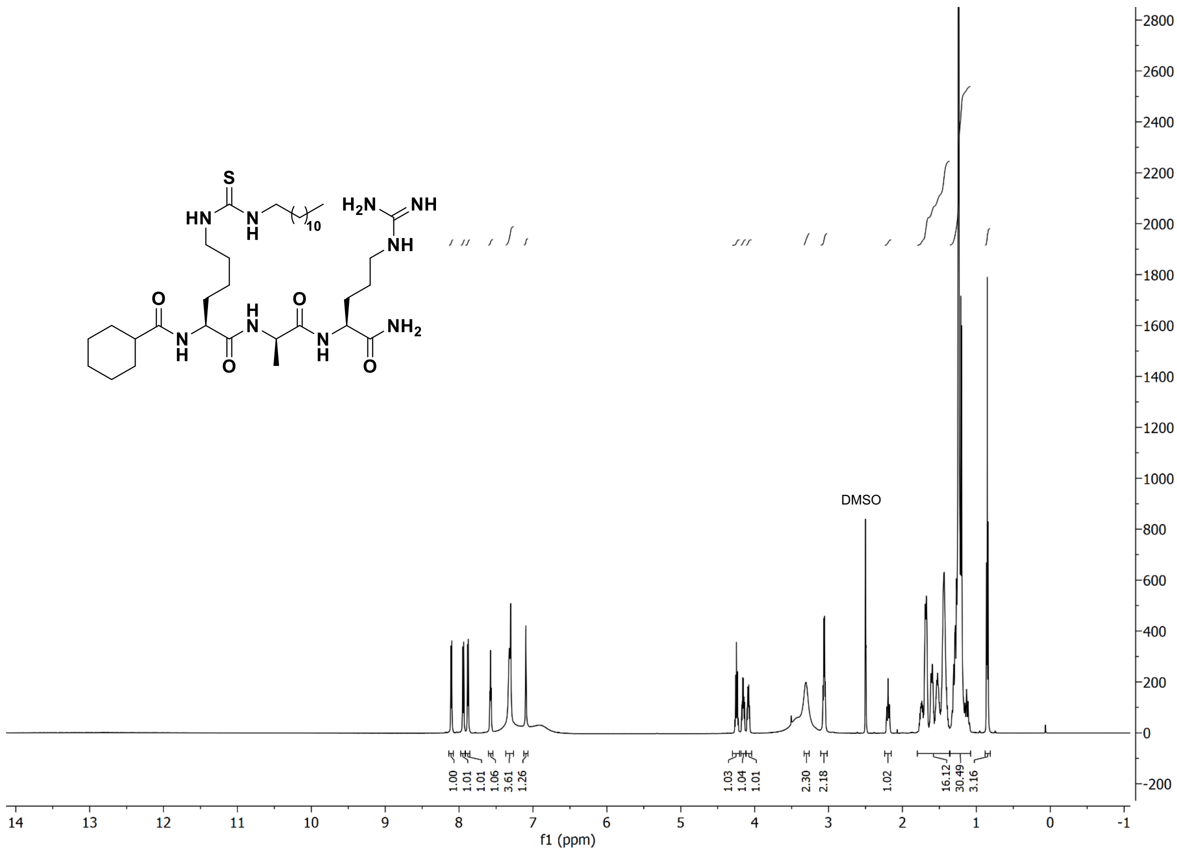

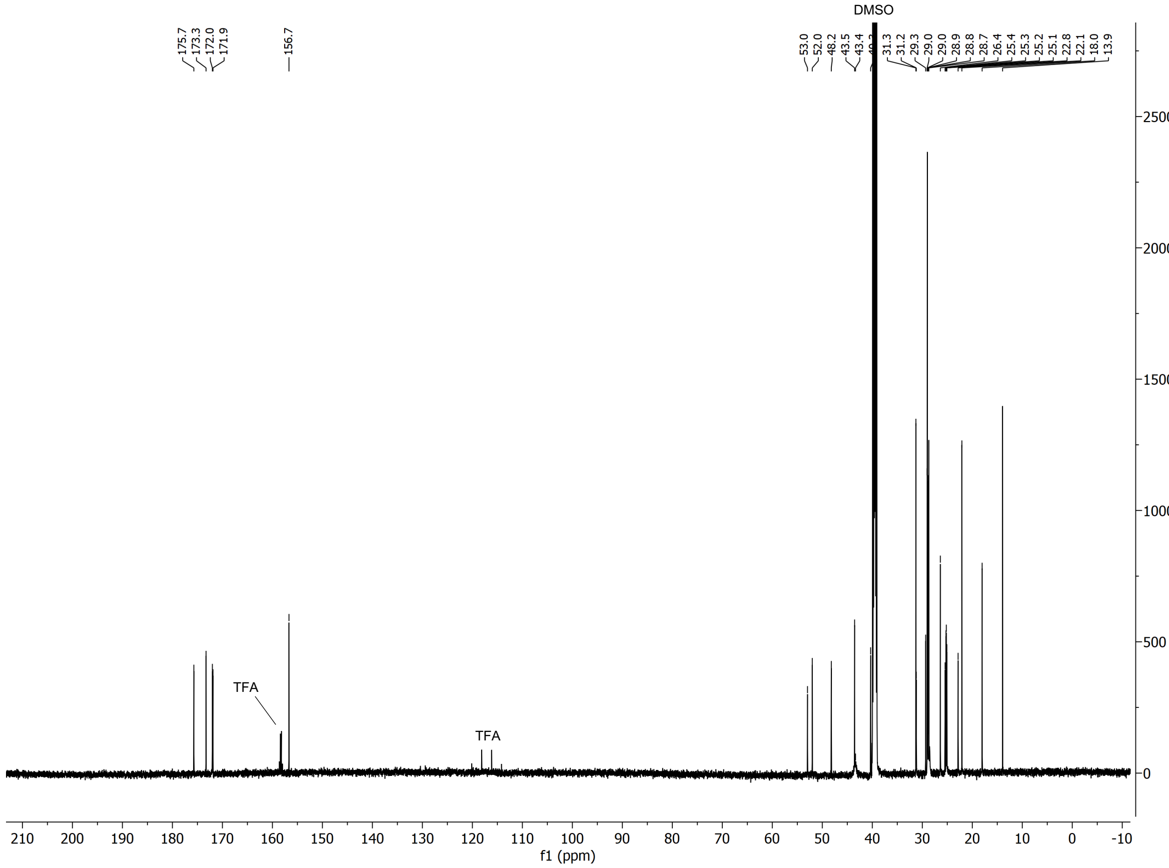

#### $^{1}$H and $^{13}$C spectra of compound **S9**

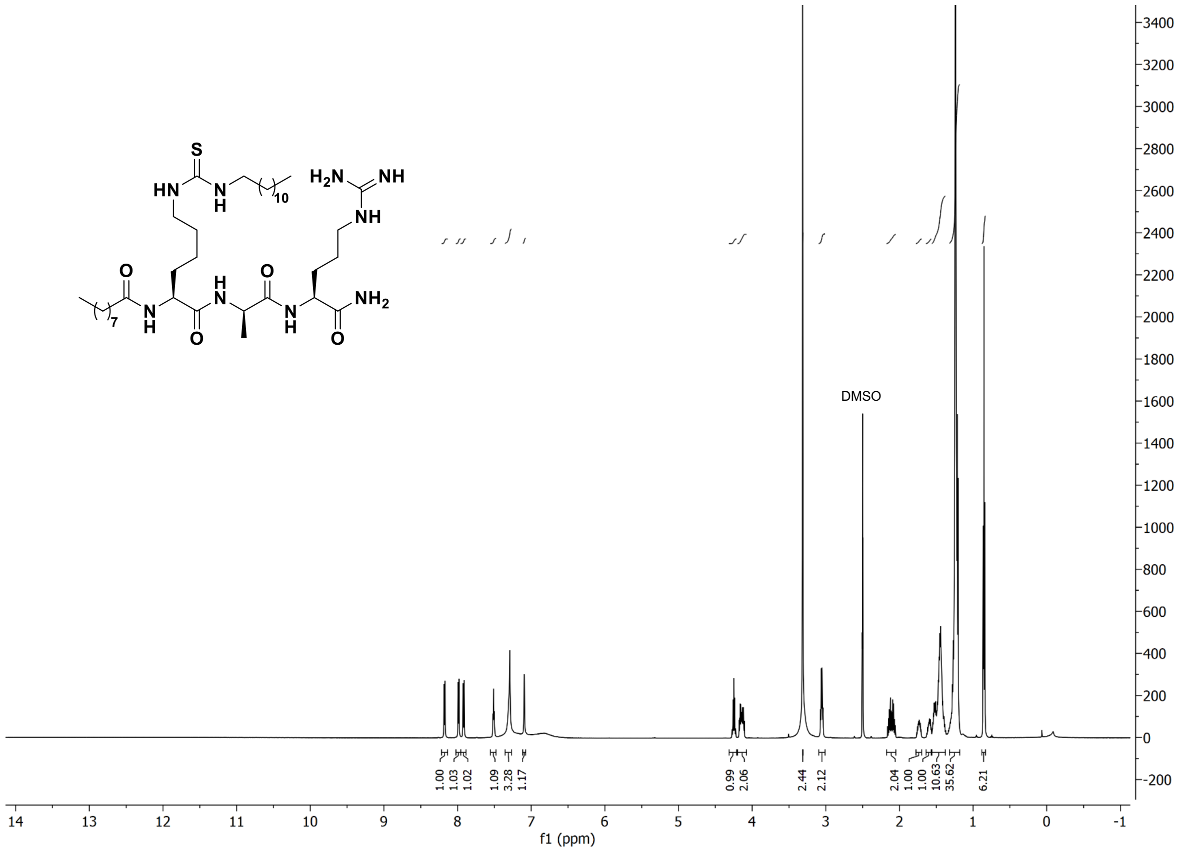

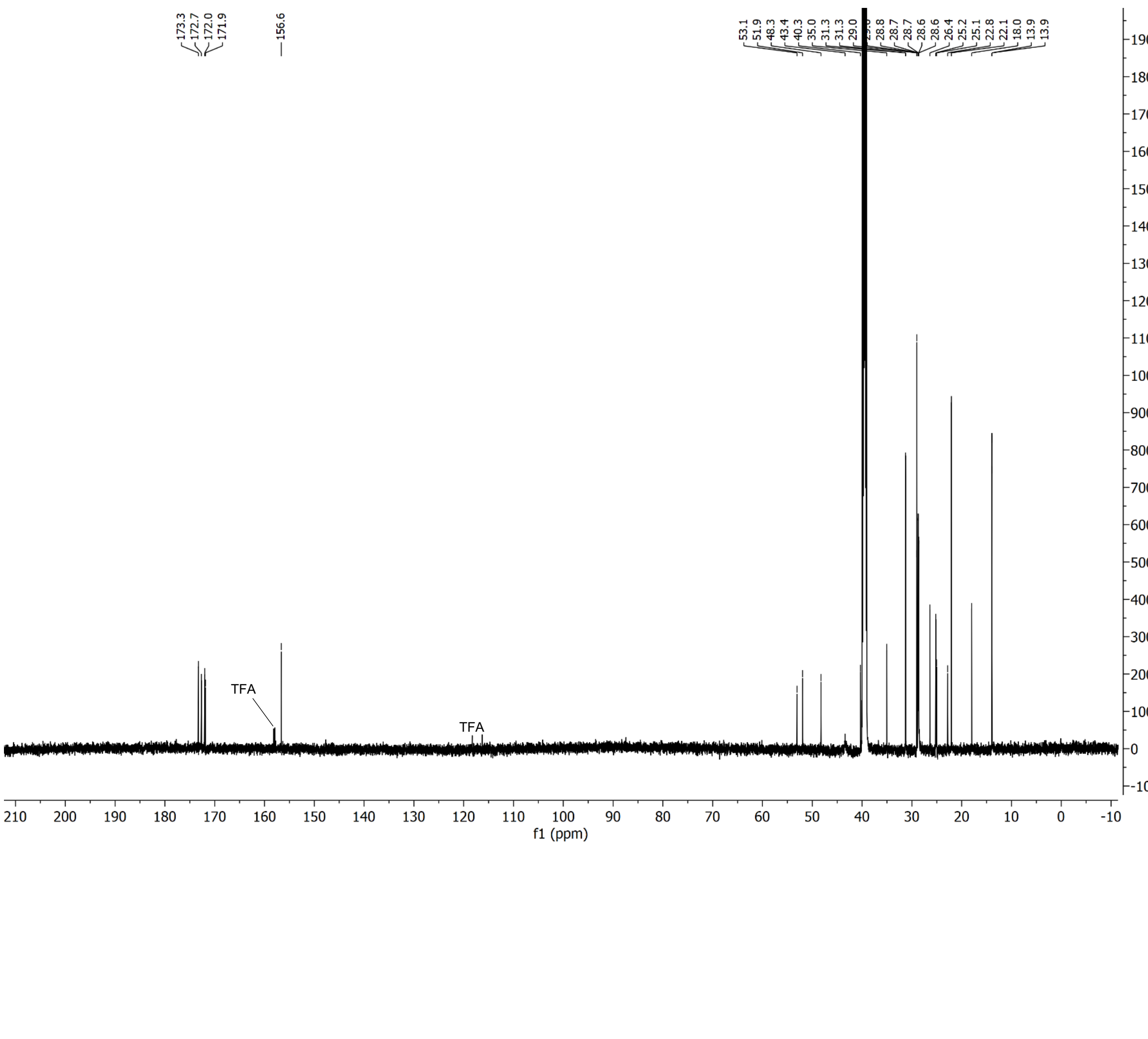

[1]H and [13]C spectra of compound **S10**

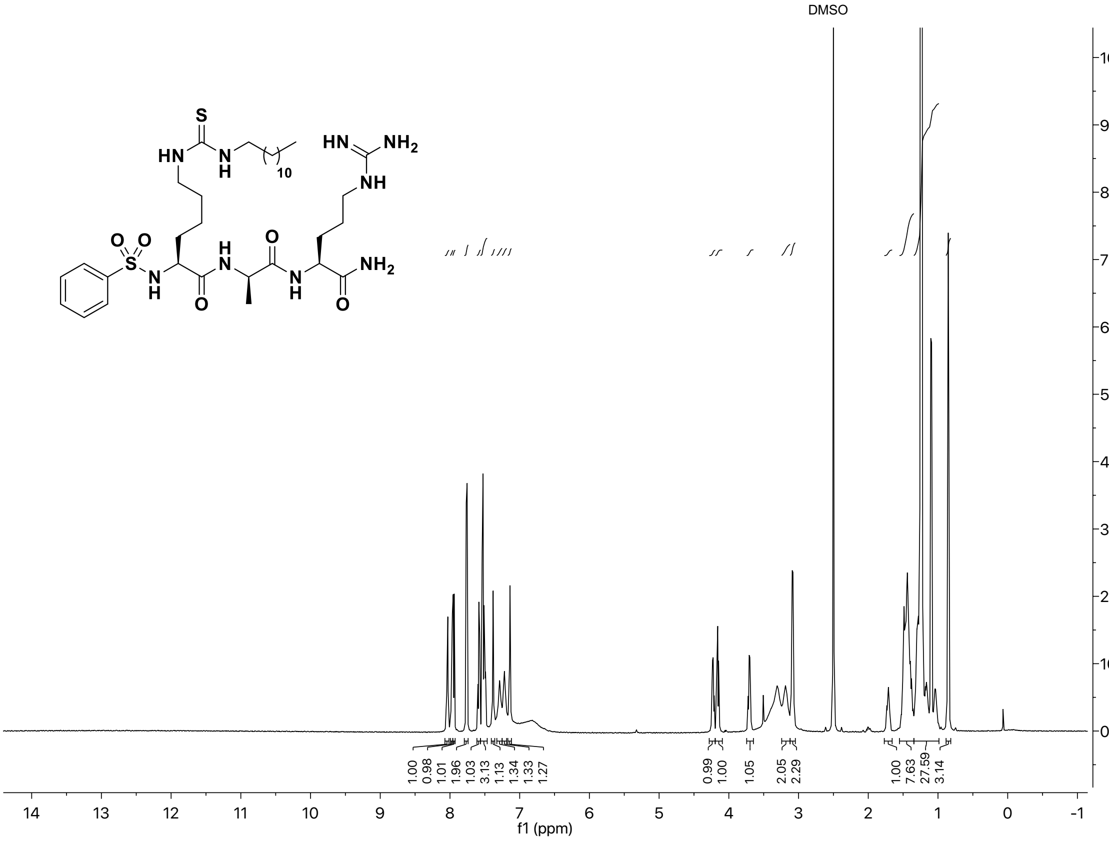

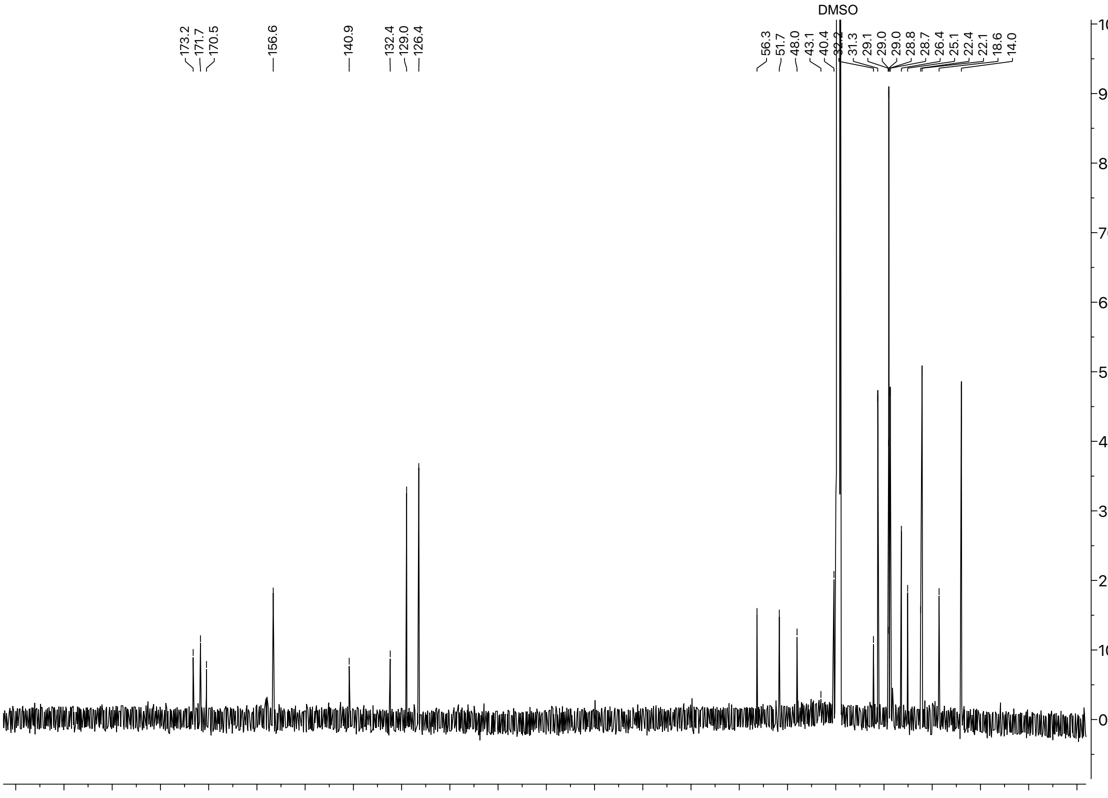

#### [1]H and [13]C spectra of compound **S11**

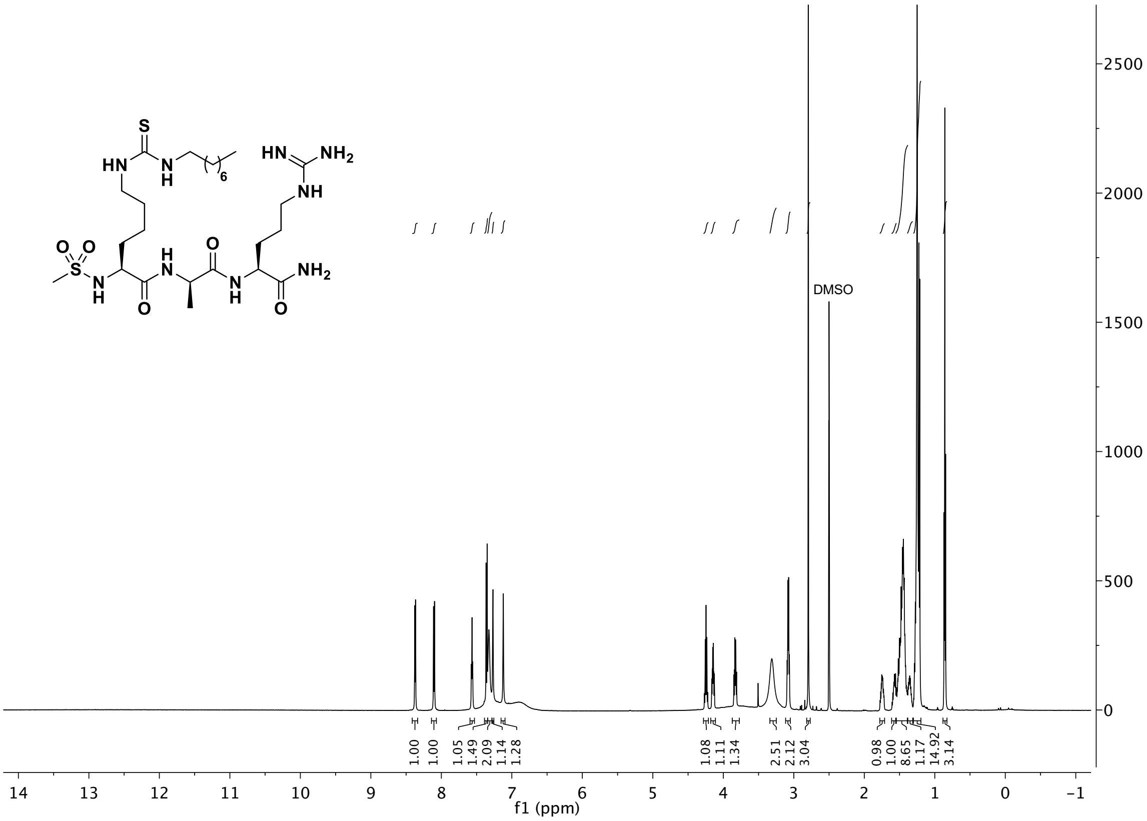

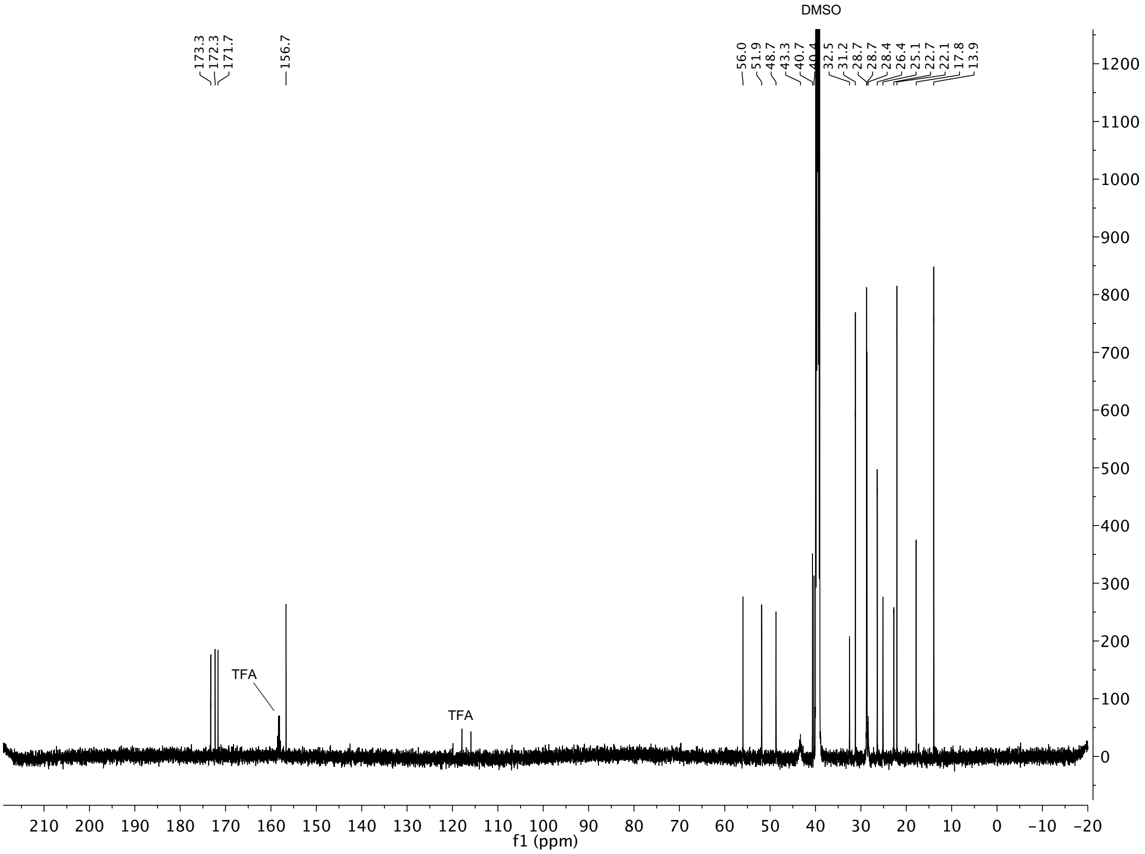

### $^{1}H$ and $^{13}C$ spectra of compound **S12**

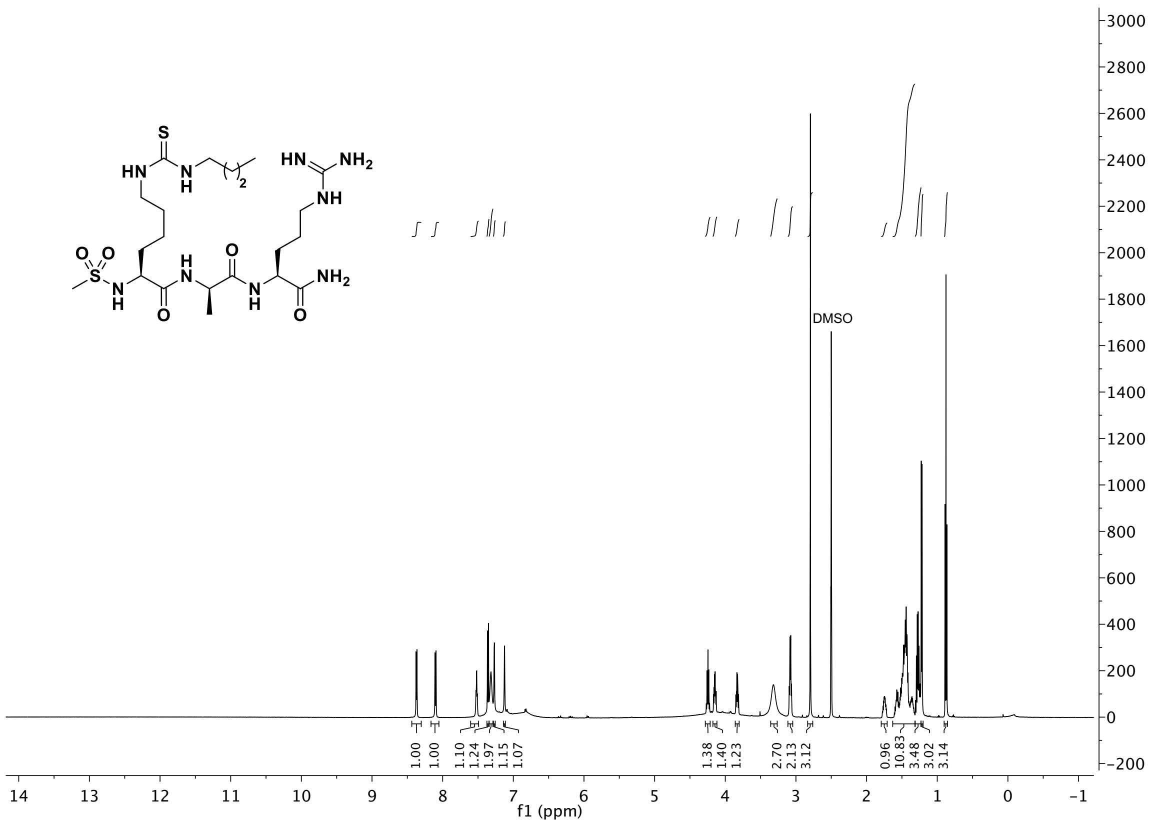

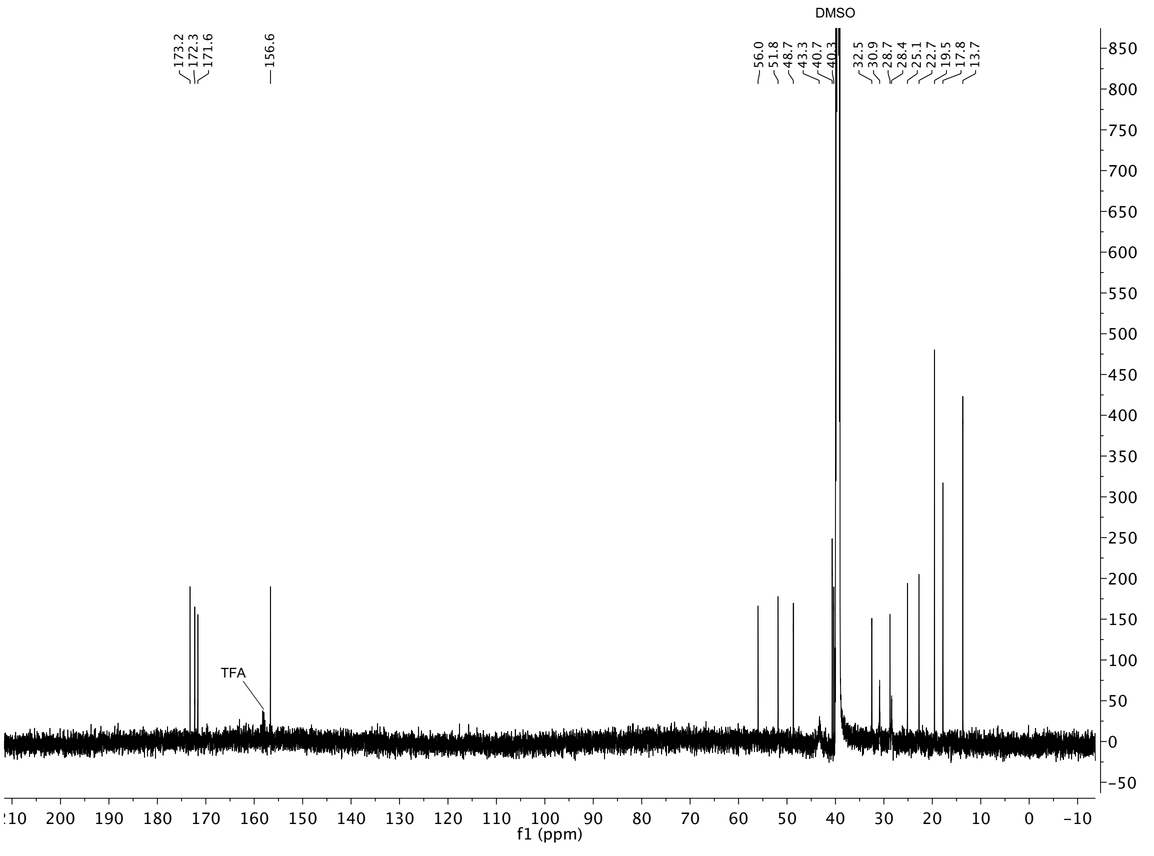

### $^{1}H$ and $^{13}C$ spectra of compound **S13**

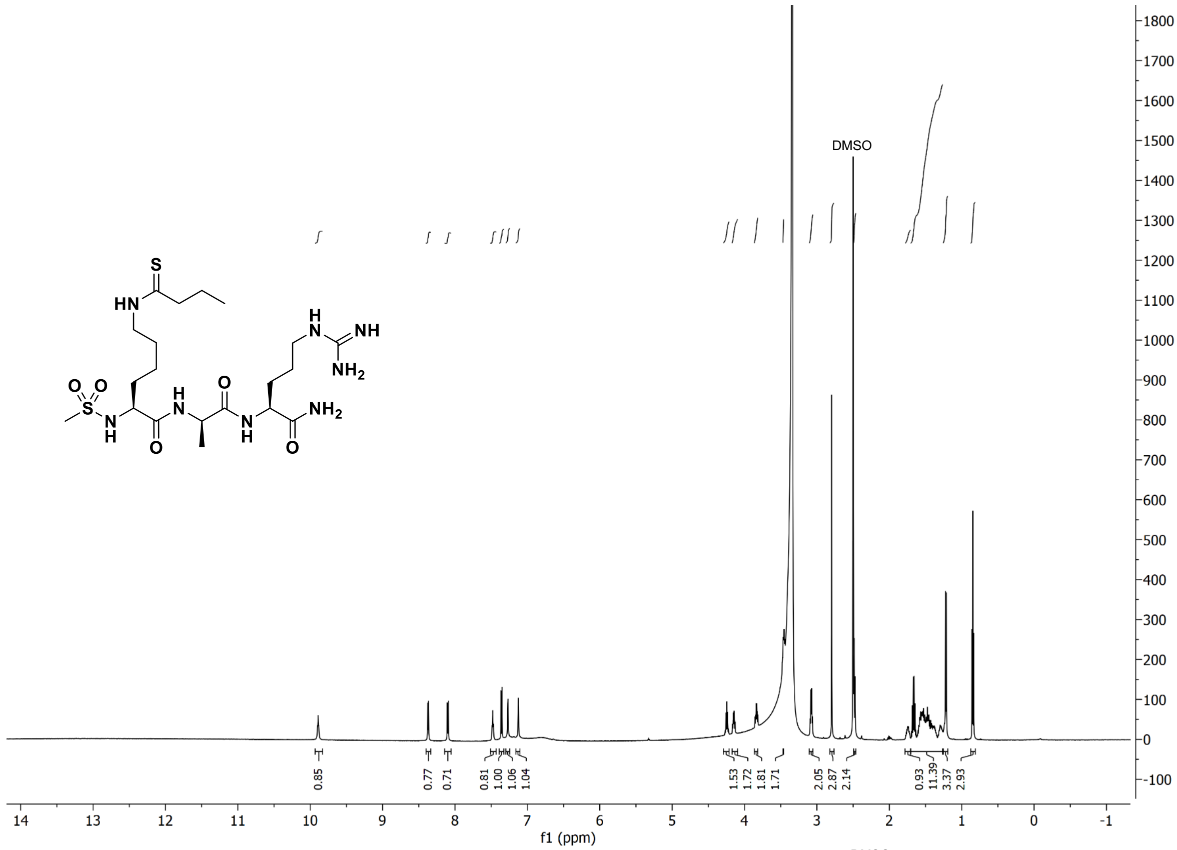

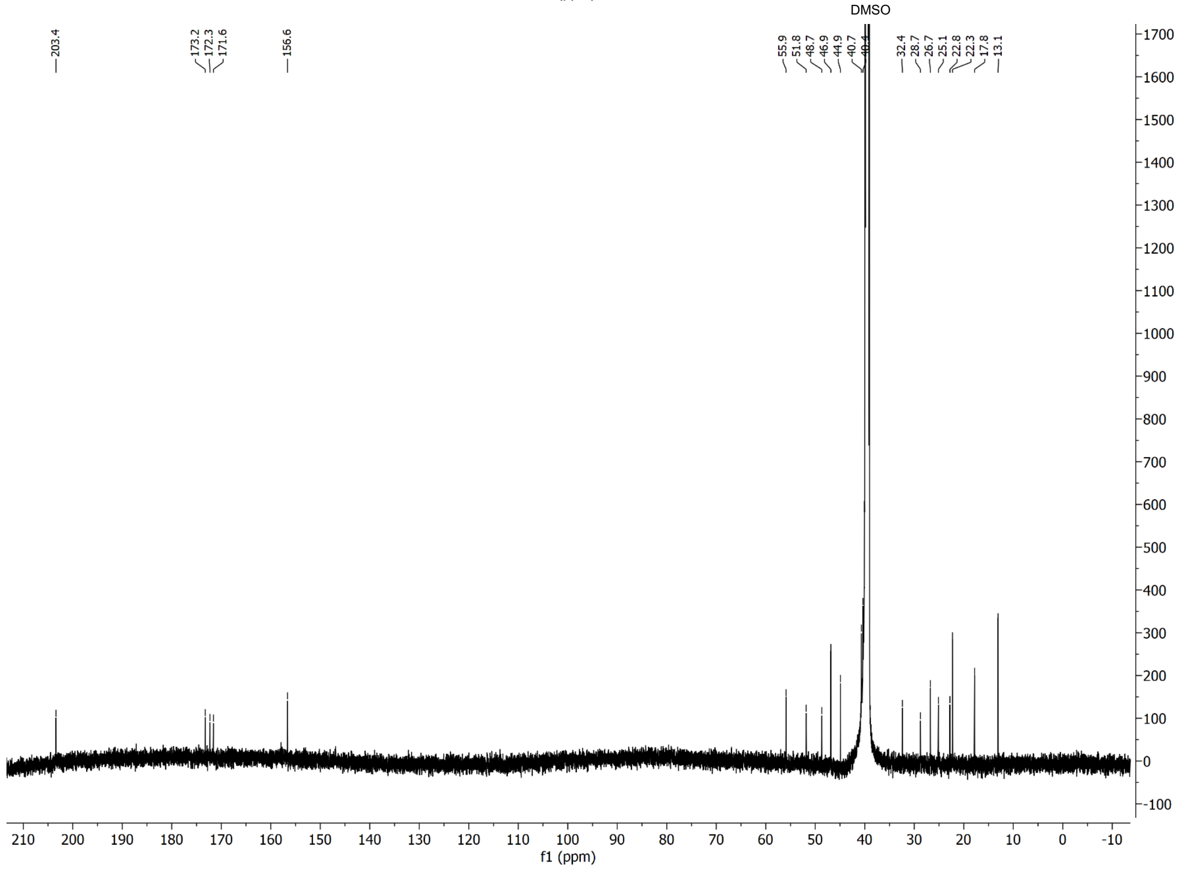

### $^{1}H$ and $^{13}C$ spectra of compound **S14**

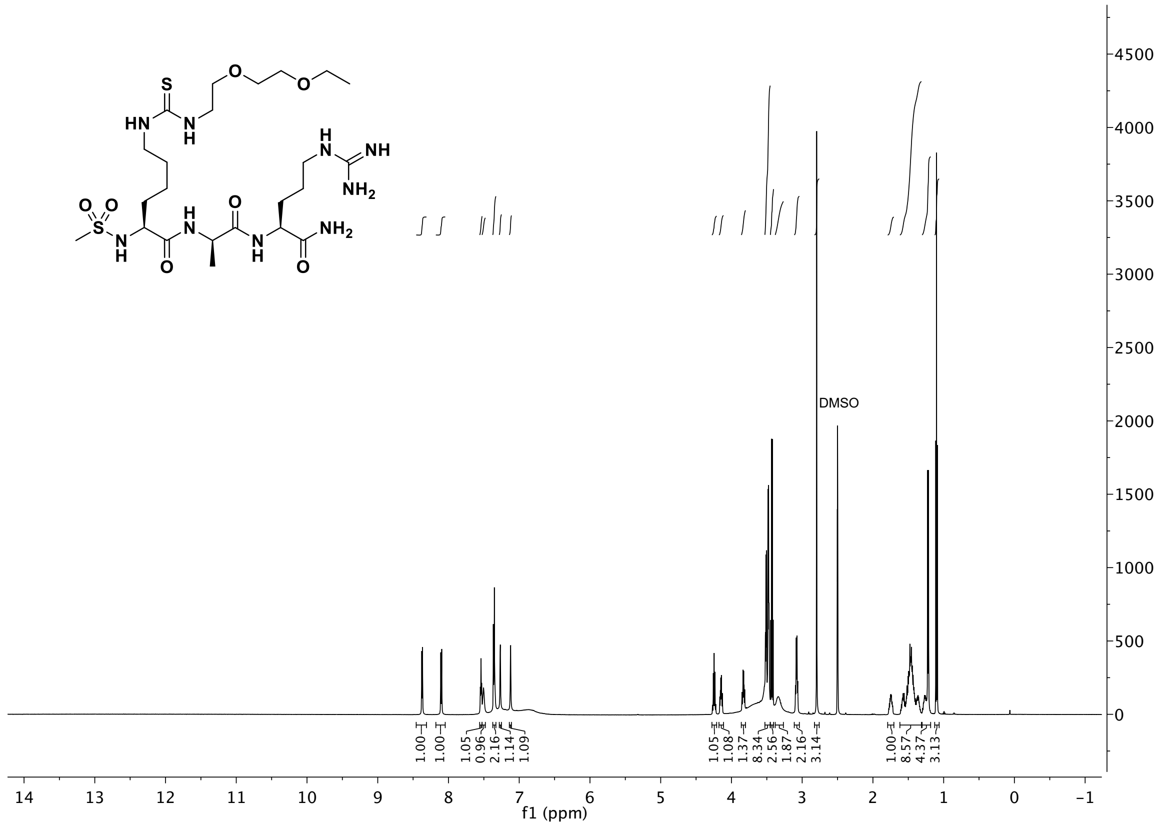

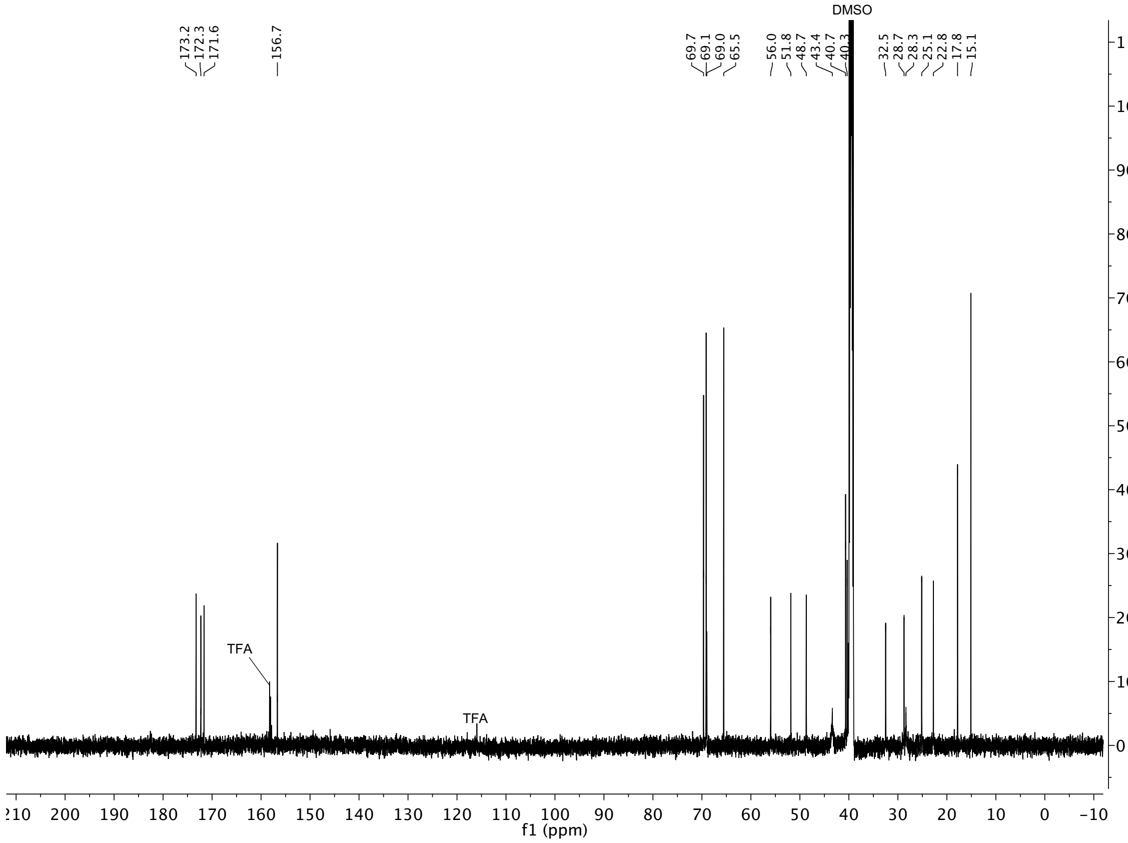

#### $^{1}H$ and $^{13}C$ spectra of compound **S15**

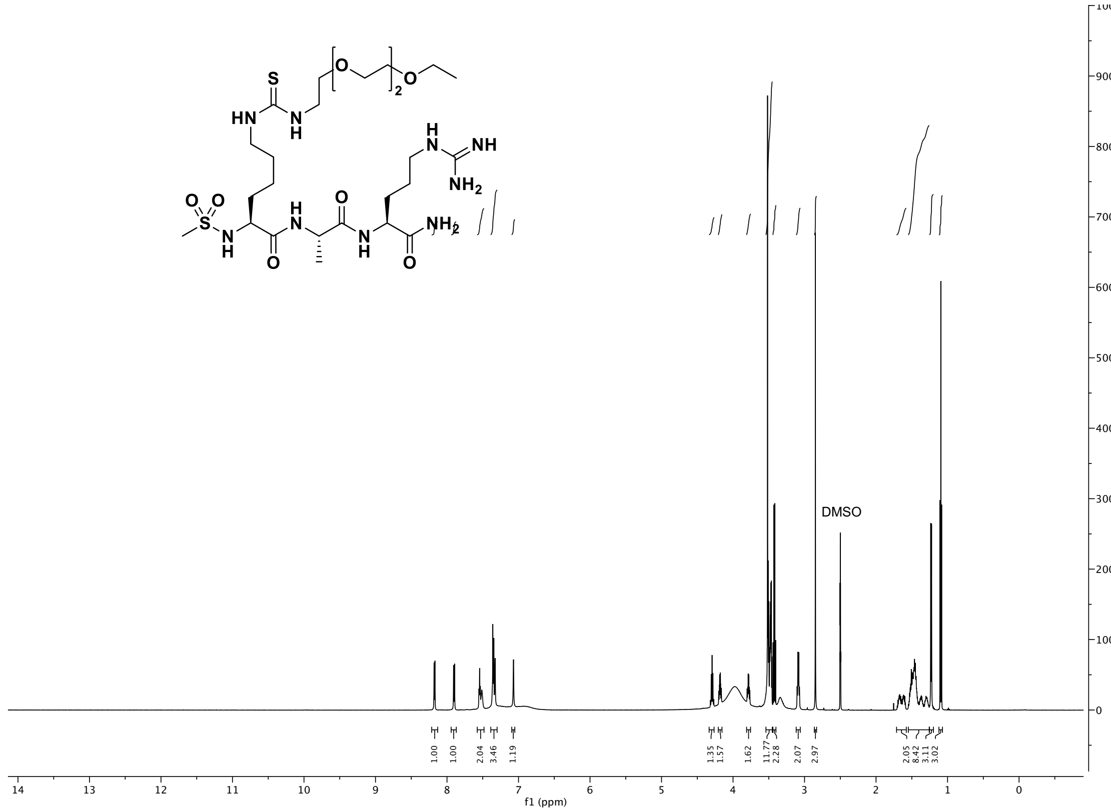

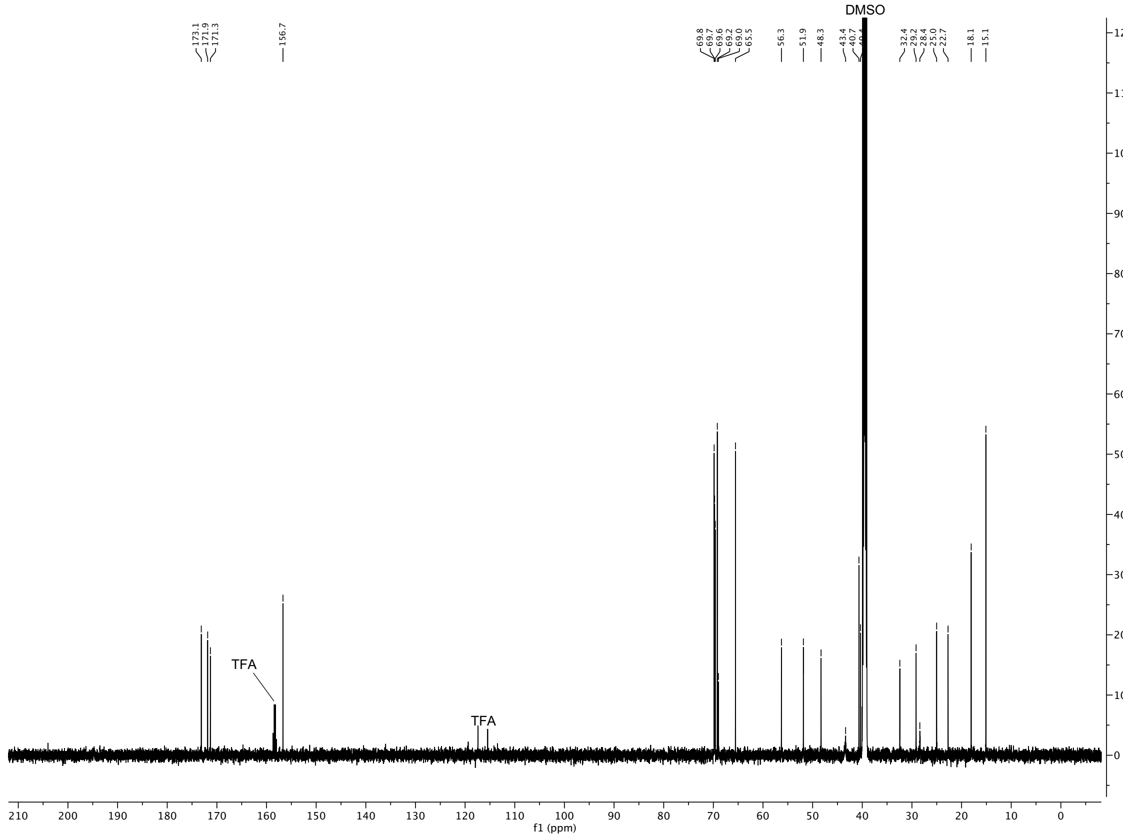

#### $^1$H and $^{13}$C spectra of compound **S16**

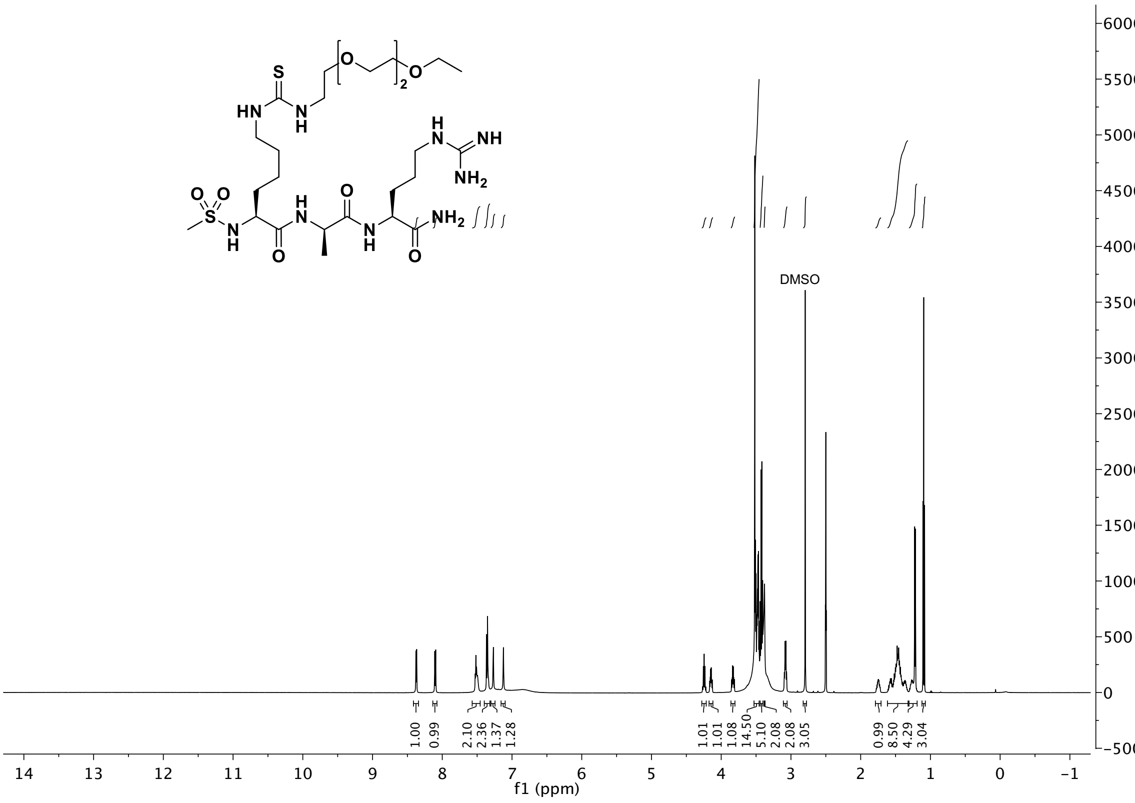

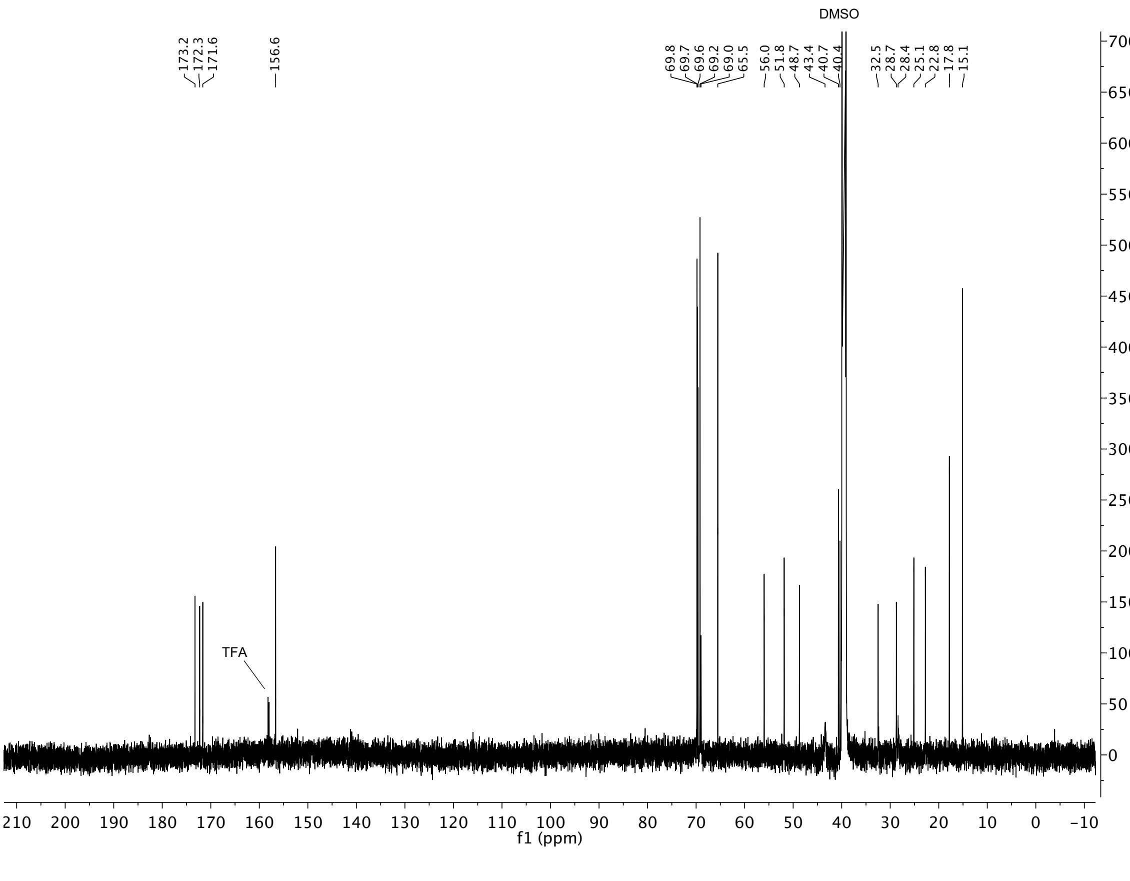

$^{1}H$ and $^{13}C$ spectra of compound **S17**

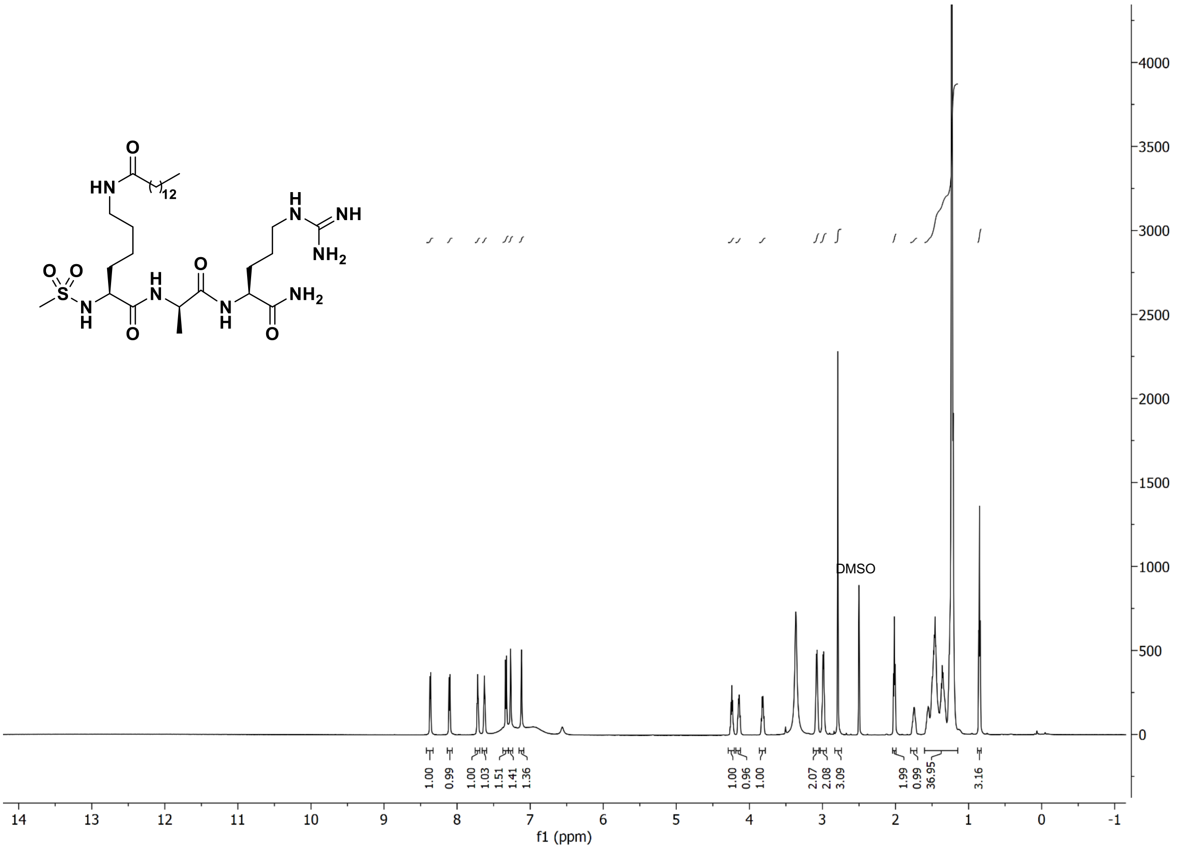

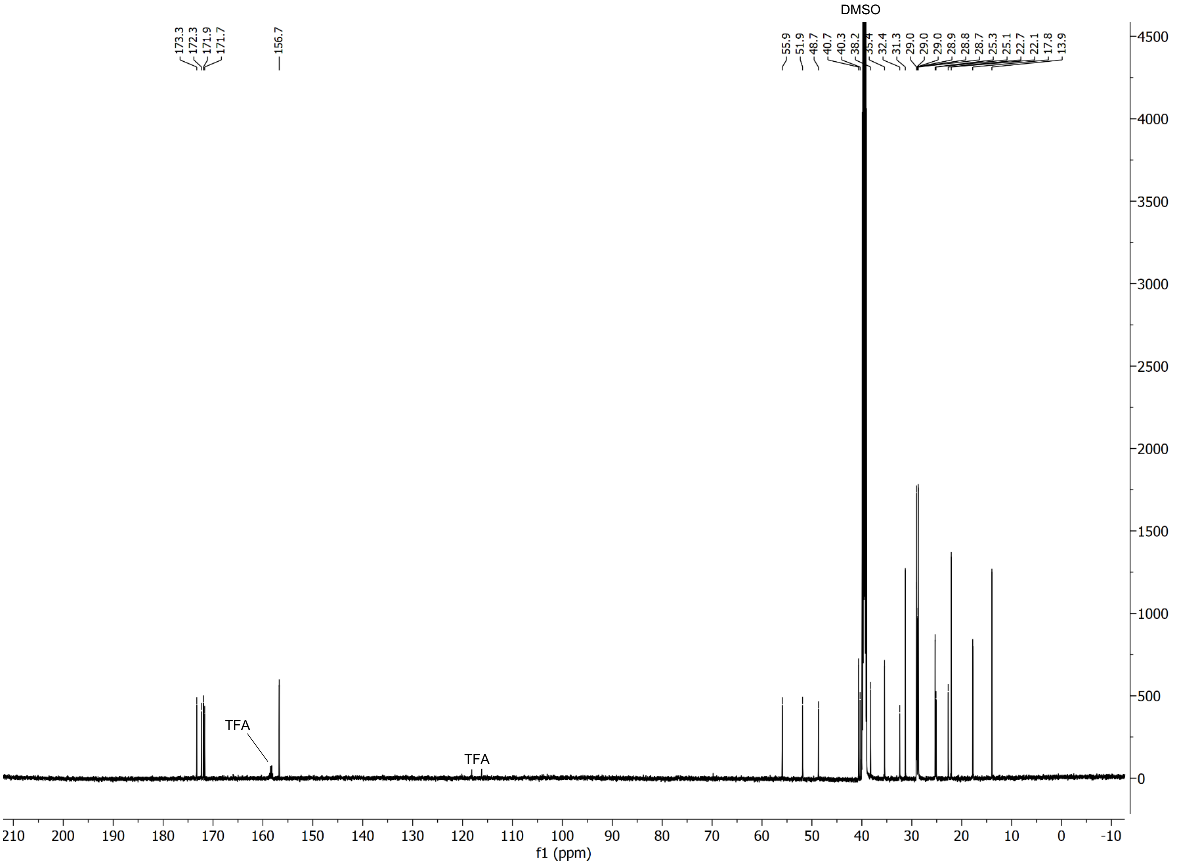

#### [1]H and [13]C spectra of compound **S18**

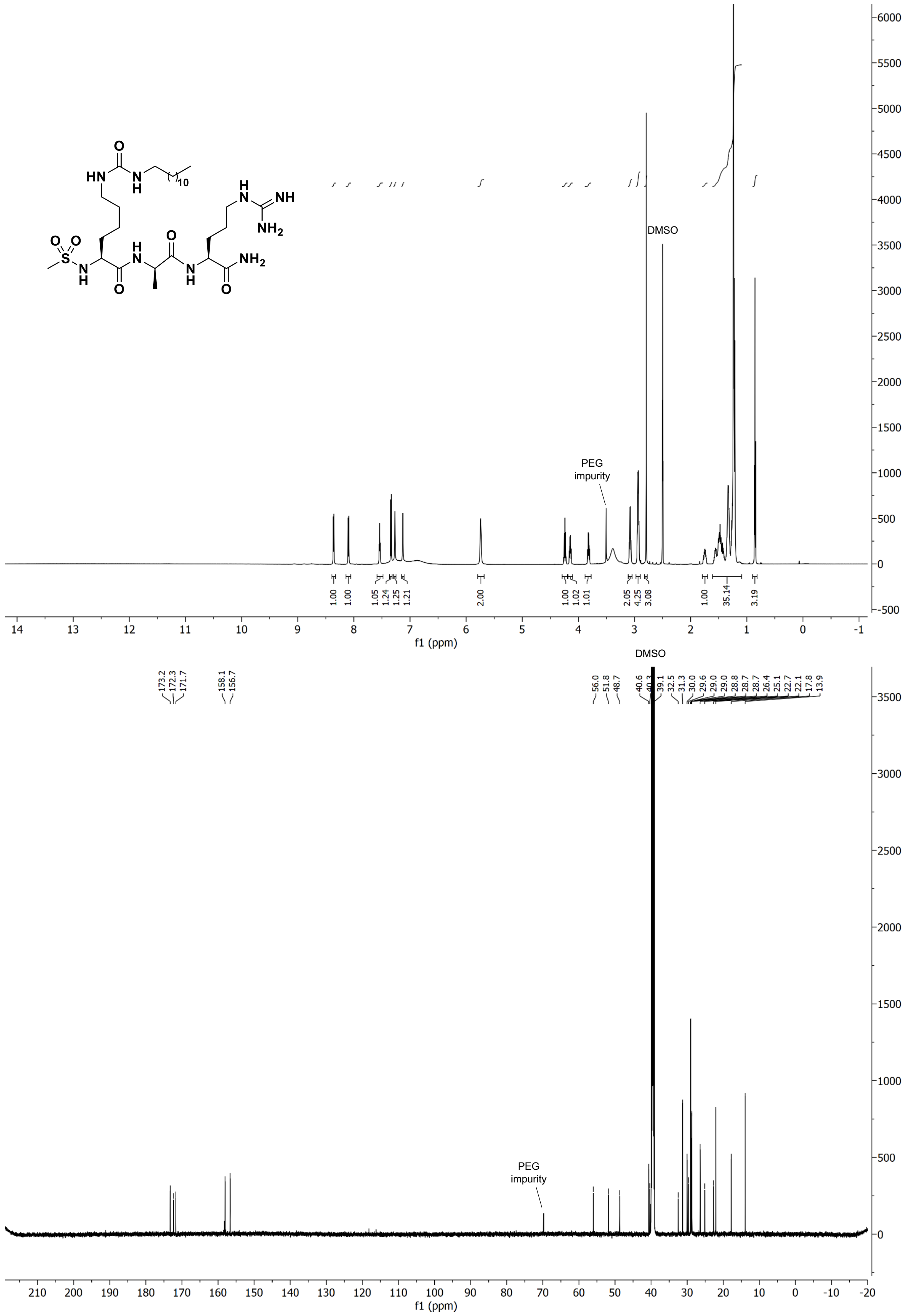

### $^{1}H$ and $^{13}C$ spectra of compound **S19**

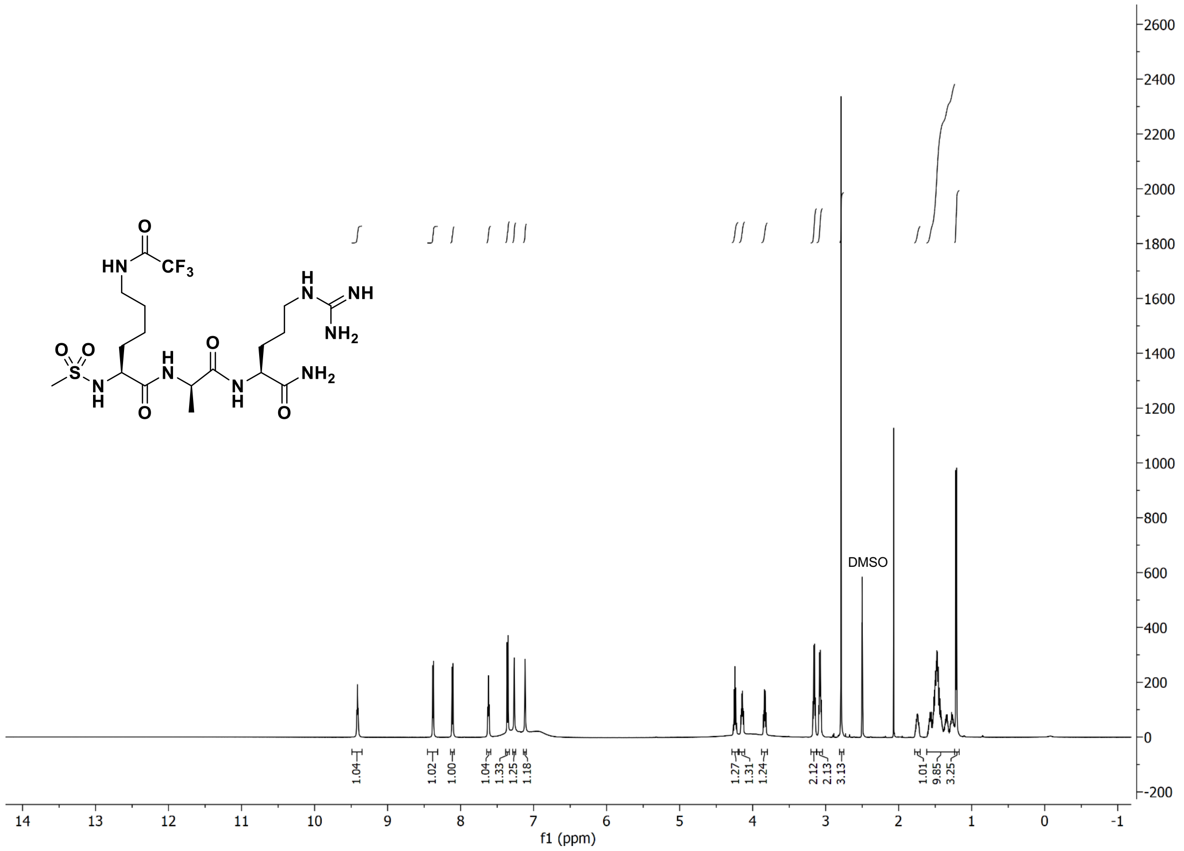

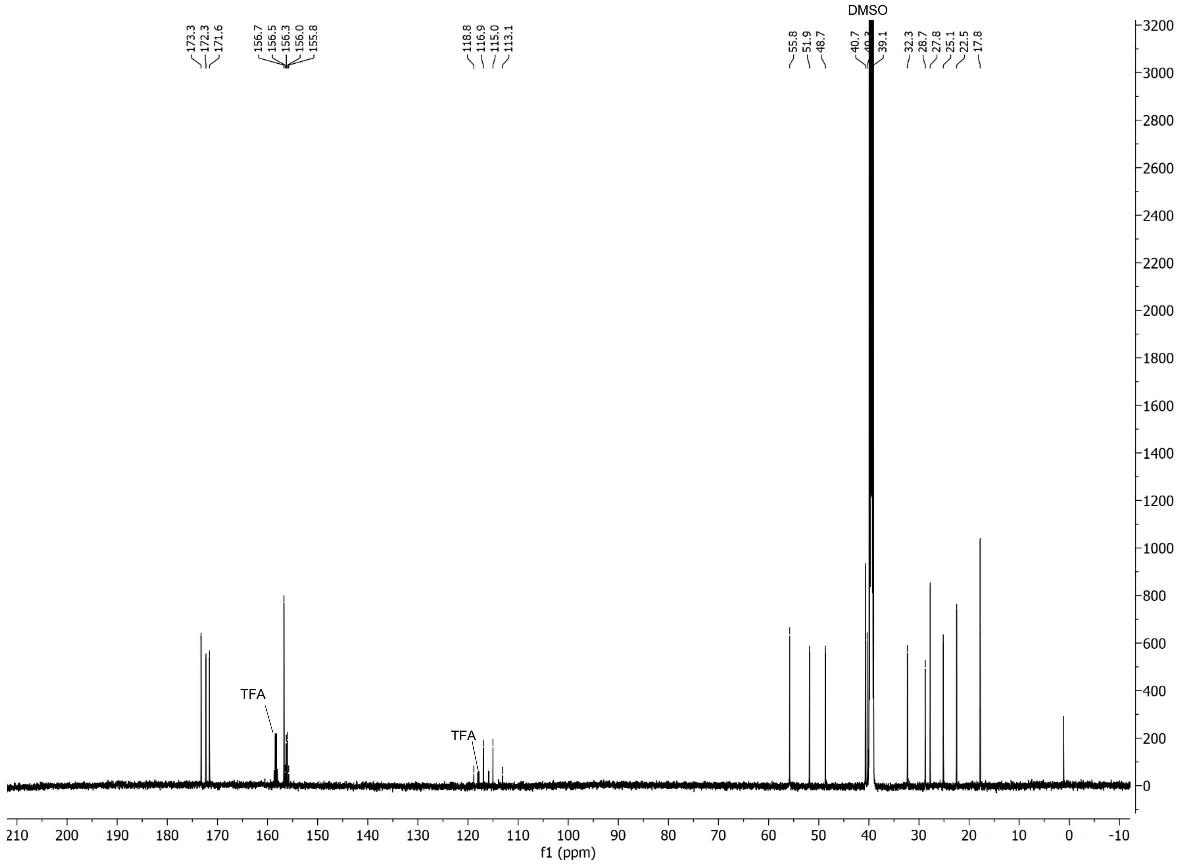

#### $^{19}$F spectrum of compound **S19**

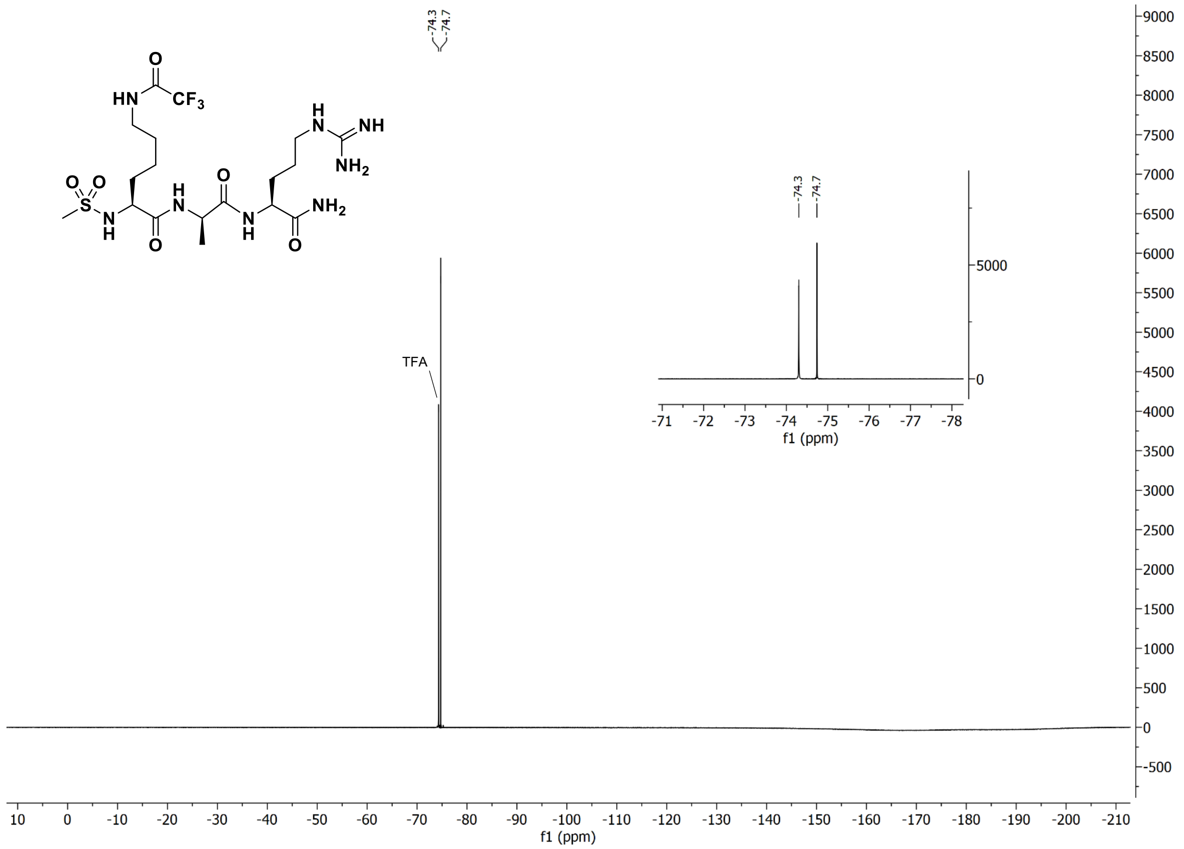

### $^{1}$H and $^{13}$C spectra of compound **S20**

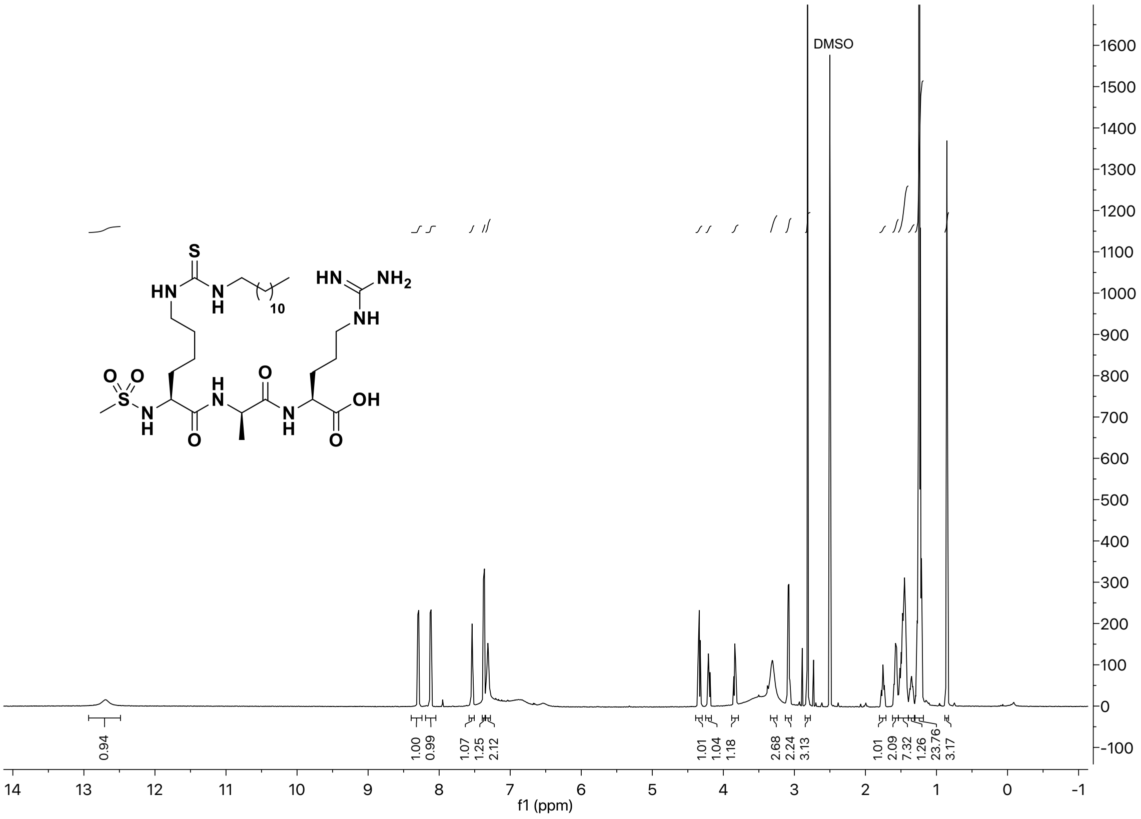

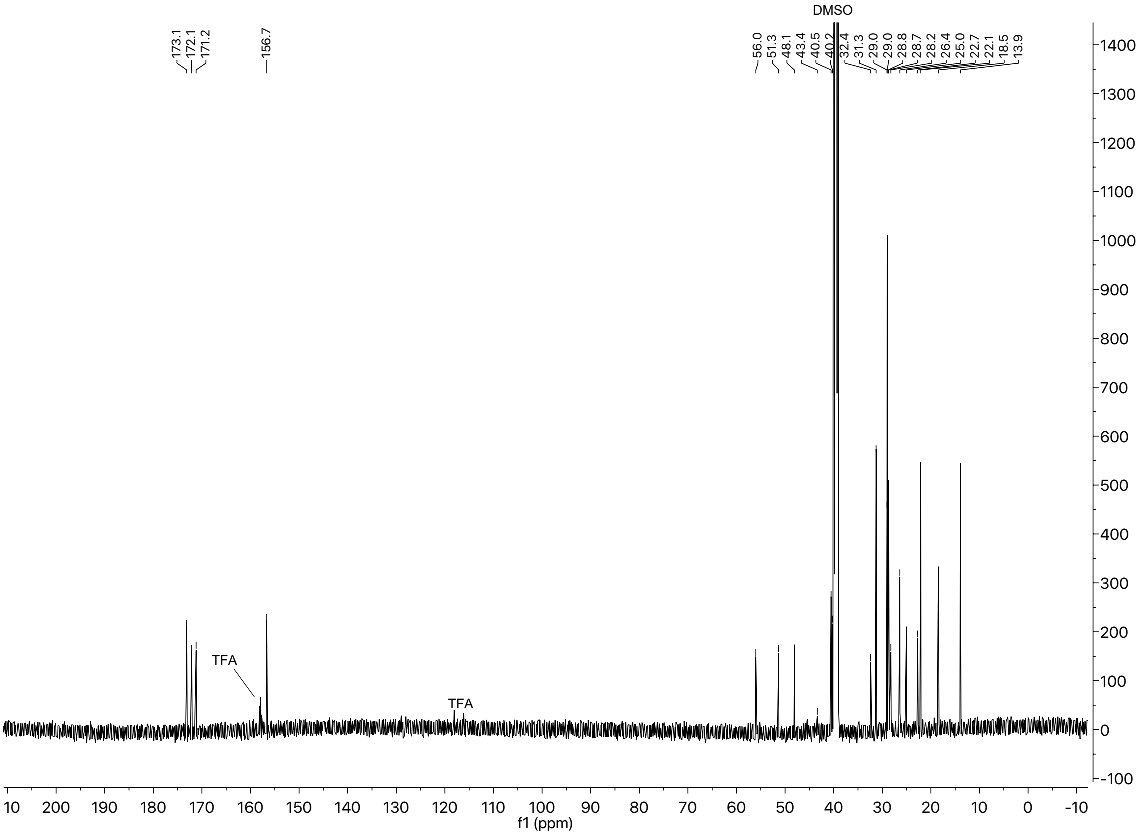

### $^{1}H$ and $^{13}C$ spectra of compound **S21**

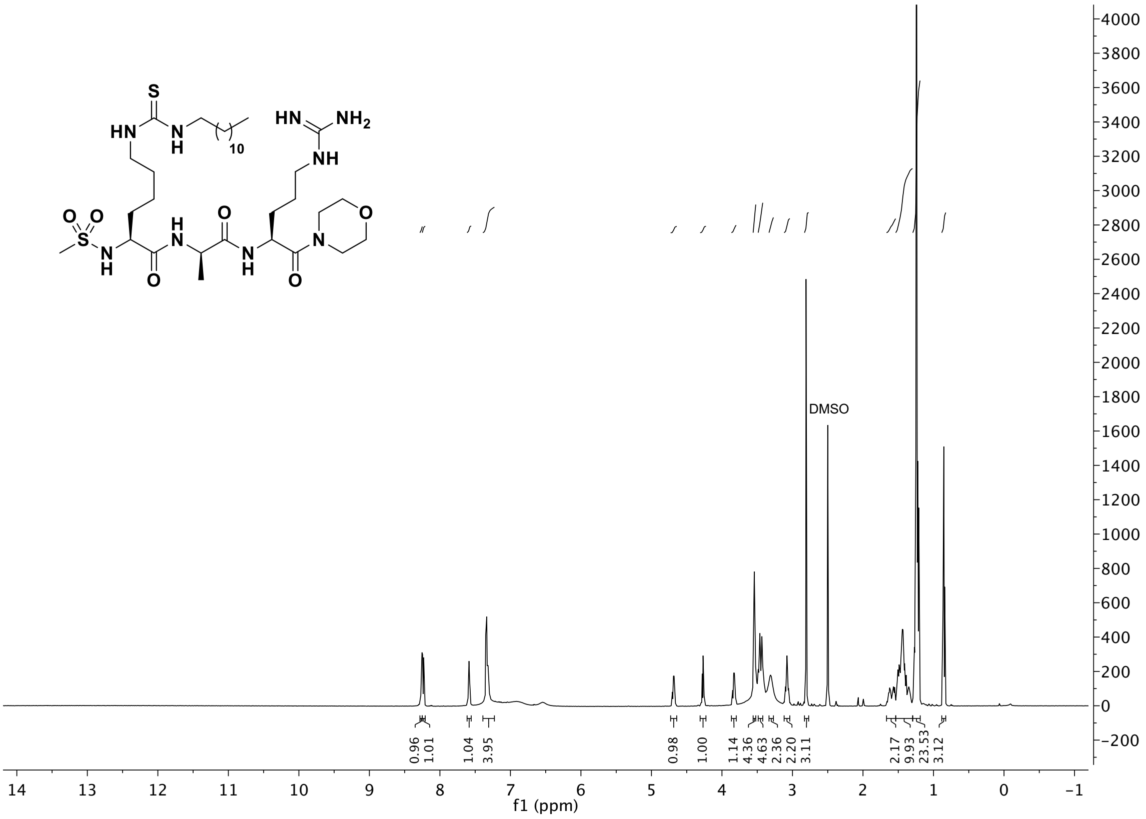

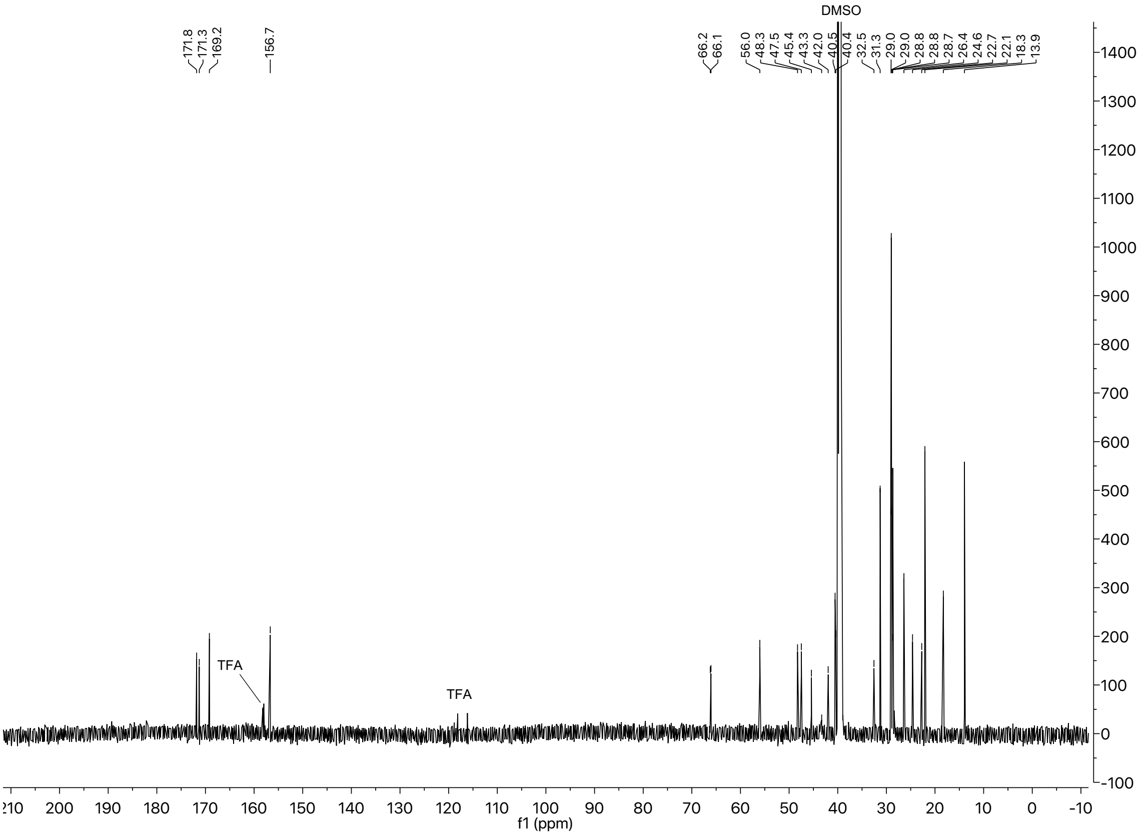

### $^{1}$H and $^{13}$C spectra of compound **S22**

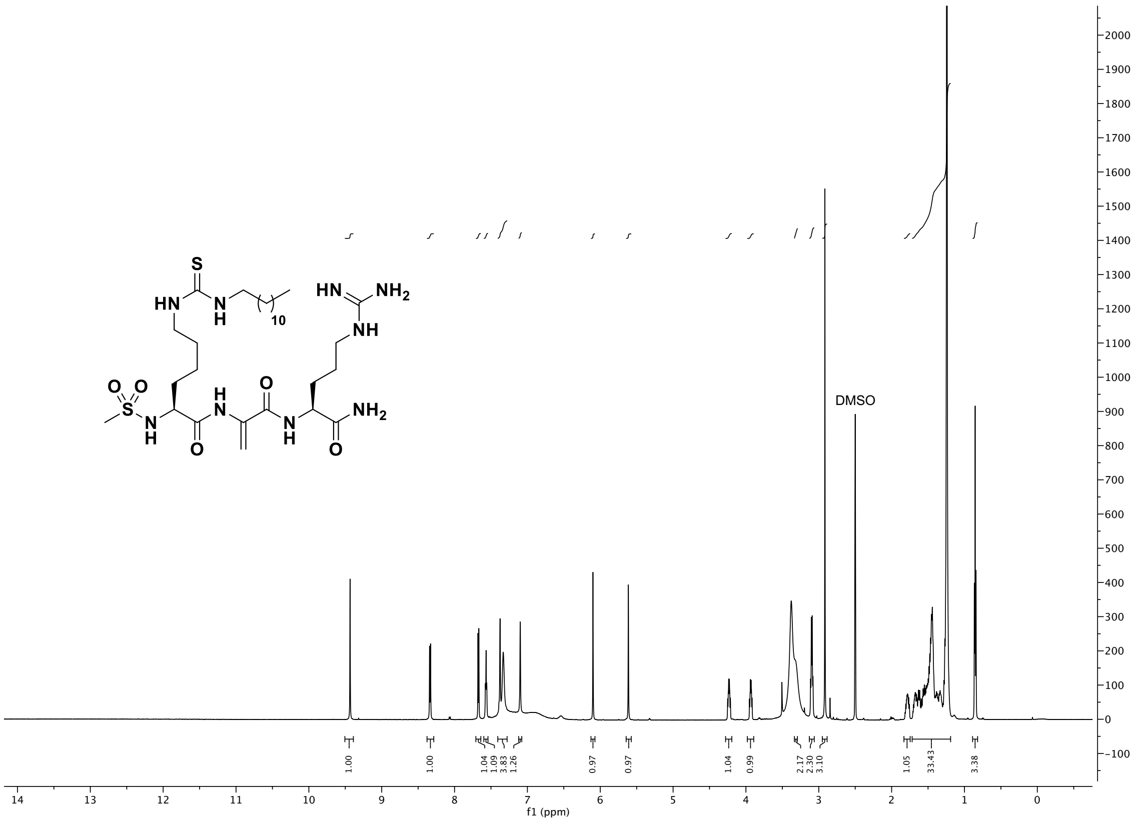

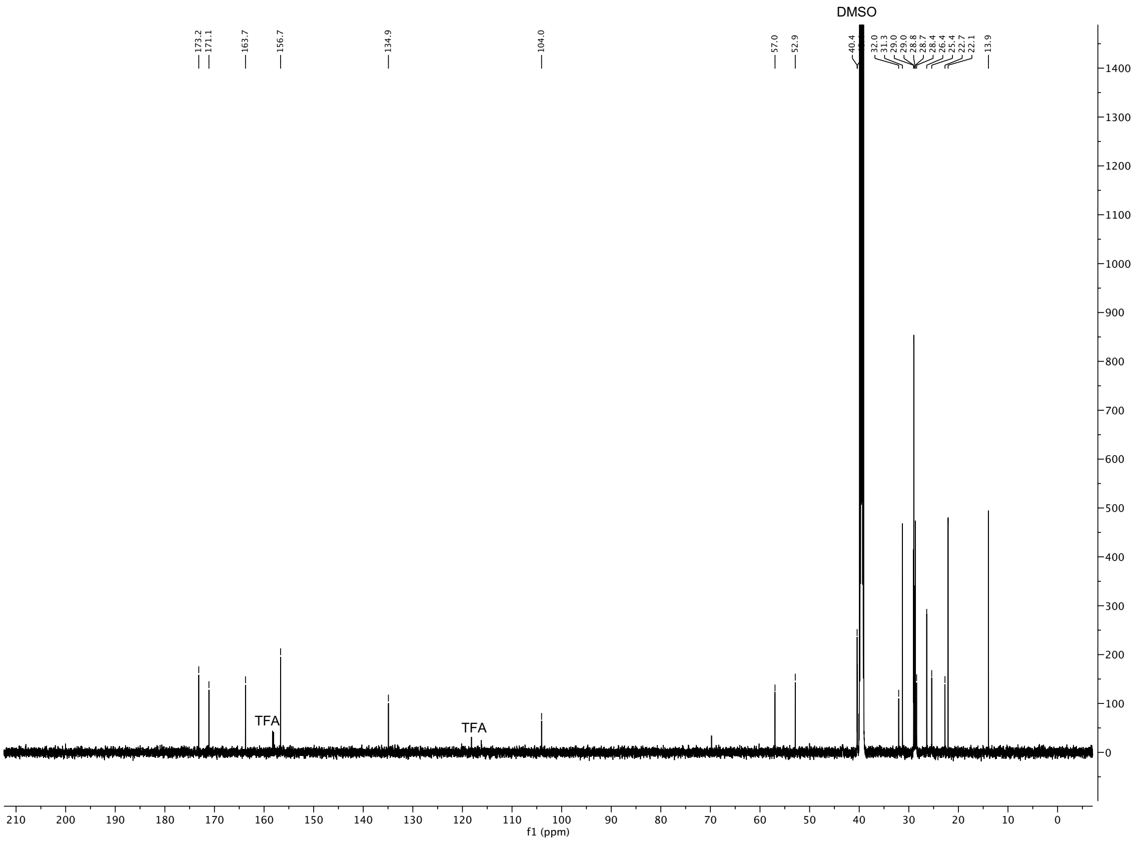

### $^1$H and $^{13}$C spectra of compound **S23**

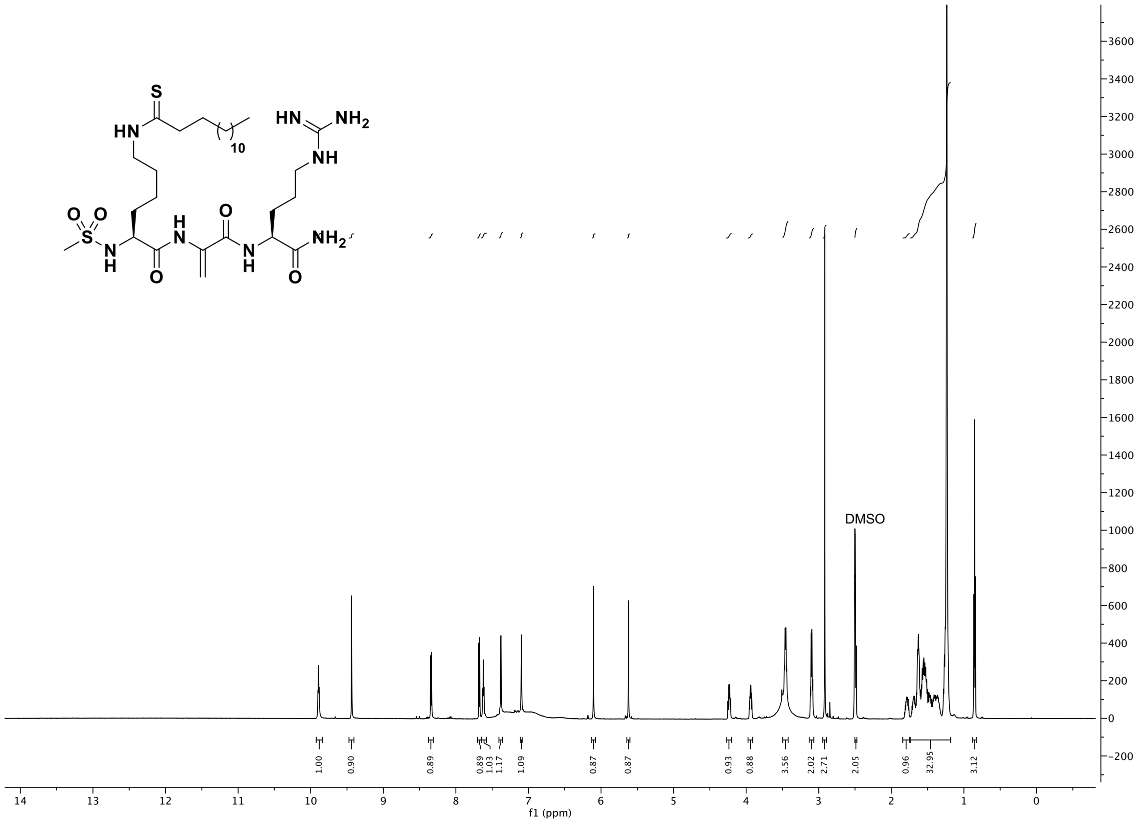

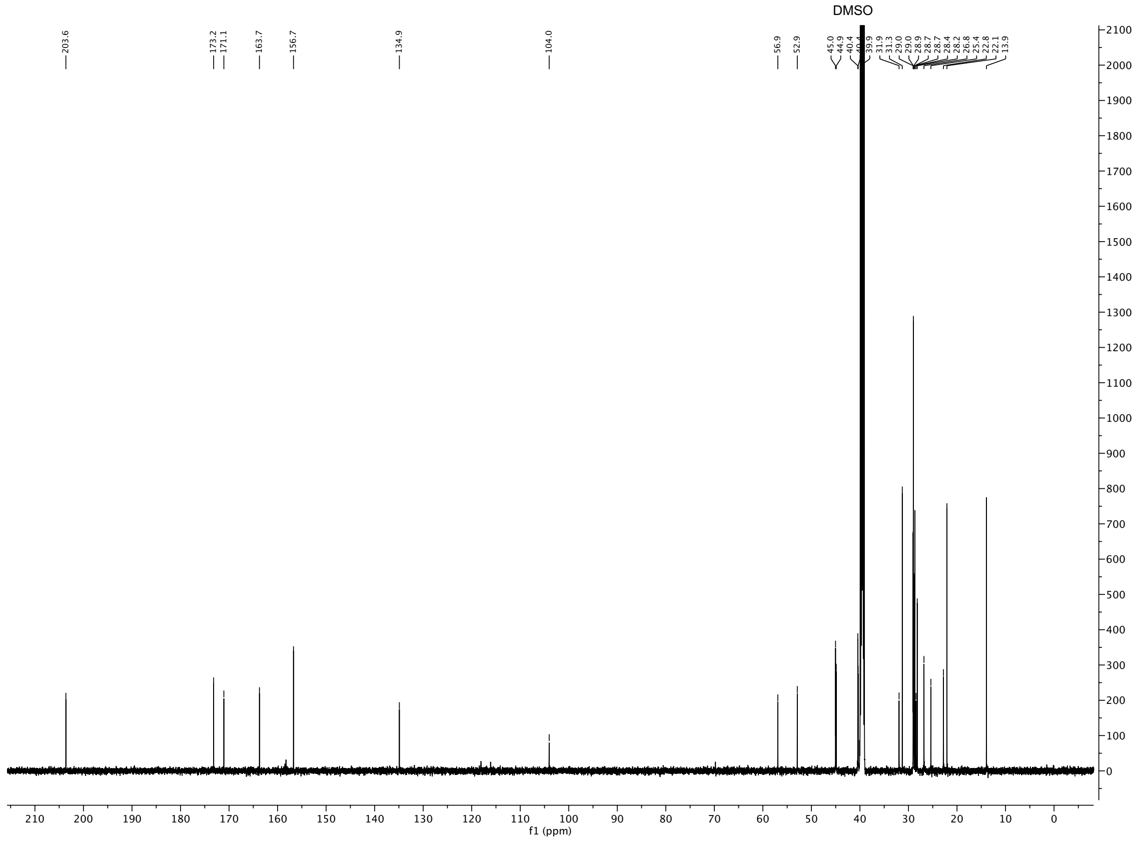

### $^{1}H$ and $^{13}C$ spectra of compound **S24**

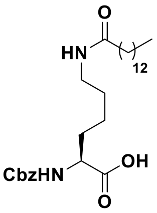

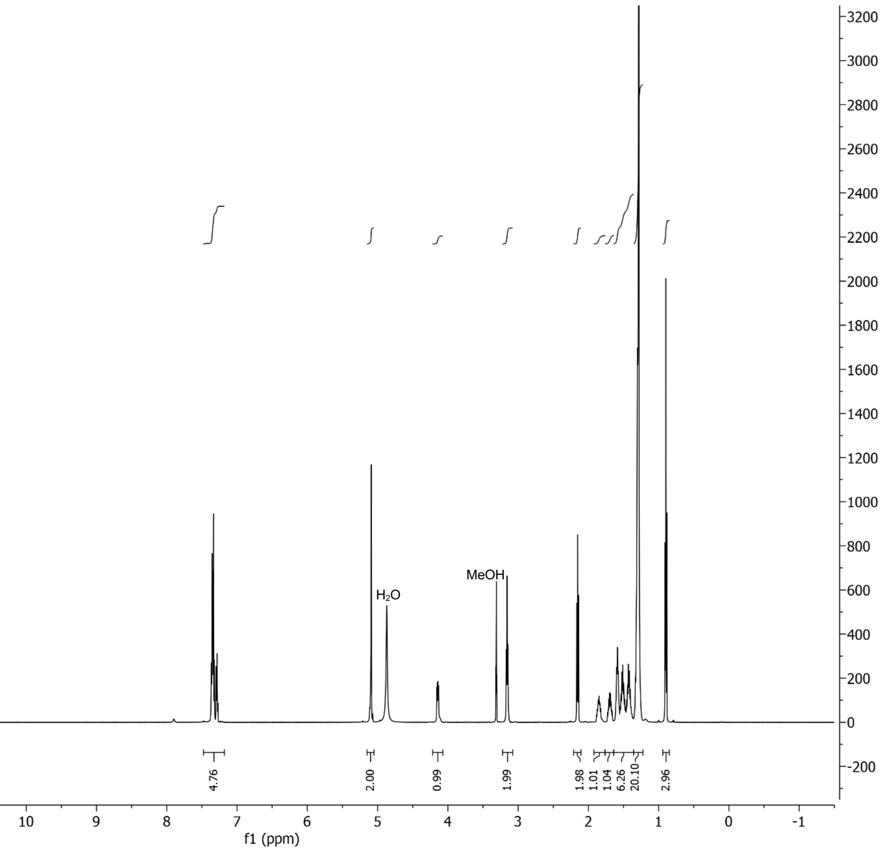

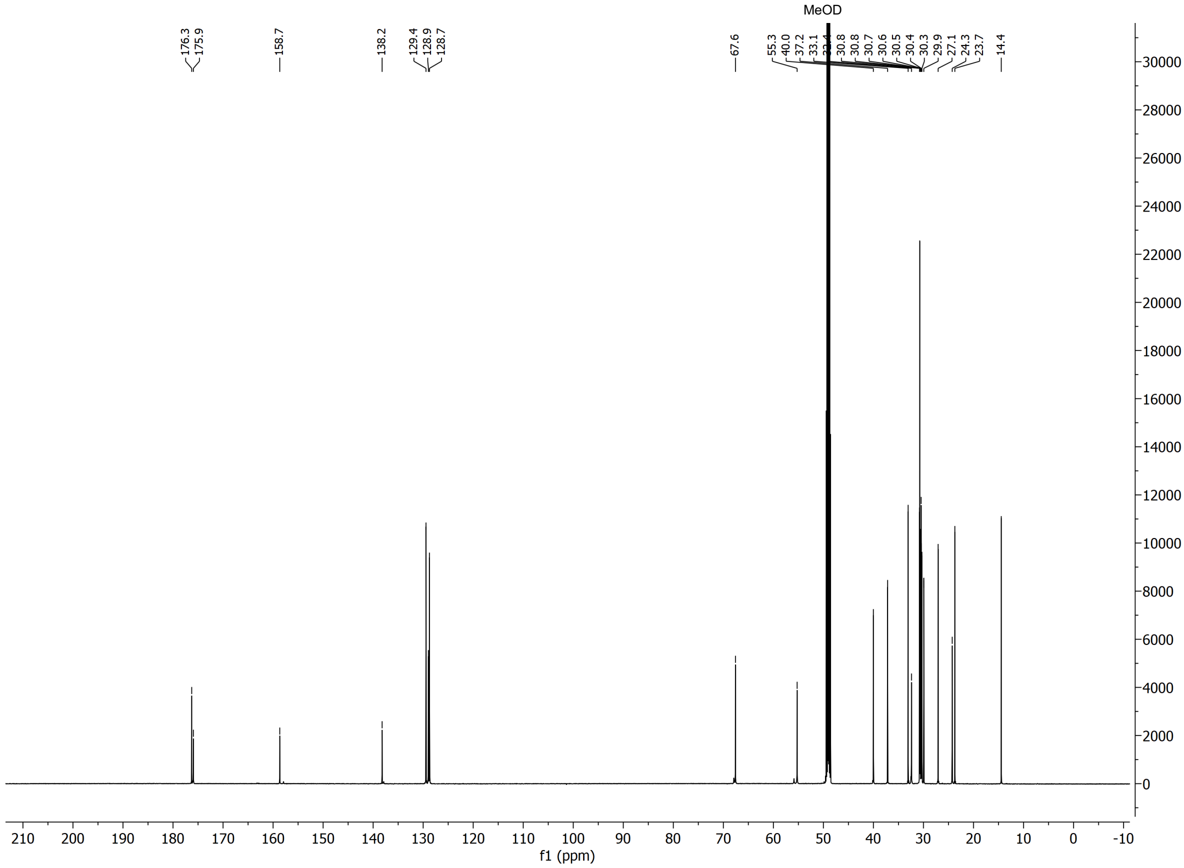

#### $^1H$ and $^{13}C$ spectra of compound **S25**

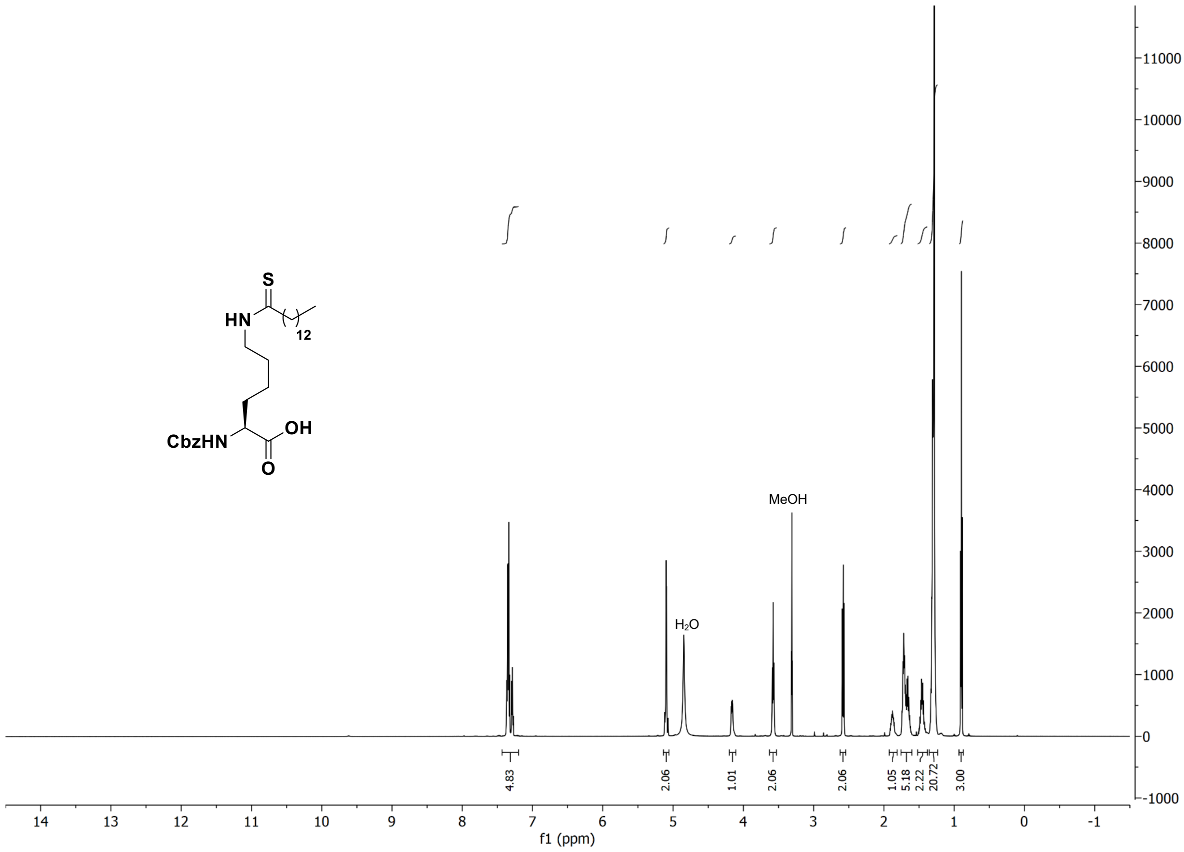

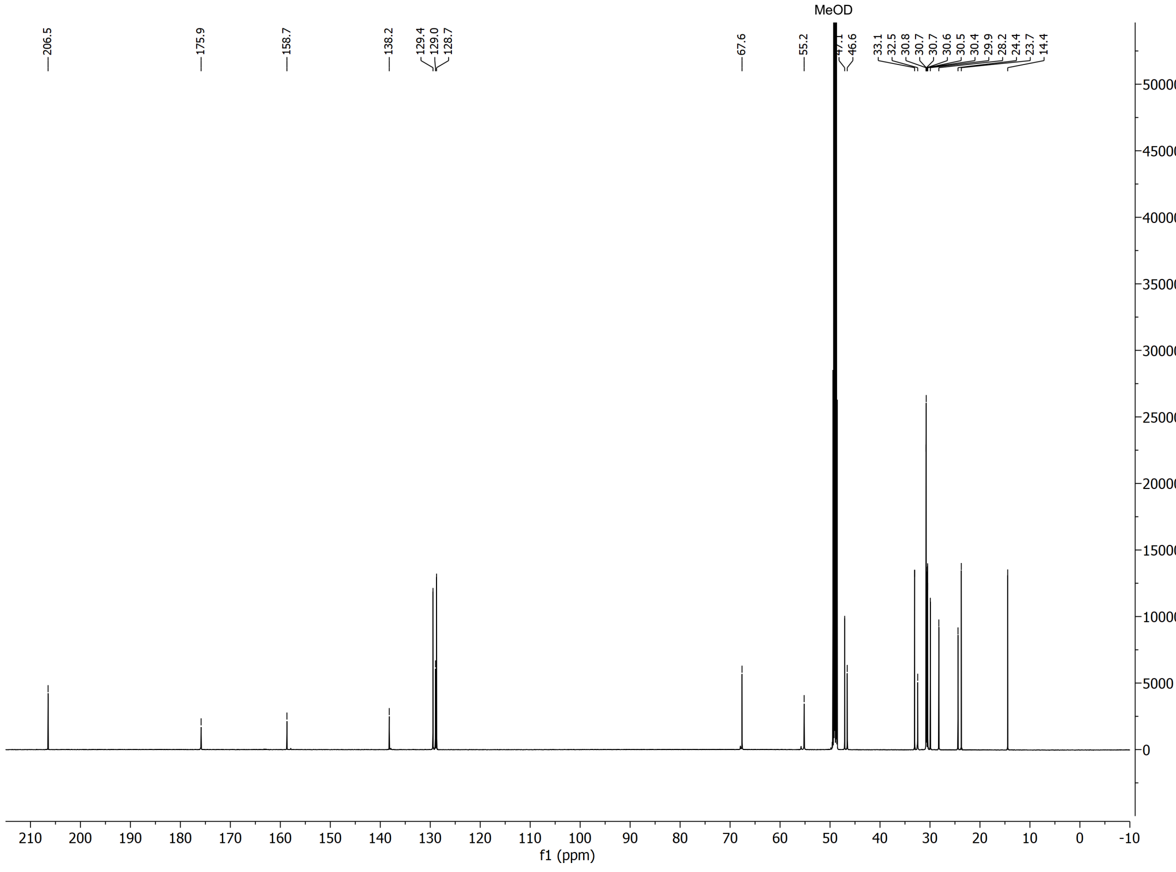

### $^{1}$H and $^{13}$C spectra of compound **S26**

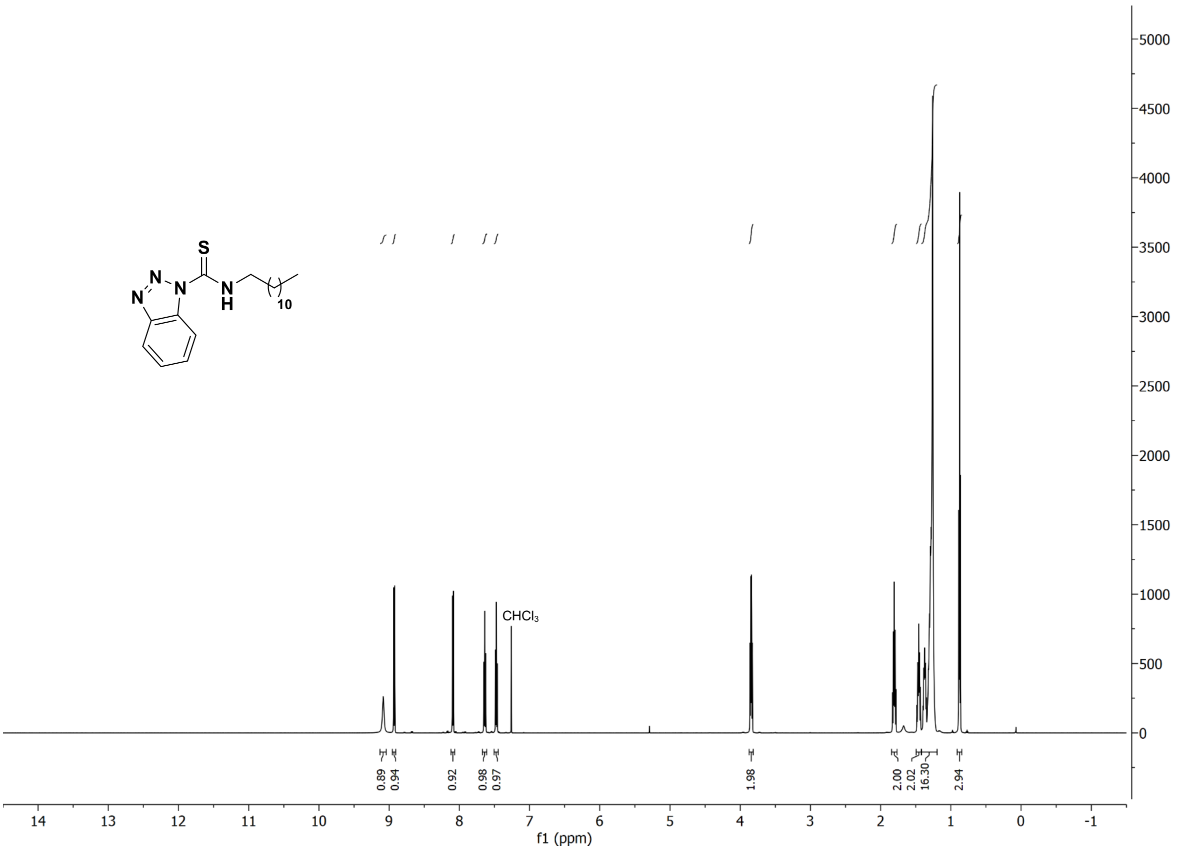

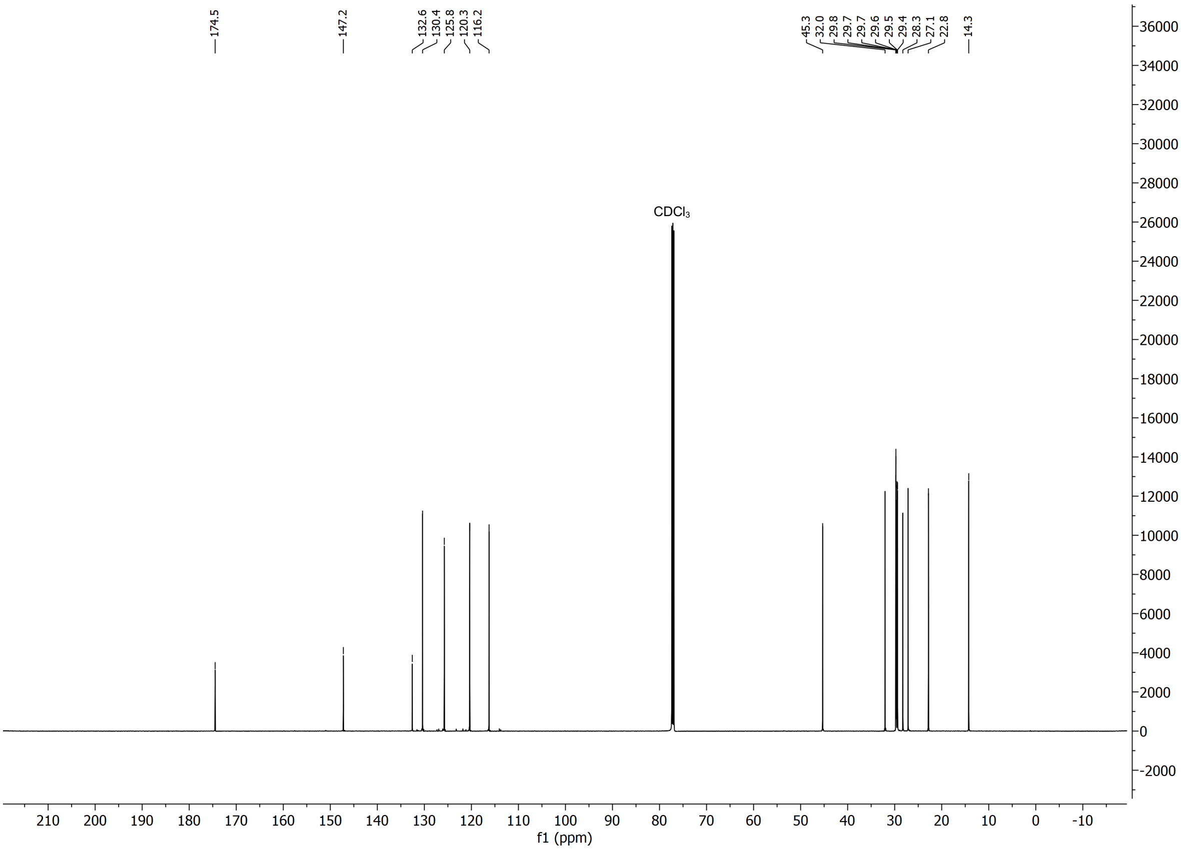

### $^{1}$H and $^{13}$C spectra of compound **S28**

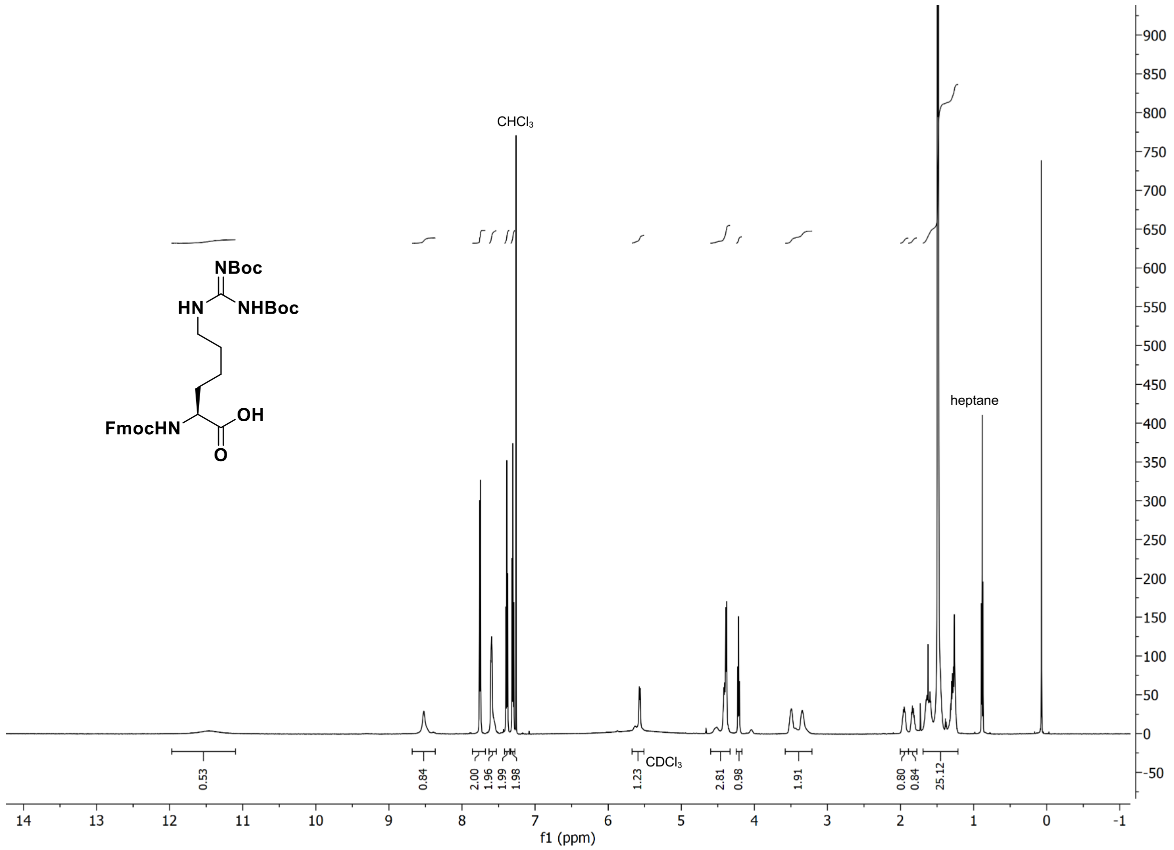

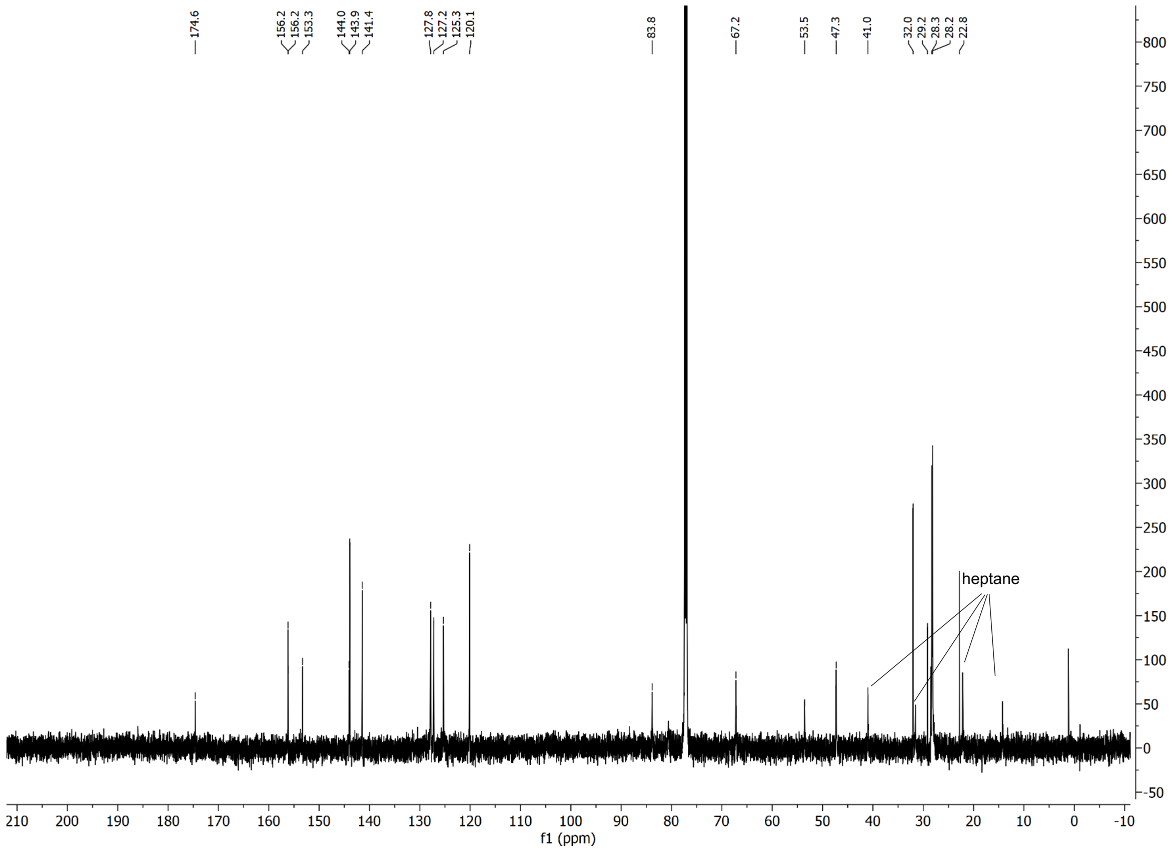

$^1H$ and $^{13}C$ spectra of **TM**

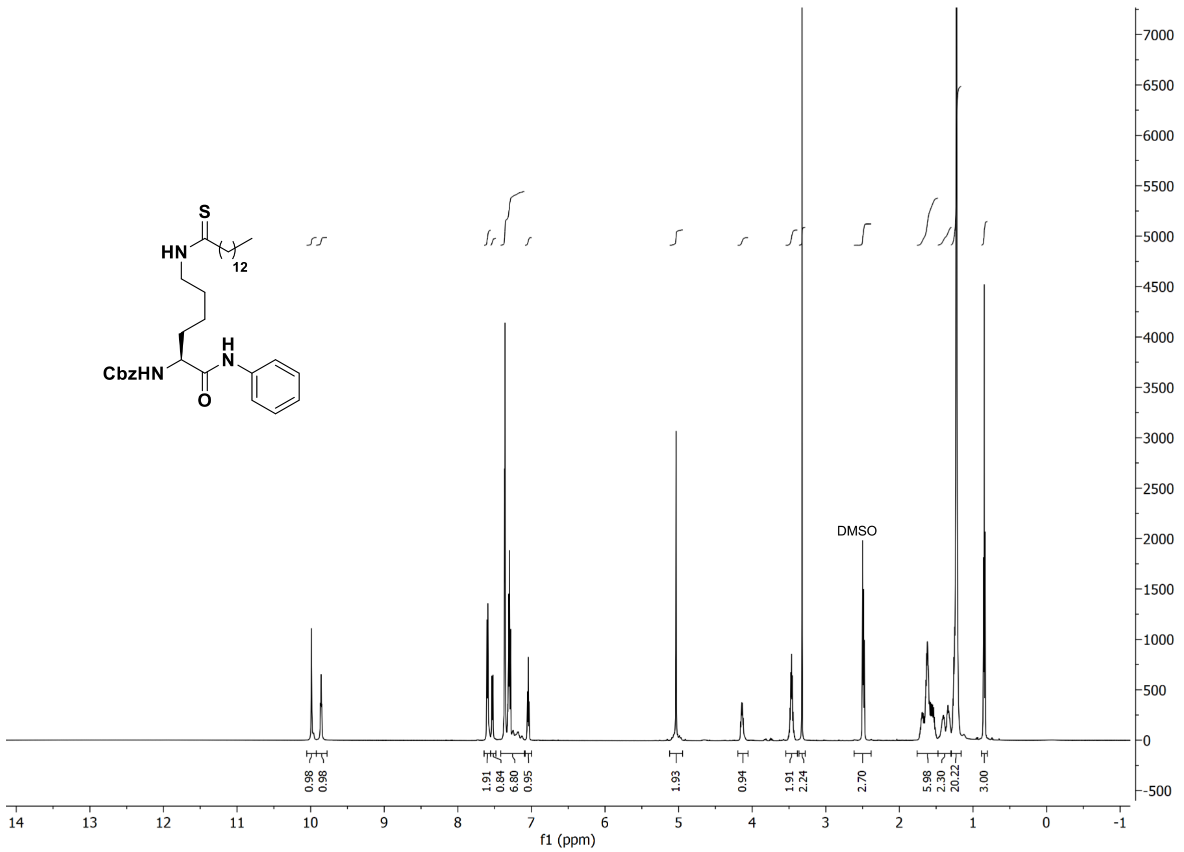

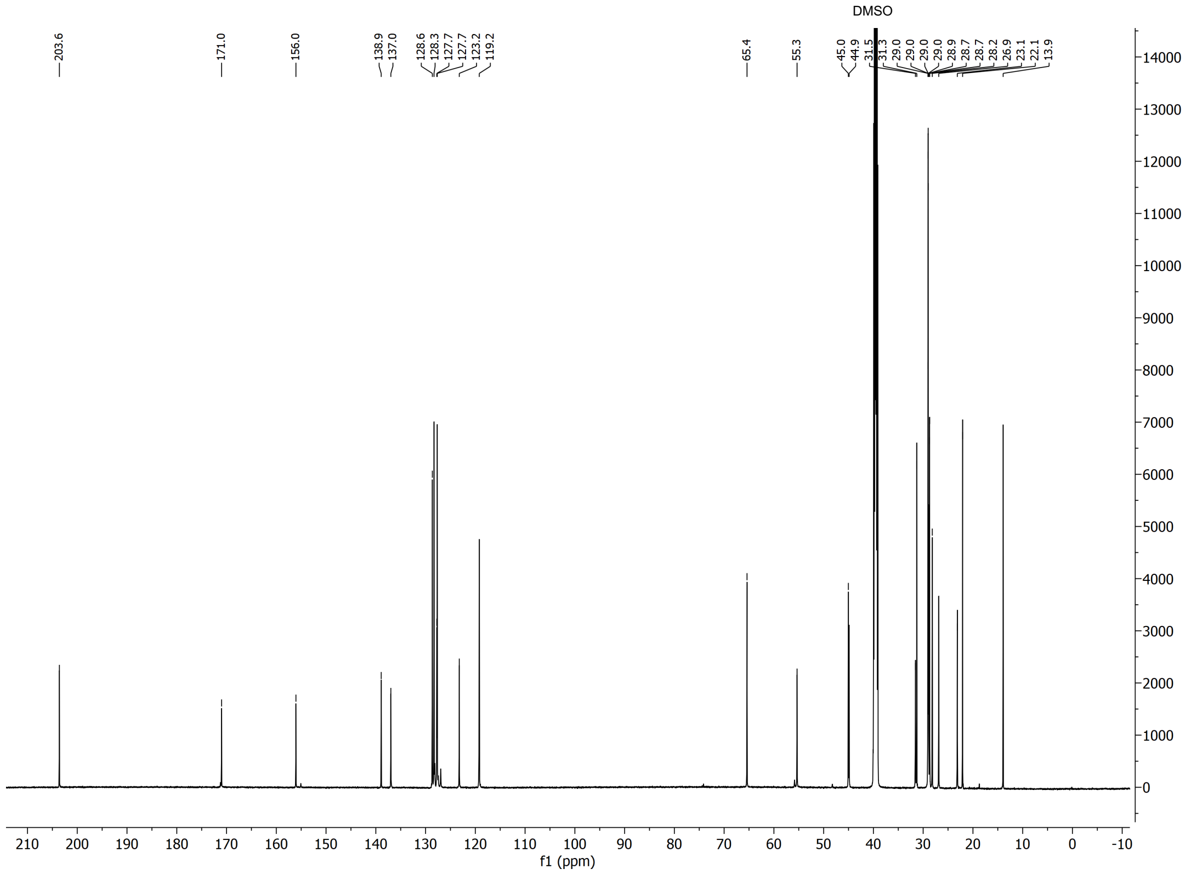

#### $^{1}H$ and $^{13}C$ spectra of **tenovin-6**

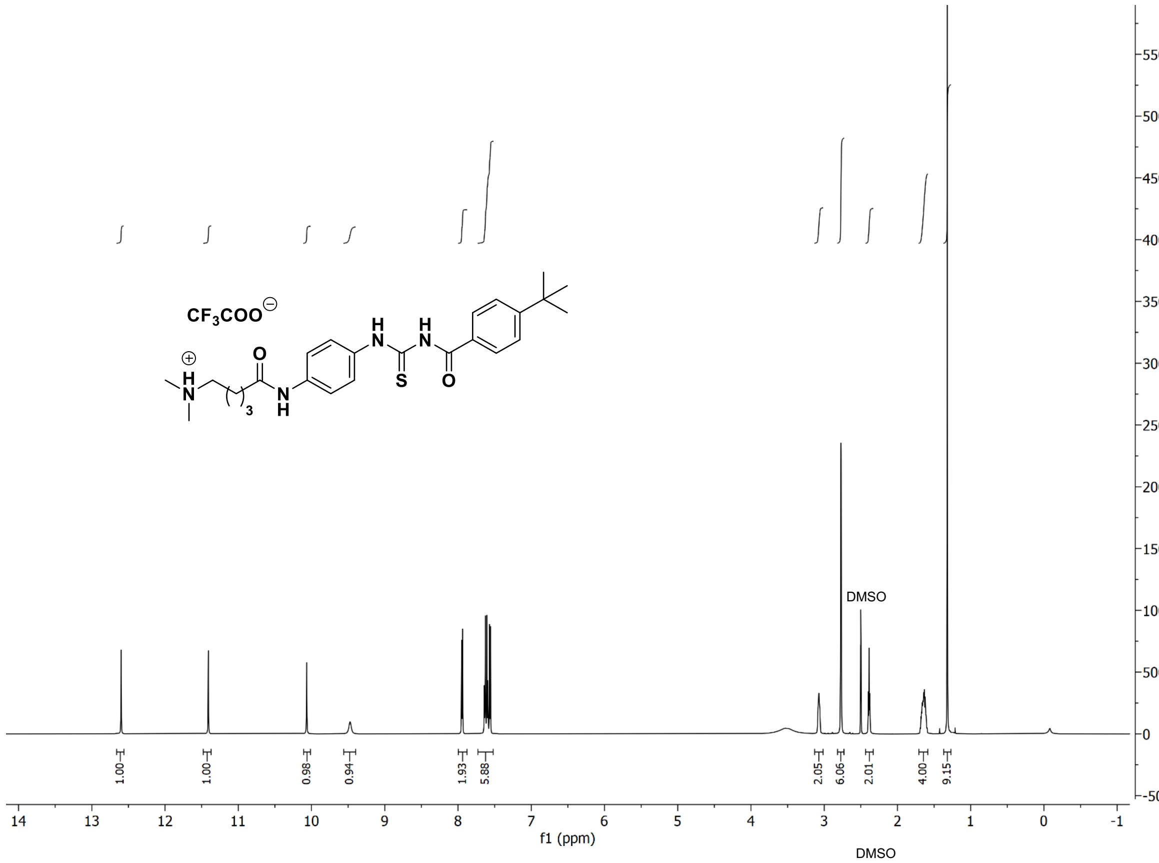

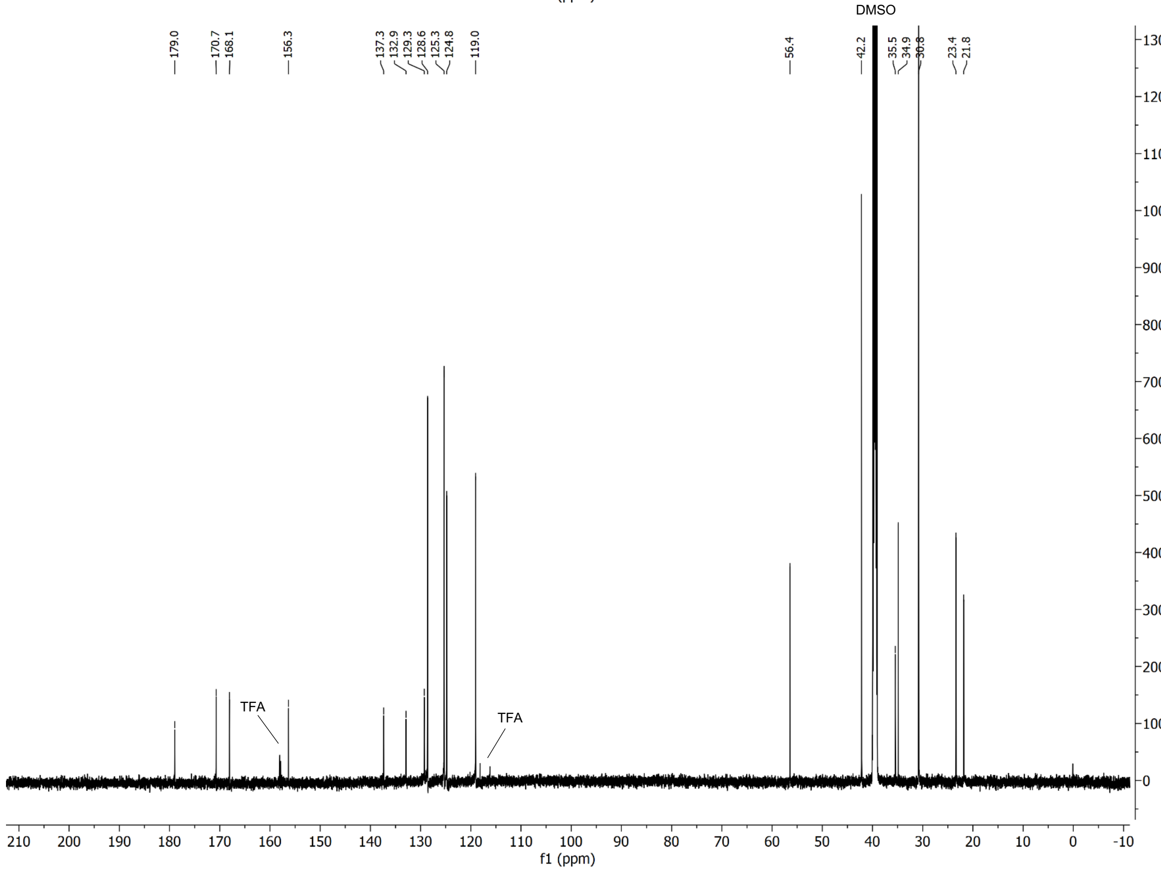
